# ChromBPNet: bias factorized, base-resolution deep learning models of chromatin accessibility reveal cis-regulatory sequence syntax, transcription factor footprints and regulatory variants

**DOI:** 10.1101/2024.12.25.630221

**Authors:** Anusri Pampari, Anna Shcherbina, Evgeny Z. Kvon, Michael Kosicki, Surag Nair, Soumya Kundu, Arwa S. Kathiria, Viviana I. Risca, Kristiina Kuningas, Kaur Alasoo, William James Greenleaf, Len A. Pennacchio, Anshul Kundaje

**Affiliations:** Department of Computer Science, Stanford University, Stanford CA, 94305; Department of Biomedical Data Sciences, Stanford University, Stanford CA, 94305; Department of Genetics, Stanford University, Stanford CA, 94305; Environmental Genomics & System Biology Division, Lawrence Berkeley National Laboratory, Berkeley, CA 94720, USA; Department of Developmental and Cell Biology, University of California, Irvine, CA 92697, USA; Department of Applied Physics, Stanford University, Stanford, California 94305, USA; Institute of Computer Science, University of Tartu, Tartu, Estonia

## Abstract

Despite extensive mapping of cis-regulatory elements (cREs) across cellular contexts with chromatin accessibility assays, the sequence syntax and genetic variants that regulate transcription factor (TF) binding and chromatin accessibility at context-specific cREs remain elusive. We introduce ChromBPNet, a deep learning DNA sequence model of base-resolution accessibility profiles that detects, learns and deconvolves assay-specific enzyme biases from regulatory sequence determinants of accessibility, enabling robust discovery of compact TF motif lexicons, cooperative motif syntax and precision footprints across assays and sequencing depths. Extensive benchmarks show that ChromBPNet, despite its lightweight design, is competitive with much larger contemporary models at predicting variant effects on chromatin accessibility, pioneer TF binding and reporter activity across assays, cell contexts and ancestry, while providing interpretation of disrupted regulatory syntax. ChromBPNet also helps prioritize and interpret regulatory variants that influence complex traits and rare diseases, thereby providing a powerful lens to decode regulatory DNA and genetic variation.

## INTRODUCTION

Transcription factors (TFs) bind specific DNA sequence motifs encoded in cis-regulatory elements (cREs) to modulate transcription of genes. While these motifs are abundant in the genome, TFs selectively occupy only a small fraction of potential sites in a cell context-specific manner^1,2^. Selective TF occupancy is governed by factors beyond the intrinsic sequence specificity and concentration of TFs, including cooperative and competitive interactions between TFs and the nucleosomes that package DNA into chromatin^3–6^. These interactions are mediated by regulatory sequence syntax of cREs, consisting of flexible arrangements of TF motifs with variable composition, affinity, density, spacing, and orientation^7–21^. Disease-associated genetic variants frequently disrupt regulatory sequence syntax, altering TF occupancy, chromatin state and gene expression^22–24^. Hence, deciphering the context-specific sequence code of cREs is crucial for understanding gene regulation and the genetic basis of traits and diseases.

Technical limitations of assays that directly profile genome-wide TF binding have restricted comprehensive binding maps of human TFs to a few cellular contexts^25–36^. Chromatin accessibility assays provide a scalable alternative to profile cREs bound by TFs that often exhibit a remodeled “open” chromatin state devoid of nucleosomes, rendering them more accessible to enzymatic cleavage or transposition^37–42^. DNase-seq and ATAC-seq assays use the DNase-I nuclease and Tn5 transposase respectively to preferentially enrich sequencing coverage within accessible regions, enabling genome-wide identification of putative cREs in a single experiment^43–50^. ATAC-seq has been further adapted to profile chromatin accessibility at single cell resolution^51–55^. These assays have been deployed across thousands of cellular contexts resulting in comprehensive maps of millions of putative cREs, including promoters, enhancers, and insulators, with diverse context-specific accessibility patterns reflecting dynamic TF-mediated regulation^29,30,49,56–63^. Further, thousands of genetic variants have been associated with variation of chromatin accessibility in molecular quantitative trait locus (QTL) studies and disease-associated genetic variants identified by genome-wide association studies (GWAS) are highly enriched in accessible cREs of disease-relevant cell types, highlighting the utility of chromatin accessibility in mapping regulatory genetic variants^23,24,64–71^.

While accessible cREs encompass a large proportion of context-specific TF binding sites and are highly enriched for their cognate motifs, the mere presence of an enriched TF motif in an accessible region is insufficient evidence to infer whether or not it is occupied by a TF or its role in regulating chromatin accessibility^29,49,72–74^. TFs exhibit distinct abilities to engage with chromatin – “pioneer” factors can bind their target motifs within closed chromatin and initiate accessibility through nucleosome remodeling, while other “settler” TFs require pre-accessible regions for binding^5,75–77^. However, even this dichotomization of TFs into pioneers and settlers oversimplifies the regulation of chromatin accessibility by not accounting for sequence-mediated cooperation and competition between multiple TFs and nucleosomes, which depends on genomic context, TF concentration and cellular state. The hierarchical and combinatorial complexity of regulatory syntax also implies that naive overlap of non-coding genetic variants with accessible regions or TF motif instances is not sufficient to understand their pleiotropic, quantitative impact on chromatin accessibility and downstream phenotypes. Several computational approaches have been developed to address these challenges.

Computational footprinting methods attempt to identify occupied TF binding sites in accessible regions by exploiting the observation that some bound proteins protect the DNA from enzymatic cleavage^37–42^. These methods typically employ unsupervised statistical models to scan base-resolution chromatin accessibility profiles to identify characteristic “footprint” patterns of reduced enzymatic cleavage that align with TF motifs^47,77–90^. Some footprinting methods such as HINT-

ATAC, TOBIAS and PRINT explicitly correct for sequence-specific enzymatic biases that can significantly distort footprints in accessible regions by leveraging sequence models of enzyme bias trained on cleavage profiles from genomic background or naked DNA libraries^78,79,90,91^. While these methods have successfully revealed TF binding dynamics and helped prioritize regulatory variants^92,93^, they face key limitations: they typically require high read depth, rely on pre-defined motifs thereby missing novel sequence motifs and cooperative syntax, struggle to detect TFs with low residence times with weak footprints or non-canonical effects on profile shape, and cannot directly infer quantitative effects of variants on chromatin accessibility.

An alternate approach to overcome some of these limitations, involves training and interpreting supervised machine learning models such as support vector machines (SVMs) and deep neural networks to predict chromatin accessibility as a function of local DNA sequence context^94–103^. Deep learning models of regulatory DNA learn de novo predictive sequence features and higher-order syntax, enabling novel insights via various model interpretation frameworks^103–114^. Model predictions can denoise sparse coverage profiles, making them applicable to datasets across a wide range of sequencing depths^104^. Unlike traditional footprinting methods, they can detect TF motifs that influence accessibility without relying on heuristic footprint definitions, potentially identifying TFs with weak or unconventional footprints^94,95,97,99,100,102^. Additionally, in silico mutagenesis experiments can predict the regulatory impact of genetic variants not observed in the training samples^94,95,97–101,115^. However, all contemporary models, with the exception of AlphaGenome^116^, train on either binarized peak level data or coarse-resolution profiles (32-100 bp), thereby not leveraging fine-grained, base-resolution patterns that have been shown to enhance interpretability and predictive accuracy for other biochemical readouts^94–103^. Further, large, multi-task, long-context models trained on diverse regulatory datasets have shown improved predictive performance but require substantial computational resources and cannot be easily trained on individual datasets^97,115,116^. While transfer learning approaches are enabling efficient fine-tuning of pretrained models on new datasets^117–119^, robust and efficient interpretation of large, fine-tuned models remains a challenge.

To address these challenges, we introduce ChromBPNet, a deep learning model augmented with robust model interpretation methods to accurately predict base-resolution chromatin accessibility profiles from local DNA sequence. ChromBPNet features a novel end-to-end framework that automatically detects, learns and deconvolves assay-specific enzyme biases from regulatory sequence determinants of accessibility, thereby significantly improving concordance between DNase-seq and ATAC-seq experiments and enabling imputation of high-resolution, denoised TF footprints and predictive sequence syntax even from low coverage datasets. ChromBPNet models of accessibility across diverse cell-lines reveal relatively compact lexicons of shared and context-specific predictive TF motifs and cooperative composite elements that directly influence chromatin accessibility and reflect TF occupancy. We introduce new measures for predicting base-resolution effects of variants on chromatin accessibility and show through extensive benchmarks that ChromBPNet performs competitively or better than much larger, multi-task models at predicting and interpreting effects of genetic variants on chromatin accessibility, TF binding and reporter activity. Our models exhibit robust variant effect predictions across experimental assays, bulk and single cell datasets, ancestry groups, and cellular contexts. ChromBPNet enables prioritization and interpretation of fine mapped variants in GWAS loci associated with complex traits and putative causal variants in rare disease, with support from validation experiments. ChromBPNet thus offers a powerful but lightweight predictive model for dissecting context-specific sequence determinants of chromatin accessibility and prioritizing regulatory genetic variants.

## RESULTS

### Convolutional neural networks accurately predict base-resolution chromatin accessibility profiles from local DNA sequence context

We trained a convolutional neural network (CNN) to model unstranded, base-resolution ATAC-seq or DNase-seq cleavage profiles in 1 Kb genomic segments as a function of local (2114 bp) regulatory DNA sequence context **(Fig. 1a)**. We adapted the BPNet architecture, originally developed to predict base-resolution binding profiles of individual TFs, by increasing the number of pattern detectors (filters) in each convolutional layer from 64 to 512, to accommodate the more complex sequence lexicon and syntax required to predict genome-wide chromatin accessibility^104^.

**Figure 1:**
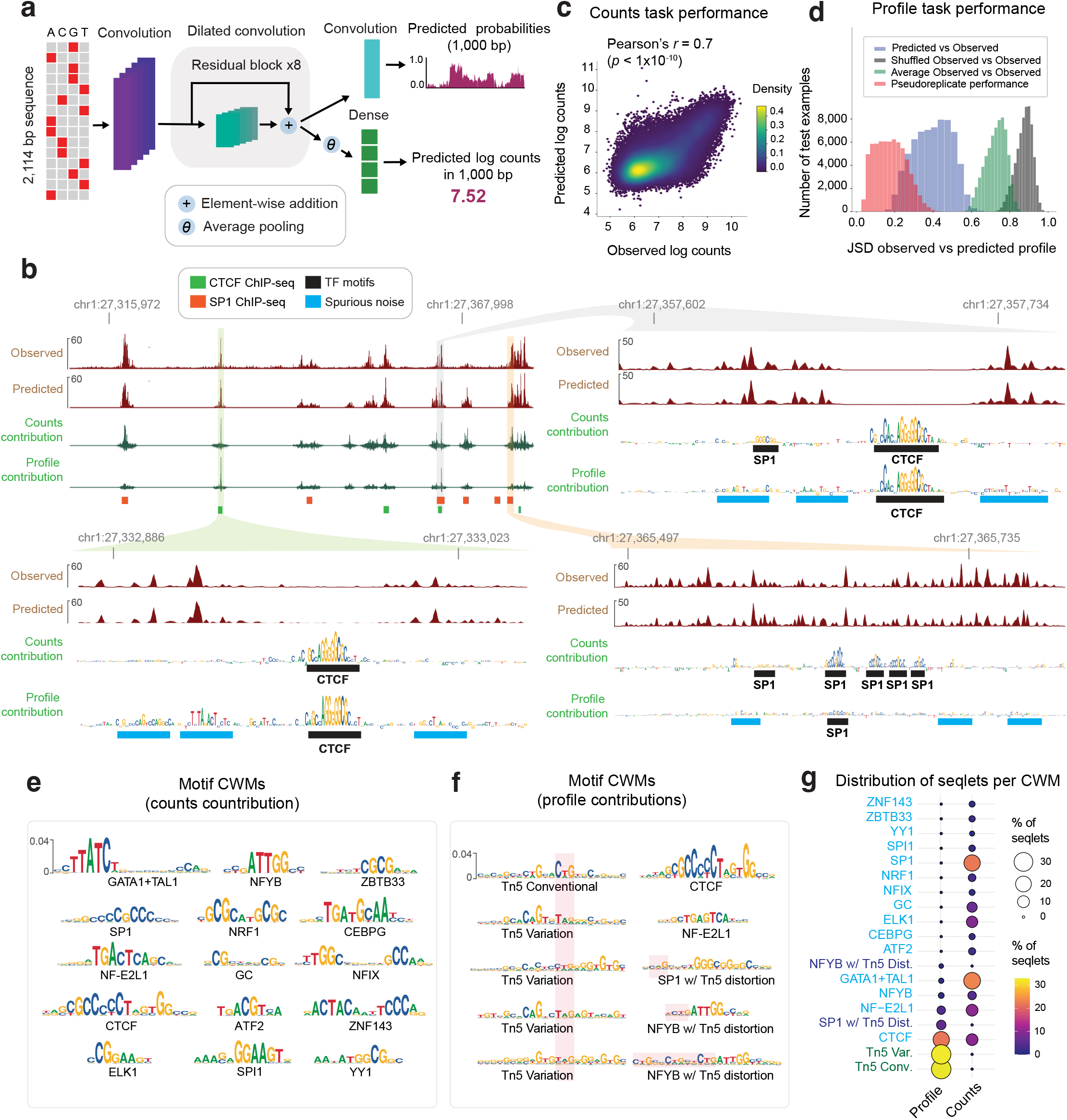
Convolutional neural networks accurately predict base-resolution ATAC-seq profiles from local DNA sequence context but are confounded by Tn5 sequence bias. **(a)** Schematic of the expanded BPNet architecture that maps 2114 bp of input sequence to 1 Kb base-resolution DNase-seq or ATAC-seq cleave profiles as two complementary outputs - a count head capturing the total coverage (log scale) and a profile head that captures the base-resolution multinomial probability distribution of reads **(b)** Exemplar locus on chr11 (held-out test chromosome) showing observed (Obs) ATAC-seq profiles, BPNet predicted (Pred) profiles at multiple resolutions and DeepLIFT sequence contribution scores to count and profile shape predictions. High count contribution scores highlight TF motif instances, whereas high profile contributions also capture spurious features that resemble Tn5 sequence preference. **(c)** Measured and predicted log(total counts) from K562 ATAC-seq peaks in held-out chromosomes exhibit high and statistically significant correlation. **(d)** Distribution of Jensen-Shannon Distance (JSD) between measured and predicted base-resolution K562 ATAC-seq profiles from peak regions in held-out chromosomes (blue) partially overlaps with distribution of concordance of pseudo-replicate profiles (red) which acts an upper bound, and is substantially better that baseline concordance between observed profiles and the average profile over all peaks (green) and between observed profiled and scrambled versions of the observed profiles (grey). **(e)** Top 15 TF-MODISCO contribution weight matrix (CMW) motifs derived from count contribution scores of the K562 ATAC-seq BPNet model map to well known TF motifs. **(f)** TF-MODISCO CWM motifs derived from profile contribution scores map to Tn5 bias motifs and distorted TF motifs. **(g)** Comparison of the frequency of high-confidence predictive instances (seqlets) of each TF-MODISCO motif identified from the count and profile heads show that Tn5 bias motifs dominate the profile contribution scores over TF motifs.

The model predicts the expected base-resolution coverage profile over each segment as two complementary outputs: the “count head” of the model predicts total coverage (total counts) over the segment, which captures the magnitude of accessibility, while the “profile head” predicts the profile shape in the form of a base-resolution multinomial probability distribution across the segment. Model parameters were optimized using the BPNet multiscale loss function composed of a linear combination of a mean-squared error (MSE) loss for the total coverage (in *log* scale) and a multinomial negative log-likelihood loss for profile shape^104^. Model training, tuning and performance evaluation were performed using reproducible peaks and G/C content-matched background regions from non-overlapping chromosome partitions in a 5-fold cross-validation set up.

We first trained separate models on ATAC-seq and DNase-seq profiles in the K562 cell line^30^. Visual inspection of measured and predicted profiles over representative peak regions in held-out chromosomes highlighted strong concordance at multiple resolutions (**Fig. 1b, Extended Fig. 1a)**. Systematic evaluation of prediction performance across the held-out peak regions in test chromosomes corroborated the strong correlation (Pearson correlation *r* = 0.70+/-0.02 *p* < 1e-10 for ATAC-seq and *r* = 0.71+/-0.02, *p* < 1e-10 for DNase-seq) between measured and predicted total counts **(Fig. 1c, Extended Fig. 2b).** Measured and predicted profile shapes were also highly concordant, as measured by Jensen-Shannon Distance (JSD), approaching the upper bound of pseudoreplicate concordance (**Fig. 1d, Extended Fig. 2c**). The models also accurately discriminate high-confidence, reproducible peaks from background regions across entire test chromosomes (area under the Receiver Operating Curve (auROC) = 0.98+/-0.001, average precision (AP) = 0.42+/-0.01 for ATAC-seq and auROC = 0.98+/-0.001, AP = 0.46+/-0.02 for DNase-seq).

### Model interpretation reveals the influence of Tn5/DNase-I sequence preferences on chromatin accessibility profiles

Next, we applied our well-established model interpretation framework to identify predictive sequence features^97,120^. First, we used DeepLIFT/DeepSHAP to estimate the contribution scores of individual bases in peaks to the model’s predictions of total counts (count contribution) and profile shape (profile contribution) separately^121,9797,120,97^. Visual inspection of count contribution scores from the K562 ATAC-seq models at three representative ATAC-seq peaks in K562 clearly highlighted predictive motif instances of key TFs such as CTCF and SP1, supported by ChIP-seq peaks (**Fig. 1b**). In contrast, profile contribution scores at these loci were diminished for several TF motifs (e.g. SP1 motifs) that had high count contributions and highlighted several spurious bases that did not match known TF motifs.

To investigate the discrepancy between count and profile contribution scores more globally, we derived count and profile contribution scores within 30,000 randomly sampled ATAC-seq peak sequences in K562. We then used the TF-MODISCO motif discovery algorithm to align and cluster subsequences with high contribution scores (called seqlets) from all the scored sequences into a consolidated, non-redundant repertoire of motifs in the form of Contribution Weight Matrices (CWMs) and Position Frequency matrices (PFMs)^106,97^. We derived TF-MODISCO motifs separately for count and profile contribution score profiles and assigned putative TF labels for each motif based on their similarity to previously annotated TF motifs and overlap enrichment with ENCODE TF ChIP-seq datasets in K562 (**Fig. 1e,f**). Motifs derived from count contribution scores mapped to well-known TF drivers of chromatin accessibility in K562 including GATA-TAL1, NFYB, STAT5A, CTCF, SP1, NFIX, ATF2, and ELK1 (**Fig. 1e**). In contrast, several motifs derived from profile contribution scores did not match known TFs, while those that did match were less prevalent and distorted (**Fig. 1f,g**). A closer examination of the top ranking spurious motifs suggested a strong resemblance to the intrinsic sequence preference of the Tn5 transposase used in ATAC-seq experiments (**Fig. 1f**)^79^.

Analogous interpretation of DNase-seq models in K562 revealed greater similarity of profile and count contribution scores than observed for ATAC-seq models. Both types of contribution scores at representative DNase-seq peaks highlighted known TF motif instances and fewer spurious bases **(Extended Fig. 1a)**. TF-MODISCO motifs derived from count and profile contribution scores were also similar to each other and to motifs derived from count contribution scores of the ATAC-seq models **(Extended Fig. 1b,c)**. However, TF-MODISCO identified one motif matching the intrinsic sequence preference of the DNase-I enzyme from the DNase-seq profile contribution scores but not count contribution scores **(Extended Fig. 1c)**. Nevertheless, its prevalence and influence was less pronounced compared to Tn5’s motif identified from ATAC-seq profile contribution scores **(Extended Fig. 1b-d)**.

Collectively, these results demonstrate that deep learning models trained naively on base-resolution chromatin accessibility profiles conflate the influence of TF motif syntax with the intrinsic sequence preferences of enzymes like Tn5 and DNase-I. While total counts (overall accessibility) in peaks are not affected by enzyme sequence preferences and exclusively determined by TF motif syntax, the base-resolution positional distribution of reads (profile shape) in peaks is impacted by enzyme sequence preferences, with Tn5 causing stronger distortions in ATAC-seq profiles than DNase-I in DNase-seq profiles. These insights led us to develop new model training strategies that explicitly decouple the influence of enzyme sequence preferences from the contribution of TF motif syntax on base-resolution chromatin accessibility profiles.

### CNNs learn Tn5/DNase-I sequence preferences from background cleavage profiles of chromatin accessibility experiments

To correct the impact of intrinsic enzymatic biases on base-resolution positional distribution of reads, we first developed new predictive sequence models of enzyme sequence preference. For ATAC-seq, unstranded Tn5 insertion profiles are typically obtained by aggregating appropriately offset 5’ read starts from both strands. The canonical offsets used to correct for Tn5’s dimeric binding and the insertion of two adaptors, spaced 9 bp apart, are +4 bp for reads mapping to the + strand and-5 bp for reads mapping on the - strand^122^. However, other offsets, such as +4/-4 bp shifts, have also been reported to improve models of Tn5 sequence preference^79,91^. To reconcile these reports, we systematically evaluated the different offset choices based on alignment of simple position weight matrix (PWM) motif models of Tn5 sequence preference derived from *k*-mers centered on shifted 5’ read-starts in background regions from the + and - strand separately. +4/-4 bp offsets were found to result in superior alignment of Tn5 PWMs derived from the two strands, justifying the use of this offset to derive unstranded coverage profiles of Tn5 insertions aggregated over both strands. (**Extended Fig. 3a-b**). Analogous analysis of strand-specific PWMs derived from DNase-I cleavage profiles, identified shifts of 0 bp for reads on the + strand and +1 bp for reads on the - strand for optimal alignment of DNase-I sequence preferences across strands^79,91^.

Previous models of Tn5 and DNase-I sequence preference have used mononucleotide and dinucleotide weight matrix models, and enumerated or ensemble *k*-mer models^78,79,85,88,91,92,123–126^. Given the prior success of CNNs at modeling sequence specificity of TFs, we decided to test whether BPNet models trained only on ATAC-seq or DNase-seq profiles from non-accessible background genomic regions devoid of TF binding might learn accurate models of Tn5 and DNase-I sequence preferences. A modified BPNet “bias model” with 128 filters and an effective receptive field of 100 bp trained on sparse background ATAC-seq profiles in GM12878 was surprisingly effective at predicting total counts and profile shapes (Pearson correlation *r =* 0.60 (*p* < 1e-10), median JSD = 0.74) (**Extended Fig. 3c-d**), suggesting a strong influence of intrinsic Tn5 sequence preference in background regions. Since accidental training on weakly accessible regions occupied by sequence-specific TFs could result in contaminated bias models that learn TF motifs alongside enzymatic sequence preferences, we used our interpretation framework to systematically discover all sequence motifs learned by bias models trained on background regions passing different signal thresholds. We introduced a background signal coverage threshold as an adjustable hyperparameter with conservative defaults ensuring that bias models only learned motifs resembling Tn5 bias or other low complexity sequence patterns that did not resemble any known TF motifs.

We then used the bias model trained on background regions to predict Tn5 insertion bias profiles in peak regions. A representative peak locus in GM12878 revealed remarkable similarity between the predicted bias profile, the measured ATAC-seq profile and the predicted ATAC-seq profile from the expanded BPNet model. The exception was around the peak summit, where there was stronger protection against Tn5 insertion in measured and predicted ATAC-seq profiles compared to the predicted bias profile (**Fig. 2b)**. DeepLIFT profile contribution scores from the expanded BPNet model highlighted a CTCF binding site at the peak center, likely conferring this protection, as well as other spurious bases in the rest of the sequence. Profile contribution scores of the same sequence from the Tn5 bias model highlighted many of the same spurious bases suggesting that these likely reflect Tn5 sequence preference. However, the CTCF motif had no detectable contribution to the Tn5 bias profile confirming that the bias model had not accidentally learned the CTCF motif.

**Figure 2:**
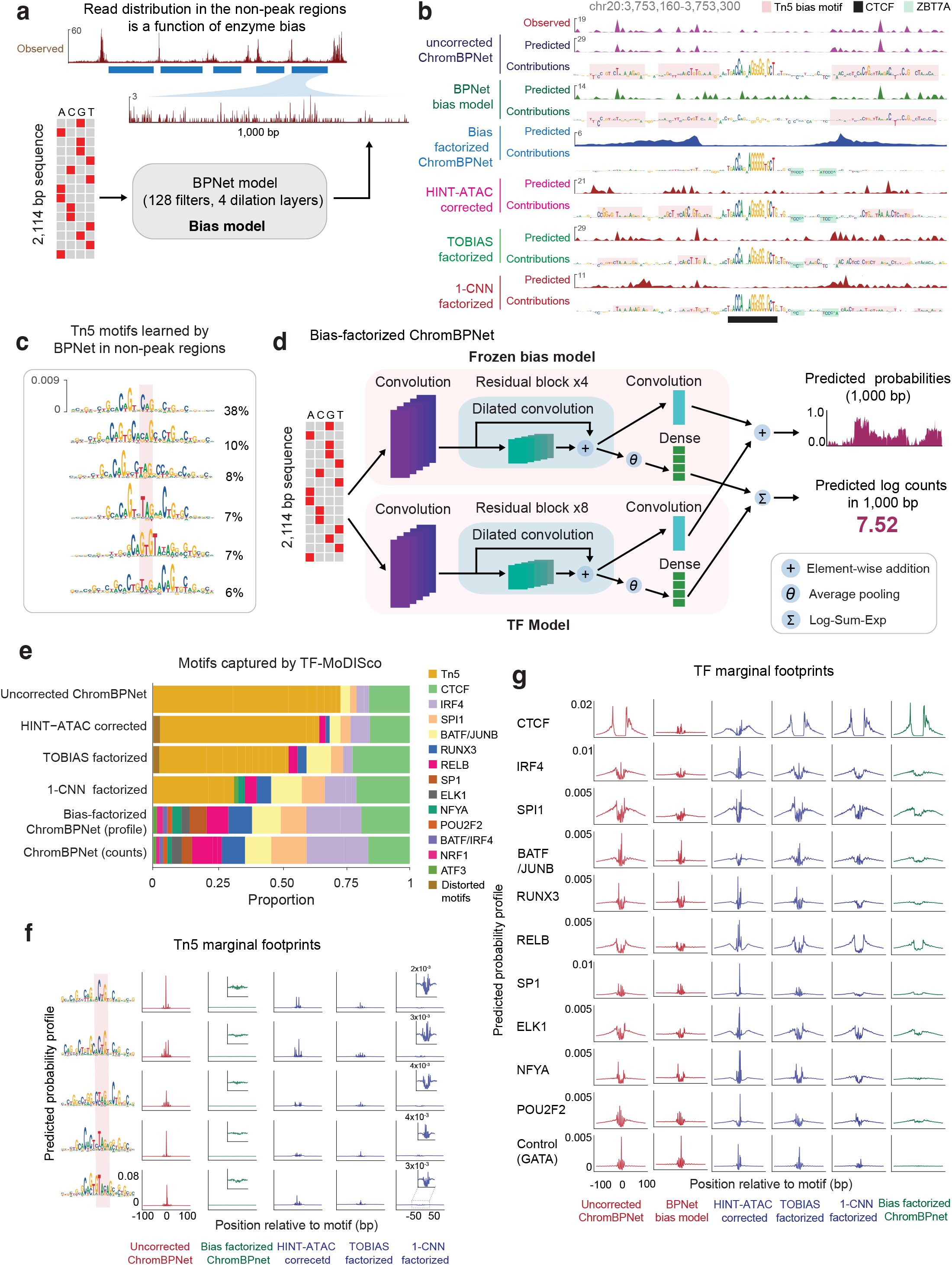
Bias-factorized ChromBPNet models of ATAC-seq profiles coupled to neural network models of Tn5 bias deconvolve Tn5 bias from regulatory TF motif syntax. **(a)** Schematic of BPNet models used to learn Tn5 bias from ATAC-seq profiles from background chromatin that excludes ATAC-seq peaks. **(b)** Exemplar locus at chr20:3,753,160-3,753,300 showing several tracks from top to bottom: observed GM12878 ATAC-seq profile; uncorrected predicted profile from ChromBPNet; uncorrected profile contribution scores from ChromBPNet shows spurious Tn5 bias; predicted Tn5 bias profiles from BPNet bias model shows strong resemblance to observed and uncorrected profiles; profile contribution scores from bias model match spurious Tn5 bias features; bias-corrected predicted profile from ChromBPNet shows a strong, denoised latent footprint; bias-corrected profile contribution scores from ChromBPNet are devoid of spurious Tn5 bias highlighting CTCF and ZBTB7A motifs; predicted profiles from ChromBPNet model using HINT-ATAC’s bias correction method; profile contribution scores from HINT-ATAC ChromBPNet model show spurious Tn5 bias; predicted profiles from ChromBPNet model using TOBIAS’s bias correction method; profile contribution scores from TOBIAS ChromBPNet model shows spurious Tn5 bias; predicted profiles from ChromBPNet model that uses a simplified 1-filter CNN bias model; profile contribution scores from ChromBPNet model with 1-filter CNN bias model shows spurious Tn5 bias. **(c)** TF-MODISCO motifs derived from profile contribution scores of the BPNet bias model trained on chromatin background capture different variations of the canonical Tn5 sequence preference motif. **(d)** Schematic of the bias-factorized architecture of ChromBPNet consisting of a pre-trained bias model and residual TF sub-model with trainable parameters. **(e)** Relative frequency of high-confidence predictive instances (seqlets) of TF-MODISCO motifs derived from profile contribution scores of various ChromBPNet models trained on GM12878 ATAC-seq data using different bias models (no bias model, TOBIAS bias model, 1-filter CNN bias model and BPNet bias model) or pre-corrected profiles (HINT-ATAC) shows that Tn5 bias motifs dominate all but the ChromBPNet model coupled with the BPNet bias model. **(f)** Marginal footprints of various TF-MODISCO Tn5 bias motifs (rows) using predicted profiles from GM12878 ATAC-seq ChromBPNet models using different bias models or pre-corrected profiles (cols) show that only the BPNet bias model enables near optimal correction of Tn5 bias. **(g)** Marginal footprints of various predictive TF motifs (rows) using predicted profiles from GM12878 ATAC-seq ChromBPNet models that use different bias models or pre-corrected profiles (cols) show that only the BPNet bias model removes confounding Tn5 bias.

Systematic comparison of the predicted Tn5 bias profiles against measured ATAC-seq profiles across all peak regions in each of the cell-lines, revealed high concordance of profile shapes (median JSD=0.48) but poor prediction of total counts (*r* =-0.16) (**Extended Fig. 3e-h**), corroborating our previous results from the expanded BPNet ATAC-seq models that Tn5 bias strongly influences profile shape but not total counts in ATAC-seq peaks. TF-MODISCO further confirmed this conclusion by discovering pervasive Tn5 bias motifs in profile contribution scores but not in count contribution scores from the Tn5 bias model across ATAC-seq peak sequences (**Fig. 2c, Supp Files 3)**. Further, no known TF motifs were identified by TF-MODISCO in count and profile contribution scores from the Tn5 bias model across ATAC-seq peak sequences, affirming that the Tn5 bias model did not accidentally learn any known TF motifs.

Analogous results and conclusions were obtained for bias models trained on DNase-I cleavage profiles in background regions from DNase-seq experiments (**Extended Fig. 3g, 3h, 5a, 5b**). Interestingly, TF-MODISCO applied to profile contribution scores of the DNase bias models in background regions revealed multiple subtle variations of the canonical DNase-I sequence preference motif, suggesting that even for DNase-I, a single PWM motif model is insufficient to fully represent its sequence preference (**Extended Fig. 5a**). TF-MODISCO did not discover any DNase-I bias motifs from count contribution scores of the DNase bias model, supporting our previous observation that DNase-I sequence preference also exclusively affects profile shape (**Supp Files 3**).

Finally, we explicitly quantified the sensitivity of ATAC-seq and DNase-seq profiles to the various variations of Tn5 and DNase-I bias motifs with a “marginal footprinting” approach (**Extended Fig. 3i-k, 5d)**^104,109,127^. We sampled the highest affinity *k*-mer from each bias motif, embedded it into a library of randomized background sequences, and averaged the normalized profile predictions from the bias models over the entire library to obtain marginal footprints. Each of the Tn5 and DNase-I bias motifs produced distinct marginal footprints, confirming the sensitivity of the bias model to these motifs. However, marginal Tn5 bias footprints were orders of magnitude stronger than the marginal DNase-I bias footprints, supporting our previous conclusions that Tn5 introduces stronger distortion of profile shapes than DNase-I.

### A bias-factorized neural network (ChromBPNet) deconvolves enzyme sequence preferences from regulatory sequence determinants of chromatin accessibility

Next, we developed a novel “bias-factorized” neural network architecture, named ChromBPNet, to deconvolve enzyme sequence preferences from TF motif syntax and other sequence features influencing base-resolution ATAC-seq and DNase-seq profiles in peak regions (**Fig. 2d**). ChromBPNet processes 2114 bp input sequences to produce a 1 Kb base-resolution profiles, decomposed into total counts and base-resolution positional probability distribution of reads through two parallel submodels. It combines a pre-trained BPNet enzyme bias model whose parameters are frozen with an expanded BPNet architecture which has learnable parameters. Training occurs in two stages. The first stage fits and scales the predicted enzyme bias profiles from the frozen bias submodel to the measured total counts and profile shapes across peaks and background regions in the training set. The second stage trains the expanded BPNet submodel on the residual variance in the total counts and profile shape not explained by the bias submodel from the first stage fit. The model predicts uncorrected accessibility coverage profiles by combining outputs from both submodels. To obtain a “bias-corrected” coverage profile, we disconnect the bias submodel and only use the residual submodel’s predictions. Uncorrected or bias-corrected count and profile sequence contribution scores are derived by applying DeepLIFT/SHAP to the predictions from count and profile heads of the combined model or residual submodel, respectively.

We trained separate ChromBPNet models on ATAC-seq and DNase-seq profiles from 5 ENCODE Tier 1 cell-lines: K562, HepG2, GM12878, IMR90, and H1-hESC. In K562, the prediction performance of uncorrected predictions from the ChromBPNet models matched that of the expanded BPNet models without bias factorization (**Extended Fig. 2a**). ChromBPNet models from the other 4 cell-lines also achieved high predictive performance for total counts (median *r* = 0.69) and profile shape (median JSD = 0.62) across held-out peaks, on par with the K562 models (**Extended Fig. 2b-c**).

To evaluate the efficacy of bias correction, we first examined the bias-corrected ATAC-seq profile predicted by the GM12878 ATAC-seq ChromBPNet model at the exemplar ATAC-seq peak bound by CTCF (**Fig. 2b**). The bias-corrected predicted profile revealed a distinctive, latent footprint tightly bookending the CTCF motif, obscured in the measured and uncorrected predicted profiles. Bias-corrected profile contribution scores exclusively highlighted the CTCF motif and two ZBTB7A motifs with weak repressive (negative contribution scores) effects that were not discernable in uncorrected profile contribution scores. Bias correction also suppressed spurious flanking bases representing Tn5 bias that were highlighted by uncorrected profile contribution scores. Examination of analogous tracks from the DNase-seq ChromBPNet models showed that DNase-I bias produced milder distortions of predicted profiles and contribution scores at this locus (**Extended Fig. 5b**).

Comparison of TF-MODISCO motifs derived from uncorrected and bias-corrected profile contribution scores across 30,000 randomly sampled GM12878 ATAC-seq peaks showed that bias correction completely eliminated Tn5 bias motifs that dominated uncorrected profile contribution scores and clearly revealed motifs of key LCL TFs, consistent with motifs derived from count contribution scores (**Fig. 2e**). DNase-seq models also showed complete elimination of the DNase-I bias motifs from profile contribution scores (**Extended Fig. 5c**).

Marginal footprinting showed that bias-corrected profiles were insensitive to Tn5 and DNase-I bias motifs, in contrast to uncorrected profiles (**Fig. 2f**, **Extended Fig. 5d**). Furthermore, bias-corrected marginal ATAC-seq footprints of predictive TF motifs lacked high-frequency distortions present in uncorrected footprints that strongly matched enzyme bias interference at these motifs, and revealed clear differences in footprint morphology (depth and shapes) between TF motifs (**Fig. 2g**). Marginal footprints from bias-corrected profiles were absent at GATA1 motifs, used as a negative control since GATA TFs lack expression in GM12878. In contrast, uncorrected profiles produced detectable GATA1 marginal footprints, identical to those derived from bias model profiles. Analogous analysis of TF footprints from DNase-seq ChromBPNet models revealed discernible but subtle effects of bias correction (**Extended Fig. 5e**).

These results demonstrate ChromBPNet’s effectiveness at deconvolving and eliminating enzyme sequence preferences, revealing high fidelity, bias-corrected chromatin accessibility profiles, TF footprints and the underlying sequence drivers at base-resolution.

### ChromBPNet outperforms alternative approaches for enzyme bias correction

Next, we leveraged ChromBPNet’s modular design for bias correction to systematically evaluate alternative approaches for measuring and correcting enzyme bias.

First, we compared Tn5 bias models trained on cleavage profiles from naked genomic DNA versus chromatinized background regions from ATAC-seq experiments^79^. TF-MODISCO motifs from the naked DNA bias model’s profile contribution scores showed stronger A/T content at specific positions compared to those derived from chromatinized background despite being trained on compositionally matched regions, indicating differences in learned enzyme sequence preferences (**Extended Fig. 4a**). In contrast, chromatin background models from different cell-lines (HepG2 and GM12878) did not show any discernible motif differences (**Extended Fig. 4a**). Further, ChromBPNet models trained on GM12878 ATAC-seq data augmented with the naked DNA bias model produced TF-MODISCO motifs from bias-corrected profile contribution scores contaminated with strong G/C bias (**Extended Fig. 4b**). In contrast, training the GM12878 model with a HepG2 chromatin-background bias model avoided this contamination. Marginal footprinting at Tn5 bias motifs and TF motifs showed that naked DNA bias-corrected profiles performed incomplete bias correction compared to chromatin-background bias-corrected profiles (**Extended Fig. 4c,d**).

We verified these results by comparing to the recent seq2PRINT method that also aims to use a neural network trained on a higher quality naked DNA transposition experiment to learn and correct Tn5 biases^128^. Comparing their observed naked DNA Tn5 transposition profiles with chromatin background GM12878 ATAC-seq profiles across 1,000 high-coverage 100bp BAC regions revealed substantial variability of correlation (Pearson *r*:-0.011 to 0.93, mean = 0.56; Spearman *ρ*:-0.013 to 0.90, mean = 0.39), suggesting that naked DNA profiles do not consistently capture chromatin background bias (**Supp Fig. 1a**). While predicted profiles from the seq2PRINT naked DNA bias model performed comparably to our ChromBPNet chromatin background model when evaluated against naked DNA profiles, it consistently performed worse against chromatin background profiles (**Supp Fig. 1a**). Furthermore, ChromBPNet models trained on keratinocyte and GM12878 ATAC-seq data using the seq2PRINT naked DNA bias model failed to correct Tn5 bias. TF-MoDISco applied to the corrected profile contribution scores revealed persistent GC bias and uncorrected Tn5 bias motifs (**Supp Fig. 1b-c)**. In contrast, experiment-matched chromatin background models achieved superior bias correction with no detectable artifacts (**Supp Fig. 1d, Figure 2e**). These results demonstrate that Tn5 bias manifests in an experiment-specific manner and that chromatin background models matched to each experiment are necessary for successful bias correction.

Our results support previous findings that naked DNA enzyme cleavage profiles do not fully capture the enzyme sequence preferences in chromatin, and that chromatin-background bias models enable superior bias correction^79^. While our results suggest transferability of chromatin-background bias models across cell-lines, we advise performing careful verification of bias correction via model interpretation, as presented above, since unknown differences in experimental protocols and other dataset-specific artifacts could impact the efficacy of bias correction.

Next, we used ChromBPNet’s bias-correction verification framework to evaluate established bias correction methods, TOBIAS^78^ and HINT-ATAC^79^. Expanded BPNet models (i.e. ChromBPNet without the bias-correction module) trained directly on GM12878 ATAC-seq and DNase-seq profiles pre-corrected by these methods showed only partial bias correction in profile predictions and contribution scores at exemplar loci, TF-MODISCO motifs derived from profile contribution scores, and marginal footprints (**Fig. 2b,2e-g** for ATAC-seq, **Extended Fig. 5b-e** for DNase-seq, **Supp Files 2**). ChromBPNet models augmented with TOBIAS’s bias model also failed to fully eliminate enzyme bias, underperforming even simple baseline CNN bias models trained on chromatin background (**Fig. 2b,2e-g** for ATAC-seq, **Extended Fig. 5b-e** for DNase-seq, **Supp Files 2**).

In summary, ChromBPNet models augmented with BPNet bias models trained on chromatin background provide superior bias correction compared to alternative approaches.

### ChromBPNet substantially improves concordance between ATAC-seq and DNase-seq profiles & predictive sequence features

While DNase-seq and ATAC-seq experiments identify largely overlapping peak regions in matched biosamples, their base-resolution cleavage profiles often differ substantially (**Fig. 3a**). We hypothesized that these differences stem from profile sparsity and enzyme-specific biases distorting profile shapes. Given that ChromBPNet predicts denoised, bias-corrected profiles at base resolution, we expected higher concordance between predicted profiles from independently trained ATAC-seq and DNase-seq models compared to measured profiles.

**Figure 3:**
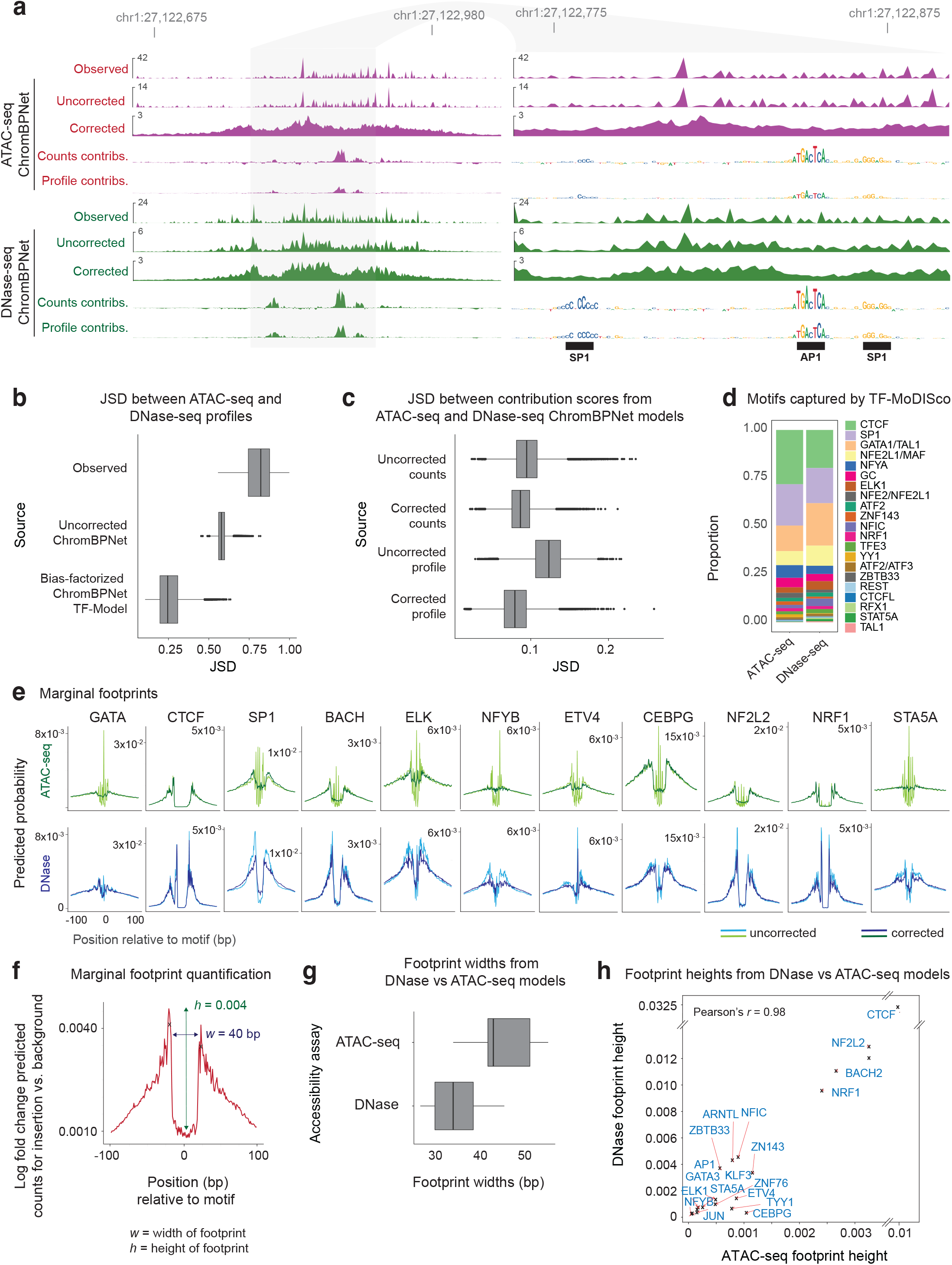
ChromBPNet’s bias correction improves concordance between ATAC-seq and DNase-seq experiments. **(a)** Exemplar locus at chr1:27,122,675-27,122,980 showing several tracks from top to bottom: observed K562 ATAC-seq profile; uncorrected predicted ATAC-seq profile from ATAC-seq ChromBPNet; bias-corrected predicted ATAC-seq profile; bias-corrected count and profile contribution scores from ATAC-seq ChromBPNet highlighting AP1 and SP1 motifs (on right zoomed in panel); observed K562 DNase-seq profile shows poor similarity to observed ATAC-seq profile; uncorrected predicted DNase-seq profile from DNase-seq ChromBPNet; bias-corrected predicted DNase-seq profile from DNase-seq ChromBPNet is more similar to bias-corrected ATAC-seq profile but shows narrower footprints; bias-corrected count and profile contribution scores from DNase-seq ChromBPNet highlighting AP1 and SP1 motifs that are highly concordant with contribution scores derived from ATAC-seq ChromBPNet; **(b)** Distribution of discordance (measured using JSD) between pairs of observed K562 DNase-seq and ATAC-seq profiles over peak regions, and predicted (uncorrected and bias-corrected) ATAC-seq and DNase-seq profiles. **(c)** Distribution of discordance (JSD) between pairs of different types of contribution score profiles from ATAC-seq and DNase-seq ChromBPNet models over peak regions. From top to bottom: count contribution scores from uncorrected ChromBPNet; count contribution scores from bias-corrected ChromBPNet; profile contribution scores from uncorrected ChromBPNet; profile contribution scores from bias-corrected ChromBPNet. Bias-correction significantly improves concordance of corrected profile contribution scores. **(d)** Relative frequency of high-confidence predictive motif instances (seqlets) of TF-MODISCO motifs derived from bias-corrected profile contribution scores derived from K562 ATAC-seq and DNase-seq ChromBPNet models show high similarity. **(e)** Comparison of marginal footprints at TF-MODISCO motifs derived from uncorrected and bias-corrected normalized probability profiles from K562 ATAC-seq (top) and DNase-seq (bottom) ChromBPNet models (y-axis scales are the same for top and bottom panels of each column). **(f)** Schematic showing definition of height and width of marginal footprints. **(g)** Distribution of widths of bias-corrected marginal footprints at TF-MODISCO motifs from ATAC-seq and DNase-seq ChromBPNet models shows that DNase-seq produces narrower footprints. **(h)** In contrast, the heights of the bias-corrected marginal footprints from DNase-seq and ATAC-seq models are highly correlated across TF motifs.

To test this hypothesis, we compared predictions from K562 ATAC-seq and DNase-seq models across 30K randomly sampled overlapping peaks. Uncorrected predicted profiles, which alleviate coverage sparsity but retain enzyme biases, showed improved similarity (JSD = 0.58 ± 0.03, lower is better) compared to measured profiles (JSD = 0.81 ± 0.08) (**Fig. 3b**). Bias-corrected predicted profiles demonstrated substantially higher concordance (JSD = 0.26 ± 0.08).

The improved concordance of bias-corrected predicted profiles extended to the corresponding predictive sequence features highlighted by sequence contribution scores derived from the ATAC-seq and DNase-seq models. Profile contribution scores from bias-corrected models showed much higher similarity (JSD = 0.1 ± 0.03) than those from uncorrected profiles (JSD = 0.15 ± 0.03) (**Fig. 3c**). Count contribution scores remained unaffected by bias correction, supporting our finding that enzyme bias primarily impacts profile shape prediction and associated contribution scores. The improved concordance was not only evident at individual loci (**Fig. 3a**) but also resulted in strong similarity of predictive TF-MODISCO motifs and the proportion of predictive motif instances identified from profile contribution scores of both models (**Fig. 3d**).

Despite improved concordance of bias-corrected profiles and contribution scores, subtle differences persisted between bias-corrected ATAC-seq and DNase-seq footprints at predictive motif instances (**Fig. 3a**). Systematic comparison of marginal bias-corrected footprints at key TF-MODISCO motifs revealed that DNase-seq footprints were consistently narrower (25-35 bp vs. 35-40 bp) and deeper (0-0.0325 vs 0-0.01 probability of cleavage) than ATAC-seq footprints, often showing more refined structure (**Fig. 3e-h**). These differences are consistent with DNase-I’s smaller size (∼31 kDa), direct DNA cleavage mechanism, and higher sensitivity to local chromatin structure, compared to the larger Tn5 transposase (53.3 kDa) used in ATAC-seq^129,130^.

Nevertheless, we observed high correlation of bias-corrected marginal footprint depths between assays across predictive TF motifs (Pearson *r* = 0.98, **Fig. 3h**). This demonstrates that ChromBPNet’s bias-corrected footprints from both ATAC-seq and DNase-seq consistently reveal differences in TF residence times and occupancy, ranging from deep (CTCF, NRF1, BACH) to moderate (GATA, ELK, SP1) and shallow (NFYB, STA5) footprints^131,132^.

Thus, ChromBPNet eliminates sequence-specific enzyme preferences while preserving intrinsic structural differences between Tn5 and DNase-I, revealing highly concordant predictive motif syntax and co-localized footprints from both assays.

### ChromBPNet can impute base-resolution profiles, footprints and predictive sequence motifs from low coverage ATAC-seq and DNase-seq datasets

Sequencing coverage in ATAC-seq/DNase-seq experiments significantly impacts the sensitivity and specificity of typical analysis workflows. For example, traditional footprinting methods often require >100M reads for sensitive and high-fidelity footprint identification at individual loci ^73,74^. While the ChromBPNet models analyzed above were trained on such deeply sequenced bulk datasets, deep sequencing is frequently constrained by cost or library complexity. For instance, rare cell populations identified from single-cell or single-nucleus ATAC-seq experiments may yield shallow pseudobulk samples with only 5-10M reads from as few as 100 nuclei. We sought to investigate whether ChromBPNet models could be successfully trained on such shallow datasets and how read depth affects model performance and interpretation.

To investigate ChromBPNet’s performance as a function of read depth, we created subsampled datasets (250M, 100M, 50M, 25M, 15M, 10M, 5M and 2M reads) from our deepest ATAC-seq dataset (572M reads, GM12878). As expected, base-resolution profiles deteriorated with decreasing read depth (JSDs ranging from 0.1-1.0, median JSD increasing from 0.3 (250M), 0.5 (100M), 0.6 (50M), 0.7 (25M), 0.9 (5M)) (**Fig. 4a,b, Supp Fig. 2a**). At 5M reads, profiles were often too sparse to visually identify peaks or footprints (**Fig. 4a**).

**Figure 4:**
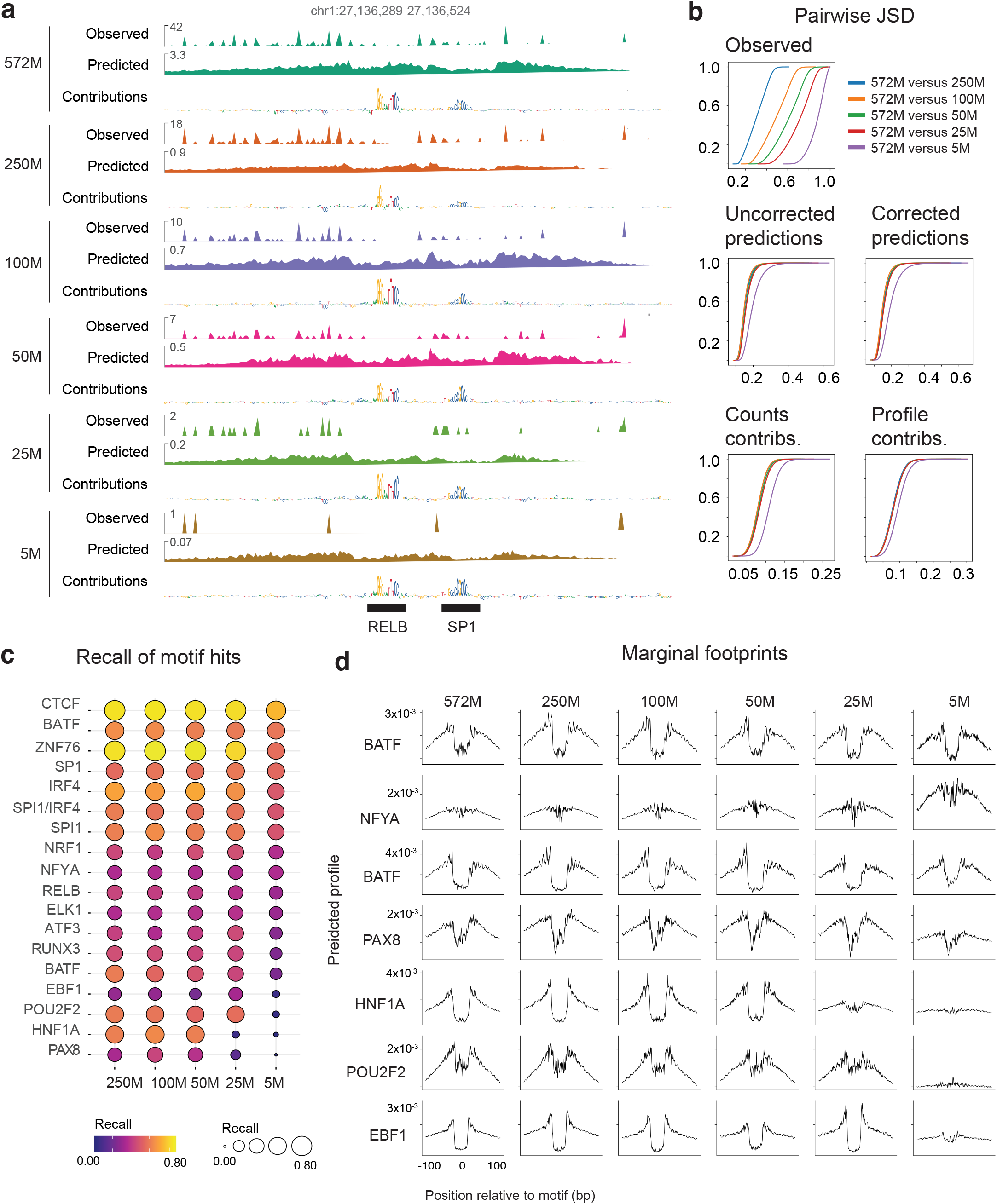
ChromBPNet can impute base-resolution profiles, TF footprints and predictive sequence motifs from low coverage ATAC-seq and DNase-seq datasets. **(a)** Exemplar locus on chr1 (held-out test chromosome) showing observed (Obs) GM12878 ATAC-seq profiles deteriorating with progressively subsampled datasets (250M, 100M, 50M, 25M, and 5M total reads). However, bias-corrected predicted profiles and contribution scores from GM12878 ATAC-seq ChromBPNet models trained on datasets of all read depths are highly concordant. **(b)** Distribution of discordance (JSD) between pairs (maximal read depth vs. progressively subsampled data) of various types of profiles over peak regions. Observed ATAC-seq profiles (which exhibit increasing discordance with reducing read depth). In contrast, uncorrected and bias-corrected predicted profiles, bias-corrected count and profile contribution scores exhibit very similar distributions as low as 25M. Some deterioration is only visible at 5M reads. **(c)** Comparison of recall of predictive instances of top ranking TF-MODISCO motifs in profile contribution scores from the GM12878 ChromBPNet models trained on various subsampled datasets, using the instances identified at 572M reads (full read-depth) as ground truth. Motif recall is high and consistent across most read depths. Rare motifs are lost at low read depths (5M reads). **(d)** Comparison of marginal footprints at representative TF-MODISCO motifs from ChromBPNet models trained on GM12878 ATAC-seq data at different subsampled read-depths.

In contrast, ChromBPNet models trained independently on each subsampled dataset imputed uncorrected and bias-corrected base-resolution profiles with remarkably high fidelity (JSDs ranging from 0.1-0.6, median JSD ranging from 0.147 (250M), 0.145 (100M), 0.153 (50M), 0.156 (25M), 0.190 (5M)) relative to the reference high-depth model, showing visible degradation only at 5M reads (**Fig. 4b, Supp Fig. 2a**). Count and profile contribution scores also remained highly concordant across read depths, with some degradation at 5M reads. Accurate footprint and sequence syntax imputation was evident at individual loci across all read depths (**Fig. 4a**).

To estimate genome-wide recovery of predictive sequence motifs as a function of read depth, we compared predictive instances of TF-MODISCO motifs derived from profile contribution scores from the reference high-depth model to those from models trained on subsampled datasets. As expected, recall rates were higher for top-ranked, more frequent motifs (e.g., IRF, SPI, RUNX) compared to less frequent ones (e.g., PAX, COE, HNF1) (**Fig. 4c, Supp Fig. 2b**). Recall rates remained stable across subsampled models down to 25M reads, below which we observed substantial loss for the least frequent motifs. Recovery of bias-corrected marginal footprints showed similar trends (**Fig. 4d**).

To corroborate these results in single-cell contexts, we trained ChromBPNet on pseudobulk GM12878 scATAC-seq profiles aggregated from progressively fewer cells (11,000 down to ∼100 cells, corresponding to 250M down to 2M reads). These models revealed consistent trends in profile imputation quality and motif recall that closely parallel bulk ATAC-seq read subsampling results (**Methods, Supp Fig. 2**).

Hence, ChromBPNet accurately imputes base-resolution profiles, predictive sequence motifs, and their marginal footprints from moderate coverage (25M reads) chromatin accessibility profiles, comparable to much deeper samples. Even at very low coverage (5M reads), despite some expected information loss, ChromBPNet’s predicted profiles and sequence annotations offer substantial benefits over directly using measured profiles. These results indicate that ChromBPNet can be effectively applied to typical bulk ATAC-seq and DNase-seq datasets, as well as cell-type resolved, pseudo-bulk scATAC-seq profiles, to decipher TF motif lexicons and footprints.

### ChromBPNet reveals compact TF motif lexicons, cooperative composite elements and predictive motif instances that influence chromatin accessibility and TF occupancy

To decipher comprehensive TF motif lexicons that regulate chromatin accessibility in each of the five ENCODE Tier-1 cell-lines, we applied TF-MODISCO to contribution scores from ATAC-seq and DNase-seq ChromBPNet models across all peak regions of each cell-line. Count contribution scores revealed between 26 to 47 predictive motifs supported by over 100 seqlets, while profile contribution scores yielded 49 to 114 predictive motifs across cell lines and assays (**Table 5**, **Supp Files 5**). Further clustering of motifs derived from both count and profile contributions across both assays revealed compact, non-redundant lexicons of predictive motifs in each cell line: GM12878 (51 motifs), HepG2 (60 motifs), IMR90 (47 motifs), H1-hESC (41 motifs), and K562 (47 motifs). These findings are supported by previous reports of a limited repertoire of TFs influencing chromatin accessibility^133^. These included motifs of ubiquitous TFs (e.g. CTCF, SP1 and AP1) and well-known cell-type specific TFs (e.g. HNF4 and FOXA2 in HepG2; IRF4, SPI1, RUNX3, and RELB in GM12878; GATA1-TAL1 in K562; POU5F1-SOX2, ZIC3, TEAD4 and SOX2 in H1-hESC; and FOXL1 in IMR90) (**Fig. 5a; Extended Fig. 6a and 7a-b**).

**Figure 5:**
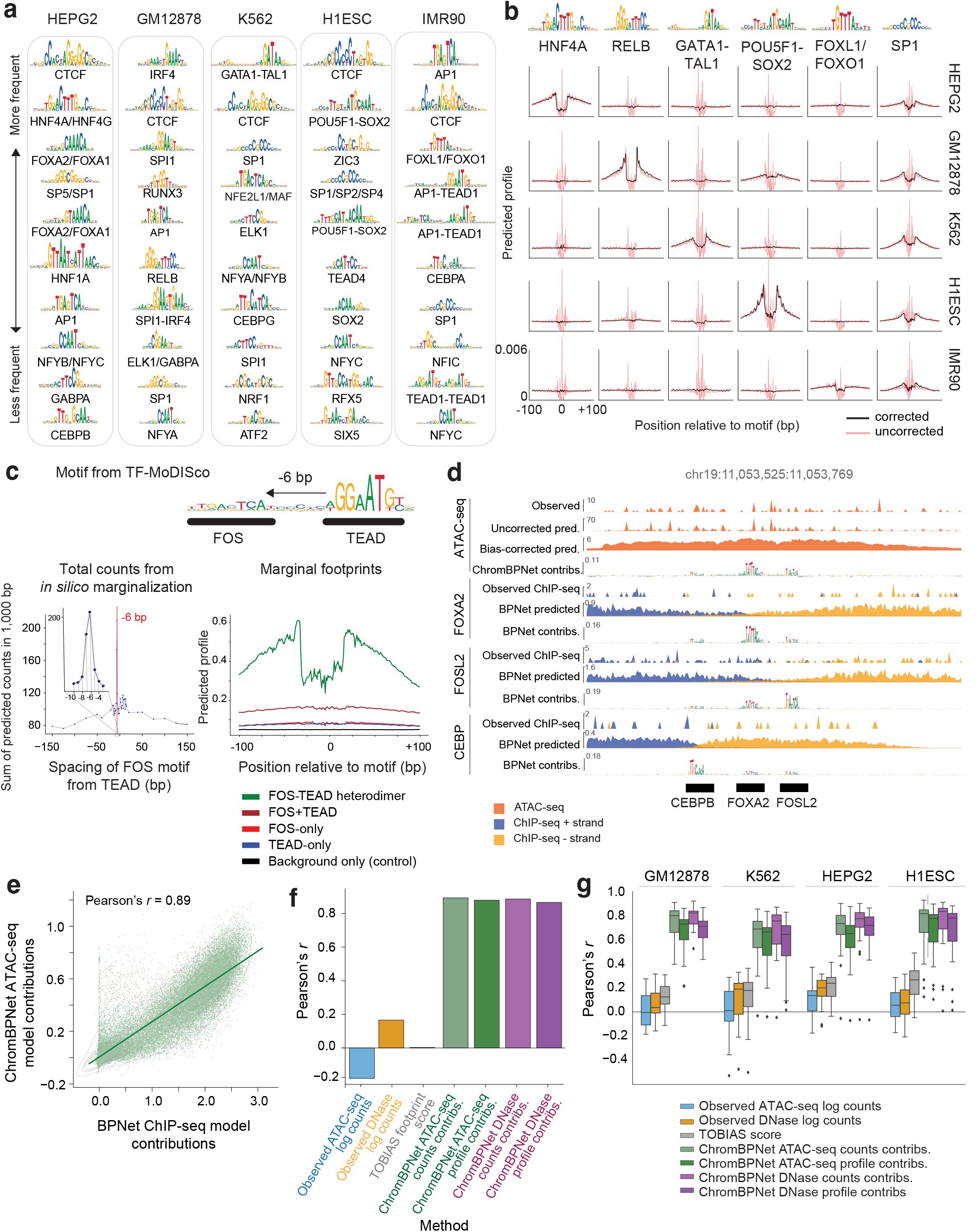
ChromBPNet models of ATAC-seq data reveals compact TF motif lexicons, cooperative composite elements and predictive motif instances that influence chromatin accessibility and TF occupancy. **(a)** Top 10 TF-MODISCO motifs (ranked by number of predictive motif instances) derived from count contribution scores from ATAC-seq ChromBPNet models of diverse cell-lines. **(b)** Marginal footprints of TF-MODISCO motifs using normalized uncorrected (red) and bias-corrected profile predictions (black) from ATAC-seq ChromBPNet models of diverse cell-lines. Bias-correction reveals cell-type specificity of footprints. (**c)** TF-MODISCO applied to the IMR90 ATAC-seq ChromBPNet model reveals a FOS-TEAD composite heterodimer motif (top). Strength (maximum) of marginal footprints of paired FOS and TEAD motifs with variable spacing shows strong cooperative effects only at the fixed 6 bp spacing (bottom-left). Marginal footprints of the FOS motif, TEAD motif, FOS-TEAD composite motif and the sum of FOS and TEAD marginal footprints demonstrates the super-additive cooperative effect of the FOS-TEAD composite (bottom-right). **(d)** Exemplar accessible region in chr19 near the LDLR gene in HEPG2 displaying the observed profile from ATAC-seq and FOXA2, FOSL2 and CEPBP ChIP-seq data, alongside uncorrected and bias corrected predicted profiles, and count contribution scores from ATAC-seq ChromBPNet model and TF ChIP-seq BPNet models. Predictive motif instances from ChromBPNet and BPNet models are in strong agreement. **(e)** Count contribution scores derived from HepG2 ATAC-seq ChromBPNet model and CTCF TF ChIP-seq BPNet model are highly correlated at CTCF motif matches in CTCF peaks overlapping ATAC-seq peaks in HepG2. **(f)** Comparison of correlation of CTCF ChIP-seq BPNet model count contribution scores at CTCF motif matches with various types of motif activity scores derived from ATAC-seq data in HEPG2. Count and profile contribution scores from ATAC-seq and DNase-seq ChromBPNet models substantially outperform observed peak-level ATAC-seq and DNase-seq counts and TOBIAS footprint scores. **(g)** Similar comparisons as in (f) over motif instances of various TF-MODISCO motifs derived from ChromBPNet models of different cell-lines show that motif contribution scores from ATAC-seq and DNase-seq ChromBPNet models show much stronger correlations with motif contribution scores from matched TF ChIP-seq BPNet models compared to the alternative measures (observed peak-level ATAC-seq and DNase-seq counts and TOBIAS footprint scores) of motif activity.

Bias-corrected marginal footprints of motifs verified their cell-type specific or ubiquitous effects on chromatin accessibility. For example, the HNF4A, RELB, GATA1-TAL1, POU5F1-SOX2 and FOXL1 motifs were identified only in HepG2, GM12878, K562, H1-hESC and IMR90 models, respectively, and displayed corresponding cell-type-specific marginal footprints, matching their expression patterns (**Fig. 5b; Extended Fig. 7c**). In contrast, the ubiquitous CTCF and SP1 motifs exhibited consistent and reproducible marginal footprints across models from all five cell lines. Notably, the shape of uncorrected marginal ATAC-seq footprints lacked cell-type specificity, underscoring the importance of bias correction for accurately inferring TF footprints (**Fig. 5b**).

TF-MODISCO motifs included several composite elements—homotypic or heterotypic combinations of TF motifs—exhibiting strict syntax constraints, including spacing, order, and orientation. Examples included the FOS-TEAD composite in IMR90^134^ and IRF1-SPI1 and IRF1-AP1 composites in GM12878^135^ (**Extended Fig. 6b**). These composite elements typically mediate cooperative effects of multiple TFs through precise binding configurations. Using ChromBPNet, we tested whether the models learned these syntax-dependent cooperative effects. For instance, we used the IMR90 ATAC-seq and DNase-seq models to predict and compare bias-corrected marginal footprints of the FOS-TEAD composite element against those of the individual TEAD and FOS motifs. The composite footprint was substantially stronger than the footprints of either motifs as well as their sum, suggesting a super-additive cooperative effect (**Fig. 5c; Extended Fig. 7d**). Furthermore, altering the spacing or orientation of TEAD and FOS motifs abolished the cooperative effect, confirming strict syntax requirements (**Fig. 5c; Extended Fig. 7d**). This behavior was corroborated using BPNet models trained on FOS ChIP-seq data from IMR90 (**Extended Fig. 6c**).

We next used the extensive compendium of TF ChIP-seq datasets in these cell-lines to benchmark the predictive motif instances of TF-MODISCO motifs identified by ChromBPNet against occupancy of putative TFs that might bind the motif. However, directly comparing ChromBPNet derived motif instances to TF ChIP-seq peaks presents several challenges: ChIP-seq peaks (∼200 bp) are wider than individual motifs (4–25 bp), often lack the direct binding motif of the target TF and may contain motifs of multiple TFs that contribute to binding of the target TF. Additionally, ChIP-seq coverage does not provide reliable estimates of each motif’s contribution to TF occupancy. To overcome these limitations, we instead used contribution scores derived from BPNet models trained on the TF ChIP-seq datasets as a high-resolution estimate of their contribution to occupancy of the target TF.

In exploratory analyses of specific enhancer and promoter regions, we found strong agreement between ChromBPNet and BPNet-predicted motif instances. For example, the HepG2 ATAC-seq and DNase-seq ChromBPNet models identified CEBP, FOXA, and AP1 motifs within an accessible enhancer 35 Kb downstream of the LDLR gene, aligning with motif instances highlighted by BPNet models trained on CEBPB, FOXA2, and FOSL2 ChIP-seq data (**Fig. 5d; Extended Fig. 7e**). Similarly, the K562 ATAC-seq and DNase-seq models identified GATA-TAL1 heterodimer motif in the PKLR promoter, matching GATA-TAL1 heterodimer motifs from GATA1 and TAL1 ChIP-seq BPNet models (**Extended Fig. 6d, 7f**).

We then systematically benchmarked all TF-MODISCO motifs derived from ChromBPNet against reference BPNet models of ChIP-seq datasets of putative TFs that could bind these motifs. For each TF-MODISCO motif assigned to putative target TFs for which TF ChIP-seq data was available, we identified motif instances based on sequence similarity to the TF-MODISCO PWMs (*p* ≤ 0.05, MOODS PWM scanner) in all ATAC-seq/DNase-seq peaks overlapping ChIP-seq peaks of the target TF^136^. For each motif instance, we estimated the (1) total count contribution score from ChromBPNet, (2) measured accessibility (total coverage) of the overlapping ATAC-seq/DNase-seq peak, and (3) footprint scores from the TOBIAS footprinting method^78^. We assessed how well these metrics correlated with the reference count contribution scores from BPNet models of the target TF ChIP-seq data.

Using the CTCF motif as a case study, we observed a strong correlation (Pearson *r* > 0.89) between ChromBPNet contribution scores and BPNet-derived scores from CTCF ChIP-seq models in HepG2 (**Fig. 5e**). In contrast, total accessibility scores (r =-0.1) and TOBIAS footprinting scores (*r* = 0.2) exhibited poor correlation (**Fig. 5f**). This pattern was consistent across a broader set of motifs and TF ChIP-seq datasets (**Fig. 5g**). These results indicate that ChromBPNet’s contribution scores accurately capture the quantitative effect sizes of motif instances on TF occupancy, outperforming alternative footprinting approaches.

### ChromBPNet predicts effects of genetic variants on chromatin accessibility across assays, ancestry groups, cell-lines and primary cells

Predictive models of regulatory DNA could potentially enable in silico prioritization and interpretation of non-coding genetic variants that influence regulatory activity. Hence, we evaluated the ability of ChromBPNet models trained using the reference genome to predict the counterfactual effects of genetic variants on chromatin accessibility. Using cell context-specific DNase-seq or ATAC-seq models, we predicted 1 Kb bias-corrected, base-resolution coverage profiles for reference and alternate alleles of any query variant, centered within its 2,114 bp genomic sequence context. From the pair of predicted profiles, we computed several complementary measures of variant effects.

(1) **Log fold-change (logFC)** of total predicted coverage between the reference and alternate alleles, providing a canonical effect size of the variant on local accessibility.
(2) **Active Allele Quantile (AAQ)** is the percentile of the predicted total coverage of the stronger allele relative to the distribution of predicted total coverage across all ATAC-seq/DNase-seq peaks.
(3) **Jensen-Shannon distance (JSD)** between the bias-corrected base-resolution probability profiles of the two alleles, which captures effects on profile shape, such as changes in TF footprints.
(4) **Integrative Effect Size (IES)** is the product of logFC and JSD, capturing both total coverage change and profile shape differences.
(5) **Integrative Prioritization Score (IPS)** is the product of logFC, JSD, and AAQ, integrating all aspects of variant impact to prioritize biologically relevant variants.

We first benchmarked variant effect prediction performance against DNase-seq quantitative trait loci (dsQTLs) identified in 70 lymphoblastoid cell-lines (LCLs) of Yoruban African ancestry^65^, using a curated dataset^137^ of ∼500 statistically significant dsQTLs and a 50x larger negative set of matched control variants. Both sets are restricted to variants that overlap DNase-seq peaks from the LCL cohort, to ensure that the positive set is enriched for likely causal, local cis-dsQTLs while minimizing the inclusion of spurious variants in linkage disequilibrium.

We computed the five types of variant effect scores from ChromBPNet models trained on ATAC-seq and DNase-seq data from a single reference LCL (GM12878) of European ancestry. We first tested these scores in terms of their classification performance relative to the positive and negative sets (**Extended Fig. 8a**). Among individual scores, JSD generally performed on par with or better than logFC, suggesting that changes in profile shape alone were as discriminative as changes in total coverage. Combining JSD and logFC through the Integrative Effect Size (IES) further improved performance. However, the Integrative Prioritization Score (IPS), which incorporates the Active Allele Quantile (AAQ) along with logFC and JSD, yielded the best results, with an average precision (AP) of 0.54 for ATAC-seq and 0.43 for DNase-seq (**Fig. 6a**).

**Figure 6:**
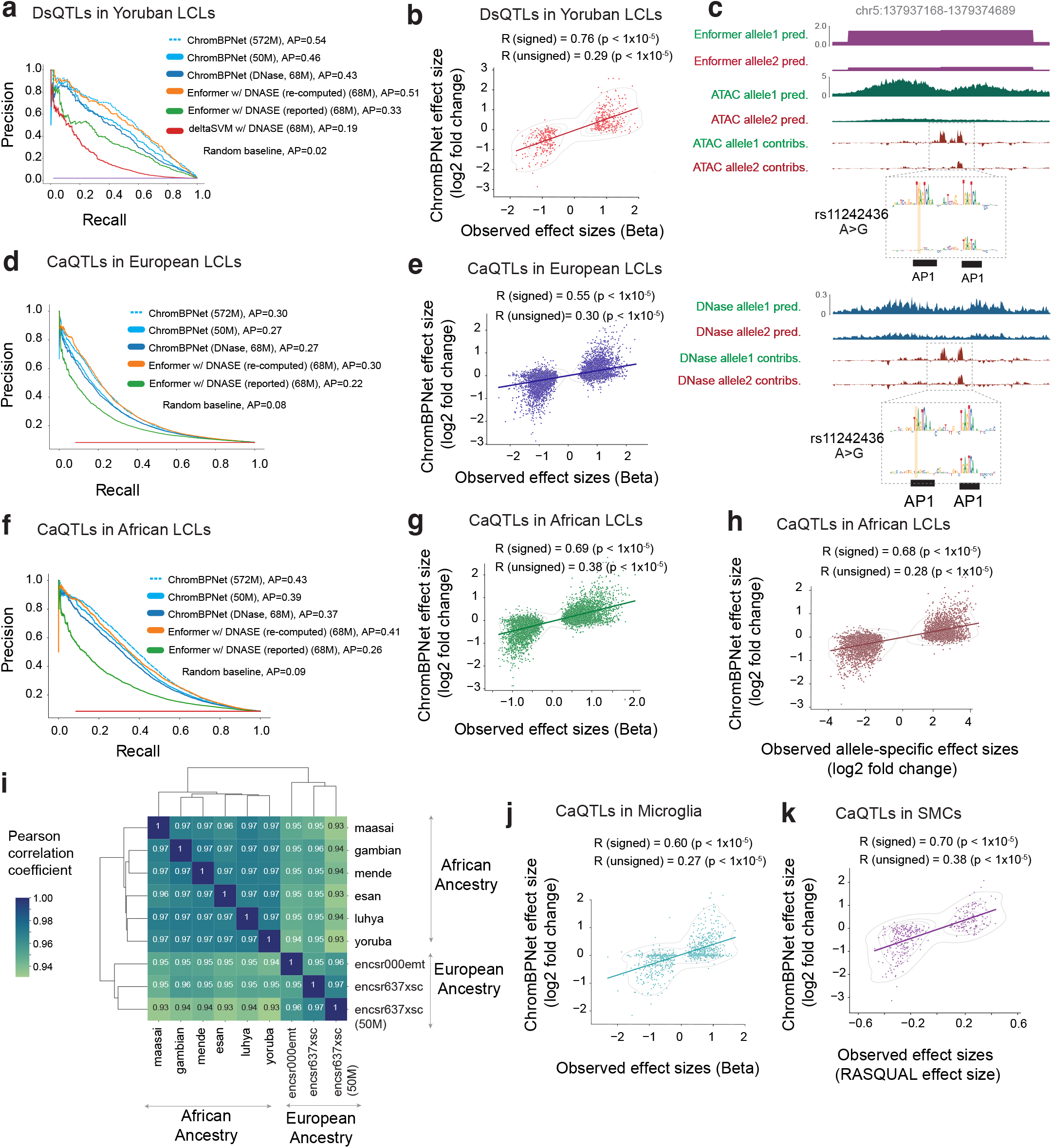
ChromBPNet predicts effects of genetic variants on chromatin accessibility across assays, ancestry groups, cell-lines and primary cells. **(a)** Comparison of variant classification performance (Precision-recall curves and average precision (AP)) of different models (gkmSVM, ChromBPNet and Enformer) trained on GM12878 DNase-seq or ATAC-seq at discriminating significant DNase-seq QTLs (dsQTLs) in Yoruban lymphoblastoid cell-lines (LCLs) from controls variants. ChromBPNet is highly competitive with Enformer and substantially outperforms gkmSVM. **(b)** The observed effect sizes (x-axis, regression betas) of significant dsQTLs are significantly correlated with GM12878 ATAC-seq ChromBPNet’s predicted effect sizes (y-axis, log fold changes). **(c)** Example locus around dsQTL SNP rs11242436. Tracks displayed include bias-corrected predicted profiles from ChromBPNet models trained on GM12878 ATAC-seq and DNase-seq data and predicted profiles from Enformer’s GM12878 DNase-seq head for both alleles. Also shown are ChromBPNet’s contribution scores for both alleles illustrating the disruption of an AP1 motif by the variant and a cooperative effect on another adjacent AP1 motif. **(d)** Comparison of variant classification performance of ChromBPNet and Enformer trained on GM12878 DNase-seq or ATAC-seq data at discriminating significant ATAC-seq QTLs (caQTLs) in European LCLs from control variants. **(e)** The observed effect sizes (x-axis, regression betas) of significant European LCL caQTLs are significantly correlated with GM12878 ATAC-seq ChromBPNet’s predicted effect sizes (y-axis, log fold changes). **(f)** Comparison of variant classification performance of ChromBPNet and Enformer trained on GM12878 DNase-seq or ATAC-seq data at discriminating significant ATAC-seq QTLs (caQTLs) in African LCLs from control variants. **(g)** The observed effect sizes (x-axis, regression betas) of significant African LCL caQTLs are significantly correlated with GM12878 ATAC-seq ChromBPNet’s predicted effect sizes (y-axis, log fold changes). **(h)** The observed allelic imbalance of ATAC-seq coverage (x-axis, log fold changes) at heteroyzgous variants in the African caQTL LCL cohort is significantly correlated with GM12878 ATAC-seq ChromBPNet’s predicted effect sizes (y-axis, log fold changes). **(i)** Predicted effect sizes (log fold changes) for the significant African LCL caQTLs from ChromBPNet models trained on ATAC-seq data from reference LCLs from different ancestry groups/subgroups (each row/column in the matrix) are highly correlated. **(j)** The observed effect sizes (x-axis, regression betas) of significant caQTLs in microglia are significantly correlated with predicted effect sizes from a ChromBPNet model trained on a reference microglia scATAC-seq pseudobulk dataset (y-axis, log fold changes). **(k)** The observed effect sizes (x-axis, effect sizes from the RASQUAL method) of significant caQTLs in coronary artery smooth muscle cell-lines are significantly correlated with predicted effect sizes from a ChromBPNet model trained on a reference caSMC scATAC-seq pseudobulk dataset (y-axis, log fold changes).

We suspected this performance gap between the ATAC-seq and DNase-seq could be simply due to the substantial difference in read depth of the ATAC-seq (572M reads) and DNase-seq (68M reads) datasets used to train the models. To investigate further, we rescored the variants with ChromBPNet models trained on subsampled ATAC-seq datasets (250M, 100M, 50M, 25M and 5M). AP remained above 0.5 for the 100M reads model and dropped to 0.46 at 50M reads, which is comparable to that of the 68M DNase-seq model (AP = 0.43) (**Fig. 6a, Extended Fig. 8b**). Below 50M reads, AP declined more sharply, reaching 0.29 at 5M reads (**Extended Fig. 8b**). The ATAC-seq and DNase-seq ChromBPNet models substantially outperformed the gkm-SVM model (AP = 0.19) trained on the same 68M DNase-seq dataset. Surprisingly, both ChromBPNet models also substantially outperformed published variant effect scores on GM12878 DNase-seq from the Enformer model (AP = 0.33), which is a state-of-the-art, long-context (200 Kb), multi-task transformer model trained on chromatin accessibility, TF binding and gene expression tracks from diverse cell types in human and mouse, including the 68M GM12878 DNase-seq dataset (**Fig. 6a**, **Extended Fig. 8b**).

Next, we evaluated the correlation between predicted logFC effect sizes from ChromBPNet models and the observed effect sizes of the significant dsQTLs in the positive set. The ChromBPNet models trained on ATAC-seq (*r* = 0.76 at 572M, *r* = 0.73 at 50M) and DNase-seq (*r* = 0.73 and at 68M) outperformed the gkm-SVM model (*r* = 0.73 at 68M DNase-seq) (**Fig. 6b, Extended Fig. 8c**). All ChromBPNet models also substantially outperformed Enformer (*r* = 0.56 at 68M DNase-seq reads) (**Extended Fig. 8c**). Remarkably, subsampled ATAC-seq models maintained high correlations even at very low read depths (*r* = 0.67 at 5M reads). Similar trends were observed when comparing unsigned predicted and measured effect sizes.

We suspected that Enformer’s poor performance relative to ChromBPNet may be due to potential flaws in their published variant scores. Upon investigation, we realized that Enformer’s reported variant scores were computed by comparing the total coverage of predicted profiles spanning ∼100 Kb around each variant for both alleles, which can greatly diminish the local effect of variants. To address this issue, we recomputed Enformer variant effect scores as the allelic log fold change of coverage over local 2 Kb profiles around each variant. These revised local variant effect scores showed substantial improvements in variant prioritization (AP = 0.53 at 68M DNase-seq), outperforming matched ChromBPNet models (AP = 0.43 at 68M DNase-seq, AP = 0.46 at 50M ATAC-seq). dsQTL effect size prediction also improved substantially (*r* = 0.73, |*r*| = 0.20 at 68M DNase-seq), but slightly underperformed matched ChromBPNet models (*r* = 0.73, |*r*| = 0.25). However, ChromBPNet trained on the higher coverage (572M) ATAC-seq data (AP = 0.54, *r* = 0.76, |*r*| = 0.29) outperformed Enformer, highlighting the importance of high quality, deeply sequenced training data.

Beyond predicting effects of variants on accessibility, ChromBPNet can also provide insights into the disrupted regulatory syntax, via contrasts of contribution scores of reference and alternate allele sequences. For example, for dsQTL rs11242436, both ChromBPNet and Enformer predicted strong effects on accessibility. Comparing the contribution scores of the reference and alternative sequences derived from the DNase-seq and ATAC-seq ChromBPNet models showed that the “A” allele not only directly disrupts a predictive AP1 motif instance but also reduces the contribution of a neighboring AP1 motif, suggesting disruption of cooperative syntax (**Fig. 6c**). Additionally, ChromBPNet’s base-resolution predictions reveal a loss of footprints, providing a more detailed characterization of the variant’s impact compared to Enformer’s binned predictions.

To verify the robustness of our benchmarks, we performed additional comparative evaluations using two other independent, well powered chromatin accessibility QTL (caQTL) studies based on ATAC-seq profiling in ∼100 LCLs of European ancestry^66^ and ∼100 LCLs of diverse African ancestry^67^. The overall conclusions from both benchmarks were consistent with those of the dsQTL benchmark and remained robust across different significance thresholds for defining significant caQTLs (**Fig. 6d-g, Extended Fig. 9a-d**).

The African caQTL study also identified 5,897 heterozygous variants across all individuals with significant intra-individual allelic imbalance of accessibility. Variant effect scores (logFC) from the GM12878 ATAC-seq ChromBPNet model showed strong directional correlation (*r* = 0.68) with the reported allelic imbalance effect sizes (consensus average over all individuals) on par with the cross-individual QTL effect sizes from the same study (*r* = 0.69) (**Fig. 6g,h, Extended Fig. 9f**).

The African caQTL dataset, spanning six ancestry subgroups (Esan, Maasai, Mende, Gambian, Luhya, and Yoruba), enabled assessment of variant effect score robustness across models trained on different reference datasets. We observed very high correlations (*r* = 0.96-0.97) in variant effect scores (logFC and JSD) between all pairs of African subgroup models (**Fig. 6i, Extended Fig. 9e**). Models trained on European LCL reference datasets showed slightly lower but still high correlation (*r* = 0.93-0.95) with all African ancestry subgroup models, suggesting strong cross-ancestry generalization of variant effect predictions.

Finally, we evaluated ChromBPNet models trained on pseudobulk scATAC-seq profiles from two disease-relevant primary cell types, human microglia and coronary smooth muscle cells, against cell-context matched caQTL datasets^69,70^. Using reference ChromBPNet models trained on an independent scATAC-seq pseudobulk dataset of microglia^138^ and models trained on pooled smooth muscle cell (SMC) profiles^70^, we observed strong directional correlations between predicted and reported effect sizes for significant caQTLs (*r* = 0.6 for microglia, *r* = 0.7 for SMCs) (**Figure 6j,k, Extended Fig. 9g,h**).

These comprehensive analyses demonstrate that ChromBPNet models, despite being trained on single-reference ATAC-seq/DNase-seq profiles using only the reference genome and its short local context, can robustly prioritize causal variants and predict variant effects on chromatin accessibility, performing on par with much larger state-of-the-art, multi-task deep learning models. ChromBPNet exhibits strong generalization across assays (DNase-seq and ATAC-seq), QTLs identified across diverse ancestry groups, and cellular contexts, maintaining robust performance in both cell lines and primary cells.

### ChromBPNet predicts effects of sequence variation influencing pioneer TF occupancy, massively parallel reporter activity, complex traits and rare disease

Having established that ChromBPNet can predict variant effects on chromatin accessibility and that motif contribution scores reflect their influence on TF occupancy, we hypothesized that ChromBPNet’s variant effect predictions should correlate with their upstream effects on TF occupancy, particularly for lineage-specific pioneer factors. We tested this hypothesis using a comprehensive binding QTL (bQTL) study that assessed effects of over 1 million variants on genome-wide SPI1 pioneer TF occupancy via a pooled ChIP-seq experiment across 60 Yoruban LCLs^139^. Using GM12878 ChromBPNet models to predict variant effects on DNase-seq and ATAC-seq profiles, we observed that bins of bQTLs with higher statistical significance showed stronger predicted effect sizes on chromatin accessibility, concordant with observed bQTL effect size distributions (**Extended Fig. 10a**).

Using the rs5764238 (C>G) SPI1 bQTL as a representative example, we visualized and compared the allele-specific predicted profiles and contribution scores from both ChromBPNet models to those derived from BPNet models trained on SPI1 ChIP-seq profiles from GM12878 **(Fig. 7a)**. The BPNet model predicted low TF occupancy for the C allele, regulated by a weak flanking SPI1 motif. The G allele significantly increased predicted occupancy by amplifying an overlapping SP1 motif’s contribution score. DNase-seq and ATAC-seq ChromBPNet predictions corroborated these findings, matching the measured bQTL allelic effect direction.

**Figure 7:**
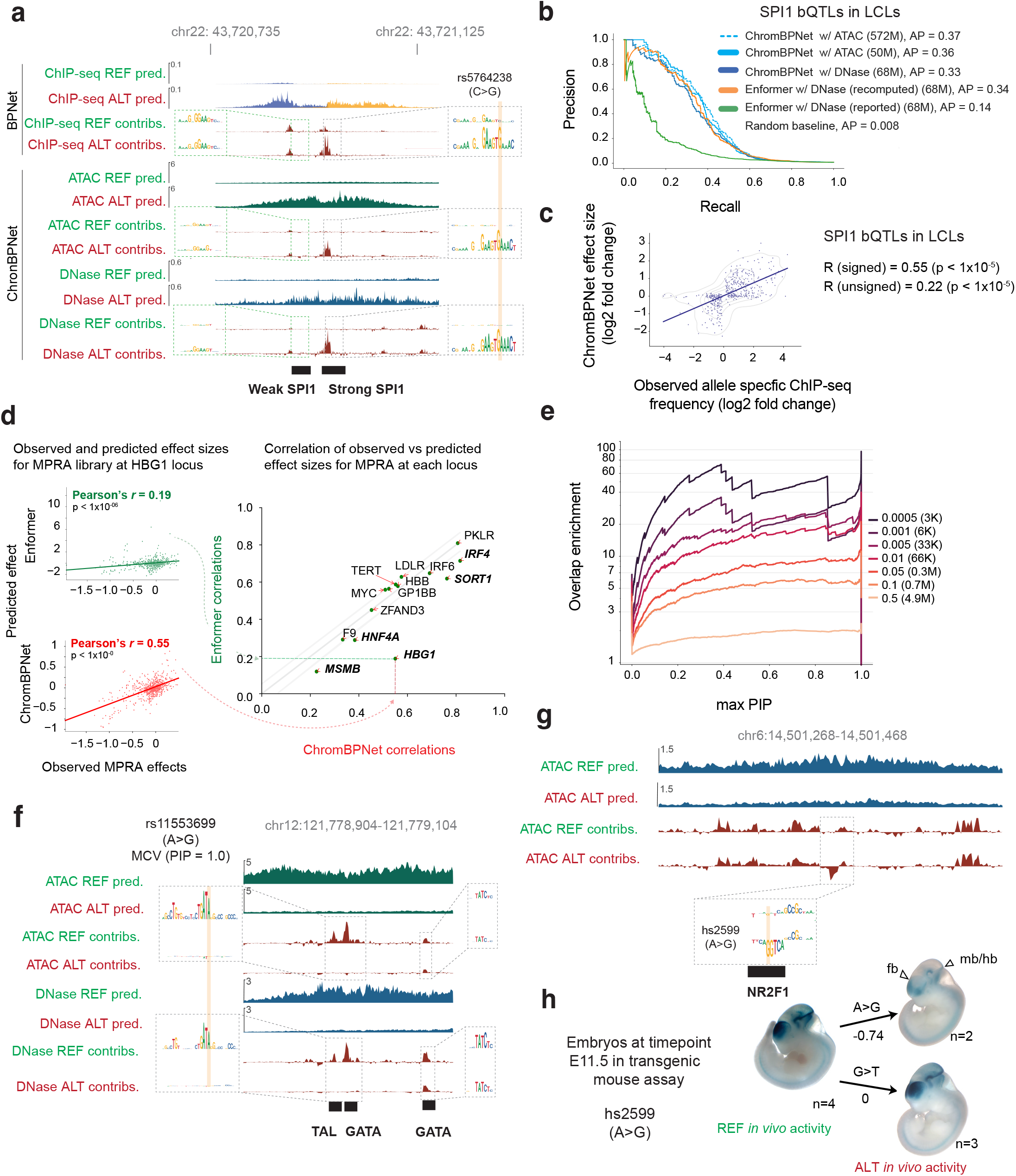
ChromBPNet predicts effects of genetic variants influencing pioneer TF occupancy, massively parallel reporter activity of cRE mutagenesis, complex traits and rare disease. **(a)** Example locus around SNP rs5764238, which is a significant SPI1 binding QTL (bQTL) in LCLs. Predicted profiles and contribution scores from various models are shown. From top to bottom: SPI1 BPNet ChIP-seq model in GM12878, GM12878 ATAC-seq ChromBPNet and GM12878 DNase-seq ChromBPNet. Contributions consistently illustrate the creation of a strong SPI1 motif by the G allele, with a neighboring weak SPI1 motif driving weak accessibility for the C allele. **(b)** Comparison of variant classification performance (Precision-recall curves and average precision (AP)) of different models (ChromBPNet and Enformer) trained on GM12878 DNase-seq or ATAC-seq at discriminating SPI1 bQTLs in LCLs from controls variants shows that ChromBPNet is competitive with Enformer. **(c)** The observed effect sizes (x-axis, allele specific ChIP frequencies) of significant SPI1 bQTLs are significantly correlated with GM12878 ATAC-seq ChromBPNet’s predicted effect sizes (y-axis, log fold changes). **(d)** Benchmark against massively parallel reporter assays (MPRAs) that measure effects of saturated mutagenesis of various cREs (enhancers and promoters) in various cell-lines. The cREs are named based on the nearest gene locus. For each cRE mutagenesis experiment, Pearson correlation is computed between the measured MPRA effect and the predicted effect from DNase-seq ChromBPNet models trained in the closest matched cell-line, over all mutants. Pearson correlation was also estimated using predicted effects from Enformer DNase-seq heads in the closest matched cell-lines. The scatter plot on the right compares the Pearson correlations of ChromBPNet (x-axis) to the Pearson correlations of Enformer (y-axis) for all cREs. Grey lines flanking the diagonal represent +/-0.5 from the diagonal. ChromBPNet outperforms Enformer at several cREs (marked in bold, *IRF4, SORT1, HNF4A, MSMB, HBG1*). The scatter plots on the left compare the observed MPRA effects (x-axis) against the predicted effects from Enformer (top plot) or ChromBPNet (bottom plot) for all mutants of the HBG1 cRE. **(e)** Benchmark testing overlap enrichment (y-axis) of fine mapped variants from GWAS loci associated with red blood cell traits that pass various posterior probability (PIP) thresholds (x-axis) with variants predicted to have strong effects (at various thresholds) by ChromBPNet model trained on K562 ATAC-seq data. Each curve corresponds to a different ChromBPNet effect size threshold (the right panel shows the threshold scores and the number of variants that pass each threshold). Enrichments increase with increasing PIP and more stringent ChromBPNet thresholds. **(f)** Example locus around SNP rs11553699, which is a high-PIP fine mapped variant in a GWAS locus associated with the Mean Corpuscular Volume (MCV) read blood cell trait. Bias-corrected predicted profiles and contribution scores for each allele from K562 ATAC-seq and DNase-seq ChromBPNet model show that the G allele disrupts a GATA motif. **(g)** Example locus around variant hs2599, which is a *de novo* variant from a patient with severe, undiagnosed neurodevelopmental disorder (NDD). Bias-corrected predicted profiles and contribution scores are shown for both alleles from a ChromBPNet model trained on scATAC-seq pseudobulk data from glutamatergic neuron cell-type from human fetal brain tissue. The G allele is predicted to reduce accessibility due to the creation of a repressive NR2F1 motif with strong negative contribution scores. **(h)** Transgenic mouse reporter assay in the E11.5 developmental stage to test the de novo NDD variant shows mice with the activating A allele showing strong enhancer activity in various brain regions, consistent with our model’s predictions in (g). In contrast, mice with the alternate G allele showed significantly reduced enhancer activity, in the forebrain (fb), hindbrain (hb), eyes and midbrain (mb).

We evaluated discriminative performance using 447 high-confidence bQTLs against a negative set (125x larger) of control variants from the African caQTL benchmark. This negative set was carefully chosen to exclude variants that were not SPI1 bQTLs but could still act as significant chromatin QTLs, ensuring a fair comparison. The DNase-seq (68M reads) ChromBPNet model (AP = 0.33) substantially outperformed Enformer’s published scores (AP = 0.14) but slightly underperformed revised local effect scores from Enformer (AP = 0.34) (**Fig. 7b**). ATAC-seq ChromBPNet models surpassed Enformer’s revised scores (AP = 0.37) even at matched read depths (**Fig. 7b**). All ChromBPNet models outperformed Enformer in predicting signed and unsigned bQTL effect sizes (*r* = 0.59 for 572M ATAC-seq reads, *r* = 0.58 for 50M ATAC-seq & r = 0.59 for 68M DNase-seq, versus *r* = 0.53 for Enformer revised scores) (**Fig. 7c, Extended Fig. 10c**). These findings remained robust across different bQTL significance thresholds (**Extended Fig. 10b,c**).

We further investigated whether ChromBPNet’s predicted effects of sequence variation on chromatin accessibility related to experimentally measured effects on downstream regulatory activity as measured by massively parallel reporter assays (MPRAs). We used a benchmark dataset from the CAGI5 competition that used MPRAs to profile the effects of over 30,000 single nucleotide substitutions and deletions in 20 disease-associated regulatory elements including nine promoters (e.g., IRF4, IRF6, MYC, SORT1) and five enhancer elements (e.g., TERT, LDLR, F9, HBGI) in relevant cell-lines^140,141^. Using ChromBPNet models trained on ATAC-seq/DNase-seq data in matched cell-lines, we compared ChromBPNet’s predicted effect sizes for each of the variants with measured MPRA scores and benchmarked against Enformer’s predictions. ChromBPNet matched Enformer’s performance within a margin of +/-0.05 (Pearson correlation) for 9 loci and exceeded it substantially for 5 out of 14 loci (**Fig. 7d, Extended Fig. 10d**). At the HBG1 locus where ChromBPNet showed the largest advantage, Enformer’s contribution scores highlighted strong repressive effects for a TTCTTCAT motif which was strongly activating in the MPRA, while ChromBPNet’s contribution scores were more neutral (**Supp Fig. 3**), likely explaining the performance gap between models (**Fig. 7d**).

Next, we tested whether ChromBPNet’s variant effect scores could be used to prioritize potentially causal variants within GWAS loci associated with complex traits and diseases. We focused on a collection of GWAS studies of 9 red blood cell traits for which reliable statistical fine mapping was available^142^. Among these traits, we identified three - Mean corpuscular volume (MCV), Mean corpuscular hemoglobin (MCH) and Mean corpuscular hemoglobin concentration (MCHC) - for which over 50% of heritability was explained by chromatin accessible regions in K562. Using K562 ATAC-seq and DNase-seq ChromBPNet models, we predicted variant effects for 11,916,124 variants including 12,951 variants in loci associated with these three traits. We then estimated the enrichment of overlap of high scoring variants from ChromBPNet (across a range of score thresholds) with fine mapped variants passing various posterior probability (PIP) thresholds (**Fig. 7e, Extended Fig. 10e**). We observed a strong increase in enrichment with higher PIP thresholds across all ChromBPNet score thresholds. Notably, the enrichment became more pronounced at more stringent ChromBPNet score thresholds, reaching up to 70-fold enrichment (at a PIP threshold of 0.4).

ChromBPNet can also provide insights into the regulatory syntax disrupted by fine mapped variants predicted to have strong effects on accessibility. For example, rs11553699 is a fine mapped SNV (PIP = 1) strongly associated with mean corpuscular volume (MCV). Contribution scores of reference and alternate allele sequences from ChromBPNet revealed that the A to G transition disrupts a GATA-TAL1 composite element, representing binding sites of GATA1 and TAL1 which are critical regulators of erythroid development (**Fig. 7f**). These results suggest that ChromBPNet’s variant effect scores can help prioritize and interpret potentially causal variants in GWAS loci.

ChromBPNet’s sequence-based predictions of variant effects are independent of population allele frequencies, enabling evaluation of both common and rare non-coding variants. We previously demonstrated this capability using BPNet models trained on cell-type resolved scATAC-seq pseudobulk data from fetal heart and brain cell types to prioritize de novo variants in patients with congenital heart disease and neurodevelopmental disorders^94,95^. Here, we showcase the utility of ChromBPNet models trained on scATAC-seq pseudobulk profiles from human fetal brain glutamatergic neurons to analyze a *de novo* variant (chr6:14501369 A>G) from a patient in the Deciphering Developmental Disorders (DDD) cohort with severe, undiagnosed neurodevelopmental disorder^143,144^. The variant lies within a strong scATAC-seq peak in the reference glutamatergic neuron sample. ChromBPNet predicts the risk allele (G) to decrease chromatin accessibility by 40% (**Fig. 7g**). Further, contribution scores of reference and alternate sequences reveals that this reduction stems from the risk allele creating a binding site for the NR2F1 repressor with strong negative contribution scores, which is known to suppress alternative cell fates during neuronal specification and glutamatergic neuron development^145,146^. The variant lies between the CD83 (384 Kb) and JARID2 (748 Kb) genes, with JARID2 being the likely target given its expression in glutamatergic neurons, critical role in neural development through PRC2 and previous associations with autism and intellectual disability^147–150^.

To validate these predictions, we tested both alleles in transgenic mouse reporter assays at embryonic day E11.5. The risk allele abolished enhancer activity in the forebrain, hindbrain and eyes, while reducing activity in the midbrain (**Fig. 7h**). As a control, we tested another nearby DDD de novo variant (chr6:14501675 G>T) that ChromBPNet predicted would not affect accessibility (**Extended Fig. 10f)**. This control variant showed no effect on enhancer activity, supporting ChromBPNet’s ability to distinguish functional from neutral variants (**Fig. 7g**).

Overall, our analyses suggest that despite training ChromBPNet models only on chromatin accessibility profiles, their variant effect scores generalize to upstream and downstream regulatory layers including TF occupancy and reporter activity and for prioritization of fine mapped variants in GWAS loci and *de novo* variants in rare disease.

## DISCUSSION

ChromBPNet combines the predictive power of deep learning, the precision of footprinting methods, and systematic bias correction to enhance inference of regulatory sequence syntax and genetic variation.

### ChromBPNet introduces several technical innovations that enhance predictive models of chromatin accessibility

ChromBPNet directly models raw base-resolution coverage profiles without requiring ad-hoc pre-processing or normalization. Its bias-factorized architecture and loss functions, combined with stage-wise residual training, implicitly account for profile sparsity while disentangling enzyme preferences from regulatory syntax. This model design was guided by systematic model interpretation, revealing insights that performance metrics alone would miss. By separately modeling total coverage and profile shape, we discovered through DeepLIFT, TF-MODISCO, and marginal footprinting analyses that initial models learned confounding enzyme biases that affected profile shape but not total coverage.

ChromBPNet’s modular design also facilitates systematic evaluation of different bias models and external bias correction strategies within a predictive framework. Rigorous benchmarks using our integrated interpretation framework demonstrate that BPNet-like CNNs trained on background chromatin accessibility profiles consistently outperform established methods like HINT-ATAC and TOBIAS, as well as CNN models such as seq2PRINT trained on naked DNA^128,79^. While chromatin-background bias models can usually transfer between experiments using identical protocols, experiment-specific bias models guarantee automatic adaptation to different assays, protocol variations and experiment-specific artifacts without requiring any additional data collection. To ensure robust use across diverse experimental settings, the ChromBPNet package not only provides carefully validated pretrained reference bias models but also includes checkpoints with diagnostic reports for vetting bias correction.

Comparative analysis of ATAC-seq and DNase-seq ChromBPNet models support the robustness and generalizability of our bias correction approach. Despite the distinct enzymatic biases of these assays, after bias-correction, the models predict highly consistent predicted profiles, contribution scores and motifs. Bias-corrected marginal footprints preserve the structural differences between assays while showing strong correlation in footprint depth across TF motifs.

### ChromBPNet improves interpretation of sequence determinants of chromatin accessibility

ChromBPNet provides insights into a fundamental question about which motifs mediate the influence of TFs on chromatin accessibility across cellular contexts. Models from multiple cell-lines consistently reveal that a relatively compact lexicon of both cell-type specific and ubiquitous motifs, including composite elements, is sufficient to accurately predict both overall accessibility and base-resolution profiles, supporting earlier findings that a limited set of TFs directly regulate chromatin accessibility^133^. While some motifs map to established pioneer factors like FOXA2 in HepG2, GATA1 in K562, and OCT4-SOX2 in H1-ESCs, others may prove to be novel pioneers or influence accessibility through alternative mechanisms. For example, in our recent study of Drosophila embryogenesis, ChromBPNet models applied to dynamic chromatin accessibility profiles reveal distinct roles and hierarchical relationships between pioneer factors like Zelda and context-dependent activators, each governed by specific cis-regulatory rules^112^.

ChromBPNet also enables systematic in silico analysis of how regulatory syntax influences accessibility through precise perturbation of motif combinations, affinity, spacing, and orientation. Our analysis of the FOS-TEAD composite element demonstrates how specific spacing and orientation constraints between motifs produce super-additive effects on accessibility^134^. ChromBPNet can also reveal insights into how TF motif affinity and concentration jointly regulate accessibility. In Drosophila embryogenesis, ChromBPNet revealed that Zelda pioneers accessibility proportional to its motif affinity^112^. In our recent fibroblast reprogramming study, ChromBPNet revealed how supraphysiological concentrations of reprogramming factors engage cryptic low-affinity sites flanked by AP1 motifs, redirecting AP1 away from somatic regulatory elements^151^. Similarly, in cranial neural crest cells, ChromBPNet uncovered how low-affinity TWIST1 motifs drive dosage-sensitive responses through TF-nucleosome competition, while SOX9 dosage sensitivity is mediated through AP-1 motifs^152^.

Further, ChromBPNet not only enables precision identification of predictive motif instances in accessible cREs but also helps quantify their effects on accessibility. We demonstrate that contribution scores of predictive motif instances strongly reflect their direct effects on TF occupancy, substantially outperforming traditional footprinting approaches. ChromBPNet’s predicted motif instances and effect sizes are also strongly corroborated by systematic motif mutagenesis experiments via CRISPR VariantFlowFISH experiments in human cell lines and large-scale in vivo reporter experiments in transgenic mouse models^153,154^. Key insights from additional studies are summarized in a Supplementary Note in Methods Section 1.27.

### Our study advances both the methodology and benchmarking standards for predicting and interpreting regulatory genetic variation

We introduce novel measures of variant effects that complement traditional measures of allelic imbalance of total accessibility (log-fold changes), including the Jensen-Shannon Distance (JSD) for changes in base-resolution profile shape and the Active Allele Quantile (AAQ) that calibrates variant effects against genome-wide accessibility. While profile shape changes alone are surprisingly predictive, combining these measures into an integrative score further improves variant prioritization.

To enable rigorous evaluation of variant effect prediction, we contribute carefully curated benchmarking datasets spanning chromatin QTLs from diverse ancestry groups^67,66^, pioneer TF binding QTLs^139^, reporter assays^140,141^, and fine-mapped GWAS variants for red blood cell traits^155^. These benchmarks systematically assess prediction accuracy across multiple regulatory layers, ancestry groups, and experimental assays. Our evaluation framework examines both variant classification and effect size prediction across significance thresholds. To ensure fair comparisons with existing methods like deltaSVM and Enformer we incorporate benchmarks reported in their respective publications. We also identified and corrected flaws in Enformer’s published variant effect scores, significantly improving its performance.

ChromBPNet accurately predicts directional effects of chromatin QTLs robustly across assays and cell types. These results stand in stark contrast to recent reports showing that long-context expression models, including Enformer, struggle to achieve directionally concordant predictions of expression QTL effects^156,157^. Notably, we find that Enformer’s DNase-seq predictions, like ChromBPNet’s, accurately capture directional effects of chromatin QTLs. This suggests both models can effectively predict causal effects of sequence variation on local chromatin accessibility despite training only on reference genome sequences. The challenge in expression QTL prediction likely stems from the difficulty of modeling distal regulatory interactions with current long-context gene expression models^158–160^. Further, the high concordance between predictions from ChromBPNet models trained on data from diverse ancestry groups suggests that the model learns causal regulatory mechanisms independent of the confounding effects of population-specific linkage disequilibrium.

However, despite substantial differences in architecture (6M vs 250M parameters), sequence context (2Kb vs >100Kb), and training data (single task vs massively multi-task), ChromBPNet is highly competitive with Enformer across multiple benchmarks. ChromBPNet generally outperforms Enformer in predicting quantitative effect sizes for chromatin QTLs, binding QTLs, and reporter assays but underperforms at variant classification. However, even this advantage largely disappears when ChromBPNet is trained on higher-depth data, suggesting that Enformer’s edge at variant classification stems less from its longer context window and more from indirect access to reinforcing regulatory signals from TF and histone modification datasets via multi-task training, especially in LCLs. Models trained on deeper datasets are more sensitive to rarer motifs and are likely more effective at learning cooperative effects of motif epistasis within local sequence contexts, thereby improving sensitivity and specificity of variant prioritization (**Fig. 4c-d)**. In contrast, ChromBPNet consistently underperforms AlphaGenome on our benchmarks^116^, suggesting that bonafide improvements in learning distal regulatory effects enable improved variant effect prediction. Nevertheless, ChromBPNet serves a strong baseline that significantly outperforms large annotation-agnostic DNA language models (DNALMs) in zero-shot, probed and fine-tuned settings at predicting accessibility QTLs^161^.

Our evaluations across multiple, independent chromatin QTL benchmark datasets also reveals important considerations for benchmarking variant effect prediction models. While absolute performance varies across datasets and significance thresholds, particularly for variant classification, the relative performance between methods is quite consistent, and quantitative effect size predictions are more comparable across datasets. These variations likely stem from differences in data quality, sequencing depth, peak-calling and QTL analysis approaches. Importantly, we find that restricting chromatin QTLs to variants within peaks is quite critical to enrich for likely causal variants. Without this filtering, performance decreases due to apparent false negatives – variants with strong observed but no predicted effects that likely tag the true causal variants^67^.

Finally, ChromBPNet provides interpretable predictions for disease-relevant variants across the allele frequency spectrum. Predicted variant effects on chromatin accessibility in disease-relevant cell types enrich for fine-mapped common GWAS variants and also help prioritize de novo variants in rare diseases for downstream validation experiments ^95,94^. Further, ChromBPNet delivers stable and robust interpretation of the context-specific regulatory syntax disrupted by prioritized non-coding variants, which not only provides mechanistic insights but can also help weed out potential false positives that may disrupt spurious sequence features.

### Limitations, Caveats and future opportunities

ChromBPNet’s predictions of total accessibility still lag behind pseudo-replicate concordance and the latest long-context models^115,116^, suggesting that while local sequence context can accurately predict profile shape distal context is critical for modeling total accessibility. This means that ChromBPNet’s predictions and derived interpretations are also less reliable outside peak regions. Future improvements may come from stage-wise training approaches that progressively build long-context models from pre-trained local models like ChromBPNet, and training objectives that leverage multiple axes of regulatory variation across the genome and across cell states^119^.

ChromBPNet demonstrates the versatility of BPNet’s architectural backbone and loss functions - minimal modifications enable robust modeling of chromatin accessibility profiles. While we have not extensively explored alternative architectures, recent benchmarks show that CNNs are extremely competitive with alternative architectures^162^. Replacing ChromBPNet’s dilated residual layers with biLSTMs yield marginal performance gains, suggesting that further architecture optimizations are unlikely to deliver major improvements^162^. The key contribution of ChromBPNet lies in the bias-factorized model architecture and bias correction approach. The modular design of the model allows for seamlessly testing alternative backbone architectures. Future work could also explore complementary signal representations beyond count and profile decomposition, such as fragment length-aware V-plots, to enhance TF footprint and nucleosome positioning signal^163,164^.

Our systematic evaluations show that read-depth and data quality can impact several model applications. ChromBPNet effectively imputes profiles from sparse datasets (≥5M reads), enabling analysis of rare cell populations from single-cell data. However, sensitivity to rare motifs and variant classification performance decrease notably below 25M reads, though effect size predictions remain robust. While multi-tasking across auxiliary datasets can boost performance, as seen with AlphaGenome^116^, careful consideration is needed as we have observed that multi-task models trained across diverse contexts learn causally inconsistent features affecting counterfactual predictions and sequence design applications.

ChromBPNet models cis-regulatory sequence determinants of chromatin accessibility in bulk or pseudobulk profiles, requiring separate models for distinct cell states and biosamples. Future extensions could integrate sequence with regulator expression to jointly model cis and trans effects at single base and single cell resolution^118,165^. Such an approach could enable interpretable sequence-anchored “foundation” models trained on large-scale multi-omic single cell atlases.

ChromBPNet is useful to decipher TF motif lexicons, syntax and genetic variants that specifically influence chromatin accessibility. However, it cannot reveal sequence determinants of many other “settler” TFs that bind open chromatin but do not influence accessibility. Models trained on TF ChIP-seq data that integrate sequence with measured accessibility profiles or bias-corrected predicted profiles from ChromBPNet could potentially recover binding sites of settler TFs. Further, in this study, we explicitly focus our analysis on retrieving predictive TF motifs that influence accessibility. ATAC-seq assays also capture nucleosome footprints proximal to accessible sites that fall within ChromBPNet’s receptive field^128,164^. Hence, ChromBPNet could also be used to study the impact of TF motifs and other sequence features on nucleosome occupancy and positioning. More broadly, regulatory DNA sequences encode multiple, intertwined, pleiotropic cis-regulatory codes that have affected different biochemical readouts of regulatory interactions and activity^166^. This means that to comprehensively decipher all layers of the cis-regulatory code and types of regulatory variants, it will be critical to train and interpret predictive models of diverse regulatory and transcriptional readouts within and across diverse cellular contexts.

Comparative analysis of ChromBPNet models of different cell types and states can provide intricate insights into the context-specificity of global and local sequence determinants of chromatin accessibility^151^. However, explicitly fine tuning these models on differential accessibility would enable more sensitive detection of subtle differential effects especially across closely related cellular contexts^119^.

Further, while ChromBPNet is quite effective at predicting directionally concordant effects of regulatory variants on chromatin accessibility, its performance could be further boosted by explicitly exposing the model to genetic variation and matched accessibility profiles across multiple individuals^167^. The extensive variant prediction benchmarks we provide in this study also have several limitations. For the chromatin accessibility QTL classification benchmarks, we conservatively restrict control variants to peak regions to match the context of significant QTLs. Future evaluations should examine other types of control variants, such as those that may create or destroy isolated motif instances in background chromatin that could yield spurious predictions. Our current benchmarks are restricted to local QTLs and do not evaluate the ability to predict distal chromatin QTLs and eQTLs. Future evaluations will also need to test prediction of other classes of variants beyond SNVs, such as indels and STRs^168,169^.

Finally, while ChromBPNet can prioritize trait and disease-relevant non-coding variants that affect chromatin accessibility, these predictions will need to be integrated with measured and predicted effects of variants on diverse molecular and cellular phenotypes in disease-relevant cellular contexts to improve precision and sensitivity of mapping causal variants and understanding their mechanism of action^168,170^.

In conclusion, ChromBPNet provides a lightweight, versatile tool for enhancing the utility of chromatin accessibility data to decipher regulatory syntax and genetic variation at unprecedented resolution. The principled framework we introduce for bias correction and interpretation provides a generalizable template for disentangling technical and biological effects from sequencing-based molecular profiling assays. Early release of a stable, well-documented package implementing ChromBPNet has already enabled broad adoption by the community ^112,151,159,161,166,170–179^. We remain committed to maintaining and extending ChromBPNet to address the challenges outlined above.

## DATA AND CODE AVAILABILITY

### ChromBPNet code repository

https://github.com/kundajelab/chrombpnet Includes code, documentation and tutorials for training, prediction and interpretation

### Data, model and model outputs

For DNase-seq and ATAC-seq data from the ENCODE cell lines (except for H1-hESC ATAC-seq), all data, ChromBPNet models and model-derived outputs are at the ENCODE portal http://encodeproject.org. ENCODE File IDs (ENCIDs) and other metadata are documented in Table 1. H1ESC ATAC-seq data is deposited at GEO https://www.ncbi.nlm.nih.gov/geo/query/acc.cgi?acc=GSE267154. The corresponding ChromBPnet models and model-derived outputs are hosted at the Synapse repository https://www.synapse.org/chrombpnet (syn59449898).

The Synapse repository https://www.synapse.org/chrombpnet (syn59449898) also hosts several other models, datasets and analysis outputs: These include

- Alternative bias correction baseline models (Expanded BPNet, TOBIAS, HINT-ATAC, naked DNA and HEPG2 background bias) in GM12878
- ChromBPNet models trained on GM12878 subsampled datasets (250M to 5M reads)
- AFGR ChromBPNet across ancestries
- ChromBPNet models for microglia and smooth muscle cells scATAC-seq
- Motifs paired with corresponding TF ChIP-seq; corresponding TF ChIP-seq ENCODE IDs and BPNet Model ENCIDs
- Variant effect prediction scores (for dsQTLs, caQTLs, bQTLs, and fine-mapped GWAS variants for blood traits).
- CAGI5 MPRA QTLs were obtained from Kricher, Martin (correspondence) and corresponding Enformer variant effect predictions were obtained from Avsec, Ziga (correspondence). ChromBPNet models used in scoring CAGI5 MPRA QTLs are uploaded to ENCODE or synapse and their corresponding IDs can be found in Table 4.

### Code to reproduce results and generate figures

https://github.com/kundajelab/chrombpnet-figures This code uses the models and outputs listed above

## AUTHOR CONTRIBUTIONS

A.K., A.P., and A.S. conceived the project. A.P. developed ChromBPNet including the bias models and bias correction approaches, trained all models, developed benchmarks and performed all analyses with guidance from A.K. A.S. and S.N. provided conceptual advice during ChromBPNet’s architecture development. A.S. developed preliminary versions of the expanded BPNet and ChromBPNet models and trained preliminary models. A.P. performed all data preprocessing with help from A.S. S.K trained ChromBPNet models in smooth muscle cells and microglia. A.S.K., V.R., and W.J.G. generated the H1ESC ATAC-seq experimental data. E.K., M.K., and L.A.P. performed transgenic mouse reporter experiments. K.K. and K.A. reprocessed and called caQTLs for the European LCL caQTL dataset and provided peak calls for the African LCL caQTL dataset. A.P. drafted the manuscript and figures with guidance from A.K. A.K. contributed to writing and revisions. All authors read and provided feedback on the manuscript.

## ACKNOWLEDGEMENTS

A.P. was supported by a Stanford Bio-X Fellowship. A.S. was supported by the Stanford BioX Bowes fellowship. This work was supported by NIH grants U01HG009431, U01HG012069 and U24HG012343 awarded to A.K. K.K. and K.A. were supported by the Estonian Research Council (grant no. PSG415). Research was supported by National Institutes of Health (NIH) grant R01HG003988 (to L.A.P.) and conducted at the E.O. Lawrence Berkeley National Laboratory under the US Department of Energy Contract DE-AC02-05CH11231, University of California.

We thank Georgi Marinov and Jacob Schreiber for their feedback on the paper, Austin Wang for providing GC matching speedup code, Aman Patel for assistance with model uploads and docker, Selin Jessa for helping with figure aesthetics, Rosa Ma and Jesse Engrietz for providing the fine-mapped GWAS variants for blood traits. We would like to thank Jennifer Jou, Idan Gabdank and Ben Hitz and other members of the ENCODE DCC for supporting ChromBPNet models at the ENCODE portal. We would like to also thank the ENCODE data production centers particularly the labs of John Stamatoyannopoulos for DNase-seq data, Michael Snyder and William Greenleaf for ATAC-seq data, Michael Snyder and Rick Myers for TF ChIP-seq data. We also acknowledge Stephanie Nevins and Guangwen Gavin from the hESC’s core technical assistance team for help with H1ESC ATAC-seq experiments. Some of the computing for this project was performed on the Sherlock cluster. We would like to thank Stanford University and the Stanford Research Computing Center for providing computational resources and support that contributed to these research results. We would also like to thank all members of the ENCODE and IGVF consortia for their helpful feedback. We would also like to thank all early adopters of ChromBPNet.

## METHODS

### 1.0 ATAC-seq of frozen H1 Human Embryonic Stem Cells (hESC)

Due to the lack of high quality ATAC-seq data for H1 Human Embryonic Stem Cells (hESCs), we performed the ATAC-seq protocol as follows: Cryopreserved cells were thawed using standard procedures and treated with DNase I for 30 minutes to remove extracellular DNA from dead cells, then Ficoll purified. ATAC-seq experiments was performed following the omniATAC protocol. Briefly, ∼70,000 cells were centrifuged at 500 g, then resuspended in 1 mL 1x PBS and centrifuged again. Cells were then resuspended in 50 uL ATAC-RSB-Lysis buffer (10 mM Tris-HCl pH 7.4, 10 mM NaCl, 3 mM MgCl2, 0.1% IGEPAL CA-630, 0.1% Tween-20) and incubated on ice for 3 minutes. Subsequently 1 mL ATAC-RSB-Wash buffer (10 mM Tris-HCl pH 7.4, 10 mM NaCl, 3 mM MgCl2, 0.1% Tween-20) was added, the tubes were inverted several times, and nuclei were centrifuged at 500 g for 5 min at 4C. Transposition was carried out by resuspending nuclei in a mix of 25 uL 2x TD buffer (20 mM Tris-HCl pH 7.6, 10 mM MgCl2, 20% Dimethyl Formamide), 2.5 uL transposase (Illumina) and 22.5 uL nuclease-free H2O, and incubating at 37C for 30 min in a Thermomixer at 300 rpm. Transposed DNA was isolated using the MinElute PCR Purification Kit (Qiagen Cat# 28004/28006), and PCR amplified for a custom number of cycles for each library that yields ∼1/4 of plateau SYBR green fluorescene in a pilot 5 µL qPCR reaction (∼10 cycles) using the usual ATAC-seq settings. The resulting data has been deposited and is publicly available at the Gene Expression Omnibus (GEO) under the accession number GSE267154 (https://www.ncbi.nlm.nih.gov/geo/query/acc.cgi?acc=GSE267154).

### 1.1 Data pre-processing for ATAC-seq and DNase-seq

#### Bulk ATAC/DNase-seq

We download bam files from ENCODE website https://www.encodeproject.org based on the accession ids in Table 1. We start with filtered bams for paired-end datasets and unfiltered-bams for single end datasets. Reads from single end datasets are filtered using SAMtools view (1.7) to remove unmapped reads and mates, nonprimary alignments and reads that failed platform or vendor quality checks (-F 780) or had poor mapping quality (<30 MAPQ score). The total number of final filtered reads are reported in Table 1. The final filtered BAM file is converted to bed format using bedtools ‘bamtobed’ (v.2.29.2) and the reads are shifted using awk (version 4.1.4) to +4 on the positive strand and-4 on the negative for ATAC-seq and +1 on the negative strand for DNase-seq. BigWig tracks containing the strand-specific number of aligned 5′ read ends were generated using bedtools genomecov-bg-5, followed by bedGraph to BigWig conversion using UCSC bedGraphToBigWig v.4.

For ATAC-seq datasets we use the default peak set (pseudoreplicated peaks) from the ENCODE portal. For DNase-seq we start from filtered bams and use caper (V2.1.2) to do peak-calling on the paired-end samples (HEPG2 and K562). For the single-end DNase-seq samples we call peaks directly from the filtered bams using the following macs2 command. *macs2 callpeak-g “hs”-p 0.01 --shift-75 --extsize 150 --nomodel --keep-dup all --call-summits.* We further exclude peaks that overlap with ENCODE black-list regions and rank the macs2 peaks by p-value and cap the number of peaks to 200K.

#### Naked DNA ATAC/DNase-seq

The naked DNA ATAC-seq data was downloaded from the NCBI Sequence Read Archive (SRA) under accession number SRX030445, and the naked DNA DNase-seq data was retrieved with accession numbers SRR1565781 and SRR1565782. These FASTQs were then processed using caper (version 2.1.2) to generate corresponding BAM files. Subsequently, we applied the same processing pipeline used for our bulk datasets to these BAM files, resulting in the creation of bigwig files.

#### Single-cell ATAC-seq (scATAC-seq)

For the smooth muscle cell scATAC-seq analysis, we utilized fragment files available from the Gene Expression Omnibus (GEO) under accession number GSE175621 ^70^. We obtained filtered reads aggregated by cell type and corresponding cell-type-specific peaks directly from the authors of the study. The filtered BAM files were converted to BED format using bedtools’bamtobed’ (version 2.29.2). For ATAC-seq data, reads were shifted using awk (version 4.1.4) by +4 bases on the positive strand and-4 bases on the negative strand. We generated BigWig tracks containing strand-specific counts of aligned 5’ read ends using bedtools genomecov with the-bg and-5 options, followed by conversion from bedGraph to BigWig format using UCSC bedGraphToBigWig (version 4).

For microglia, we accessed raw data from GEO (accession GSE147672). Pre-processing of peaks and raw data followed protocols outlined by ^180^ and the accompanying GitHub repository (https://github.com/kundajelab/alzheimers_parkinsons/tree/master).

#### African ancestry bulk ATAC-seq

To analyze chromatin accessibility patterns across diverse African ancestries, we focused on samples from six populations: Gambian, Luhya, Esan, Maasai, Mende, and Yoruba ^67^. For each ancestry, we selected the five deepest sequenced individual samples, as listed in Table 3 with their corresponding ENCODE IDs. We downloaded these samples from the ENCODE portal and pooled their filtered BAM files. Using caper v2.0.3, we then called overlap peaks on these pooled bams. iles. We then applied the same processing pipeline used for our bulk datasets to these BAM files, resulting in the creation of bigwig files.

#### Non-peaks generation

To generate non-peak regions (or background regions) with GC distribution matching the peak regions, we employed a multi-step process. First, we divided the entire genome into overlapping bins of 2114 bp (the input length for ChromBPNet) with a 1000 bp stride. We then excluded bin regions that intersected with peaks and blacklisted regions. This step yielded a candidate set of negative regions. Finally, we selected candidate negatives (twice the size of the peak set) that closely matched the GC distribution of the peak regions.

### 1.2 Enzymatic shifts in ATAC-seq and DNase-seq

The Tn5 transposase functions as a dimer, inserting two adapters separated by 9 bp ^50,181^. Conventionally, to identify the center of the dimer complex for each transposition event, reads aligning to the + strand are shifted by +4 bp, while those aligning to the - strand are shifted by-5 bp. However, we observed that this +4/-5 shift for ATAC-seq leads to a single-base misalignment between the positive and negative strands. To demonstrate this, we constructed 24-mer position weight matrices (PWMs) from sequences flanking the shifted reads, separately for positive and negative strand profiles, as well as for collapsed unstranded profiles. While PWM motifs from both positive and negative shifted reads resembled the known Tn5 motif, they were offset by 1 bp, indicating a disagreement in the Tn5 transposase dimer centers between strands (Extended Figure 3a). Consequently, the PWM from collapsed unstranded reads, which averages the two stranded motifs, did not resemble the Tn5 sequence bias motif. This misalignment is concerning because a model trained to predict the unstranded +4/-5 shifted profile from the sequence would attribute inaccurate sequence preferences to the enzyme. We demonstrate this in Extended Figure 3a, where a simple CNN model trained to predict the +4/-5 shifted profile learns the distorted motif instead of the conventional Tn5 motif. We resolved this disagreement by shifting the negative strand reads by-4 bp instead of-5 bp. The PWM constructed on +4/-4 shifted unstranded ATAC-seq reads then matches the Tn5 motif. Following a similar analysis for DNase-seq assays, we shift reads on the - strand by +1 bp and leave the + strand unshifted, in contrast to the conventional unshifted (i.e., 0/0) reads in DNase-seq assays (Extended Figure 3b).

### 1.3 ChromBPNet (without bias correction) architecture

The ChromBPNet architecture, inspired by BPNet models 97. for TF ChIP-seq datasets, employs a sequence-to-profile design. This fully convolutional neural network takes a 2114 bp one-hot-encoded DNA sequence as input (A = [1,0,0,0], C = [0,1,0,0], G = [0,0,1,0], T = [0,0,0,1]) and maps it to chromatin accessibility read count profiles at base-resolution for 1000 bp of the output. The architecture begins with a convolution layer using 512 filters of width 21 bp, followed by dilated convolutional layers, each with 512 filters of width 3. The dilation rate, which represents the number of skipped positions in the convolutional filter, doubles at each layer. Through systematic architecture search using HEPG2 ATAC-seq models, we determined that 8 dilated layers, corresponding to an effective receptive field of 1041 bp, provided optimal performance (**Supp Fig. 4**).

ChromBPNet’s final convolutional layer output, known as the bottleneck activation map, feeds into two distinct output heads: (1) A deconvolutional layer with a filter width of 75 bp predicts the probabilities of observing a particular read at a specific position in the input sequence, contributing to shape or profile prediction. (2) A global average pooling layer, followed by a fully connected layer, predicts the total number of read counts aligned to the input sequence, encapsulating total read count prediction This dual-head approach allows ChromBPNet to capture both the spatial distribution and overall magnitude of chromatin accessibility signals, providing a comprehensive model of DNA sequence-chromatin accessibility relationships.

### 1.4 ChromBPNet (without bias correction) training and hyper-parameter tuning

Utilizing an Adam optimizer with a learning rate set at 0.001, our model undergoes training on peak regions with random shifts (jittering) of 500bp around the summit during each training epoch. To ensure the quality of training data, we exclude peak-regions falling into three categories: (1) those beyond the 0.99 threshold of total counts, (2) regions located on the chromosomal edges, making it impossible to construct the full 2114 bp length regions, and (3) regions that intersect with the blacklist region obtained from https://www.encodeproject.org/files/ENCFF356LFX.

### 1.5 BPNet based bias model architecture

The bias model, responsible for capturing the enzymatic bias contribution, adopts a sequence-to-profile architecture akin to BPNet and ChromBPNet (without bias correction). However, it is characterized by a significantly smaller capacity and a shallow architecture. This fully convolutional neural network processes a 2114 bp one-hot-encoded DNA sequence (A = [1,0,0,0], C = [0,1,0,0], G = [0,0,1,0], T = [0,0,0,1]) as input, mapping it to chromatin accessibility read count profiles at base-resolution for a 1000 bp output. In its design, the architecture features an initial convolution layer employing 128 filters with a width of 21 bp. Subsequently, four dilated convolutional layers follow, each equipped with 128 filters of width 3. Notably, the dilation rate, indicating the number of skipped positions in the convolutional filter, doubles at each layer. This cumulative approach results in a receptive field of 81 bp for any given position in the sequence.

The output of the final convolutional layer within BPNet, often referred to as the bottleneck activation map, serves as the input for two distinct output heads. Firstly, a deconvolutional layer with a filter width of 75 bp predicts the probabilities of observing a specific read at a particular position in the input sequence, contributing to shape or profile prediction. Secondly, a global average pooling layer, followed by a fully connected layer, predicts the total number of read counts aligned to the input sequence, encapsulating total read count prediction.

### 1.6 BPNet based bias model training and hyper-parameter tuning

#### Model Training

For training the bias model, we employ an Adam optimizer with a learning rate of 0.001. The training process is conducted using the filtered set of non-peak region obtained as follows-

#### Data Preparation

First, acquire non-peak regions, twice the number of peak regions, GC-matched with the peak set. Second, filter out non-peak regions whose total counts in the 1000 bp profile exceed a threshold. The threshold is determined by the 0.01th quantile of total counts in peak regions, multiplied by a fraction less than 1 (quantile_of_total_counts_in_peaks * fraction). This filtering aims to eliminate potential false-positives in non-peak regions that may have resulted from peak-calling algorithms. By removing high-signal regions from the non-peak set, particularly those associated with open chromatin and TF-binding, we ensure the bias model captures only the bias contribution and not the TF contribution to the accessibility profile.

#### Hyperparameter Tuning

The background signal coverage threshold is a critical hyperparameter tuned empirically for each dataset to ensure the bias model captures only enzymatic preferences, not biological TF binding signals. Our tuning process is iterative and interpretation-guided: We begin with a conservative default threshold (maximum background signal set to 80% of the signal at the 1st percentile of peak regions) and train a preliminary bias model. We then apply DeepLIFT and TF-MoDISco to analyze learned sequence motifs. If any known TF motifs are identified, it indicates that weakly accessible peaks with TF occupancy have infiltrated the background regions used to the train the bias mode. Hence, to obtain a more conservative background set, we reduce the maximum background signal threshold (e.g. to 70%) and retrain. We repeat this until the bias model learns only enzymatic preference motifs (e.g., Tn5 or DNase-I patterns) with no TF motif contamination.

This process intentionally excludes background regions that may contain weak TF binding sites to ensure bias model purity. A potential limitation arises when datasets are particularly noisy: overly stringent thresholds can remove many background regions, skewing the GC-content distribution relative to the peak set and potentially impairing bias model performance. For such cases, we recommend using a robust pre-trained bias model from a higher-quality dataset of the same assay type, as enzymatic biases are largely consistent across experiments. Table 1 lists the final threshold parameters for each model, and Supplementary File 3 contains TF-MoDISco reports validating bias model purity.

### 1.7 Bias-factorized ChromBPNet architecture

The Bias Factorized ChromBPNet leverages a dual-architecture approach for assay bias correction, integrating two distinct models:

#### TF Model with ChromBPNet (without bias) Architecture

The first is a TF Model, which mirrors the structure of ChromBPNet without bias correction from Section 1.4. This model is initialized with random weights and equipped with trainable parameters. It takes a 2114 bp sequence as input, has a receptive field of 1041 bp and predicts the profile and total coverage of a 1000 bp profile. Let *p^tf^* be a vector of length 1000 denoting the predicted probabilities along the sequence, ensuring ∑ *p^tf^_i_*= 1, and *n ^tf^* be the total number of predicted counts for the same sequence.

#### Bias model with BPNet architecture

The second component is a Bias Model with BPNet architecture, which adopts a shallow structure with a receptive field of 81 bp from Section 1.5. This model is pre-trained on background regions, and its weights are frozen during the training of the bias-factorized ChromBPNet model. Similar to the TF Model, it outputs probabilities of vector length 1000 denoted by *p^bias^*, ensuring ∑ *p_i_^bias^* = 1, with *n ^bias^* representing the total number of predicted counts for the same sequence.

The final probabilities *p^pred^* and counts *n^pred^* of the Bias Factorized ChromBPNet are then determined as follows:

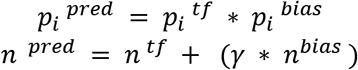

Where *γ* is the scaling factor that accounts for the difference in total bias coverage in peaks versus non-peaks. We calculate this scaling factor as follows 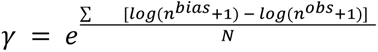 where N is the number of examples in the training and validation set.

### 1.8 ChromBPNet loss

The ChromBPNet loss function closely mirrors the loss function employed by BPNet, minimizing errors in both predicting the profile shape and estimating the total signal across all training sequences. Here we delve into the definitions of these loss terms:

#### Multinomial Negative Log-Likelihood (MNLL) Loss for Profile Shape

- Consider *k^obs^* a vector of length L representing the observed read counts along a sequence of the same length L.
- Define *n ^obs^* = ∑*_i_ k_i_* as the total number of observed read counts for the length L
- Let *p^pred^* be a vector of length L denoting the predicted probabilities along the sequence, ensuring that ∑ *p_i_* = 1
- The error in profile shape is quantified by the Multinomial Negative Log-Likelihood (MNLL) loss – 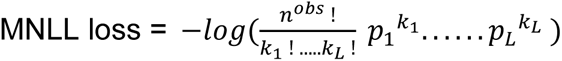

#### Error in Total Signal

- Define *n ^obs^* = ∑*_i_ k_i_* as the total number of observed read counts for the length L
- And *n ^pred^* be the total number of predicted counts for the sequence of length L
- The error in total signal is quantified by the means squared error (MSE) loss - MSE loss = (*log*(1 + *n^obs^*) − *log*(1 + *n^pred^*))^2^

The loss function of ChromBPNet integrates both the MNLL loss term and the MSE loss term using the hyperparameter λ, which is set to 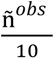 where ñ*^obs^* is the median of the total counts across all sequences in our validation set. This overall loss, employed for training ChromBPNet is summarized here – 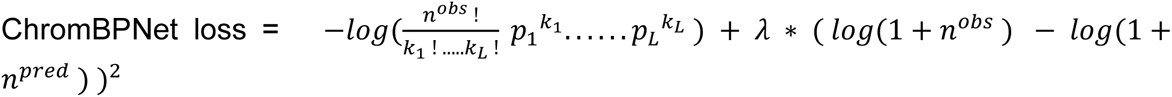

### 1.9 Bias-factorized ChromBPNet training and hyperparameter tuning

Our training process incorporates both peaks and GC-matched negative samples for each epoch, maintaining a 1:10 ratio of non-peaks to peaks. To enhance model generalization, we apply random jitter to the peaks during training, shifting them by up to 500 bp around the summit, while non-peaks remain stationary. We utilize an Adam optimizer with a learning rate of 0.001 for optimization. The training employs a five-fold cross-validation strategy, with train/test and validation splits as detailed in Table 2. The validation set plays a crucial role in determining when to halt training; we cease the process when the validation loss shows no improvement for five consecutive epochs. At this point, we restore the model weights from the last known best checkpoint.

### 1.10 Performance metrics

The evaluation of ChromBPNet models involves assessing the performance of its two output heads independently: profile shape and total signal.

For profile shape evaluation, we employ the Jensen Shannon Distance (JSD) between observed and predicted base-resolution probability profiles for each region. Let *p^pred^* be a vector of length L representing the predicted probabilities along the sequence, where ∑ *p_i_* = 1. Similarly, let *k^obs^* be a vector of length L representing the observed read counts. The JSD between two probability vectors *p^pred^* and 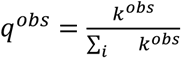 is defined as:

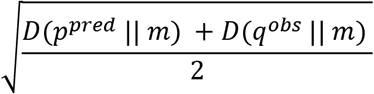

where m is the pointwise mean of *p^pred^* and *q^obs^*, and D is the Kullback-Leibler divergence computed with logarithm base 2. JSD ranges from 0 to 1, with lower values indicating better performance.

For total signal evaluation, we calculate the Pearson correlation coefficient (Pearson’s r) between the predicted and observed log counts for all regions. Higher values of Pearson’s r indicate better performance.

#### Genomewide AUPRC / AUROC

We performed chromosome-wide predictions using ChromBPNet models on sequences centered around 1000 bp bins with a 250 bp stride. We then created genome-wide bins (B) of 100 bp with a 100 bp stride.

To evaluate AUPRC and AUROC metrics, we define positive and negative regions using the following procedure:

To define positive regions, we considered the intersection of IDR peaks from ATAC-seq and DNase-seq data, centered at the summit and expanded by 100 bp on both sides. A bin B was labeled positive if it had 100% overlap with this IDR peak set.

We identified and removed ambiguous bins based on the following criteria:

- Bins overlapping an IDR peak by < 100%
- Bins overlapping an overlap-peak by any percentage without being a positive bin
- Bins overlapping blacklist regions by any percentage
- Bins with < 50% positions uniquely mappable for 100 bp reads (https://bismap.hoffmanlab.org/raw/uint/hg38all/umap/)

Negative regions for evaluation were defined as the genome-wide bins excluding both positive and ambiguous bins.

### 1.11 DeepLIFT contribution scores for sequence-to-profile models

We employ the DeepLIFT ^121, 97^. algorithm to extract sequence features predictive of accessibility. DeepLIFT assesses the contribution of each feature in the input sequence (one-hot encoded) to predict the difference between the output prediction for that sequence and a reference. This relationship can be expressed as:

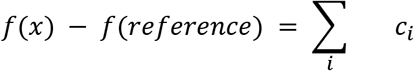

where *f* is the model output, and *c_i_* represents the contribution scores derived for a feature *i* in an input sequence *x* using DeepLIFT. For generating reference sequences, we utilize a di-nucleotide shuffling approach on our input sequence x. This technique preserves the di-nucleotide frequency of the original sequence while randomizing its overall structure. We create 20 such shuffled sequences for each input, serving as our reference set. We then compute DeepLIFT contribution scores for the original sequence relative to each of these 20 reference sequences. The final contribution scores are obtained by averaging across these multiple references. This procedure is applied independently when generating contribution scores for the counts and profile heads. The query sequence and the 20 references each require their own forward passes. Each (query-reference) pair requires its own DeepLIFT multiplier backpropagation pass to estimate contribution scores.

The application of this approach differs for the two output heads of our model. For the counts head, which involves a single output scalar value, this approach is straightforward. However, for the profile head, where the same input feature contributes to predicting a vector of probability values, we adopt a strategy similar to BPNet. Here, contribution scores derived for each value in the output vector are weighted based on the probabilities predicted in the output profile. Let *P^pred^* be the output of the profile head with a vector of length L denoting the predicted probabilities along the sequence, ensuring ∑ *p_j_* = 1.. If *C_i_*_,*j*_ represents the contribution scores for each feature *i* with respect to the profile prediction at *j*, then the profile contribution score of a feature with respect to the entire output profile is given by:

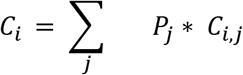

The DeepLIFT contribution scores were computed using TensorFlow v.1.6 and the DeepExplain implementation of DeepLIFT is available at https://github.com/kundajelab/shap/tree/master.

### 1.12 Motif discovery using TF–Modisco

To systematically identify recurring sequence patterns with high contribution scores and generate Contribution Weight Matrices (CWMs), we employ the de novo motif discovery tool TF-Modisco (https://github.com/kundajelab/tfmodisco (version 0.5.16.0)) ^106, 97^. We apply this tool to contribution scores derived independently from both the counts and profile output heads of the base pair models, centered 500 bp around the summit regions in our peaks. The application of TF-Modisco varies based on specific analysis goals. For model Quality Control (QC) (Fig 1-2), we conduct smaller scale runs utilizing contribution scores from 30K peaks and 50K seqlets to determine the presence or absence of bias motifs. For large-scale analyses aimed at studying the detailed syntax governing the cis-regulatory landscape in specific cell types (Fig 3-5), we utilize a more efficient implementation called modisco-lite (https://github.com/jmschrei/tfmodisco-lite (Version 2.2.0). In these cases, we run modisco-lite on all peaks with 1,000,000 seqlets, employing the command “*modisco motifs-n 1,000,000-w 500*”.

### 1.13 Annotating Contribution Weight Matrices (CWMs) from TF-Modisco with Transcription Factors

We annotate our Contribution Weight Matrices (CWMs) with corresponding Transcription Factor (TF) labels using a two-pronged approach: leveraging existing TF-motif databases via TOMTOM and integrating statistical TF-ChIP enrichment analysis when applicable.

For TOMTOM (Version 5.5.5)^182^ annotation, we first obtain the PWM matrices for the seqlets representing the CWMs in TF-Modisco. These matrices are trimmed by removing edges where the CWM signal is less than 30% of the max-score and stored in MEME format. We then use TOMTOM to annotate each TF-Modisco pattern with the top matching TF motif from the JASPAR database, using the command: *tomtom-no-ssc-oc. --verbosity 1-text-min-overlap 5-mi 1-dist pearson-evalue-thresh 10.0*.

For TF-ChIP enrichment analysis, we consider the genomic landscape defined by all bases covered by peaks in an ATAC-seq or DNase-seq assay. We assess association significance using

Fisher’s exact test, comparing bases covered by seqlets constituting a TFModiso CWM with those covered by TF ChIP-seq peaks. To determine the bases covered by TF ChIP-seq peaks, we aggregate all peaks for each TF from various ENCODE portal experiments, considering a 20bp region around each peak summit and merging these regions using bedtools.

Following the association analysis, we compute corrected p-values for each motif using the Benjamini-Hochberg correction, filtering them at a false discovery rate (FDR) of 0.1. We rank TFs associated with a motif using the log-odds ratio. Manual annotation occurs by assigning a TF label when one of the five highest-ranked TFs aligns with existing databases such as TOMTOM. In cases where TF ChIP-seq datasets are unavailable, we rely solely on TOMTOM for annotation.

### 1.14 Unifying count and profile TF-MODISCO motifs from DNase-seq and ATAC-seq ChromBPNet models

To identify a unified, non-redundant set of motifs for each cell-line, we followed a three-step process. First, we compiled TF-MODISCO motifs (PFMs) from both profile and count heads across ATAC-seq and DNase-seq assays. Next, we filtered out any motifs with less than 100 seqlets support to ensure robustness. Finally, we use the motif clustering tool from the Gimmemotifs^183^ toolkit with a threshold of 0.99 (*gimme cluster-t 0.99*) to cluster the motifs into a unified, non-redundant set for each cell-line.

### 1.15 Marginal footprinting

We introduce a method to obtain marginal footprints, which allows us to generate footprints for a TF/bias motif by marginalizing over the contribution of other TF motifs that may influence the footprint due to co-occurrence. The procedure involves the following steps:

- Sequence library Generation: We craft a library of sequences by inserting TF/bias motif instances of interest within the center of background sequences that are GC-content matched with the peaks. Unless otherwise stated, we usually insert the **highest affinity** motif instance (kmer with highest contribution score according to CWM) in background sequences.
- Model Predictions: We generate model predictions for each sequence in the library and its reverse complement. These predictions comprise a logits vector of length L (denoted as *f*) and scalar values for log-counts (*S*).
- Profile Calculation: To derive the **probability profile predictions**, we apply *softmax* (*f*) to the predicted logits for each sequence. We then average the per-base predicted profile values between the sequence and its reverse complement (after flipping the prediction made from the reverse complement sequence).
- Marginal Footprint: We average the above probability profiles across the entire library of sequences to obtain the “marginal footprint” for the motif of interest.

#### Interpretation (shape vs magnitude)

In this work, “marginal footprints” are computed from the model’s predicted probability profiles (per-sequence profiles normalized to sum to 1) for the highest affinity motif instances. Consequently, marginal footprints primarily characterize motif-associated changes in the *shape* of the cut distribution (i.e., local redistribution of cuts around the motif) and should not be interpreted as the motif’s marginal effect on total accessibility magnitude. A motif can therefore exhibit a strong footprint-like pattern in the averaged probability profile even when its marginal effect on total accessibility in an inert background is small. When we assess accessibility magnitude, we do so separately using the model’s count head (or equivalently the total predicted counts), rather than the normalized probability profile. Changes in affinity of the inserted motif can affect marginal footprints and total marginal accessibility.

To determine the height and width of these marginal footprints and to use them for comparison across DNase/ATAC-seq assays, we generate controls by first creating similar “marginal footprints” on synthetic sequences without any motif insertions. We compute log-fold changes in total counts coverage between the final “marginal footprint” of the TF of interest and controls. This is then multiplied with the total probability predictions obtained for the marginal footprints of the TF of interest. The resulting footprint is smoothed using a convolution with an identity filter of width 5, denoted as vector *M*. We then analyze the gradient of this profile M to determine the footprint characteristics: The positions of maximum gradient change on either side of the center of the TF/motif insertion (*rs*, *ls*) provide coordinates for the valley of the footprint.

Height is calculated as: *Max*(*M*[: *rs*], *M*[*ls*:]) − *min*(*M*[*rs*, *ls*])

Width is determined as: *ls* − *rs*

### 1.16 ChromBPNet model baselines with HINT-ATAC correction

We applied HINT-correction ^79^ to the downloaded BAM files and overlapped peak-set of GM12878 ENCODE datasets described in Section 1.1 to obtain bias-corrected profiles. This was achieved using the command “rgt-hint tracks --bc --bigWig --organism=hg38”. Using these HINT bias-corrected tracks, we trained BPNet models following the architecture outlined in Section 1.3. These models incorporate 512 filters and 8 dilation layers, and are designed to process a 2114 bp input sequence to predict both total counts and profile shape for a 1000 bp region within the peaks. For model training, we followed the procedure detailed in Section 1.4, specifically for fold 0. The resulting trained models are available in the synapse repository syn59480573, with the corresponding TF-MODISCO reports provided in Supp Files 1 and 2

### 1.17 ChromBPNet model with TOBIAS correction

We employed the TOBIAS ^78^ method for benchmarking using two distinct approaches:

#### Direct TOBIAS Correction

We applied TOBIAS to the filtered BAM and overlapped peak-set of the GM12878 ENCODE ATAC-seq dataset from Section 1.1 using the command TOBIAS ATACorrect. The resulting bias-corrected profile contained both positive and negative values, incompatible with ChromBPNet’s requirement for positive-only inputs to its multinomial loss function. To address this, we explored two transformations: a) Softmax b) Shifting by the lowest value to ensure all values are positive, then normalizing to create a probability vector.

We then multiplied these transformed probability profiles with the total counts in 1000bp of the observed profile to prepare them for training. Using these processed profiles, we trained BPNet models as described in Section 1.3, featuring 512 filters and 8 dilation layers. These models process 2114 bp input sequences to predict total counts and profile shape for 1000 bp regions within peaks. Training followed the procedure outlined in Section 1.4 for fold 0.

#### TOBIAS-factorized ChromBPNet

We obtained unnormalized bias tracks from TOBIAS (Equation 1 from TOBIAS supplementary paper) and input them to a 1 filter-deconvolution layer (kernel width 75, matching the last layer in the BPNet architecture). This layer’s weights were learned by training in peak regions. The resulting smoothed bias tracks were used in place of the BPNet bias model output described in Section 1.7 to train a TOBIAS-factorized ChromBPNet. We trained this model on fold 0 of the observed profiles, following the training procedure in Section 1.4.

The trained models from both approaches are available in the synapse repository syn59480573, with corresponding TF-MODISCO reports provided in Supp Files 1.

### 1.18 ChromBPNet models with naked DNA BPNet bias models

We developed a naked DNA-factorized ChromBPNet model through a multi-step process. Initially, we trained BPNet-based bias models following the architecture outlined in Section 1.5, using peak regions from bulk ATAC-seq and DNase-seq data, but incorporating signals from the Naked DNA bigwigs obtained in Section 1.1. We then froze this trained bias model and integrated it as the bias component described in Section 1.7 to train a Naked DNA-factorized ChromBPNet,

To evaluate the efficacy of this bias correction method, we obtained TF-MODISCO consensus motifs from contribution scores of the same 30K peaks used for the GM12878 ChromBPNet model analysis in Extended Figure 4, ensuring direct comparison between the analyses. The resulting trained models are available in the synapse repository syn59510977, with the corresponding TF-MODISCO reports provided in Supp Files 4.

### 1.19 ChromBPNet models on African ancestry bulk ATAC-seq

To develop bias-factorized ChromBPNet models for each of the African ancestry datasets, we utilized the peaks and bigwigs generated for each ancestry sub-groups from Section 1.2. These models were trained using the hg38 reference genome as the sequence input. Instead of training a new bias model for each ancestry, we leveraged the existing K562 ATAC-seq bias model across all ancestry-specific ChromBPNet models^67^. The resulting trained models are available in the synapse repository syn59651013,

### 1.20 ChromBPNet models on single-cell ATAC-seq

To develop bias-factorized ChromBPNet models for the Microglia and Smooth Muscle Cell (SMC) single-cell ATAC-seq datasets, we utilized the peaks and bigwigs generated for each cell type as described in Section 1.1. For the Microglia ChromBPNet model, we employed a bias model trained on cluster 1 (isocortical excitatory cell type) from the scATAC-seq pseudobulk dataset, while the ChromBPNet model itself was trained on cluster 24 (microglia)^138^. In contrast, for the Smooth Muscle Cell ChromBPNet model, we used an in vivo trained bias model on the same SMC background profile. The resulting trained models are available in the synapse repository syn59479965 (SMC) and syn59479966 (Microglia),

### 1.21 Subsampling

To investigate the impact of sequencing depth on our analysis, we performed a systematic subsampling of our complete deep GM12878 ATAC-seq dataset (ENCSR637XSC), originally sequenced at 572M reads. The subsampling process was executed using the following command:

“*sambamba view-f bam-t 40 --subsampling-seed=1234-s frac”,* where’frac’ was appropriately set to achieve BAM files with read depths of 250M, 100M, 50M, 25M, and 5M.

For each subsampled dataset, we employed the caper tool (version 2.1.2) for peak-calling and generated GC-matched negatives. When training bias-factorized ChromBPNet models on these subsampled datasets, we transferred the HEPG2 bias model across all read depths instead of training new bias models for each subsample.

To assess the consistency of motif identification across different sequencing depths in Figure 4c, we employed the FiNeMo method (https://github.com/austintwang/finemo_gpu). This approach allowed us to identify occurrences of TF-MODISCO motifs, which were initially generated from the ChromBPNet model trained on the full 572M read-depth dataset. We then searched for these motifs within the contribution scores of ChromBPNet models trained on subsampled datasets at various read depths.

### 1.22 Comparative Analysis of ChromBPNet and BPNet TF ChIP-seq Contribution Scores

To establish a baseline for comparing ChromBPNet model contribution scores from ATAC-seq/DNase-seq with TF-ChIP BPNet model contribution scores, we implemented a systematic approach. First, we annotated the CWM motifs identified by TF-Modisco in our contribution scores with corresponding TFs, using the method detailed in Section 1.13. For each TF and its associated ENCODE experiments, we extracted a BPNet model along with its corresponding peaks and contribution scores from the ENCODE portal. We then used MOODS (p-value = 0.001) to identify significant sequence hits corresponding to the CWM motif representation. This set was further refined by selecting only those hits that lie within an ATAC-seq peak and are within a 20 bp window around the ChIP-seq summit. For these filtered instances, we extracted the sum of contribution scores from both ChromBPNet and BPNet models and computed the Pearson correlation between them. For each CWM-TF pair, we report the highest Pearson correlation coefficient obtained across all experiments. Our analysis focused on GM12878, HEPG2, K562, and H1ESC cell lines, which have the most abundant ChIP-seq datasets, excluding IMR90. In total, we evaluated 72 unique CWM-TF pairs and 138 CWM-experiment pairs across these four cell types. The CWMs used for MOODS and the specific TF ChIP-seq ENCIDs considered for each motif are provided in the synapse repository syn63887196.

### 1.23 ChromBPNet variant effect prediction metrics

To assess the impact of variants on chromatin accessibility, we developed a systematic approach using our ChromBPNet model. Consider a SNP with alleles *a*_1_ and *a*_2_, alongside a reference sequence. We generate two sequences, *S*_1_ and *S*_2_, by inserting alleles *a*_1_ and *a*_2_ respectively into the reference genome sequence. We then utilize a ChromBPNet model trained on fold *i*, which has two output heads:

1. A total counts output head, represented by the function *f_i_*(*x*) and
2. A profile probability vector, represented by *P_i_*(*x*).

For both sequences *S*_1_ and *S*_2_, centered on their respective alleles, we compute: the predicted total counts: *f_i_*(*S*_1_) and *f_i_*(*S*_2_) the predicted profile probability vectors: *P_i_*(*S*_1_) and *P_i_*(*S*_2_)

Using these predictions, we calculate the following orthogonal scoring measures to quantify the impact of allele *a*_2_ relative to *a*_1_:

1. **Log-Fold Change:** *logFC_i_*(*a*_2_, *a*_1_) = *log*(*f_i_*(*S*_2_)) - *log*(*f_i_*(*S*_1_))
2. **Jensen Shannon Distance** between *P_i_*(*S*_1_) and *P_i_*(*S*_2_): 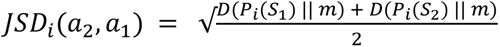

where m is the pointwise mean of *P_i_*(*S*_1_) and *P_i_*(*S*_2_), and *D* is the Kullback-Leibler divergence calculated with base log e. As JSD is unsigned, we multiply it with

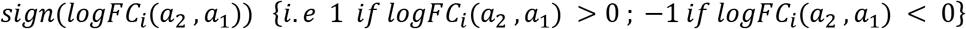

to determine the direction of effect while using it as an effect size measure.

1. **Active Allele Quantile:** *AAQ_i_*(*a*_2_, *a*_1_) = *max*(*pp_a_*_1_, *pp_a_*_2_) where *pp_a_*_1_ and *pp_a_*_2_ is the percentile of *log*(*f_i_*(*S*_1_)) and *log*(*f_i_*(*S*_2_)) with respect to log counts distribution in peaks.

We then use all these three measures computed across 5-folds as follows to define:

- **Integrative Effect Size (IES)** combining effect sizes from both total coverage and shape

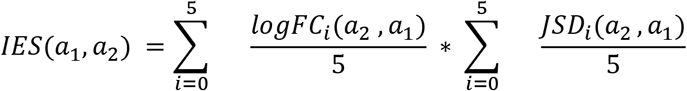

- Integrative Prioritization Score (IPS) to differentiate significant from non-significant variants:

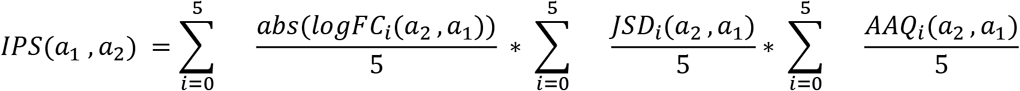

- **Significance scores (p-value):** To assess the statistical significance of our IPS scores, we begin by establishing a null distribution of IPS scores through the generation of 1,000,000 synthetic examples. For each example, we perform dinucleotide shuffling on the input reference sequence intended for SNP insertion, followed by the reintroduction of variant alleles into these shuffled sequences. We then compute IPS scores for each of these synthetic sequences, forming a comprehensive null distribution. This distribution is subsequently utilized to calculate empirical p-values for the IPS scores obtained from allele insertions in the actual reference sequences of interest.

These scores provide a comprehensive assessment of the impact of genetic variants on predicted chromatin accessibility. The Active Allele Quantile (AAQ) provides a confidence measure for predicted variant effects. Rather than using a null distribution, AAQ compares the predicted signal at a variant’s active allele to the distribution of predicted signals in peak regions. Higher AAQ values indicate greater confidence that the variant resides in accessible chromatin. We recommend using the composite IPS score for variant screening and prioritization, as it combines effect magnitude with prediction confidence. For quantifying the magnitude and direction of variant effects, we recommend using logFC and JSD scores without AAQ. For ease of use and reproducibility, the software for calculating these scores is publicly available at: https://github.com/kundajelab/variant-scorer

### 1.24 Enformer variant effect prediction metrics

Enformer’s (reported) variant scores: To obtain Enformer’s reported variant effect prediction scores, we follow a systematic process^97^. We begin by downloading all h5 files from the provided link - https://console.cloud.google.com/storage/browser/dm-enformer/variant-scores/1000-genomes/enformer;tab=objects?prefix=&forceOnObjectsSortingFiltering=false. Next, we search these files based on RSID or SNP positions. We then retrieve the corresponding SAR (SNP Activity Ratio) and SAD (SNP Activity Difference) Enformer predicted scores for the appropriate DNase heads, as outlined in the target sheet (https://raw.githubusercontent.com/calico/basenji/master/manuscripts/cross2020/targets_human.txt). Finally, we match the chromosome, position, and ref/alt allele fields to assign Enformer scores to each SNP.

Enformer’s (re-computed) variant scores: Enformer’s initial underperformance in predicting variant effects can be attributed to the methodology used to compute the reported scores. The original approach calculated log-fold changes in counts over a large 114,688 bp region, aiming to capture global accessibility changes. However, this broad focus inadvertently diluted local variant effect scores, reducing prediction accuracy. To address this limitation, we implemented a refined methodology. We now insert variant alleles into reference sequences and obtain predictions from the relevant DNase head. Focusing on the center 8 bins, covering a context of 1024 bp around the alleles, we sum these counts to emphasize local accessibility changes. The log-fold change between the alternate and reference alleles is then used to calculate the revised variant effect scores.

### 1.25 Variant effect prediction datasets

The following is a comprehensive list of variant effect datasets used to benchmark ChromBPNet. We have meticulously standardized these datasets into an easily accessible resource to streamline future benchmarking efforts for machine learning models focused on variant effect prediction. The standardized datasets can be accessed at syn64126763. Below, we provide detailed information on the preprocessing steps applied to each dataset-

**DNase QTLs in Yoruba LCLs:** To prepare the deltaSVM dataset for benchmarking, we downloaded Supplementary Table 1 from the deltaSVM study^137^, available at https://staticcontent.springer.com/esm/art%3A10.1038%2Fng.3331/MediaObjects/41588_2015_BFng3331_MOESM26_ESM.xlsx The original dataset comprised 574 dsQTL SNPs and 27,735 control SNPs, along with deltaSVM predicted effect sizes. We then filtered the dataset to include only SNPs with pre-computed Enformer predictions, resulting in 560 dsQTL SNPs and 26,813 control SNPs. To obtain the actual effect sizes, we extracted data from https://www.ncbi.nlm.nih.gov/geo/query/acc.cgi?acc=GSE31388 (file: GSE31388_dsQtlTable.txt.gz).

**Chromatin QTLs in European LCLs:** ATAC-seq data from^66^ was processed with the nf-core/atacseq v2.1.2 pipeline using Nextflow v23.09.3. We aligned raw ATAC-seq reads to the GRCh38 reference genome (Homo_sapiens.GRCh38.dna.primary_assembly.fa downloaded from Ensembl) with BWA v0.7.17. We called broad peaks with MACS2 v2.2.7.1 and defined consensus peaks as the union of all peaks that were present in at least 5% of the samples. We then quantified read overlaps with the set of consensus peaks with featureCounts v2.0.1. Finally, we normalized the read counts (counts per million) and then used the inverse normal transformation to standardize the data distribution. Genotype data for the 91 overlapping samples were downloaded from 1000 Genomes 30x on GRCh38 website. Finally, we used the eQTL-Catalogue/qtlmap v24.01.1 workflow to perform chromatin accessibility QTL analysis. We set cis window size to 200,000 bp and excluded peaks that had less than 25 variants within that window. The resulting summary statistics from this analysis are publicly available at https://zenodo.org/records/13848268 (file: QTD100018.all.tsv.gz). For those interested in the detailed methodology of our association testing workflow, we refer to the comprehensive description provided in^185^.

To refine our analysis, we expanded the broad peaks called across individuals to ±100 bp of the summit of the top 50K peaks for each individual, using this to filter the SNPs further. This yielded 130,268 variants with associations. We then focused on variants with pre-computed Enformer variant effect scores, resulting in 7,900 significant QTLs (-log10 p > 6) and 87,165 control QTLs (-log10 p < 3), establishing a baseline ratio of 0.08. Additional analysis varying the threshold for significant QTLs (-log10 p from 6 to 24) while keeping the control set constant is presented in our Extended Figure 9.

**Chromatin QTLs in African LCLs**: ^67^ identified 11,098 significant SNPs out of 219,382 tested for caQTLs across 100 representative individuals from 6 African populations. We obtained these 219,382 SNPs and their measured effect sizes (Beta) from the authors for our analysis. To complement this data and standardize the analysis with the European caQTLs processed earlier, we applied the same workflow to the ATAC-seq data from the 100 African individuals. We utilized the nf-core/atacseq v2.1.2 pipeline with Nextflow v23.09.3. Raw ATAC-seq reads were aligned to the GRCh38 reference genome (Homo_sapiens.GRCh38.dna.primary_assembly.fa from Ensembl) using BWA v0.7.17. Subsequently, we called broad peaks using MACS2 v2.2.7.1

We then filtered the QTLs dataset to include only SNPs within ±100bp of the summit of the top 50K peaks across all 100 individuals, ensuring they had pre-computed Enformer scores and fell within the provided regions attribute. This resulted in 79,026 variants, of which 6,821 were significant (-log10 p > 5). We defined our control set as SNPs with-log10 p < 3, yielding 6,821 significant and 72,205 control SNPs, establishing a baseline ratio of 0.09. Additional analyses varying the significance threshold (-log10 p from 5 to 23) while keeping the control set constant are presented in our Extended Figure 9.

The original study also identified 7,559 high-confidence Allele-Specific Chromatin Accessibility (ASC) sites from ATAC-Seq. We further filtered these to sites within ±100bp of the summit of the top 50K peaks across all African populations, resulting in 5,220 SNPs for comparison with ChromBPNet predictions.

**Chromatin QTLS in smooth muscle cells (SMCs):** We extracted 1,984 significant caQTLs, along with their RASQUAL effect sizes, from Smooth Muscle Cells (SMCs) as reported by ^70^. This data was sourced from Supplementary Data 6, available at https://static-content.springer.com/esm/art%3A10.1038%2Fs41588-022-01069-0/MediaObjects/41588_2022_1069_MOESM10_ESM.xlsx. To refine our analysis and reduce false positives, we further filtered these caQTLs. We included only those SNPs falling within ±100 bp of the peak summits identified in our SMC ATAC-seq data. This stringent filtering process resulted in a final set of 386 caQTL SNPs, providing a focused dataset for evaluating chromatin accessibility changes in SMCs

**Chromatin QTLS in Microglia:** We processed the significant caQTLs in Microglia from ^69^ through a multi-step approach. First, we downloaded SNPs from https://www.synapse.org/#!Synapse:syn30863713 and obtained effect sizes by merging “PeakID” and “Top_SNP_perPeak” columns with the microglia_macrophage_meta-caQTL_summary_result (https://www.synapse.org/#!Synapse:syn30308248) based on “Peak” and “Variant” columns respectively. This resulted in 4,978 caQTL effects, including effects of single variants on multiple peaks. We then retrieved caQTL positions using rsid attributes as queries on dbSNP or by splitting attributes to obtain chromosome and position. Next, we filtered these caQTLs to include only SNPs falling within microglia peaks provided at https://www.synapse.org/#!Synapse:syn269491355, yielding 956 caQTL effect sizes. To address instances where a single SNP was attributed to multiple effect sizes, we retained only the maximum effect size for each SNP. This final refinement resulted in a set of 877 unique caQTLs with corresponding effect sizes, which were then used for benchmarking ChromBPNet predictions.

**SPI1 binding QTLs in Yoruba LCLs:** The SPI1 binding QTLs dataset^139^ comprises 999,799 SPI1 bQTLs. We obtained significance p-values and allele frequencies directly from the authors and computed effect sizes as the log fold change in allele chip frequencies. Among these, we identified 4,834 significant bQTLs using a threshold of-log10 p > 4. We further filtered this set to include only those with available Enformer predictions, which maintained the count at 3,818 bQTLs. To establish a suitable control set, we avoided using the same bQTLs, as they could potentially include bQTLs for other transcription factors. Instead, we utilized SNPs with-log10 p > 1 from the African chromatin QTLs processed earlier, resulting in 57,612 control SNPs. This approach yielded a random baseline of 0.06 when compared with the 3,818 significant bQTLs.

**MPRA variants:** We obtained test sets from the CAGI5 (Critical Assessment of Genome Interpretation 5) competition ^140,141^ through personal communication with M. Kircher. Additionally, we acquired Enformer accuracies and their predicted effect sizes from Ziga Avsec ^97^, also through personal communication. To ensure a fair comparison with Enformer’s cell-type-matched scores, we trained ChromBPNet models on the same cell types used by Enformer for each test set. We extracted ENCODE DNase experiments for these specific cell types, which included:’HepG2’ for F9, LDLR, and SORT1;’K562’ for GP1BB, HBB, HBG1, and PKLR;’HEK293’ for HNF4A, MSMB, TERT (performed in HEK293T cells), and MYCrs6983267;’pancreas’ for ZFAND3;’glioblastoma’ for TERT (performed in GBM cells);’keratinocyte’ for IRF6; and’SK-MEL’ for IRF4. Table 4 provides a detailed summary of the specific ENCODE experiment IDs (ENCIDs) used for each test set. We then directly (in a training-free setting as reported in Enformer) compared the predicted effect sizes from both ChromBPNet and Enformer with the observed effect sizes reported in the CAGI5 test set.

**Fine-mapped GWAS variants for blood traits:** We downloaded fine-mapped UK Biobank data from https://www.finucanelab.org/data, focusing on nine blood traits: Hb, HbA1c, Plt, RBC, WBC, MCH, MCV, and MCHC ^155^. To identify regulatory regions in K562 cells, we employed the Activity-By-Contact (ABC) model using reference files ENCFF860XAE and ENCFF790GF from ENCODE^160,186^. We then tested these regulatory regions for blood-trait enrichment using summary statistics from fine-mapped variants for each UK Biobank trait. This analysis involved running LDSC (https://github.com/bulik/ldsc) with 1000 Genome EUR Phase 3 genotype data to estimate LD scores, utilizing baseline v.2.2 annotations as recommended by LDSC developers, and employing HapMap 3 SNPs (excluding the Major Histocompatibility Complex region) as regression SNPs. Our analysis identified three traits (MCH, MCV, and MCHC) that explained more than 50% of their heritability through variants in K562 cells, resulting in 12,951 GWAS variants associated with blood traits. For each variant, we assigned a maxPIP score, representing the maximum of the Posterior Inclusion Probability (PIP) scores across the identified traits.

For our control set, we collated common background variants from https://alkesgroup.broadinstitute.org/LDSCORE/baseline_v1.1_hg38_annots/. We annotated the genome into different regions using GENCODEv29, retaining only variants that were neither coding, nor in transcription start sites (TSS), nor in splice sites. To identify alleles based on rsids, we used the database available at https://ftp.ncbi.nih.gov/snp/organisms/human_9606_b151_GRCh38p7/VCF/00-common_all.vcf.gz, filtering out variants without allele information. We separated entries with multiple alternate alleles, resulting in 11,903,173 control variants.

Using the full set of 11,916,124 variants, we computed the enrichment of high-scoring ChromBPNet variants overlapping with fine-mapped variants at various posterior probability (PIP) thresholds. We defined *fine*. *mapped_p_* as all variants with *PIP* > *p* for any blood trait and ℎ*ig*ℎ. *scoring_t_* as all variants with ChromBPNet *pval* > *t*. The enrichment was calculated as:

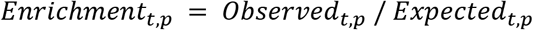

Where, at a given thresholds (t,p), the observed intersection between GWAS fine-mapped variants and significant ChromBPNet variants is represented by

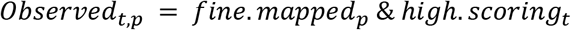

Additionally, the expected intersection between GWAS fine-mapped variants and significant ChromBPNet variants can be expressed as

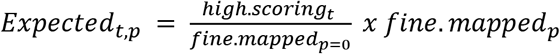

### 1.26 Transgenic E11.5 embryo generation

Transgenic E11.5 mouse embryos were generated as described previously^187^. Briefly, super-ovulating female FVB mice were mated with FVB males and fertilized embryos were collected from the oviducts. Regulatory elements sequences were synthesized by Twist Biosciences. Inserts generated in this way were cloned into the donor plasmid containing minimal Shh promoter, lacZ reporter gene and H11 locus homology arms (Addgene, 139098) using NEBuilder HiFi DNA Assembly Mix (NEB, E2621). The sequence identity of donor plasmids was verified using long-read sequencing (Primordium). Plasmids are available upon request. A mixture of Cas9 protein (Alt-R SpCas9 Nuclease V3, IDT, Cat#1081058, final concentration 20 ng/μL), hybridized sgRNA against H11 locus (Alt-R CRISPR-Cas9 tracrRNA, IDT, cat#1072532 and Alt-R CRISPR-Cas9 locus targeting crRNA, gctgatggaacaggtaacaa, total final concentration 50 ng/μL) and donor plasmid (12.5 ng/μL) was injected into the pronucleus of donor FVB embryos. The efficiency of targeting and the gRNA selection process is described in detail in Osterwalder 202215. Embryos were cultured in M16 with amino acids at 37oC, 5% CO2 for 2 hours and implanted into pseudopregnant CD-1 mice. Embryos were collected at E11.5 for lacZ staining as described previously^187^. Briefly, embryos were dissected from the uterine horns, washed in cold PBS, fixed in 4% PFA for 30 min and washed three times in embryo wash buffer (2 mM MgCl2, 0.02% NP-40 and 0.01% deoxycholate in PBS at pH 7.3). They were subsequently stained overnight at room temperature in X-gal stain (4 mM potassium ferricyanide, 4 mM potassium ferrocyanide, 1 mg/mL X-gal and 20 mM Tris pH 7.5 in embryo wash buffer). PCR using genomic DNA extracted from embryonic sacs digested with DirectPCR Lysis Reagent (Viagen, 301-C) containing Proteinase K (final concentration 6 U/mL) was used to confirm integration at the H11 locus and test for presence of tandem insertions ^187^ Only embryos with donor plasmid insertion at H11 were used. The stained transgenic embryos were washed three times in PBS and imaged from both sides using a Leica MZ16 microscope and Leica DFC420 digital camera. Data related to transgenic assays can be found at VISTA Enhancer Browser (enhancer.lbl.gov).

Bias factorized fetal brain ChromBPNet models^94^ trained on glutameric neurons (cluster 4 in scATAC-seq) were downloaded from https://www.synapse.org/Synapse:syn63395628/files/.

### 1.27 Independent Validation by the Community in Diverse Contexts

Beyond the benchmarks in our manuscript, ChromBPNet’s core claims about its ability to reveal diverse facets of the cis-regulatory code have now been stress-tested in several studies including some by independent groups in exactly the kinds of settings the reviewer highlights. These studies apply ChromBPNet (or fine-tuned variants) to new experiments, tissues, and organisms, and consistently recover novel insights into interpretable regulatory syntax and biologically meaningful variant prioritizations.:

**Primary tissues and scATAC-seq–derived pseudobulks:** ChromBPNet has been applied to cell-type/state–resolved pseudobulk accessibility profiles across a wide range of adult and fetal primary tissues, including human heart^184^ and vasculature^188^, adult and fetal human brain^184^, a massive human fetal tissue atlas^189^, human retina^190^, and human cranial neural crest cells^152^. Across these settings, ChromBPNet-derived motif syntax and sequence features are reported to be cell-type specific and useful for prioritizing candidate regulatory variants.

**Dynamic cell-state transitions**: The framework has also been used to analyze time-varying regulatory programs, including work dissecting temporal hierarchies of factors during Drosophila embryogenesis^112^ and and mapping cis-regulatory rewiring during fibroblast reprogramming that supports an intriguing repression by theft model of fibroblast de-programming mediated by non-canonical low affinity sites detected by the models^151^.

**General sequence “syntax” principles and experimental follow-up:** ChromBPNet-style motif grammar inferences have been connected to large-scale perturbation/mutagenesis efforts^153,154^ extended to cross-species settings spanning large evolutionary distances^173^, and used to highlight cooperative contributions of weaker/low-affinity motif instances in accessibility models^191^. Fine-tuned ChromBPNet models have additionally been used to study TF dosage-sensitive responses and associated cis-regulatory logic^152^.

### 1.28 Comparative Analysis of ChromBPNet scATAC-seq and bulk ATAC-seq models

We supplemented our original bulk ATAC-seq downsampling series (572M down to 5M reads) with additional datasets at 15M, 10M, and 2M reads (**Supp Fig. 2**), and trained parallel ChromBPNet models on pseudo-bulk GM12878 scATAC-seq data (ENCSR371ZNU). For this analysis, we selected high-quality cells using stringent filtering thresholds (TSS enrichment > 6 and >5,000 UMIs/cell), resulting in a dataset of 217 million fragments (434 million reads) from 19,000 cells. From this high-quality pool, we created a downsampling series by subsampling cells to generate pseudo-bulk profiles with read depths equivalent to our bulk ATAC-seq experiments (250M, 100M, 50M, 25M, 15M, 10M, 5M, and 2M reads), corresponding to approximately 11k, 4k, 2k, 1k, 663, 463, 212, and 103 cells for each respective data point.

### Imputation and Motif Recall Performance

Using these newly trained models, we extended our analysis in **Supp Fig. 2a** comparing the distribution of discordance (JSD) between profiles from the maximal read depth sample versus profiles from progressively subsampled datasets, with and without bias correction, over peak regions. While observed ATAC-seq profiles exhibit increasing discordance with reducing read depth, uncorrected and bias-corrected predicted profiles, along with bias-corrected contribution scores, exhibit very similar distributions down to 25M reads, with deterioration only visible at 10M reads. We also compared recall of predictive instances of top-ranking TF-MODISCO motifs in profile contribution scores from GM12878 ChromBPNet models trained on various subsampled datasets, using instances identified at 572M reads as ground truth (**Supp Fig. 2b**). Motif recall remains high and consistent across most read depths, with rare motifs lost only at low read depths (5-10M reads for bulk ATAC-seq and equivalent to ∼100 cells for scATAC-seq) (**Supp Fig. 2c)**.

### Variant Prediction Performance and Recommendation

Our extended analysis also provided recommendations for read depth requirements for variant effect prediction (**Supp Fig. 5**). The ideal depth is task-dependent, with quantitative effect size prediction being much more robust to low coverage than high-precision variant classification. We also found that models trained on pseudo-bulk scATAC-seq data consistently outperform bulk models at equivalent read depths, likely due to the higher signal-to-noise ratio from upstream single-cell quality control. For bulk ATAC-seq, we recommend a minimum of 25M reads for classification and 10-15M reads for effect size quantification. For higher-quality pseudo-bulk scATAC-seq, these minimums are lower, requiring only ∼15M reads for classification and as few as 2-5M reads for robust effect size prediction

### Important Caveats

These recommendations assume that data quality (signal-to-noise ratio e.g. TSS enrichment and fraction of reads in peaks) is comparable to pseudo-bulk GM12878 scATAC-seq data (ENCSR371ZNU). However, the feasibility of this approach has been demonstrated by several studies that successfully applied ChromBPNet to scATAC-seq data from complex primary tissues and rare developmental cell states: Ma et al.^178^ used ChromBPNet on human fetal heart scATAC-seq pseudobulk profiles to prioritize cell-type-specific variants in congenital heart disease; Naqvi et al.^152^ modeled transcription factor dosage responses in primary human cranial neural crest cells; and Marderstein et al.^184^ applied similar approaches to fetal brain neurons for interpreting de novo variants in neurodevelopmental disorders.

We emphasize that performance at minimal depths (5-10M reads) shows substantial degradation compared to optimal conditions, and users should carefully evaluate whether this performance level meets their analytical needs.

### 1.29 ChromBPNet model baselines with seq2PRINT bias correction

We applied seq2PRINT bias correction by downloading the h5 files containing the genome-wide Tn5 bias pre-computed using their Tn5 convolutional neural net model provided at https://zenodo.org/records/15170402. The resulting bias tracks were used in place of the BPNet bias model output described in Section 1.7 to train a seq2PRINT-factorized ChromBPNet. We trained this model on fold 0 of the observed profiles, following the training procedure in Section 1.4.

## TABLES

**Table 1:** Experimental configurations and training parameters for ChromBPNet models across ENCODE cell lines and chromatin accessibility assays. Table includes model identifiers and download locations for both ChromBPNet and associated bias models. The columns in this table provide detailed information on:

- Assay: ATAC-seq or DNase-seq
- Cell line: ENCODE canonical cell line (GM12878, HEPG2, IMR90, H1ESC, K562)
- Experiment ID: ENCODE/GEO accession for training data (BAMs and peaks)
- Read-depth: Post-processing read depth
- Sequencing: Paired-end or single-end characteristics
- Bias model: In vivo (same assay background) or transferred
- Filter parameter for backgrounds: Hyperparameter used to determine the threshold for filtering background regions in bias model training. Represents the fraction multiplied by the 0.01th quantile of total counts in peak regions.
- Bias model ID: ENCODE/Synapse ID for bias model download
- ChromBPNet model ID: ENCODE/Synapse ID for ChromBPNet model download

**Table 2:** Chromosome partitioning for 5-fold cross-validation of ChromBPNet models. This table outlines the division of chromosomes into training, validation, and test sets for each fold of the model training process. The columns provide the following information:

- Fold #: The identifier for each cross-validation fold
- Train chromosomes: Chromosomes used for training the model in each fold
- Validation chromosomes: Chromosomes used for model validation in each fold
- Test chromosomes: Chromosomes reserved for final model testing in each fold

**Table 3:** Metadata for training AFGR (African Genome Resources) ChromBPNet models across six African ancestries. The columns provide the following information:

- ATAC-seq expt. (ENCODE ENCID): The ENCODE experiment identifier for each ATAC-seq dataset.
- Cell-Line Name: The specific cell line of the experiment
- Ancestry: The African ancestry group associated with each cell line, including Gambian, Luhya, Esan, Maasai, Mende, and Yoruba

**Table 4:** This table presents the mapping between the regulatory elements studied in CAGI5 MPRA experiments to the ChromBPNet models trained on relevant celltypes.

- Locus (gene) name: The specific enhancer or promoter element assessed in the CAGI5 MPRA experiments.
- DNase-seq expt. (ENCODE ENCID): The ENCODE experiment identifier for the DNase-seq dataset used to train the ChromBPNet model.
- ChromBPNet model (ENCODE ENCID): The ENCODE or Synapse identifier for downloading the ChromBPNet model used for variant predictions at each gene locus.
- Celltype / Tissue: The cell type or tissue corresponding to the ENCODE experiment

**Table 5:** This table presents a comprehensive overview of the count and profile motifs identified by TF-MODISCO for various cell types and assay types (ATAC-seq or DNase-seq). Motifs are categorized based on their contribution to chromatin accessibility: “+” indicates activating motifs and “-” indicates repressing motifs. Columns show the number of motifs identified under different criteria: count and profile motifs for activating, repressing and both combined as well as those with at least 100 seqlets.

## SUPPLEMENTARY FILES

**Supp Files 1:** Series of TF-MODISCO reports, each generated using 30,000 seqlets derived from both the counts and profile heads of six different ATAC-seq models (1) BPNet trained on raw ATAC-seq data, (2) BPNet trained on bias-corrected signals from HINT, (3) BPNet trained with a bias track from TOBIAS, (4) BPNet trained on bias-corrected signals from TOBIAS (using softmax transformations and shifting the minimum to account for negative signals), (5) BPNet trained with a bias track from a simple CNN bias model, and (6) ChromBPNet. For motif identification, we employed TOMTOM to compare against the JASPAR database, reporting the top hit for each motif along with its corresponding q-value.

**Supp Files 2:** Series of TF-MODISCO reports, each generated using 30,000 seqlets derived from both the counts and profile heads of six different DNase-seq models: (1) BPNet trained on raw DNase-seq data, (2) BPNet trained on bias-corrected signals from HINT, (4) BPNet trained with a bias track from a simple CNN bias model, and (5) ChromBPNet. For motif identification, we employed TOMTOM to compare against the JASPAR database, reporting the top hit for each motif along with its corresponding q-value.

**Supp Files 3:** Series of TF-MODISCO reports, each generated using 30,000 seqlets derived from both the counts and profile heads of BPNet bias models trained in ATAC-seq and DNase-seq in the following canonical cell-lines GM12878, HEPG2, IMR90, H1ESC, K562. For motif identification, we employed TOMTOM to compare against the JASPAR database, reporting the top hit for each motif along with its corresponding q-value.

**Supp Files 4:** Series of TF-MODISCO reports, each generated using 30,000 seqlets derived from both the counts and profile heads of the following models: (1) BPNet bias model trained on raw naked DNA ATAC-seq, (2) BPNet bias model trained on raw naked DNA DNase-seq, (3) BPNet trained in GM12878 ATAC-seq with a naked DNA bias model (4) BPNet trained in GM12878 ATAC-seq with a HEPG2 chromatin bias model (5) BPNet trained in GM12878 DNase-seq with a naked DNA bias model and (6) BPNet trained in GM12878 DNase-seq with a HEPG2 chromatin bias model. For motif identification, we employed TOMTOM to compare against the JASPAR database, reporting the top hit for each motif along with its corresponding *q*-value.

**Supp Files 5:** Series of TF-MODISCO report table, each generated using 1 million seqlets derived from both the counts and profile heads of ChromBPNet models trained in ATAC-seq and DNase-seq in the following canonical cell-lines GM12878, HEPG2, IMR90, H1ESC, K562. For motif identification and annotation, we report two approaches: (1) TOMTOM comparison against the JASPAR database, reporting the top three hits for each motif along with their corresponding *q*-values and (2) Enrichment analysis using ENCODE TF ChIP-seq datasets, listing the top 10 TFs with the highest enrichment of peak overlap with predictive motif instances.

---

**Extended Figure 1:**
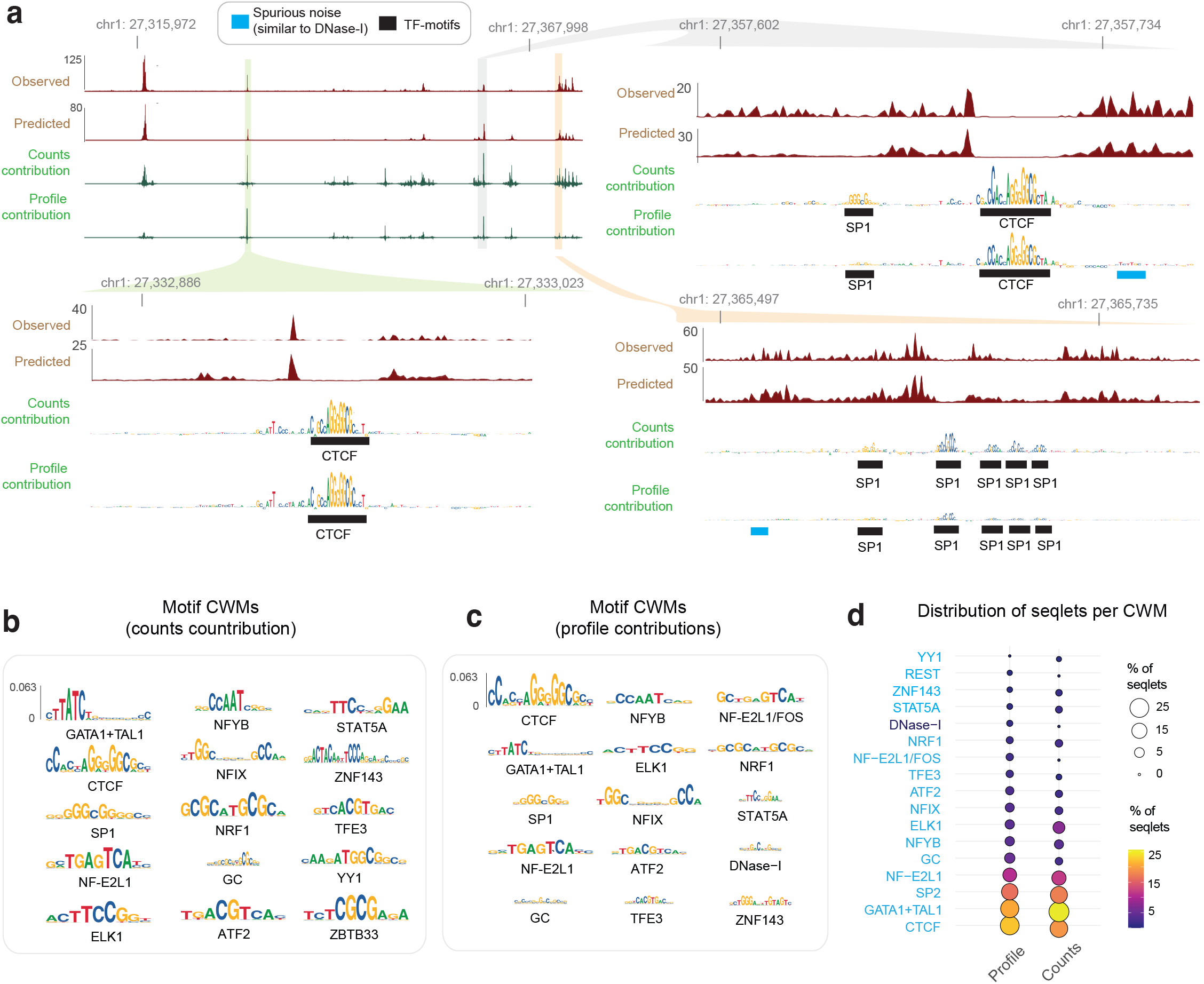
Convolutional neural networks accurately predict base-resolution DNase-seq profiles from local DNA sequence context but are confounded by DNase-I sequence bias. **(a)** Exemplar locus on chr11 (held-out test chromosome) showing observed (Obs) DNase-seq profiles, BPNet predicted (Pred) profiles at multiple resolutions and DeepLIFT sequence contribution scores to count and profile shape predictions. High count contribution scores highlight TF motif instances, whereas high profile contributions also capture spurious features that resemble DNase-I sequence preference. **(b)** Top 15 TF-MODISCO contribution weight matrix (CMW) motifs derived from count contribution scores of the K562 DNase-seq BPNet model map to well known TF motifs. **(c)** Top 15 TF-MODISCO contribution weight matrix (CMW) motifs derived from profile contribution scores of the K562 DNase-seq BPNet model map to well known TF motifs. **(d)** Comparison of the frequency of high-confidence predictive instances (seqlets) of each TF-MODISCO motif identified from the count and profile heads show that Dnase-I bias motifs moderately dominate the profile contribution scores over TF motifs.

**Extended Figure 2:**
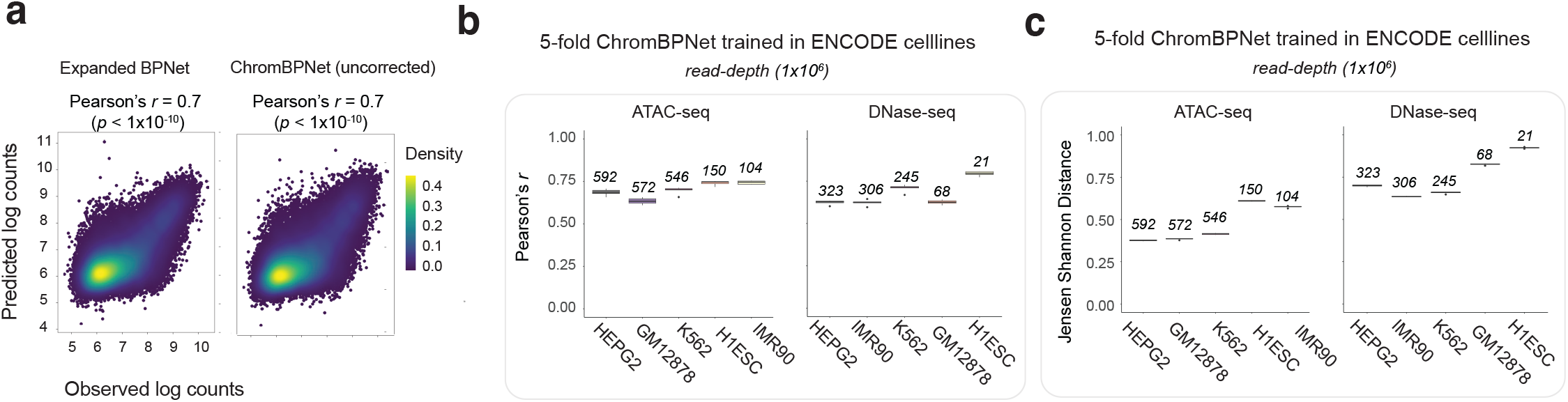
Comparison of count and profile head performance across different models, assays and cell-lines. **(a)** Uncorrected total count predictions from expanded BPNet models and bias-factorized ChromBPNet models of K562 ATAC-seq data show identical similarity to the measured total counts. **(b,c)** ChromBPNet models trained on ATAC-seq and DNase-seq data from diverse cell-lines show comparable and stable total count prediction performance (b) and profile prediction performance (c) across folds (box-plots), assays and cell-lines despite differences in read depth.

**Extended Figure 3:**
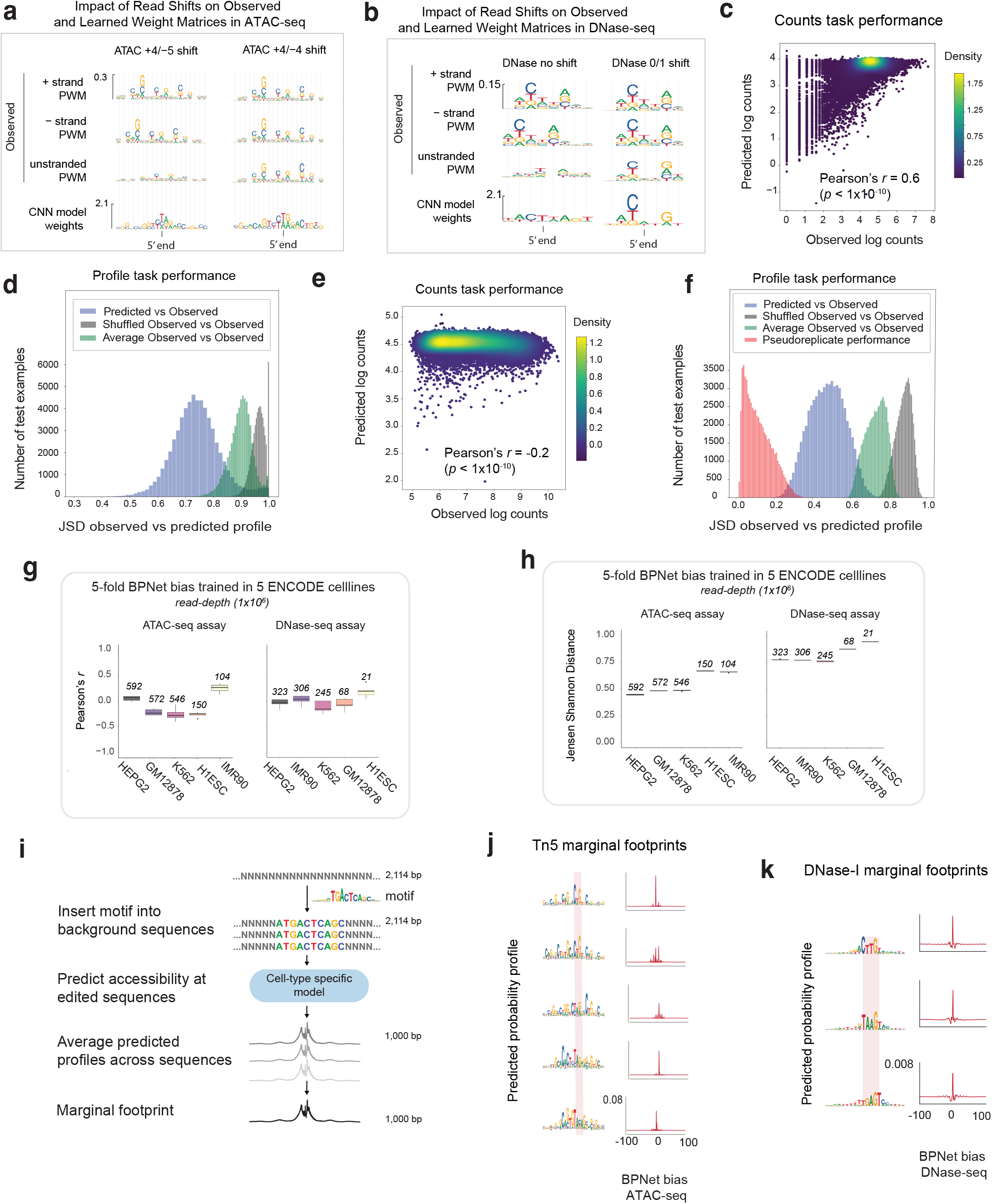
Performance evaluation of BPNet bias models: **(a,b)** Comparison of position frequency matrices derived from *k*-mers centered on 5’ ends of GM12878 ATAC-seq reads (a) and DNase-seq reads (b) (from chr20) mapping to the + strand only, - strand only, or both strands with different strand-specific shifts (+4/-5 or +4/-4 for ATAC-seq and 0/0 or 0/+1 for DNase-seq). The +4/-4 shift and the 0/+1 shifts correct the misalignment of strand-specific Tn5 and DNase-I PWMs respectively. Filter weights from the 1-filter CNN chromatin background bias model shows correct Tn5 and DNase-I bias motifs when trained on ATAC-seq reads with +4/-4 shift and DNase-seq reads with 0/+1 shifts respectively. **(c,d)** GM12878 ATAC-seq chromatin background BPNet bias model shows strong prediction of total counts in background regions of held-out test chromosomes (c) and strong prediction of profile shape (d) (blue distribution) relative to shuffled observed profiles (grey distribution) and average observed profiles (green distribution) in background regions of held-out test chromosomes. **(e)** GM12878 ATAC-seq chromatin background BPNet bias model shows poor prediction of total counts in ATAC-seq peak regions in held-out test chromosomes, indicating that bias does not affect total counts in peaks. **(f)** GM12878 ATAC-seq chromatin background BPNet bias model achieves high performance (blue distribution) at predicting profile shapes in peak regions in held-out test chromosomes compared to pseudoreplicate concordance (red), outperforming average observed profiles (green) and shuffled observed profiles (grey) baselines, suggesting Tn5 bias strongly influences profile shapes in peaks. **(g,h)** Performance of chromatin background BPNet bias models at predicting total counts (g) and profile shapes (h) in peak regions in held-out test chromosomes for ATAC-seq and DNase-seq data from diverse cell-lines. **(i)** Schematic of the marginal footprinting approach that averages model predictions over a library of background sequences containing a centrally embedded query motif. **(j,k)** Marginal footprints of various Tn5 (j) and DNase-I (k) TF-MODISCO bias motifs (rows) from predicted profiles of GM12878 ATAC-seq (j) and DNase-seq (k) chromatin background BPNet bias models.

**Extended Figure 4:**
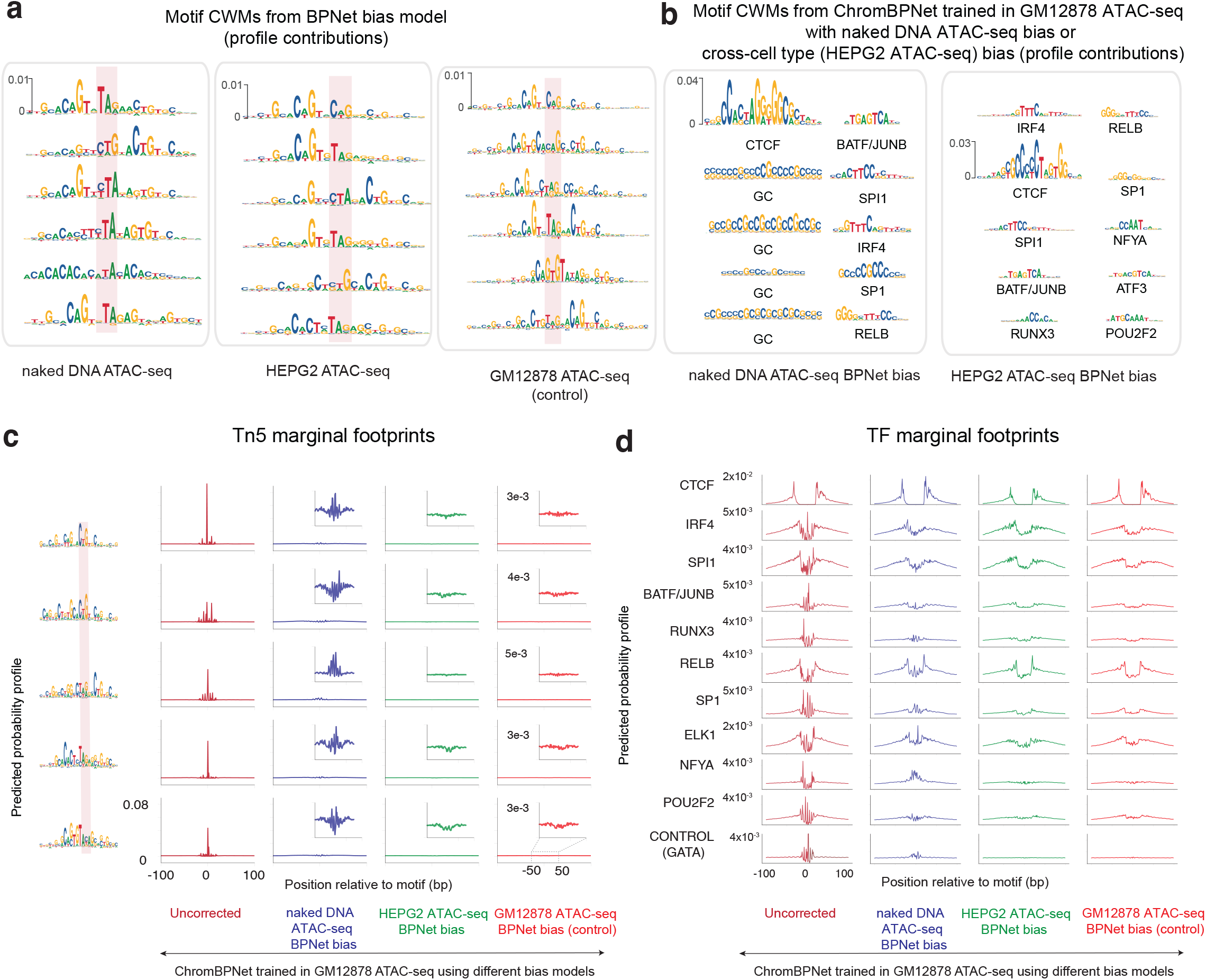
Tn5 bias models derived from naked DNA transposition libraries differ from those derived from chromatin background resulting in incomplete bias correction. **(a)** TF-MODISCO motifs derived from profile contribution scores of BPNet bias models trained on naked DNA transposition library (column 1) show stronger A/T content compared to those derived from bias models trained on HepG2 (col2) and GM12878 (col3) ATAC-seq background chromatin profiles. **(b)** Top ranked TF-MODISCO motifs derived from profile contribution scores of peak regions from GM12878 ATAC-seq ChromBPNet models coupled to BPNet bias model trained on naked DNA library (col1) or chromatin background from HepG2 ATAC-seq experiment (col2). The former show strong G/C contamination of learned TF motifs showing incomplete bias correction. **(c)** Marginal footprints of various TF-MODISCO Tn5 bias motifs (rows) using predicted profiles from GM12878 ATAC-seq ChromBPNet models using different bias models (no bias model, naked DNA bias model, HepG2 chromatin background bias model, GM12878 chromatin background bias model) show that the naked DNA BPNet bias model does not completely eliminate Tn5 bias and that chromatin bias models can provide very effective bias correction even across cell-lines. **(d)** Marginal footprints of various predictive TF motifs (rows) using predicted profiles from GM12878 ATAC-seq ChromBPNet models that use different bias models show that naked DNA BPNet bias model does not completely eliminate Tn5 bias (especially visible for NFYA, POU2F2 and the control motif of GATA1 that is not expressed in GM12878).

**Extended Figure 5:**
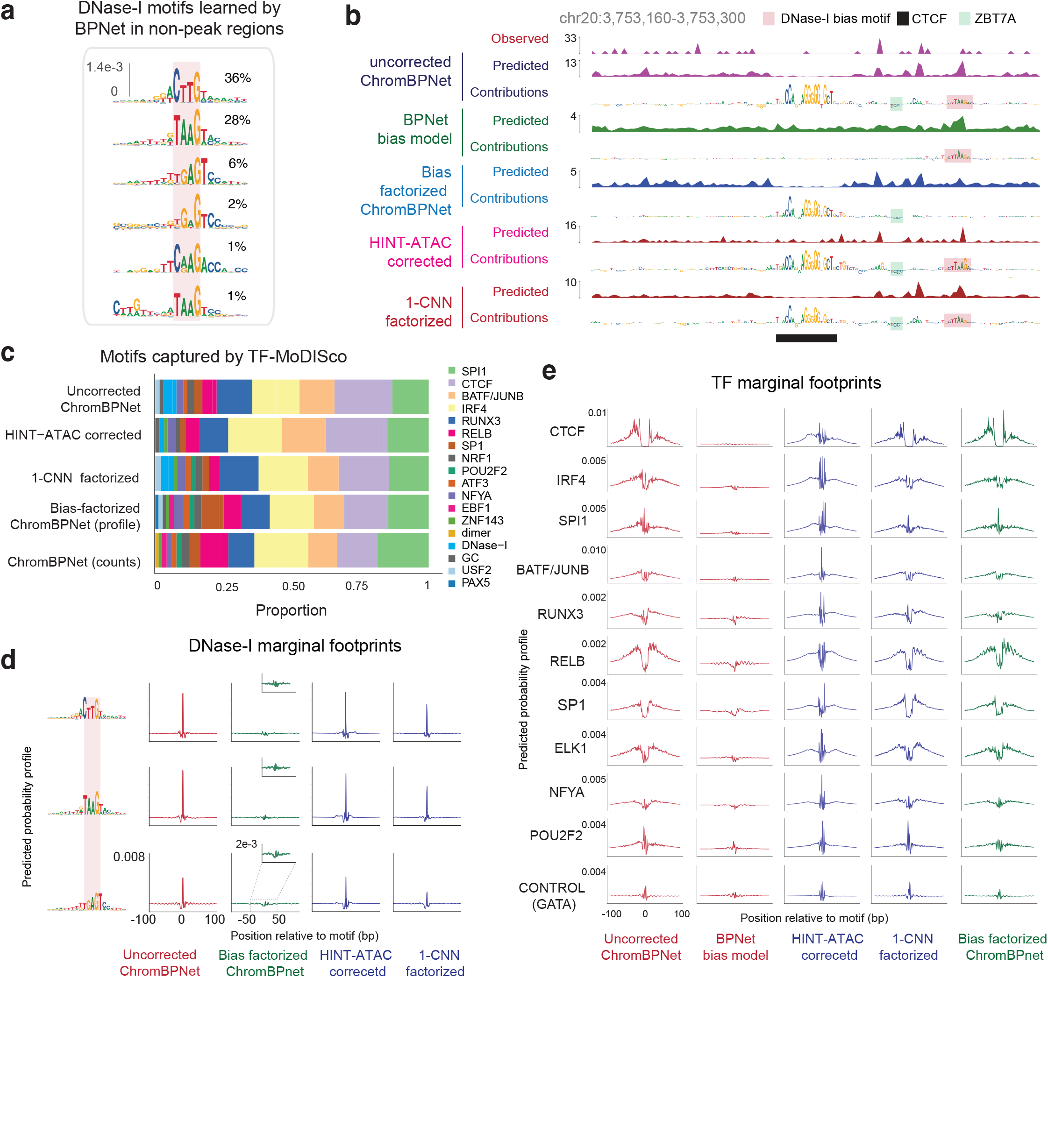
Bias-factorized ChromBPNet models of DNase-seq profiles coupled to neural network models of DNase-I bias deconvolve DNase-I bias from regulatory TF motif syntax. **(a)** TF-MODISCO motifs derived from profile contribution scores of HEPG2 BPNet bias model trained on chromatin background capture different variations of the canonical DNase-I sequence preference motif **(b)** Exemplar locus at chr20:3,753,160-3,753,300 showing several tracks from top to bottom: observed GM12878 DNase-seq profile; uncorrected predicted profile from ChromBPNet; uncorrected profile contribution scores from ChromBPNet shows spurious DNase-I bias; predicted DNase-I bias profiles from BPNet bias model shows moderate resemblance to observed and uncorrected profiles; profile contribution scores from bias model match spurious DNase-I bias features; bias-corrected predicted profile from ChromBPNet shows a strong, denoised latent footprint; bias-corrected profile contribution scores from ChromBPNet are devoid of spurious DNase-I bias highlighting CTCF and ZBTB7A motifs; predicted profiles from ChromBPNet model using HINT-ATAC’s bias correction method; profile contribution scores from HINT-ATAC ChromBPNet model show spurious DNase-I bias; predicted profiles from ChromBPNet model that uses a simplified 1-filter CNN bias model; profile contribution scores from ChromBPNet model with 1-filter CNN bias model shows spurious DNase-I bias **(c)** Relative frequency of high-confidence predictive instances (seqlets) of TF-MODISCO motifs derived from profile contribution scores of various ChromBPNet models trained on GM12878 DNase-seq data using different bias models (no bias model, 1-filter CNN bias model and BPNet bias model) or pre-corrected profiles (HINT-ATAC) shows that DNase-I bias motifs dominate all but the ChromBPNet model coupled with the BPNet bias model. **(d)** Marginal footprints of various TF-MODISCO DNase-I bias motifs (rows) using predicted profiles from GM12878 DNase-seq ChromBPNet models that use different bias models or pre-corrected profiles (cols) show that only the BPNet bias model enables near optimal correction of DNase-I bias. **(e)** Marginal footprints of various predictive TF motifs (rows) using predicted profiles from GM12878 DNase-seq ChromBPNet models that us different bias models or pre-corrected profiles (cols) show that only the BPNet bias model removes confounding DNase-I bias.

**Extended Figure 6:**
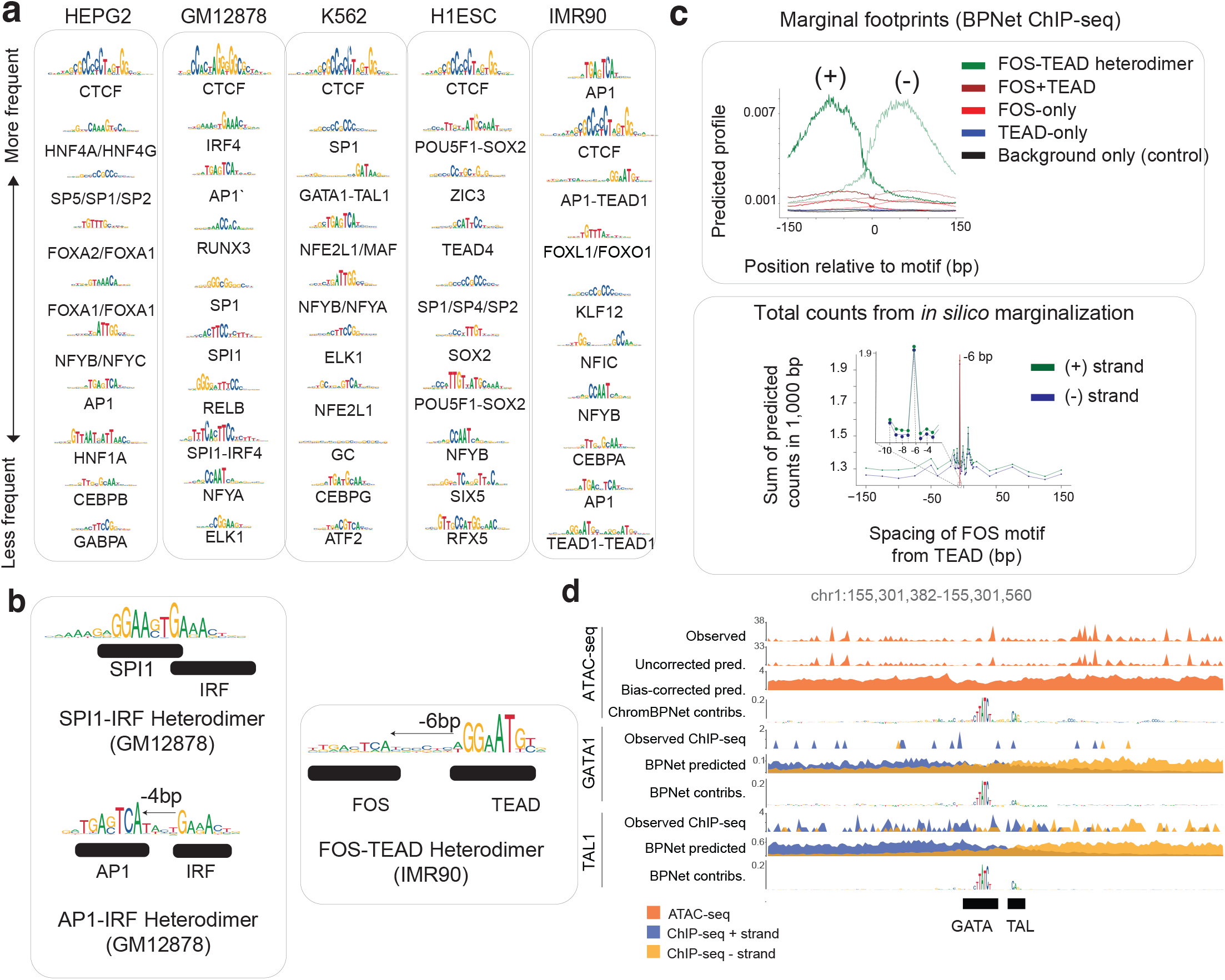
Additional evidence for ChromBPNet derived TF motif lexicons, cooperative composite elements and predictive motif instances with support from TF ChIP-seq. **(a)** Top 10 TF-MODISCO motifs (ranked by number of predictive motif instances) derived from profile contribution scores from ATAC-seq ChromBPNet models of diverse cell-lines. Motifs names containing / refer to multiple TFs of the same family that match the motif. Motif names containing - refer to composite elements. **(b)** Exemplar composite motifs identified by TF-MODISCO from contribution scores derived from ChromBPNet models trained on ATAC-seq and DNase-seq data from IMR90 (FOS-TEAD composite) and GM12878 (SPI1-IRF and AP1-IRF composites). **(c)** Marginal profiles at FOS motif, TEAD motif, FOS-TEAD composite motif and the sum of FOS and TEAD marginal profiles from BPNet model of FOS TF ChIP-seq data from IMR90 demonstrates the super-additive cooperative effect of the FOS-TEAD composite on FOS binding (top). Variation of the strength (maximum) of predicted marginal FOS ChIP-seq profiles anchored at FOS and TEAD motifs with variable spacing inserted in background sequences shows strong cooperative effects on FOS binding only at the fixed 6 bp spacing (bottom). **(d)** Exemplar accessible region in chr1 near the PKLR gene in K562 displaying the observed profile from ATAC-seq and GATA1, TAL1 ChIP-seq data, alongside uncorrected and bias corrected predicted profiles, and count contribution scores from ATAC-seq ChromBPNet model and TF ChIP-seq BPNet models. Predictive motif instances from ChromBPNet and BPNet models are in strong agreement.

**Extended Figure 7:**
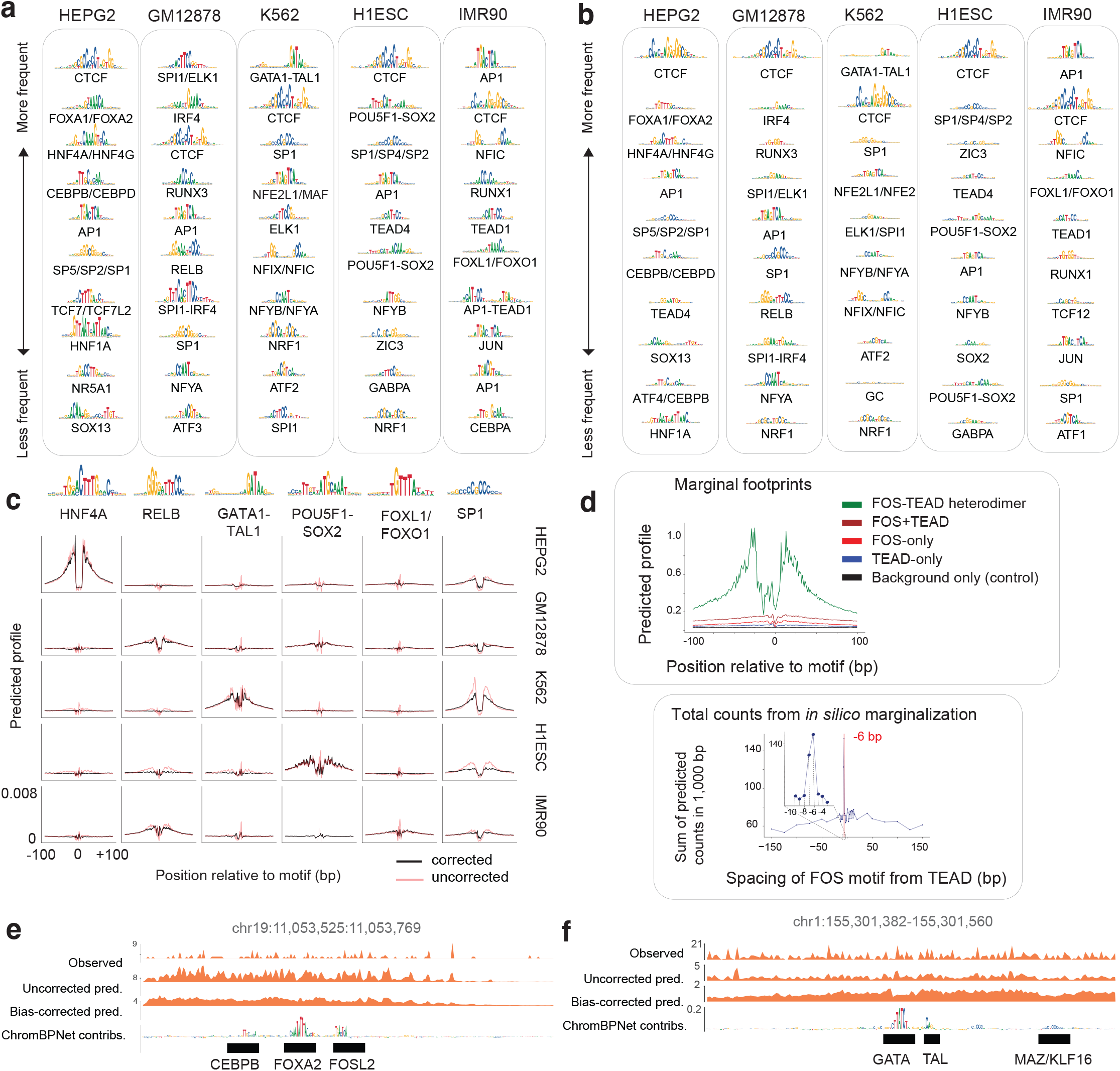
ChromBPNet models of DNase-seq data reveal compact TF motif lexicons, cooperative composite elements and predictive motif instances that influence chromatin accessibility and TF occupancy. **(a,b)** Top 10 TF-MODISCO motifs (ranked by number of predictive motif instances) derived from count contribution scores (a) and profile contribution scores (b) from DNase-seq ChromBPNet models of diverse cell-lines. Motifs names containing / refer to multiple TFs of the same family that match the motif. Motif names containing - refer to composite elements. **(c)** Marginal footprints for TF-MODISCO motifs using normalized uncorrected (red) and bias-corrected profile predictions (black) from DNase-seq ChromBPNet models of diverse cell-lines. Bias-correction reveals cell-type specificity of footprints. **(d)** Marginal footprints for the FOS motif, TEAD motif, FOS-TEAD composite motif and the sum of FOS and TEAD marginal footprints demonstrates the super-additive cooperative effect of the FOS-TEAD composite (top). Variation of the strength (maximum) of marginal footprints anchored at FOS and TEAD TF motifs with variable spacing inserted in background sequences shows strong cooperative effects only at the fixed 6 bp spacing (bottom). **(e)** Exemplar accessible region in chr19 near the LDLR gene in HEPG2 displaying the observed profile from DNase-seq, alongside uncorrected and bias corrected predicted profiles, and count contribution scores from DNase-seq ChromBPNet model. **(f)** Exemplar accessible region in chr1 near the PKLR gene in K562 displaying the observed profile from DNase-seq, alongside uncorrected and bias corrected predicted profiles, and count contribution scores from DNase-seq ChromBPNet model.

**Extended Figure 8:**
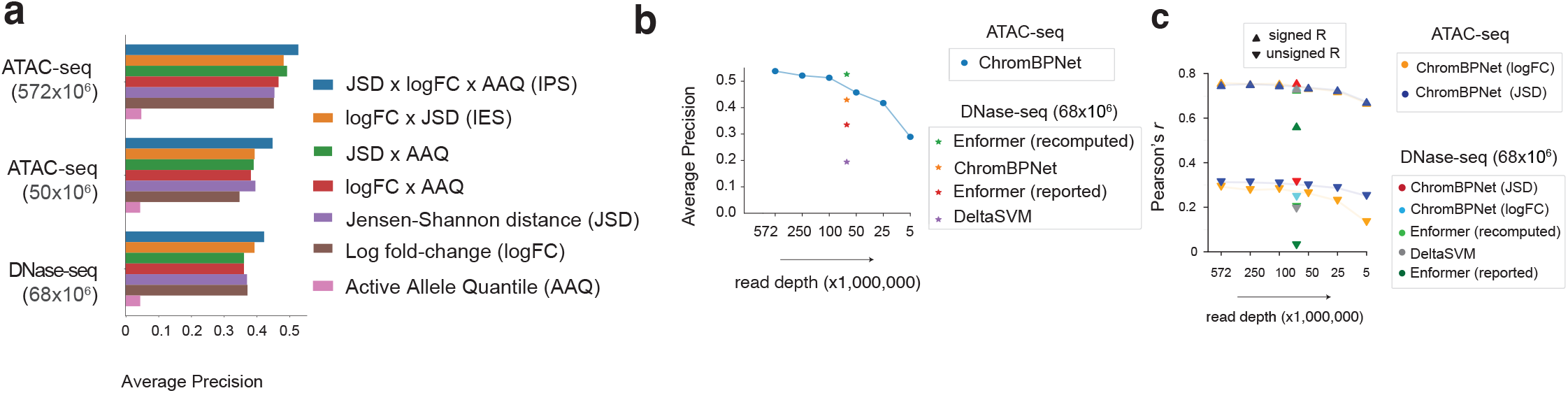
Impact of training data read depth and variant scoring measures on ChromBPNet’s variant classification and effect prediction performance for DNase-seq QTLs (dsQTLs) in Yoruban lymphoblastoid cell lines (LCLs) **(a)** Comparison of variant classification performance (average precision (AP)) of ChromBPNet models trained on GM12878 DNase-seq and ATAC-seq data (with two different read depths) at discriminating significant dsQTLs in Yoruban LCLs from control variants. Performance evaluated using different variant effect measures shows that the Integrative Prioritization Score (IPS) outperforms Active Allele Quantile (AAQ), log-fold change in predicted coverage (logFC), Jensen Shanon distance (JSD) of predicted allelic profiles, JSD x AAQ and logFC x AAQ. **(b)** LCL dsQTL variant classification performance (y-axis) of ChromBPNet models trained on GM12878 ATAC-seq data decreases with read depth (572M, 250M, 100M, 50M, 25M, and 5M reads) of the training dataset (x-axis). Performance of GM12878 DNase-seq models (gkmSVM, ChromBPNet and Enformer) are also shown for comparison. **(c)** Correlation between LCL dsQTL observed effect sizes (signed and unsigned) and GM12878 ATAC-seq ChromBPNet model’s predicted allelic log-fold changes of total counts or changes in profile shape (JSD) remain stable across most read depths (572M, 250M, 100M, 50M, 25M, and 5M reads) of the training set. Effect size prediction performance of various GM12878 DNase-seq models (gkmSVM, ChromBPNet and Enformer) are shown for comparison.

**Extended Figure 9:**
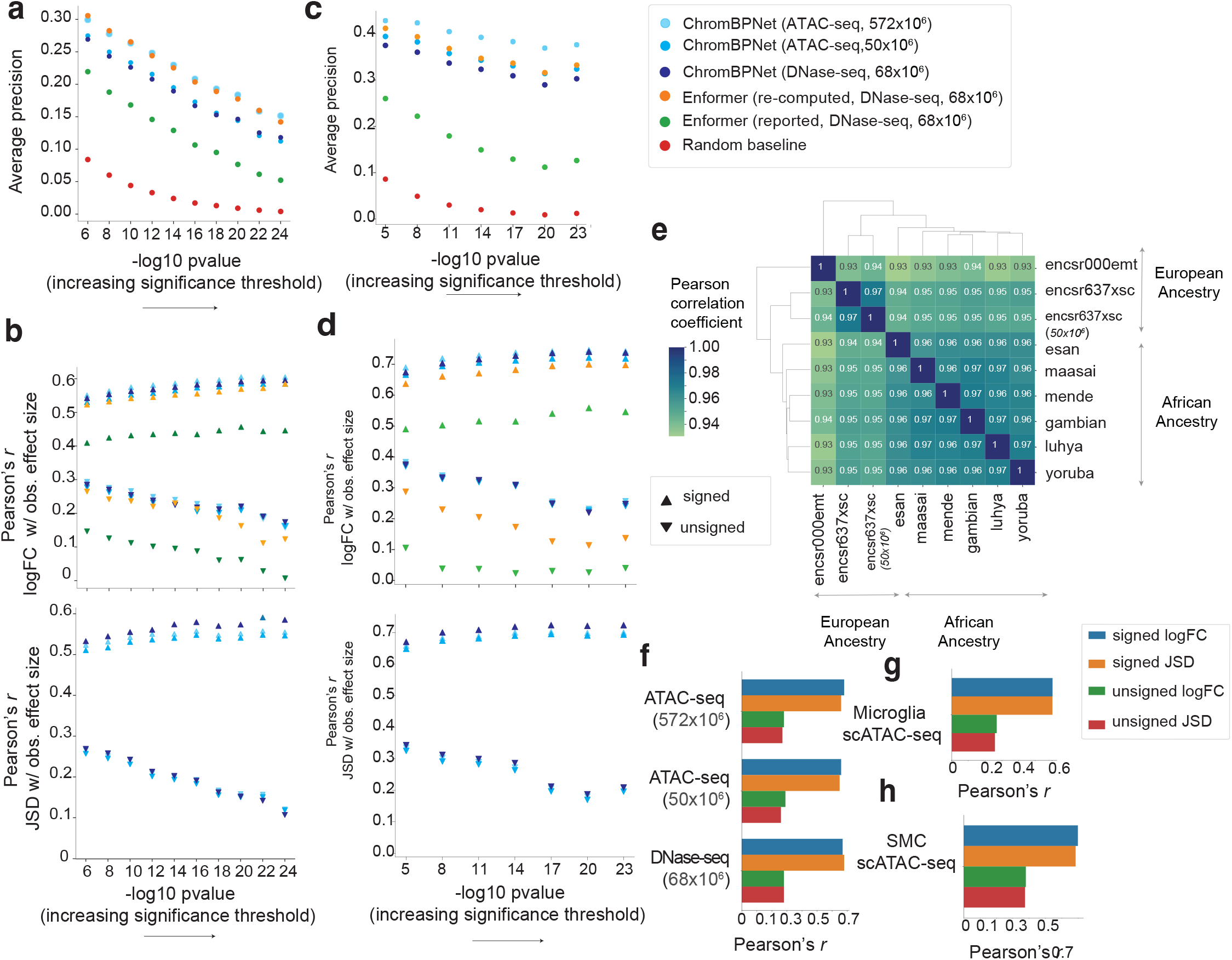
Comparison of ChromBPNet and Enformer models for variant classification and effect size prediction caQTLs across various significance thresholds. **(a)** Comparison of variant classification performance (average precision (AP)) of ChromBPNet and Enformer trained on GM12878 DNase-seq or ATAC-seq data at discriminating significant ATAC-seq QTLs (caQTLs) from European LCLs at varying caQTL significance thresholds (x-axis) versus control variants. **(b)** Comparison of correlation between observed effect sizes (signed and unsigned) of significant caQTLs from European LCLs selected at varying thresholds (x-axis) and predicted effect sizes (top: allelic log fold change of total counts, bottom: JSD between allelic profiles) of ChromBPNet and Enformer models trained on GM12878 DNase-seq or ATAC-seq data. **(c)** Comparison of variant classification performance of ChromBPNet and Enformer trained on GM12878 DNase-seq or ATAC-seq data at discriminating significant ATAC-seq QTLs (caQTLs) from African LCLs at varying caQTL significance thresholds (x-axis) versus control variants. **(d)** Comparison of correlation between observed effect sizes (signed and unsigned) of significant caQTLs from African LCLs selected at varying thresholds (x-axis) and predicted effect sizes (top: allelic log fold change of total counts, bottom: JSD between allelic profiles) of ChromBPNet and Enformer models trained on GM12878 DNase-seq or ATAC-seq data. **(e)** Predicted allelic effects on profile shape (Jensen Shannon Distance (JSD)) for the significant African LCL caQTLs from ChromBPNet models trained on ATAC-seq data from reference LCLs from different ancestry groups/subgroups (each row/column in the matrix) are highly correlated. **(f)** Correlation of observed allelic imbalance (signed and unsigned) of ATAC-seq coverage (log fold changes) at heteroyzgous variants in the African caQTL LCL cohort with predicted effect sizes (logFC and JSD) from ChromBPNet models trained on GM12878 ATAC-seq (at two read depths) or DNase-seq. **(g)** Correlation of observed effect sizes (regression betas) of significant caQTLs in microglia with various predicted effect sizes (logFC and JSD) from a ChromBPNet model trained on a reference microglia scATAC-seq pseudobulk dataset **(h)** Correlation of the observed effect sizes (effect sizes from the RASQUAL method) of significant caQTLs in coronary artery smooth muscle cell-lines with various predicted effect sizes (logFC and JSD) from a ChromBPNet model trained on a reference caSMC scATAC-seq pseudobulk dataset.

**Extended Figure 10:**
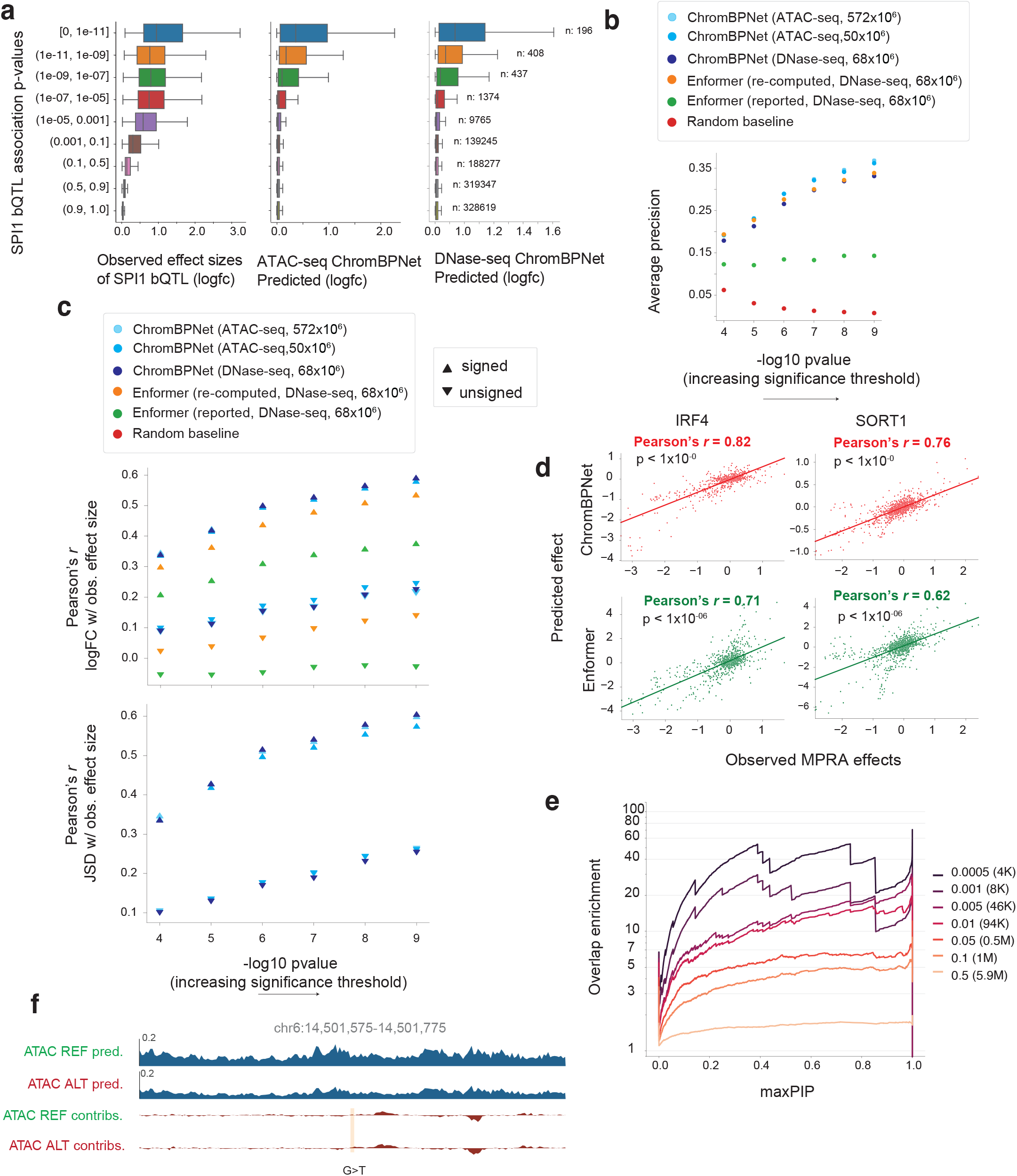
Additional evidence for benchmarks of ChromBPNet’s variant effect prediction performance against SPI1 binding QTLs, massively parallel reporter activity of cRE mutagenesis and fine mapped GWAS variants. **(a)** Comparison of distribution of observed SPI1 bQTL effect sizes (left), predicted effect sizes from ChromBPNet model of GM12878 ATAC-seq data (middle) and GM12878 DNase-seq data (right) stratified by p-value of the bQTL association significance (y-axis). More significant bQTL associations show stronger effect sizes. **(b)** Comparison of variant classification performance (average precision (AP)) of ChromBPNet and Enformer trained on GM12878 DNase-seq or ATAC-seq data at discriminating significant SPI1 bQTLs in LCLs at varying bQTL significance thresholds (x-axis) versus control variants. **(c)** Comparison of correlation between observed effect sizes (signed and unsigned) of significant bQTLs from LCLs selected at varying thresholds (x-axis) and predicted effect sizes (top: allelic log fold change of total counts, bottom: JSD between allelic profiles) of ChromBPNet and Enformer models trained on GM12878 DNase-seq or ATAC-seq data. **(d)** Comparison of the observed MPRA effects (x-axis) against the predicted effects from Enformer (top plot) or ChromBPNet (bottom plot) for all mutants of the IRF4 and SORT1 cREs. **(e)** Benchmark testing overlap enrichment (y-axis) of fine mapped variants from GWAS loci associated with red blood cell traits that pass various posterior probability (PIP) thresholds (x-axis) with variants predicted to have strong effects (at various thresholds) by ChromBPNet model trained on K562 DNase-seq data. Each curve corresponds to a different ChromBPNet effect size threshold (the right panel shows the threshold scores and the number of variants that pass each threshold). Enrichments increase with increasing PIP and more stringent ChromBPNet thresholds. (f) Example locus around variant hs2599, which is a de novo variant from a patient with severe, undiagnosed neurodevelopmental disorder (NDD). Bias-corrected predicted profiles and contribution scores are shown for both alleles from a ChromBPNet model trained on scATAC-seq pseudobulk data from glutamatergic neuron cell-type from human fetal brain tissue. The G allele and T allele are predicted to result in no change in accessibility.

**Supplementary Figure 1:**
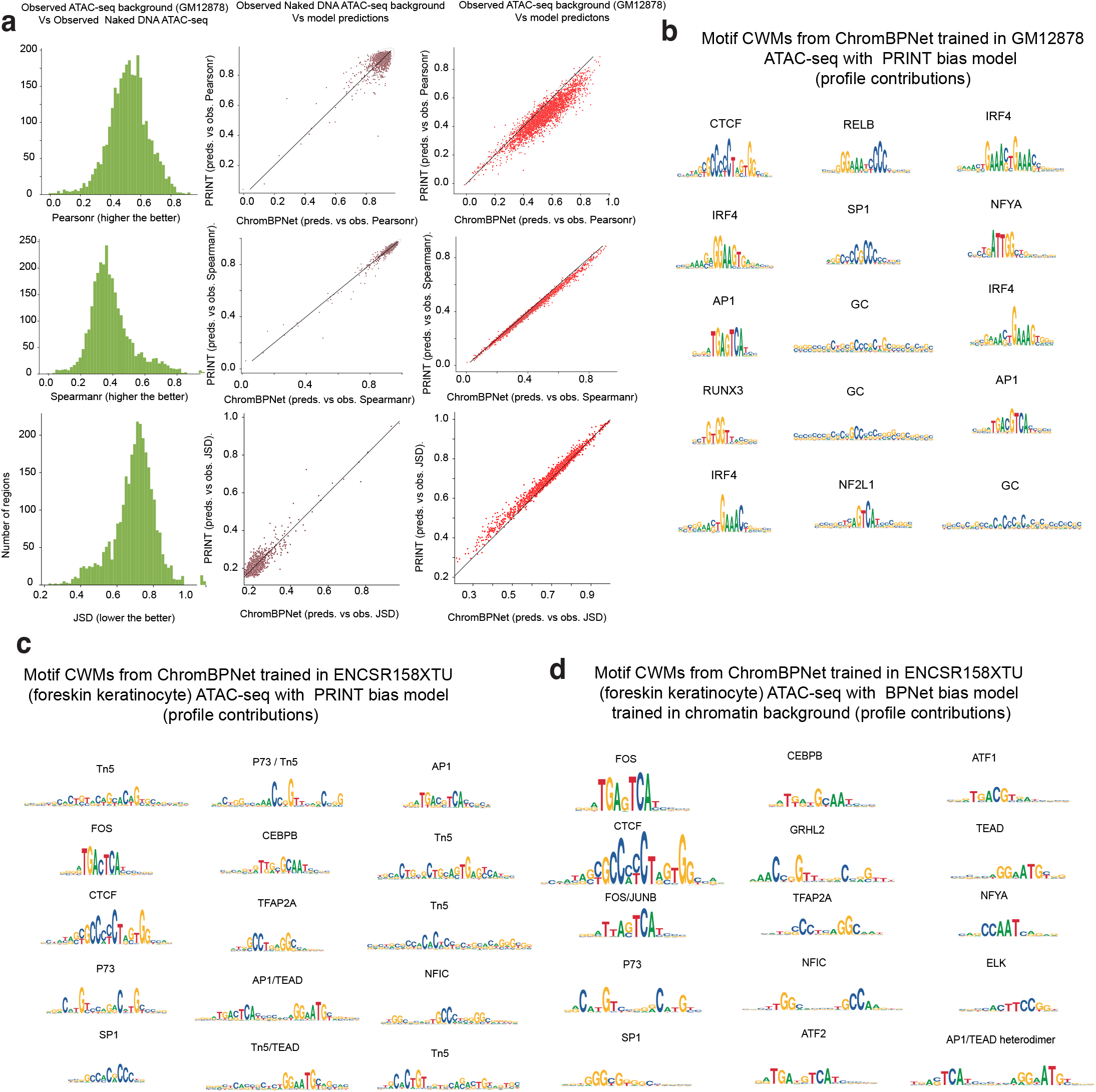
seq2PRINT comparison: ChromBPNet provides stronger experiment-matched bias correction and bias-corrected learned motifs. (a) Bias-model evaluation across 1,000 high-coverage 100-bp BAC regions. Left: distributions of Pearson (r), Spearman (ρ), and JSD between observed naked DNA and GM12878 ATAC-seq profiles, showing that naked DNA does not consistently match chromatin background. Middle/right: region-wise comparisons of seq2PRINT versus ChromBPNet; both are similar on naked DNA, but ChromBPNet consistently improves agreement to chromatin background (higher r/ρ, lower JSD). (b–d) TF-MoDISco contribution weight matrices (CWMs) from ChromBPNet models trained with different bias corrections: using seq2PRINT on GM12878 ATAC-seq (b) or foreskin keratinocyte ATAC-seq (c) leaves residual GC/Tn5-like artifacts, whereas an experiment-matched chromatin-background bias model (d) removes technical patterns and yields cleaner biological motifs.

**Supplementary Figure 2:**
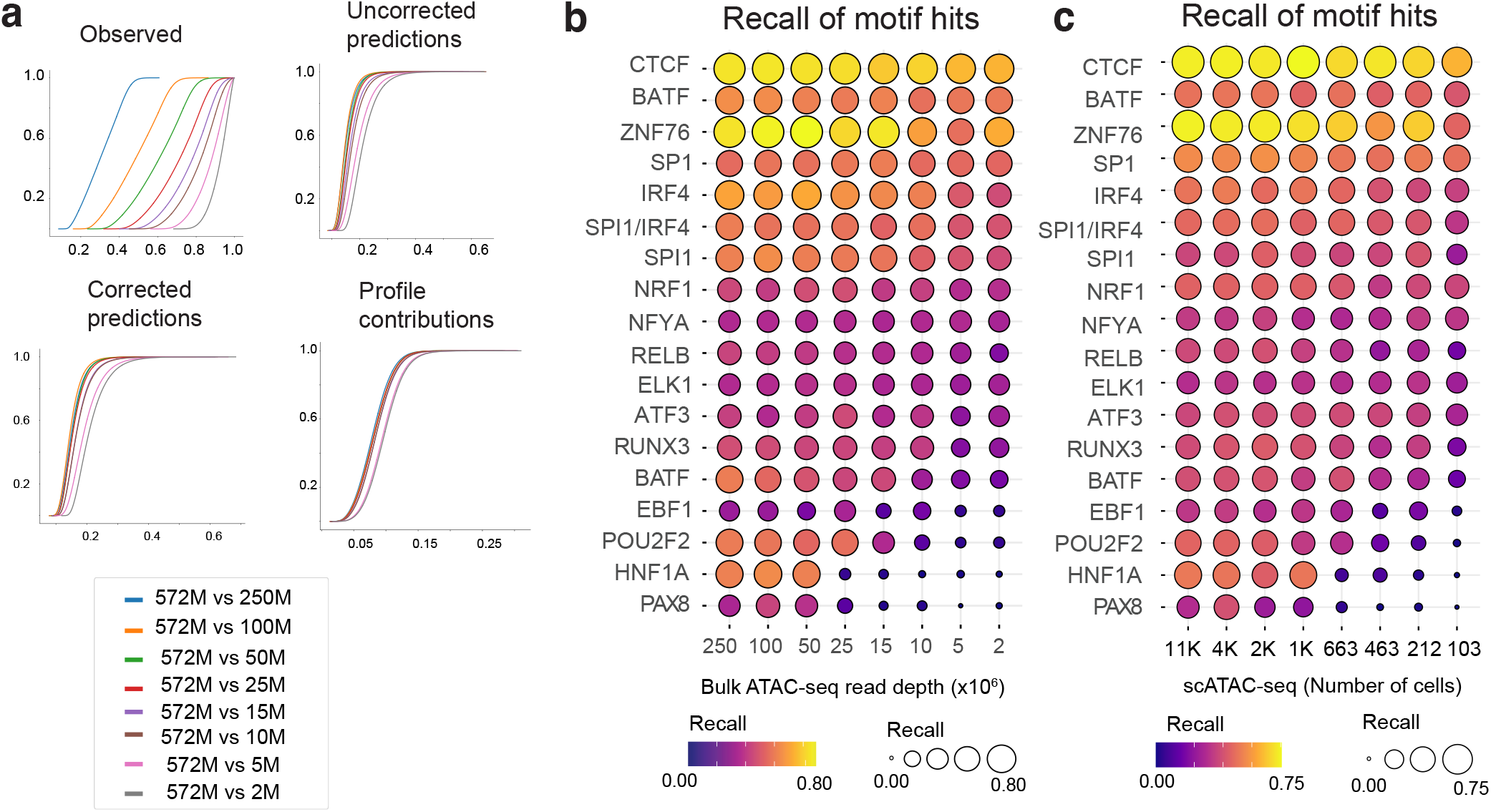
Granular downsampling in bulk and pseudobulk scATAC-seq shows stable profile imputation and motif instance recovery until low coverage. (a) Jensen–Shannon divergence (JSD) between profiles from the maximal-depth dataset and progressively downsampled datasets over peak regions. Curves summarize discordance for observed profiles, uncorrected ChromBPNet predictions, bias-corrected ChromBPNet predictions, and bias-corrected profile contribution tracks (each compared to the maximal-depth reference). (b) Recall of predictive motif instances for top TF-MoDISco motifs, computed from profile contribution scores of models trained at each bulk ATAC-seq read depth (250M, 100M, 50M, 25M, 15M, 10M, 5M, 2M reads). Ground-truth instances are defined as the predictive motif hits identified in the maximal-depth bulk model (572M reads). (c) The same motif-instance recall analysis for pseudobulk GM12878 scATAC-seq (ENCSR371ZNU) generated by aggregating progressively fewer high-quality cells (TSS enrichment > 6; >5,000 UMIs/cell). Cell subsampling yields pseudobulks with read depths matched to the bulk series (250M → 2M reads), corresponding to ∼11k, 4k, 2k, 1k, 663, 463, 212, and 103 cells, respectively. Across both bulk and pseudobulk series, predicted profiles and contribution tracks remain similar to the maximal-depth reference down to ∼25M reads, with clear degradation emerging at ∼10M reads; rare motifs show reduced recall only at the lowest depths (5–10M reads in bulk and ∼100–200 cells in scATAC pseudobulk).

**Supplementary Figure 3:**
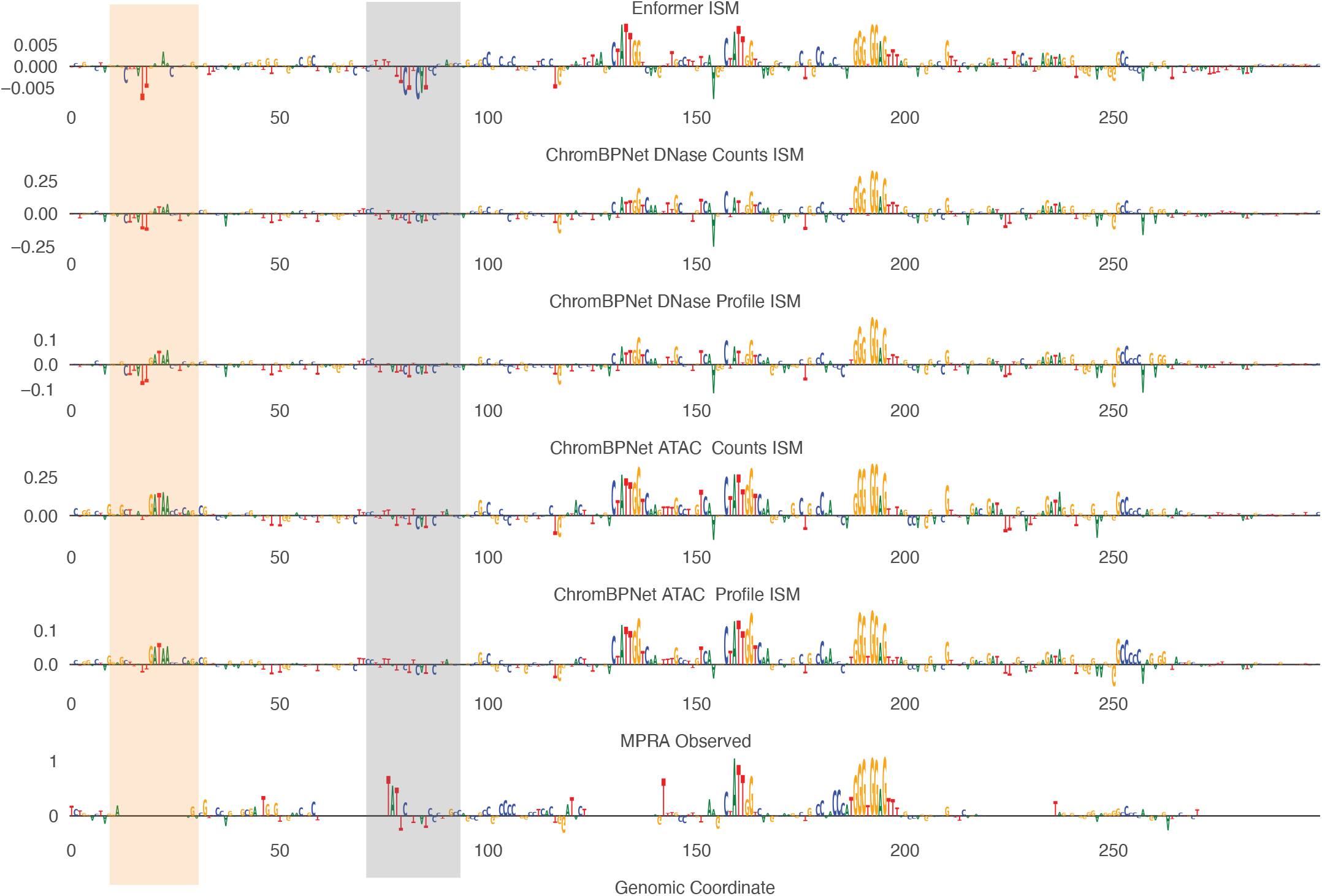
In silico saturation mutagenesis of the HBG1 MPRA element for ChromBPNet and Enformer. DNase head compared to experimental mutagenesis MPRA data. ISM contribution profiles (profile and count heads) for ChromBPNet (ATAC-and DNase-trained) and Enformer across the HBG1 MPRA sequence show that Enformer identifies a strong repressive motif ∼85 bp from the left that is much weaker in ChromBPNet and has a positive effect in the MPRA. ChromBPNet exclusively localizes an activating GATA motif within the first ∼50 bp which Enformer does not. The MPRA did not measure effects at this motif.

**Supplementary Figure 4:**
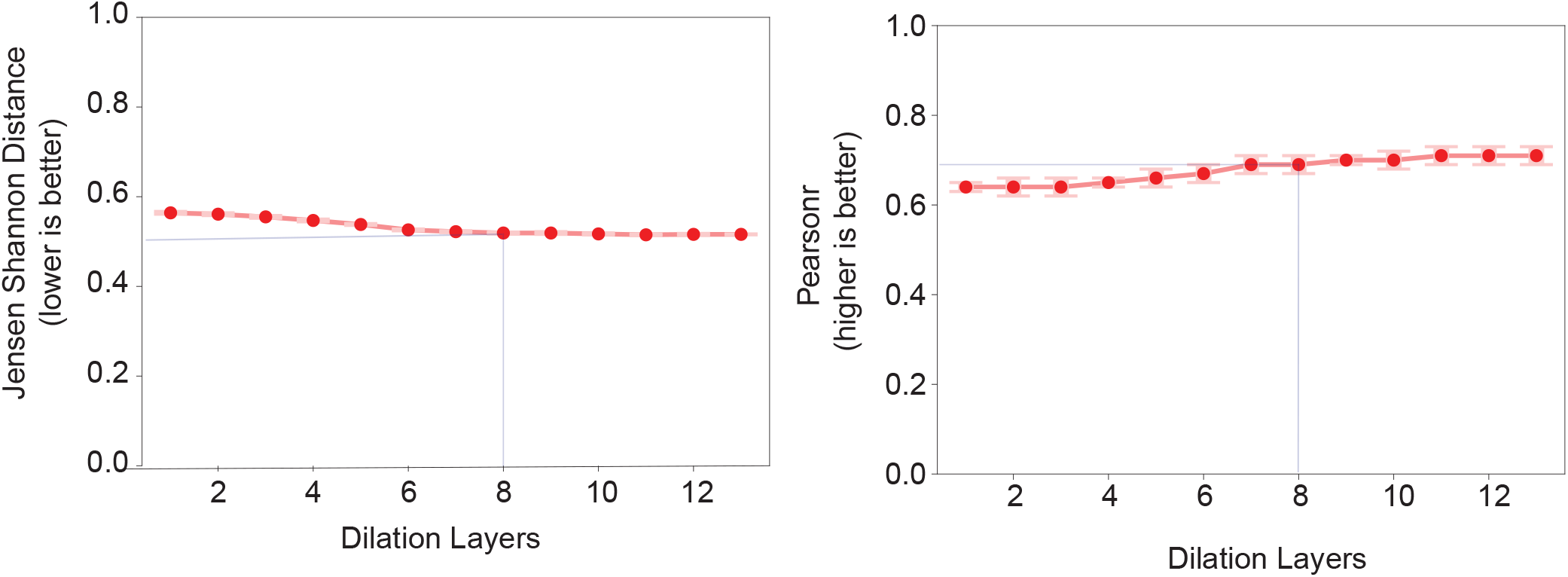
Choosing ChromBPNet receptive field size by varying dilated-convolution depth. Performance as a function of the number of dilated convolutional layers. Left: profile-shape accuracy (JSD; lower is better) improves rapidly and then saturates. Right: total-count accuracy (Pearson r; higher is better) similarly saturates. The selected 8-layer architecture (vertical line) balances receptive field with diminishing returns from additional layers.

**Supplementary Figure 5:**
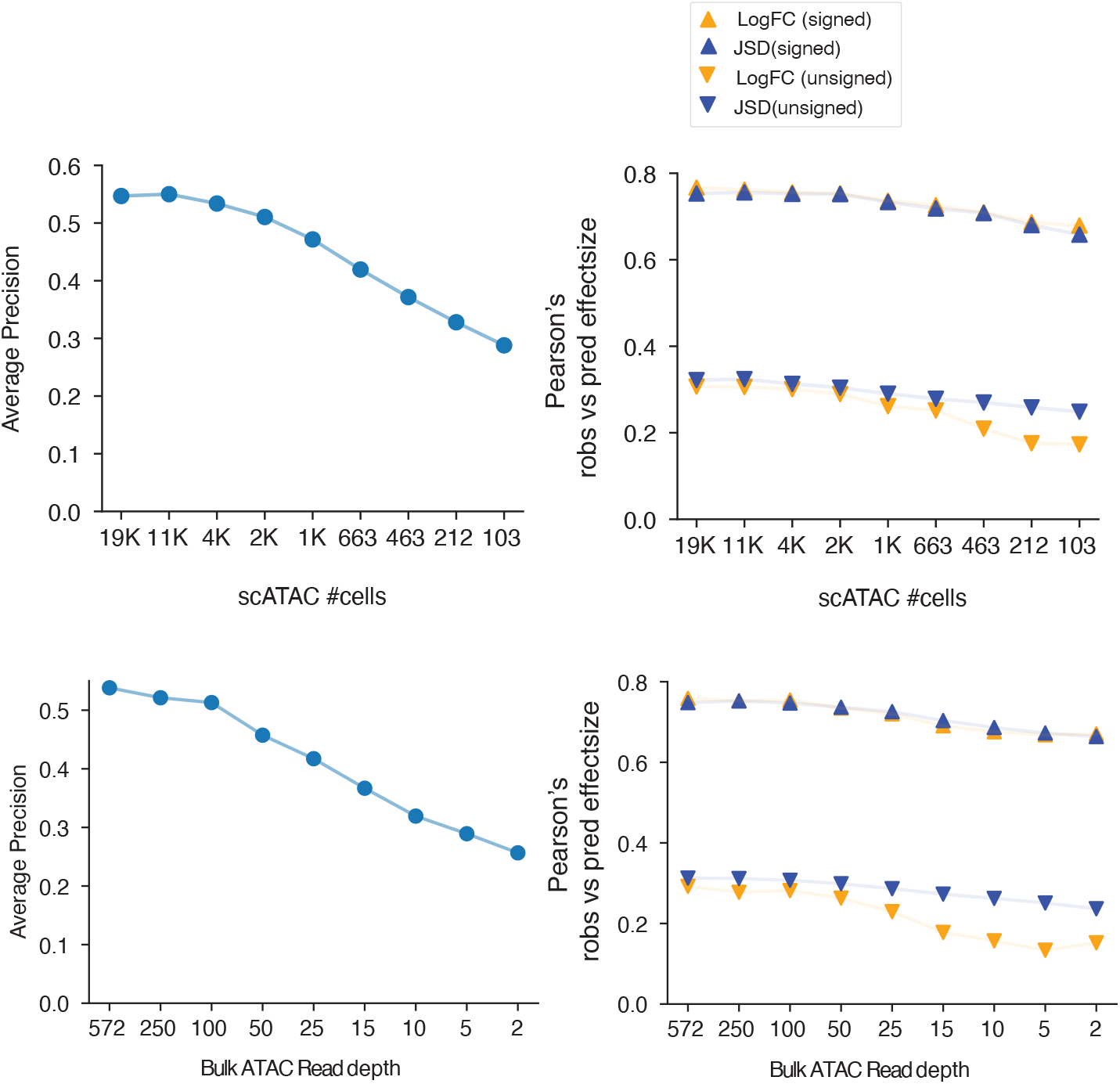
Sequencing depth requirements differ for dsQTL classification versus quantitative effect-size prediction, and pseudobulk scATAC models are more robust at matched read depth. ChromBPNet models were trained on GM12878 pseudobulk scATAC-seq profiles aggregated from decreasing numbers of high-quality cells (top row; 19k → 103 cells) and on bulk ATAC-seq profiles downsampled to matched read depths (bottom row; 250M → 2M reads). Left panels: average precision for classifying LCL dsQTLs using ChromBPNet-predicted allelic effects. Right panels: Pearson correlation between observed dsQTL effect sizes and predicted variant effects, shown for both signed and unsigned metrics. Predicted effects are computed either as allelic log-fold change in total accessibility (total-count head) or as allelic change in profile shape (JSD between allele-specific predicted profiles), as indicated. Quantitative effect-size correlations are comparatively stable down to low depth, whereas high-precision classification degrades earlier. At matched read depth, models trained on pseudobulk scATAC-seq consistently outperform bulk-trained models, consistent with improved signal-to-noise from upstream single-cell QC.

