## Supplementary Files 1 for "ChromBPNet: bias factorized, base-resolution deep learning models of chromatin accessibility reveal cis-regulatory sequence syntax, transcription factor footprints and regulatory variants": fig1a_k562_ATAC_raw_bpnet_uncorrected_counts_modisco.pdf

| pattern | num_seqlets | cwm_fwd | cwm_rev | TOMTOM_match | TOMTOM_qval | TOMTOM_match_logo |
|---|---|---|---|---|---|---|
| pos_patterns.pattern_0 | 3831 | | | GATA4_HUMAN.H11MO.0.A | 9.627200e-03 | |
| pos_patterns.pattern_1 | 3616 | | | KLF12_HUMAN.H11MO.0.C | 2.789230e-05 | |
| pos_patterns.pattern_2 | 1654 | | | BACH2_MOUSE.H11MO.0.A | 2.755450e-04 | |
| pos_patterns.pattern_3 | 1639 | | | CTCF_MOUSE.H11MO.0.A | 4.015900e-16 | |
| pos_patterns.pattern_4 | 1513 | | | ELK1_ETS_1 | 1.183960e-01 | |
| pos_patterns.pattern_5 | 700 | | | NFYB_HUMAN.H11MO.0.A | 9.192150e-03 | |
| pos_patterns.pattern_6 | 518 | | | NRF1_MOUSE.H11MO.0.A | 6.193500e-07 | |
| pos_patterns.pattern_7 | 447 | | | NRF1_NRF_1 | 7.521700e-03 | |
| pos_patterns.pattern_8 | 422 | | | FOSL2+JUN_MA1131.1 | 1.396980e-04 | |
| pos_patterns.pattern_9 | 376 | | | TAF1_HUMAN.H11MO.0.A | 1.155700e-01 | |
| pos_patterns.pattern_10 | 318 | | | SPIB_MOUSE.H11MO.0.A | 9.220600e-04 | |
| pos_patterns.pattern_11 | 282 | | | E2F2_HUMAN.H11MO.0.B | 1.321570e-01 | |
| pos_patterns.pattern_12 | 274 | | | ATF4_MOUSE.H11MO.0.A | 9.997960e-06 | |
| pos_patterns.pattern_13 | 239 | | | NFIC_HUMAN.H11MO.0.A | 2.630340e-07 | |
| pos_patterns.pattern_14 | 204 | | | ZN143_MOUSE.H11MO.0.A | 5.362850e-20 | |
| pos_patterns.pattern_15 | 169 | | | SP2_HUMAN.H11MO.0.A | 8.109120e-04 | |
| pos_patterns.pattern_16 | 144 | | | NRF1_MA0506.1 | 8.529800e-02 | |
| pos_patterns.pattern_17 | 135 | | | TYY1_HUMAN.H11MO.0.A | 7.331310e-03 | |
| pos_patterns.pattern_18 | 78 | | | STAT1_HUMAN.H11MO.0.A | 1.615170e-02 | |
| pos_patterns.pattern_19 | 75 | | | ETV6_ETS_1 | 5.495850e-05 | |
| pos_patterns.pattern_20 | 40 | | | ZNF76_HUMAN.H11MO.0.C | 4.157780e-13 | |
| pos_patterns.pattern_21 | 38 | | | SRY_MA0084.1 | 1.000000e+00 | |
| pos_patterns.pattern_22 | 37 | | | FLI1_ETS_3 | 2.185180e-01 | |
| pos_patterns.pattern_23 | 36 | | | KLF5_MA0599.1 | 5.070640e-03 | |
| pos_patterns.pattern_24 | 27 | | | NaN | NaN | |
| pos_patterns.pattern_25 | 21 | | | NaN | NaN | |
| neg_patterns.pattern_0 | 1188 | | | CTCF_MA0139.1 | 1.172910e-08 | |
| neg_patterns.pattern_1 | 171 | | | HIC2_C2H2_1 | 9.044890e-01 | |
| neg_patterns.pattern_2 | 170 | | | REST_MA0138.2 | 2.159730e-11 | |
| neg_patterns.pattern_3 | 125 | | | NFYA_MA0060.3 | 1.558440e-02 | |

| pattern | num_seqlets | cwm_fwd | cwm_rev | TOMTOM_match | TOMTOM_qval | TOMTOM_match_logo |
|---|---|---|---|---|---|---|
| neg_patterns.pattern_4 | 60 | 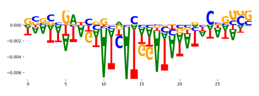 | 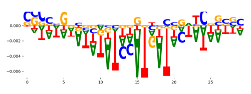 | Z354A_HUMAN.H11MO.0.C | 1.083500e-01 | 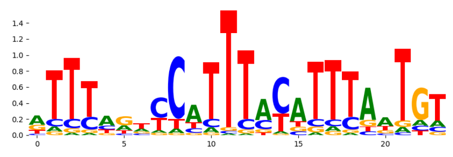 |
| neg_patterns.pattern_5 | 57 | 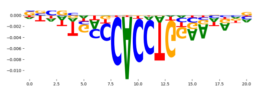 | 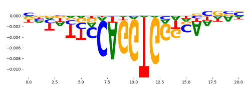 | SNAI1_HUMAN.H11MO.0.C | 1.380430e-03 | 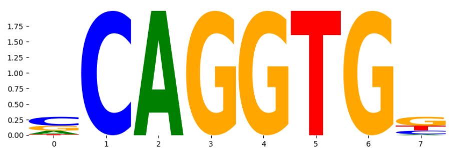 |
| neg_patterns.pattern_6 | 36 | 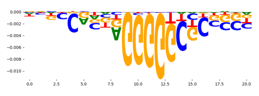 | 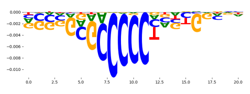 | CTCFL_MOUSE.H11MO.0.A | 4.582550e-05 | 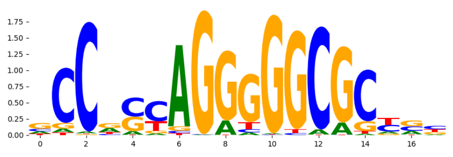 |
| neg_patterns.pattern_7 | 31 | 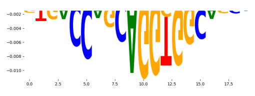 | 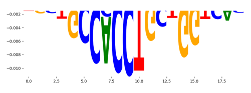 | CTCFL_HUMAN.H11MO.0.A | 7.590780e-04 | 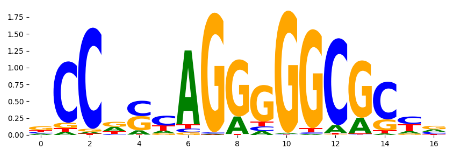 |
| neg_patterns.pattern_8 | 29 | 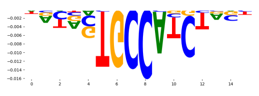 | 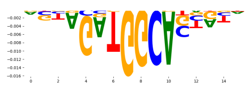 | NEUROD1_MA1109.1 | 5.070610e-01 | 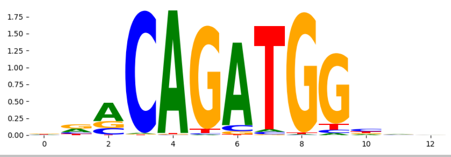 |
