## Supplementary Files 1 for "ChromBPNet: bias factorized, base-resolution deep learning models of chromatin accessibility reveal cis-regulatory sequence syntax, transcription factor footprints and regulatory variants": fig1b_k562_ATAC_raw_bpnet_uncorrected_profile_modisco.pdf

| pattern | num_seqlets | cwm_fwd | cwm_rev | TOMTOM_match | TOMTOM_qval | TOMTOM_match_logo |
|---|---|---|---|---|---|---|
| pos_patterns.pattern_0 | 6900 | | | TN5_2 | 6.560760e-08 | |
| pos_patterns.pattern_1 | 5483 | | | TN5_2 | 2.815270e-04 | |
| pos_patterns.pattern_2 | 4514 | | | CTCF_MA0139.1 | 5.112140e-16 | |
| pos_patterns.pattern_3 | 845 | | | Bach1+Mafk_MA0591.1 | 2.250170e-06 | |
| pos_patterns.pattern_4 | 473 | | | TN5_3 | 2.936680e-08 | |
| pos_patterns.pattern_5 | 382 | | | TN5_6 | 1.881390e-20 | |
| pos_patterns.pattern_6 | 380 | | | SP2_HUMAN.H11MO.0.A | 2.601250e-08 | |
| pos_patterns.pattern_7 | 375 | | | SP5_MOUSE.H11MO.0.C | 2.782670e-03 | |
| pos_patterns.pattern_8 | 286 | | | SP2_HUMAN.H11MO.0.A | 2.561090e-06 | |
| pos_patterns.pattern_9 | 285 | | | NFYC_HUMAN.H11MO.0.A | 1.534090e-04 | |
| pos_patterns.pattern_10 | 243 | | | KLF5_MA0599.1 | 1.671040e-05 | |
| pos_patterns.pattern_11 | 194 | | | TAL1_HUMAN.H11MO.0.A | 4.687400e-08 | |
| pos_patterns.pattern_12 | 131 | | | KLF5_MA0599.1 | 1.147150e-02 | |
| pos_patterns.pattern_13 | 119 | | | KLF9_MA1107.1 | 6.118750e-04 | |
| pos_patterns.pattern_14 | 114 | | | NFYA_MA0060.3 | 1.603150e-07 | |
| pos_patterns.pattern_15 | 90 | | | ZN143_MOUSE.H11MO.0.A | 1.170090e-10 | |
| pos_patterns.pattern_16 | 74 | | | ATF4_MOUSE.H11MO.0.A | 1.631440e-06 | |
| pos_patterns.pattern_17 | 69 | | | REST_HUMAN.H11MO.0.A | 4.836510e-12 | |
| pos_patterns.pattern_18 | 61 | | | GATA3_GATA_2 | 2.123370e-03 | |
| pos_patterns.pattern_19 | 31 | | | KLF4_HUMAN.H11MO.0.A | 5.955100e-04 | |
| pos_patterns.pattern_20 | 30 | | | TN5_1 | 3.216320e-01 | |
| pos_patterns.pattern_21 | 30 | | | NRF1_MA0506.1 | 4.268540e-04 | |
| pos_patterns.pattern_22 | 30 | | | Gata1_MA0035.3 | 2.141770e-02 | |
