## Supplementary Files 1 for "ChromBPNet: bias factorized, base-resolution deep learning models of chromatin accessibility reveal cis-regulatory sequence syntax, transcription factor footprints and regulatory variants": fig1c_gm12878_ATAC_raw_bpnet_uncorrected_counts_modisco.pdf

| pattern | num_seqlets | cwm_fwd | cwm_rev | TOMTOM_match | TOMTOM_qval | TOMTOM_match_logo |
|---|---|---|---|---|---|---|
| pos_patterns.pattern_0 | 3928 | | | IRF1_MOUSE.H11MO.0.A | 3.923590e-03 | |
| pos_patterns.pattern_1 | 3289 | | | ELF5_HUMAN.H11MO.0.A | 7.641830e-04 | |
| pos_patterns.pattern_2 | 2562 | | | NFKB1_HUMAN.H11MO.1.B | 8.201100e-05 | |
| pos_patterns.pattern_3 | 2267 | | | FOS_HUMAN.H11MO.0.A | 2.146600e-03 | |
| pos_patterns.pattern_4 | 2039 | | | RUNX1_HUMAN.H11MO.0.A | 1.367720e-03 | |
| pos_patterns.pattern_5 | 1466 | | | ELK1_ETS_1 | 9.781250e-02 | |
| pos_patterns.pattern_6 | 1071 | | | KLF12_HUMAN.H11MO.0.C | 1.145100e-04 | |
| pos_patterns.pattern_7 | 1029 | | | CTCF_HUMAN.H11MO.0.A | 6.543820e-13 | |
| pos_patterns.pattern_8 | 563 | | | NFYA_MA0060.3 | 4.821560e-01 | |
| pos_patterns.pattern_9 | 554 | | | Atf3_MA0605.1 | 2.034030e-02 | |
| pos_patterns.pattern_10 | 462 | | | NRF1_HUMAN.H11MO.0.A | 1.946740e-07 | |
| pos_patterns.pattern_11 | 364 | | | Pou2f2.mouse_POU_2 | 7.033640e-03 | |
| pos_patterns.pattern_12 | 346 | | | EBF1_EBF_1 | 2.177400e-05 | |
| pos_patterns.pattern_13 | 200 | | | MEF2D_MOUSE.H11MO.0.A | 4.565780e-07 | |
| pos_patterns.pattern_14 | 185 | | | PAX5_MOUSE.H11MO.0.A | 2.481870e-14 | |
| pos_patterns.pattern_15 | 168 | | | HNF1B_HUMAN.H11MO.0.A | 3.538430e-06 | |
| pos_patterns.pattern_16 | 160 | | | SPI1_ETS_1 | 1.479060e-02 | |
| pos_patterns.pattern_17 | 151 | | | JUND_MA0492.1 | 3.078890e-03 | |
| pos_patterns.pattern_18 | 136 | | | KAISO_HUMAN.H11MO.0.A | 2.645110e-03 | |
| pos_patterns.pattern_19 | 132 | | | TYY1_HUMAN.H11MO.0.A | 5.620560e-06 | |
| pos_patterns.pattern_20 | 130 | | | PAX1_MA0779.1 | 2.194780e-06 | |
| pos_patterns.pattern_21 | 125 | | | ZN143_HUMAN.H11MO.0.A | 5.216290e-10 | |
| pos_patterns.pattern_22 | 50 | | | FOSL1_MA0477.1 | 4.760960e-03 | |
| pos_patterns.pattern_23 | 39 | | | RFX2_HUMAN.H11MO.0.A | 2.563000e-09 | |
| pos_patterns.pattern_24 | 31 | | | ZNF76_HUMAN.H11MO.0.C | 1.010970e-10 | |
| pos_patterns.pattern_25 | 26 | | | Rfx3.mouse_RFX_1 | 3.691570e-02 | |
| pos_patterns.pattern_26 | 22 | | | IRF7_HUMAN.H11MO.0.C | 2.195690e-03 | |
| pos_patterns.pattern_27 | 20 | | | ZNF76_HUMAN.H11MO.0.C | 3.744990e-09 | |
| neg_patterns.pattern_0 | 116 | | | NFATC2_MA0152.1 | 7.336560e-01 | |

| pattern | num_seqlets | cwm_fwd | cwm_rev | TOMTOM_match | TOMTOM_qval | TOMTOM_match_logo |
|---|---|---|---|---|---|---|
| neg_patterns.pattern_1 | 69 | 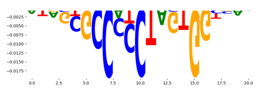 | 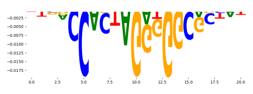 | CTCF_MA0139.1 | 4.306850e-07 | 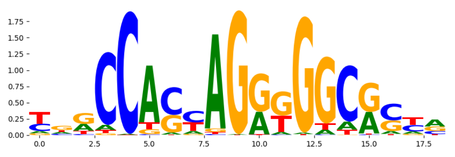 |
| neg_patterns.pattern_2 | 46 | 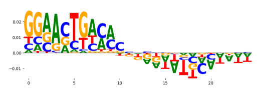 | 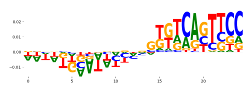 | STAT1+STAT2_MA0517.1 | 6.559600e-03 | 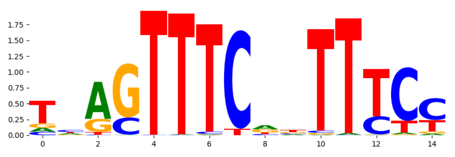 |
| neg_patterns.pattern_3 | 29 | 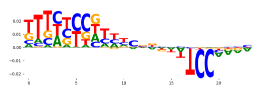 | 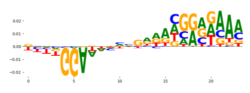 | NFATC3_MA0625.1 | 1.162920e-01 | 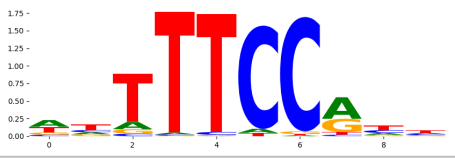 |
