## Supplementary Files 1 for "ChromBPNet: bias factorized, base-resolution deep learning models of chromatin accessibility reveal cis-regulatory sequence syntax, transcription factor footprints and regulatory variants": fig1d_gm12878_ATAC_raw_bpnet_uncorrected_profile_modisco.pdf

| pattern | num_seqlets | cwm_fwd | cwm_rev | TOMTOM_match | TOMTOM_qval | TOMTOM_match_logo |
|---|---|---|---|---|---|---|
| pos_patterns.pattern_0 | 7085 | | | TN5_1 | 2.170570e-09 | |
| pos_patterns.pattern_1 | 4939 | | | TN5_2 | 1.456940e-02 | |
| pos_patterns.pattern_2 | 3628 | | | CTCF_MA0139.1 | 3.028070e-14 | |
| pos_patterns.pattern_3 | 1634 | | | TN5_3 | 1.245300e-12 | |
| pos_patterns.pattern_4 | 897 | | | KLF4_MA0039.3 | 1.000000e+00 | |
| pos_patterns.pattern_5 | 856 | | | BATF+JUN_MA0462.1 | 1.740240e-06 | |
| pos_patterns.pattern_6 | 673 | | | IRF1_MOUSE.H11MO.0.A | 1.321160e-05 | |
| pos_patterns.pattern_7 | 651 | | | TN5_4 | 3.105130e-04 | |
| pos_patterns.pattern_8 | 632 | | | TN5_4 | 8.225450e-04 | |
| pos_patterns.pattern_9 | 577 | | | SPIB_MOUSE.H11MO.0.A | 1.448930e-04 | |
| pos_patterns.pattern_10 | 427 | | | IRF4_HUMAN.H11MO.0.A | 7.396960e-07 | |
| pos_patterns.pattern_11 | 417 | | | TN5_4 | 1.145880e-01 | |
| pos_patterns.pattern_12 | 385 | | | TN5_6 | 2.160850e-18 | |
| pos_patterns.pattern_13 | 195 | | | NFKB1_HUMAN.H11MO.1.B | 9.020920e-06 | |
| pos_patterns.pattern_14 | 95 | | | NFKB2_HUMAN.H11MO.0.B | 2.098310e-04 | |
| pos_patterns.pattern_15 | 90 | | | RUNX2_MA0511.2 | 1.000000e+00 | |
| pos_patterns.pattern_16 | 88 | | | TN5_7 | 1.235670e-06 | |
| pos_patterns.pattern_17 | 67 | | | TN5_6 | 5.583200e-06 | |
| pos_patterns.pattern_18 | 46 | | | NFKB1_HUMAN.H11MO.1.B | 3.280860e-04 | |
| pos_patterns.pattern_19 | 24 | | | NFKB1_MA0105.4 | 2.501510e-04 | |
| pos_patterns.pattern_20 | 20 | | | STAT1+STAT2_MA0517.1 | 1.265250e-01 | |
