## Supplementary Files 1 for "ChromBPNet: bias factorized, base-resolution deep learning models of chromatin accessibility reveal cis-regulatory sequence syntax, transcription factor footprints and regulatory variants": fig1e_gm12878_ATAC_hint_corrected_bpnet_counts_modisco.pdf

| pattern | num_seqlets | cwm_fwd | cwm_rev | TOMTOM_match | TOMTOM_qval | TOMTOM_match_logo |
|---|---|---|---|---|---|---|
| pos_patterns.pattern_0 | 3860 | | | IRF1_MOUSE.H11MO.0.A | 4.563880e-03 | |
| pos_patterns.pattern_1 | 3463 | | | SPIB_MOUSE.H11MO.0.A | 2.153090e-06 | |
| pos_patterns.pattern_2 | 2344 | | | FOSL2+JUN_MA1130.1 | 1.388970e-03 | |
| pos_patterns.pattern_3 | 1831 | | | RUNX3_HUMAN.H11MO.0.A | 2.386630e-03 | |
| pos_patterns.pattern_4 | 1530 | | | Elk3.mouse_ETS_1 | 1.035640e-04 | |
| pos_patterns.pattern_5 | 1515 | | | TF65_MOUSE.H11MO.0.A | 1.629520e-05 | |
| pos_patterns.pattern_6 | 1381 | | | CTCF_MA0139.1 | 3.875760e-16 | |
| pos_patterns.pattern_7 | 1336 | | | KLF12_HUMAN.H11MO.0.C | 5.353910e-04 | |
| pos_patterns.pattern_8 | 660 | | | NFKB2_HUMAN.H11MO.0.B | 5.708390e-04 | |
| pos_patterns.pattern_9 | 523 | | | ATF1_HUMAN.H11MO.0.B | 1.046210e-04 | |
| pos_patterns.pattern_10 | 519 | | | NFYB_HUMAN.H11MO.0.A | 1.056160e-02 | |
| pos_patterns.pattern_11 | 484 | | | BATF3_HUMAN.H11MO.0.B | 2.836750e-08 | |
| pos_patterns.pattern_12 | 430 | | | NRF1_MOUSE.H11MO.0.A | 1.867060e-06 | |
| pos_patterns.pattern_13 | 353 | | | POU5F1_MA1115.1 | 3.183180e-03 | |
| pos_patterns.pattern_14 | 307 | | | EBF1_EBF_1 | 1.949320e-05 | |
| pos_patterns.pattern_15 | 161 | | | SPI1_ETS_1 | 2.228180e-02 | |
| pos_patterns.pattern_16 | 148 | | | TYY1_HUMAN.H11MO.0.A | 1.164150e-07 | |
| pos_patterns.pattern_17 | 144 | | | ZBTB33_MA0527.1 | 5.212800e-02 | |
| pos_patterns.pattern_18 | 120 | | | HNF1B_MOUSE.H11MO.0.A | 8.984210e-06 | |
| pos_patterns.pattern_19 | 110 | | | ZNF76_HUMAN.H11MO.0.C | 1.296790e-20 | |
| pos_patterns.pattern_20 | 55 | | | MEF2D_HUMAN.H11MO.0.A | 1.008160e-03 | |
| pos_patterns.pattern_21 | 50 | | | FOXJ3_HUMAN.H11MO.0.A | 2.261780e-01 | |
| pos_patterns.pattern_22 | 45 | | | THA11_MOUSE.H11MO.0.B | 5.394040e-11 | |
| pos_patterns.pattern_23 | 39 | | | PAX2_PAX_1 | 9.979950e-07 | |
| pos_patterns.pattern_24 | 31 | | | MEF2B_MA0660.1 | 1.470570e-04 | |
| pos_patterns.pattern_25 | 30 | | | JUN_MA0489.1 | 3.706570e-03 | |
| neg_patterns.pattern_0 | 127 | | | TEAD1_MOUSE.H11MO.0.A | 5.340050e-01 | |
