## Supplementary Files 1 for "ChromBPNet: bias factorized, base-resolution deep learning models of chromatin accessibility reveal cis-regulatory sequence syntax, transcription factor footprints and regulatory variants": fig1f_gm12878_ATAC_hint_corrected_bpnet_profile_modisco.pdf

| pattern | num_seqlets | cwm_fwd | cwm_rev | TOMTOM_match | TOMTOM_qval | TOMTOM_match_logo |
|---|---|---|---|---|---|---|
| pos_patterns.pattern_0 | 9994 | 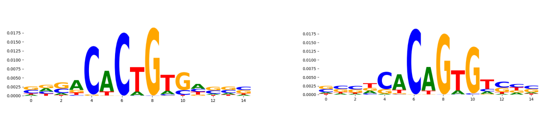 | | NFIB_NFI_1 | 1.000000e+00 | 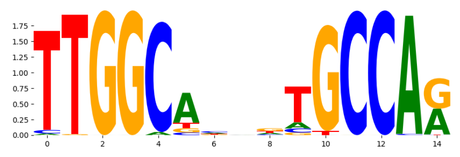 |
| pos_patterns.pattern_1 | 3447 | 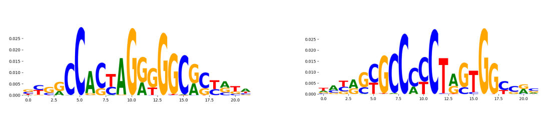 | | CTCF_MA0139.1 | 7.712760e-15 | 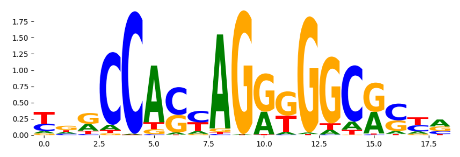 |
| pos_patterns.pattern_2 | 2589 | 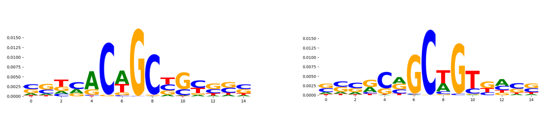 | | MSC_MA0665.1 | 2.364090e-02 | 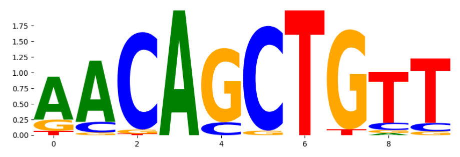 |
| pos_patterns.pattern_3 | 1666 | 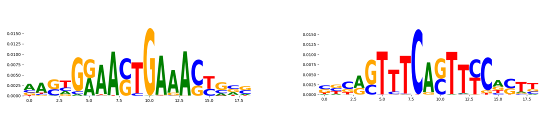 | | IRF1_MOUSE.H11MO.0.A | 6.778660e-05 | 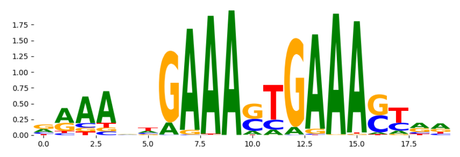 |
| pos_patterns.pattern_4 | 942 | 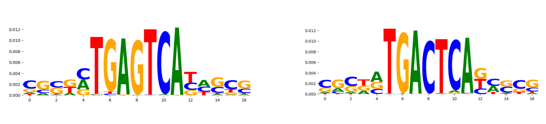 | | FOSL1+JUN_MA1128.1 | 3.929150e-04 | 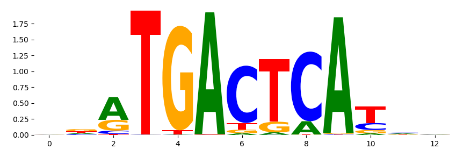 |
| pos_patterns.pattern_5 | 841 | 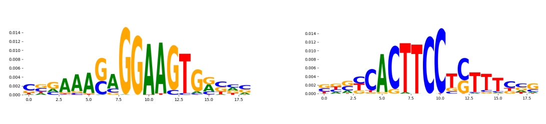 | | SPIB_MOUSE.H11MO.0.A | 5.683400e-05 | 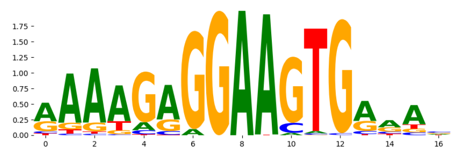 |
| pos_patterns.pattern_6 | 713 | 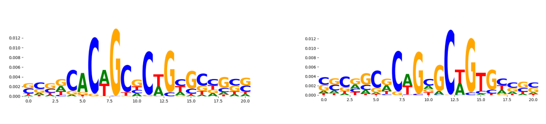 | | TBX1_TBX_4 | 7.178580e-01 | 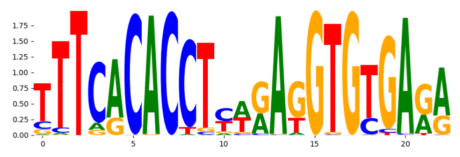 |
| pos_patterns.pattern_7 | 581 | 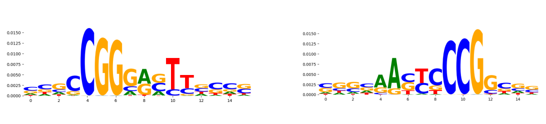 | | TF65_MOUSE.H11MO.0.A | 7.645550e-01 | 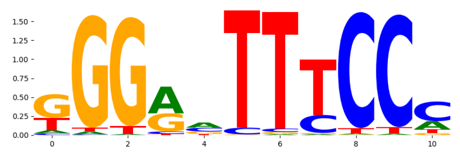 |
| pos_patterns.pattern_8 | 555 | 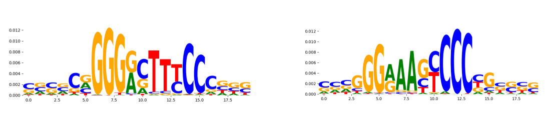 | | TF65_HUMAN.H11MO.0.A | 8.829450e-05 | 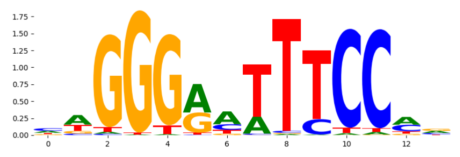 |
| pos_patterns.pattern_9 | 405 | 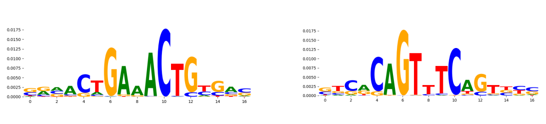 | | STAT1_MOUSE.H11MO.0.A | 3.632700e-01 | 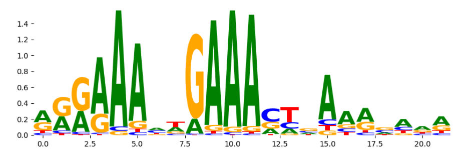 |
| pos_patterns.pattern_10 | 343 | 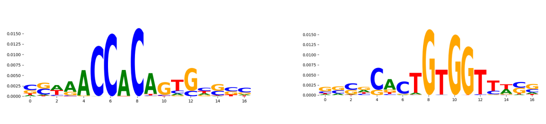 | | GLI2_C2H2_2 | 5.547620e-01 | 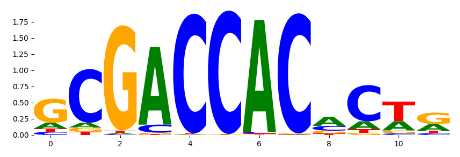 |
| pos_patterns.pattern_11 | 219 | 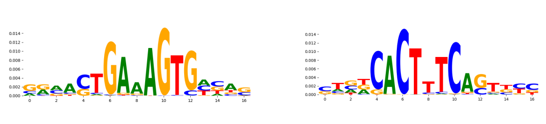 | | IRF1_MA0050.2 | 2.263410e-02 | 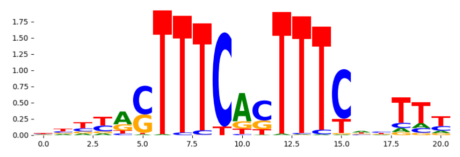 |
| pos_patterns.pattern_12 | 197 | 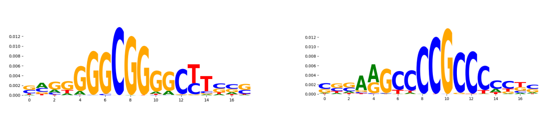 | | KLF3_HUMAN.H11MO.0.B | 2.732320e-04 | 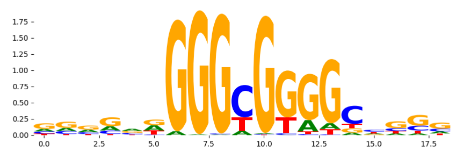 |
| pos_patterns.pattern_13 | 157 | 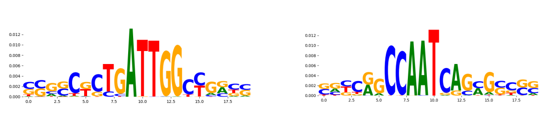 | | NFYB_HUMAN.H11MO.0.A | 1.703180e-09 | 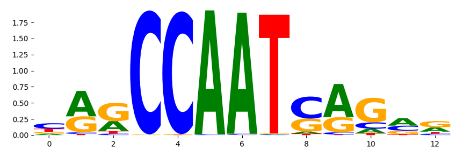 |
| pos_patterns.pattern_14 | 152 | 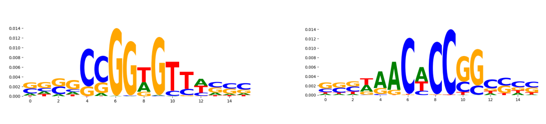 | | MESP1_bHLH_1 | 6.930440e-01 | 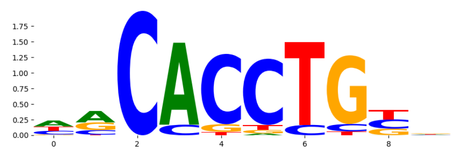 |
| pos_patterns.pattern_15 | 68 | 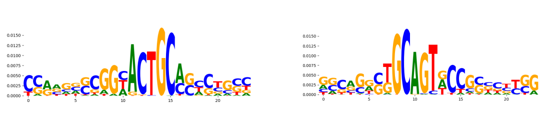 | | GABPA_HUMAN.H11MO.0.A | 1.668740e-01 | 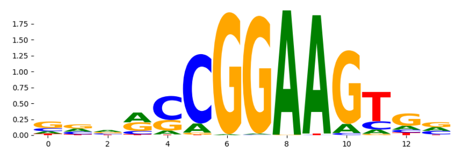 |
| pos_patterns.pattern_16 | 64 | 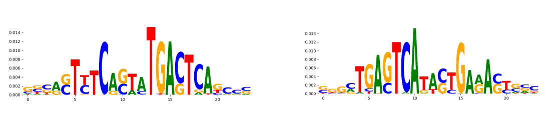 | | BATF_HUMAN.H11MO.0.A | 5.450350e-05 | 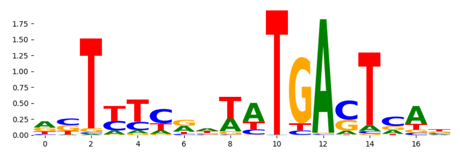 |
| pos_patterns.pattern_17 | 56 | 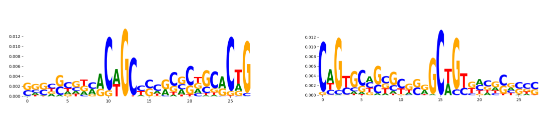 | | TN5_1 | 1.686230e-02 | 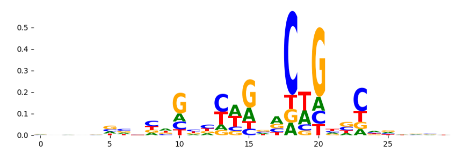 |
| pos_patterns.pattern_18 | 55 | 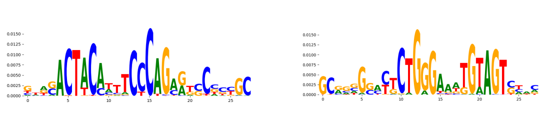 | | ZNF76_HUMAN.H11MO.0.C | 4.258900e-08 | 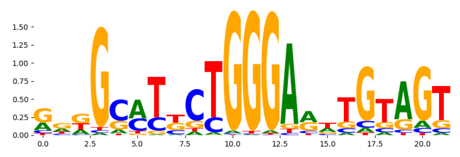 |
| pos_patterns.pattern_19 | 27 | 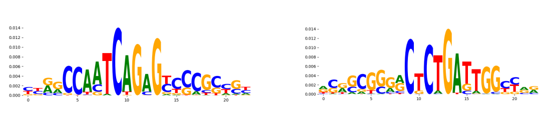 | | FOXI1_HUMAN.H11MO.0.B | 6.648740e-02 | 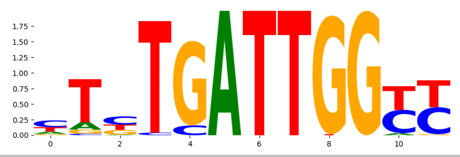 |
