## Supplementary Files 1 for "ChromBPNet: bias factorized, base-resolution deep learning models of chromatin accessibility reveal cis-regulatory sequence syntax, transcription factor footprints and regulatory variants": fig1g_gm12878_ATAC_raw_bpnet_w_tobias_bias_counts_modisco.pdf

| pattern | num_seqlets | cwm_fwd | cwm_rev | TOMTOM_match | TOMTOM_qval | TOMTOM_match_logo |
|---|---|---|---|---|---|---|
| pos_patterns.pattern_0 | 4566 | | | IRF1_MOUSE.H11MO.0.A | 1.760010e-04 | |
| pos_patterns.pattern_1 | 4137 | | | CTCF_MA0139.1 | 1.931650e-15 | |
| pos_patterns.pattern_2 | 3065 | | | ELF5_HUMAN.H11MO.0.A | 2.072470e-07 | |
| pos_patterns.pattern_3 | 2206 | | | RUNX3_HUMAN.H11MO.0.A | 3.068200e-03 | |
| pos_patterns.pattern_4 | 2175 | | | NFKB1_HUMAN.H11MO.1.B | 5.736500e-04 | |
| pos_patterns.pattern_5 | 1832 | | | ATF3_MOUSE.H11MO.0.A | 3.824140e-03 | |
| pos_patterns.pattern_6 | 726 | | | BC11A_HUMAN.H11MO.0.A | 2.651010e-07 | |
| pos_patterns.pattern_7 | 697 | | | KLF12_HUMAN.H11MO.0.C | 2.164240e-04 | |
| pos_patterns.pattern_8 | 626 | | | BATF_MOUSE.H11MO.0.A | 8.122900e-03 | |
| pos_patterns.pattern_9 | 305 | | | NRF1_HUMAN.H11MO.0.A | 1.397670e-06 | |
| pos_patterns.pattern_10 | 301 | | | NFKB2_HUMAN.H11MO.0.B | 8.742870e-02 | |
| pos_patterns.pattern_11 | 299 | | | BATF_HUMAN.H11MO.0.A | 4.902450e-06 | |
| pos_patterns.pattern_12 | 287 | | | IRF4_MOUSE.H11MO.0.A | 1.275080e-02 | |
| pos_patterns.pattern_13 | 281 | | | CREB1_HUMAN.H11MO.0.A | 6.425180e-07 | |
| pos_patterns.pattern_14 | 264 | | | NFYB_HUMAN.H11MO.0.A | 1.239030e-03 | |
| pos_patterns.pattern_15 | 197 | | | IRF9_IRF_1 | 1.062330e-03 | |
| pos_patterns.pattern_16 | 157 | | | Pou2f2.mouse_POU_2 | 1.603020e-04 | |
| pos_patterns.pattern_17 | 151 | | | ZNF76_HUMAN.H11MO.0.C | 9.513880e-18 | |
| pos_patterns.pattern_18 | 96 | | | PAX5_MOUSE.H11MO.0.A | 8.247180e-08 | |
| pos_patterns.pattern_19 | 72 | | | HNF1A_HUMAN.H11MO.0.C | 7.857330e-06 | |
| pos_patterns.pattern_20 | 40 | | | PAX2_PAX_1 | 5.599190e-04 | |
| pos_patterns.pattern_21 | 35 | | | ZBTB33_MA0527.1 | 3.479680e-04 | |
| pos_patterns.pattern_22 | 29 | | | Rfx1_MA0509.1 | 9.618820e-09 | |
| neg_patterns.pattern_0 | 99 | | | TF65_HUMAN.H11MO.0.A | 5.033530e-01 | |
| neg_patterns.pattern_1 | 53 | | | ZN770_HUMAN.H11MO.0.C | 1.642020e-01 | |
| neg_patterns.pattern_2 | 48 | | | FOXB1_forkhead_2 | 5.103930e-01 | |
| neg_patterns.pattern_3 | 34 | | | ZNF384_MA1125.1 | 4.184840e-02 | |
| neg_patterns.pattern_4 | 20 | | | Alx4.mouse_homeodomain_1 | 2.879690e-01 | |

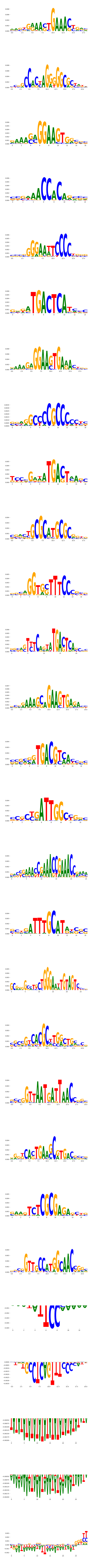

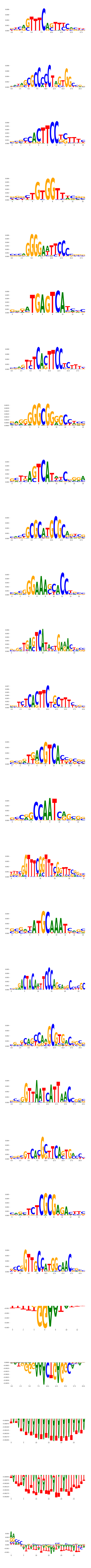

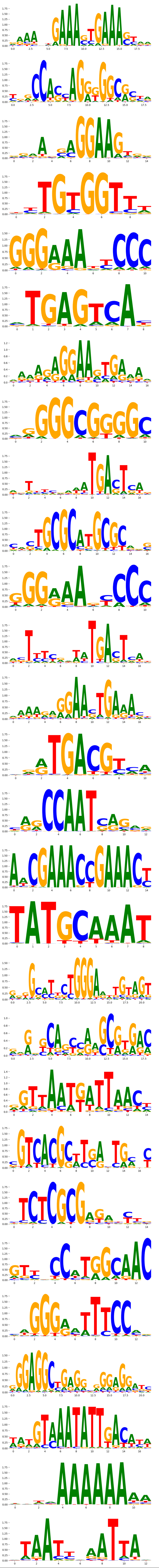
