## Supplementary Files 1 for "ChromBPNet: bias factorized, base-resolution deep learning models of chromatin accessibility reveal cis-regulatory sequence syntax, transcription factor footprints and regulatory variants": fig1h_gm12878_ATAC_raw_bpnet_w_tobias_bias_profile_modisco.pdf

| pattern | num_seqlets | cwm_fwd | cwm_rev | TOMTOM_match | TOMTOM_qval | TOMTOM_match_logo |
|---|---|---|---|---|---|---|
| pos_patterns.pattern_0 | 4348 | | | CTCF_MA0139.1 | 4.109980e-14 | |
| pos_patterns.pattern_1 | 3584 | | | TN5_8 | 2.094890e-02 | |
| pos_patterns.pattern_2 | 1461 | | | FOSL1+JUN_MA1128.1 | 4.101000e-04 | |
| pos_patterns.pattern_3 | 1240 | | | TN5_8 | 1.000000e+00 | |
| pos_patterns.pattern_4 | 1213 | | | TN5_7 | 1.774580e-04 | |
| pos_patterns.pattern_5 | 1115 | | | TN5_3 | 8.297080e-02 | |
| pos_patterns.pattern_6 | 928 | | | SPIB_MOUSE.H11MO.0.A | 8.512980e-07 | |
| pos_patterns.pattern_7 | 699 | | | IRF4_HUMAN.H11MO.0.A | 1.934270e-08 | |
| pos_patterns.pattern_8 | 689 | | | RUNX2_HUMAN.H11MO.0.A | 1.112250e-01 | |
| pos_patterns.pattern_9 | 672 | | | RELB_HUMAN.H11MO.0.C | 8.182670e-08 | |
| pos_patterns.pattern_10 | 625 | | | TN5_3 | 1.388560e-01 | |
| pos_patterns.pattern_11 | 616 | | | IRF1_HUMAN.H11MO.0.A | 1.789220e-09 | |
| pos_patterns.pattern_12 | 523 | | | TN5_8 | 4.741650e-02 | |
| pos_patterns.pattern_13 | 508 | | | TBX21_MOUSE.H11MO.0.A | 1.181510e-02 | |
| pos_patterns.pattern_14 | 492 | | | Klf1_MA0493.1 | 6.737760e-01 | |
| pos_patterns.pattern_15 | 482 | | | MAZ_HUMAN.H11MO.0.A | 6.974330e-03 | |
| pos_patterns.pattern_16 | 465 | | | IRF1_MOUSE.H11MO.0.A | 1.186230e-03 | |
| pos_patterns.pattern_17 | 447 | | | TN5_6 | 7.206550e-04 | |
| pos_patterns.pattern_18 | 290 | | | TN5_7 | 4.677140e-02 | |
| pos_patterns.pattern_19 | 284 | | | MEF2B_MA0660.1 | 1.000000e+00 | |
| pos_patterns.pattern_20 | 238 | | | Zfp740.mouse_C2H2_1 | 6.316440e-02 | |
| pos_patterns.pattern_21 | 210 | | | NFYB_HUMAN.H11MO.0.A | 2.563190e-07 | |
| pos_patterns.pattern_22 | 113 | | | ZN490_HUMAN.H11MO.0.C | 1.000000e+00 | |
| pos_patterns.pattern_23 | 45 | | | RUNX1_HUMAN.H11MO.0.A | 2.950580e-01 | |
| pos_patterns.pattern_24 | 37 | | | RUNX2_RUNX_2 | 4.670560e-01 | |
| pos_patterns.pattern_25 | 36 | | | ZN143_MOUSE.H11MO.0.A | 7.631770e-09 | |
| pos_patterns.pattern_26 | 31 | | | MEF2D_MA0773.1 | 1.000000e+00 | |
| pos_patterns.pattern_27 | 23 | | | RUNX3_MOUSE.H11MO.0.A | 3.420230e-01 | |
| pos_patterns.pattern_28 | 21 | | | SP1_MA0079.3 | 1.409690e-03 | |
