## Supplementary Files 1 for "ChromBPNet: bias factorized, base-resolution deep learning models of chromatin accessibility reveal cis-regulatory sequence syntax, transcription factor footprints and regulatory variants": fig1i_gm12878_ATAC_tobias_corrected_softmax_bpnet_counts_modisco.pdf

| pattern | num_seqlets | cwm_fwd | cwm_rev | TOMTOM_match | TOMTOM_qval | TOMTOM_match_logo |
|---|---|---|---|---|---|---|
| pos_patterns.pattern_0 | 3553 | | | ELK1_ETS_1 | 1.866050e-01 | |
| pos_patterns.pattern_1 | 2804 | | | SP1_HUMAN.H11MO.0.A | 1.768760e-02 | |
| pos_patterns.pattern_2 | 2235 | | | NRF1_MOUSE.H11MO.0.A | 3.845280e-03 | |
| pos_patterns.pattern_3 | 1628 | | | PAX9_MA0781.1 | 3.302860e-01 | |
| pos_patterns.pattern_4 | 1495 | | | NFYB_HUMAN.H11MO.0.A | 4.304610e-05 | |
| pos_patterns.pattern_5 | 1100 | | | MYC_MOUSE.H11MO.0.A | 1.000000e+00 | |
| pos_patterns.pattern_6 | 1033 | | | SPI1_MOUSE.H11MO.0.A | 9.429020e-04 | |
| pos_patterns.pattern_7 | 1015 | | | STAT1_MOUSE.H11MO.0.A | 4.208770e-04 | |
| pos_patterns.pattern_8 | 625 | | | NRF1_MA0506.1 | 1.000000e+00 | |
| pos_patterns.pattern_9 | 625 | | | CTCF_MOUSE.H11MO.0.A | 1.717520e-09 | |
| pos_patterns.pattern_10 | 565 | | | MYCN_MA0104.4 | 1.737630e-01 | |
| pos_patterns.pattern_11 | 532 | | | FOSL2+JUN_MA1130.1 | 1.565650e-04 | |
| pos_patterns.pattern_12 | 482 | | | MYC_MOUSE.H11MO.0.A | 4.834650e-01 | |
| pos_patterns.pattern_13 | 481 | | | SPIB_MOUSE.H11MO.0.A | 1.595660e-05 | |
| pos_patterns.pattern_14 | 460 | | | ELK1_ETS_3 | 1.182600e-01 | |
| pos_patterns.pattern_15 | 449 | | | KLF12_HUMAN.H11MO.0.C | 3.898350e-01 | |
| pos_patterns.pattern_16 | 407 | | | NRF1_MA0506.1 | 3.097940e-01 | |
| pos_patterns.pattern_17 | 394 | | | TAF1_HUMAN.H11MO.0.A | 1.157600e-01 | |
| pos_patterns.pattern_18 | 353 | | | NRF1_MOUSE.H11MO.0.A | 2.475430e-01 | |
| pos_patterns.pattern_19 | 329 | | | USF2_HUMAN.H11MO.0.A | 1.649940e-01 | |
| pos_patterns.pattern_20 | 293 | | | TAF1_HUMAN.H11MO.0.A | 3.194060e-01 | |
| pos_patterns.pattern_21 | 239 | | | AP2B_HUMAN.H11MO.0.B | 4.523980e-01 | |
| pos_patterns.pattern_22 | 220 | | | RFX1_HUMAN.H11MO.0.B | 1.000000e+00 | |
| pos_patterns.pattern_23 | 174 | | | CTCF_MA0139.1 | 3.856570e-07 | |
| pos_patterns.pattern_24 | 152 | | | SP2_HUMAN.H11MO.0.A | 1.215810e-05 | |
| pos_patterns.pattern_25 | 150 | | | Gabpa_MA0062.2 | 1.199510e-03 | |
| pos_patterns.pattern_26 | 135 | | | ASCL1_MA1100.1 | 1.888670e-01 | |
| pos_patterns.pattern_27 | 131 | | | Gabpa_MA0062.2 | 4.930940e-02 | |
| pos_patterns.pattern_28 | 97 | | | THAP1_HUMAN.H11MO.0.C | 4.050360e-01 | |

| pattern | num_seqlets | cwm_fwd | cwm_rev | TOMTOM_match | TOMTOM_qval | TOMTOM_match_logo |
|---|---|---|---|---|---|---|
| pos_patterns.pattern_29 | 88 | | | THA11_HUMAN.H11MO.0.B | 3.665950e-12 | |
| pos_patterns.pattern_30 | 51 | | | THAP1_HUMAN.H11MO.0.C | 9.952360e-02 | |
| pos_patterns.pattern_31 | 39 | | | THAP1_HUMAN.H11MO.0.C | 9.337530e-02 | |
| neg_patterns.pattern_0 | 843 | | | POU3F2_POU_1 | 4.098810e-01 | |
| neg_patterns.pattern_1 | 529 | | | ZNF384_MA1125.1 | 4.397650e-02 | |
| neg_patterns.pattern_2 | 227 | | | SP2_HUMAN.H11MO.0.A | 2.033890e-07 | |
| neg_patterns.pattern_3 | 195 | | | SP2_HUMAN.H11MO.0.A | 1.983480e-06 | |
| neg_patterns.pattern_4 | 121 | | | FOXD2_forkhead_1 | 2.697460e-01 | |
| neg_patterns.pattern_5 | 110 | | | ZNF384_MA1125.1 | 3.631520e-02 | |
| neg_patterns.pattern_6 | 109 | | | SP1_MOUSE.H11MO.0.A | 1.712820e-05 | |
| neg_patterns.pattern_7 | 101 | | | TEAD4_MA0809.1 | 3.824400e-01 | |
| neg_patterns.pattern_8 | 94 | | | SP2_HUMAN.H11MO.0.A | 7.532500e-06 | |
| neg_patterns.pattern_9 | 92 | | | EGR1_MOUSE.H11MO.0.A | 6.416160e-01 | |
| neg_patterns.pattern_10 | 90 | | | SP2_HUMAN.H11MO.0.A | 1.104700e-02 | |
| neg_patterns.pattern_11 | 89 | | | SP3_HUMAN.H11MO.0.B | 2.046470e-04 | |
| neg_patterns.pattern_12 | 81 | | | RREB1_MA0073.1 | 8.505600e-01 | |
| neg_patterns.pattern_13 | 68 | | | ZFX_MOUSE.H11MO.0.B | 7.971570e-03 | |
| neg_patterns.pattern_14 | 66 | | | SP2_MA0516.1 | 1.765490e-03 | |
| neg_patterns.pattern_15 | 66 | | | SP2_MA0516.1 | 6.206960e-03 | |
| neg_patterns.pattern_16 | 64 | | | SP2_HUMAN.H11MO.0.A | 6.791570e-02 | |
| neg_patterns.pattern_17 | 60 | | | SP1_HUMAN.H11MO.0.A | 1.619120e-03 | |
| neg_patterns.pattern_18 | 57 | | | NRF1_MOUSE.H11MO.0.A | 4.263370e-01 | |
| neg_patterns.pattern_19 | 53 | | | SP2_HUMAN.H11MO.0.A | 2.360460e-03 | |
| neg_patterns.pattern_20 | 43 | | | SP1_HUMAN.H11MO.0.A | 2.831540e-04 | |
| neg_patterns.pattern_21 | 38 | | | IRF9_HUMAN.H11MO.0.C | 1.000000e+00 | |
| neg_patterns.pattern_22 | 35 | | | MEF2A_MA0052.3 | 5.509490e-02 | |
| neg_patterns.pattern_23 | 35 | | | PRDM6_HUMAN.H11MO.0.C | 4.227550e-02 | |
