## Supplementary Files 1 for "ChromBPNet: bias factorized, base-resolution deep learning models of chromatin accessibility reveal cis-regulatory sequence syntax, transcription factor footprints and regulatory variants": fig1j_gm12878_ATAC_tobias_corrected_softmax_bpnet_profile_modisco.pdf

| pattern | num_seqlets | cwm_fwd | cwm_rev | TOMTOM_match | TOMTOM_qval | TOMTOM_match_logo |
|---|---|---|---|---|---|---|
| pos_patterns.pattern_0 | 12552 | | | TN5_1 | 5.663620e-09 | |
| pos_patterns.pattern_1 | 5748 | | | TN5_1 | 4.147250e-07 | |
| pos_patterns.pattern_2 | 1322 | | | SP4_HUMAN.H11MO.0.A | 2.359510e-04 | |
| pos_patterns.pattern_3 | 1079 | | | NFYB_HUMAN.H11MO.0.A | 4.048520e-05 | |
| pos_patterns.pattern_4 | 1040 | | | CTCF_MA0139.1 | 5.939280e-12 | |
| pos_patterns.pattern_5 | 746 | | | SP2_HUMAN.H11MO.0.A | 1.788920e-03 | |
| pos_patterns.pattern_6 | 643 | | | ELK1_ETS_1 | 1.436010e-03 | |
| pos_patterns.pattern_7 | 437 | | | JUND_MOUSE.H11MO.0.A | 5.345460e-07 | |
| pos_patterns.pattern_8 | 434 | | | ETS1_HUMAN.H11MO.0.A | 2.060740e-03 | |
| pos_patterns.pattern_9 | 362 | | | NRF1_MA0506.1 | 5.675820e-06 | |
| pos_patterns.pattern_10 | 288 | | | USF2_HUMAN.H11MO.0.A | 6.085520e-03 | |
| pos_patterns.pattern_11 | 232 | | | IRF4_HUMAN.H11MO.0.A | 4.422980e-03 | |
| pos_patterns.pattern_12 | 202 | | | IRF4_HUMAN.H11MO.0.A | 8.420850e-13 | |
| pos_patterns.pattern_13 | 168 | | | SP3_HUMAN.H11MO.0.B | 3.366770e-03 | |
| pos_patterns.pattern_14 | 151 | | | SP2_HUMAN.H11MO.0.A | 4.925510e-03 | |
| pos_patterns.pattern_15 | 99 | | | NFYB_HUMAN.H11MO.0.A | 3.037540e-04 | |
| pos_patterns.pattern_16 | 91 | | | ELK1_ETS_4 | 5.428270e-04 | |
| pos_patterns.pattern_17 | 57 | | | TN5_6 | 6.151010e-08 | |
| pos_patterns.pattern_18 | 49 | | | ZN143_MOUSE.H11MO.0.A | 1.217090e-03 | |
| pos_patterns.pattern_19 | 47 | | | NaN | NaN | |
| pos_patterns.pattern_20 | 39 | | | Gabpa_MA0062.2 | 1.149790e-01 | |
| pos_patterns.pattern_21 | 29 | | | JUND_MOUSE.H11MO.0.A | 4.139050e-03 | |
| pos_patterns.pattern_22 | 20 | | | TN5_1 | 9.834180e-01 | |
| neg_patterns.pattern_0 | 2365 | | | NFAC4_HUMAN.H11MO.0.C | 1.000000e+00 | |
| neg_patterns.pattern_1 | 1768 | | | SP1_HUMAN.H11MO.0.A | 1.451810e-06 | |
| neg_patterns.pattern_2 | 1354 | | | ZNF384_MA1125.1 | 4.172670e-03 | |
| neg_patterns.pattern_3 | 991 | | | NFATC1_NFAT_1 | 1.000000e+00 | |
| neg_patterns.pattern_4 | 981 | | | SP2_HUMAN.H11MO.0.A | 1.182470e-07 | |
| neg_patterns.pattern_5 | 945 | | | ZFX_MOUSE.H11MO.0.B | 3.481630e-04 | |
| neg_patterns.pattern_6 | 864 | | | ZFX_MOUSE.H11MO.0.B | 8.140090e-04 | |

| pattern | num_seqlets | cwm_fwd | cwm_rev | TOMTOM_match | TOMTOM_qval | TOMTOM_match_logo |
|---|---|---|---|---|---|---|
| neg_patterns.pattern_7 | 773 | 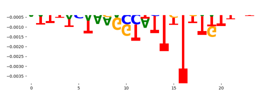 | 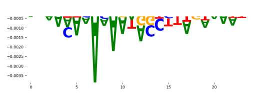 | ZNF713_C2H2_1 | 1.000000e+00 | 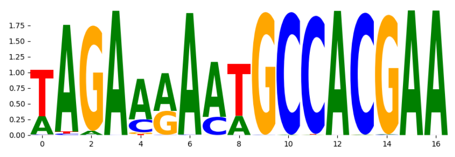 |
| neg_patterns.pattern_8 | 759 | 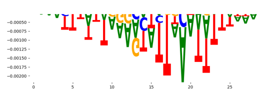 | 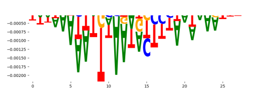 | NFAC2_HUMAN.H11MO.0.B | 1.000000e+00 | 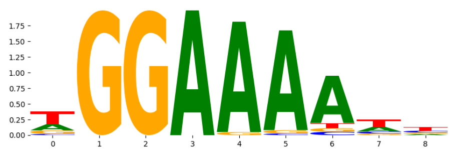 |
| neg_patterns.pattern_9 | 650 | 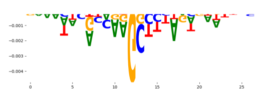 | 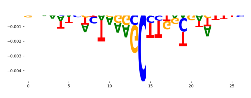 | PATZ1_HUMAN.H11MO.0.C | 1.924410e-03 | 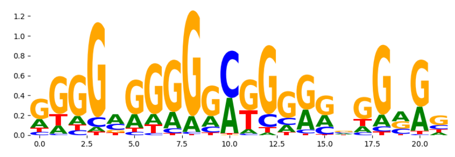 |
| neg_patterns.pattern_10 | 630 | 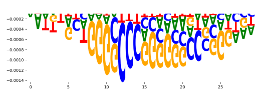 | 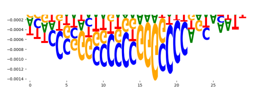 | SP1_HUMAN.H11MO.0.A | 2.269010e-04 | 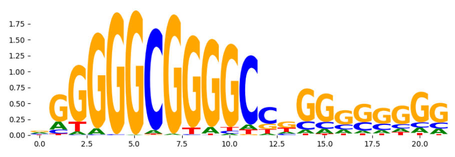 |
| neg_patterns.pattern_11 | 625 | 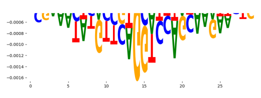 | 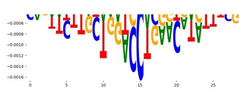 | ZFX_MOUSE.H11MO.0.B | 7.234320e-05 | 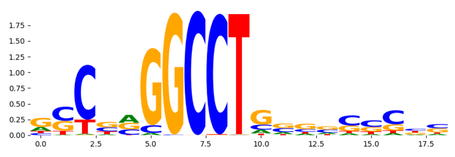 |
| neg_patterns.pattern_12 | 586 | 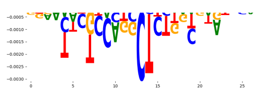 | 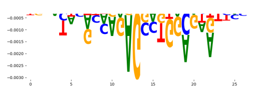 | WT1_HUMAN.H11MO.0.C | 1.638130e-05 | 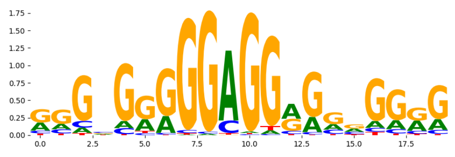 |
| neg_patterns.pattern_13 | 407 | 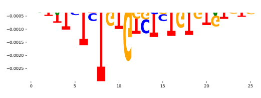 | 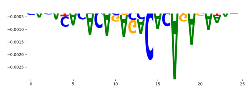 | RREB1_MA0073.1 | 9.233300e-01 | 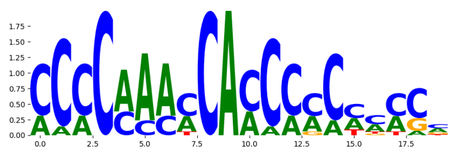 |
| neg_patterns.pattern_14 | 377 | 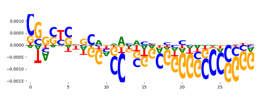 | 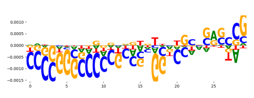 | SP2_HUMAN.H11MO.0.A | 8.961810e-06 | 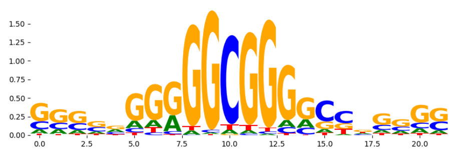 |
| neg_patterns.pattern_15 | 376 | 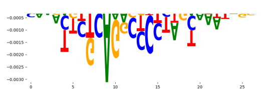 | 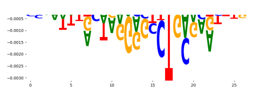 | ZFX_MOUSE.H11MO.0.B | 1.339560e-08 | 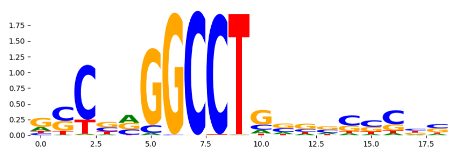 |
| neg_patterns.pattern_16 | 358 | 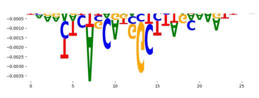 | 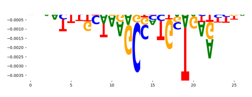 | ZFX_MOUSE.H11MO.0.B | 4.109220e-04 | 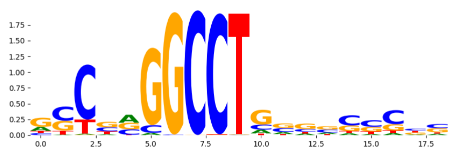 |
| neg_patterns.pattern_17 | 140 | 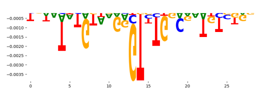 | 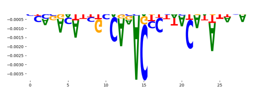 | RREB1_MA0073.1 | 9.889390e-01 | 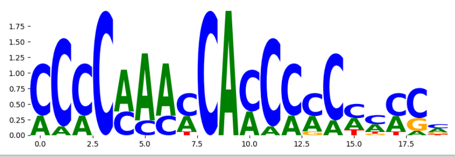 |
