## Supplementary Files 1 for "ChromBPNet: bias factorized, base-resolution deep learning models of chromatin accessibility reveal cis-regulatory sequence syntax, transcription factor footprints and regulatory variants": fig1k_gm12878_ATAC_tobias_corrected_shifted_bpnet_counts_modisco.pdf

| pattern | num_seqlets | cwm_fwd | cwm_rev | TOMTOM_match | TOMTOM_qval | TOMTOM_match_logo |
|---|---|---|---|---|---|---|
| pos_patterns.pattern_0 | 3666 | | | IRF1_MOUSE.H11MO.0.A | 1.230180e-03 | |
| pos_patterns.pattern_1 | 2970 | | | BC11A_HUMAN.H11MO.0.A | 2.846230e-04 | |
| pos_patterns.pattern_2 | 2743 | | | ELK1_ETS_1 | 7.783460e-02 | |
| pos_patterns.pattern_3 | 2019 | | | ATF3_MOUSE.H11MO.0.A | 4.503730e-03 | |
| pos_patterns.pattern_4 | 1920 | | | KLF12_HUMAN.H11MO.0.C | 1.621440e-05 | |
| pos_patterns.pattern_5 | 1729 | | | RELA_MA0107.1 | 1.783870e-04 | |
| pos_patterns.pattern_6 | 1221 | | | RUNX1_HUMAN.H11MO.0.A | 2.627720e-04 | |
| pos_patterns.pattern_7 | 854 | | | IRF4_HUMAN.H11MO.0.A | 2.717980e-11 | |
| pos_patterns.pattern_8 | 783 | | | NFKB2_HUMAN.H11MO.0.B | 3.426330e-03 | |
| pos_patterns.pattern_9 | 626 | | | CTCF_HUMAN.H11MO.0.A | 3.118780e-12 | |
| pos_patterns.pattern_10 | 586 | | | Atf1_MA0604.1 | 4.157550e-03 | |
| pos_patterns.pattern_11 | 583 | | | NRF1_HUMAN.H11MO.0.A | 3.174730e-07 | |
| pos_patterns.pattern_12 | 567 | | | IRF5_IRF_2 | 4.258940e-02 | |
| pos_patterns.pattern_13 | 530 | | | NFYB_HUMAN.H11MO.0.A | 2.407970e-04 | |
| pos_patterns.pattern_14 | 219 | | | THA11_HUMAN.H11MO.0.B | 9.242580e-12 | |
| pos_patterns.pattern_15 | 39 | | | JUN_HUMAN.H11MO.0.A | 2.541160e-01 | |
| pos_patterns.pattern_16 | 32 | | | NRF1_MA0506.1 | 5.971230e-01 | |
| pos_patterns.pattern_17 | 27 | | | RUNX2_MA0511.2 | 3.848450e-01 | |
| neg_patterns.pattern_0 | 440 | | | CTCF_MA0139.1 | 2.022710e-07 | |
| neg_patterns.pattern_1 | 74 | | | NFAC2_HUMAN.H11MO.0.B | 1.933690e-02 | |
| neg_patterns.pattern_2 | 38 | | | SP2_HUMAN.H11MO.0.A | 7.257830e-09 | |

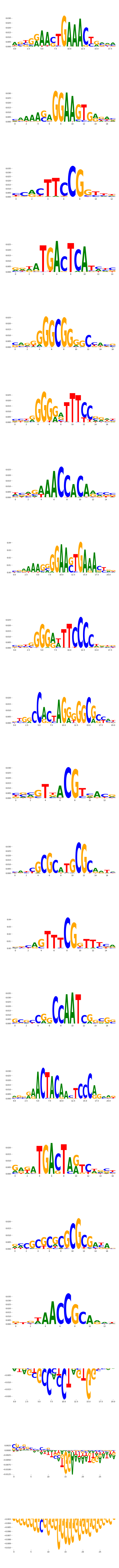

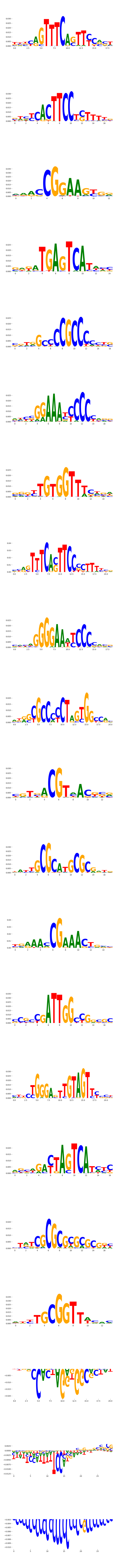

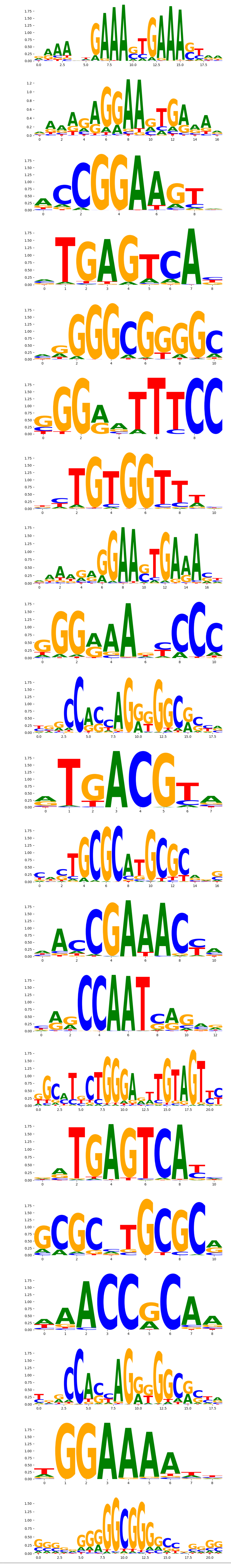
