## Supplementary Files 1 for "ChromBPNet: bias factorized, base-resolution deep learning models of chromatin accessibility reveal cis-regulatory sequence syntax, transcription factor footprints and regulatory variants": fig1l_gm12878_ATAC_tobias_corrected_shifted_bpnet_profile_modisco.pdf

| pattern | num_seqlets | cwm_fwd | cwm_rev | TOMTOM_match | TOMTOM_qval | TOMTOM_match_logo |
|---|---|---|---|---|---|---|
| pos_patterns.pattern_0 | 7247 | | | TN5_1 | 5.966190e-07 | |
| pos_patterns.pattern_1 | 6293 | | | TN5_1 | 4.587720e-09 | |
| pos_patterns.pattern_2 | 2916 | | | CTCF_MA0139.1 | 3.954930e-14 | |
| pos_patterns.pattern_3 | 1182 | | | SP1_HUMAN.H11MO.0.A | 2.542470e-06 | |
| pos_patterns.pattern_4 | 1098 | | | IRF1_MOUSE.H11MO.0.A | 1.347470e-05 | |
| pos_patterns.pattern_5 | 966 | | | ZFX_MOUSE.H11MO.0.B | 9.952670e-05 | |
| pos_patterns.pattern_6 | 964 | | | ATF3_MOUSE.H11MO.0.A | 4.114900e-03 | |
| pos_patterns.pattern_7 | 956 | | | ETV4_MOUSE.H11MO.0.B | 1.387850e-04 | |
| pos_patterns.pattern_8 | 673 | | | TN5_1 | 1.052150e-03 | |
| pos_patterns.pattern_9 | 606 | | | NFYB_HUMAN.H11MO.0.A | 5.641940e-05 | |
| pos_patterns.pattern_10 | 540 | | | SPIB_MOUSE.H11MO.0.A | 6.272120e-08 | |
| pos_patterns.pattern_11 | 422 | | | NFKB1_HUMAN.H11MO.1.B | 4.463950e-06 | |
| pos_patterns.pattern_12 | 410 | | | SP2_HUMAN.H11MO.0.A | 6.773570e-05 | |
| pos_patterns.pattern_13 | 383 | | | ATF1_MOUSE.H11MO.0.B | 1.533150e-03 | |
| pos_patterns.pattern_14 | 383 | | | NRF1_HUMAN.H11MO.0.A | 6.465060e-08 | |
| pos_patterns.pattern_15 | 225 | | | ZFX_MOUSE.H11MO.0.B | 1.450120e-04 | |
| pos_patterns.pattern_16 | 225 | | | THA11_HUMAN.H11MO.0.B | 1.142100e-07 | |
| pos_patterns.pattern_17 | 210 | | | NRF1_MA0506.1 | 1.000000e+00 | |
| pos_patterns.pattern_18 | 140 | | | TN5_3 | 7.479780e-04 | |
| pos_patterns.pattern_19 | 105 | | | ZFX_MOUSE.H11MO.0.B | 2.247100e-05 | |
| pos_patterns.pattern_20 | 101 | | | SP2_HUMAN.H11MO.0.A | 5.687900e-05 | |
| pos_patterns.pattern_21 | 50 | | | BACH1_HUMAN.H11MO.0.A | 5.408480e-03 | |
| pos_patterns.pattern_22 | 44 | | | TF65_HUMAN.H11MO.0.A | 3.422850e-04 | |
| pos_patterns.pattern_23 | 34 | | | Gabpa_MA0062.2 | 3.521410e-01 | |
| pos_patterns.pattern_24 | 27 | | | SP1_HUMAN.H11MO.0.A | 4.457970e-04 | |
| pos_patterns.pattern_25 | 20 | | | RUNX1_MA0002.2 | 3.934640e-03 | |
