## Supplementary Files 1 for "ChromBPNet: bias factorized, base-resolution deep learning models of chromatin accessibility reveal cis-regulatory sequence syntax, transcription factor footprints and regulatory variants": fig1m_gm12878_ATAC_raw_bpnet_w_simple_cnn_bias_counts_modisco.pdf

| pattern | num_seqlets | cwm_fwd | cwm_rev | TOMTOM_match | TOMTOM_qval | TOMTOM_match_logo |
|---|---|---|---|---|---|---|
| pos_patterns.pattern_0 | 4336 | | | IRF1_MOUSE.H11MO.0.A | 1.834170e-04 | |
| pos_patterns.pattern_1 | 3248 | | | CTCF_MA0139.1 | 2.421050e-15 | |
| pos_patterns.pattern_2 | 2992 | | | SPI1_HUMAN.H11MO.0.A | 7.559290e-07 | |
| pos_patterns.pattern_3 | 2777 | | | JDP2_MA0655.1 | 2.685390e-03 | |
| pos_patterns.pattern_4 | 2230 | | | RUNX1_HUMAN.H11MO.0.A | 1.528440e-02 | |
| pos_patterns.pattern_5 | 1639 | | | NFKB2_HUMAN.H11MO.0.B | 3.018970e-06 | |
| pos_patterns.pattern_6 | 581 | | | ELK4_MA0076.2 | 1.510160e-05 | |
| pos_patterns.pattern_7 | 566 | | | NFKB1_HUMAN.H11MO.1.B | 4.476840e-03 | |
| pos_patterns.pattern_8 | 488 | | | KLF12_HUMAN.H11MO.0.C | 8.851910e-05 | |
| pos_patterns.pattern_9 | 446 | | | REL_MA0101.1 | 1.874080e-01 | |
| pos_patterns.pattern_10 | 417 | | | Mafb.mouse_bZIP_3 | 1.206350e-04 | |
| pos_patterns.pattern_11 | 331 | | | NFYB_HUMAN.H11MO.0.A | 1.284900e-03 | |
| pos_patterns.pattern_12 | 309 | | | POU5F1_MA1115.1 | 4.193960e-03 | |
| pos_patterns.pattern_13 | 243 | | | EBF1_EBF_1 | 5.866960e-06 | |
| pos_patterns.pattern_14 | 204 | | | NRF1_MOUSE.H11MO.0.A | 3.730520e-07 | |
| pos_patterns.pattern_15 | 197 | | | HNF1B_HUMAN.H11MO.0.A | 1.156310e-06 | |
| pos_patterns.pattern_16 | 164 | | | PAX5_HUMAN.H11MO.0.A | 9.524420e-09 | |
| pos_patterns.pattern_17 | 163 | | | MEF2A_MOUSE.H11MO.0.A | 3.959170e-08 | |
| pos_patterns.pattern_18 | 114 | | | ZNF76_HUMAN.H11MO.0.C | 6.548520e-19 | |
| pos_patterns.pattern_19 | 104 | | | SPI1_ETS_1 | 4.771300e-03 | |
| pos_patterns.pattern_20 | 82 | | | PAX2_PAX_1 | 2.871660e-06 | |
| pos_patterns.pattern_21 | 55 | | | ZBTB33_MA0527.1 | 3.952770e-05 | |
| pos_patterns.pattern_22 | 50 | | | PAX2_PAX_1 | 4.706490e-04 | |
| pos_patterns.pattern_23 | 20 | | | CTCF_MOUSE.H11MO.0.A | 2.120740e-03 | |
| neg_patterns.pattern_0 | 42 | | | NFAC2_HUMAN.H11MO.0.B | 2.377100e-02 | |
| neg_patterns.pattern_1 | 38 | | | TAF1_MOUSE.H11MO.0.A | 5.239090e-01 | |
| neg_patterns.pattern_2 | 35 | | | STAT1_HUMAN.H11MO.0.A | 5.974570e-01 | |
| neg_patterns.pattern_3 | 28 | | | RELB_MA1117.1 | 4.317820e-01 | |
