## Supplementary Files 1 for "ChromBPNet: bias factorized, base-resolution deep learning models of chromatin accessibility reveal cis-regulatory sequence syntax, transcription factor footprints and regulatory variants": fig1n_gm12878_ATAC_raw_bpnet_w_simple_cnn_bias_profile_modisco.pdf

| pattern | num_seqlets | cwm_fwd | cwm_rev | TOMTOM_match | TOMTOM_qval | TOMTOM_match_logo |
|---|---|---|---|---|---|---|
| pos_patterns.pattern_0 | 4237 | | | ZKSC1_HUMAN.H11MO.0.B | 5.527610e-01 | |
| pos_patterns.pattern_1 | 4113 | | | CTCF_MA0139.1 | 1.703100e-12 | |
| pos_patterns.pattern_2 | 2360 | | | JDP2_MA0655.1 | 4.416050e-04 | |
| pos_patterns.pattern_3 | 1775 | | | SPIB_MOUSE.H11MO.0.A | 5.722130e-07 | |
| pos_patterns.pattern_4 | 1189 | | | FOXL1_forkhead_2 | 7.336600e-01 | |
| pos_patterns.pattern_5 | 1120 | | | RUNX1_HUMAN.H11MO.0.A | 1.091480e-06 | |
| pos_patterns.pattern_6 | 1071 | | | IRF1_MOUSE.H11MO.0.A | 9.615410e-04 | |
| pos_patterns.pattern_7 | 898 | | | REL_MA0101.1 | 1.281250e-04 | |
| pos_patterns.pattern_8 | 795 | | | NaN | NaN | |
| pos_patterns.pattern_9 | 521 | | | NFYB_HUMAN.H11MO.0.A | 9.825340e-10 | |
| pos_patterns.pattern_10 | 518 | | | IRF1_MOUSE.H11MO.0.A | 3.547410e-07 | |
| pos_patterns.pattern_11 | 474 | | | IRF4_HUMAN.H11MO.0.A | 7.017240e-06 | |
| pos_patterns.pattern_12 | 352 | | | IRF1_HUMAN.H11MO.0.A | 2.010720e-04 | |
| pos_patterns.pattern_13 | 306 | | | FOSL2+JUN_MA1131.1 | 6.691600e-05 | |
| pos_patterns.pattern_14 | 176 | | | KLF12_HUMAN.H11MO.0.C | 3.817100e-04 | |
| pos_patterns.pattern_15 | 120 | | | Pou2f2.mouse_POU_2 | 2.084390e-03 | |
| pos_patterns.pattern_16 | 116 | | | RUNX2_MA0511.2 | 5.678670e-01 | |
| pos_patterns.pattern_17 | 111 | | | NRF1_MOUSE.H11MO.0.A | 9.917260e-05 | |
| pos_patterns.pattern_18 | 89 | | | ELK1_MOUSE.H11MO.0.B | 7.022940e-05 | |
| pos_patterns.pattern_19 | 83 | | | ZNF76_HUMAN.H11MO.0.C | 1.241290e-10 | |
| pos_patterns.pattern_20 | 63 | | | PEBB_HUMAN.H11MO.0.C | 3.851560e-03 | |
| pos_patterns.pattern_21 | 37 | | | Hic1.mouse_C2H2_1 | 4.453450e-01 | |
| pos_patterns.pattern_22 | 33 | | | TN5_8 | 1.000000e+00 | |
| pos_patterns.pattern_23 | 30 | | | PAX6_HUMAN.H11MO.0.C | 1.253830e-01 | |
| neg_patterns.pattern_0 | 26 | | | SPIB_MOUSE.H11MO.0.A | 1.395230e-01 | |
