## Supplementary Files 1 for "ChromBPNet: bias factorized, base-resolution deep learning models of chromatin accessibility reveal cis-regulatory sequence syntax, transcription factor footprints and regulatory variants": fig1o_gm12878_ATAC_raw_chrombpnet_counts_modisco.pdf

| pattern | num_seqlets | cwm_fwd | cwm_rev | TOMTOM_match | TOMTOM_qval | TOMTOM_match_logo |
|---|---|---|---|---|---|---|
| pos_patterns.pattern_0 | 3247 | | | CTCF_MA0139.1 | 5.308560e-13 | |
| pos_patterns.pattern_1 | 2898 | | | IRF4_MOUSE.H11MO.0.A | 4.201300e-02 | |
| pos_patterns.pattern_2 | 2751 | | | ELF5_HUMAN.H11MO.0.A | 2.734420e-04 | |
| pos_patterns.pattern_3 | 2049 | | | FOS+JUND_MA1141.1 | 2.575030e-03 | |
| pos_patterns.pattern_4 | 1904 | | | IRF1_MOUSE.H11MO.0.A | 9.016690e-06 | |
| pos_patterns.pattern_5 | 1821 | | | RUNX3_HUMAN.H11MO.0.A | 1.828630e-02 | |
| pos_patterns.pattern_6 | 1612 | | | RELB_HUMAN.H11MO.0.C | 3.794730e-06 | |
| pos_patterns.pattern_7 | 807 | | | SP1_HUMAN.H11MO.0.A | 1.591760e-05 | |
| pos_patterns.pattern_8 | 799 | | | ELK4_HUMAN.H11MO.0.A | 1.005590e-05 | |
| pos_patterns.pattern_9 | 375 | | | REL_MA0101.1 | 1.826090e-01 | |
| pos_patterns.pattern_10 | 335 | | | NFYC_HUMAN.H11MO.0.A | 6.717320e-03 | |
| pos_patterns.pattern_11 | 322 | | | NFKB1_HUMAN.H11MO.1.B | 6.422330e-03 | |
| pos_patterns.pattern_12 | 309 | | | Pou2f2.mouse_POU_2 | 5.568920e-03 | |
| pos_patterns.pattern_13 | 282 | | | BATF_HUMAN.H11MO.0.A | 1.126030e-06 | |
| pos_patterns.pattern_14 | 277 | | | NRF1_MOUSE.H11MO.0.A | 2.666620e-07 | |
| pos_patterns.pattern_15 | 217 | | | Mafb.mouse_bZIP_3 | 1.999360e-04 | |
| pos_patterns.pattern_16 | 212 | | | ETS1_HUMAN.H11MO.0.A | 2.905130e-04 | |
| pos_patterns.pattern_17 | 200 | | | COE1_MOUSE.H11MO.0.A | 2.760570e-06 | |
| pos_patterns.pattern_18 | 171 | | | ZN143_MOUSE.H11MO.0.A | 4.361840e-14 | |
| pos_patterns.pattern_19 | 163 | | | MEF2D_HUMAN.H11MO.0.A | 2.501050e-10 | |
| pos_patterns.pattern_20 | 149 | | | SPIB_ETS_1 | 1.969570e-02 | |
| pos_patterns.pattern_21 | 123 | | | PAX2_PAX_1 | 3.134400e-08 | |
| pos_patterns.pattern_22 | 89 | | | HNF1B_HUMAN.H11MO.0.A | 9.561310e-06 | |
| pos_patterns.pattern_23 | 85 | | | PAX5_MOUSE.H11MO.0.A | 8.230710e-08 | |
| pos_patterns.pattern_24 | 83 | | | TYY1_HUMAN.H11MO.0.A | 2.496250e-07 | |
| pos_patterns.pattern_25 | 35 | | | ZBTB33_MA0527.1 | 6.928390e-06 | |
| pos_patterns.pattern_26 | 34 | | | PAX2_PAX_1 | 4.710900e-04 | |
| pos_patterns.pattern_27 | 21 | | | THRA_nuclearreceptor_1 | 2.809530e-02 | |
| neg_patterns.pattern_0 | 33 | | | NFATC2_MA0152.1 | 3.798600e-02 | |
