## Supplementary Files 1 for "ChromBPNet: bias factorized, base-resolution deep learning models of chromatin accessibility reveal cis-regulatory sequence syntax, transcription factor footprints and regulatory variants": fig1p_gm12878_ATAC_raw_chrombpnet_profile_modisco.pdf

| pattern | num_seqlets | cwm_fwd | cwm_rev | TOMTOM_match | TOMTOM_qval | TOMTOM_match_logo |
|---|---|---|---|---|---|---|
| pos_patterns.pattern_0 | 4606 | | | CTCF_MA0139.1 | 3.632500e-14 | |
| pos_patterns.pattern_1 | 3246 | | | STAT1+STAT2_MA0517.1 | 1.433520e-01 | |
| pos_patterns.pattern_2 | 2725 | | | JDP2_MA0655.1 | 2.768140e-03 | |
| pos_patterns.pattern_3 | 2423 | | | BC11A_HUMAN.H11MO.0.A | 6.951580e-05 | |
| pos_patterns.pattern_4 | 2254 | | | RUNX1_HUMAN.H11MO.0.A | 5.247210e-02 | |
| pos_patterns.pattern_5 | 2050 | | | TF65_HUMAN.H11MO.0.A | 1.136670e-05 | |
| pos_patterns.pattern_6 | 1980 | | | IRF8_HUMAN.H11MO.0.B | 6.938850e-06 | |
| pos_patterns.pattern_7 | 1632 | | | KLF12_HUMAN.H11MO.0.C | 4.112910e-05 | |
| pos_patterns.pattern_8 | 885 | | | NFYB_HUMAN.H11MO.0.A | 3.936070e-04 | |
| pos_patterns.pattern_9 | 775 | | | Gabpa_MA0062.2 | 6.718830e-05 | |
| pos_patterns.pattern_10 | 590 | | | NRF1_MA0506.1 | 1.852890e-08 | |
| pos_patterns.pattern_11 | 453 | | | POU5F1_MA1115.1 | 4.135300e-03 | |
| pos_patterns.pattern_12 | 409 | | | BATF_HUMAN.H11MO.0.A | 7.153110e-08 | |
| pos_patterns.pattern_13 | 336 | | | FOSL2+JUN_MA1131.1 | 1.302120e-04 | |
| pos_patterns.pattern_14 | 268 | | | TYY1_MOUSE.H11MO.0.A | 8.967910e-07 | |
| pos_patterns.pattern_15 | 259 | | | JUN_MA0488.1 | 4.398200e-03 | |
| pos_patterns.pattern_16 | 244 | | | IRF9_IRF_1 | 4.245900e-04 | |
| pos_patterns.pattern_17 | 243 | | | ZN143_MOUSE.H11MO.0.A | 2.222570e-18 | |
| pos_patterns.pattern_18 | 160 | | | MITF_HUMAN.H11MO.0.A | 5.099030e-05 | |
| pos_patterns.pattern_19 | 138 | | | HNF1A_HUMAN.H11MO.0.C | 7.704190e-05 | |
| pos_patterns.pattern_20 | 118 | | | SPI1_ETS_1 | 1.635400e-02 | |
| pos_patterns.pattern_21 | 113 | | | RELB_MA1117.1 | 7.435450e-01 | |
| pos_patterns.pattern_22 | 108 | | | ATF3_MOUSE.H11MO.0.A | 3.099670e-01 | |
| pos_patterns.pattern_23 | 85 | | | PAX2_PAX_1 | 7.215390e-07 | |
| pos_patterns.pattern_24 | 51 | | | Hic1.mouse_C2H2_1 | 7.661810e-01 | |
| pos_patterns.pattern_25 | 43 | | | PAX6_PAX_1 | 1.275950e-04 | |
| pos_patterns.pattern_26 | 42 | | | Rfx1_MA0509.1 | 8.271380e-09 | |
| pos_patterns.pattern_27 | 34 | | | MEF2A_MA0052.3 | 9.347320e-07 | |
| pos_patterns.pattern_28 | 32 | | | KLF12_HUMAN.H11MO.0.C | 2.487710e-05 | |

| pattern | num_seqlets | cwm_fwd | cwm_rev | TOMTOM_match | TOMTOM_qval | TOMTOM_match_logo |
|---|---|---|---|---|---|---|
| pos_patterns.pattern_29 | 31 | | | PRDM1_HUMAN.H11MO.0.A | 5.607540e-01 | |
| pos_patterns.pattern_30 | 27 | | | PAX5_MOUSE.H11MO.0.A | 3.016880e-07 | |
| pos_patterns.pattern_31 | 27 | | | ZBTB33_MA0527.1 | 4.313920e-05 | |
| pos_patterns.pattern_32 | 27 | | | POU2F2_MA0507.1 | 1.456120e-01 | |
| pos_patterns.pattern_33 | 22 | | | JUN_MA0488.1 | 4.814100e-03 | |
| neg_patterns.pattern_0 | 796 | | | NFATC2_MA0152.1 | 3.082730e-01 | |
| neg_patterns.pattern_1 | 476 | | | ZBT7A_HUMAN.H11MO.0.A | 8.705380e-04 | |
| neg_patterns.pattern_2 | 276 | | | RREB1_MA0073.1 | 1.530550e-02 | |
| neg_patterns.pattern_3 | 190 | | | IRF4_HUMAN.H11MO.0.A | 9.010000e-03 | |
| neg_patterns.pattern_4 | 189 | | | NFAC1_HUMAN.H11MO.0.B | 5.006260e-01 | |
| neg_patterns.pattern_5 | 160 | | | NFAC3_HUMAN.H11MO.0.B | 2.500410e-01 | |
| neg_patterns.pattern_6 | 150 | | | SPIB_MOUSE.H11MO.0.A | 2.806570e-04 | |
| neg_patterns.pattern_7 | 139 | | | ZNF384_MA1125.1 | 5.876630e-03 | |
| neg_patterns.pattern_8 | 136 | | | DNASE_2 | 1.400110e-01 | |
| neg_patterns.pattern_9 | 133 | | | SMCA5_MOUSE.H11MO.0.C | 1.223470e-01 | |
| neg_patterns.pattern_10 | 129 | | | RELA_MA0107.1 | 1.777080e-03 | |
| neg_patterns.pattern_11 | 93 | | | JDP2_bZIP_1 | 4.234990e-05 | |
| neg_patterns.pattern_12 | 90 | | | ZNF384_MA1125.1 | 3.263780e-01 | |
| neg_patterns.pattern_13 | 86 | | | TBX3_HUMAN.H11MO.0.C | 1.000000e+00 | |
| neg_patterns.pattern_14 | 85 | | | ZNF384_MA1125.1 | 4.805640e-01 | |
| neg_patterns.pattern_15 | 76 | | | ZBT7A_HUMAN.H11MO.0.A | 1.071990e-01 | |
| neg_patterns.pattern_16 | 75 | | | IRF3_HUMAN.H11MO.0.B | 1.635960e-01 | |
| neg_patterns.pattern_17 | 70 | | | IRF8_HUMAN.H11MO.0.B | 5.473260e-05 | |
| neg_patterns.pattern_18 | 69 | | | ELK4_ETS_1 | 7.451550e-04 | |
| neg_patterns.pattern_19 | 66 | | | SPI1_HUMAN.H11MO.0.A | 4.871920e-02 | |
| neg_patterns.pattern_20 | 58 | | | ONECUT3_CUT_1 | 2.024900e-01 | |
| neg_patterns.pattern_21 | 56 | | | NaN | NaN | |
| neg_patterns.pattern_22 | 51 | | | CTCF_HUMAN.H11MO.0.A | 1.000000e+00 | |
| neg_patterns.pattern_23 | 44 | | | BCL6B_C2H2_1 | 1.000000e+00 | |
| neg_patterns.pattern_24 | 44 | | | SOX7_HMG_3 | 2.522630e-01 | |

| pattern | num_seqlets | cwm_fwd | cwm_rev | TOMTOM_match | TOMTOM_qval | TOMTOM_match_logo |
|---|---|---|---|---|---|---|
| neg_patterns.pattern_25 | 36 | 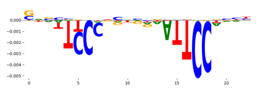 | 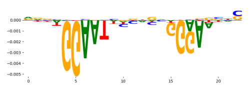 | TEAD3_MA0808.1 | 3.810550e-01 | 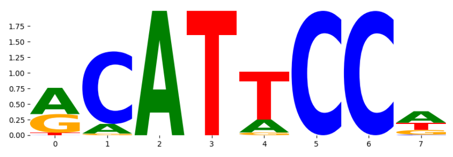 |
| neg_patterns.pattern_26 | 32 | 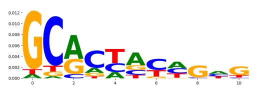 | 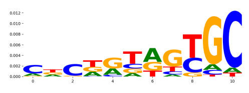 | ZN264_HUMAN.H11MO.0.C | 1.000000e+00 | 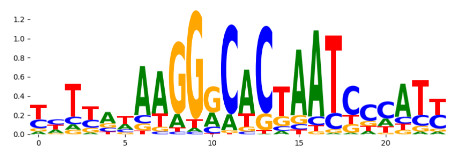 |
| neg_patterns.pattern_27 | 29 | 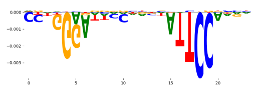 | 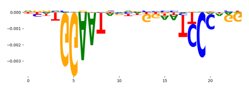 | TEAD3_MA0808.1 | 3.227880e-01 | 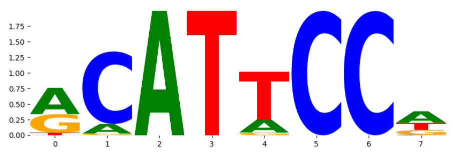 |
| neg_patterns.pattern_28 | 28 | 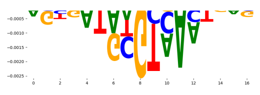 | 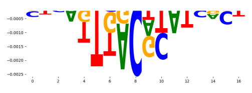 | Foxg1.mouse_forkhead_1 | 3.832810e-01 | 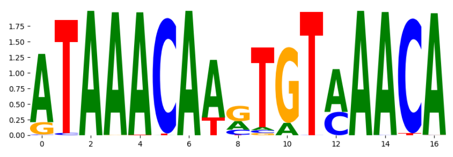 |
| neg_patterns.pattern_29 | 27 | 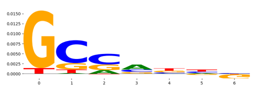 | 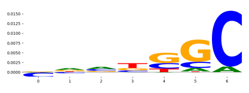 | ZEB1_MA0103.3 | 1.000000e+00 | 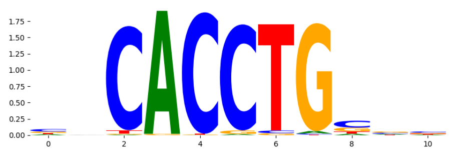 |
| neg_patterns.pattern_30 | 26 | 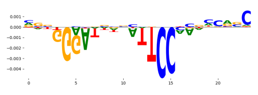 | 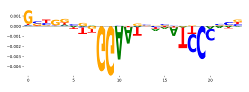 | SOX18_HMG_3 | 7.999790e-01 | 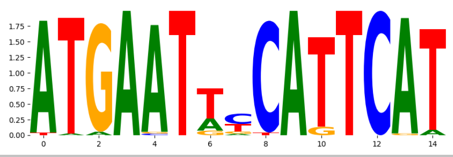 |
