## Supplementary Files 2 for "ChromBPNet: bias factorized, base-resolution deep learning models of chromatin accessibility reveal cis-regulatory sequence syntax, transcription factor footprints and regulatory variants": fig2a_gm12878_DNASE_raw_bpnet_uncorrected_counts_modisco.pdf

| pattern | num_seqlets | cwm_fwd | cwm_rev | TOMTOM_match | TOMTOM_qval | TOMTOM_match_logo |
|---|---|---|---|---|---|---|
| pos_patterns.pattern_0 | 4409 | | | IRF1_MOUSE.H11MO.0.A | 7.444490e-04 | |
| pos_patterns.pattern_1 | 3684 | | | SPIB_MOUSE.H11MO.0.A | 1.480820e-05 | |
| pos_patterns.pattern_2 | 3044 | | | RUNX1_HUMAN.H11MO.0.A | 4.132100e-03 | |
| pos_patterns.pattern_3 | 2536 | | | JUNB_HUMAN.H11MO.0.A | 2.812630e-03 | |
| pos_patterns.pattern_4 | 2111 | | | NFKB1_HUMAN.H11MO.1.B | 2.768810e-08 | |
| pos_patterns.pattern_5 | 2085 | | | CTCF_MA0139.1 | 1.274450e-11 | |
| pos_patterns.pattern_6 | 1796 | | | SP1_HUMAN.H11MO.0.A | 5.770860e-05 | |
| pos_patterns.pattern_7 | 1684 | | | Gabpa_MA0062.2 | 4.774250e-05 | |
| pos_patterns.pattern_8 | 1089 | | | ATF3_HUMAN.H11MO.0.A | 4.449200e-03 | |
| pos_patterns.pattern_9 | 982 | | | NRF1_MA0506.1 | 5.413410e-07 | |
| pos_patterns.pattern_10 | 802 | | | NFYB_HUMAN.H11MO.0.A | 1.421390e-03 | |
| pos_patterns.pattern_11 | 382 | | | RELB_HUMAN.H11MO.0.C | 1.000000e+00 | |
| pos_patterns.pattern_12 | 339 | | | POU5F1_MA1115.1 | 3.582870e-03 | |
| pos_patterns.pattern_13 | 299 | | | EBF1_EBF_1 | 1.746020e-05 | |
| pos_patterns.pattern_14 | 290 | | | ZNF76_HUMAN.H11MO.0.C | 8.680800e-22 | |
| pos_patterns.pattern_15 | 261 | | | IRF4_MOUSE.H11MO.0.A | 6.509460e-04 | |
| pos_patterns.pattern_16 | 187 | | | PAX1_MA0779.1 | 2.123530e-06 | |
| pos_patterns.pattern_17 | 144 | | | ZBTB33_MA0527.1 | 1.845290e-04 | |
| pos_patterns.pattern_18 | 111 | | | MEF2D_MOUSE.H11MO.0.A | 1.276620e-06 | |
| pos_patterns.pattern_19 | 65 | | | NFIA_NFI_1 | 7.049680e-04 | |
| pos_patterns.pattern_20 | 63 | | | TYY1_HUMAN.H11MO.0.A | 4.536160e-05 | |
| pos_patterns.pattern_21 | 53 | | | RFX2_HUMAN.H11MO.0.A | 2.041250e-08 | |
| pos_patterns.pattern_22 | 41 | | | Gabpa_MA0062.2 | 1.960340e-01 | |
| pos_patterns.pattern_23 | 41 | | | PAX2_PAX_1 | 4.355510e-07 | |
| pos_patterns.pattern_24 | 39 | | | HNF1B_HUMAN.H11MO.0.A | 9.153840e-06 | |
| pos_patterns.pattern_25 | 37 | | | PRDM6_HUMAN.H11MO.0.C | 4.408250e-02 | |
| pos_patterns.pattern_26 | 37 | | | SP1_HUMAN.H11MO.0.A | 5.460660e-07 | |
| pos_patterns.pattern_27 | 23 | | | PAX2_PAX_1 | 2.247030e-03 | |
| pos_patterns.pattern_28 | 21 | | | PRD14_HUMAN.H11MO.0.A | 1.000000e+00 | |

| pattern | num_seqlets | cwm_fwd | cwm_rev | TOMTOM_match | TOMTOM_qval | TOMTOM_match_logo |
|---|---|---|---|---|---|---|
| pos_patterns.pattern_29 | 20 | 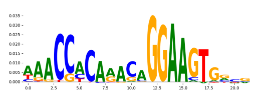 | 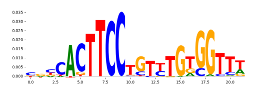 | ETV4_MOUSE.H11MO.0.B | 9.482910e-03 | 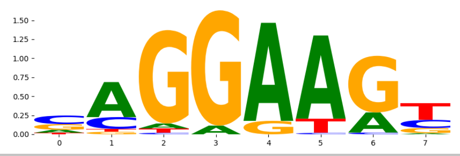 |
