## Supplementary Files 2 for "ChromBPNet: bias factorized, base-resolution deep learning models of chromatin accessibility reveal cis-regulatory sequence syntax, transcription factor footprints and regulatory variants": fig2b_gm12878_DNASE_raw_bpnet_uncorrected_profile_modisco.pdf

| pattern | num_seqlets | cwm_fwd | cwm_rev | TOMTOM_match | TOMTOM_qval | TOMTOM_match_logo |
|---|---|---|---|---|---|---|
| pos_patterns.pattern_0 | 4753 | | | CTCF_MA0139.1 | 2.748960e-12 | |
| pos_patterns.pattern_1 | 2984 | | | ELF5_HUMAN.H11MO.0.A | 9.836690e-05 | |
| pos_patterns.pattern_2 | 2932 | | | RUNX1_HUMAN.H11MO.0.A | 8.685900e-02 | |
| pos_patterns.pattern_3 | 2884 | | | ATF3_MOUSE.H11MO.0.A | 4.129770e-03 | |
| pos_patterns.pattern_4 | 1979 | | | IRF8_HUMAN.H11MO.0.B | 1.040800e-05 | |
| pos_patterns.pattern_5 | 1883 | | | SIX2_MA1119.1 | 5.779520e-02 | |
| pos_patterns.pattern_6 | 825 | | | RELB_HUMAN.H11MO.0.C | 1.352190e-07 | |
| pos_patterns.pattern_7 | 785 | | | RARA_nuclearreceptor_5 | 1.000000e+00 | |
| pos_patterns.pattern_8 | 658 | | | KLF12_HUMAN.H11MO.0.C | 1.278430e-06 | |
| pos_patterns.pattern_9 | 584 | | | NFYC_HUMAN.H11MO.0.A | 8.614680e-06 | |
| pos_patterns.pattern_10 | 571 | | | NRF1_MOUSE.H11MO.0.A | 1.211210e-05 | |
| pos_patterns.pattern_11 | 355 | | | MITF_MA0620.2 | 3.577560e-06 | |
| pos_patterns.pattern_12 | 341 | | | NFKB1_HUMAN.H11MO.1.B | 1.266870e-03 | |
| pos_patterns.pattern_13 | 319 | | | FOSL2+JUN_MA1131.1 | 4.061170e-05 | |
| pos_patterns.pattern_14 | 310 | | | NaN | NaN | |
| pos_patterns.pattern_15 | 277 | | | MAZ_HUMAN.H11MO.0.A | 1.695200e-07 | |
| pos_patterns.pattern_16 | 240 | | | COE1_MOUSE.H11MO.0.A | 4.520040e-05 | |
| pos_patterns.pattern_17 | 233 | | | PO2F2_HUMAN.H11MO.0.A | 9.060280e-04 | |
| pos_patterns.pattern_18 | 232 | | | PAX1_MA0779.1 | 1.601050e-05 | |
| pos_patterns.pattern_19 | 188 | | | EGR2_HUMAN.H11MO.0.A | 2.405910e-04 | |
| pos_patterns.pattern_20 | 173 | | | ZN143_HUMAN.H11MO.0.A | 4.017130e-15 | |
| pos_patterns.pattern_21 | 133 | | | NFIA_HUMAN.H11MO.0.C | 8.056880e-05 | |
| pos_patterns.pattern_22 | 88 | | | RFX3_MOUSE.H11MO.0.C | 1.898380e-10 | |
| pos_patterns.pattern_23 | 62 | | | SPIB_ETS_1 | 5.293210e-02 | |
| pos_patterns.pattern_24 | 48 | | | ZBTB33_MA0527.1 | 6.110090e-03 | |
| pos_patterns.pattern_25 | 25 | | | ZN770_HUMAN.H11MO.0.C | 4.335230e-02 | |
| pos_patterns.pattern_26 | 20 | | | PAX6_PAX_1 | 2.708400e-04 | |
| neg_patterns.pattern_0 | 88 | | | NFATC2_MA0152.1 | 6.944530e-01 | |
| neg_patterns.pattern_1 | 72 | | | DNASE_2 | 1.447250e-01 | |
| neg_patterns.pattern_2 | 70 | | | DNASE_2 | 2.132820e-01 | |

| pattern | num_seqlets | cwm_fwd | cwm_rev | TOMTOM_match | TOMTOM_qval | TOMTOM_match_logo |
|---|---|---|---|---|---|---|
| neg_patterns.pattern_3 | 39 | 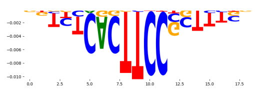 | 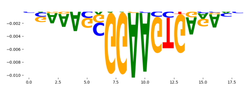 | SPIB_MOUSE.H11MO.0.A | 2.046650e-05 | 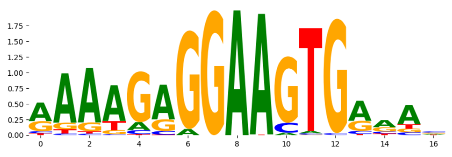 |
| neg_patterns.pattern_4 | 27 | 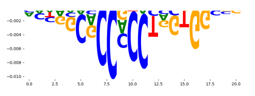 | 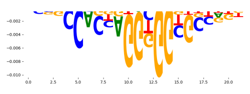 | CTCF_MA0139.1 | 7.212070e-04 | 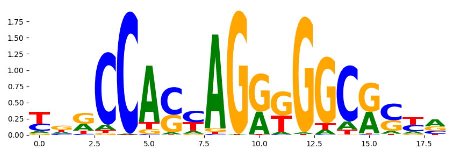 |
| neg_patterns.pattern_5 | 22 | 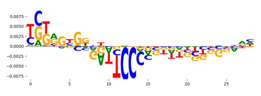 | 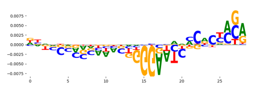 | FLI1_ETS_4 | 1.000000e+00 | 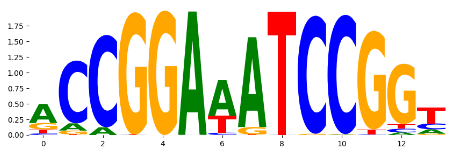 |
| neg_patterns.pattern_6 | 22 | 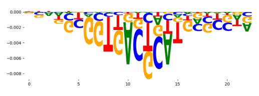 | 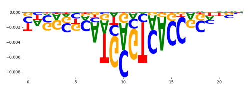 | FOS_MA0476.1 | 3.004900e-02 | 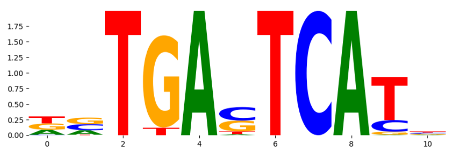 |
| neg_patterns.pattern_7 | 21 | 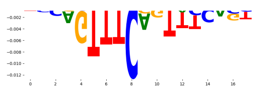 | 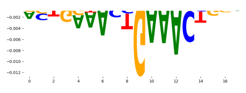 | STAT1+STAT2_MA0517.1 | 2.546030e-05 | 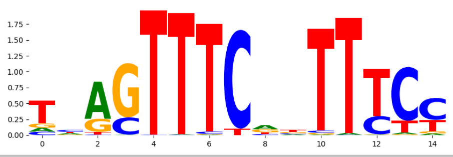 |
