## Supplementary Files 2 for "ChromBPNet: bias factorized, base-resolution deep learning models of chromatin accessibility reveal cis-regulatory sequence syntax, transcription factor footprints and regulatory variants": fig2c_gm12878_DNASE_hint_corrected_bpnet_counts_modisco.pdf

| pattern | num_seqlets | cwm_fwd | cwm_rev | TOMTOM_match | TOMTOM_qval | TOMTOM_match_logo |
|---|---|---|---|---|---|---|
| pos_patterns.pattern_0 | 4738 | | | IRF1_MOUSE.H11MO.0.A | 9.414170e-04 | |
| pos_patterns.pattern_1 | 3534 | | | ETV4_MOUSE.H11MO.0.B | 3.151370e-03 | |
| pos_patterns.pattern_2 | 2787 | | | FOS+JUND_MA1141.1 | 1.577420e-04 | |
| pos_patterns.pattern_3 | 2637 | | | RUNX3_HUMAN.H11MO.0.A | 1.304710e-03 | |
| pos_patterns.pattern_4 | 2212 | | | TF65_MOUSE.H11MO.0.A | 3.804810e-08 | |
| pos_patterns.pattern_5 | 2139 | | | CTCF_MA0139.1 | 2.293180e-13 | |
| pos_patterns.pattern_6 | 1991 | | | ELK4_HUMAN.H11MO.0.A | 3.103140e-04 | |
| pos_patterns.pattern_7 | 1796 | | | KLF12_HUMAN.H11MO.0.C | 1.251460e-05 | |
| pos_patterns.pattern_8 | 1426 | | | IRF4_HUMAN.H11MO.0.A | 6.666610e-06 | |
| pos_patterns.pattern_9 | 794 | | | NFYB_HUMAN.H11MO.0.A | 3.717810e-03 | |
| pos_patterns.pattern_10 | 780 | | | ATF3_HUMAN.H11MO.0.A | 5.651600e-03 | |
| pos_patterns.pattern_11 | 739 | | | NRF1_HUMAN.H11MO.0.A | 2.481590e-06 | |
| pos_patterns.pattern_12 | 530 | | | COE1_MOUSE.H11MO.0.A | 8.134510e-07 | |
| pos_patterns.pattern_13 | 507 | | | TF65_HUMAN.H11MO.0.A | 1.923230e-01 | |
| pos_patterns.pattern_14 | 484 | | | POU5F1_MA1115.1 | 3.022450e-03 | |
| pos_patterns.pattern_15 | 295 | | | ZN143_MOUSE.H11MO.0.A | 5.551070e-16 | |
| pos_patterns.pattern_16 | 211 | | | MEF2D_HUMAN.H11MO.0.A | 9.715190e-09 | |
| pos_patterns.pattern_17 | 205 | | | TYY1_HUMAN.H11MO.0.A | 2.217760e-05 | |
| pos_patterns.pattern_18 | 180 | | | SPIB_ETS_1 | 9.319850e-03 | |
| pos_patterns.pattern_19 | 173 | | | ZBTB33_MA0527.1 | 1.252540e-03 | |
| pos_patterns.pattern_20 | 121 | | | PAX2_PAX_1 | 1.150330e-07 | |
| pos_patterns.pattern_21 | 100 | | | PAX1_MA0779.1 | 2.645540e-02 | |
| pos_patterns.pattern_22 | 94 | | | PAX2_PAX_1 | 5.707590e-14 | |
| pos_patterns.pattern_23 | 84 | | | ETV2_HUMAN.H11MO.0.B | 3.544090e-01 | |
| pos_patterns.pattern_24 | 83 | | | HNF1A_HUMAN.H11MO.0.C | 1.495850e-05 | |
| pos_patterns.pattern_25 | 57 | | | RFX2_MOUSE.H11MO.0.A | 1.580220e-11 | |
| pos_patterns.pattern_26 | 45 | | | NFIA_HUMAN.H11MO.0.C | 1.386330e-04 | |
| pos_patterns.pattern_27 | 40 | | | PAX5_MOUSE.H11MO.0.A | 1.032360e-04 | |
| pos_patterns.pattern_28 | 39 | | | MEF2D_HUMAN.H11MO.0.A | 7.279740e-03 | |

| pattern | num_seqlets | cwm_fwd | cwm_rev | TOMTOM_match | TOMTOM_qval | TOMTOM_match_logo |
|---|---|---|---|---|---|---|
| pos_patterns.pattern_29 | 34 | 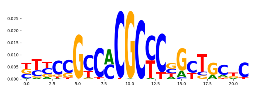 | 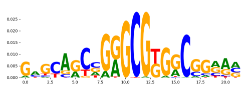 | KLF12_HUMAN.H11MO.0.C | 4.235060e-04 | 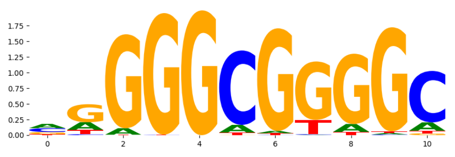 |
| pos_patterns.pattern_30 | 32 | 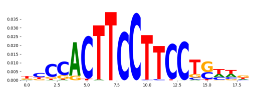 | 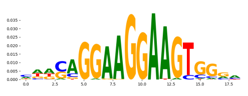 | EWSR1-FLI1_MA0149.1 | 7.417030e-02 | 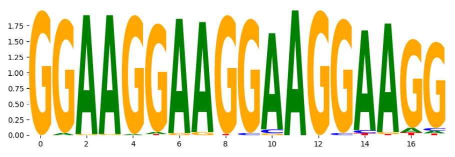 |
| pos_patterns.pattern_31 | 32 | 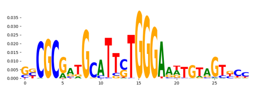 | 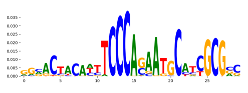 | ZNF76_HUMAN.H11MO.0.C | 4.371540e-09 | 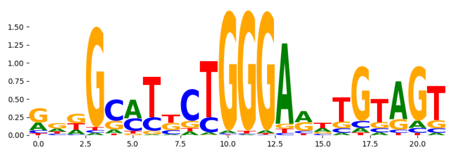 |
| pos_patterns.pattern_32 | 29 | 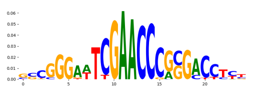 | 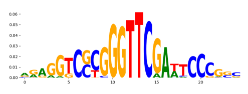 | ZNF524_C2H2_2 | 7.440270e-01 | 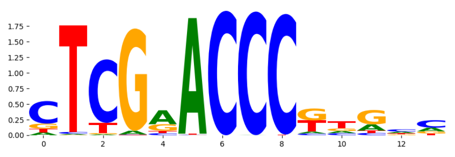 |
| pos_patterns.pattern_33 | 21 | 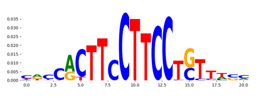 | 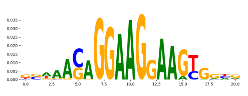 | ETV6_MOUSE.H11MO.0.C | 2.074270e-01 | 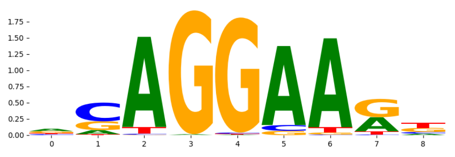 |
| neg_patterns.pattern_0 | 212 | 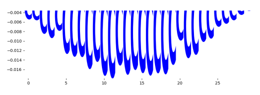 | 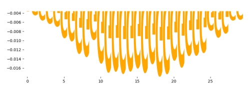 | SP5_MOUSE.H11MO.0.C | 2.232230e-07 | 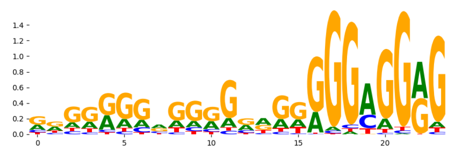 |
| neg_patterns.pattern_1 | 25 | 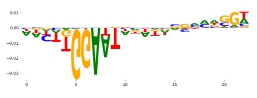 | 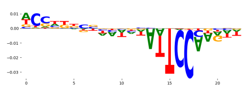 | HXB7_HUMAN.H11MO.0.C | 7.349900e-01 | 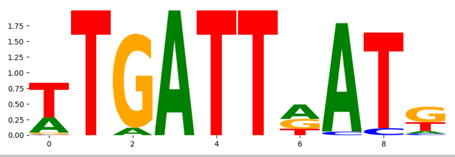 |
