## Supplementary Files 2 for "ChromBPNet: bias factorized, base-resolution deep learning models of chromatin accessibility reveal cis-regulatory sequence syntax, transcription factor footprints and regulatory variants": fig2d_gm12878_DNASE_hint_corrected_bpnet_profile_modisco.pdf

| pattern | num_seqlets | cwm_fwd | cwm_rev | TOMTOM_match | TOMTOM_qval | TOMTOM_match_logo |
|---|---|---|---|---|---|---|
| pos_patterns.pattern_0 | 4122 | | | CTCF_MA0139.1 | 3.109480e-16 | |
| pos_patterns.pattern_1 | 3559 | | | IRF1_MOUSE.H11MO.0.A | 4.111150e-04 | |
| pos_patterns.pattern_2 | 3414 | | | ATF3_MOUSE.H11MO.0.A | 3.275040e-03 | |
| pos_patterns.pattern_3 | 3216 | | | ELF5_HUMAN.H11MO.0.A | 7.345470e-05 | |
| pos_patterns.pattern_4 | 2265 | | | RUNX3_HUMAN.H11MO.0.A | 1.645520e-03 | |
| pos_patterns.pattern_5 | 1064 | | | NFKB1_HUMAN.H11MO.1.B | 3.300750e-08 | |
| pos_patterns.pattern_6 | 702 | | | CTCF_MA0139.1 | 6.732770e-06 | |
| pos_patterns.pattern_7 | 644 | | | IRF4_MOUSE.H11MO.0.A | 3.099870e-04 | |
| pos_patterns.pattern_8 | 594 | | | NFYC_HUMAN.H11MO.0.A | 7.145140e-06 | |
| pos_patterns.pattern_9 | 393 | | | KLF12_HUMAN.H11MO.0.C | 1.394290e-04 | |
| pos_patterns.pattern_10 | 359 | | | BATF_HUMAN.H11MO.0.A | 1.063880e-05 | |
| pos_patterns.pattern_11 | 353 | | | NRF1_HUMAN.H11MO.0.A | 1.034210e-06 | |
| pos_patterns.pattern_12 | 326 | | | MAZ_HUMAN.H11MO.0.A | 2.388140e-07 | |
| pos_patterns.pattern_13 | 260 | | | ZNF76_HUMAN.H11MO.0.C | 1.329620e-16 | |
| pos_patterns.pattern_14 | 220 | | | POU5F1_MA1115.1 | 4.623340e-03 | |
| pos_patterns.pattern_15 | 202 | | | ARNTL_bHLH_1 | 2.866610e-04 | |
| pos_patterns.pattern_16 | 198 | | | RREB1_MA0073.1 | 2.131110e-03 | |
| pos_patterns.pattern_17 | 169 | | | FOSB+JUN_MA1127.1 | 1.237610e-04 | |
| pos_patterns.pattern_18 | 160 | | | COE1_HUMAN.H11MO.0.A | 9.799740e-07 | |
| pos_patterns.pattern_19 | 158 | | | PAX6_PAX_1 | 3.842930e-04 | |
| pos_patterns.pattern_20 | 123 | | | RFX2_MA0600.2 | 6.283760e-09 | |
| pos_patterns.pattern_21 | 69 | | | ETV6_ETS_1 | 1.059990e-01 | |
| pos_patterns.pattern_22 | 48 | | | PAX2_PAX_1 | 5.927890e-05 | |
| pos_patterns.pattern_23 | 39 | | | Pax6_MA0069.1 | 1.374240e-01 | |
| pos_patterns.pattern_24 | 28 | | | HNF1B_HUMAN.H11MO.0.A | 9.419460e-06 | |
