## Supplementary Files 2 for "ChromBPNet: bias factorized, base-resolution deep learning models of chromatin accessibility reveal cis-regulatory sequence syntax, transcription factor footprints and regulatory variants": fig2e_gm12878_DNASE_raw_bpnet_w_simple_cnn_bias_counts_modisco.pdf

| pattern | num_seqlets | cwm_fwd | cwm_rev | TOMTOM_match | TOMTOM_qval | TOMTOM_match_logo |
|---|---|---|---|---|---|---|
| pos_patterns.pattern_0 | 5233 | | | ELF5_HUMAN.H11MO.0.A | 6.305480e-05 | |
| pos_patterns.pattern_1 | 4788 | | | STAT2_HUMAN.H11MO.0.A | 3.986260e-03 | |
| pos_patterns.pattern_2 | 3148 | | | CTCF_MA0139.1 | 1.280350e-10 | |
| pos_patterns.pattern_3 | 3127 | | | RUNX1_HUMAN.H11MO.0.A | 5.697370e-04 | |
| pos_patterns.pattern_4 | 2375 | | | ATF3_MOUSE.H11MO.0.A | 3.701810e-03 | |
| pos_patterns.pattern_5 | 1886 | | | IRF4_HUMAN.H11MO.0.A | 2.849450e-07 | |
| pos_patterns.pattern_6 | 1098 | | | RELB_HUMAN.H11MO.0.C | 2.919520e-05 | |
| pos_patterns.pattern_7 | 683 | | | NFKB2_HUMAN.H11MO.0.B | 8.284950e-04 | |
| pos_patterns.pattern_8 | 627 | | | NRF1_MA0506.1 | 7.182100e-06 | |
| pos_patterns.pattern_9 | 602 | | | KLF12_HUMAN.H11MO.0.C | 8.946070e-05 | |
| pos_patterns.pattern_10 | 582 | | | ATF3_HUMAN.H11MO.0.A | 7.083360e-04 | |
| pos_patterns.pattern_11 | 459 | | | NFYB_HUMAN.H11MO.0.A | 5.790940e-03 | |
| pos_patterns.pattern_12 | 386 | | | ZN143_MOUSE.H11MO.0.A | 4.892370e-19 | |
| pos_patterns.pattern_13 | 362 | | | POU5F1_MA1115.1 | 3.450920e-03 | |
| pos_patterns.pattern_14 | 147 | | | COE1_HUMAN.H11MO.0.A | 1.096840e-04 | |
| pos_patterns.pattern_15 | 83 | | | ZBTB33_MA0527.1 | 2.438900e-04 | |
| pos_patterns.pattern_16 | 38 | | | RUNX3_RUNX_1 | 1.218610e-02 | |
| pos_patterns.pattern_17 | 37 | | | MEF2D_MOUSE.H11MO.0.A | 5.839710e-05 | |
| pos_patterns.pattern_18 | 27 | | | RFX2_HUMAN.H11MO.0.A | 2.804740e-09 | |
| pos_patterns.pattern_19 | 23 | | | Gabpa_MA0062.2 | 6.330340e-01 | |
| pos_patterns.pattern_20 | 20 | | | RUNX3_HUMAN.H11MO.0.A | 3.905670e-01 | |
| pos_patterns.pattern_21 | 20 | | | RELA_MA0107.1 | 2.682180e-01 | |
