## Supplementary Files 2 for "ChromBPNet: bias factorized, base-resolution deep learning models of chromatin accessibility reveal cis-regulatory sequence syntax, transcription factor footprints and regulatory variants": fig2f_gm12878_DNASE_raw_bpnet_w_simple_cnn_bias_profile_modisco.pdf

| pattern | num_seqlets | cwm_fwd | cwm_rev | TOMTOM_match | TOMTOM_qval | TOMTOM_match_logo |
|---|---|---|---|---|---|---|
| pos_patterns.pattern_0 | 4101 | | | CTCF_MA0139.1 | 2.287820e-12 | |
| pos_patterns.pattern_1 | 3241 | | | ELF5_HUMAN.H11MO.0.A | 4.308260e-05 | |
| pos_patterns.pattern_2 | 3222 | | | RUNX1_HUMAN.H11MO.0.A | 8.906750e-02 | |
| pos_patterns.pattern_3 | 2513 | | | JDP2_MA0655.1 | 3.125490e-03 | |
| pos_patterns.pattern_4 | 1469 | | | IRF5_IRF_2 | 2.736630e-01 | |
| pos_patterns.pattern_5 | 1241 | | | IRF4_HUMAN.H11MO.0.A | 1.934230e-09 | |
| pos_patterns.pattern_6 | 847 | | | IRF1_MOUSE.H11MO.0.A | 7.514000e-06 | |
| pos_patterns.pattern_7 | 827 | | | NFKB2_HUMAN.H11MO.0.B | 3.858840e-03 | |
| pos_patterns.pattern_8 | 643 | | | NFYB_HUMAN.H11MO.0.A | 4.380140e-08 | |
| pos_patterns.pattern_9 | 612 | | | NaN | NaN | |
| pos_patterns.pattern_10 | 537 | | | KLF12_HUMAN.H11MO.0.C | 3.008810e-05 | |
| pos_patterns.pattern_11 | 528 | | | NRF1_HUMAN.H11MO.0.A | 1.700930e-06 | |
| pos_patterns.pattern_12 | 471 | | | JUN+JUNB_MA1133.1 | 8.663770e-05 | |
| pos_patterns.pattern_13 | 462 | | | RARA_nuclearreceptor_5 | 1.249140e-02 | |
| pos_patterns.pattern_14 | 447 | | | STAT2_HUMAN.H11MO.0.A | 3.660880e-01 | |
| pos_patterns.pattern_15 | 444 | | | MITF_MA0620.2 | 2.793130e-05 | |
| pos_patterns.pattern_16 | 420 | | | PO2F2_HUMAN.H11MO.0.A | 1.096130e-03 | |
| pos_patterns.pattern_17 | 256 | | | ZNF76_HUMAN.H11MO.0.C | 1.962140e-15 | |
| pos_patterns.pattern_18 | 218 | | | SP5_MOUSE.H11MO.0.C | 2.378940e-07 | |
| pos_patterns.pattern_19 | 202 | | | SP1_C2H2_1 | 2.616200e-03 | |
| pos_patterns.pattern_20 | 195 | | | COE1_MOUSE.H11MO.0.A | 1.109220e-06 | |
| pos_patterns.pattern_21 | 93 | | | RFX2_HUMAN.H11MO.0.A | 7.245930e-08 | |
| pos_patterns.pattern_22 | 91 | | | ZBTB33_MA0527.1 | 8.818680e-05 | |
| pos_patterns.pattern_23 | 66 | | | NFIC_HUMAN.H11MO.0.A | 9.682840e-02 | |
| pos_patterns.pattern_24 | 53 | | | IRF1_HUMAN.H11MO.0.A | 3.473330e-05 | |
| pos_patterns.pattern_25 | 36 | | | ZN770_HUMAN.H11MO.0.C | 1.595400e-02 | |
| pos_patterns.pattern_26 | 21 | | | SOX9_HMG_3 | 1.000000e+00 | |
| neg_patterns.pattern_0 | 136 | | | NFATC2_MA0152.1 | 9.297200e-01 | |
| neg_patterns.pattern_1 | 92 | | | DNASE_2 | 7.579940e-03 | |
| neg_patterns.pattern_2 | 30 | | | IRF4_MOUSE.H11MO.0.A | 1.192910e-03 | |

| pattern | num_seqlets | cwm_fwd | cwm_rev | TOMTOM_match | TOMTOM_qval | TOMTOM_match_logo |
|---|---|---|---|---|---|---|
| neg_patterns.pattern_3 | 29 | 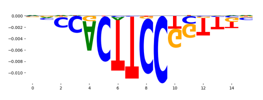 | 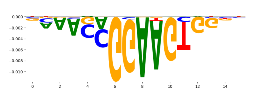 | ETS1_MOUSE.H11MO.0.A | 2.164150e-05 | 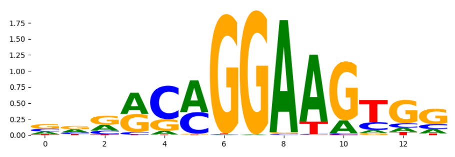 |
| neg_patterns.pattern_4 | 29 | 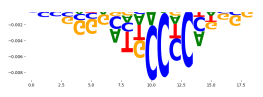 | 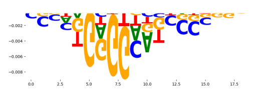 | REL_MA0101.1 | 3.466780e-03 | 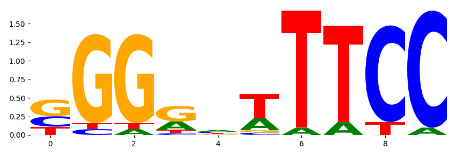 |
| neg_patterns.pattern_5 | 27 | 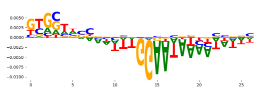 | 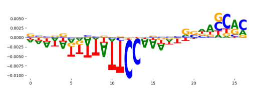 | NFATC2_MA0152.1 | 5.477360e-01 | 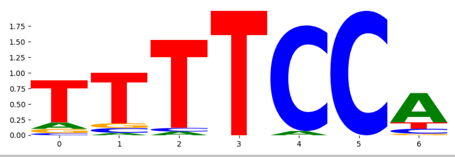 |
