## Supplementary Files 2 for "ChromBPNet: bias factorized, base-resolution deep learning models of chromatin accessibility reveal cis-regulatory sequence syntax, transcription factor footprints and regulatory variants": fig2g_gm12878_DNASE_raw_chrombpnet_profile_modisco.pdf

| pattern | num_seqlets | cwm_fwd | cwm_rev | TOMTOM_match | TOMTOM_qval | TOMTOM_match_logo |
|---|---|---|---|---|---|---|
| pos_patterns.pattern_0 | 5066 | | | ELF5_HUMAN.H11MO.0.A | 1.428730e-05 | |
| pos_patterns.pattern_1 | 3975 | | | CTCF_MA0139.1 | 1.062330e-12 | |
| pos_patterns.pattern_2 | 2871 | | | FOS_HUMAN.H11MO.0.A | 1.183150e-03 | |
| pos_patterns.pattern_3 | 2799 | | | STAT2_HUMAN.H11MO.0.A | 7.865400e-03 | |
| pos_patterns.pattern_4 | 2576 | | | RUNX1_HUMAN.H11MO.0.A | 1.778140e-02 | |
| pos_patterns.pattern_5 | 2534 | | | IRF4_MOUSE.H11MO.0.A | 3.367440e-05 | |
| pos_patterns.pattern_6 | 2333 | | | NFKB1_HUMAN.H11MO.1.B | 2.431960e-08 | |
| pos_patterns.pattern_7 | 1064 | | | KLF12_HUMAN.H11MO.0.C | 1.081220e-05 | |
| pos_patterns.pattern_8 | 762 | | | NRF1_MA0506.1 | 1.646800e-06 | |
| pos_patterns.pattern_9 | 548 | | | Pou2f2.mouse_POU_2 | 5.314080e-03 | |
| pos_patterns.pattern_10 | 497 | | | FOSB+JUNB_MA1136.1 | 4.589460e-05 | |
| pos_patterns.pattern_11 | 466 | | | NFYB_HUMAN.H11MO.0.A | 1.320010e-03 | |
| pos_patterns.pattern_12 | 465 | | | COE1_MOUSE.H11MO.0.A | 1.073500e-06 | |
| pos_patterns.pattern_13 | 420 | | | RELB_HUMAN.H11MO.0.C | 6.833930e-01 | |
| pos_patterns.pattern_14 | 339 | | | ZN143_MOUSE.H11MO.0.A | 5.872660e-15 | |
| pos_patterns.pattern_15 | 314 | | | ETS1_HUMAN.H11MO.0.A | 2.869920e-02 | |
| pos_patterns.pattern_16 | 216 | | | ARNTL_bHLH_1 | 4.374330e-05 | |
| pos_patterns.pattern_17 | 149 | | | SPIB_ETS_1 | 4.166820e-02 | |
| pos_patterns.pattern_18 | 132 | | | PAX9_MA0781.1 | 6.841950e-08 | |
| pos_patterns.pattern_19 | 107 | | | MEF2D_HUMAN.H11MO.0.A | 1.326780e-04 | |
| pos_patterns.pattern_20 | 107 | | | ZBTB33_MA0527.1 | 5.588200e-06 | |
| pos_patterns.pattern_21 | 101 | | | HNF1B_MOUSE.H11MO.0.A | 1.047060e-05 | |
| pos_patterns.pattern_22 | 56 | | | TYY1_HUMAN.H11MO.0.A | 1.989540e-05 | |
| pos_patterns.pattern_23 | 43 | | | BATF+JUN_MA0462.1 | 1.396300e-03 | |
| pos_patterns.pattern_24 | 38 | | | PAX6_PAX_1 | 1.729180e-02 | |
| pos_patterns.pattern_25 | 27 | | | ETV6_ETS_1 | 1.637700e-04 | |
| pos_patterns.pattern_26 | 25 | | | Rfx1_MA0509.1 | 9.727740e-09 | |
| pos_patterns.pattern_27 | 21 | | | ZNF524_C2H2_2 | 1.000000e+00 | |
