## Supplementary Files 2 for "ChromBPNet: bias factorized, base-resolution deep learning models of chromatin accessibility reveal cis-regulatory sequence syntax, transcription factor footprints and regulatory variants": fig2h_gm12878_DNASE_raw_chrombpnet_profile_modisco.pdf

| pattern | num_seqlets | cwm_fwd | cwm_rev | TOMTOM_match | TOMTOM_qval | TOMTOM_match_logo |
|---|---|---|---|---|---|---|
| pos_patterns.pattern_0 | 5225 | | | CTCF_MA0139.1 | 3.548440e-14 | |
| pos_patterns.pattern_1 | 4768 | | | ELF5_HUMAN.H11MO.0.A | 3.100350e-06 | |
| pos_patterns.pattern_2 | 3569 | | | ATF3_MOUSE.H11MO.0.A | 3.449910e-03 | |
| pos_patterns.pattern_3 | 3408 | | | RUNX1_HUMAN.H11MO.0.A | 2.641150e-03 | |
| pos_patterns.pattern_4 | 2744 | | | IRF4_MOUSE.H11MO.0.A | 1.880380e-06 | |
| pos_patterns.pattern_5 | 2439 | | | IRF8_IRF_1 | 1.441590e-01 | |
| pos_patterns.pattern_6 | 2136 | | | KLF12_HUMAN.H11MO.0.C | 1.086750e-04 | |
| pos_patterns.pattern_7 | 2042 | | | NFKB1_HUMAN.H11MO.1.B | 3.254690e-07 | |
| pos_patterns.pattern_8 | 1089 | | | NFYB_HUMAN.H11MO.0.A | 7.289610e-04 | |
| pos_patterns.pattern_9 | 849 | | | NRF1_MA0506.1 | 1.822270e-06 | |
| pos_patterns.pattern_10 | 799 | | | FOSB+JUNB_MA1136.1 | 5.258220e-05 | |
| pos_patterns.pattern_11 | 592 | | | COE1_MOUSE.H11MO.0.A | 4.403920e-06 | |
| pos_patterns.pattern_12 | 534 | | | Pou2f2.mouse_POU_2 | 4.226150e-03 | |
| pos_patterns.pattern_13 | 504 | | | USF2_MA0526.2 | 4.436500e-06 | |
| pos_patterns.pattern_14 | 465 | | | EGR2_HUMAN.H11MO.0.A | 4.591040e-04 | |
| pos_patterns.pattern_15 | 374 | | | SP5_MOUSE.H11MO.0.C | 2.360100e-07 | |
| pos_patterns.pattern_16 | 364 | | | PAX1_MA0779.1 | 2.336640e-04 | |
| pos_patterns.pattern_17 | 354 | | | ZN143_MOUSE.H11MO.0.A | 1.300800e-13 | |
| pos_patterns.pattern_18 | 222 | | | Rfx1_MA0509.1 | 3.656680e-09 | |
| pos_patterns.pattern_19 | 105 | | | NFIC_HUMAN.H11MO.0.A | 2.077250e-03 | |
| pos_patterns.pattern_20 | 97 | | | Gabpa_MA0062.2 | 1.498330e-01 | |
| pos_patterns.pattern_21 | 89 | | | HNF1B_HUMAN.H11MO.0.A | 1.030430e-05 | |
| pos_patterns.pattern_22 | 89 | | | ZBTB33_MA0527.1 | 1.767420e-05 | |
| pos_patterns.pattern_23 | 81 | | | ZNF384_MA1125.1 | 4.805280e-02 | |
| pos_patterns.pattern_24 | 69 | | | SPIB_ETS_1 | 4.978250e-02 | |
| pos_patterns.pattern_25 | 43 | | | TYY1_HUMAN.H11MO.0.A | 1.083250e-04 | |
| pos_patterns.pattern_26 | 32 | | | NFKB2_HUMAN.H11MO.0.B | 1.000000e+00 | |
| pos_patterns.pattern_27 | 26 | | | SPIC_ETS_1 | 1.298670e-02 | |
| pos_patterns.pattern_28 | 24 | | | GABPA_HUMAN.H11MO.0.A | 2.609580e-03 | |

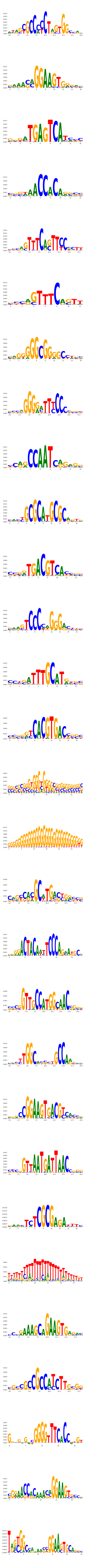

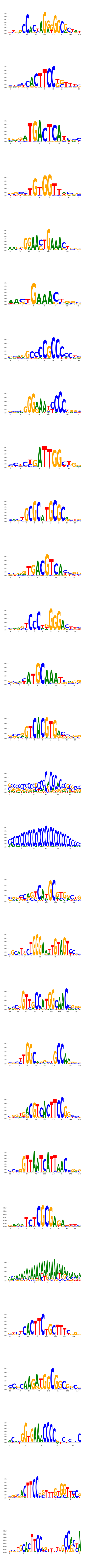

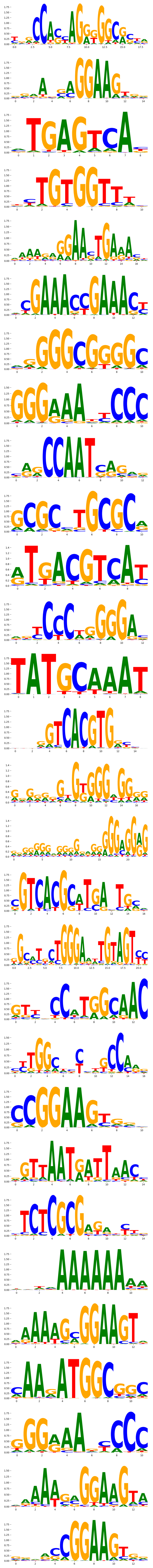

| pattern | num_seqlets | cwm_fwd | cwm_rev | TOMTOM_match | TOMTOM_qval | TOMTOM_match_logo |
|---|---|---|---|---|---|---|
| neg_patterns.pattern_0 | 108 | 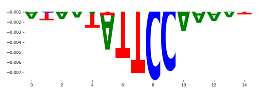 | 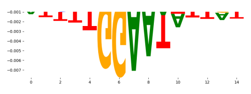 | NFATC2_MA0152.1 | 5.058770e-01 | 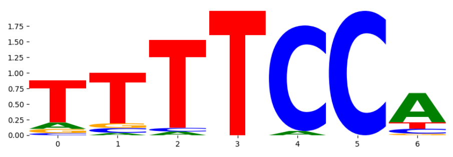 |
| neg_patterns.pattern_1 | 102 | 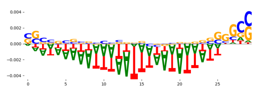 | 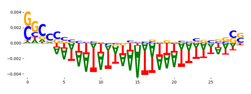 | DNASE_2 | 1.031660e-01 | 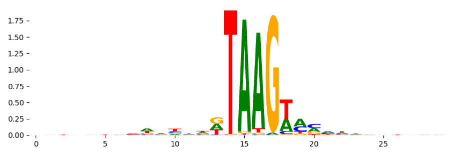 |
| neg_patterns.pattern_2 | 79 | 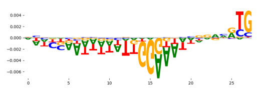 | 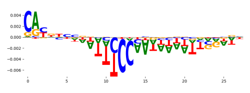 | NFATC2_MA0152.1 | 3.617970e-01 | 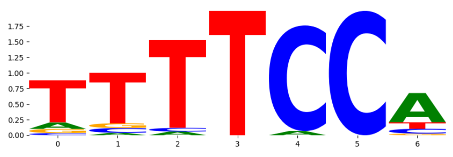 |
| neg_patterns.pattern_3 | 72 | 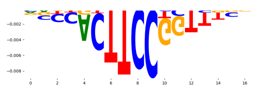 | 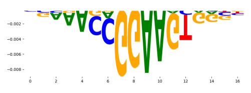 | ERG_MOUSE.H11MO.0.A | 1.909290e-05 | 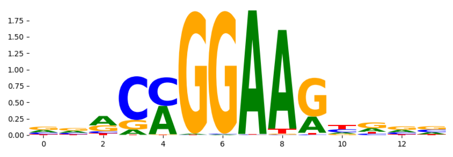 |
| neg_patterns.pattern_4 | 52 | 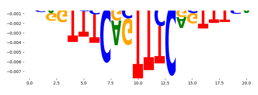 | 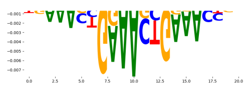 | IRF1_MOUSE.H11MO.0.A | 4.698710e-05 | 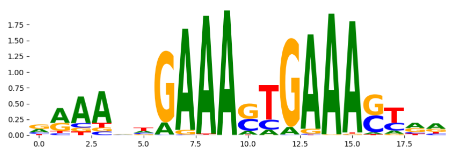 |
| neg_patterns.pattern_5 | 34 | 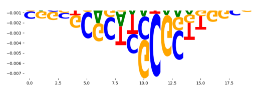 | 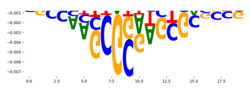 | ETV6_ETS_1 | 6.391250e-02 | 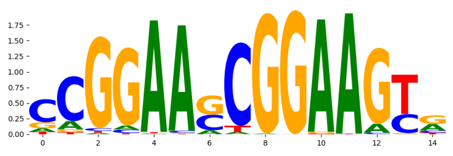 |
| neg_patterns.pattern_6 | 33 | 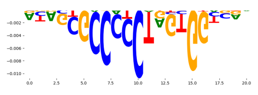 | 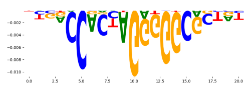 | CTCF_MA0139.1 | 5.470420e-08 | 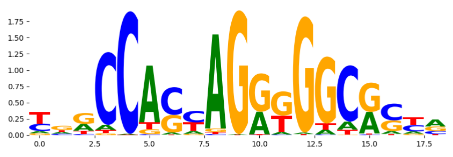 |
| neg_patterns.pattern_7 | 32 | 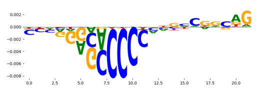 | 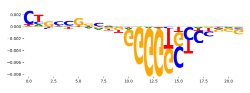 | ZBT7A_HUMAN.H11MO.0.A | 2.069190e-02 | 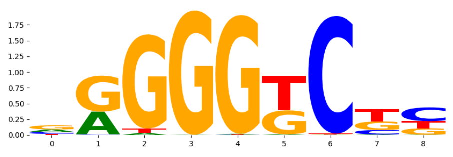 |
| neg_patterns.pattern_8 | 31 | 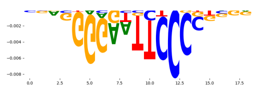 | 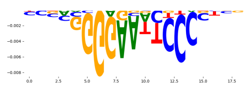 | NFKB1_HUMAN.H11MO.1.B | 3.623150e-03 | 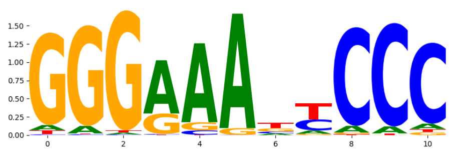 |
| neg_patterns.pattern_9 | 21 | 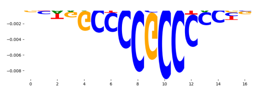 | 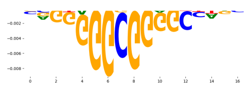 | SP1_HUMAN.H11MO.0.A | 1.952870e-04 | 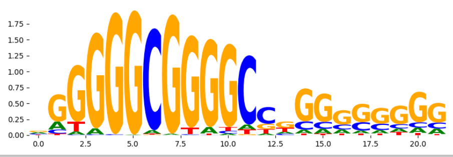 |
