## Supplementary Files 3 for "ChromBPNet: bias factorized, base-resolution deep learning models of chromatin accessibility reveal cis-regulatory sequence syntax, transcription factor footprints and regulatory variants": gm12878_ATAC_raw_bpnet_bias_fold0_counts_modisco.pdf

| pattern | num_seqlets | cwm_fwd | cwm_rev | TOMTOM_match | TOMTOM_qval | TOMTOM_match_logo |
|---|---|---|---|---|---|---|
| pos_patterns.pattern_0 | 3167 | | | ZBT17_HUMAN.H11MO.0.A | 7.240880e-01 | |
| pos_patterns.pattern_1 | 2396 | | | ZFX_MOUSE.H11MO.0.B | 1.578660e-02 | |
| pos_patterns.pattern_2 | 2380 | | | ZN436_HUMAN.H11MO.0.C | 1.253010e-01 | |
| pos_patterns.pattern_3 | 2290 | | | GMEB2_SAND_2 | 1.000000e+00 | |
| pos_patterns.pattern_4 | 2255 | | | SP3_HUMAN.H11MO.0.B | 7.758650e-03 | |
| pos_patterns.pattern_5 | 1933 | | | SOX2_HUMAN.H11MO.0.A | 1.000000e+00 | |
| pos_patterns.pattern_6 | 1552 | | | NFATC2_MA0152.1 | 1.860260e-01 | |
| pos_patterns.pattern_7 | 1502 | | | RUNX2_HUMAN.H11MO.0.A | 1.000000e+00 | |
| pos_patterns.pattern_8 | 1443 | | | RREB1_MA0073.1 | 3.905240e-07 | |
| pos_patterns.pattern_9 | 1270 | | | PAX5_MA0014.3 | 1.000000e+00 | |
| pos_patterns.pattern_10 | 1254 | | | RUNX2_HUMAN.H11MO.0.A | 1.074140e-01 | |
| pos_patterns.pattern_11 | 1119 | | | TFAP4_HUMAN.H11MO.0.A | 4.682400e-02 | |
| pos_patterns.pattern_12 | 1081 | | | STAT1_MOUSE.H11MO.0.A | 1.000000e+00 | |
| pos_patterns.pattern_13 | 1011 | | | PRD16_MOUSE.H11MO.0.B | 1.000000e+00 | |
| pos_patterns.pattern_14 | 821 | | | SP1_HUMAN.H11MO.0.A | 5.556300e-01 | |
| pos_patterns.pattern_15 | 816 | | | NR4A2_nuclearreceptor_1 | 1.000000e+00 | |
| pos_patterns.pattern_16 | 703 | | | ZFX_MOUSE.H11MO.0.B | 6.031190e-01 | |
| pos_patterns.pattern_17 | 679 | | | GCM1_GCM_3 | 1.514020e-01 | |
| pos_patterns.pattern_18 | 595 | | | PBX2_MA1113.1 | 1.000000e+00 | |
| pos_patterns.pattern_19 | 589 | | | KLF5_MOUSE.H11MO.0.A | 1.000000e+00 | |
| pos_patterns.pattern_20 | 564 | | | NFAC2_HUMAN.H11MO.0.B | 4.804290e-01 | |
| pos_patterns.pattern_21 | 516 | | | NFATC2_MA0152.1 | 2.844080e-02 | |
| pos_patterns.pattern_22 | 431 | | | SOX8_HMG_7 | 1.000000e+00 | |
| pos_patterns.pattern_23 | 412 | | | PRD14_HUMAN.H11MO.0.A | 5.033940e-01 | |
| pos_patterns.pattern_24 | 353 | | | ZN140_HUMAN.H11MO.0.C | 1.170690e-01 | |
| pos_patterns.pattern_25 | 324 | | | SOX8_HMG_7 | 3.262770e-01 | |
| pos_patterns.pattern_26 | 309 | | | KLF4_MA0039.3 | 5.791810e-01 | |
| pos_patterns.pattern_27 | 294 | | | KLF5_MOUSE.H11MO.0.A | 8.831600e-01 | |
| pos_patterns.pattern_28 | 303 | | | NFKB2_MA0778.1 | 2.552380e-01 | |

| pattern | num_seqlets | cwm_fwd | cwm_rev | TOMTOM_match | TOMTOM_qval | TOMTOM_match_logo |
|---|---|---|---|---|---|---|
| pos_patterns.pattern_29 | 294 | | | BHA15_HUMAN.H11MO.0.B | 1.000000e+00 | |
| pos_patterns.pattern_30 | 287 | | | NFAC3_HUMAN.H11MO.0.B | 1.000000e+00 | |
| pos_patterns.pattern_31 | 269 | | | ASCL1_MOUSE.H11MO.0.A | 6.632690e-01 | |
| pos_patterns.pattern_32 | 252 | | | SRBP2_HUMAN.H11MO.0.B | 4.771640e-01 | |
| pos_patterns.pattern_33 | 237 | | | COE1_HUMAN.H11MO.0.A | 1.000000e+00 | |
| pos_patterns.pattern_34 | 193 | | | SRY_HMG_2 | 6.872720e-01 | |
| pos_patterns.pattern_35 | 144 | | | ASCL1_MOUSE.H11MO.0.A | 3.599420e-01 | |
| pos_patterns.pattern_36 | 139 | | | AP2A_MOUSE.H11MO.0.A | 6.657940e-01 | |
| pos_patterns.pattern_37 | 132 | | | NaN | NaN | |
| pos_patterns.pattern_38 | 130 | | | AP2C_MOUSE.H11MO.0.A | 3.008310e-01 | |
| pos_patterns.pattern_39 | 112 | | | ZN547_HUMAN.H11MO.0.C | 1.000000e+00 | |
| pos_patterns.pattern_40 | 113 | | | ZN331_HUMAN.H11MO.0.C | 6.761010e-03 | |
| pos_patterns.pattern_41 | 90 | | | FOXO3_forkhead_3 | 5.862900e-01 | |
| pos_patterns.pattern_42 | 84 | | | FOXO3_forkhead_3 | 1.000000e+00 | |
| pos_patterns.pattern_43 | 70 | | | SPDEF_ETS_2 | 1.000000e+00 | |
| pos_patterns.pattern_44 | 71 | | | MYCN_HUMAN.H11MO.0.A | 2.040050e-02 | |
| pos_patterns.pattern_45 | 81 | | | TFAP2B_MA0812.1 | 1.838620e-01 | |
| pos_patterns.pattern_46 | 66 | | | NRF1_MOUSE.H11MO.0.A | 3.170360e-01 | |
| pos_patterns.pattern_47 | 55 | | | SOX4_HMG_1 | 7.582180e-01 | |
| pos_patterns.pattern_48 | 42 | | | FOXO3_forkhead_3 | 8.100940e-01 | |
| pos_patterns.pattern_49 | 42 | | | Hoxa9_MA0594.1 | 1.000000e+00 | |
| pos_patterns.pattern_50 | 42 | | | ZN436_HUMAN.H11MO.0.C | 5.236760e-01 | |
| pos_patterns.pattern_51 | 33 | | | NFAC3_HUMAN.H11MO.0.B | 6.859780e-01 | |
| pos_patterns.pattern_52 | 35 | | | TFAP2C_TFAP_6 | 6.529020e-01 | |
| pos_patterns.pattern_53 | 26 | | | ZNF282_C2H2_1 | 5.043650e-01 | |
| neg_patterns.pattern_0 | 674 | | | TN5_7 | 3.319080e-03 | |
| neg_patterns.pattern_1 | 412 | | | PRDM6_HUMAN.H11MO.0.C | 1.326790e-01 | |
| neg_patterns.pattern_2 | 316 | | | TN5_2 | 1.090470e-01 | |
| neg_patterns.pattern_3 | 281 | | | TN5_2 | 7.529050e-05 | |
| neg_patterns.pattern_4 | 221 | | | SP2_HUMAN.H11MO.0.A | 3.044040e-08 | |

| pattern | num_seqlets | cwm_fwd | cwm_rev | TOMTOM_match | TOMTOM_qval | TOMTOM_match_logo |
|---|---|---|---|---|---|---|
| neg_patterns.pattern_5 | 227 | | | ZN667_HUMAN.H11MO.0.C | 6.987780e-01 | |
| neg_patterns.pattern_6 | 125 | | | SP2_HUMAN.H11MO.0.A | 3.498470e-04 | |
| neg_patterns.pattern_7 | 119 | | | TN5_2 | 1.160430e-04 | |
| neg_patterns.pattern_8 | 78 | | | TN5_2 | 1.601800e-04 | |
| neg_patterns.pattern_9 | 67 | | | SP2_HUMAN.H11MO.0.A | 7.469830e-08 | |
| neg_patterns.pattern_10 | 81 | | | Zic3.mouse_C2H2_1 | 1.000000e+00 | |
| neg_patterns.pattern_11 | 66 | | | SP2_HUMAN.H11MO.0.A | 1.543380e-01 | |
| neg_patterns.pattern_12 | 74 | | | ZNF384_MA1125.1 | 1.467600e-02 | |
| neg_patterns.pattern_13 | 59 | | | PBX3_MA1114.1 | 2.259020e-01 | |
| neg_patterns.pattern_14 | 51 | | | RREB1_MA0073.1 | 1.000000e+00 | |
| neg_patterns.pattern_15 | 49 | | | ELF1_HUMAN.H11MO.0.A | 8.238480e-01 | |
| neg_patterns.pattern_16 | 49 | | | TN5_2 | 2.175870e-04 | |
| neg_patterns.pattern_17 | 38 | | | ZN431_MOUSE.H11MO.0.C | 1.000000e+00 | |
| neg_patterns.pattern_18 | 33 | | | TN5_8 | 3.674490e-02 | |
| neg_patterns.pattern_19 | 40 | | | DNASE_5 | 1.504450e-06 | |
| neg_patterns.pattern_20 | 30 | | | SP1_HUMAN.H11MO.0.A | 2.234150e-05 | |
| neg_patterns.pattern_21 | 33 | | | NR1D1_HUMAN.H11MO.0.B | 9.290610e-02 | |
| neg_patterns.pattern_22 | 41 | | | TBX20_MOUSE.H11MO.0.C | 1.847560e-01 | |
| neg_patterns.pattern_23 | 37 | | | TN5_6 | 2.173160e-12 | |
| neg_patterns.pattern_24 | 25 | | | TP63_MA0525.2 | 1.000000e+00 | |
| neg_patterns.pattern_25 | 40 | | | Foxd3_MA0041.1 | 9.308280e-02 | |
