## Supplementary Files 3 for "ChromBPNet: bias factorized, base-resolution deep learning models of chromatin accessibility reveal cis-regulatory sequence syntax, transcription factor footprints and regulatory variants": gm12878_ATAC_raw_bpnet_bias_fold0_profile_modisco.pdf

| pattern | num_seqlets | cwm_fwd | cwm_rev | TOMTOM_match | TOMTOM_qval | TOMTOM_match_logo |
|---|---|---|---|---|---|---|
| pos_patterns.pattern_0 | 16571 | | | TN5_1 | 7.273770e-06 | |
| pos_patterns.pattern_1 | 4565 | | | TN5_2 | 2.013520e-20 | |
| pos_patterns.pattern_2 | 3604 | | | TN5_1 | 3.957760e-06 | |
| pos_patterns.pattern_3 | 3200 | | | TN5_1 | 1.591750e-06 | |
| pos_patterns.pattern_4 | 3057 | | | TN5_4 | 6.834380e-09 | |
| pos_patterns.pattern_5 | 2439 | | | TN5_7 | 5.285130e-07 | |
| pos_patterns.pattern_6 | 2336 | | | TN5_3 | 2.476830e-04 | |
| pos_patterns.pattern_7 | 2300 | | | TN5_3 | 2.384440e-17 | |
| pos_patterns.pattern_8 | 1444 | | | TN5_3 | 9.988090e-04 | |
| pos_patterns.pattern_9 | 1046 | | | TN5_6 | 7.668230e-14 | |
| pos_patterns.pattern_10 | 1041 | | | TN5_8 | 2.961390e-19 | |
| pos_patterns.pattern_11 | 938 | | | TN5_8 | 7.516760e-06 | |
| pos_patterns.pattern_12 | 479 | | | ZNF384_MA1125.1 | 4.203370e-02 | |
| pos_patterns.pattern_13 | 280 | | | TCF4_bHLH_2 | 8.339860e-01 | |
| pos_patterns.pattern_14 | 257 | | | TCF4_bHLH_2 | 4.585190e-01 | |
| pos_patterns.pattern_15 | 185 | | | TN5_4 | 4.439380e-01 | |
| pos_patterns.pattern_16 | 165 | | | EGR2_HUMAN.H11MO.0.A | 1.000000e+00 | |
| pos_patterns.pattern_17 | 97 | | | OVOL1_HUMAN.H11MO.0.C | 1.000000e+00 | |
| pos_patterns.pattern_18 | 49 | | | TN5_6 | 1.168370e-08 | |
| pos_patterns.pattern_19 | 37 | | | ZNF18_HUMAN.H11MO.0.C | 4.381050e-01 | |
| pos_patterns.pattern_20 | 39 | | | ZN549_HUMAN.H11MO.0.C | 1.000000e+00 | |
