## Supplementary Files 3 for "ChromBPNet: bias factorized, base-resolution deep learning models of chromatin accessibility reveal cis-regulatory sequence syntax, transcription factor footprints and regulatory variants": gm12878_ATAC_raw_bpnet_bias_fold1_counts_modisco.pdf

| pattern | num_seqlets | cwm_fwd | cwm_rev | TOMTOM_match | TOMTOM_qval | TOMTOM_match_logo |
|---|---|---|---|---|---|---|
| pos_patterns.pattern_0 | 3468 | | | SP2_HUMAN.H11MO.0.A | 9.197300e-03 | |
| pos_patterns.pattern_1 | 2224 | | | ZN384_HUMAN.H11MO.0.C | 1.000000e+00 | |
| pos_patterns.pattern_2 | 807 | | | RREB1_MA0073.1 | 7.863040e-03 | |
| pos_patterns.pattern_3 | 802 | | | RHOXF1_homeodomain_1 | 1.000000e+00 | |
| pos_patterns.pattern_4 | 757 | | | RORA_nuclearreceptor_1 | 1.000000e+00 | |
| pos_patterns.pattern_5 | 739 | | | RELB_HUMAN.H11MO.0.C | 2.754320e-01 | |
| pos_patterns.pattern_6 | 699 | | | TBX20_MA0689.1 | 4.031450e-01 | |
| pos_patterns.pattern_7 | 684 | | | RXRG_nuclearreceptor_4 | 9.522160e-01 | |
| pos_patterns.pattern_8 | 633 | | | STAT2_HUMAN.H11MO.0.A | 8.953190e-01 | |
| pos_patterns.pattern_9 | 567 | | | THB_HUMAN.H11MO.0.C | 1.082970e-01 | |
| pos_patterns.pattern_10 | 563 | | | RREB1_MA0073.1 | 2.341980e-02 | |
| pos_patterns.pattern_11 | 547 | | | BHA15_HUMAN.H11MO.0.B | 9.723000e-01 | |
| pos_patterns.pattern_12 | 540 | | | ZBTB6_HUMAN.H11MO.0.C | 1.000000e+00 | |
| pos_patterns.pattern_13 | 516 | | | Ddit3+Cebpa_MA0019.1 | 1.000000e+00 | |
| pos_patterns.pattern_14 | 516 | | | Nr5a2_MA0505.1 | 4.599300e-01 | |
| pos_patterns.pattern_15 | 505 | | | ZBTB7A_C2H2_1 | 8.056770e-01 | |
| pos_patterns.pattern_16 | 490 | | | ZNF384_MA1125.1 | 6.929320e-02 | |
| pos_patterns.pattern_17 | 446 | | | RREB1_MA0073.1 | 2.350810e-02 | |
| pos_patterns.pattern_18 | 430 | | | WT1_HUMAN.H11MO.0.C | 6.992450e-02 | |
| pos_patterns.pattern_19 | 428 | | | RREB1_MA0073.1 | 9.972970e-01 | |
| pos_patterns.pattern_20 | 407 | | | SP1_HUMAN.H11MO.0.A | 4.917680e-01 | |
| pos_patterns.pattern_21 | 343 | | | ID4_MA0824.1 | 9.237330e-01 | |
| pos_patterns.pattern_22 | 331 | | | ASCL1_MA1100.1 | 8.585940e-01 | |
| pos_patterns.pattern_23 | 261 | | | ERR1_MOUSE.H11MO.0.A | 1.000000e+00 | |
| pos_patterns.pattern_24 | 238 | | | ZN250_HUMAN.H11MO.0.C | 1.000000e+00 | |
| pos_patterns.pattern_25 | 233 | | | RREB1_MA0073.1 | 9.153480e-05 | |
| pos_patterns.pattern_26 | 193 | | | ZNF384_MA1125.1 | 2.969120e-02 | |
| pos_patterns.pattern_27 | 143 | | | ZBTB6_HUMAN.H11MO.0.C | 2.469610e-01 | |
| pos_patterns.pattern_28 | 85 | | | IRF8_IRF_1 | 8.774610e-01 | |

| pattern | num_seqlets | cwm_fwd | cwm_rev | TOMTOM_match | TOMTOM_qval | TOMTOM_match_logo |
|---|---|---|---|---|---|---|
| pos_patterns.pattern_29 | 84 | | | Nr5a2_MA0505.1 | 1.000000e+00 | |
| pos_patterns.pattern_30 | 83 | | | RREB1_MA0073.1 | 5.745850e-01 | |
| pos_patterns.pattern_31 | 56 | | | SP3_HUMAN.H11MO.0.B | 4.332280e-01 | |
| pos_patterns.pattern_32 | 50 | | | HXA1_HUMAN.H11MO.0.C | 4.626460e-01 | |
| pos_patterns.pattern_33 | 49 | | | RREB1_MA0073.1 | 8.210420e-01 | |
| pos_patterns.pattern_34 | 41 | | | ZNF282_C2H2_1 | 1.000000e+00 | |
| pos_patterns.pattern_35 | 36 | | | SP2_HUMAN.H11MO.0.A | 3.423870e-01 | |
| pos_patterns.pattern_36 | 32 | | | NHLH1_bHLH_1 | 1.000000e+00 | |
| pos_patterns.pattern_37 | 28 | | | TBX20_TBX_5 | 1.000000e+00 | |
| neg_patterns.pattern_0 | 184 | | | SP2_HUMAN.H11MO.0.A | 1.106890e-04 | |
| neg_patterns.pattern_1 | 158 | | | ZN331_HUMAN.H11MO.0.C | 8.456850e-03 | |
| neg_patterns.pattern_2 | 156 | | | SP2_HUMAN.H11MO.0.A | 7.873710e-05 | |
| neg_patterns.pattern_3 | 150 | | | TN5_1 | 5.538670e-04 | |
| neg_patterns.pattern_4 | 131 | | | RREB1_MA0073.1 | 1.000000e+00 | |
| neg_patterns.pattern_5 | 116 | | | NR4A2_nuclearreceptor_3 | 2.080070e-01 | |
| neg_patterns.pattern_6 | 104 | | | ZN341_HUMAN.H11MO.0.C | 6.459850e-02 | |
| neg_patterns.pattern_7 | 104 | | | ZN667_HUMAN.H11MO.0.C | 1.445040e-01 | |
| neg_patterns.pattern_8 | 102 | | | ZFX_MOUSE.H11MO.0.B | 7.238720e-04 | |
| neg_patterns.pattern_9 | 93 | | | PRDM1_MA0508.2 | 1.334930e-01 | |
| neg_patterns.pattern_10 | 88 | | | SP2_HUMAN.H11MO.0.A | 2.679590e-08 | |
| neg_patterns.pattern_11 | 79 | | | MECP2_HUMAN.H11MO.0.C | 1.424340e-01 | |
| neg_patterns.pattern_12 | 77 | | | TN5_2 | 9.999990e-01 | |
| neg_patterns.pattern_13 | 71 | | | TN5_1 | 1.000000e+00 | |
| neg_patterns.pattern_14 | 70 | | | ZFX_MOUSE.H11MO.0.B | 1.348220e-04 | |
| neg_patterns.pattern_15 | 69 | | | NR2C2_MA0504.1 | 5.361540e-02 | |
| neg_patterns.pattern_16 | 68 | | | ZN331_HUMAN.H11MO.0.C | 3.358310e-01 | |
| neg_patterns.pattern_17 | 41 | | | EGR1_HUMAN.H11MO.0.A | 4.039400e-02 | |
| neg_patterns.pattern_18 | 36 | | | ZN322_MOUSE.H11MO.0.B | 1.000000e+00 | |
