## Supplementary Files 3 for "ChromBPNet: bias factorized, base-resolution deep learning models of chromatin accessibility reveal cis-regulatory sequence syntax, transcription factor footprints and regulatory variants": gm12878_ATAC_raw_bpnet_bias_fold1_profile_modisco.pdf

| pattern | num_seqlets | cwm_fwd | cwm_rev | TOMTOM_match | TOMTOM_qval | TOMTOM_match_logo |
|---|---|---|---|---|---|---|
| pos_patterns.pattern_0 | 7822 | | | TN5_1 | 8.270390e-09 | |
| pos_patterns.pattern_1 | 5272 | | | TN5_4 | 2.229310e-02 | |
| pos_patterns.pattern_2 | 4320 | | | TN5_1 | 5.971440e-05 | |
| pos_patterns.pattern_3 | 2739 | | | TN5_2 | 6.309330e-13 | |
| pos_patterns.pattern_4 | 2423 | | | KLF4_MA0039.3 | 4.298560e-01 | |
| pos_patterns.pattern_5 | 2224 | | | TN5_3 | 5.846380e-05 | |
| pos_patterns.pattern_6 | 1812 | | | TN5_3 | 1.512530e-12 | |
| pos_patterns.pattern_7 | 1128 | | | TN5_3 | 2.447420e-04 | |
| pos_patterns.pattern_8 | 603 | | | TN5_6 | 2.250980e-15 | |
| pos_patterns.pattern_9 | 556 | | | TN5_3 | 5.321250e-02 | |
| pos_patterns.pattern_10 | 185 | | | ZNF384_MA1125.1 | 8.226180e-02 | |
| pos_patterns.pattern_11 | 134 | | | EGR2_HUMAN.H11MO.0.A | 1.000000e+00 | |
| pos_patterns.pattern_12 | 98 | | | TN5_3 | 1.509640e-01 | |
| pos_patterns.pattern_13 | 50 | | | TN5_6 | 2.004820e-06 | |
| pos_patterns.pattern_14 | 43 | | | PRDM6_HUMAN.H11MO.0.C | 4.299110e-02 | |
