## Supplementary Files 3 for "ChromBPNet: bias factorized, base-resolution deep learning models of chromatin accessibility reveal cis-regulatory sequence syntax, transcription factor footprints and regulatory variants": gm12878_ATAC_raw_bpnet_bias_fold2_counts_modisco.pdf

| pattern | num_seqlets | cwm_fwd | cwm_rev | TOMTOM_match | TOMTOM_qval | TOMTOM_match_logo |
|---|---|---|---|---|---|---|
| pos_patterns.pattern_0 | 5009 | | | Zfx_MA0146.2 | 3.692740e-01 | |
| pos_patterns.pattern_1 | 3048 | | | PRD16_MOUSE.H11MO.0.B | 2.560730e-03 | |
| pos_patterns.pattern_2 | 1014 | | | TBX21_TBX_4 | 6.613920e-01 | |
| pos_patterns.pattern_3 | 955 | | | KLF5_MOUSE.H11MO.0.A | 9.115280e-01 | |
| pos_patterns.pattern_4 | 865 | | | ZN467_HUMAN.H11MO.0.C | 1.522520e-08 | |
| pos_patterns.pattern_5 | 696 | | | ZNF85_HUMAN.H11MO.0.C | 1.000000e+00 | |
| pos_patterns.pattern_6 | 620 | | | ZNF143_C2H2_1 | 2.731160e-01 | |
| pos_patterns.pattern_7 | 568 | | | ZIM3_HUMAN.H11MO.0.C | 6.123900e-02 | |
| pos_patterns.pattern_8 | 552 | | | ZN382_HUMAN.H11MO.0.C | 1.414040e-01 | |
| pos_patterns.pattern_9 | 504 | | | SIX1_HUMAN.H11MO.0.A | 9.999980e-01 | |
| pos_patterns.pattern_10 | 493 | | | ZNF354C_MA0130.1 | 1.000000e+00 | |
| pos_patterns.pattern_11 | 490 | | | TF65_HUMAN.H11MO.0.A | 2.609800e-01 | |
| pos_patterns.pattern_12 | 459 | | | ZNF524_C2H2_1 | 1.557100e-01 | |
| pos_patterns.pattern_13 | 453 | | | PRDM6_HUMAN.H11MO.0.C | 9.410920e-03 | |
| pos_patterns.pattern_14 | 404 | | | SOX17_HUMAN.H11MO.0.C | 7.245020e-01 | |
| pos_patterns.pattern_15 | 365 | | | FOXO1_HUMAN.H11MO.0.A | 5.786890e-01 | |
| pos_patterns.pattern_16 | 333 | | | BATF3_HUMAN.H11MO.0.B | 1.000000e+00 | |
| pos_patterns.pattern_17 | 296 | | | ARI5B_HUMAN.H11MO.0.C | 1.000000e+00 | |
| pos_patterns.pattern_18 | 289 | | | ZNF384_MA1125.1 | 8.713580e-02 | |
| pos_patterns.pattern_19 | 275 | | | PITX1_MOUSE.H11MO.0.C | 2.281660e-01 | |
| pos_patterns.pattern_20 | 251 | | | ZNF384_MA1125.1 | 1.496470e-02 | |
| pos_patterns.pattern_21 | 231 | | | BATF_MOUSE.H11MO.0.A | 1.000000e+00 | |
| pos_patterns.pattern_22 | 225 | | | NFAC2_HUMAN.H11MO.0.B | 5.257230e-01 | |
| pos_patterns.pattern_23 | 73 | | | SMAD3_HUMAN.H11MO.0.B | 1.000000e+00 | |
| pos_patterns.pattern_24 | 67 | | | RELA_MA0107.1 | 9.228360e-01 | |
| pos_patterns.pattern_25 | 39 | | | THAP1_HUMAN.H11MO.0.C | 1.845960e-01 | |
| pos_patterns.pattern_26 | 25 | | | GLIS1_C2H2_1 | 1.000000e+00 | |
| neg_patterns.pattern_0 | 357 | | | ZN331_HUMAN.H11MO.0.C | 1.027600e-01 | |
| neg_patterns.pattern_1 | 152 | | | ERG_HUMAN.H11MO.0.A | 3.322090e-01 | |

| pattern | num_seqlets | cwm_fwd | cwm_rev | TOMTOM_match | TOMTOM_qval | TOMTOM_match_logo |
|---|---|---|---|---|---|---|
| neg_patterns.pattern_2 | 149 | | | TN5_1 | 4.369900e-01 | |
| neg_patterns.pattern_3 | 145 | | | ZN667_HUMAN.H11MO.0.C | 1.000000e+00 | |
| neg_patterns.pattern_4 | 138 | | | TN5_2 | 1.724740e-02 | |
| neg_patterns.pattern_5 | 131 | | | TN5_2 | 4.913220e-03 | |
| neg_patterns.pattern_6 | 131 | | | ETV5_HUMAN.H11MO.0.C | 3.620440e-02 | |
| neg_patterns.pattern_7 | 122 | | | TN5_1 | 1.199890e-05 | |
| neg_patterns.pattern_8 | 112 | | | ZN667_HUMAN.H11MO.0.C | 1.000000e+00 | |
| neg_patterns.pattern_9 | 111 | | | TN5_2 | 7.222020e-02 | |
| neg_patterns.pattern_10 | 111 | | | PRDM6_HUMAN.H11MO.0.C | 5.925330e-02 | |
| neg_patterns.pattern_11 | 109 | | | TN5_1 | 1.482230e-01 | |
| neg_patterns.pattern_12 | 108 | | | TN5_2 | 1.000000e+00 | |
| neg_patterns.pattern_13 | 105 | | | NR1H3_MOUSE.H11MO.0.A | 5.086500e-01 | |
| neg_patterns.pattern_14 | 82 | | | ERR1_HUMAN.H11MO.0.A | 1.000000e+00 | |
| neg_patterns.pattern_15 | 77 | | | Z324A_HUMAN.H11MO.0.C | 1.648230e-01 | |
| neg_patterns.pattern_16 | 54 | | | TAL1_HUMAN.H11MO.0.A | 5.554190e-01 | |
| neg_patterns.pattern_17 | 46 | | | SP2_HUMAN.H11MO.0.A | 3.132240e-02 | |
| neg_patterns.pattern_18 | 41 | | | RREB1_MA0073.1 | 1.000000e+00 | |
