## Supplementary Files 3 for "ChromBPNet: bias factorized, base-resolution deep learning models of chromatin accessibility reveal cis-regulatory sequence syntax, transcription factor footprints and regulatory variants": gm12878_ATAC_raw_bpnet_bias_fold2_profile_modisco.pdf

| pattern | num_seqlets | cwm_fwd | cwm_rev | TOMTOM_match | TOMTOM_qval | TOMTOM_match_logo |
|---|---|---|---|---|---|---|
| pos_patterns.pattern_0 | 7437 | | | TN5_1 | 7.658590e-10 | |
| pos_patterns.pattern_1 | 5352 | | | TN5_4 | 2.418690e-02 | |
| pos_patterns.pattern_2 | 4089 | | | TN5_1 | 1.024400e-06 | |
| pos_patterns.pattern_3 | 2994 | | | TN5_2 | 4.821820e-15 | |
| pos_patterns.pattern_4 | 2552 | | | TN5_3 | 2.025430e-01 | |
| pos_patterns.pattern_5 | 2006 | | | TN5_3 | 2.619100e-13 | |
| pos_patterns.pattern_6 | 1154 | | | TN5_3 | 1.152600e-05 | |
| pos_patterns.pattern_7 | 788 | | | TN5_4 | 6.802640e-06 | |
| pos_patterns.pattern_8 | 659 | | | TN5_4 | 5.366920e-06 | |
| pos_patterns.pattern_9 | 550 | | | TN5_6 | 4.211230e-16 | |
| pos_patterns.pattern_10 | 522 | | | TN5_7 | 1.324550e-07 | |
| pos_patterns.pattern_11 | 389 | | | TN5_3 | 5.795100e-04 | |
| pos_patterns.pattern_12 | 383 | | | TN5_3 | 5.830120e-04 | |
| pos_patterns.pattern_13 | 191 | | | ZNF384_MA1125.1 | 8.557220e-02 | |
| pos_patterns.pattern_14 | 138 | | | EGR2_HUMAN.H11MO.0.A | 1.000000e+00 | |
| pos_patterns.pattern_15 | 110 | | | TN5_3 | 2.166000e-04 | |
| pos_patterns.pattern_16 | 75 | | | TBX1_TBX_1 | 2.409750e-01 | |
| pos_patterns.pattern_17 | 74 | | | TN5_6 | 8.135070e-07 | |
| pos_patterns.pattern_18 | 55 | | | TN5_6 | 6.431080e-01 | |
| pos_patterns.pattern_19 | 38 | | | TN5_6 | 9.888360e-03 | |
| pos_patterns.pattern_20 | 27 | | | TN5_3 | 3.540550e-02 | |
| pos_patterns.pattern_21 | 20 | | | NaN | NaN | |
