## Supplementary Files 3 for "ChromBPNet: bias factorized, base-resolution deep learning models of chromatin accessibility reveal cis-regulatory sequence syntax, transcription factor footprints and regulatory variants": gm12878_ATAC_raw_bpnet_bias_fold3_counts_modisco.pdf

| pattern | num_seqlets | cwm_fwd | cwm_rev | TOMTOM_match | TOMTOM_qval | TOMTOM_match_logo |
|---|---|---|---|---|---|---|
| pos_patterns.pattern_0 | 2898 | | | RARA_HUMAN.H11MO.0.A | 1.000000e+00 | |
| pos_patterns.pattern_1 | 2446 | | | FOXG1_forkhead_1 | 1.000000e+00 | |
| pos_patterns.pattern_2 | 1909 | | | ZFX_MOUSE.H11MO.0.B | 1.642920e-01 | |
| pos_patterns.pattern_3 | 1444 | | | TN5_6 | 1.000000e+00 | |
| pos_patterns.pattern_4 | 1125 | | | ZFX_MOUSE.H11MO.0.B | 8.413770e-01 | |
| pos_patterns.pattern_5 | 845 | | | ESR1_MA0112.3 | 1.000000e+00 | |
| pos_patterns.pattern_6 | 716 | | | CPEB1_RRM_1 | 7.049260e-01 | |
| pos_patterns.pattern_7 | 609 | | | NKX32_HUMAN.H11MO.0.C | 2.201210e-02 | |
| pos_patterns.pattern_8 | 593 | | | RREB1_MA0073.1 | 4.349290e-01 | |
| pos_patterns.pattern_9 | 588 | | | SP3_HUMAN.H11MO.0.B | 3.953510e-01 | |
| pos_patterns.pattern_10 | 497 | | | ARI5B_HUMAN.H11MO.0.C | 5.935980e-01 | |
| pos_patterns.pattern_11 | 495 | | | SUH_HUMAN.H11MO.0.A | 3.182160e-01 | |
| pos_patterns.pattern_12 | 434 | | | GLI2_C2H2_2 | 3.161670e-02 | |
| pos_patterns.pattern_13 | 422 | | | NFAC3_HUMAN.H11MO.0.B | 7.273110e-01 | |
| pos_patterns.pattern_14 | 414 | | | SP1_HUMAN.H11MO.0.A | 7.495330e-01 | |
| pos_patterns.pattern_15 | 400 | | | KLF9_HUMAN.H11MO.0.C | 8.648230e-01 | |
| pos_patterns.pattern_16 | 380 | | | RXRG_nuclearreceptor_4 | 5.101380e-02 | |
| pos_patterns.pattern_17 | 376 | | | DNASE_2 | 2.645940e-01 | |
| pos_patterns.pattern_18 | 357 | | | RREB1_MA0073.1 | 1.742160e-01 | |
| pos_patterns.pattern_19 | 357 | | | SMAD4_MOUSE.H11MO.0.A | 1.000000e+00 | |
| pos_patterns.pattern_20 | 342 | | | NRL_MA0842.1 | 1.816000e-01 | |
| pos_patterns.pattern_21 | 338 | | | IRF3_IRF_1 | 2.686040e-01 | |
| pos_patterns.pattern_22 | 333 | | | SP1_HUMAN.H11MO.0.A | 1.000000e+00 | |
| pos_patterns.pattern_23 | 277 | | | HOXA13_homeodomain_1 | 1.263750e-01 | |
| pos_patterns.pattern_24 | 263 | | | RUNX2_HUMAN.H11MO.0.A | 1.041400e-01 | |
| pos_patterns.pattern_25 | 223 | | | ZFX_MOUSE.H11MO.0.B | 9.965160e-01 | |
| pos_patterns.pattern_26 | 175 | | | NFIB_MOUSE.H11MO.0.C | 6.670110e-02 | |
| pos_patterns.pattern_27 | 85 | | | ZFP28_HUMAN.H11MO.0.C | 1.000000e+00 | |
| pos_patterns.pattern_28 | 78 | | | ZN770_HUMAN.H11MO.0.C | 9.184030e-01 | |

| pattern | num_seqlets | cwm_fwd | cwm_rev | TOMTOM_match | TOMTOM_qval | TOMTOM_match_logo |
|---|---|---|---|---|---|---|
| pos_patterns.pattern_29 | 34 | | | GLI1_MOUSE.H11MO.0.C | 2.626020e-01 | |
| neg_patterns.pattern_0 | 142 | | | SPIB_HUMAN.H11MO.0.A | 4.290230e-01 | |
| neg_patterns.pattern_1 | 116 | | | ZFX_MOUSE.H11MO.0.B | 8.505990e-04 | |
| neg_patterns.pattern_2 | 114 | | | ZFX_MOUSE.H11MO.0.B | 4.904940e-04 | |
| neg_patterns.pattern_3 | 109 | | | ZFX_MOUSE.H11MO.0.B | 1.706840e-03 | |
| neg_patterns.pattern_4 | 106 | | | SP2_HUMAN.H11MO.0.A | 6.364530e-07 | |
| neg_patterns.pattern_5 | 102 | | | ZN667_HUMAN.H11MO.0.C | 1.000000e+00 | |
| neg_patterns.pattern_6 | 98 | | | SP2_HUMAN.H11MO.0.A | 2.372990e-03 | |
| neg_patterns.pattern_7 | 93 | | | TCF7_MOUSE.H11MO.0.A | 1.000000e+00 | |
| neg_patterns.pattern_8 | 88 | | | ZFX_MOUSE.H11MO.0.B | 3.690310e-05 | |
| neg_patterns.pattern_9 | 81 | | | SP1_HUMAN.H11MO.0.A | 1.025250e-03 | |
| neg_patterns.pattern_10 | 73 | | | FOXP2_MA0593.1 | 2.113070e-01 | |
| neg_patterns.pattern_11 | 72 | | | TN5_1 | 9.992260e-01 | |
| neg_patterns.pattern_12 | 68 | | | TN5_2 | 1.624960e-02 | |
| neg_patterns.pattern_13 | 57 | | | TN5_2 | 4.790880e-02 | |
| neg_patterns.pattern_14 | 56 | | | TN5_2 | 4.887350e-02 | |
| neg_patterns.pattern_15 | 55 | | | TN5_7 | 6.611330e-01 | |
| neg_patterns.pattern_16 | 54 | | | TN5_2 | 2.910640e-02 | |
| neg_patterns.pattern_17 | 43 | | | COT2_HUMAN.H11MO.0.A | 8.533560e-02 | |
| neg_patterns.pattern_18 | 38 | | | TN5_2 | 1.379840e-03 | |
| neg_patterns.pattern_19 | 37 | | | TN5_2 | 1.552500e-03 | |
| neg_patterns.pattern_20 | 34 | | | TN5_2 | 2.809280e-01 | |
| neg_patterns.pattern_21 | 31 | | | TN5_8 | 2.540890e-02 | |
