## Supplementary Files 3 for "ChromBPNet: bias factorized, base-resolution deep learning models of chromatin accessibility reveal cis-regulatory sequence syntax, transcription factor footprints and regulatory variants": gm12878_ATAC_raw_bpnet_bias_fold3_profile_modisco.pdf

| pattern | num_seqlets | cwm_fwd | cwm_rev | TOMTOM_match | TOMTOM_qval | TOMTOM_match_logo |
|---|---|---|---|---|---|---|
| pos_patterns.pattern_0 | 9191 | 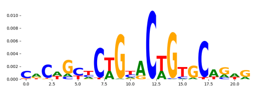 | 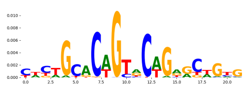 | TN5_2 | 8.122110e-09 | 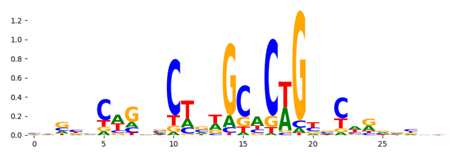 |
| pos_patterns.pattern_1 | 5255 | 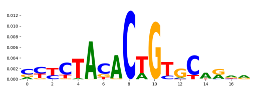 | 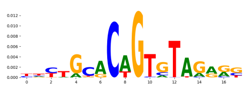 | TN5_4 | 1.493170e-02 | 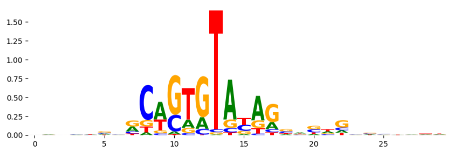 |
| pos_patterns.pattern_2 | 4201 | 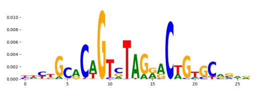 | 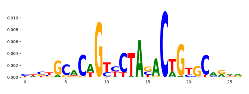 | TN5_1 | 1.115290e-06 | 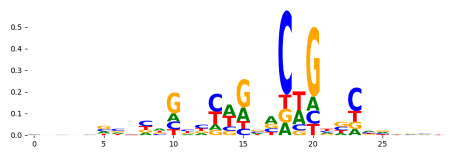 |
| pos_patterns.pattern_3 | 3464 | 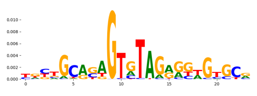 | 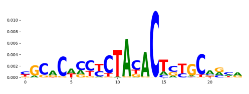 | TN5_3 | 9.726860e-02 | 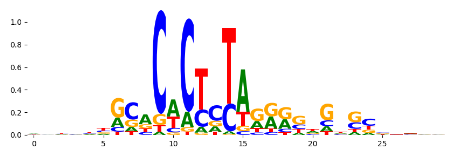 |
| pos_patterns.pattern_4 | 2934 | 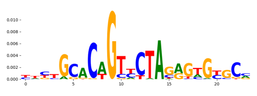 | 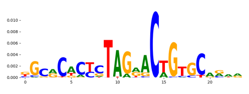 | TN5_3 | 1.030630e-04 | 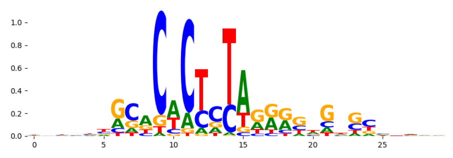 |
| pos_patterns.pattern_5 | 2895 | 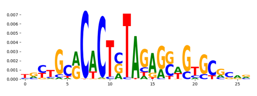 | 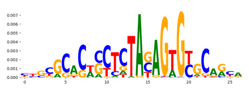 | TN5_3 | 4.175510e-14 | 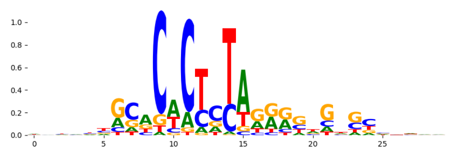 |
| pos_patterns.pattern_6 | 606 | 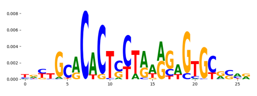 | 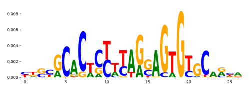 | TN5_3 | 1.062170e-06 | 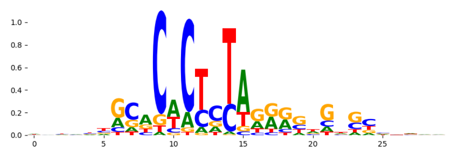 |
| pos_patterns.pattern_7 | 480 | 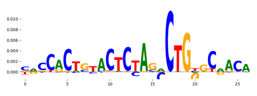 | 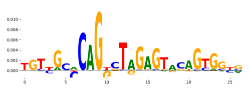 | TN5_6 | 2.994360e-19 | 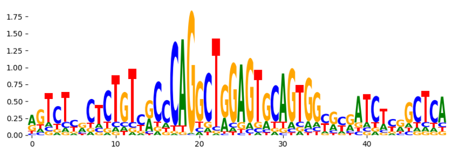 |
| pos_patterns.pattern_8 | 219 | 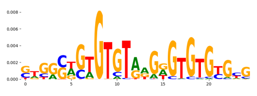 | 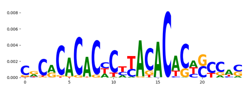 | EGR2_HUMAN.H11MO.0.A | 1.000000e+00 | 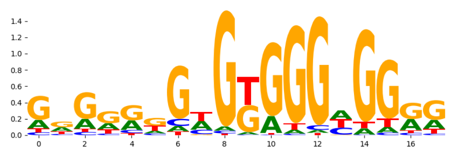 |
| pos_patterns.pattern_9 | 47 | 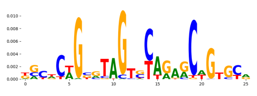 | 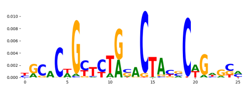 | TN5_1 | 8.274060e-03 | 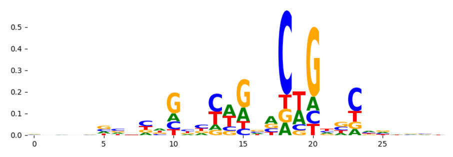 |
| pos_patterns.pattern_10 | 40 | 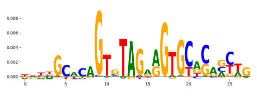 | 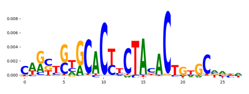 | ID4_MA0824.1 | 1.000000e+00 | 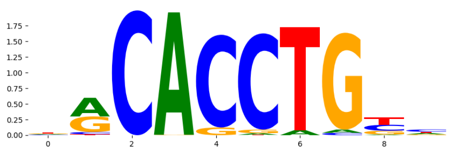 |
| pos_patterns.pattern_11 | 23 | 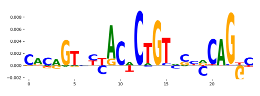 | 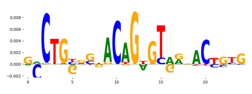 | TN5_6 | 7.961920e-08 | 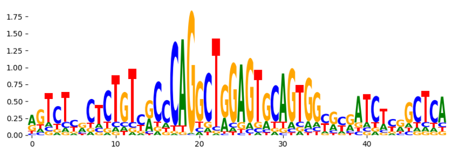 |
| pos_patterns.pattern_12 | 20 | 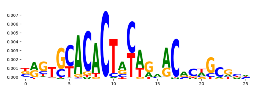 | 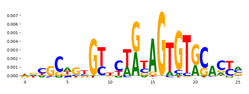 | ZN554_HUMAN.H11MO.0.C | 1.288120e-01 | 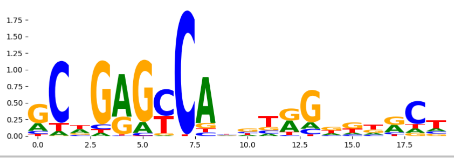 |
