## Supplementary Files 3 for "ChromBPNet: bias factorized, base-resolution deep learning models of chromatin accessibility reveal cis-regulatory sequence syntax, transcription factor footprints and regulatory variants": gm12878_ATAC_raw_bpnet_bias_fold4_counts_modisco.pdf

| pattern | num_seqlets | cwm_fwd | cwm_rev | TOMTOM_match | TOMTOM_qval | TOMTOM_match_logo |
|---|---|---|---|---|---|---|
| pos_patterns.pattern_0 | 3970 | | | NaN | NaN | |
| pos_patterns.pattern_1 | 1938 | | | ZBT17_MOUSE.H11MO.0.A | 5.572700e-01 | |
| pos_patterns.pattern_2 | 1263 | | | NaN | NaN | |
| pos_patterns.pattern_3 | 993 | | | SP1_HUMAN.H11MO.0.A | 3.986790e-02 | |
| pos_patterns.pattern_4 | 778 | | | TBX21_TBX_4 | 2.394480e-01 | |
| pos_patterns.pattern_5 | 762 | | | HIC1_HUMAN.H11MO.0.C | 3.479560e-01 | |
| pos_patterns.pattern_6 | 695 | | | RUNX3_RUNX_3 | 4.701590e-01 | |
| pos_patterns.pattern_7 | 655 | | | TF65_HUMAN.H11MO.0.A | 3.217300e-01 | |
| pos_patterns.pattern_8 | 552 | | | STAT1_MOUSE.H11MO.0.A | 1.846030e-03 | |
| pos_patterns.pattern_9 | 457 | | | RELB_HUMAN.H11MO.0.C | 1.016070e-01 | |
| pos_patterns.pattern_10 | 457 | | | RREB1_MA0073.1 | 7.600580e-02 | |
| pos_patterns.pattern_11 | 450 | | | RXRG_nuclearreceptor_4 | 1.000000e+00 | |
| pos_patterns.pattern_12 | 447 | | | RREB1_MA0073.1 | 2.713940e-03 | |
| pos_patterns.pattern_13 | 418 | | | SMAD4_MOUSE.H11MO.0.A | 1.000000e+00 | |
| pos_patterns.pattern_14 | 417 | | | ZBT17_MOUSE.H11MO.0.A | 1.036140e-01 | |
| pos_patterns.pattern_15 | 392 | | | ZN436_HUMAN.H11MO.0.C | 1.000000e+00 | |
| pos_patterns.pattern_16 | 389 | | | RREB1_MA0073.1 | 1.768800e-02 | |
| pos_patterns.pattern_17 | 375 | | | ZFX_HUMAN.H11MO.0.A | 1.000000e+00 | |
| pos_patterns.pattern_18 | 372 | | | Bach1+Mafk_MA0591.1 | 5.560770e-01 | |
| pos_patterns.pattern_19 | 356 | | | RREB1_MA0073.1 | 2.065060e-01 | |
| pos_patterns.pattern_20 | 310 | | | RREB1_MA0073.1 | 1.504670e-01 | |
| pos_patterns.pattern_21 | 303 | | | NFAC3_HUMAN.H11MO.0.B | 6.362900e-01 | |
| pos_patterns.pattern_22 | 291 | | | SMAD2_HUMAN.H11MO.0.A | 2.561830e-01 | |
| pos_patterns.pattern_23 | 288 | | | ZFX_MOUSE.H11MO.0.B | 5.131410e-01 | |
| pos_patterns.pattern_24 | 186 | | | SOX8_HMG_6 | 1.000000e+00 | |
| pos_patterns.pattern_25 | 181 | | | FOXO1_HUMAN.H11MO.0.A | 1.000000e+00 | |
| pos_patterns.pattern_26 | 164 | | | RUNX3_RUNX_2 | 1.000000e+00 | |
| pos_patterns.pattern_27 | 147 | | | HINFP1_C2H2_3 | 5.035720e-01 | |
| pos_patterns.pattern_28 | 144 | | | FEV_HUMAN.H11MO.0.B | 1.000000e+00 | |
| pos_patterns.pattern_29 | 128 | | | SALL4_HUMAN.H11MO.0.B | 1.000000e+00 | |

| pattern | num_seqlets | cwm_fwd | cwm_rev | TOMTOM_match | TOMTOM_qval | TOMTOM_match_logo |
|---|---|---|---|---|---|---|
| pos_patterns.pattern_30 | 116 | | | HINFP1_C2H2_3 | 1.000000e+00 | |
| pos_patterns.pattern_31 | 103 | | | PBX3_MA1114.1 | 3.555710e-01 | |
| pos_patterns.pattern_32 | 76 | | | FOXB1_forkhead_2 | 2.698020e-01 | |
| pos_patterns.pattern_33 | 51 | | | ZN502_HUMAN.H11MO.0.C | 1.353210e-01 | |
| pos_patterns.pattern_34 | 34 | | | Zfx_MA0146.2 | 1.000000e+00 | |
| pos_patterns.pattern_35 | 29 | | | ZN384_HUMAN.H11MO.0.C | 9.938560e-02 | |
| pos_patterns.pattern_36 | 25 | | | ZN502_HUMAN.H11MO.0.C | 1.457090e-07 | |
| pos_patterns.pattern_37 | 22 | | | USF1_MA0093.2 | 1.000000e+00 | |
| pos_patterns.pattern_38 | 21 | | | ZNF524_C2H2_2 | 1.000000e+00 | |
| neg_patterns.pattern_0 | 447 | | | TN5_2 | 3.500820e-03 | |
| neg_patterns.pattern_1 | 183 | | | TBX20_MOUSE.H11MO.0.C | 1.398580e-01 | |
| neg_patterns.pattern_2 | 167 | | | NKX28_HUMAN.H11MO.0.C | 9.664850e-01 | |
| neg_patterns.pattern_3 | 159 | | | SP2_HUMAN.H11MO.0.A | 5.926800e-08 | |
| neg_patterns.pattern_4 | 149 | | | ZFX_MOUSE.H11MO.0.B | 9.049580e-03 | |
| neg_patterns.pattern_5 | 148 | | | TN5_7 | 1.613830e-02 | |
| neg_patterns.pattern_6 | 140 | | | TF7L1_MOUSE.H11MO.0.A | 1.000000e+00 | |
| neg_patterns.pattern_7 | 133 | | | COT2_MOUSE.H11MO.0.A | 1.882980e-01 | |
| neg_patterns.pattern_8 | 128 | | | SP2_HUMAN.H11MO.0.A | 1.155020e-05 | |
| neg_patterns.pattern_9 | 119 | | | TN5_2 | 3.746920e-07 | |
| neg_patterns.pattern_10 | 116 | | | ETV2_HUMAN.H11MO.0.B | 5.796730e-03 | |
| neg_patterns.pattern_11 | 113 | | | ZFX_MOUSE.H11MO.0.B | 2.137900e-02 | |
| neg_patterns.pattern_12 | 111 | | | PRDM6_HUMAN.H11MO.0.C | 7.472100e-03 | |
| neg_patterns.pattern_13 | 110 | | | FOXO1_HUMAN.H11MO.0.A | 7.695380e-01 | |
| neg_patterns.pattern_14 | 101 | | | TP53_MA0106.3 | 1.000000e+00 | |
| neg_patterns.pattern_15 | 77 | | | SP1_MOUSE.H11MO.0.A | 2.256300e-08 | |
| neg_patterns.pattern_16 | 74 | | | SP3_HUMAN.H11MO.0.B | 1.901320e-04 | |
| neg_patterns.pattern_17 | 65 | | | NRF1_MOUSE.H11MO.0.A | 9.720380e-03 | |
| neg_patterns.pattern_18 | 44 | | | TN5_1 | 5.125290e-03 | |
| neg_patterns.pattern_19 | 40 | | | TN5_8 | 1.000000e+00 | |
