## Supplementary Files 3 for "ChromBPNet: bias factorized, base-resolution deep learning models of chromatin accessibility reveal cis-regulatory sequence syntax, transcription factor footprints and regulatory variants": gm12878_ATAC_raw_bpnet_bias_fold4_profile_modisco.pdf

| pattern | num_seqlets | cwm_fwd | cwm_rev | TOMTOM_match | TOMTOM_qval | TOMTOM_match_logo |
|---|---|---|---|---|---|---|
| pos_patterns.pattern_0 | 8115 | | | TN5_2 | 2.446840e-09 | |
| pos_patterns.pattern_1 | 5347 | | | TN5_4 | 1.633960e-02 | |
| pos_patterns.pattern_2 | 4304 | | | TN5_1 | 8.777700e-05 | |
| pos_patterns.pattern_3 | 2677 | | | TN5_2 | 2.040260e-11 | |
| pos_patterns.pattern_4 | 2464 | | | TN5_3 | 3.504910e-11 | |
| pos_patterns.pattern_5 | 2464 | | | TN5_3 | 2.144590e-01 | |
| pos_patterns.pattern_6 | 2038 | | | TN5_3 | 1.059380e-04 | |
| pos_patterns.pattern_7 | 591 | | | TN5_3 | 1.926640e-06 | |
| pos_patterns.pattern_8 | 543 | | | TN5_4 | 2.793510e-06 | |
| pos_patterns.pattern_9 | 530 | | | TN5_6 | 4.519450e-15 | |
| pos_patterns.pattern_10 | 138 | | | TN5_4 | 2.263700e-02 | |
| pos_patterns.pattern_11 | 119 | | | TN5_3 | 1.196820e-04 | |
| pos_patterns.pattern_12 | 114 | | | EGR2_HUMAN.H11MO.0.A | 1.000000e+00 | |
| pos_patterns.pattern_13 | 22 | | | TN5_3 | 2.866730e-04 | |
