## Supplementary Files 3 for "ChromBPNet: bias factorized, base-resolution deep learning models of chromatin accessibility reveal cis-regulatory sequence syntax, transcription factor footprints and regulatory variants": h1esc_ATAC_raw_bpnet_bias_fold0_counts_modisco.pdf

| pattern | num_seqlets | cwm_fwd | cwm_rev | TOMTOM_match | TOMTOM_qval | TOMTOM_match_logo |
|---|---|---|---|---|---|---|
| pos_patterns.pattern_0 | 3688 | | | Zfx_MA0146.2 | 0.417986 | |
| pos_patterns.pattern_1 | 3194 | | | SP3_HUMAN.H11MO.0.B | 0.005538 | |
| pos_patterns.pattern_2 | 2719 | | | ZFX_MOUSE.H11MO.0.B | 0.007030 | |
| pos_patterns.pattern_3 | 706 | | | NR2C2_MOUSE.H11MO.0.A | 0.687804 | |
| pos_patterns.pattern_4 | 663 | | | DNASE_4 | 0.298947 | |
| pos_patterns.pattern_5 | 560 | | | SP2_HUMAN.H11MO.0.A | 0.132577 | |
| pos_patterns.pattern_6 | 540 | | | NFATC3_MA0625.1 | 0.982323 | |
| pos_patterns.pattern_7 | 533 | | | PRDM6_HUMAN.H11MO.0.C | 0.190641 | |
| pos_patterns.pattern_8 | 528 | | | ZNF384_MA1125.1 | 0.059846 | |
| pos_patterns.pattern_9 | 480 | | | HXA13_HUMAN.H11MO.0.C | 0.773619 | |
| pos_patterns.pattern_10 | 405 | | | Z354A_HUMAN.H11MO.0.C | 1.000000 | |
| pos_patterns.pattern_11 | 320 | | | NFAC3_HUMAN.H11MO.0.B | 0.030386 | |
| pos_patterns.pattern_12 | 288 | | | CLOCK_HUMAN.H11MO.0.C | 0.502019 | |
| pos_patterns.pattern_13 | 274 | | | ZIM3_HUMAN.H11MO.0.C | 0.956574 | |
| pos_patterns.pattern_14 | 174 | | | NaN | NaN | |
| pos_patterns.pattern_15 | 164 | | | IRF5_IRF_1 | 1.000000 | |
| pos_patterns.pattern_16 | 115 | | | ZBT18_HUMAN.H11MO.0.C | 1.000000 | |
| pos_patterns.pattern_17 | 103 | | | NKX2-8_MA0673.1 | 1.000000 | |
| pos_patterns.pattern_18 | 73 | | | HIC1_HUMAN.H11MO.0.C | 0.932626 | |
| pos_patterns.pattern_19 | 59 | | | MTF1_C2H2_1 | 0.817915 | |
| pos_patterns.pattern_20 | 55 | | | SMCA5_MOUSE.H11MO.0.C | 0.018338 | |
| pos_patterns.pattern_21 | 43 | | | PKNOX2_MA0783.1 | 0.999997 | |
| pos_patterns.pattern_22 | 39 | | | HOXD13_MA0909.1 | 1.000000 | |
| pos_patterns.pattern_23 | 37 | | | ELK1_ETS_3 | 1.000000 | |
| neg_patterns.pattern_0 | 42 | | | SP1_HUMAN.H11MO.0.A | 0.006678 | |
| neg_patterns.pattern_1 | 41 | | | TN5_7 | 0.133522 | |
| neg_patterns.pattern_2 | 27 | | | SP1_MOUSE.H11MO.0.A | 0.000020 | |
| neg_patterns.pattern_3 | 22 | | | Zfx_MA0146.2 | 0.204657 | |
