## Supplementary Files 3 for "ChromBPNet: bias factorized, base-resolution deep learning models of chromatin accessibility reveal cis-regulatory sequence syntax, transcription factor footprints and regulatory variants": h1esc_ATAC_raw_bpnet_bias_fold0_profile_modisco.pdf

| pattern | num_seqlets | cwm_fwd | cwm_rev | TOMTOM_match | TOMTOM_qval | TOMTOM_match_logo |
|---|---|---|---|---|---|---|
| pos_patterns.pattern_0 | 8300 | | | TN5_1 | 2.528420e-10 | |
| pos_patterns.pattern_1 | 6387 | | | TN5_4 | 2.016040e-02 | |
| pos_patterns.pattern_2 | 4160 | | | TN5_1 | 8.705690e-08 | |
| pos_patterns.pattern_3 | 4034 | | | TN5_2 | 7.075140e-12 | |
| pos_patterns.pattern_4 | 3402 | | | KLF4_MA0039.3 | 9.051140e-02 | |
| pos_patterns.pattern_5 | 1151 | | | TN5_3 | 1.227700e-03 | |
| pos_patterns.pattern_6 | 1056 | | | TN5_3 | 2.715650e-07 | |
| pos_patterns.pattern_7 | 982 | | | TN5_7 | 3.901690e-08 | |
| pos_patterns.pattern_8 | 842 | | | TN5_3 | 3.273350e-04 | |
| pos_patterns.pattern_9 | 828 | | | TN5_3 | 6.993060e-08 | |
| pos_patterns.pattern_10 | 755 | | | TN5_3 | 1.695140e-06 | |
| pos_patterns.pattern_11 | 625 | | | TN5_3 | 3.409020e-05 | |
| pos_patterns.pattern_12 | 533 | | | TN5_1 | 1.179610e-04 | |
| pos_patterns.pattern_13 | 261 | | | PRDM6_HUMAN.H11MO.0.C | 4.827240e-02 | |
| pos_patterns.pattern_14 | 165 | | | EGR2_HUMAN.H11MO.0.A | 6.844730e-01 | |
| pos_patterns.pattern_15 | 111 | | | PRDM6_HUMAN.H11MO.0.C | 1.041710e-01 | |
| pos_patterns.pattern_16 | 72 | | | TN5_6 | 3.254370e-17 | |
| pos_patterns.pattern_17 | 61 | | | Hoxc10.mouse_homeodomain_2 | 3.858620e-01 | |
| pos_patterns.pattern_18 | 28 | | | ZN331_HUMAN.H11MO.0.C | 1.000000e+00 | |
| pos_patterns.pattern_19 | 23 | | | FOXJ3_HUMAN.H11MO.0.A | 1.000000e+00 | |
| pos_patterns.pattern_20 | 22 | | | TN5_2 | 6.189360e-02 | |
| pos_patterns.pattern_21 | 20 | | | TN5_3 | 1.065650e-02 | |
| neg_patterns.pattern_0 | 27 | | | ZNF384_MA1125.1 | 1.786380e-03 | |
| neg_patterns.pattern_1 | 24 | | | PRDM6_HUMAN.H11MO.0.C | 2.132810e-02 | |
| neg_patterns.pattern_2 | 23 | | | SRY_HUMAN.H11MO.0.B | 1.000000e+00 | |
