## Supplementary Files 3 for "ChromBPNet: bias factorized, base-resolution deep learning models of chromatin accessibility reveal cis-regulatory sequence syntax, transcription factor footprints and regulatory variants": h1esc_ATAC_raw_bpnet_bias_fold1_counts_modisco.pdf

| pattern | num_seqlets | cwm_fwd | cwm_rev | TOMTOM_match | TOMTOM_qval | TOMTOM_match_logo |
|---|---|---|---|---|---|---|
| pos_patterns.pattern_0 | 7390 | | | ZFX_MOUSE.H11MO.0.B | 0.212187 | |
| pos_patterns.pattern_1 | 2936 | | | SP2_HUMAN.H11MO.0.A | 0.188666 | |
| pos_patterns.pattern_2 | 2406 | | | USF1_MA0093.2 | 1.000000 | |
| pos_patterns.pattern_3 | 980 | | | RUNX2_HUMAN.H11MO.0.A | 0.664537 | |
| pos_patterns.pattern_4 | 701 | | | TEAD3_TEA_1 | 1.000000 | |
| pos_patterns.pattern_5 | 672 | | | CDX2_MA0465.1 | 1.000000 | |
| pos_patterns.pattern_6 | 657 | | | ETS1_ETS_4 | 0.364602 | |
| pos_patterns.pattern_7 | 621 | | | TF65_HUMAN.H11MO.0.A | 1.000000 | |
| pos_patterns.pattern_8 | 501 | | | PRDM6_HUMAN.H11MO.0.C | 0.098119 | |
| pos_patterns.pattern_9 | 470 | | | FOXJ3_HUMAN.H11MO.0.A | 0.431864 | |
| pos_patterns.pattern_10 | 417 | | | RREB1_MA0073.1 | 0.872257 | |
| pos_patterns.pattern_11 | 303 | | | NFATC2_MA0152.1 | 1.000000 | |
| pos_patterns.pattern_12 | 251 | | | HTF4_HUMAN.H11MO.0.A | 0.424142 | |
| pos_patterns.pattern_13 | 132 | | | CTCFL_HUMAN.H11MO.0.A | 0.157937 | |
| pos_patterns.pattern_14 | 108 | | | Hic1.mouse_C2H2_1 | 1.000000 | |
| pos_patterns.pattern_15 | 87 | | | FOXB1_forkhead_2 | 0.414740 | |
| pos_patterns.pattern_16 | 86 | | | RUNX1_HUMAN.H11MO.0.A | 1.000000 | |
| pos_patterns.pattern_17 | 62 | | | ZIC3_C2H2_1 | 1.000000 | |
| pos_patterns.pattern_18 | 49 | | | SP1_HUMAN.H11MO.0.A | 0.370660 | |
| pos_patterns.pattern_19 | 33 | | | KLF12_HUMAN.H11MO.0.C | 0.091161 | |
| neg_patterns.pattern_0 | 39 | | | TN5_2 | 0.043042 | |
| neg_patterns.pattern_1 | 38 | | | TN5_1 | 0.009433 | |
| neg_patterns.pattern_2 | 36 | | | ZFX_MOUSE.H11MO.0.B | 0.003617 | |
| neg_patterns.pattern_3 | 35 | | | SP2_HUMAN.H11MO.0.A | 0.040634 | |
| neg_patterns.pattern_4 | 33 | | | TN5_2 | 0.016386 | |
| neg_patterns.pattern_5 | 28 | | | SP2_HUMAN.H11MO.0.A | 0.000033 | |
| neg_patterns.pattern_6 | 27 | | | Zfx_MA0146.2 | 0.000031 | |
| neg_patterns.pattern_7 | 25 | | | TN5_2 | 0.003152 | |
