## Supplementary Files 3 for "ChromBPNet: bias factorized, base-resolution deep learning models of chromatin accessibility reveal cis-regulatory sequence syntax, transcription factor footprints and regulatory variants": h1esc_ATAC_raw_bpnet_bias_fold1_profile_modisco.pdf

| pattern | num_seqlets | cwm_fwd | cwm_rev | TOMTOM_match | TOMTOM_qval | TOMTOM_match_logo |
|---|---|---|---|---|---|---|
| pos_patterns.pattern_0 | 8441 | | | TN5_1 | 1.554260e-09 | |
| pos_patterns.pattern_1 | 5578 | | | TN5_8 | 4.614530e-02 | |
| pos_patterns.pattern_2 | 4741 | | | TN5_1 | 4.544490e-06 | |
| pos_patterns.pattern_3 | 3948 | | | TN5_2 | 1.604460e-19 | |
| pos_patterns.pattern_4 | 3643 | | | KLF4_MA0039.3 | 3.157090e-01 | |
| pos_patterns.pattern_5 | 1089 | | | TN5_3 | 3.862770e-05 | |
| pos_patterns.pattern_6 | 1037 | | | TN5_3 | 1.330290e-03 | |
| pos_patterns.pattern_7 | 1036 | | | TN5_3 | 3.190970e-08 | |
| pos_patterns.pattern_8 | 893 | | | TN5_3 | 7.765820e-09 | |
| pos_patterns.pattern_9 | 764 | | | TN5_3 | 6.422400e-04 | |
| pos_patterns.pattern_10 | 747 | | | TN5_3 | 3.787640e-12 | |
| pos_patterns.pattern_11 | 459 | | | TN5_1 | 7.324630e-05 | |
| pos_patterns.pattern_12 | 354 | | | TN5_7 | 6.799020e-02 | |
| pos_patterns.pattern_13 | 286 | | | PRDM6_HUMAN.H11MO.0.C | 5.478950e-02 | |
| pos_patterns.pattern_14 | 126 | | | MEF2C_MA0497.1 | 5.342060e-01 | |
| pos_patterns.pattern_15 | 67 | | | TN5_8 | 1.000000e+00 | |
| pos_patterns.pattern_16 | 56 | | | VEZF1_HUMAN.H11MO.0.C | 7.340360e-02 | |
| pos_patterns.pattern_17 | 54 | | | RREB1_MA0073.1 | 1.000000e+00 | |
| neg_patterns.pattern_0 | 22 | | | CPEB1_RRM_1 | 1.770700e-01 | |
| neg_patterns.pattern_1 | 20 | | | ZNF384_MA1125.1 | 1.035220e-03 | |
