## Supplementary Files 3 for "ChromBPNet: bias factorized, base-resolution deep learning models of chromatin accessibility reveal cis-regulatory sequence syntax, transcription factor footprints and regulatory variants": h1esc_ATAC_raw_bpnet_bias_fold2_counts_modisco.pdf

| pattern | num_seqlets | cwm_fwd | cwm_rev | TOMTOM_match | TOMTOM_qval | TOMTOM_match_logo |
|---|---|---|---|---|---|---|
| pos_patterns.pattern_0 | 3323 | | | Zfx_MA0146.2 | 0.059327 | |
| pos_patterns.pattern_1 | 3011 | | | SP1_HUMAN.H11MO.0.A | 0.024532 | |
| pos_patterns.pattern_2 | 1553 | | | SP2_HUMAN.H11MO.0.A | 0.000013 | |
| pos_patterns.pattern_3 | 715 | | | ONECUT3_CUT_1 | 0.048967 | |
| pos_patterns.pattern_4 | 608 | | | SP2_HUMAN.H11MO.0.A | 0.000133 | |
| pos_patterns.pattern_5 | 590 | | | RUNX3_RUNX_3 | 1.000000 | |
| pos_patterns.pattern_6 | 552 | | | SP2_HUMAN.H11MO.0.A | 0.000054 | |
| pos_patterns.pattern_7 | 529 | | | TF7L1_HUMAN.H11MO.0.B | 0.246321 | |
| pos_patterns.pattern_8 | 503 | | | SP1_HUMAN.H11MO.0.A | 0.036730 | |
| pos_patterns.pattern_9 | 460 | | | MXI1_HUMAN.H11MO.0.A | 0.161324 | |
| pos_patterns.pattern_10 | 421 | | | TBX21_TBX_6 | 1.000000 | |
| pos_patterns.pattern_11 | 412 | | | SIX4_MOUSE.H11MO.0.C | 1.000000 | |
| pos_patterns.pattern_12 | 403 | | | SP2_HUMAN.H11MO.0.A | 0.000067 | |
| pos_patterns.pattern_13 | 382 | | | SP2_HUMAN.H11MO.0.A | 0.000073 | |
| pos_patterns.pattern_14 | 380 | | | RUNX2_HUMAN.H11MO.0.A | 0.880530 | |
| pos_patterns.pattern_15 | 368 | | | AIRE_HUMAN.H11MO.0.C | 1.000000 | |
| pos_patterns.pattern_16 | 364 | | | COE1_HUMAN.H11MO.0.A | 0.589882 | |
| pos_patterns.pattern_17 | 363 | | | NFATC2_MA0152.1 | 1.000000 | |
| pos_patterns.pattern_18 | 257 | | | MYC_MOUSE.H11MO.0.A | 0.635173 | |
| pos_patterns.pattern_19 | 218 | | | SP1_HUMAN.H11MO.0.A | 0.000144 | |
| pos_patterns.pattern_20 | 195 | | | SP2_HUMAN.H11MO.0.A | 0.000008 | |
| pos_patterns.pattern_21 | 174 | | | HOXA13_homeodomain_1 | 0.060299 | |
| pos_patterns.pattern_22 | 155 | | | SP1_HUMAN.H11MO.0.A | 0.000615 | |
| pos_patterns.pattern_23 | 137 | | | FOXC1_forkhead_1 | 0.057777 | |
| pos_patterns.pattern_24 | 55 | | | CDX2_MOUSE.H11MO.0.A | 0.904123 | |
| neg_patterns.pattern_0 | 79 | | | ZFX_MOUSE.H11MO.0.B | 0.000078 | |
| neg_patterns.pattern_1 | 65 | | | ZFX_MOUSE.H11MO.0.B | 0.000580 | |
| neg_patterns.pattern_2 | 53 | | | ZFX_MOUSE.H11MO.0.B | 0.003725 | |
| neg_patterns.pattern_3 | 42 | | | ZFX_MOUSE.H11MO.0.B | 0.004426 | |

| pattern | num_seqlets | cwm_fwd | cwm_rev | TOMTOM_match | TOMTOM_qval | TOMTOM_match_logo |
|---|---|---|---|---|---|---|
| neg_patterns.pattern_4 | 32 | 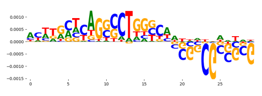 | 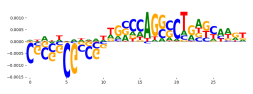 | ZFX_MOUSE.H11MO.0.B | 0.000003 | 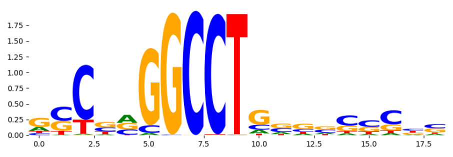 |
| neg_patterns.pattern_5 | 21 | 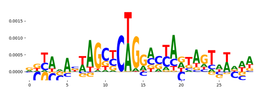 | 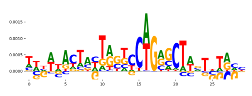 | SP1_MOUSE.H11MO.0.A | 0.008055 | 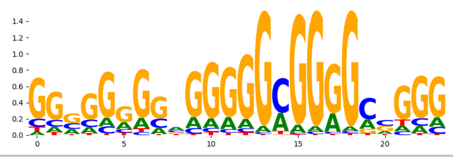 |
