## Supplementary Files 3 for "ChromBPNet: bias factorized, base-resolution deep learning models of chromatin accessibility reveal cis-regulatory sequence syntax, transcription factor footprints and regulatory variants": h1esc_ATAC_raw_bpnet_bias_fold2_profile_modisco.pdf

| pattern | num_seqlets | cwm_fwd | cwm_rev | TOMTOM_match | TOMTOM_qval | TOMTOM_match_logo |
|---|---|---|---|---|---|---|
| pos_patterns.pattern_0 | 9131 | 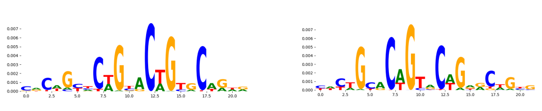 | | TN5_1 | 1.168880e-08 | 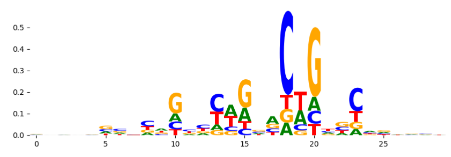 |
| pos_patterns.pattern_1 | 7828 | 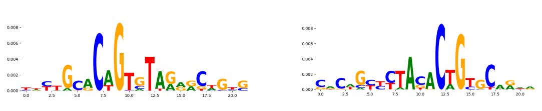 | | TN5_2 | 1.386410e-03 | 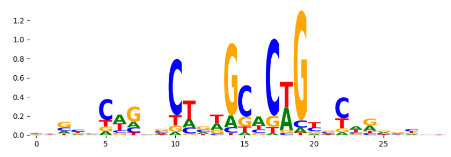 |
| pos_patterns.pattern_2 | 4650 | 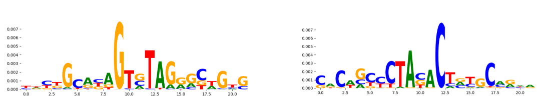 | | KLF4_MA0039.3 | 2.854600e-01 | 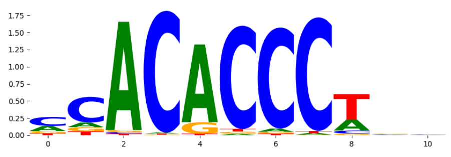 |
| pos_patterns.pattern_3 | 4395 | 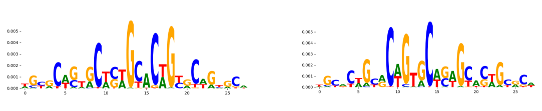 | | TN5_2 | 3.829520e-17 | 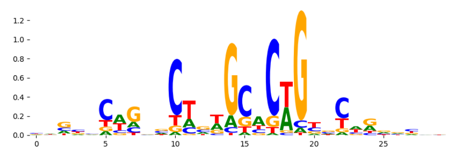 |
| pos_patterns.pattern_4 | 1439 | 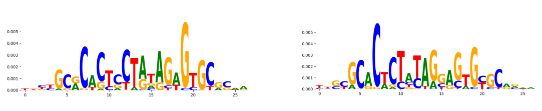 | | TN5_3 | 3.179490e-07 | 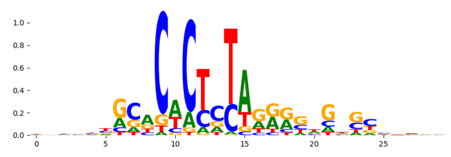 |
| pos_patterns.pattern_5 | 1076 | 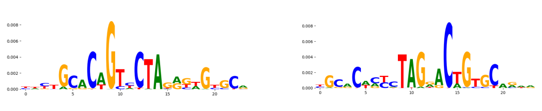 | | TN5_3 | 7.153540e-04 | 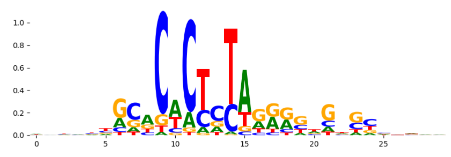 |
| pos_patterns.pattern_6 | 956 | 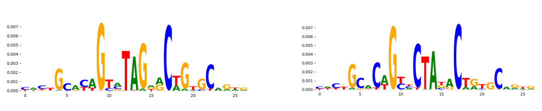 | | TN5_1 | 3.790070e-05 | 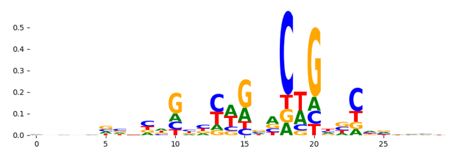 |
| pos_patterns.pattern_7 | 938 | 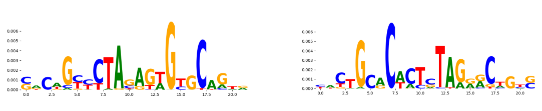 | | TN5_3 | 9.590350e-05 | 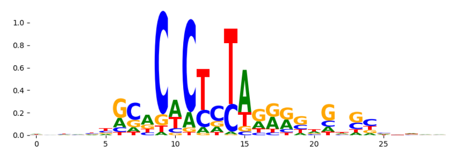 |
| pos_patterns.pattern_8 | 799 | 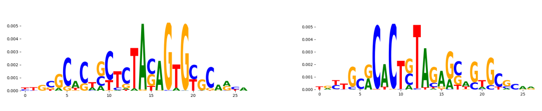 | | TN5_3 | 4.644990e-13 | 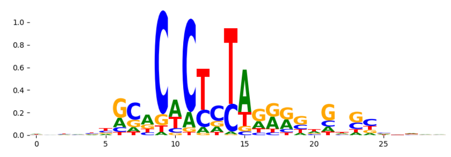 |
| pos_patterns.pattern_9 | 579 | 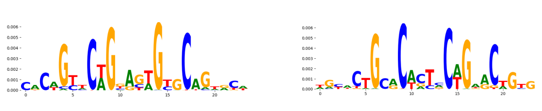 | | TN5_1 | 1.004600e-04 | 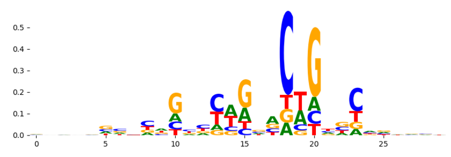 |
| pos_patterns.pattern_10 | 541 | 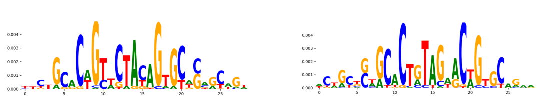 | | TN5_7 | 9.796010e-08 | 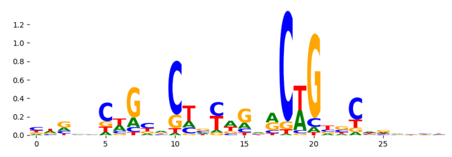 |
| pos_patterns.pattern_11 | 454 | 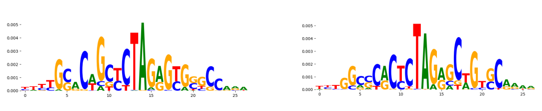 | | TN5_3 | 3.088290e-04 | 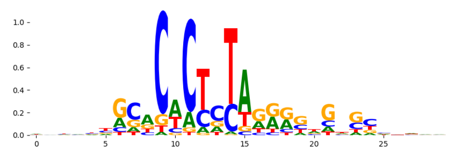 |
| pos_patterns.pattern_12 | 260 | 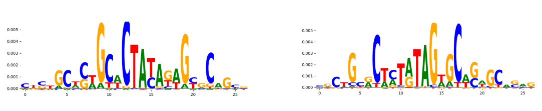 | | TN5_3 | 2.955170e-02 | 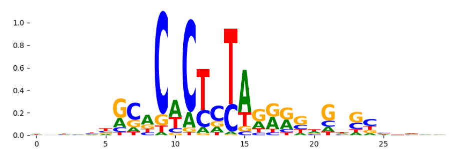 |
| pos_patterns.pattern_13 | 202 | 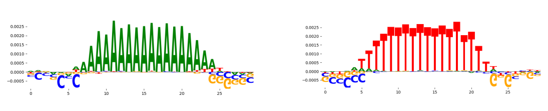 | | PRDM6_HUMAN.H11MO.0.C | 4.868180e-02 | 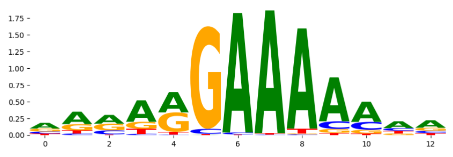 |
| pos_patterns.pattern_14 | 122 | 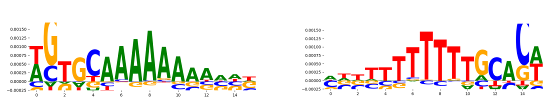 | | HOXC12_homeodomain_1 | 1.313010e-01 | 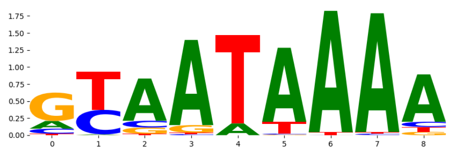 |
| pos_patterns.pattern_15 | 69 | 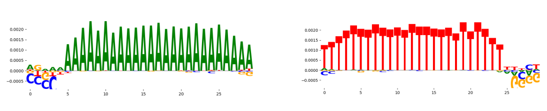 | | PRDM6_HUMAN.H11MO.0.C | 1.183380e-01 | 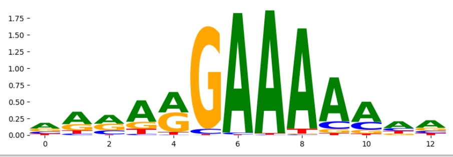 |
