## Supplementary Files 3 for "ChromBPNet: bias factorized, base-resolution deep learning models of chromatin accessibility reveal cis-regulatory sequence syntax, transcription factor footprints and regulatory variants": h1esc_ATAC_raw_bpnet_bias_fold3_counts_modisco.pdf

| pattern | num_seqlets | cwm_fwd | cwm_rev | TOMTOM_match | TOMTOM_qval | TOMTOM_match_logo |
|---|---|---|---|---|---|---|
| pos_patterns.pattern_0 | 2473 | | | MYC_MOUSE.H11MO.0.A | 1.000000e+00 | |
| pos_patterns.pattern_1 | 2120 | | | SP1_HUMAN.H11MO.0.A | 6.795050e-05 | |
| pos_patterns.pattern_2 | 1705 | | | SP1_HUMAN.H11MO.0.A | 5.390600e-04 | |
| pos_patterns.pattern_3 | 1368 | | | ZFX_MOUSE.H11MO.0.B | 3.409470e-04 | |
| pos_patterns.pattern_4 | 1245 | | | NHLH1_bHLH_1 | 4.422980e-01 | |
| pos_patterns.pattern_5 | 1018 | | | SP2_HUMAN.H11MO.0.A | 1.649610e-07 | |
| pos_patterns.pattern_6 | 838 | | | SP1_HUMAN.H11MO.0.A | 5.603110e-06 | |
| pos_patterns.pattern_7 | 745 | | | SP1_HUMAN.H11MO.0.A | 1.504240e-03 | |
| pos_patterns.pattern_8 | 615 | | | SP1_HUMAN.H11MO.0.A | 2.491780e-05 | |
| pos_patterns.pattern_9 | 582 | | | SP1_HUMAN.H11MO.0.A | 1.743420e-04 | |
| pos_patterns.pattern_10 | 579 | | | SP1_HUMAN.H11MO.0.A | 3.389240e-04 | |
| pos_patterns.pattern_11 | 557 | | | ZNF713_C2H2_1 | 3.559350e-01 | |
| pos_patterns.pattern_12 | 540 | | | NFATC1_NFAT_1 | 3.030710e-01 | |
| pos_patterns.pattern_13 | 525 | | | SP1_HUMAN.H11MO.0.A | 2.034130e-05 | |
| pos_patterns.pattern_14 | 485 | | | NFATC1_MA0624.1 | 1.000000e+00 | |
| pos_patterns.pattern_15 | 471 | | | PRDM6_HUMAN.H11MO.0.C | 9.969850e-02 | |
| pos_patterns.pattern_16 | 444 | | | MBD2_HUMAN.H11MO.0.B | 2.238740e-01 | |
| pos_patterns.pattern_17 | 442 | | | SP1_HUMAN.H11MO.0.A | 3.798030e-04 | |
| pos_patterns.pattern_18 | 438 | | | NFAC3_HUMAN.H11MO.0.B | 2.416170e-01 | |
| pos_patterns.pattern_19 | 397 | | | SP2_HUMAN.H11MO.0.A | 3.810530e-06 | |
| pos_patterns.pattern_20 | 336 | | | SP2_HUMAN.H11MO.0.A | 4.026590e-06 | |
| pos_patterns.pattern_21 | 334 | | | SP1_HUMAN.H11MO.0.A | 1.962430e-03 | |
| pos_patterns.pattern_22 | 299 | | | SP1_HUMAN.H11MO.0.A | 6.606220e-03 | |
| pos_patterns.pattern_23 | 249 | | | SP1_MOUSE.H11MO.0.A | 2.804140e-04 | |
| pos_patterns.pattern_24 | 243 | | | SP2_HUMAN.H11MO.0.A | 1.207740e-01 | |
| pos_patterns.pattern_25 | 223 | | | SP2_HUMAN.H11MO.0.A | 6.907690e-04 | |
| pos_patterns.pattern_26 | 213 | | | SP1_HUMAN.H11MO.0.A | 2.891790e-02 | |
| pos_patterns.pattern_27 | 212 | | | Hes1_MA1099.1 | 1.000000e+00 | |
| pos_patterns.pattern_28 | 202 | | | PBX3_MA1114.1 | 1.000000e+00 | |

| pattern | num_seqlets | cwm_fwd | cwm_rev | TOMTOM_match | TOMTOM_qval | TOMTOM_match_logo |
|---|---|---|---|---|---|---|
| pos_patterns.pattern_29 | 193 | 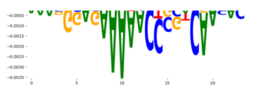 | 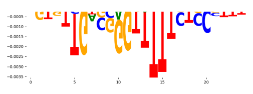 | ZN274_HUMAN.H11MO.0.A | 1.000000e+00 | 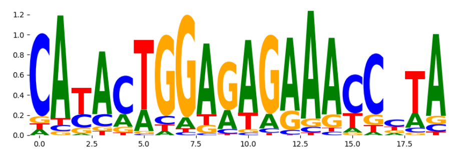 |
| pos_patterns.pattern_30 | 184 | 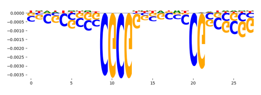 | 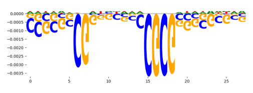 | SP2_HUMAN.H11MO.0.A | 4.056290e-04 | 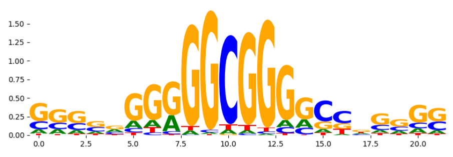 |
| pos_patterns.pattern_31 | 171 | 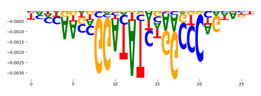 | 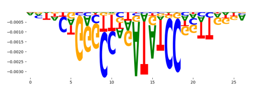 | FLI1_ETS_2 | 1.000000e+00 | 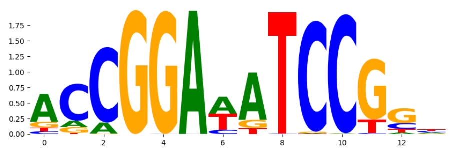 |
| pos_patterns.pattern_32 | 159 | 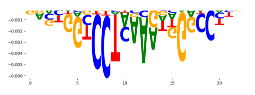 | 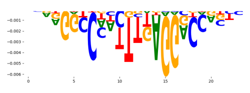 | HOXC12_MA0906.1 | 5.105430e-01 | 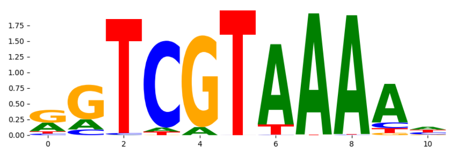 |
| pos_patterns.pattern_33 | 150 | 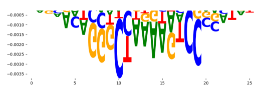 | 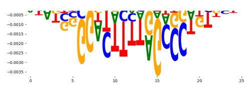 | Hoxc9_MA0485.1 | 6.949450e-01 | 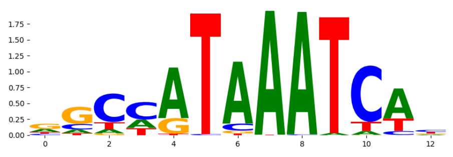 |
| pos_patterns.pattern_34 | 130 | 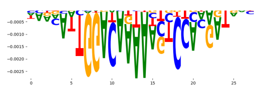 | 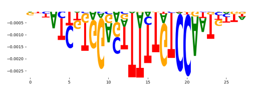 | THA11_MOUSE.H11MO.0.B | 6.762340e-01 | 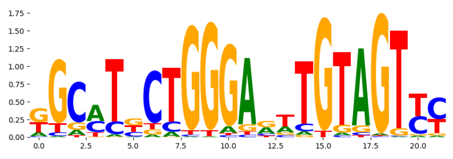 |
| pos_patterns.pattern_35 | 64 | 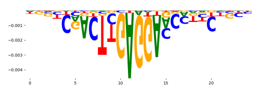 | 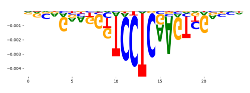 | RXRA+VDR_MA0074.1 | 1.000000e+00 | 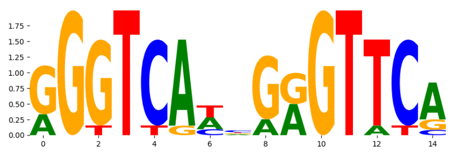 |
| pos_patterns.pattern_36 | 56 | 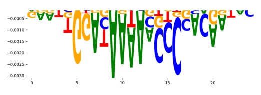 | 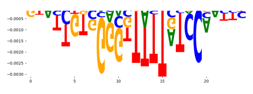 | ZN274_HUMAN.H11MO.0.A | 1.000000e+00 | 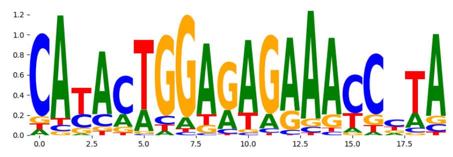 |
| pos_patterns.pattern_37 | 54 | 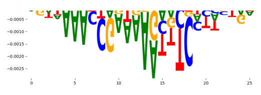 | 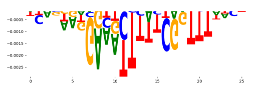 | IRF7_HUMAN.H11MO.0.C | 9.048840e-01 | 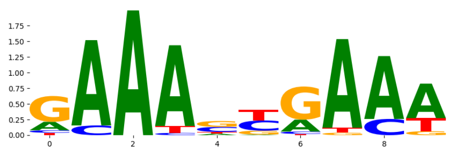 |
| pos_patterns.pattern_38 | 34 | 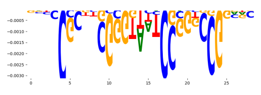 | 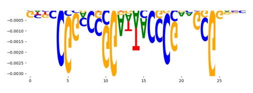 | SP1_MOUSE.H11MO.0.A | 2.611970e-01 | 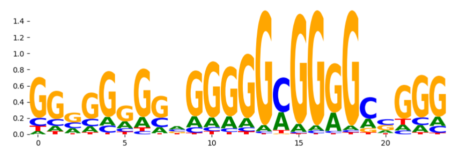 |
| neg_patterns.pattern_0 | 54 | 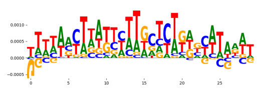 | 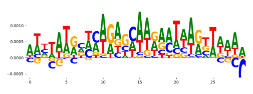 | ZN341_HUMAN.H11MO.0.C | 9.750740e-03 | 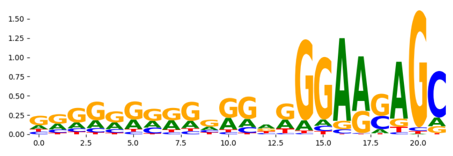 |
| neg_patterns.pattern_1 | 40 | 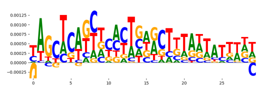 | 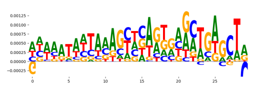 | SP1_MA0079.3 | 4.076930e-02 | 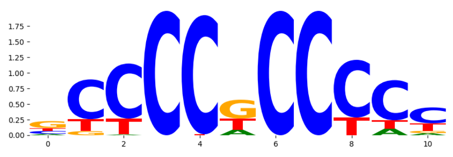 |
| neg_patterns.pattern_2 | 35 | 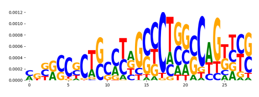 | 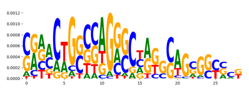 | TN5_1 | 6.171160e-03 | 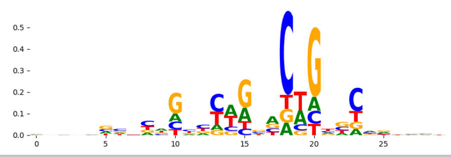 |
