## Supplementary Files 3 for "ChromBPNet: bias factorized, base-resolution deep learning models of chromatin accessibility reveal cis-regulatory sequence syntax, transcription factor footprints and regulatory variants": h1esc_ATAC_raw_bpnet_bias_fold3_profile_modisco.pdf

| pattern | num_seqlets | cwm_fwd / cwm_rev | TOMTOM_match | TOMTOM_qval | TOMTOM_match_logo |
|---|---|---|---|---|---|
| pos_patterns.pattern_0 | 8650 | 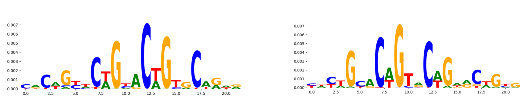 | TN5_1 | 1.589380e-08 | 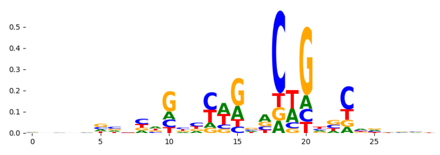 |
| pos_patterns.pattern_1 | 5671 | 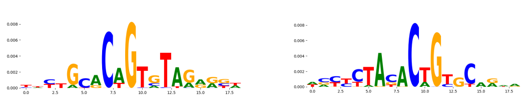 | TN5_8 | 4.565650e-02 | 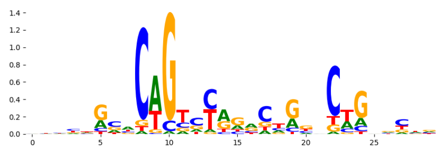 |
| pos_patterns.pattern_2 | 4599 | 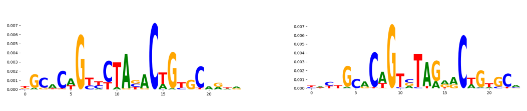 | TN5_1 | 9.031440e-06 | 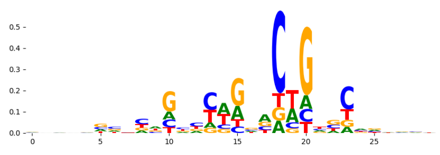 |
| pos_patterns.pattern_3 | 4067 | 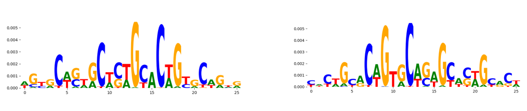 | TN5_2 | 1.602500e-10 | 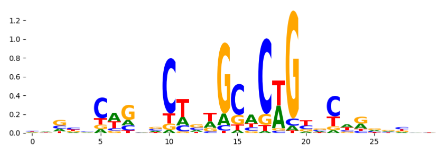 |
| pos_patterns.pattern_4 | 3103 | 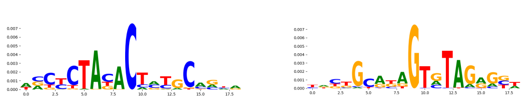 | KLF4_MA0039.3 | 1.511040e-01 | 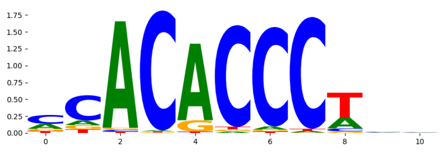 |
| pos_patterns.pattern_5 | 2835 | 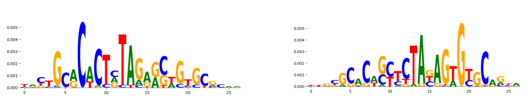 | TN5_3 | 2.571590e-18 | 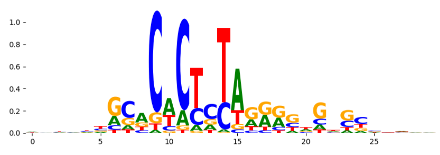 |
| pos_patterns.pattern_6 | 2306 | 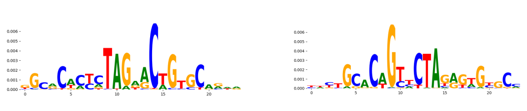 | TN5_3 | 2.587690e-05 | 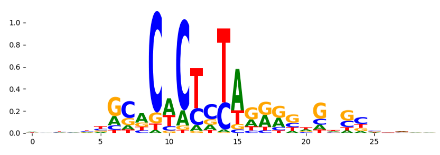 |
| pos_patterns.pattern_7 | 759 | 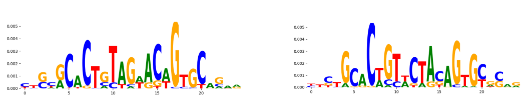 | TN5_7 | 1.719890e-06 | 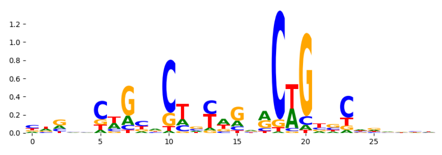 |
| pos_patterns.pattern_8 | 651 | 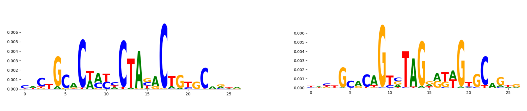 | TN5_3 | 1.172350e-04 | 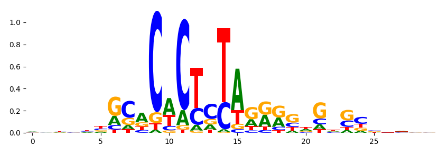 |
| pos_patterns.pattern_9 | 538 | 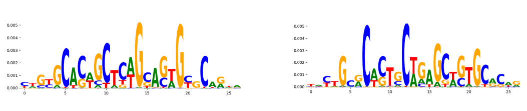 | TN5_3 | 1.437450e-05 | 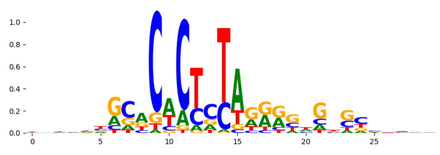 |
| pos_patterns.pattern_10 | 320 | 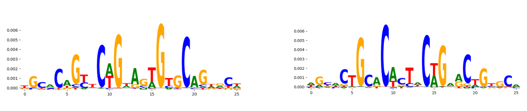 | TN5_1 | 3.460300e-06 | 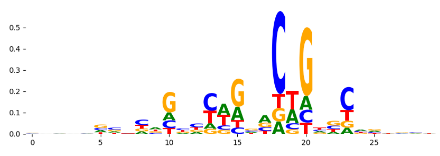 |
| pos_patterns.pattern_11 | 253 | 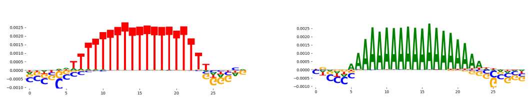 | PRDM6_HUMAN.H11MO.0.C | 6.236420e-02 | 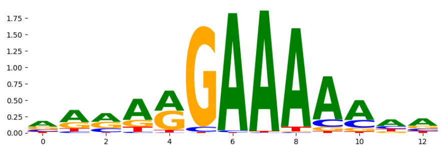 |
| pos_patterns.pattern_12 | 126 | 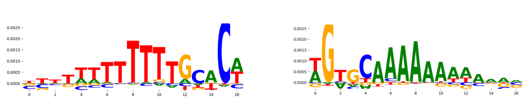 | Hoxc10.mouse_homeodomain_2 | 2.178580e-01 | 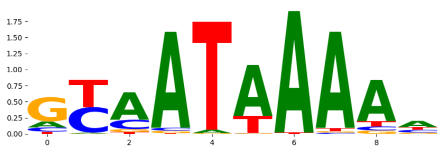 |
| pos_patterns.pattern_13 | 82 | 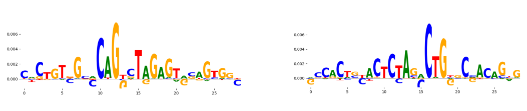 | TN5_6 | 9.169900e-17 | 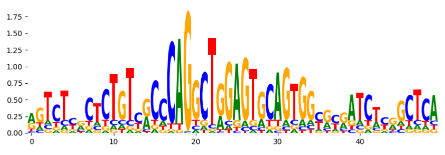 |
| pos_patterns.pattern_14 | 68 | 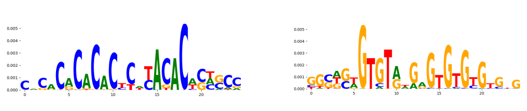 | EGR2_HUMAN.H11MO.0.A | 1.000000e+00 | 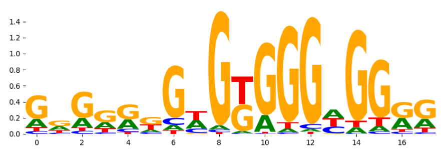 |
| pos_patterns.pattern_15 | 65 | 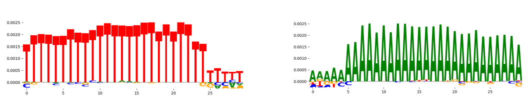 | PRDM6_HUMAN.H11MO.0.C | 1.124100e-01 | 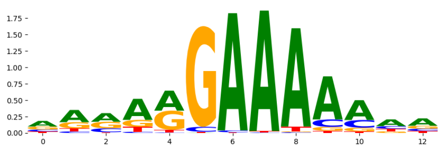 |
| neg_patterns.pattern_0 | 21 | 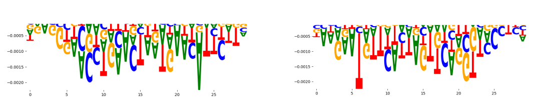 | POU3F3_POU_3 | 2.582030e-03 | 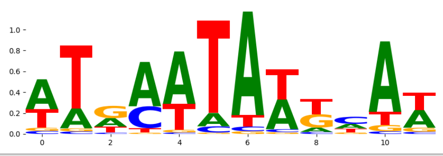 |
