## Supplementary Files 3 for "ChromBPNet: bias factorized, base-resolution deep learning models of chromatin accessibility reveal cis-regulatory sequence syntax, transcription factor footprints and regulatory variants": h1esc_ATAC_raw_bpnet_bias_fold4_counts_modisco.pdf

| pattern | num_seqlets | cwm_fwd | cwm_rev |
|---|---|---|---|
| pos_patterns.pattern_0 | 136 | 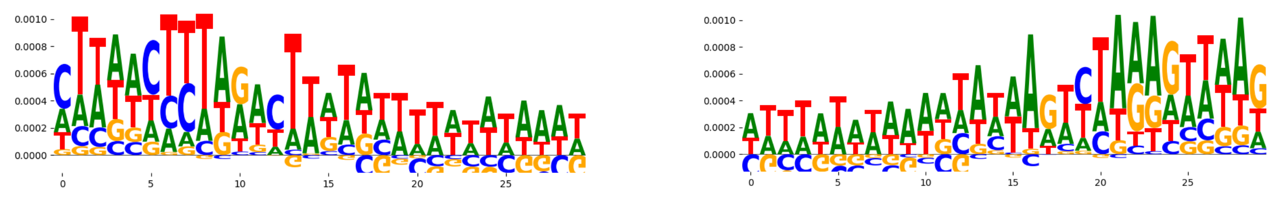 | |
| pos_patterns.pattern_1 | 127 | 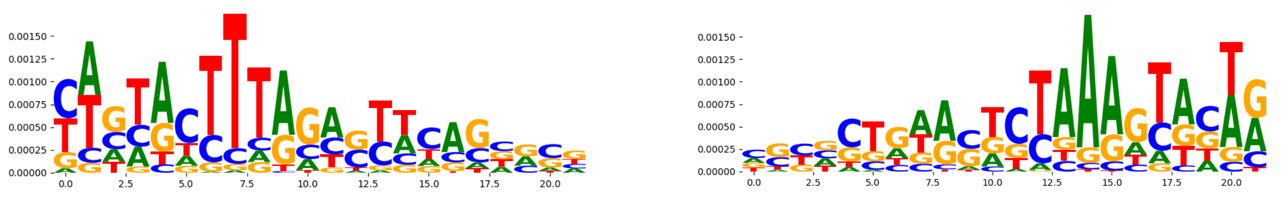 | |
| pos_patterns.pattern_2 | 81 | 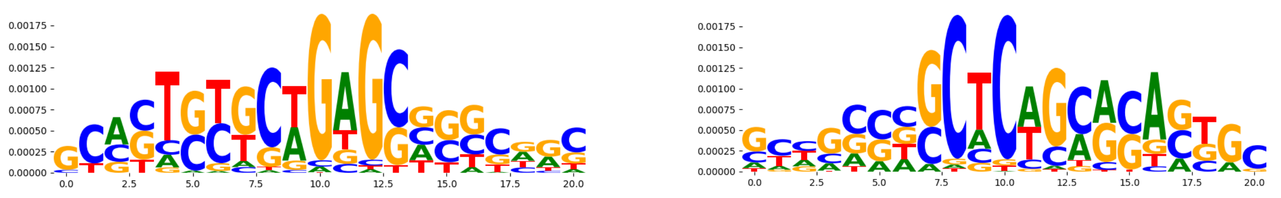 | |
| pos_patterns.pattern_3 | 79 | 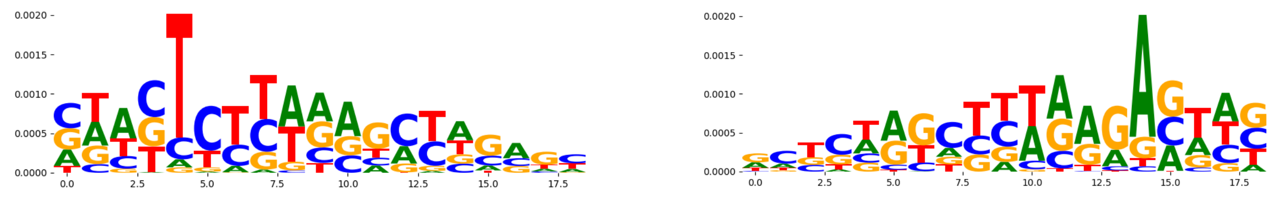 | |
| pos_patterns.pattern_4 | 67 | 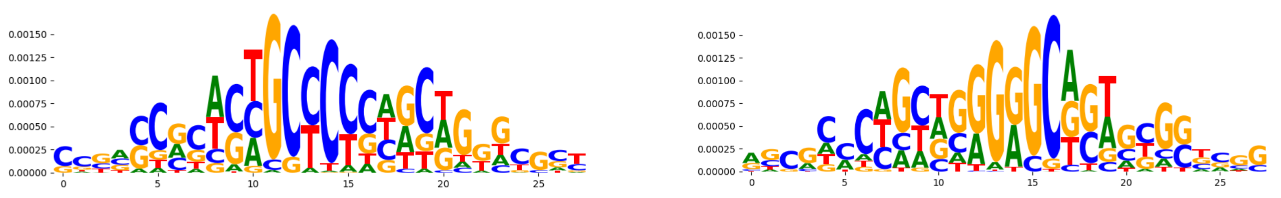 | |
| pos_patterns.pattern_5 | 64 | 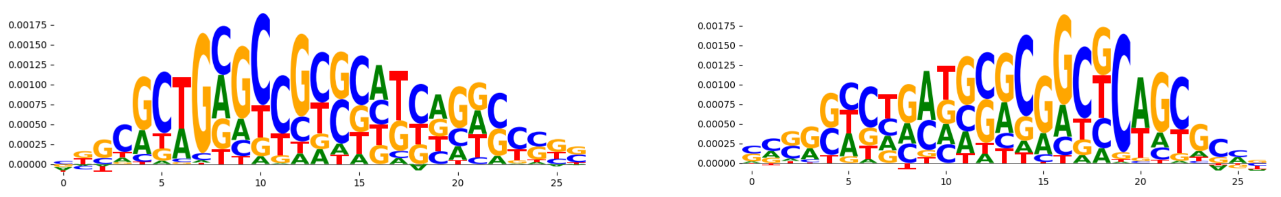 | |
| pos_patterns.pattern_6 | 61 | 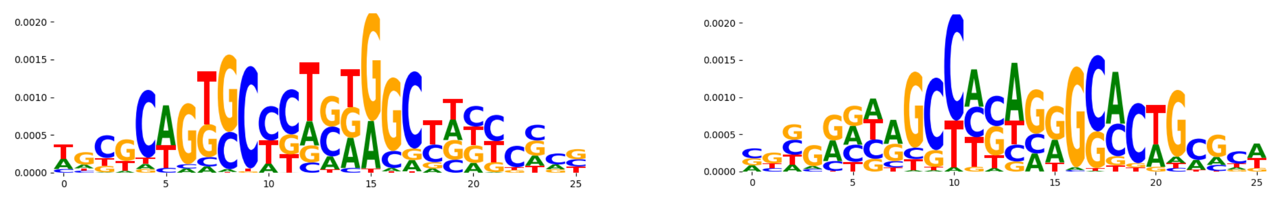 | |
| pos_patterns.pattern_7 | 55 | 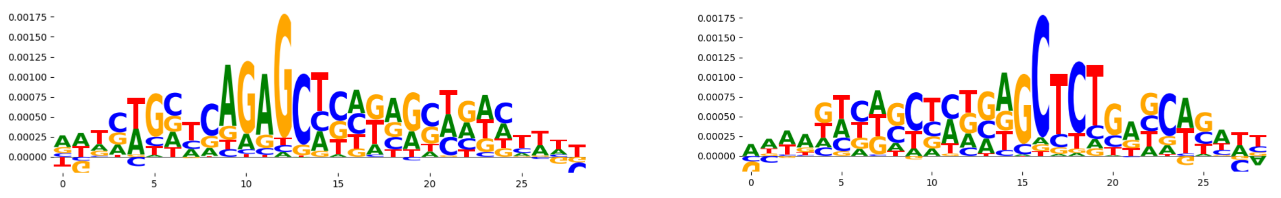 | |
| pos_patterns.pattern_8 | 45 | 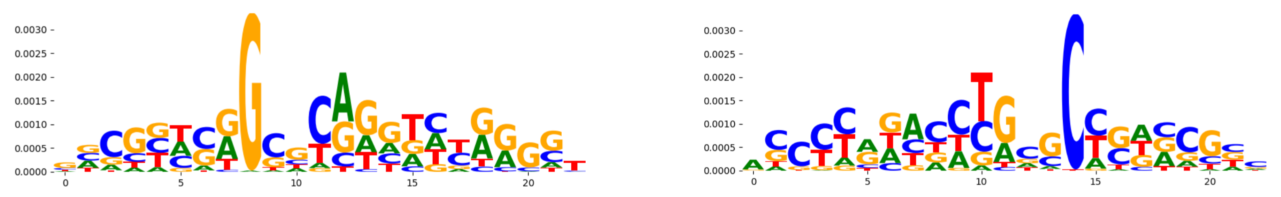 | |
| pos_patterns.pattern_9 | 45 | 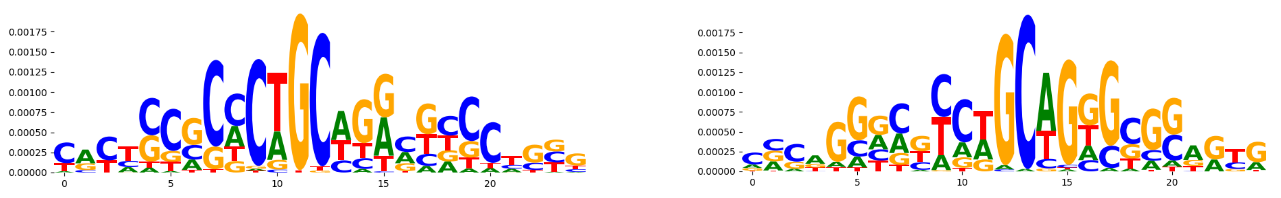 | |
| pos_patterns.pattern_10 | 44 | 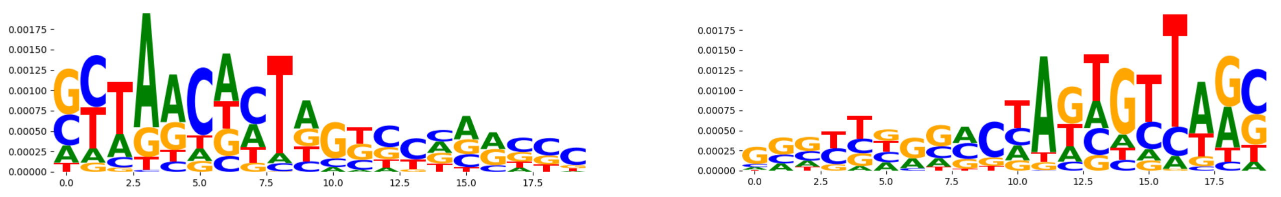 | |
| pos_patterns.pattern_11 | 28 | 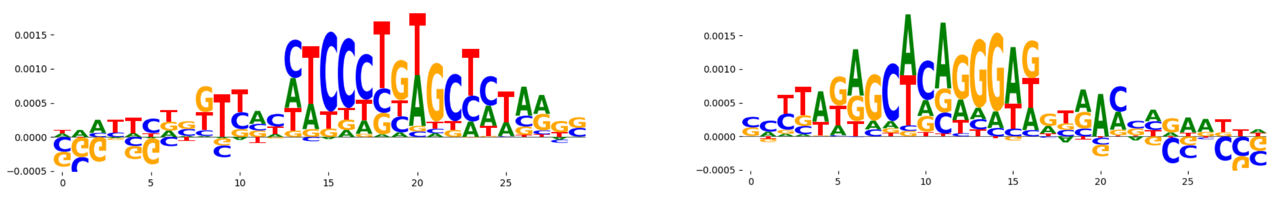 | |
| pos_patterns.pattern_12 | 22 | 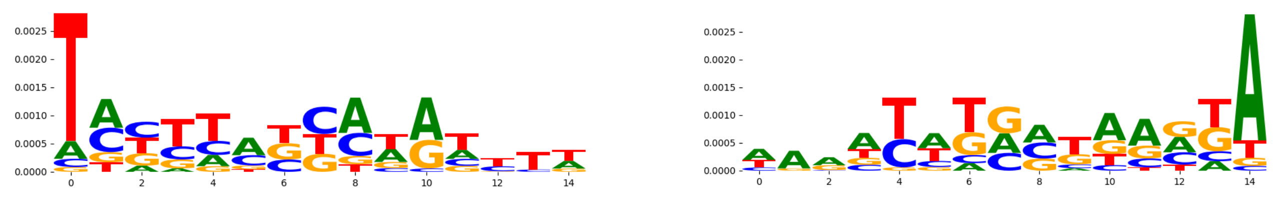 | |
| neg_patterns.pattern_0 | 2443 | 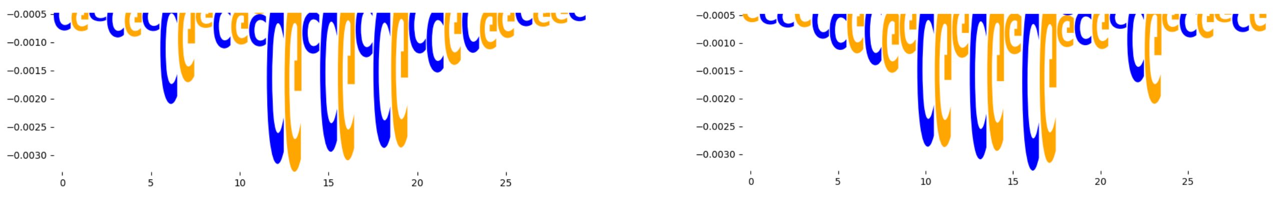 | |
| neg_patterns.pattern_1 | 2312 | 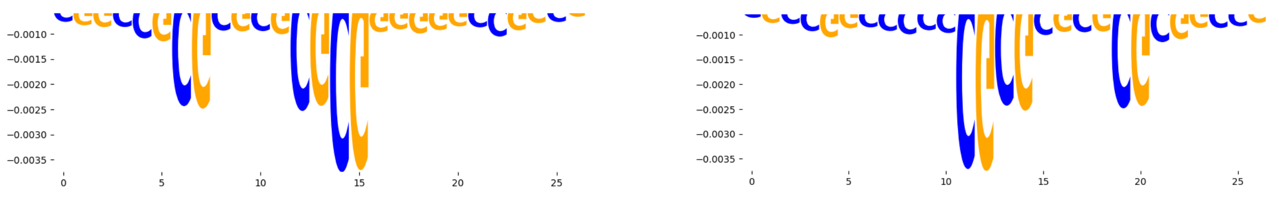 | |
| neg_patterns.pattern_2 | 1396 | 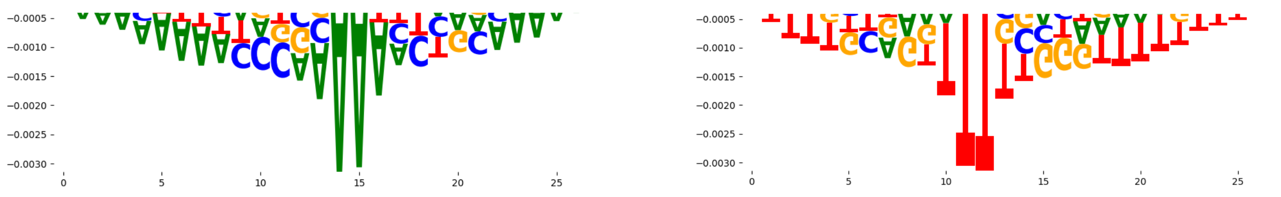 | |
| neg_patterns.pattern_3 | 1059 | 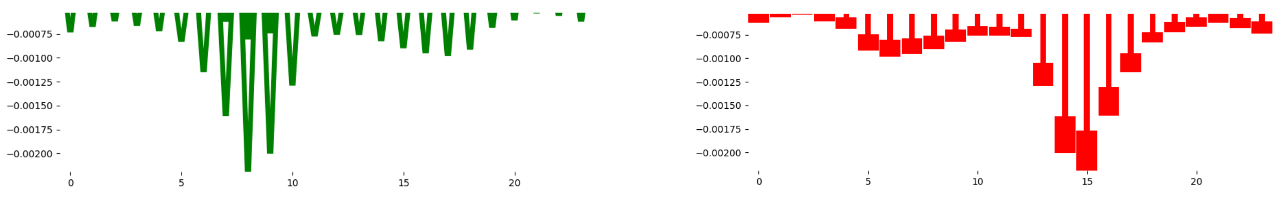 | |
| neg_patterns.pattern_4 | 953 | 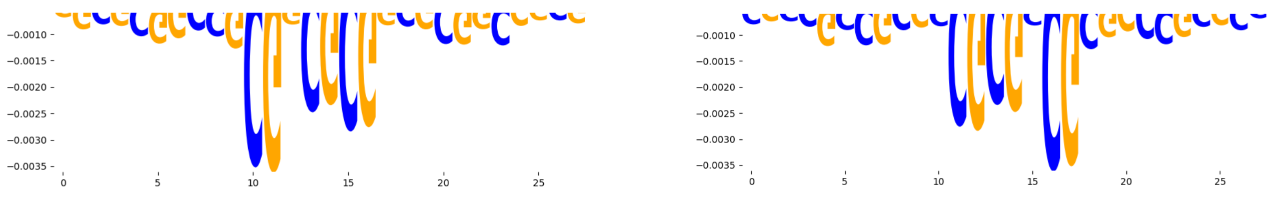 | |
| neg_patterns.pattern_5 | 788 | 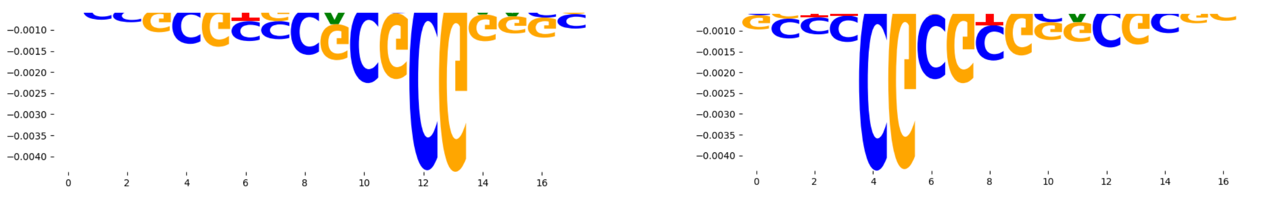 | |
| neg_patterns.pattern_6 | 737 | 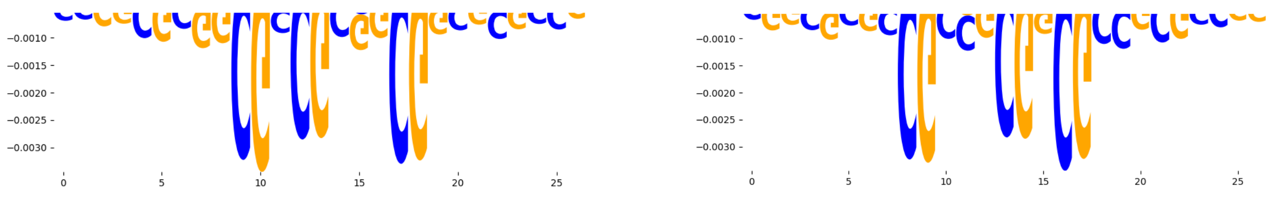 | |
| neg_patterns.pattern_7 | 440 | 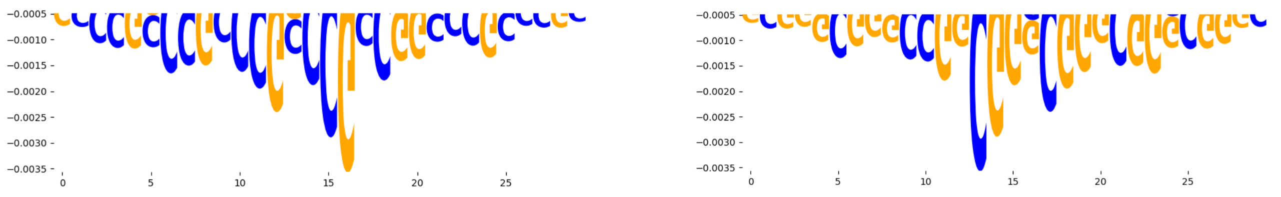 | |
| neg_patterns.pattern_8 | 400 | 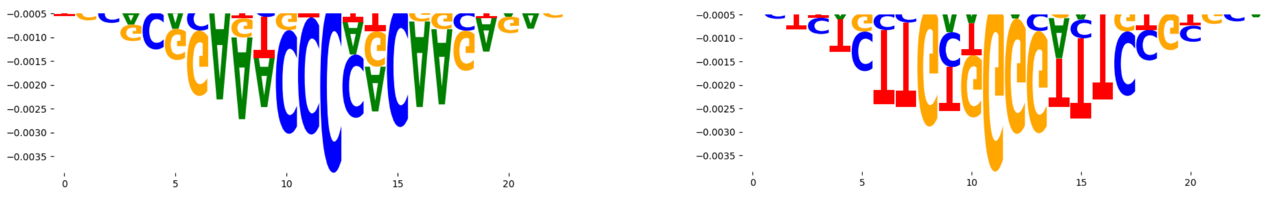 | |
| neg_patterns.pattern_9 | 373 | 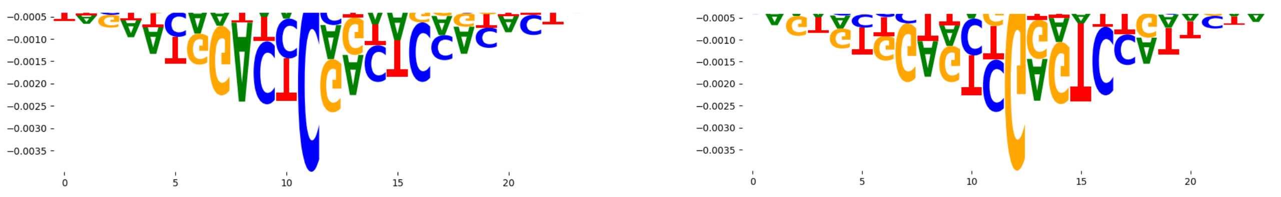 | |

| pattern | num_seqlets | cwm_fwd | cwm_rev |
| --- | --- | --- | --- |
| neg_patterns.pattern_10 | 369 | 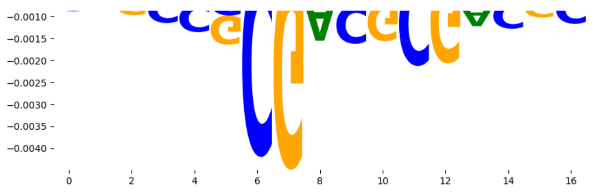 | 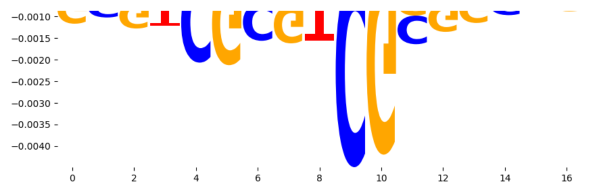 |
| neg_patterns.pattern_11 | 344 | 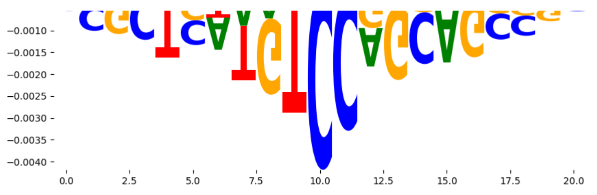 | 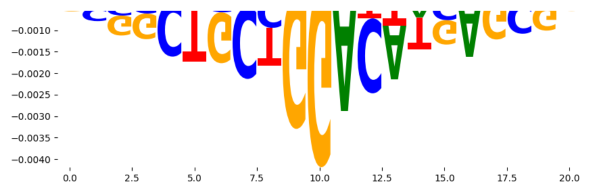 |
| neg_patterns.pattern_12 | 334 | 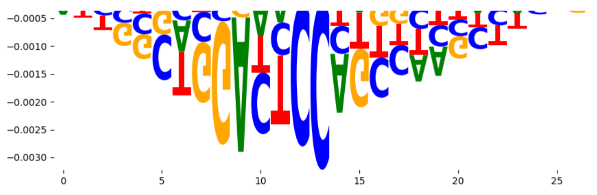 | 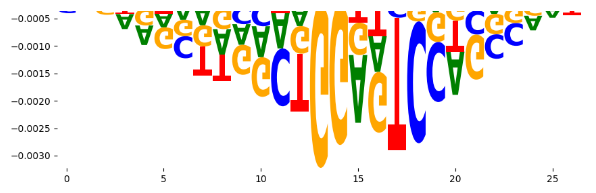 |
| neg_patterns.pattern_13 | 311 | 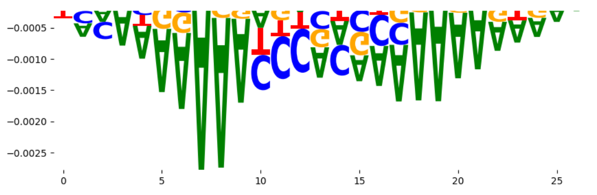 | 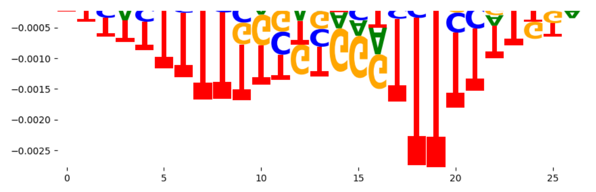 |
| neg_patterns.pattern_14 | 283 | 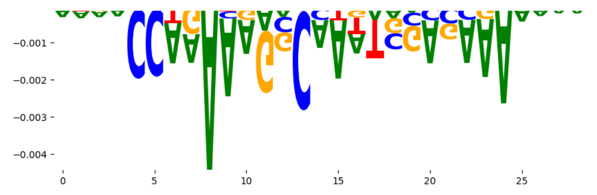 | 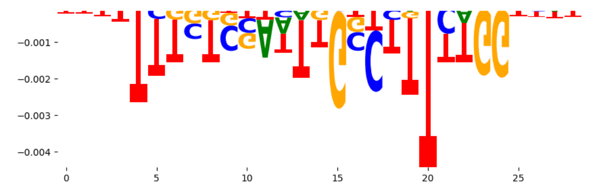 |
| neg_patterns.pattern_15 | 276 | 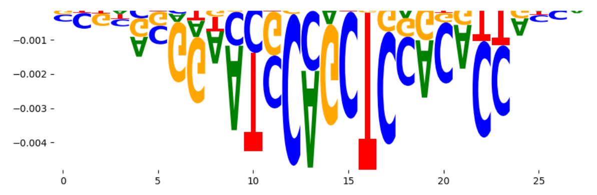 | 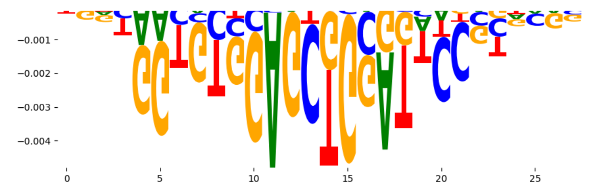 |
| neg_patterns.pattern_16 | 225 | 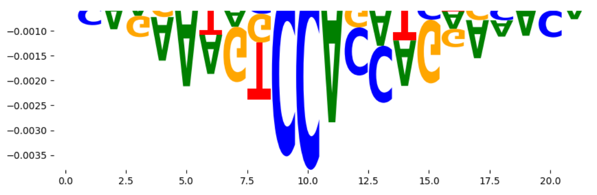 | 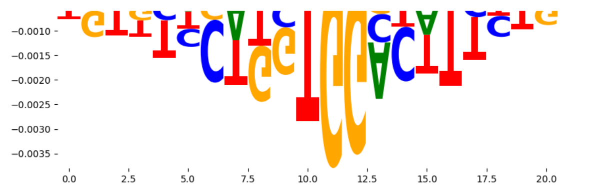 |
| neg_patterns.pattern_17 | 210 | 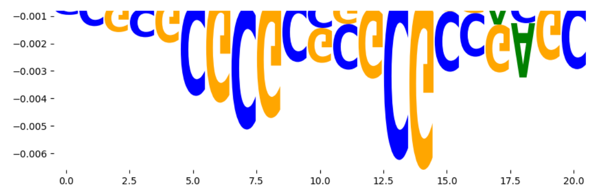 | 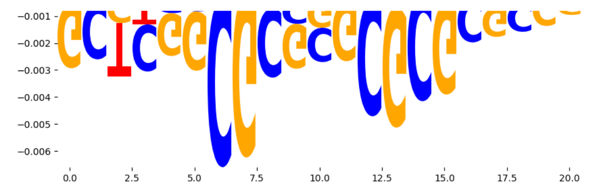 |
| neg_patterns.pattern_18 | 190 | 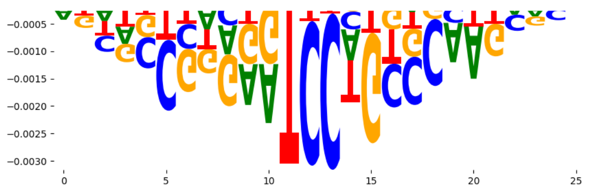 | 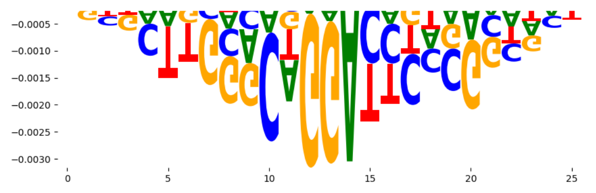 |
| neg_patterns.pattern_19 | 144 | 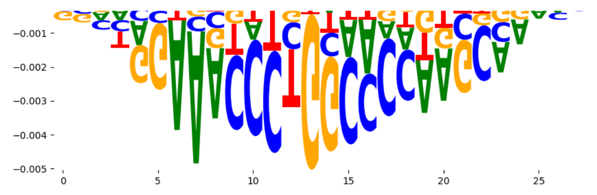 | 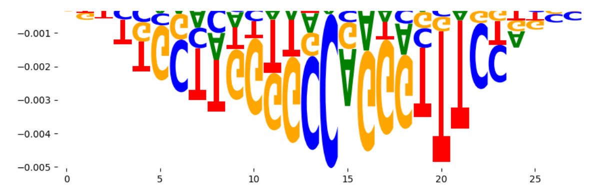 |
| neg_patterns.pattern_20 | 113 | 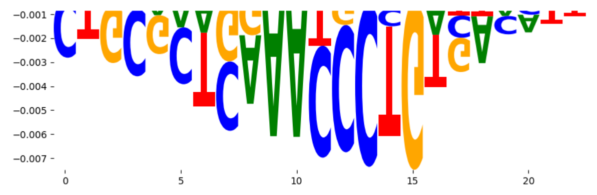 | 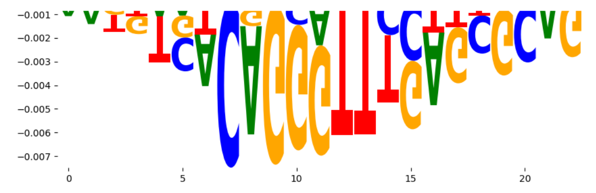 |
| neg_patterns.pattern_21 | 88 | 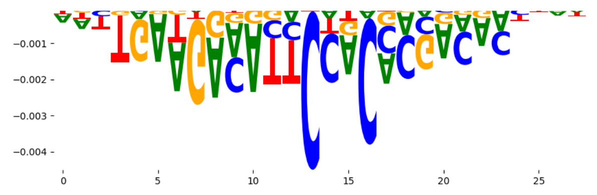 | 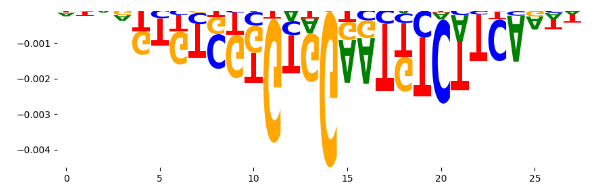 |
| neg_patterns.pattern_22 | 86 | 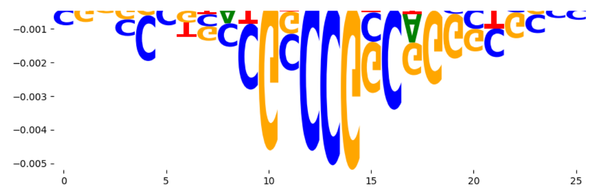 | 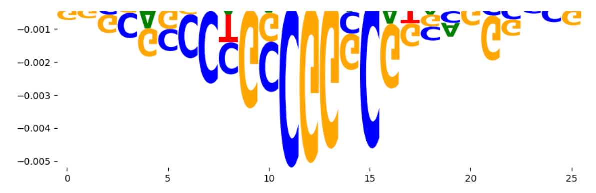 |
| neg_patterns.pattern_23 | 80 | 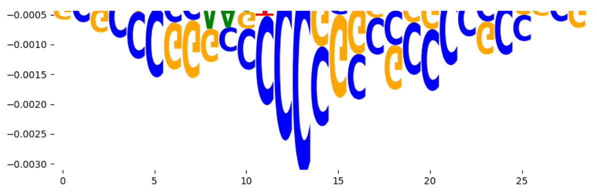 | 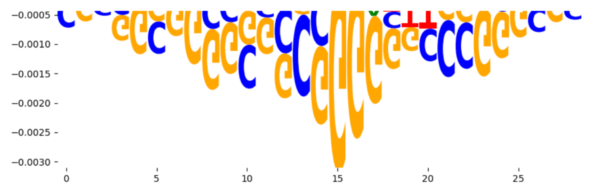 |
| neg_patterns.pattern_24 | 78 | 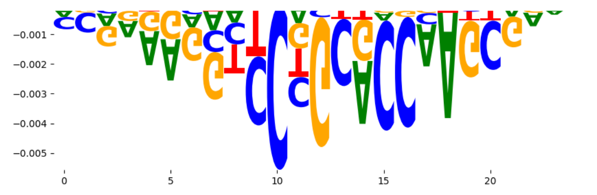 | 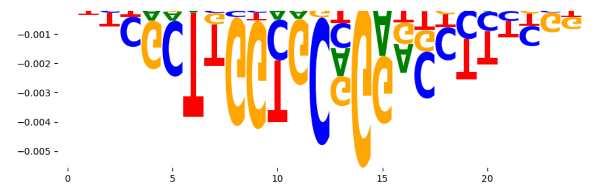 |
| neg_patterns.pattern_25 | 75 | 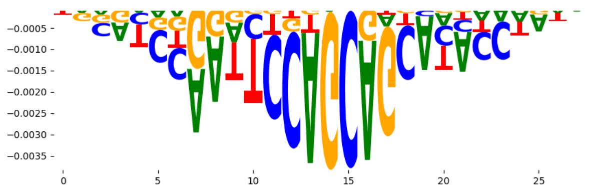 | 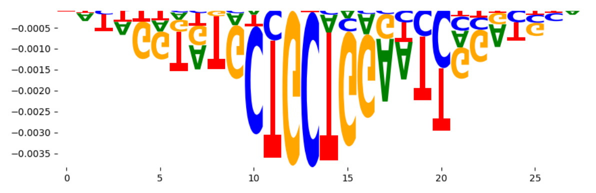 |
| neg_patterns.pattern_26 | 63 | 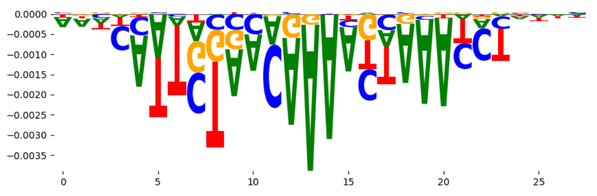 | 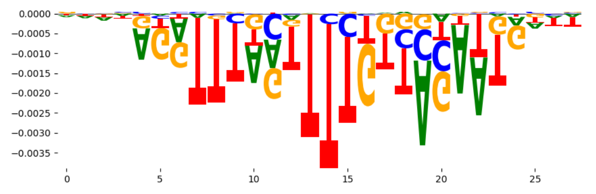 |
| neg_patterns.pattern_27 | 54 | 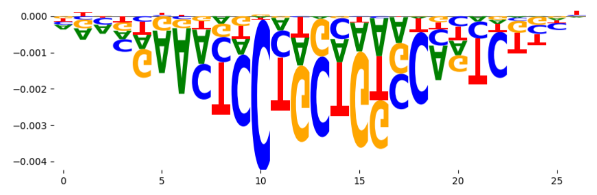 | 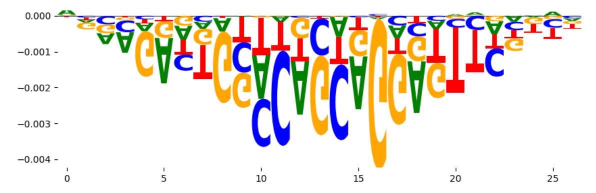 |
| neg_patterns.pattern_28 | 44 | 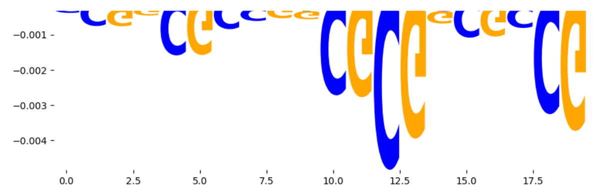 | 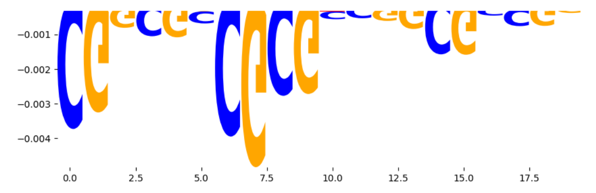 |
| neg_patterns.pattern_29 | 22 | 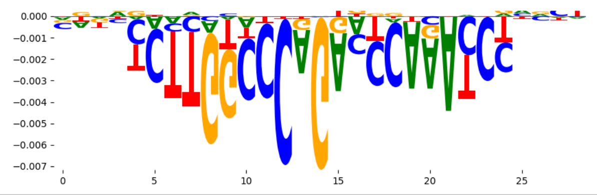 | 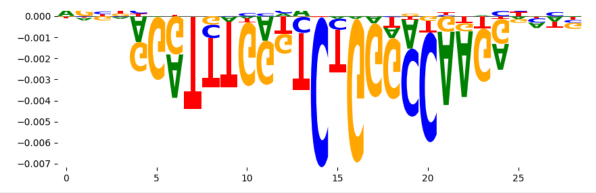 |
