## Supplementary Files 3 for "ChromBPNet: bias factorized, base-resolution deep learning models of chromatin accessibility reveal cis-regulatory sequence syntax, transcription factor footprints and regulatory variants": h1esc_ATAC_raw_bpnet_bias_fold4_profile_modisco.pdf

| pattern | num_seqlets | cwm_fwd | cwm_rev |
|---|---|---|---|
| pos_patterns.pattern_0 | 8668 | 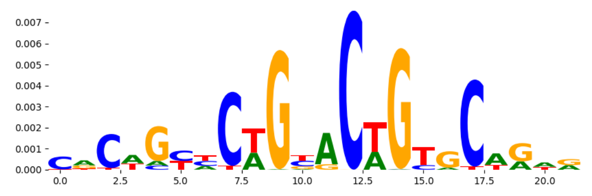 | 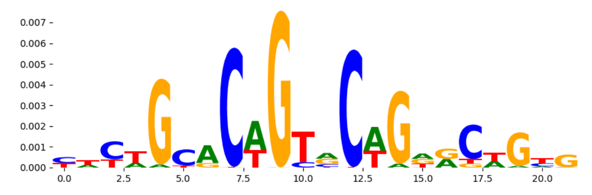 |
| pos_patterns.pattern_1 | 5608 | 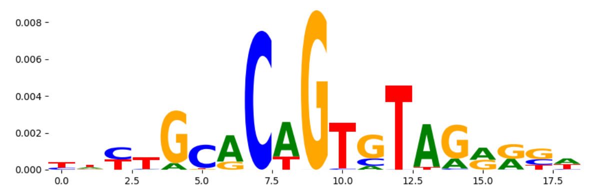 | 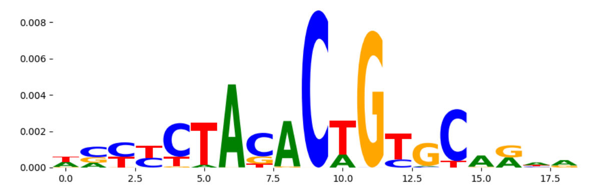 |
| pos_patterns.pattern_2 | 4298 | 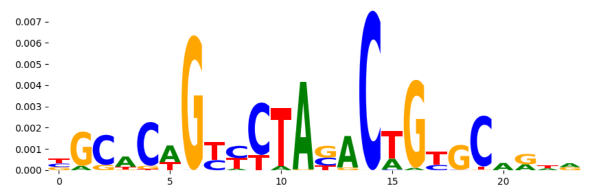 | 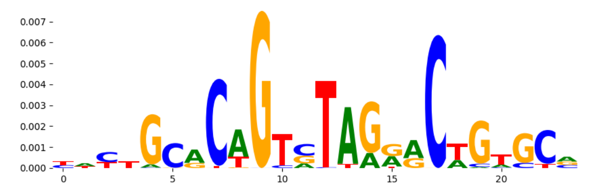 |
| pos_patterns.pattern_3 | 4027 | 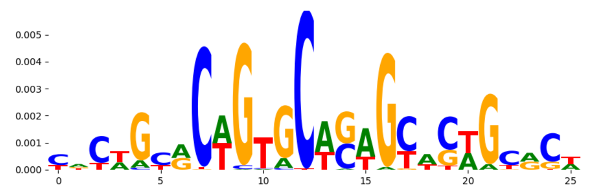 | 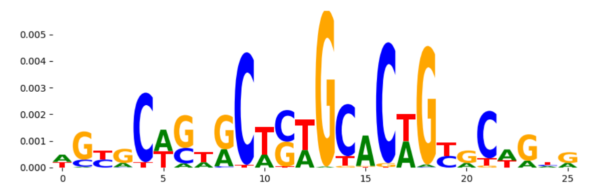 |
| pos_patterns.pattern_4 | 2732 | 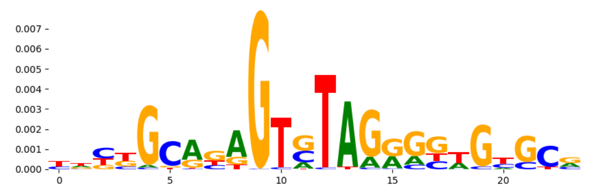 | 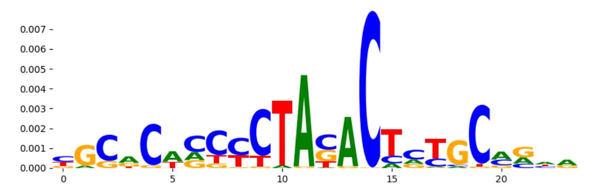 |
| pos_patterns.pattern_5 | 2347 | 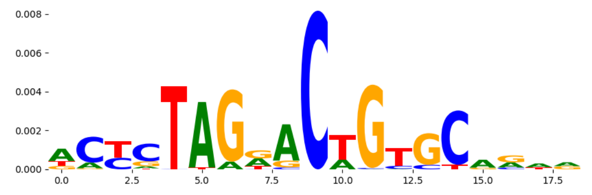 | 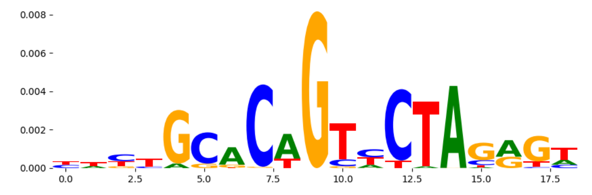 |
| pos_patterns.pattern_6 | 1908 | 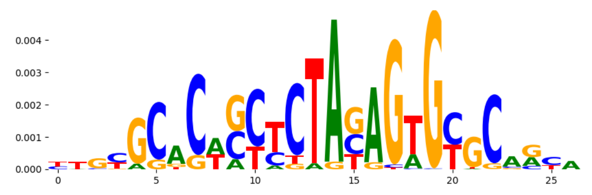 | 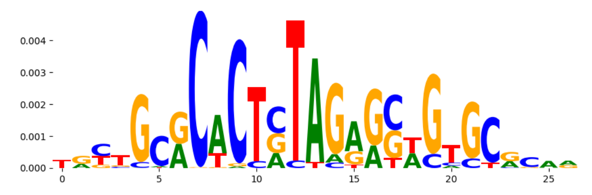 |
| pos_patterns.pattern_7 | 900 | 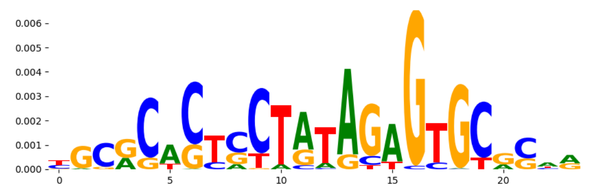 | 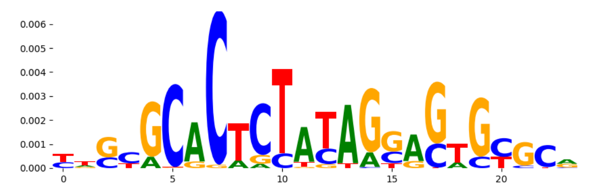 |
| pos_patterns.pattern_8 | 780 | 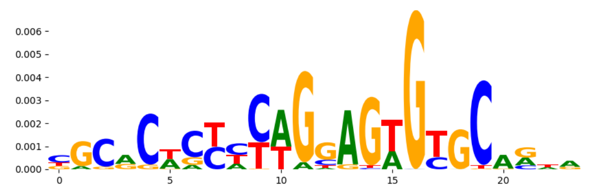 | 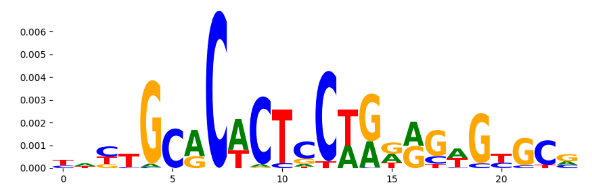 |
| pos_patterns.pattern_9 | 724 | 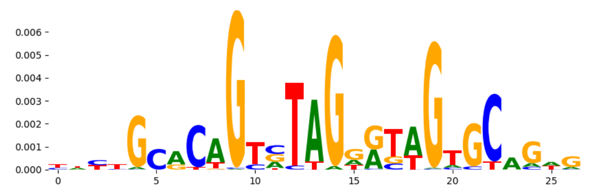 | 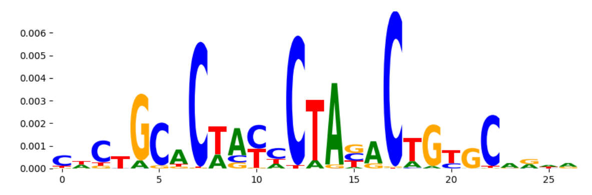 |
| pos_patterns.pattern_10 | 669 | 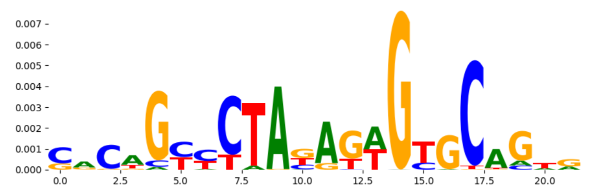 | 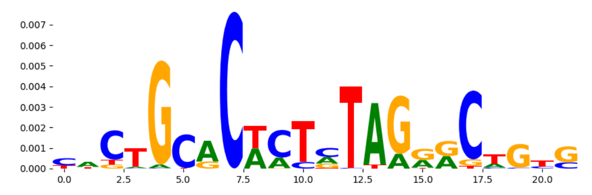 |
| pos_patterns.pattern_11 | 260 | 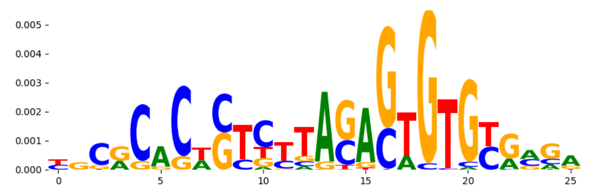 | 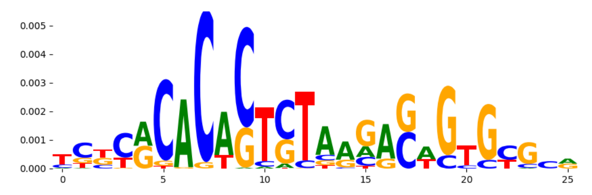 |
| pos_patterns.pattern_12 | 231 | 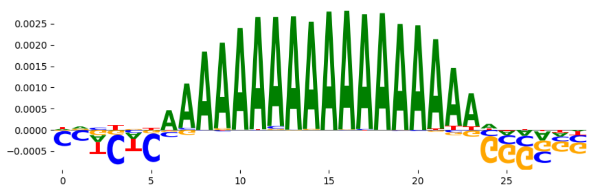 | 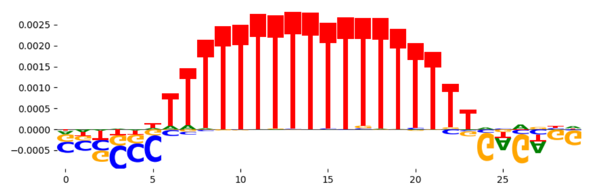 |
| pos_patterns.pattern_13 | 104 | 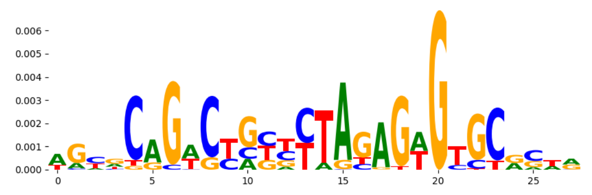 | 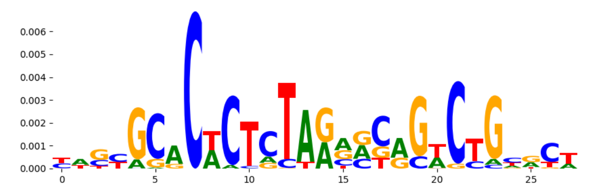 |
| pos_patterns.pattern_14 | 93 | 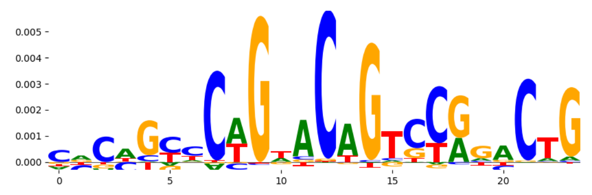 | 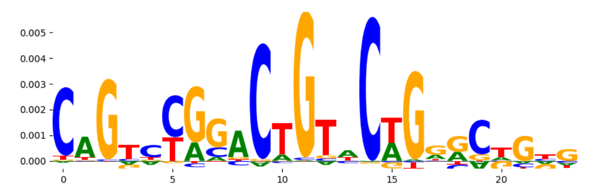 |
| pos_patterns.pattern_15 | 77 | 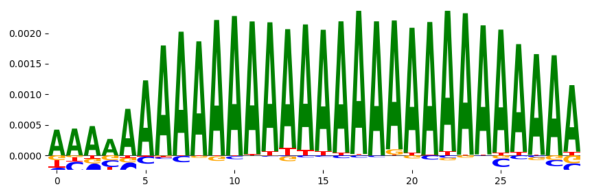 | 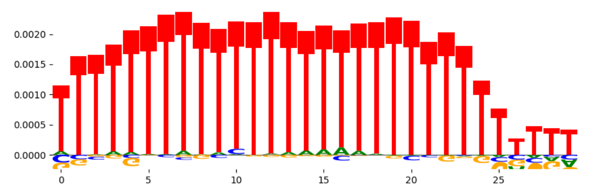 |
| pos_patterns.pattern_16 | 54 | 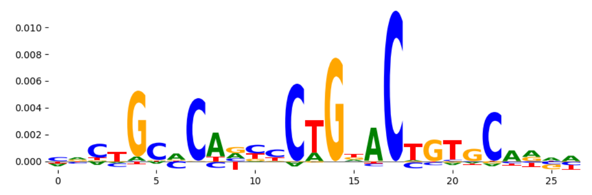 | 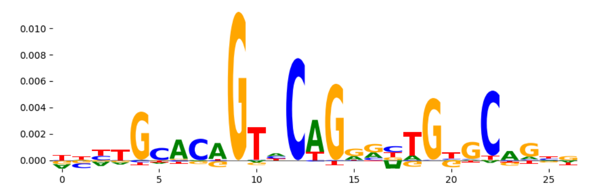 |
| pos_patterns.pattern_17 | 52 | 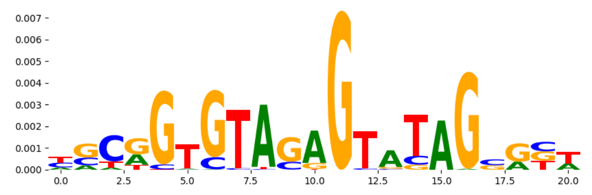 | 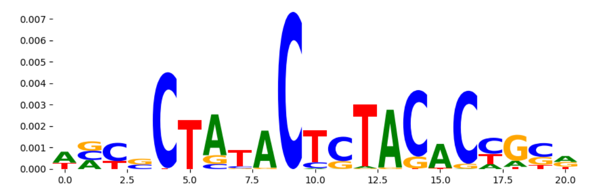 |
| pos_patterns.pattern_18 | 23 | 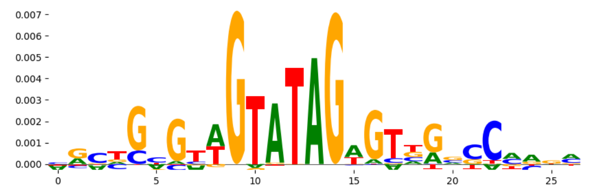 | 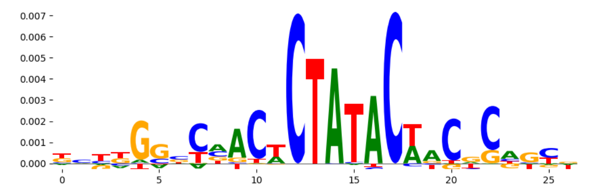 |
| pos_patterns.pattern_19 | 20 | 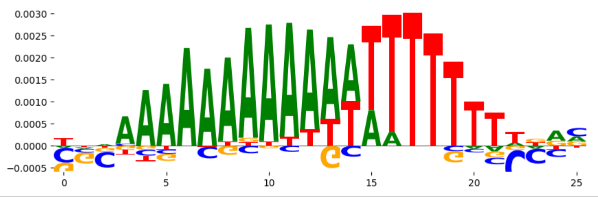 | 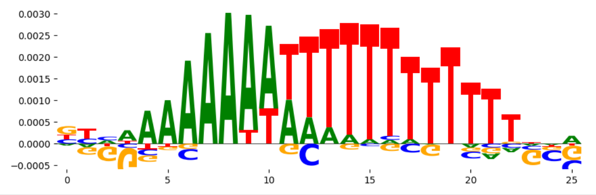 |
