## Supplementary Files 3 for "ChromBPNet: bias factorized, base-resolution deep learning models of chromatin accessibility reveal cis-regulatory sequence syntax, transcription factor footprints and regulatory variants": h1esc_DNASE_raw_bpnet_bias_fold0_counts_modisco.pdf

| pattern | num_seqlets | cwm_fwd | cwm_rev | TOMTOM_match | TOMTOM_qval | TOMTOM_match_logo |
|---|---|---|---|---|---|---|
| pos_patterns.pattern_0 | 10825 | | | CTCFL_HUMAN.H11MO.0.A | 7.392560e-06 | |
| pos_patterns.pattern_1 | 2181 | | | FOXB1_forkhead_2 | 3.328230e-01 | |
| pos_patterns.pattern_2 | 2160 | | | MEF2B_HUMAN.H11MO.0.A | 3.026800e-01 | |
| pos_patterns.pattern_3 | 1837 | | | TN5_6 | 8.274170e-02 | |
| pos_patterns.pattern_4 | 1759 | | | ZN770_HUMAN.H11MO.0.C | 4.594410e-06 | |
| pos_patterns.pattern_5 | 1657 | | | ZN770_HUMAN.H11MO.0.C | 5.683580e-08 | |
| pos_patterns.pattern_6 | 1460 | | | TN5_6 | 5.142120e-35 | |
| pos_patterns.pattern_7 | 1383 | | | SP3_HUMAN.H11MO.0.B | 4.393010e-02 | |
| pos_patterns.pattern_8 | 1323 | | | SP1_MOUSE.H11MO.0.A | 7.659200e-10 | |
| pos_patterns.pattern_9 | 1302 | | | TFAP2A_AP2_4 | 5.691270e-01 | |
| pos_patterns.pattern_10 | 1296 | | | ZNF384_MA1125.1 | 8.307610e-02 | |
| pos_patterns.pattern_11 | 1145 | | | DNASE_5 | 1.146680e-07 | |
| pos_patterns.pattern_12 | 731 | | | BARX1_MOUSE.H11MO.0.C | 7.613750e-01 | |
| pos_patterns.pattern_13 | 699 | | | ZN770_HUMAN.H11MO.0.C | 4.987530e-08 | |
| pos_patterns.pattern_14 | 595 | | | NFIC_MA0161.2 | 1.000000e+00 | |
| pos_patterns.pattern_15 | 559 | | | DNASE_5 | 1.451000e-10 | |
| pos_patterns.pattern_16 | 516 | | | DNASE_5 | 6.550720e-16 | |
| pos_patterns.pattern_17 | 464 | | | MNX1_MA0707.1 | 8.780570e-01 | |
| pos_patterns.pattern_18 | 403 | | | DNASE_5 | 2.485640e-15 | |
| pos_patterns.pattern_19 | 290 | | | ZN770_HUMAN.H11MO.0.C | 8.127400e-01 | |
| pos_patterns.pattern_20 | 260 | | | IKZF1_HUMAN.H11MO.0.C | 1.000000e+00 | |
| pos_patterns.pattern_21 | 219 | | | STAT1_MOUSE.H11MO.0.A | 7.357560e-02 | |
| pos_patterns.pattern_22 | 79 | | | DNASE_5 | 1.000000e+00 | |
| pos_patterns.pattern_23 | 49 | | | MAFB_HUMAN.H11MO.0.B | 1.000000e+00 | |
| pos_patterns.pattern_24 | 27 | | | RREB1_MA0073.1 | 1.000000e+00 | |

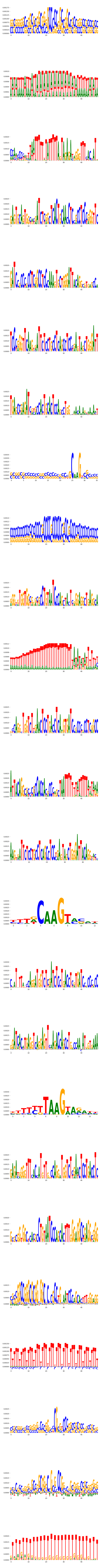

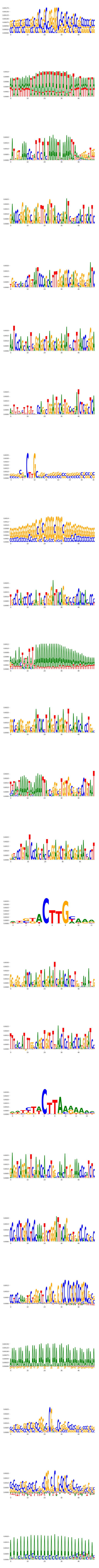

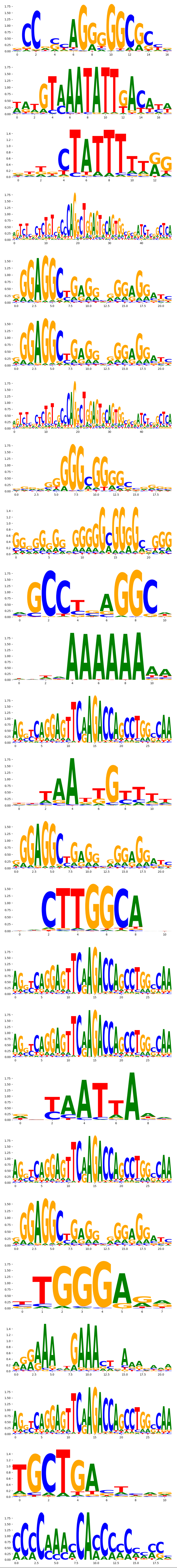
