## Supplementary Files 3 for "ChromBPNet: bias factorized, base-resolution deep learning models of chromatin accessibility reveal cis-regulatory sequence syntax, transcription factor footprints and regulatory variants": h1esc_DNASE_raw_bpnet_bias_fold1_counts_modisco.pdf

| pattern | num_seqlets | cwm_fwd | cwm_rev | TOMTOM_match | TOMTOM_qval | TOMTOM_match_logo |
|---|---|---|---|---|---|---|
| pos_patterns.pattern_0 | 4619 | | | CTCFL_MA1102.1 | 9.079450e-02 | |
| pos_patterns.pattern_1 | 4094 | | | NFIC_HUMAN.H11MO.0.A | 1.339040e-01 | |
| pos_patterns.pattern_2 | 3056 | | | SP1_HUMAN.H11MO.0.A | 6.217670e-03 | |
| pos_patterns.pattern_3 | 3030 | | | SP2_HUMAN.H11MO.0.A | 1.035430e-11 | |
| pos_patterns.pattern_4 | 885 | | | DNASE_1 | 1.100790e-01 | |
| pos_patterns.pattern_5 | 471 | | | DNASE_5 | 1.939700e-09 | |
| pos_patterns.pattern_6 | 354 | | | MEF2B_HUMAN.H11MO.0.A | 2.400440e-01 | |
| pos_patterns.pattern_7 | 315 | | | RARG_HUMAN.H11MO.0.B | 9.598430e-01 | |
| pos_patterns.pattern_8 | 311 | | | CPEB1_RRM_1 | 3.448310e-01 | |
| pos_patterns.pattern_9 | 210 | | | ZN250_HUMAN.H11MO.0.C | 1.000000e+00 | |
| pos_patterns.pattern_10 | 203 | | | VDR_MA0693.2 | 1.000000e+00 | |
| pos_patterns.pattern_11 | 196 | | | ZNF384_MA1125.1 | 3.157230e-02 | |
| pos_patterns.pattern_12 | 163 | | | ZN250_HUMAN.H11MO.0.C | 1.000000e+00 | |
| pos_patterns.pattern_13 | 158 | | | ITF2_HUMAN.H11MO.0.C | 1.000000e+00 | |
| pos_patterns.pattern_14 | 155 | | | PRDM6_HUMAN.H11MO.0.C | 1.070280e-01 | |
| pos_patterns.pattern_15 | 136 | | | EGR1_MOUSE.H11MO.0.A | 6.046960e-01 | |
| pos_patterns.pattern_16 | 54 | | | NaN | NaN | |
| neg_patterns.pattern_0 | 29 | | | SP3_HUMAN.H11MO.0.B | 3.641440e-03 | |
| neg_patterns.pattern_1 | 22 | | | CPEB1_RRM_1 | 5.270600e-02 | |
