## Supplementary Files 3 for "ChromBPNet: bias factorized, base-resolution deep learning models of chromatin accessibility reveal cis-regulatory sequence syntax, transcription factor footprints and regulatory variants": h1esc_DNASE_raw_bpnet_bias_fold2_profile_modisco.pdf

| pattern | num_seqlets | cwm_fwd | cwm_rev | TOMTOM_match | TOMTOM_qval | TOMTOM_match_logo |
|---|---|---|---|---|---|---|
| pos_patterns.pattern_0 | 4380 | | | ASCL1_MA1100.1 | 1.554320e-03 | |
| pos_patterns.pattern_1 | 3784 | | | SP1_MOUSE.H11MO.0.A | 1.540340e-01 | |
| pos_patterns.pattern_2 | 2636 | | | CTCFL_MOUSE.H11MO.0.A | 1.619870e-02 | |
| pos_patterns.pattern_3 | 2361 | | | SP1_MOUSE.H11MO.0.A | 2.901770e-11 | |
| pos_patterns.pattern_4 | 1020 | | | ERR2_HUMAN.H11MO.0.A | 1.000000e+00 | |
| pos_patterns.pattern_5 | 488 | | | DNASE_1 | 1.000000e+00 | |
| pos_patterns.pattern_6 | 472 | | | ZNF384_MA1125.1 | 6.900920e-02 | |
| pos_patterns.pattern_7 | 468 | | | DNASE_2 | 2.089980e-01 | |
| pos_patterns.pattern_8 | 374 | | | MEF2B_HUMAN.H11MO.0.A | 1.652510e-01 | |
| pos_patterns.pattern_9 | 310 | | | IKZF1_HUMAN.H11MO.0.C | 1.000000e+00 | |
| pos_patterns.pattern_10 | 303 | | | ZN770_HUMAN.H11MO.0.C | 6.116850e-01 | |
| pos_patterns.pattern_11 | 271 | | | NKX3-2_MA0122.2 | 6.596690e-01 | |
| pos_patterns.pattern_12 | 189 | | | DNASE_5 | 1.542050e-08 | |
| pos_patterns.pattern_13 | 128 | | | RREB1_MA0073.1 | 6.059040e-02 | |
| pos_patterns.pattern_14 | 102 | | | ZN770_HUMAN.H11MO.0.C | 2.338700e-07 | |
| pos_patterns.pattern_15 | 78 | | | ZN770_HUMAN.H11MO.0.C | 2.630420e-05 | |
| pos_patterns.pattern_16 | 48 | | | FOSL1_MA0477.1 | 4.572410e-01 | |
| pos_patterns.pattern_17 | 32 | | | KLF15_HUMAN.H11MO.0.A | 1.355650e-01 | |
| pos_patterns.pattern_18 | 23 | | | NKX28_HUMAN.H11MO.0.C | 1.000000e+00 | |
| neg_patterns.pattern_0 | 50 | | | ZN281_MOUSE.H11MO.0.A | 3.419830e-02 | |
| neg_patterns.pattern_1 | 26 | | | ZNF384_MA1125.1 | 3.666560e-03 | |
| neg_patterns.pattern_2 | 23 | | | ZNF384_MA1125.1 | 1.929630e-02 | |
