## Supplementary Files 3 for "ChromBPNet: bias factorized, base-resolution deep learning models of chromatin accessibility reveal cis-regulatory sequence syntax, transcription factor footprints and regulatory variants": h1esc_DNASE_raw_bpnet_bias_fold3_profile_modisco.pdf

| pattern | num_seqlets | cwm_fwd | cwm_rev | TOMTOM_match | TOMTOM_qval | TOMTOM_match_logo |
|---|---|---|---|---|---|---|
| pos_patterns.pattern_0 | 4651 | | | CTCFL_MOUSE.H11MO.0.A | 1.554180e-03 | |
| pos_patterns.pattern_1 | 3653 | | | PLAG1_MA0163.1 | 5.117070e-03 | |
| pos_patterns.pattern_2 | 2604 | | | SP1_MOUSE.H11MO.0.A | 8.787310e-08 | |
| pos_patterns.pattern_3 | 2312 | | | ZFX_MOUSE.H11MO.0.B | 8.513520e-05 | |
| pos_patterns.pattern_4 | 1836 | | | DNASE_1 | 1.000000e+00 | |
| pos_patterns.pattern_5 | 469 | | | IKZF1_HUMAN.H11MO.0.C | 1.000000e+00 | |
| pos_patterns.pattern_6 | 384 | | | DNASE_2 | 1.810630e-01 | |
| pos_patterns.pattern_7 | 364 | | | PRDM6_HUMAN.H11MO.0.C | 9.201670e-02 | |
| pos_patterns.pattern_8 | 361 | | | DNASE_2 | 2.128850e-01 | |
| pos_patterns.pattern_9 | 352 | | | DNASE_5 | 2.843870e-09 | |
| pos_patterns.pattern_10 | 292 | | | RARG_HUMAN.H11MO.0.B | 9.315280e-01 | |
| pos_patterns.pattern_11 | 228 | | | FOXJ3_HUMAN.H11MO.0.A | 3.502850e-01 | |
| pos_patterns.pattern_12 | 122 | | | ZN770_HUMAN.H11MO.0.C | 9.429680e-03 | |
| pos_patterns.pattern_13 | 86 | | | EGR1_MOUSE.H11MO.0.A | 5.061020e-01 | |
| pos_patterns.pattern_14 | 55 | | | Zfx_MA0146.2 | 9.806720e-01 | |
| pos_patterns.pattern_15 | 45 | | | TN5_6 | 1.845540e-21 | |
