## Supplementary Files 3 for "ChromBPNet: bias factorized, base-resolution deep learning models of chromatin accessibility reveal cis-regulatory sequence syntax, transcription factor footprints and regulatory variants": h1esc_DNASE_raw_bpnet_bias_fold4_profile_modisco.pdf

| pattern | num_seqlets | cwm_fwd | cwm_rev | TOMTOM_match | TOMTOM_qval | TOMTOM_match_logo |
|---|---|---|---|---|---|---|
| pos_patterns.pattern_0 | 6559 | | | CTCFL_HUMAN.H11MO.0.A | 8.883550e-02 | |
| pos_patterns.pattern_1 | 4210 | | | ERR2_HUMAN.H11MO.0.A | 1.000000e+00 | |
| pos_patterns.pattern_2 | 1806 | | | PATZ1_HUMAN.H11MO.0.C | 7.385580e-08 | |
| pos_patterns.pattern_3 | 1326 | | | MYOD1_HUMAN.H11MO.0.A | 2.152600e-02 | |
| pos_patterns.pattern_4 | 780 | | | FOXC1_HUMAN.H11MO.0.C | 6.688190e-01 | |
| pos_patterns.pattern_5 | 663 | | | IKZF1_HUMAN.H11MO.0.C | 9.924120e-01 | |
| pos_patterns.pattern_6 | 446 | | | MEF2B_HUMAN.H11MO.0.A | 2.918930e-01 | |
| pos_patterns.pattern_7 | 391 | | | RARG_HUMAN.H11MO.0.B | 6.618960e-01 | |
| pos_patterns.pattern_8 | 358 | | | ZNF384_MA1125.1 | 3.570010e-01 | |
| pos_patterns.pattern_9 | 300 | | | NaN | NaN | |
| pos_patterns.pattern_10 | 262 | | | PRDM6_HUMAN.H11MO.0.C | 6.703000e-02 | |
| pos_patterns.pattern_11 | 221 | | | ZN770_HUMAN.H11MO.0.C | 1.589580e-08 | |
| pos_patterns.pattern_12 | 207 | | | PRDM6_HUMAN.H11MO.0.C | 1.030610e-01 | |
| pos_patterns.pattern_13 | 180 | | | FOS_HUMAN.H11MO.0.A | 5.536570e-01 | |
| pos_patterns.pattern_14 | 92 | | | ZN770_HUMAN.H11MO.0.C | 2.129290e-07 | |
| pos_patterns.pattern_15 | 60 | | | IRF3_HUMAN.H11MO.0.B | 6.059640e-03 | |
| pos_patterns.pattern_16 | 50 | | | PRDM6_HUMAN.H11MO.0.C | 3.859810e-02 | |
| neg_patterns.pattern_0 | 39 | | | ZNF384_MA1125.1 | 1.707810e-01 | |
| neg_patterns.pattern_1 | 32 | | | ZNF384_MA1125.1 | 4.840300e-02 | |
| neg_patterns.pattern_2 | 29 | | | FOXD2_forkhead_1 | 1.671690e-01 | |
