## Supplementary Files 3 for "ChromBPNet: bias factorized, base-resolution deep learning models of chromatin accessibility reveal cis-regulatory sequence syntax, transcription factor footprints and regulatory variants": hepg2_ATAC_raw_bpnet_bias_fold0_counts_modisco.pdf

| pattern | num_seqlets | cwm_fwd | cwm_rev | TOMTOM_match | TOMTOM_qval | TOMTOM_match_logo |
|---|---|---|---|---|---|---|
| pos_patterns.pattern_0 | 2566 | | | SP2_HUMAN.H11MO.0.A | 1.395720e-07 | |
| pos_patterns.pattern_1 | 2393 | | | PRDM4_C2H2_1 | 8.578400e-02 | |
| pos_patterns.pattern_2 | 2215 | | | FOXC1_forkhead_1 | 2.983140e-02 | |
| pos_patterns.pattern_3 | 1327 | | | Hoxc9_MA0485.1 | 1.000000e+00 | |
| pos_patterns.pattern_4 | 1290 | | | TFAP2C_TFAP_5 | 1.000000e+00 | |
| pos_patterns.pattern_5 | 1105 | | | ONECUT1_CUT_1 | 1.099600e-01 | |
| pos_patterns.pattern_6 | 950 | | | FOXG1_forkhead_1 | 5.448140e-01 | |
| pos_patterns.pattern_7 | 875 | | | ZFX_MOUSE.H11MO.0.B | 3.373050e-05 | |
| pos_patterns.pattern_8 | 875 | | | SP1_HUMAN.H11MO.0.A | 7.741840e-03 | |
| pos_patterns.pattern_9 | 858 | | | RXRA_MOUSE.H11MO.0.A | 1.086590e-01 | |
| pos_patterns.pattern_10 | 663 | | | AP2B_HUMAN.H11MO.0.B | 7.934560e-02 | |
| pos_patterns.pattern_11 | 525 | | | HNF4A_nuclearreceptor_3 | 1.000000e+00 | |
| pos_patterns.pattern_12 | 492 | | | SP1_HUMAN.H11MO.0.A | 5.765120e-03 | |
| pos_patterns.pattern_13 | 483 | | | TGIF1_HUMAN.H11MO.0.A | 1.000000e+00 | |
| pos_patterns.pattern_14 | 315 | | | PBX3_MA1114.1 | 1.000000e+00 | |
| pos_patterns.pattern_15 | 259 | | | SP1_HUMAN.H11MO.0.A | 2.164690e-02 | |
| pos_patterns.pattern_16 | 196 | | | SP1_HUMAN.H11MO.0.A | 1.598130e-06 | |
| pos_patterns.pattern_17 | 192 | | | ZBTB6_HUMAN.H11MO.0.C | 2.099980e-04 | |
| pos_patterns.pattern_18 | 70 | | | RREB1_MA0073.1 | 1.000000e+00 | |
| pos_patterns.pattern_19 | 68 | | | COE1_HUMAN.H11MO.0.A | 4.978620e-01 | |
| neg_patterns.pattern_0 | 1073 | | | TN5_2 | 6.656500e-04 | |
| neg_patterns.pattern_1 | 302 | | | SP1_MOUSE.H11MO.0.A | 1.162680e-02 | |
| neg_patterns.pattern_2 | 164 | | | TN5_2 | 1.037090e-01 | |
| neg_patterns.pattern_3 | 152 | | | ZN331_HUMAN.H11MO.0.C | 8.921770e-01 | |
| neg_patterns.pattern_4 | 151 | | | TN5_2 | 2.758750e-02 | |
| neg_patterns.pattern_5 | 128 | | | MXI1_HUMAN.H11MO.0.A | 1.090290e-01 | |
| neg_patterns.pattern_6 | 87 | | | TN5_2 | 1.204770e-04 | |
| neg_patterns.pattern_7 | 25 | | | ZN331_HUMAN.H11MO.0.C | 2.581190e-02 | |
| neg_patterns.pattern_8 | 24 | | | TN5_7 | 6.477340e-01 | |
