## Supplementary Files 3 for "ChromBPNet: bias factorized, base-resolution deep learning models of chromatin accessibility reveal cis-regulatory sequence syntax, transcription factor footprints and regulatory variants": hepg2_ATAC_raw_bpnet_bias_fold0_profile_modisco.pdf

| pattern | num_seqlets | cwm_fwd | cwm_rev | TOMTOM_match | TOMTOM_qval | TOMTOM_match_logo |
|---|---|---|---|---|---|---|
| pos_patterns.pattern_0 | 8439 | | | TN5_1 | 2.849130e-09 | |
| pos_patterns.pattern_1 | 5533 | | | TN5_4 | 1.550140e-02 | |
| pos_patterns.pattern_2 | 3691 | | | TN5_2 | 1.503020e-18 | |
| pos_patterns.pattern_3 | 3616 | | | TN5_1 | 1.840240e-08 | |
| pos_patterns.pattern_4 | 2606 | | | TN5_3 | 9.716780e-02 | |
| pos_patterns.pattern_5 | 904 | | | TN5_3 | 2.618600e-03 | |
| pos_patterns.pattern_6 | 877 | | | TN5_3 | 1.583450e-06 | |
| pos_patterns.pattern_7 | 759 | | | TN5_3 | 4.058660e-11 | |
| pos_patterns.pattern_8 | 758 | | | TN5_7 | 1.226630e-03 | |
| pos_patterns.pattern_9 | 592 | | | TN5_3 | 1.456730e-06 | |
| pos_patterns.pattern_10 | 533 | | | TN5_3 | 1.134060e-04 | |
| pos_patterns.pattern_11 | 359 | | | TN5_4 | 2.923300e-06 | |
| pos_patterns.pattern_12 | 353 | | | PRDM6_HUMAN.H11MO.0.C | 7.689710e-02 | |
| pos_patterns.pattern_13 | 302 | | | TN5_6 | 4.560730e-17 | |
| pos_patterns.pattern_14 | 80 | | | MTF1_HUMAN.H11MO.0.C | 5.953400e-01 | |
| pos_patterns.pattern_15 | 74 | | | TN5_8 | 2.289830e-04 | |
| pos_patterns.pattern_16 | 72 | | | EGR2_HUMAN.H11MO.0.A | 1.000000e+00 | |
| pos_patterns.pattern_17 | 45 | | | TN5_7 | 5.396470e-06 | |
