## Supplementary Files 3 for "ChromBPNet: bias factorized, base-resolution deep learning models of chromatin accessibility reveal cis-regulatory sequence syntax, transcription factor footprints and regulatory variants": hepg2_ATAC_raw_bpnet_bias_fold1_counts_modisco.pdf

| pattern | num_seqlets | cwm_fwd | cwm_rev |
| --- | --- | --- | --- |
| pos_patterns.pattern_0 | 693 | 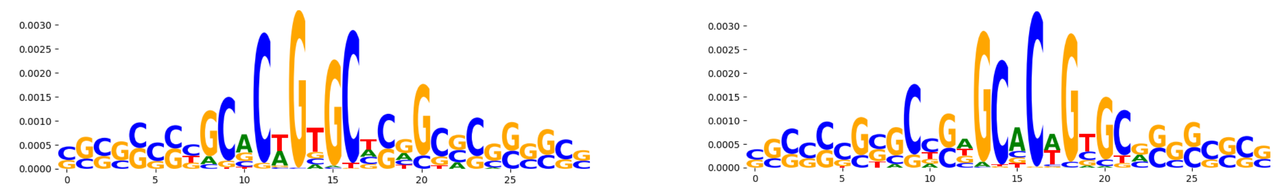 | |
| pos_patterns.pattern_1 | 478 | 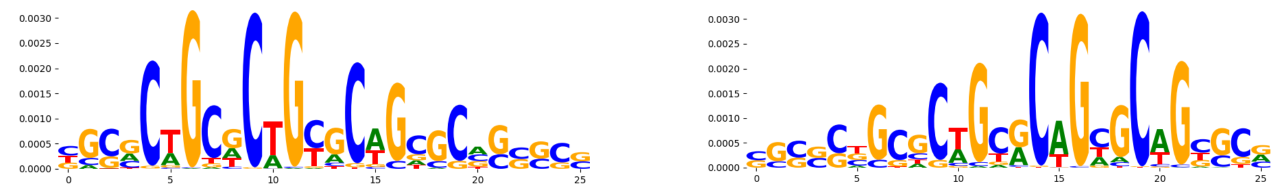 | |
| pos_patterns.pattern_2 | 173 | 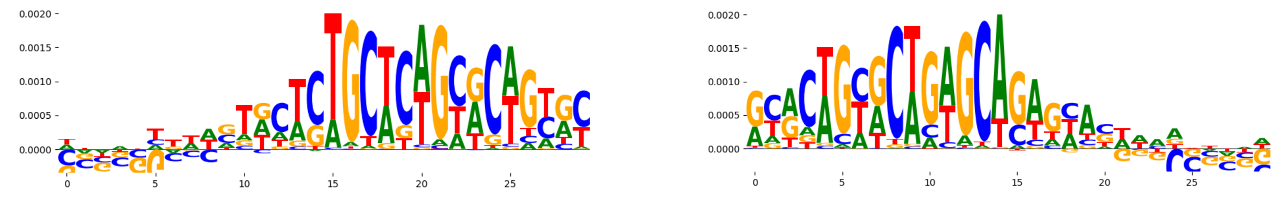 | |
| pos_patterns.pattern_3 | 172 | 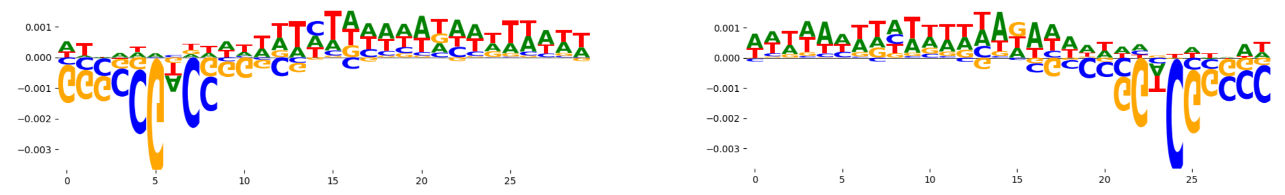 | |
| pos_patterns.pattern_4 | 133 | 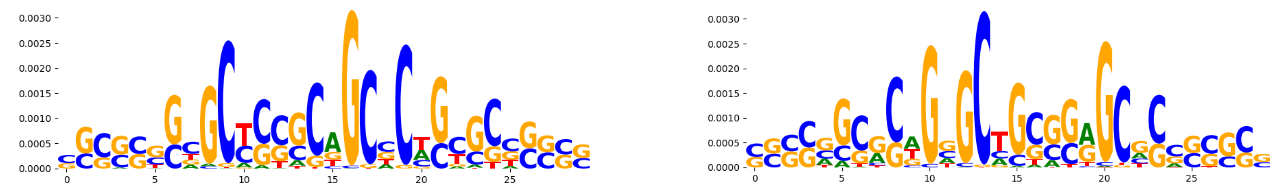 | |
| pos_patterns.pattern_5 | 129 | 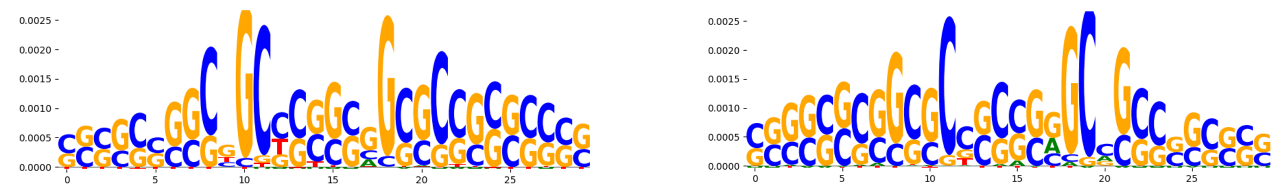 | |
| pos_patterns.pattern_6 | 120 | 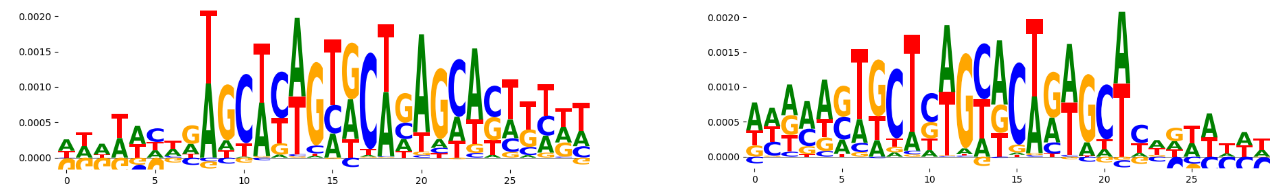 | |
| pos_patterns.pattern_7 | 118 | 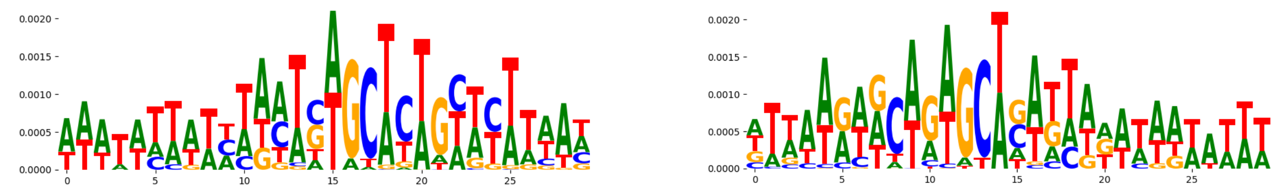 | |
| pos_patterns.pattern_8 | 115 | 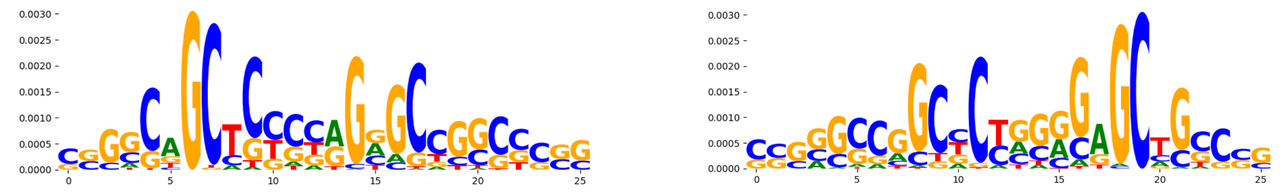 | |
| pos_patterns.pattern_9 | 96 | 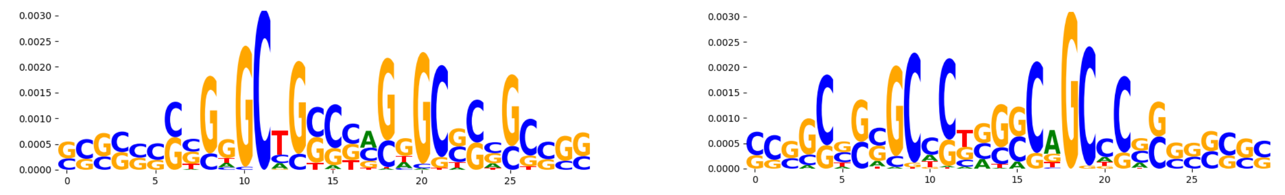 | |
| pos_patterns.pattern_10 | 69 | 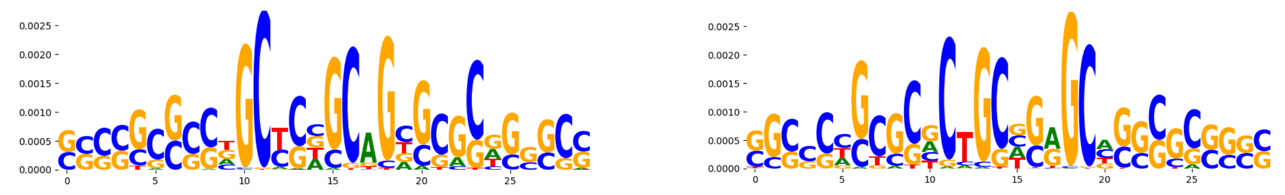 | |
| pos_patterns.pattern_11 | 61 | 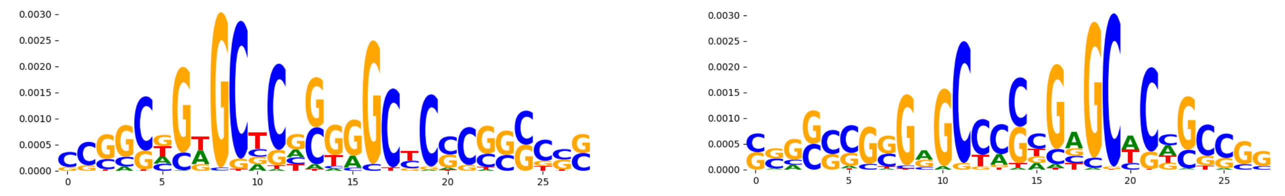 | |
| pos_patterns.pattern_12 | 36 | 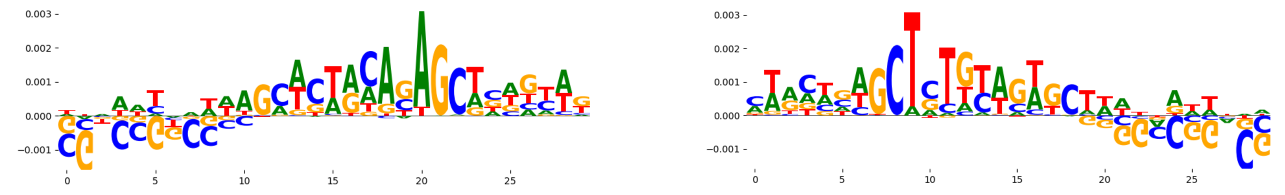 | |
| pos_patterns.pattern_13 | 28 | 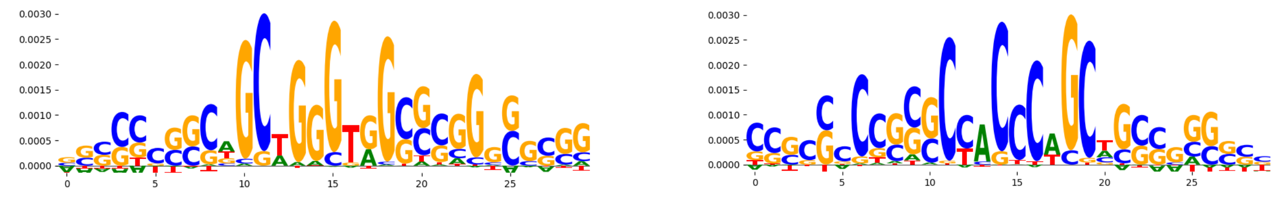 | |
| pos_patterns.pattern_14 | 25 | 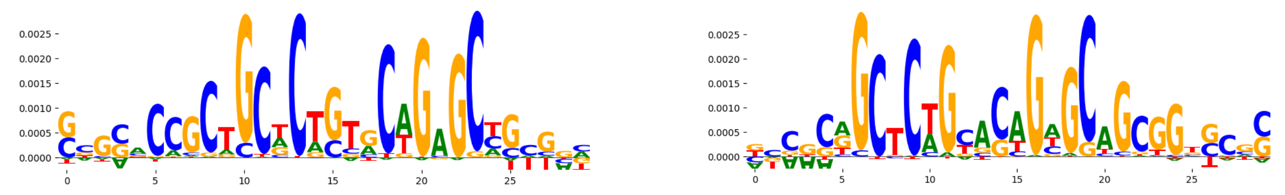 | |
| neg_patterns.pattern_0 | 8321 | 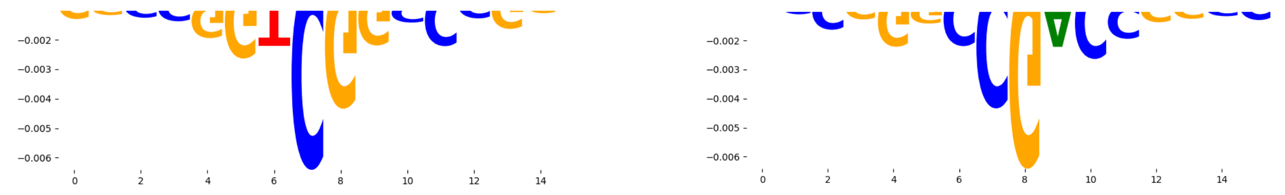 | |
| neg_patterns.pattern_1 | 1780 | 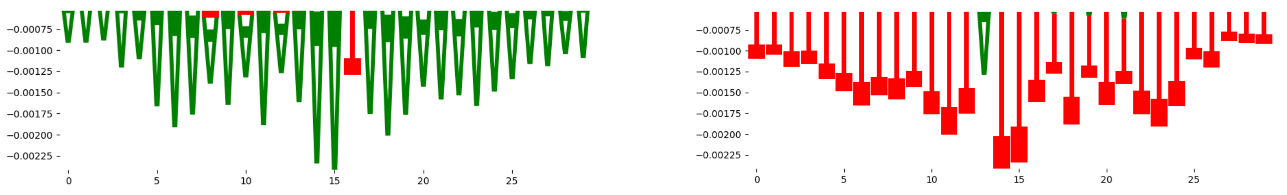 | |
| neg_patterns.pattern_2 | 1513 | 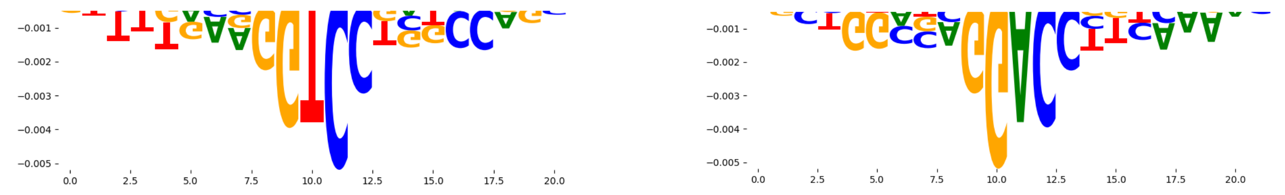 | |
| neg_patterns.pattern_3 | 1481 | 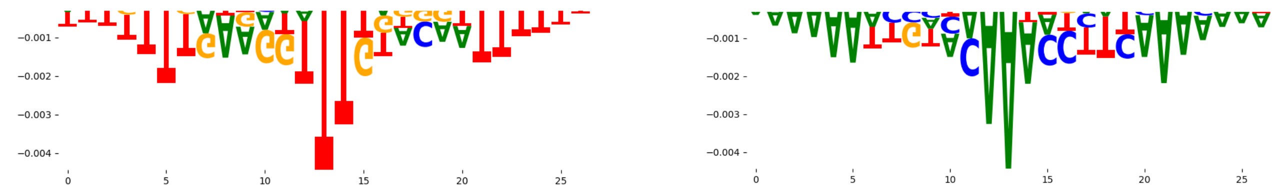 | |
| neg_patterns.pattern_4 | 1029 | 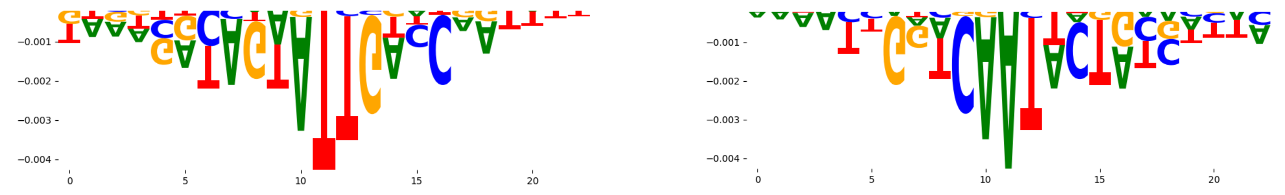 | |
| neg_patterns.pattern_5 | 962 | 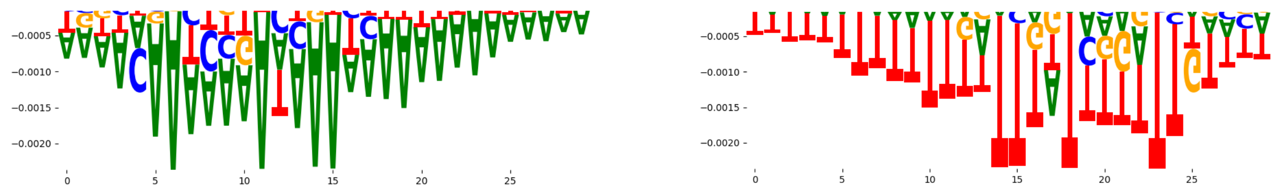 | |
| neg_patterns.pattern_6 | 863 | 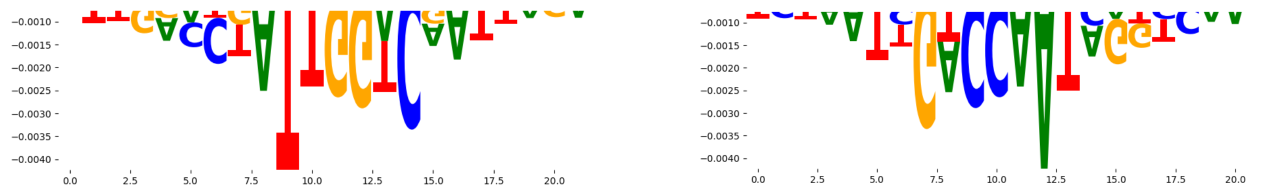 | |
| neg_patterns.pattern_7 | 786 | 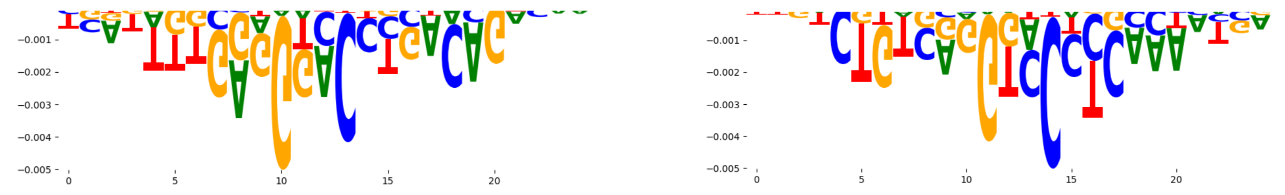 | |

| pattern | num_seqlets | cwm_fwd | cwm_rev |
|---|---|---|---|
| neg_patterns.pattern_8 | 765 | 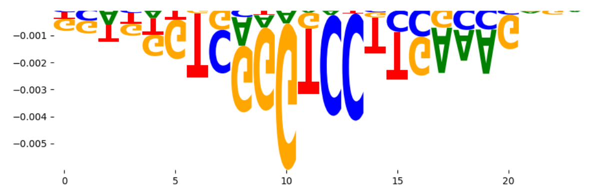 | 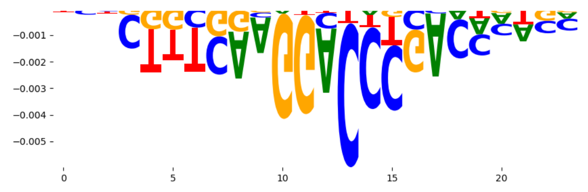 |
| neg_patterns.pattern_9 | 757 | 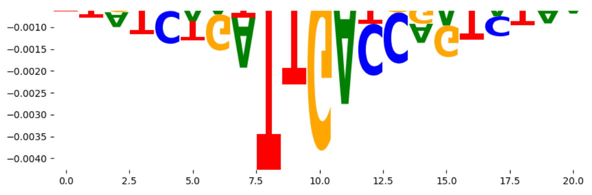 | 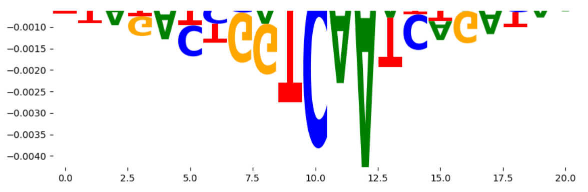 |
| neg_patterns.pattern_10 | 674 | 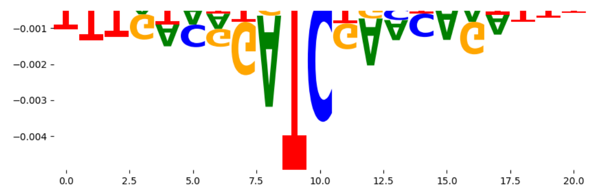 | 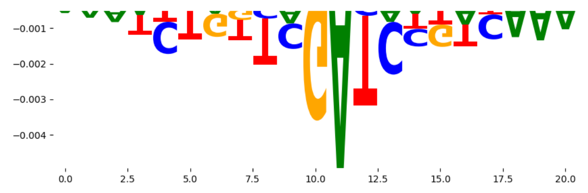 |
| neg_patterns.pattern_11 | 665 | 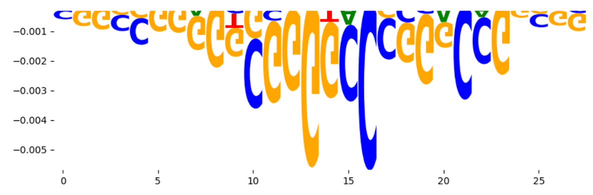 | 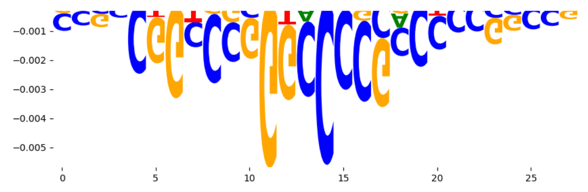 |
| neg_patterns.pattern_12 | 644 | 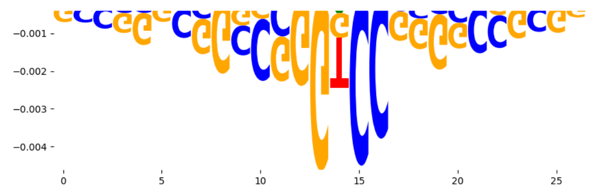 | 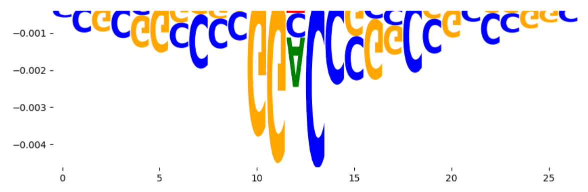 |
| neg_patterns.pattern_13 | 489 | 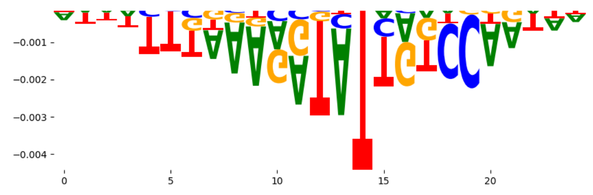 | 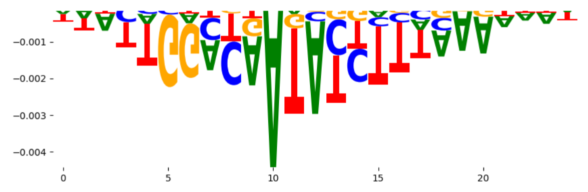 |
| neg_patterns.pattern_14 | 425 | 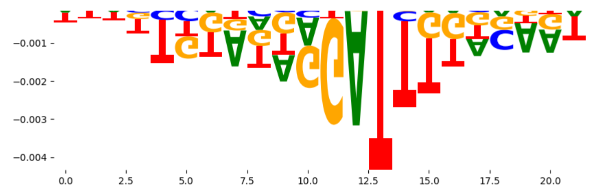 | 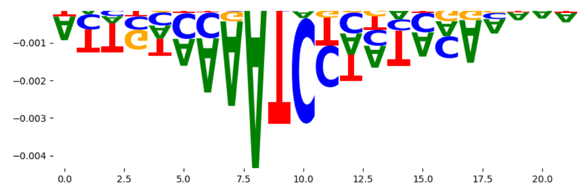 |
| neg_patterns.pattern_15 | 254 | 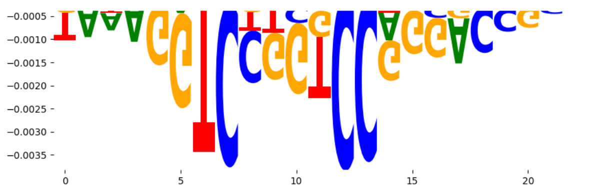 | 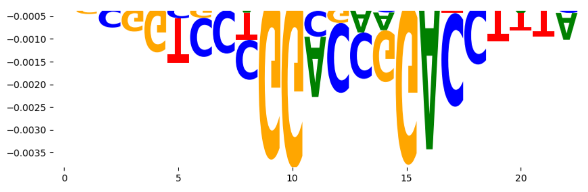 |
| neg_patterns.pattern_16 | 251 | 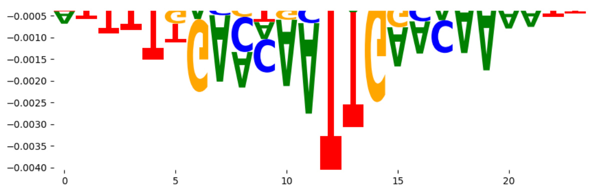 | 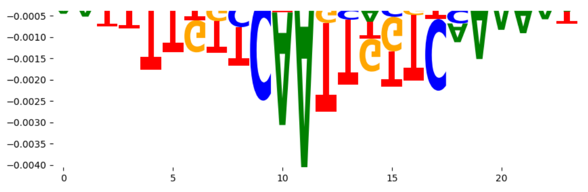 |
| neg_patterns.pattern_17 | 227 | 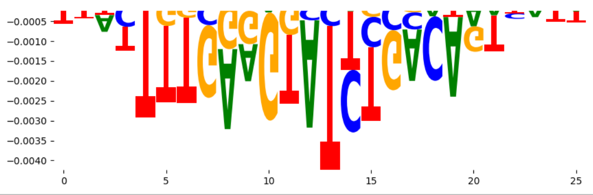 | 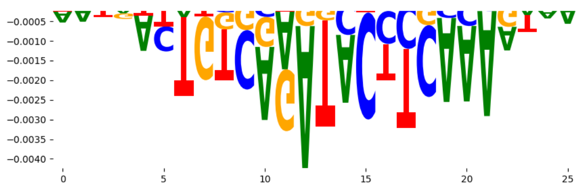 |
