## Supplementary Files 3 for "ChromBPNet: bias factorized, base-resolution deep learning models of chromatin accessibility reveal cis-regulatory sequence syntax, transcription factor footprints and regulatory variants": hepg2_ATAC_raw_bpnet_bias_fold1_profile_modisco.pdf

| pattern | num_seqlets | cwm_fwd | cwm_rev |
|---|---|---|---|
| pos_patterns.pattern_0 | 8242 | 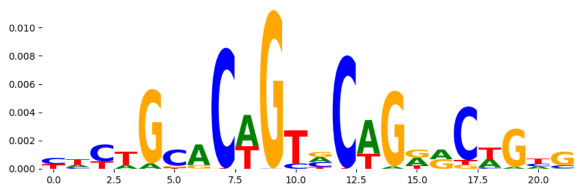 | 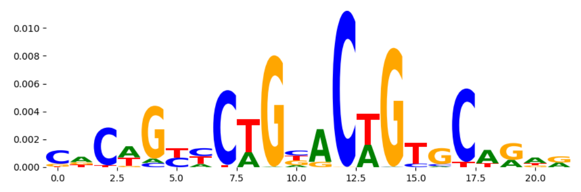 |
| pos_patterns.pattern_1 | 5518 | 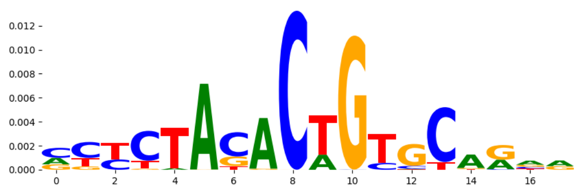 | 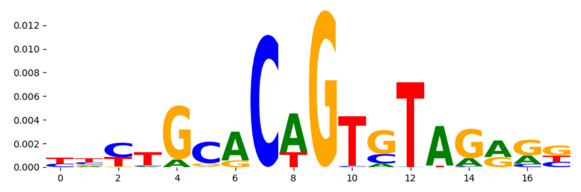 |
| pos_patterns.pattern_2 | 3906 | 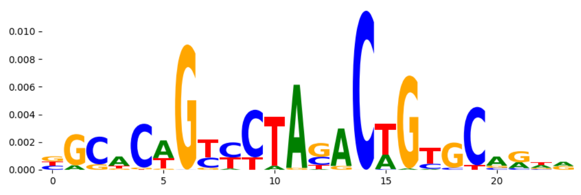 | 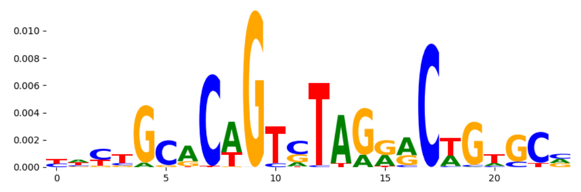 |
| pos_patterns.pattern_3 | 3728 | 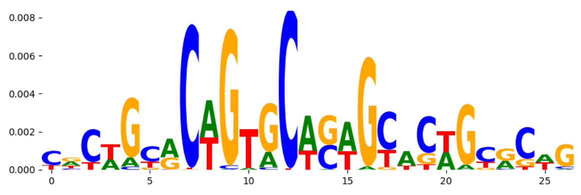 | 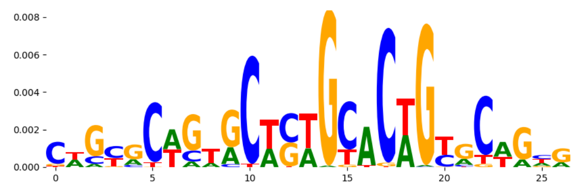 |
| pos_patterns.pattern_4 | 2671 | 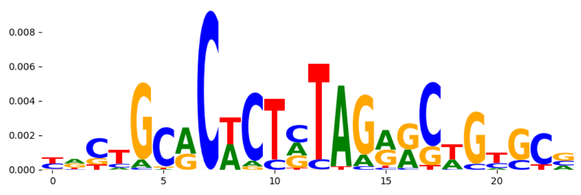 | 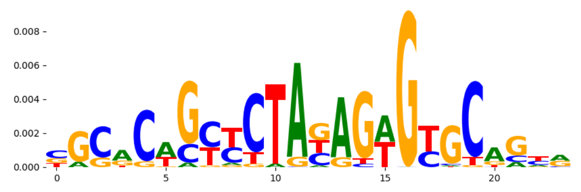 |
| pos_patterns.pattern_5 | 1975 | 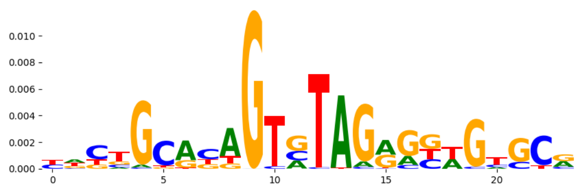 | 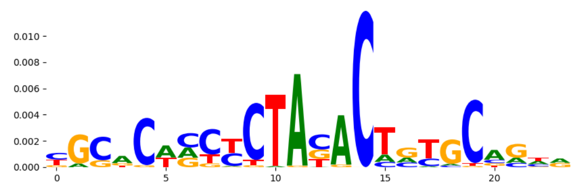 |
| pos_patterns.pattern_6 | 1745 | 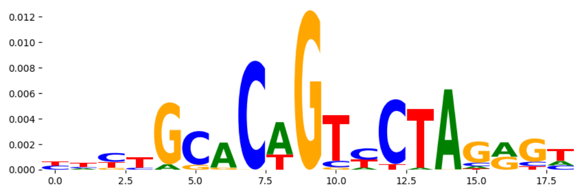 | 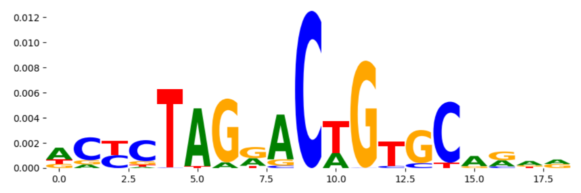 |
| pos_patterns.pattern_7 | 718 | 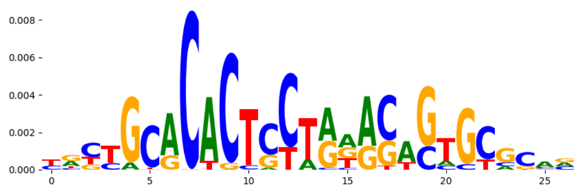 | 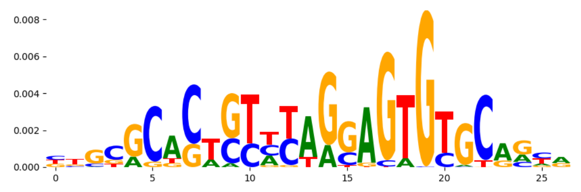 |
| pos_patterns.pattern_8 | 352 | 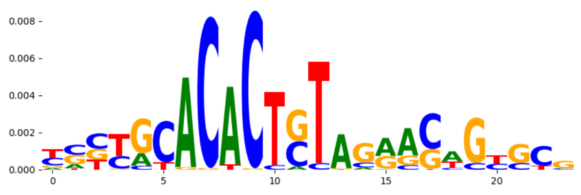 | 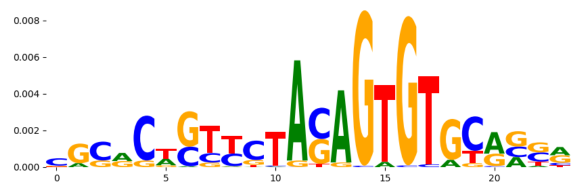 |
| pos_patterns.pattern_9 | 335 | 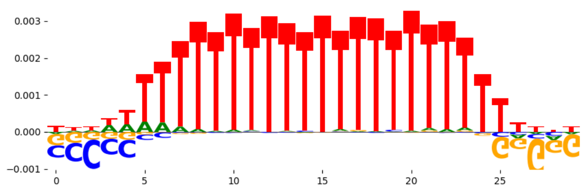 | 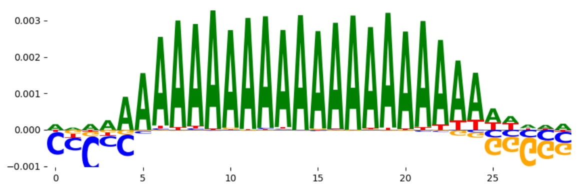 |
| pos_patterns.pattern_10 | 85 | 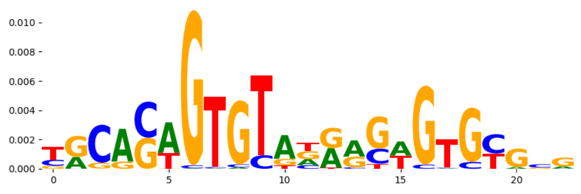 | 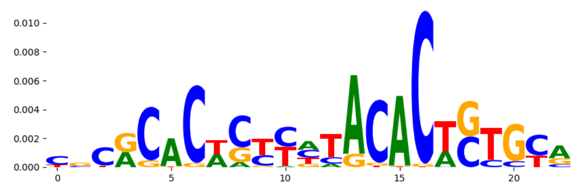 |
| pos_patterns.pattern_11 | 67 | 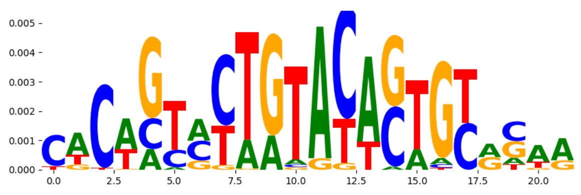 | 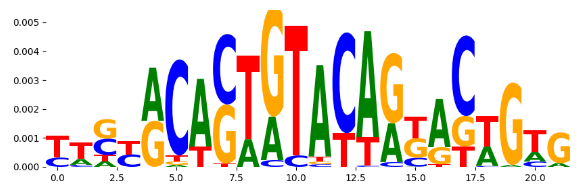 |
| pos_patterns.pattern_12 | 48 | 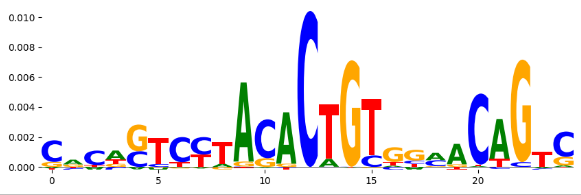 | 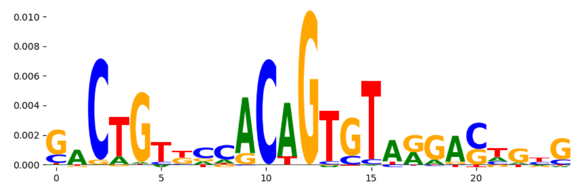 |
