## Supplementary Files 3 for "ChromBPNet: bias factorized, base-resolution deep learning models of chromatin accessibility reveal cis-regulatory sequence syntax, transcription factor footprints and regulatory variants": hepg2_ATAC_raw_bpnet_bias_fold2_counts_modisco.pdf

| pattern | num_seqlets | cwm_fwd | cwm_rev | TOMTOM_match | TOMTOM_qval | TOMTOM_match_logo |
|---|---|---|---|---|---|---|
| pos_patterns.pattern_0 | 6183 | | | SP2_HUMAN.H11MO.0.A | 4.561370e-02 | |
| pos_patterns.pattern_1 | 3409 | | | SP2_HUMAN.H11MO.0.A | 3.201500e-07 | |
| pos_patterns.pattern_2 | 1329 | | | ZNF384_MA1125.1 | 4.313160e-02 | |
| pos_patterns.pattern_3 | 861 | | | RUNX2_HUMAN.H11MO.0.A | 1.000000e+00 | |
| pos_patterns.pattern_4 | 688 | | | ZN667_HUMAN.H11MO.0.C | 1.000000e+00 | |
| pos_patterns.pattern_5 | 685 | | | SP2_MA0516.1 | 7.453490e-02 | |
| pos_patterns.pattern_6 | 562 | | | LEF1_HMG_1 | 1.233180e-01 | |
| pos_patterns.pattern_7 | 549 | | | COE1_MOUSE.H11MO.0.A | 3.453010e-02 | |
| pos_patterns.pattern_8 | 540 | | | Rara.mouse_nuclearreceptor_3 | 5.771810e-01 | |
| pos_patterns.pattern_9 | 539 | | | SOX18_HMG_3 | 9.060570e-02 | |
| pos_patterns.pattern_10 | 537 | | | RARG_nuclearreceptor_5 | 9.315850e-01 | |
| pos_patterns.pattern_11 | 503 | | | ZN320_HUMAN.H11MO.0.C | 5.317270e-01 | |
| pos_patterns.pattern_12 | 485 | | | E2F6_HUMAN.H11MO.0.A | 1.000000e+00 | |
| pos_patterns.pattern_13 | 342 | | | CDX2_MOUSE.H11MO.0.A | 5.878200e-01 | |
| pos_patterns.pattern_14 | 335 | | | ONECUT3_CUT_1 | 2.049560e-01 | |
| pos_patterns.pattern_15 | 319 | | | ZN274_HUMAN.H11MO.0.A | 1.000000e+00 | |
| pos_patterns.pattern_16 | 198 | | | ESRRA_nuclearreceptor_1 | 1.999700e-01 | |
| pos_patterns.pattern_17 | 54 | | | CPEB1_RRM_1 | 1.000000e+00 | |
| pos_patterns.pattern_18 | 20 | | | RORA_HUMAN.H11MO.0.C | 3.656980e-01 | |
| neg_patterns.pattern_0 | 587 | | | TN5_4 | 7.512680e-03 | |
| neg_patterns.pattern_1 | 204 | | | SP1_HUMAN.H11MO.0.A | 2.496090e-04 | |
| neg_patterns.pattern_2 | 171 | | | USF2_HUMAN.H11MO.0.A | 3.041830e-02 | |
| neg_patterns.pattern_3 | 153 | | | ZN331_HUMAN.H11MO.0.C | 2.041380e-03 | |
| neg_patterns.pattern_4 | 134 | | | SP2_HUMAN.H11MO.0.A | 4.150630e-03 | |
| neg_patterns.pattern_5 | 86 | | | ZFX_MOUSE.H11MO.0.B | 1.071070e-02 | |
| neg_patterns.pattern_6 | 60 | | | E2F3_MOUSE.H11MO.0.A | 1.026310e-02 | |
| neg_patterns.pattern_7 | 36 | | | TN5_2 | 7.760040e-01 | |
