## Supplementary Files 3 for "ChromBPNet: bias factorized, base-resolution deep learning models of chromatin accessibility reveal cis-regulatory sequence syntax, transcription factor footprints and regulatory variants": hepg2_ATAC_raw_bpnet_bias_fold2_profile_modisco.pdf

| pattern | num_seqlets | cwm_fwd | cwm_rev | TOMTOM_match | TOMTOM_qval | TOMTOM_match_logo |
|---|---|---|---|---|---|---|
| pos_patterns.pattern_0 | 10489 | | | TN5_2 | 1.259030e-03 | |
| pos_patterns.pattern_1 | 8038 | | | TN5_1 | 1.409980e-09 | |
| pos_patterns.pattern_2 | 4083 | | | TN5_2 | 6.249720e-12 | |
| pos_patterns.pattern_3 | 1486 | | | TN5_1 | 1.672570e-03 | |
| pos_patterns.pattern_4 | 1076 | | | TN5_3 | 3.619950e-06 | |
| pos_patterns.pattern_5 | 976 | | | TN5_1 | 3.774520e-02 | |
| pos_patterns.pattern_6 | 822 | | | TN5_3 | 2.721520e-12 | |
| pos_patterns.pattern_7 | 676 | | | TN5_7 | 2.487450e-06 | |
| pos_patterns.pattern_8 | 481 | | | TN5_3 | 6.509280e-06 | |
| pos_patterns.pattern_9 | 408 | | | TN5_3 | 9.950710e-07 | |
| pos_patterns.pattern_10 | 375 | | | TN5_6 | 5.347200e-17 | |
| pos_patterns.pattern_11 | 373 | | | TN5_1 | 2.307860e-03 | |
| pos_patterns.pattern_12 | 264 | | | PRDM6_HUMAN.H11MO.0.C | 8.077350e-02 | |
| pos_patterns.pattern_13 | 62 | | | MEF2A_MOUSE.H11MO.0.A | 9.872390e-02 | |
| pos_patterns.pattern_14 | 24 | | | TN5_3 | 4.916390e-03 | |
| pos_patterns.pattern_15 | 24 | | | ZNF384_MA1125.1 | 6.650060e-03 | |
| pos_patterns.pattern_16 | 22 | | | ZNF384_MA1125.1 | 3.137470e-02 | |
| pos_patterns.pattern_17 | 20 | | | TN5_2 | 1.072050e-02 | |
