## Supplementary Files 3 for "ChromBPNet: bias factorized, base-resolution deep learning models of chromatin accessibility reveal cis-regulatory sequence syntax, transcription factor footprints and regulatory variants": hepg2_ATAC_raw_bpnet_bias_fold3_counts_modisco.pdf

| pattern | num_seqlets | cwm_fwd | cwm_rev | TOMTOM_match | TOMTOM_qval | TOMTOM_match_logo |
|---|---|---|---|---|---|---|
| pos_patterns.pattern_0 | 4279 | | | SP1_HUMAN.H11MO.0.A | 5.256980e-01 | |
| pos_patterns.pattern_1 | 2891 | | | ZFX_MOUSE.H11MO.0.B | 4.984460e-03 | |
| pos_patterns.pattern_2 | 1631 | | | ZNF384_MA1125.1 | 9.365280e-02 | |
| pos_patterns.pattern_3 | 1628 | | | SP1_HUMAN.H11MO.0.A | 8.003940e-03 | |
| pos_patterns.pattern_4 | 892 | | | Arid5a_MA0602.1 | 1.000000e+00 | |
| pos_patterns.pattern_5 | 855 | | | HNF4A_nuclearreceptor_4 | 4.039120e-01 | |
| pos_patterns.pattern_6 | 774 | | | HXA13_HUMAN.H11MO.0.C | 9.324020e-01 | |
| pos_patterns.pattern_7 | 748 | | | SP2_HUMAN.H11MO.0.A | 3.694840e-09 | |
| pos_patterns.pattern_8 | 730 | | | ZIC3_HUMAN.H11MO.0.B | 2.580640e-01 | |
| pos_patterns.pattern_9 | 730 | | | PLAG1_MA0163.1 | 6.112310e-01 | |
| pos_patterns.pattern_10 | 706 | | | SP2_HUMAN.H11MO.0.A | 2.042260e-03 | |
| pos_patterns.pattern_11 | 703 | | | BHA15_HUMAN.H11MO.0.B | 3.479100e-01 | |
| pos_patterns.pattern_12 | 627 | | | MAZ_HUMAN.H11MO.0.A | 1.240200e-02 | |
| pos_patterns.pattern_13 | 354 | | | ZN281_MOUSE.H11MO.0.A | 6.040920e-05 | |
| pos_patterns.pattern_14 | 337 | | | ZN281_MOUSE.H11MO.0.A | 7.705600e-04 | |
| pos_patterns.pattern_15 | 298 | | | SP1_HUMAN.H11MO.0.A | 5.859200e-03 | |
| pos_patterns.pattern_16 | 294 | | | PRDM5_MOUSE.H11MO.0.A | 1.000000e+00 | |
| pos_patterns.pattern_17 | 155 | | | SP1_HUMAN.H11MO.0.A | 2.231530e-03 | |
| pos_patterns.pattern_18 | 126 | | | PRDM5_MOUSE.H11MO.0.A | 7.308600e-02 | |
| pos_patterns.pattern_19 | 115 | | | ZNF384_MA1125.1 | 1.502460e-01 | |
| pos_patterns.pattern_20 | 68 | | | BARX1_MOUSE.H11MO.0.C | 1.000000e+00 | |
| neg_patterns.pattern_0 | 252 | | | TN5_2 | 2.599430e-01 | |
| neg_patterns.pattern_1 | 217 | | | TN5_7 | 6.788550e-01 | |
| neg_patterns.pattern_2 | 211 | | | TN5_2 | 3.704520e-01 | |
| neg_patterns.pattern_3 | 190 | | | TN5_2 | 1.006620e-01 | |
| neg_patterns.pattern_4 | 180 | | | MAX_bHLH_1 | 6.772120e-03 | |
| neg_patterns.pattern_5 | 150 | | | SP2_HUMAN.H11MO.0.A | 9.060400e-03 | |
| neg_patterns.pattern_6 | 147 | | | SP2_HUMAN.H11MO.0.A | 1.204180e-01 | |
| neg_patterns.pattern_7 | 102 | | | KLF5_MOUSE.H11MO.0.A | 8.517020e-02 | |

| pattern | num_seqlets | cwm_fwd | cwm_rev | TOMTOM_match | TOMTOM_qval | TOMTOM_match_logo |
|---|---|---|---|---|---|---|
| neg_patterns.pattern_8 | 71 | 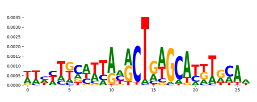 | 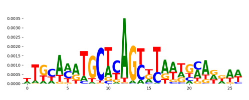 | ZFX_MOUSE.H11MO.0.B | 1.936040e-01 | 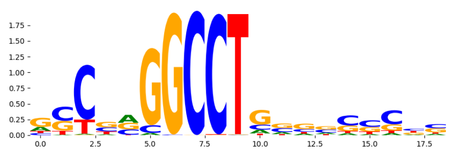 |
| neg_patterns.pattern_9 | 71 | 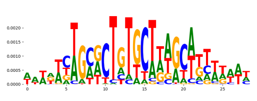 | 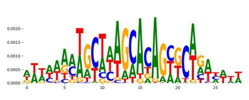 | TN5_2 | 5.279140e-03 | 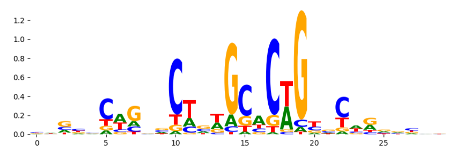 |
| neg_patterns.pattern_10 | 30 | 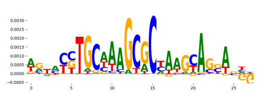 | 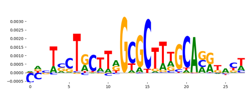 | SMAD4_HUMAN.H11MO.0.B | 3.478060e-01 | 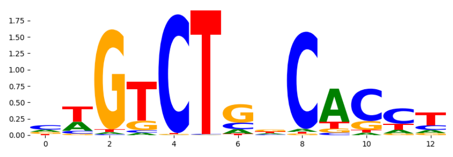 |
| neg_patterns.pattern_11 | 27 | 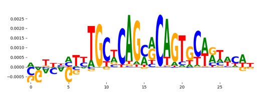 | 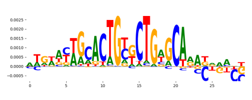 | TN5_1 | 1.748220e-01 | 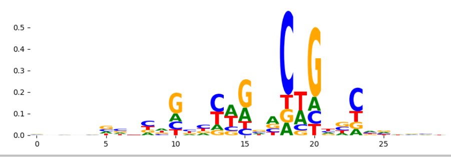 |
