## Supplementary Files 3 for "ChromBPNet: bias factorized, base-resolution deep learning models of chromatin accessibility reveal cis-regulatory sequence syntax, transcription factor footprints and regulatory variants": hepg2_ATAC_raw_bpnet_bias_fold3_profile_modisco.pdf

| pattern | num_seqlets | cwm_fwd | cwm_rev | TOMTOM_match | TOMTOM_qval | TOMTOM_match_logo |
|---|---|---|---|---|---|---|
| pos_patterns.pattern_0 | 12329 | 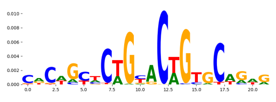 | 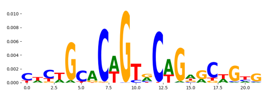 | TN5_2 | 6.058040e-09 | 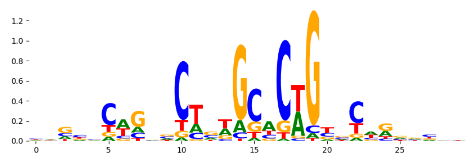 |
| pos_patterns.pattern_1 | 6736 | 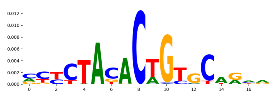 | 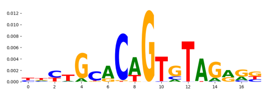 | TN5_4 | 2.121400e-02 | 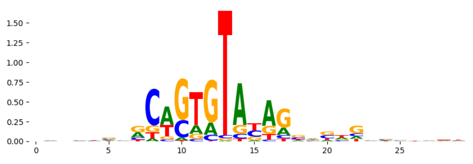 |
| pos_patterns.pattern_2 | 3753 | 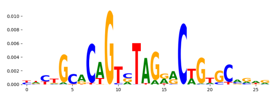 | 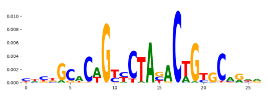 | TN5_1 | 2.584820e-08 | 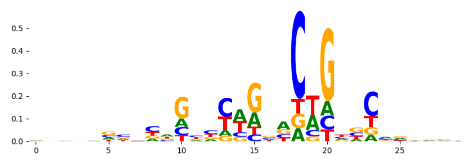 |
| pos_patterns.pattern_3 | 1653 | 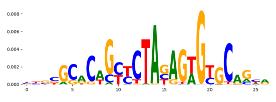 | 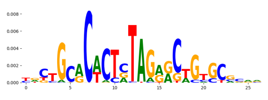 | TN5_3 | 1.433350e-10 | 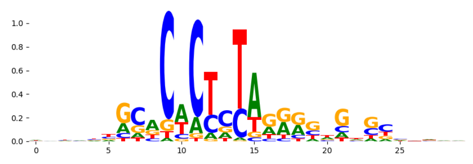 |
| pos_patterns.pattern_4 | 1349 | 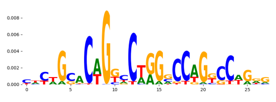 | 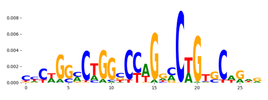 | TN5_1 | 4.617570e-05 | 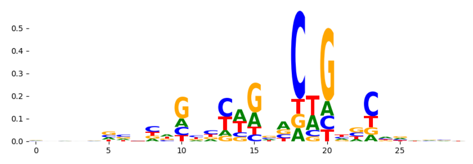 |
| pos_patterns.pattern_5 | 1164 | 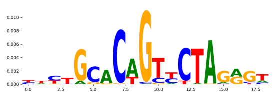 | 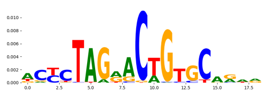 | TN5_7 | 2.616660e-02 | 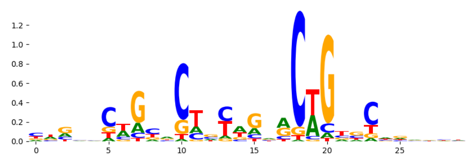 |
| pos_patterns.pattern_6 | 592 | 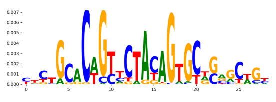 | 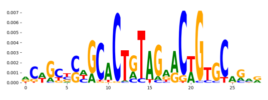 | TN5_7 | 1.742780e-10 | 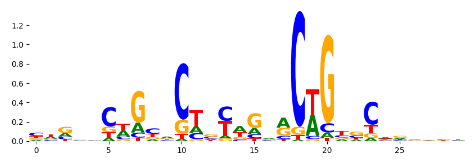 |
| pos_patterns.pattern_7 | 572 | 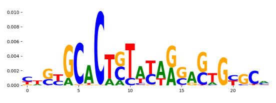 | 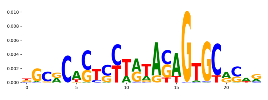 | TN5_4 | 4.545890e-04 | 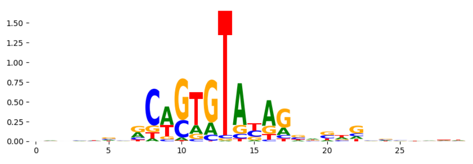 |
| pos_patterns.pattern_8 | 386 | 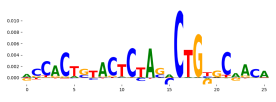 | 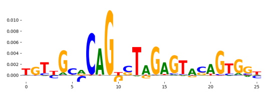 | TN5_6 | 2.613380e-13 | 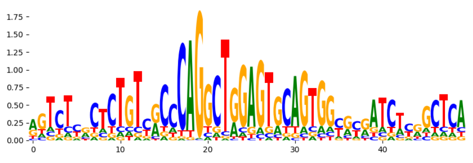 |
| pos_patterns.pattern_9 | 373 | 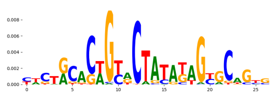 | 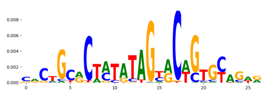 | TN5_3 | 1.141960e-02 | 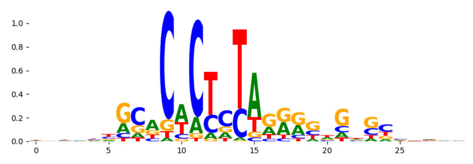 |
| pos_patterns.pattern_10 | 265 | 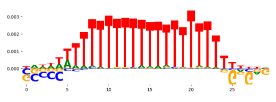 | 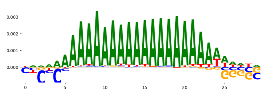 | ZNF384_MA1125.1 | 6.346600e-02 | 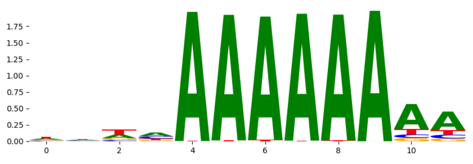 |
| pos_patterns.pattern_11 | 66 | 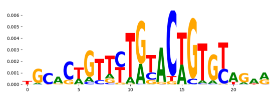 | 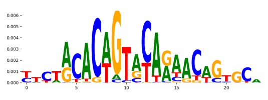 | TN5_2 | 6.194690e-02 | 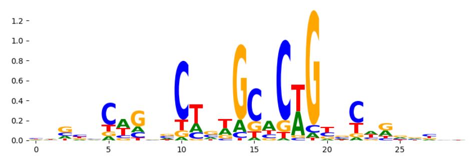 |
| pos_patterns.pattern_12 | 20 | 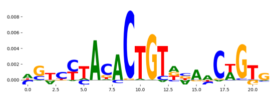 | 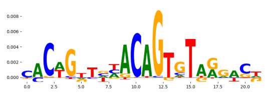 | TN5_7 | 2.104270e-01 | 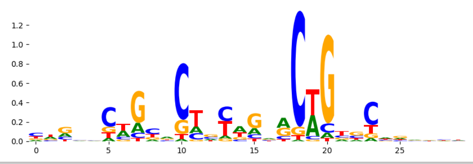 |
