## Supplementary Files 3 for "ChromBPNet: bias factorized, base-resolution deep learning models of chromatin accessibility reveal cis-regulatory sequence syntax, transcription factor footprints and regulatory variants": hepg2_ATAC_raw_bpnet_bias_fold4_counts_modisco.pdf

| pattern | num_seqlets | cwm_fwd | cwm_rev | TOMTOM_match | TOMTOM_qval | TOMTOM_match_logo |
|---|---|---|---|---|---|---|
| pos_patterns.pattern_0 | 2818 | | | ZN320_HUMAN.H11MO.0.C | 4.594080e-01 | |
| pos_patterns.pattern_1 | 1941 | | | ZFX_MOUSE.H11MO.0.B | 9.701540e-02 | |
| pos_patterns.pattern_2 | 1904 | | | ZNF384_MA1125.1 | 1.421240e-01 | |
| pos_patterns.pattern_3 | 1566 | | | SP3_HUMAN.H11MO.0.B | 4.920160e-02 | |
| pos_patterns.pattern_4 | 963 | | | GATA1+TAL1_MA0140.2 | 1.000000e+00 | |
| pos_patterns.pattern_5 | 908 | | | NFATC3_MA0625.1 | 1.632030e-01 | |
| pos_patterns.pattern_6 | 827 | | | DNASE_1 | 1.000000e+00 | |
| pos_patterns.pattern_7 | 785 | | | SNAI1_HUMAN.H11MO.0.C | 2.386940e-02 | |
| pos_patterns.pattern_8 | 706 | | | PLAG1_MA0163.1 | 1.562540e-02 | |
| pos_patterns.pattern_9 | 685 | | | NR2F1_nuclearreceptor_3 | 8.579050e-02 | |
| pos_patterns.pattern_10 | 626 | | | SP1_HUMAN.H11MO.0.A | 4.324510e-04 | |
| pos_patterns.pattern_11 | 529 | | | RREB1_MA0073.1 | 7.257090e-02 | |
| pos_patterns.pattern_12 | 522 | | | GATA1+TAL1_MA0140.2 | 1.000000e+00 | |
| pos_patterns.pattern_13 | 492 | | | SP1_MOUSE.H11MO.0.A | 7.476610e-02 | |
| pos_patterns.pattern_14 | 471 | | | Hnf4a.mouse_nuclearreceptor_1 | 9.999860e-01 | |
| pos_patterns.pattern_15 | 439 | | | NFAC1_MOUSE.H11MO.0.A | 1.000000e+00 | |
| pos_patterns.pattern_16 | 339 | | | RREB1_MA0073.1 | 2.545570e-02 | |
| pos_patterns.pattern_17 | 286 | | | NaN | NaN | |
| pos_patterns.pattern_18 | 274 | | | MYBL2_MYB_2 | 9.312390e-01 | |
| pos_patterns.pattern_19 | 243 | | | SP1_HUMAN.H11MO.0.A | 1.366920e-01 | |
| pos_patterns.pattern_20 | 222 | | | RREB1_MA0073.1 | 2.125040e-03 | |
| pos_patterns.pattern_21 | 195 | | | FOXK1_MA0852.2 | 1.000000e+00 | |
| pos_patterns.pattern_22 | 165 | | | PBX3_MA1114.1 | 1.000000e+00 | |
| pos_patterns.pattern_23 | 83 | | | Hoxa9_MA0594.1 | 1.000000e+00 | |
| pos_patterns.pattern_24 | 33 | | | SP1_MOUSE.H11MO.0.A | 5.009200e-02 | |
| neg_patterns.pattern_0 | 406 | | | TN5_2 | 3.693140e-07 | |
| neg_patterns.pattern_1 | 271 | | | TN5_2 | 1.349410e-03 | |
| neg_patterns.pattern_2 | 122 | | | ZIC1_C2H2_1 | 1.572720e-01 | |
| neg_patterns.pattern_3 | 112 | | | SP2_HUMAN.H11MO.0.A | 1.000060e-05 | |
| neg_patterns.pattern_4 | 96 | | | CTCF_MA0139.1 | 2.905270e-07 | |

| pattern | num_seqlets | cwm_fwd | cwm_rev | TOMTOM_match | TOMTOM_qval | TOMTOM_match_logo |
|---|---|---|---|---|---|---|
| neg_patterns.pattern_5 | 95 | 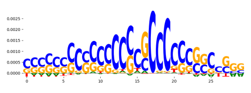 | 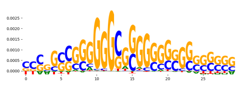 | ZBT17_HUMAN.H11MO.0.A | 4.379260e-03 | 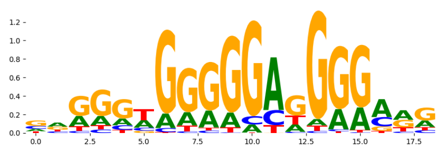 |
| neg_patterns.pattern_6 | 91 | 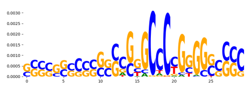 | 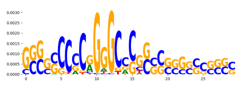 | AP2C_HUMAN.H11MO.0.A | 5.706460e-01 | 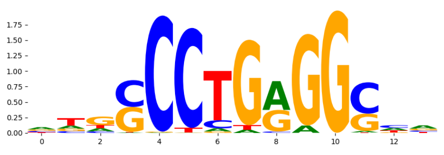 |
| neg_patterns.pattern_7 | 81 | 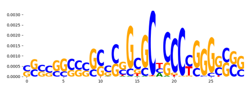 | 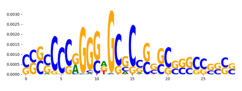 | ZN528_HUMAN.H11MO.0.C | 1.000000e+00 | 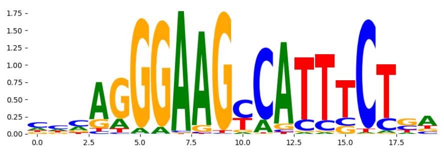 |
| neg_patterns.pattern_8 | 35 | 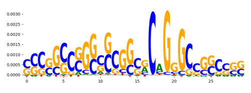 | 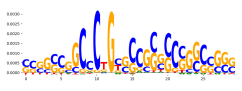 | TN5_1 | 2.591020e-01 | 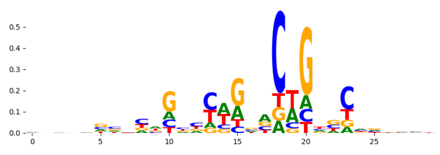 |
| neg_patterns.pattern_9 | 33 | 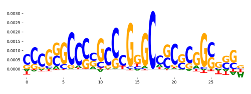 | 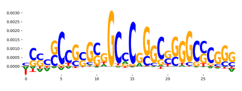 | PAX5_HUMAN.H11MO.0.A | 1.000000e+00 | 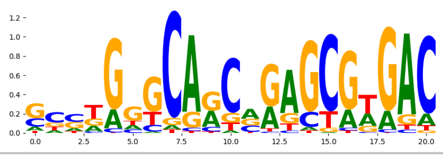 |
