## Supplementary Files 3 for "ChromBPNet: bias factorized, base-resolution deep learning models of chromatin accessibility reveal cis-regulatory sequence syntax, transcription factor footprints and regulatory variants": hepg2_ATAC_raw_bpnet_bias_fold4_profile_modisco.pdf

| pattern | num_seqlets | cwm_fwd | cwm_rev | TOMTOM_match | TOMTOM_qval | TOMTOM_match_logo |
|---|---|---|---|---|---|---|
| pos_patterns.pattern_0 | 7951 | 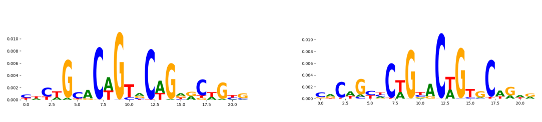 | | TN5_1 | 1.121210e-09 | 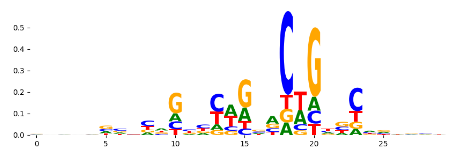 |
| pos_patterns.pattern_1 | 6047 | 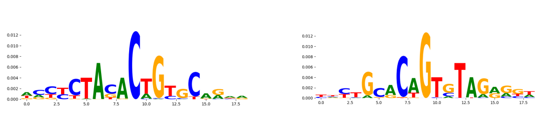 | | TN5_8 | 4.005420e-02 | 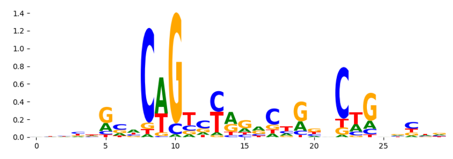 |
| pos_patterns.pattern_2 | 4203 | 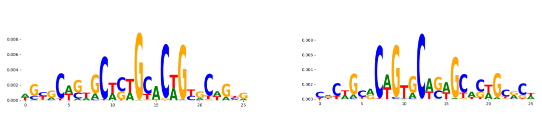 | | TN5_2 | 1.132310e-16 | 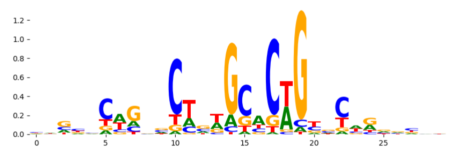 |
| pos_patterns.pattern_3 | 4050 | 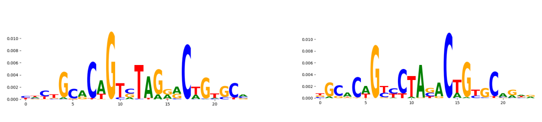 | | TN5_1 | 1.441260e-05 | 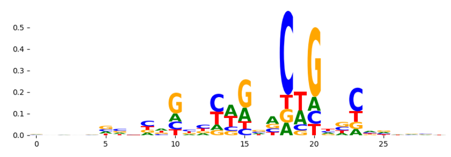 |
| pos_patterns.pattern_4 | 1755 | 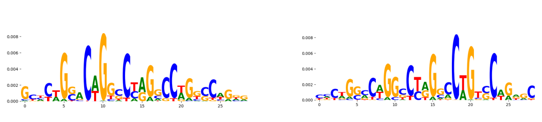 | | TN5_1 | 1.550630e-03 | 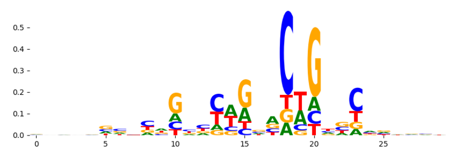 |
| pos_patterns.pattern_5 | 1283 | 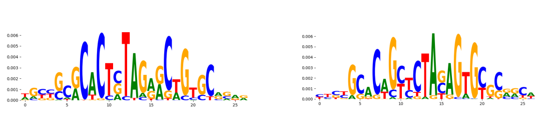 | | TN5_3 | 1.283030e-08 | 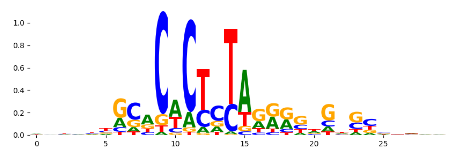 |
| pos_patterns.pattern_6 | 827 | 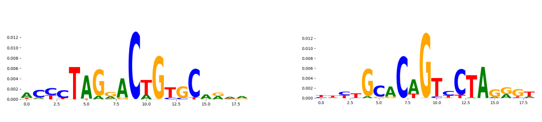 | | TN5_1 | 4.122530e-02 | 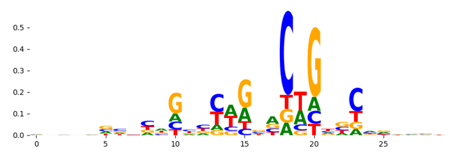 |
| pos_patterns.pattern_7 | 737 | 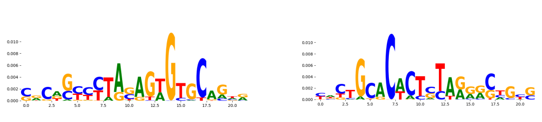 | | TN5_3 | 5.631160e-08 | 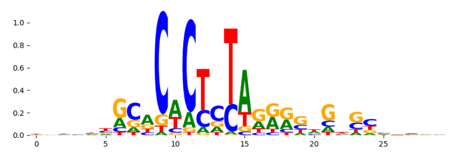 |
| pos_patterns.pattern_8 | 650 | 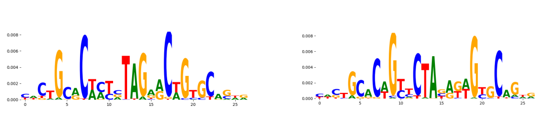 | | TN5_3 | 1.579030e-08 | 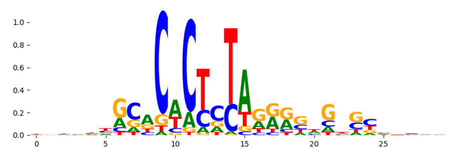 |
| pos_patterns.pattern_9 | 582 | 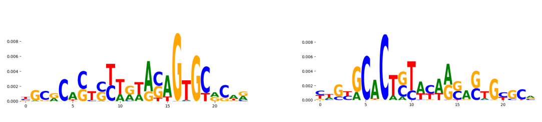 | | TN5_4 | 1.092690e-03 | 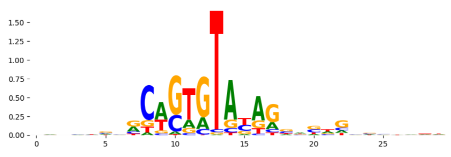 |
| pos_patterns.pattern_10 | 443 | 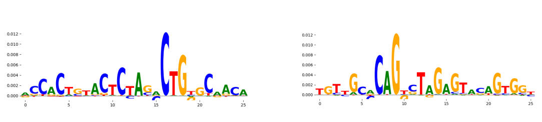 | | TN5_6 | 2.224700e-15 | 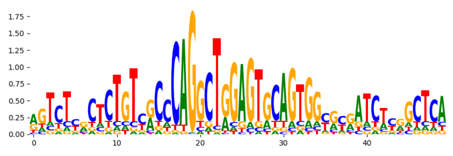 |
| pos_patterns.pattern_11 | 407 | 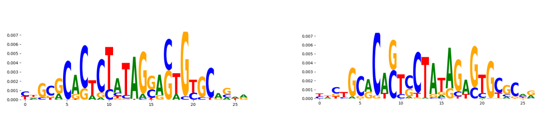 | | TN5_3 | 2.228240e-06 | 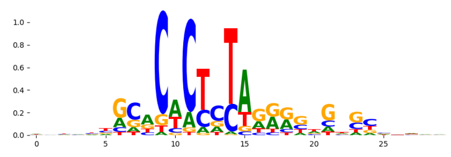 |
| pos_patterns.pattern_12 | 249 | 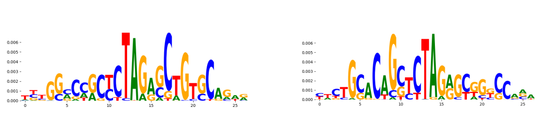 | | TN5_3 | 9.178750e-02 | 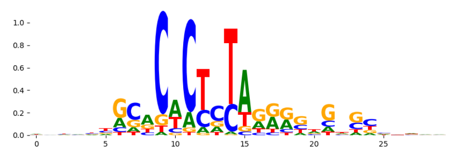 |
| pos_patterns.pattern_13 | 248 | 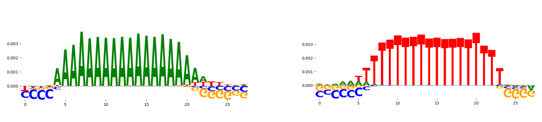 | | PRDM6_HUMAN.H11MO.0.C | 5.646740e-02 | 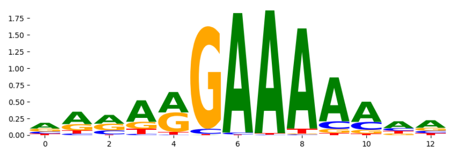 |
| pos_patterns.pattern_14 | 97 | 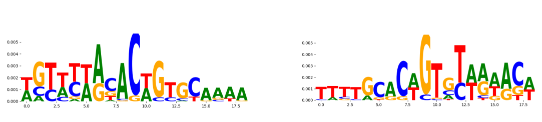 | | TN5_1 | 1.857470e-03 | 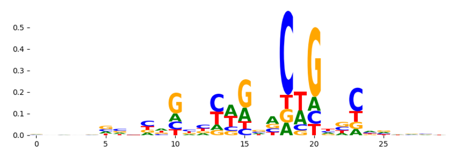 |
| pos_patterns.pattern_15 | 68 | 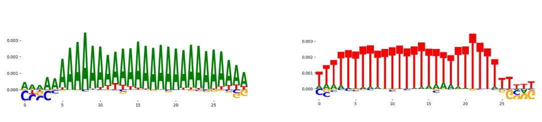 | | ZNF384_MA1125.1 | 7.701890e-02 | 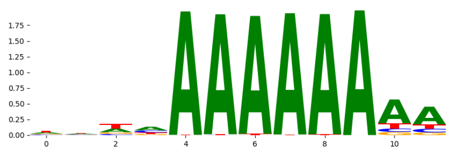 |
| pos_patterns.pattern_16 | 50 | 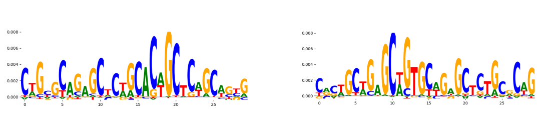 | | TN5_2 | 4.346460e-02 | 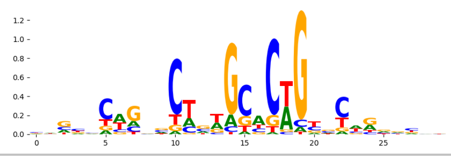 |
