## Supplementary Files 3 for "ChromBPNet: bias factorized, base-resolution deep learning models of chromatin accessibility reveal cis-regulatory sequence syntax, transcription factor footprints and regulatory variants": hepg2_DNASE_raw_bpnet_bias_fold0_profile_modisco.pdf

| pattern | num_seqlets | cwm_fwd | cwm_rev | TOMTOM_match | TOMTOM_qval | TOMTOM_match_logo |
|---|---|---|---|---|---|---|
| pos_patterns.pattern_0 | 8153 | | | DNASE_1 | 7.557850e-06 | |
| pos_patterns.pattern_1 | 6297 | | | DNASE_2 | 2.269380e-06 | |
| pos_patterns.pattern_2 | 1408 | | | DNASE_3 | 1.066990e-06 | |
| pos_patterns.pattern_3 | 973 | | | PRDM6_HUMAN.H11MO.0.C | 7.770000e-02 | |
| pos_patterns.pattern_4 | 751 | | | KLF6_HUMAN.H11MO.0.A | 4.581690e-02 | |
| pos_patterns.pattern_5 | 663 | | | MEF2B_HUMAN.H11MO.0.A | 2.418480e-01 | |
| pos_patterns.pattern_6 | 615 | | | ZNF384_MA1125.1 | 4.853220e-03 | |
| pos_patterns.pattern_7 | 530 | | | DNASE_4 | 9.556290e-03 | |
| pos_patterns.pattern_8 | 447 | | | SP1_MOUSE.H11MO.0.A | 3.542410e-09 | |
| pos_patterns.pattern_9 | 427 | | | ZNF384_MA1125.1 | 1.599120e-02 | |
| pos_patterns.pattern_10 | 328 | | | SP2_HUMAN.H11MO.0.A | 5.814950e-06 | |
| pos_patterns.pattern_11 | 307 | | | DNASE_5 | 1.986510e-15 | |
| pos_patterns.pattern_12 | 304 | | | DNASE_5 | 2.254080e-05 | |
| pos_patterns.pattern_13 | 302 | | | FOXK1_forkhead_1 | 9.335290e-01 | |
| pos_patterns.pattern_14 | 230 | | | ZN770_HUMAN.H11MO.0.C | 5.335500e-01 | |
| pos_patterns.pattern_15 | 198 | | | FOXC1_forkhead_1 | 1.091330e-01 | |
| pos_patterns.pattern_16 | 175 | | | TN5_6 | 2.307430e-20 | |
| pos_patterns.pattern_17 | 117 | | | DNASE_6 | 7.359140e-20 | |
| pos_patterns.pattern_18 | 106 | | | PRDM6_HUMAN.H11MO.0.C | 3.849030e-02 | |
| pos_patterns.pattern_19 | 94 | | | ZN770_HUMAN.H11MO.0.C | 6.576460e-01 | |
| pos_patterns.pattern_20 | 85 | | | KLF1_HUMAN.H11MO.0.A | 9.492460e-01 | |
| pos_patterns.pattern_21 | 58 | | | ZN770_HUMAN.H11MO.0.C | 3.368640e-08 | |
| pos_patterns.pattern_22 | 41 | | | IRF3_HUMAN.H11MO.0.B | 1.958740e-02 | |
| pos_patterns.pattern_23 | 32 | | | RORC_MA1151.1 | 1.000000e+00 | |
| pos_patterns.pattern_24 | 20 | | | ITF2_HUMAN.H11MO.0.C | 1.000000e+00 | |
| pos_patterns.pattern_25 | 20 | | | ZN431_MOUSE.H11MO.0.C | 1.000000e+00 | |
| neg_patterns.pattern_0 | 46 | | | ZBT17_MOUSE.H11MO.0.A | 1.353000e-03 | |
| neg_patterns.pattern_1 | 29 | | | ZNF384_MA1125.1 | 5.826750e-01 | |
| neg_patterns.pattern_2 | 21 | | | GATA3_HUMAN.H11MO.0.A | 1.457100e-01 | |
