## Supplementary Files 3 for "ChromBPNet: bias factorized, base-resolution deep learning models of chromatin accessibility reveal cis-regulatory sequence syntax, transcription factor footprints and regulatory variants": hepg2_DNASE_raw_bpnet_bias_fold1_counts_modisco.pdf

| pattern | num_seqlets | cwm_fwd | cwm_rev | TOMTOM_match | TOMTOM_qval | TOMTOM_match_logo |
|---|---|---|---|---|---|---|
| pos_patterns.pattern_0 | 9078 | | | DNASE_1 | 1.175780e-03 | |
| pos_patterns.pattern_1 | 6751 | | | DNASE_2 | 6.828250e-06 | |
| pos_patterns.pattern_2 | 2755 | | | DNASE_3 | 6.802010e-06 | |
| pos_patterns.pattern_3 | 937 | | | PRDM6_HUMAN.H11MO.0.C | 7.674420e-02 | |
| pos_patterns.pattern_4 | 691 | | | SP1_HUMAN.H11MO.0.A | 9.222830e-05 | |
| pos_patterns.pattern_5 | 480 | | | MEF2B_HUMAN.H11MO.0.A | 1.850760e-01 | |
| pos_patterns.pattern_6 | 480 | | | SP1_MOUSE.H11MO.0.A | 4.801290e-08 | |
| pos_patterns.pattern_7 | 479 | | | ZNF384_MA1125.1 | 6.805550e-05 | |
| pos_patterns.pattern_8 | 451 | | | ZNF384_MA1125.1 | 2.566890e-03 | |
| pos_patterns.pattern_9 | 435 | | | ZNF384_MA1125.1 | 3.180030e-02 | |
| pos_patterns.pattern_10 | 255 | | | SP1_HUMAN.H11MO.0.A | 6.240570e-01 | |
| pos_patterns.pattern_11 | 242 | | | SP2_HUMAN.H11MO.0.A | 1.087770e-04 | |
| pos_patterns.pattern_12 | 211 | | | TN5_6 | 7.072850e-18 | |
| pos_patterns.pattern_13 | 207 | | | DNASE_2 | 4.810540e-01 | |
| pos_patterns.pattern_14 | 200 | | | ZN770_HUMAN.H11MO.0.C | 2.351320e-08 | |
| pos_patterns.pattern_15 | 180 | | | ZN770_HUMAN.H11MO.0.C | 1.758660e-08 | |
| pos_patterns.pattern_16 | 122 | | | ZN449_HUMAN.H11MO.0.C | 6.649680e-01 | |
| pos_patterns.pattern_17 | 79 | | | ITF2_HUMAN.H11MO.0.C | 8.150400e-01 | |
| pos_patterns.pattern_18 | 60 | | | KLF4_MA0039.3 | 7.845210e-01 | |
| pos_patterns.pattern_19 | 54 | | | DNASE_2 | 1.000000e+00 | |
| pos_patterns.pattern_20 | 41 | | | PRDM6_HUMAN.H11MO.0.C | 9.679110e-02 | |
| pos_patterns.pattern_21 | 35 | | | ZNF384_MA1125.1 | 2.287250e-01 | |
| pos_patterns.pattern_22 | 30 | | | DNASE_3 | 6.597560e-01 | |
| pos_patterns.pattern_23 | 28 | | | DNASE_4 | 1.000000e+00 | |
| neg_patterns.pattern_0 | 34 | | | Foxj3.mouse_forkhead_4 | 1.192890e-01 | |
| neg_patterns.pattern_1 | 27 | | | GATA6_MA1104.1 | 6.350240e-02 | |
| neg_patterns.pattern_2 | 26 | | | EGR1_HUMAN.H11MO.0.A | 6.696370e-03 | |
| neg_patterns.pattern_3 | 26 | | | ZN281_HUMAN.H11MO.0.A | 1.594680e-04 | |
