## Supplementary Files 3 for "ChromBPNet: bias factorized, base-resolution deep learning models of chromatin accessibility reveal cis-regulatory sequence syntax, transcription factor footprints and regulatory variants": hepg2_DNASE_raw_bpnet_bias_fold2_counts_modisco.pdf

| pattern | num_seqlets | cwm_fwd | cwm_rev | TOMTOM_match | TOMTOM_qval | TOMTOM_match_logo |
|---|---|---|---|---|---|---|
| pos_patterns.pattern_0 | 8284 | | | DNASE_1 | 2.248810e-03 | |
| pos_patterns.pattern_1 | 7048 | | | DNASE_2 | 8.051420e-05 | |
| pos_patterns.pattern_2 | 1930 | | | DNASE_3 | 2.153620e-04 | |
| pos_patterns.pattern_3 | 1100 | | | SP2_HUMAN.H11MO.0.A | 2.449150e-05 | |
| pos_patterns.pattern_4 | 1011 | | | PRDM6_HUMAN.H11MO.0.C | 9.279250e-02 | |
| pos_patterns.pattern_5 | 673 | | | ZNF384_MA1125.1 | 2.531240e-02 | |
| pos_patterns.pattern_6 | 669 | | | ZNF384_MA1125.1 | 1.176740e-02 | |
| pos_patterns.pattern_7 | 461 | | | SP1_MOUSE.H11MO.0.A | 2.062780e-07 | |
| pos_patterns.pattern_8 | 422 | | | MEF2B_HUMAN.H11MO.0.A | 2.217960e-01 | |
| pos_patterns.pattern_9 | 322 | | | FOXC1_forkhead_1 | 6.186800e-02 | |
| pos_patterns.pattern_10 | 306 | | | TN5_6 | 3.650790e-19 | |
| pos_patterns.pattern_11 | 256 | | | ZN770_HUMAN.H11MO.0.C | 3.455030e-08 | |
| pos_patterns.pattern_12 | 242 | | | RARG_HUMAN.H11MO.0.B | 7.042440e-01 | |
| pos_patterns.pattern_13 | 191 | | | DNASE_5 | 1.644540e-15 | |
| pos_patterns.pattern_14 | 155 | | | KLF4_MA0039.3 | 8.473110e-01 | |
| pos_patterns.pattern_15 | 151 | | | ZN770_HUMAN.H11MO.0.C | 2.559420e-08 | |
| pos_patterns.pattern_16 | 56 | | | IKZF1_HUMAN.H11MO.0.C | 7.480660e-01 | |
| pos_patterns.pattern_17 | 53 | | | ZN770_HUMAN.H11MO.0.C | 1.505930e-06 | |
| pos_patterns.pattern_18 | 51 | | | ZN322_HUMAN.H11MO.0.B | 1.000000e+00 | |
| pos_patterns.pattern_19 | 48 | | | DNASE_2 | 2.637300e-01 | |
| pos_patterns.pattern_20 | 48 | | | EWSR1-FLI1_MA0149.1 | 8.131390e-02 | |
| pos_patterns.pattern_21 | 43 | | | ITF2_HUMAN.H11MO.0.C | 1.000000e+00 | |
| pos_patterns.pattern_22 | 30 | | | ZNF384_MA1125.1 | 3.726430e-01 | |
| neg_patterns.pattern_0 | 41 | | | MAZ_HUMAN.H11MO.0.A | 6.652480e-05 | |
| neg_patterns.pattern_1 | 22 | | | FOXJ3_HUMAN.H11MO.0.A | 7.529350e-02 | |
| neg_patterns.pattern_2 | 22 | | | ZNF384_MA1125.1 | 6.889630e-01 | |
| neg_patterns.pattern_3 | 21 | | | ZNF384_MA1125.1 | 1.970300e-02 | |

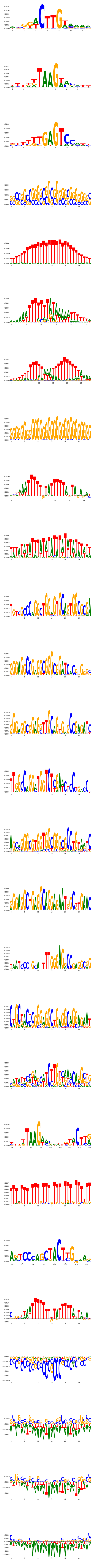

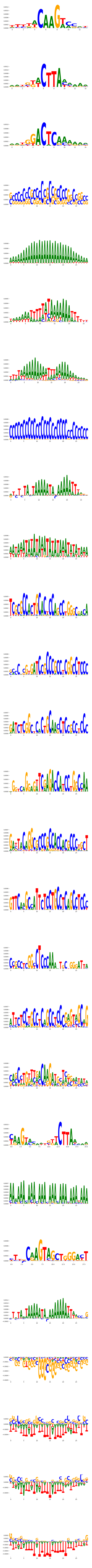

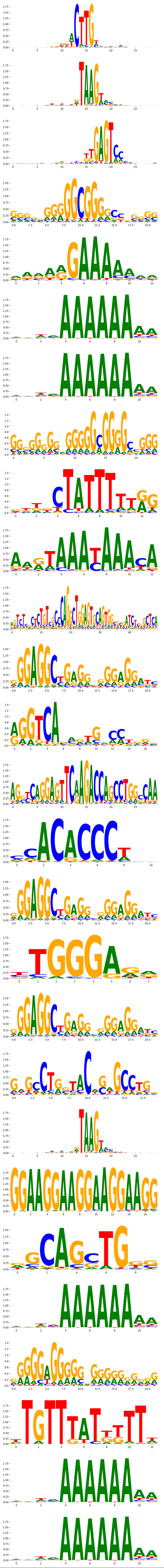
