## Supplementary Files 3 for "ChromBPNet: bias factorized, base-resolution deep learning models of chromatin accessibility reveal cis-regulatory sequence syntax, transcription factor footprints and regulatory variants": hepg2_DNASE_raw_bpnet_bias_fold3_profile_modisco.pdf

| pattern | num_seqlets | cwm_fwd | cwm_rev | TOMTOM_match | TOMTOM_qval | TOMTOM_match_logo |
|---|---|---|---|---|---|---|
| pos_patterns.pattern_0 | 8341 | | | DNASE_1 | 6.687330e-06 | |
| pos_patterns.pattern_1 | 7110 | | | DNASE_2 | 5.230500e-06 | |
| pos_patterns.pattern_2 | 1704 | | | DNASE_3 | 1.665470e-05 | |
| pos_patterns.pattern_3 | 1004 | | | PRDM6_HUMAN.H11MO.0.C | 8.248720e-02 | |
| pos_patterns.pattern_4 | 584 | | | MEF2B_HUMAN.H11MO.0.A | 1.442120e-01 | |
| pos_patterns.pattern_5 | 568 | | | DNASE_2 | 1.277190e-03 | |
| pos_patterns.pattern_6 | 560 | | | ZNF384_MA1125.1 | 2.859830e-01 | |
| pos_patterns.pattern_7 | 500 | | | SP2_HUMAN.H11MO.0.A | 1.941950e-06 | |
| pos_patterns.pattern_8 | 443 | | | SP1_MOUSE.H11MO.0.A | 5.340220e-08 | |
| pos_patterns.pattern_9 | 352 | | | RARG_HUMAN.H11MO.0.B | 1.000000e+00 | |
| pos_patterns.pattern_10 | 293 | | | DNASE_5 | 1.097480e-03 | |
| pos_patterns.pattern_11 | 286 | | | ZNF384_MA1125.1 | 6.332730e-03 | |
| pos_patterns.pattern_12 | 236 | | | ZN770_HUMAN.H11MO.0.C | 8.896830e-08 | |
| pos_patterns.pattern_13 | 228 | | | TN5_6 | 1.509230e-18 | |
| pos_patterns.pattern_14 | 206 | | | ZN770_HUMAN.H11MO.0.C | 2.245180e-08 | |
| pos_patterns.pattern_15 | 204 | | | FOXK1_forkhead_1 | 6.119230e-01 | |
| pos_patterns.pattern_16 | 105 | | | KLF4_MA0039.3 | 6.748550e-01 | |
| pos_patterns.pattern_17 | 92 | | | SP1_MOUSE.H11MO.0.A | 1.599910e-01 | |
| pos_patterns.pattern_18 | 71 | | | ITF2_HUMAN.H11MO.0.C | 1.000000e+00 | |
| pos_patterns.pattern_19 | 69 | | | DNASE_2 | 6.228950e-02 | |
| pos_patterns.pattern_20 | 64 | | | FOXB1_forkhead_2 | 8.627750e-01 | |
| pos_patterns.pattern_21 | 63 | | | PRDM6_HUMAN.H11MO.0.C | 6.924050e-02 | |
| pos_patterns.pattern_22 | 34 | | | NaN | NaN | |
| pos_patterns.pattern_23 | 33 | | | SMCA1_HUMAN.H11MO.0.C | 1.000000e+00 | |
| pos_patterns.pattern_24 | 24 | | | DNASE_4 | 1.619750e-01 | |
