## Supplementary Files 3 for "ChromBPNet: bias factorized, base-resolution deep learning models of chromatin accessibility reveal cis-regulatory sequence syntax, transcription factor footprints and regulatory variants": hepg2_DNASE_raw_bpnet_bias_fold4_counts_modisco.pdf

| pattern | num_seqlets | cwm_fwd | cwm_rev | TOMTOM_match | TOMTOM_qval | TOMTOM_match_logo |
|---|---|---|---|---|---|---|
| pos_patterns.pattern_0 | 8644 | | | DNASE_1 | 1.563260e-05 | |
| pos_patterns.pattern_1 | 7404 | | | DNASE_2 | 4.595570e-05 | |
| pos_patterns.pattern_2 | 1744 | | | DNASE_3 | 4.614350e-05 | |
| pos_patterns.pattern_3 | 1155 | | | ZNF384_MA1125.1 | 8.986210e-02 | |
| pos_patterns.pattern_4 | 604 | | | SP1_MOUSE.H11MO.0.A | 7.067970e-08 | |
| pos_patterns.pattern_5 | 433 | | | ZN770_HUMAN.H11MO.0.C | 7.469870e-07 | |
| pos_patterns.pattern_6 | 428 | | | ZNF384_MA1125.1 | 1.577380e-02 | |
| pos_patterns.pattern_7 | 424 | | | ZNF384_MA1125.1 | 6.927550e-03 | |
| pos_patterns.pattern_8 | 406 | | | ZNF384_MA1125.1 | 4.129360e-02 | |
| pos_patterns.pattern_9 | 328 | | | MEF2B_HUMAN.H11MO.0.A | 2.334880e-01 | |
| pos_patterns.pattern_10 | 294 | | | SP2_HUMAN.H11MO.0.A | 5.131330e-04 | |
| pos_patterns.pattern_11 | 278 | | | DNASE_5 | 8.018960e-04 | |
| pos_patterns.pattern_12 | 229 | | | FOXC1_forkhead_1 | 3.569100e-02 | |
| pos_patterns.pattern_13 | 161 | | | ITF2_HUMAN.H11MO.0.C | 1.000000e+00 | |
| pos_patterns.pattern_14 | 139 | | | FOXK1_forkhead_1 | 1.814010e-01 | |
| pos_patterns.pattern_15 | 133 | | | ZN770_HUMAN.H11MO.0.C | 3.435680e-08 | |
| pos_patterns.pattern_16 | 129 | | | HINFP1_C2H2_2 | 6.911260e-01 | |
| pos_patterns.pattern_17 | 115 | | | TN5_6 | 2.484630e-10 | |
| pos_patterns.pattern_18 | 109 | | | TN5_6 | 1.937960e-10 | |
| pos_patterns.pattern_19 | 104 | | | ZN770_HUMAN.H11MO.0.C | 2.052260e-08 | |
| pos_patterns.pattern_20 | 99 | | | HINFP1_C2H2_2 | 6.626120e-01 | |
| pos_patterns.pattern_21 | 81 | | | DNASE_5 | 1.908840e-03 | |
| pos_patterns.pattern_22 | 49 | | | ITF2_HUMAN.H11MO.0.C | 9.048080e-01 | |
| pos_patterns.pattern_23 | 47 | | | RARA_HUMAN.H11MO.0.A | 3.262920e-01 | |
| pos_patterns.pattern_24 | 45 | | | DNASE_5 | 1.605470e-03 | |
| pos_patterns.pattern_25 | 36 | | | TN5_6 | 6.432370e-17 | |
| pos_patterns.pattern_26 | 26 | | | DNASE_5 | 2.565630e-03 | |
| pos_patterns.pattern_27 | 20 | | | RREB1_MA0073.1 | 1.000000e+00 | |

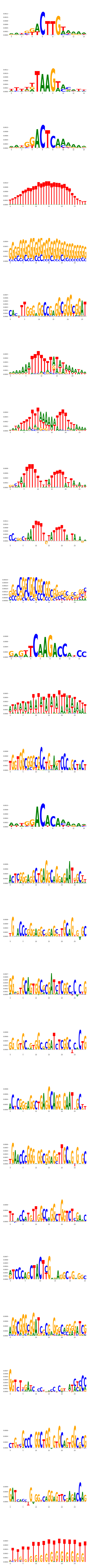
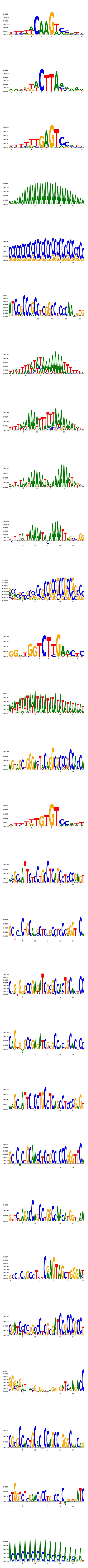
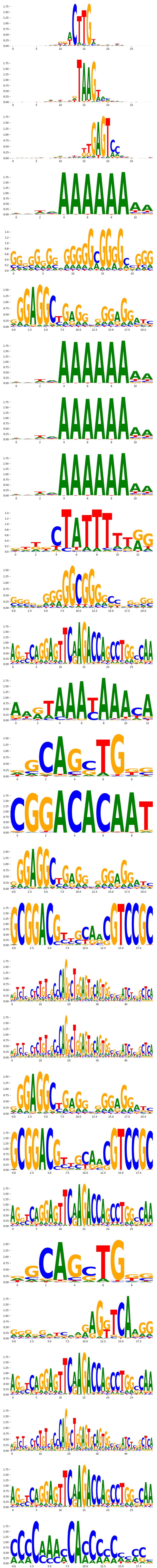
