## Supplementary Files 3 for "ChromBPNet: bias factorized, base-resolution deep learning models of chromatin accessibility reveal cis-regulatory sequence syntax, transcription factor footprints and regulatory variants": imr90_ATAC_raw_bpnet_bias_fold0_counts_modisco.pdf

| pattern | num_seqlets | cwm_fwd | cwm_rev | TOMTOM_match | TOMTOM_qval | TOMTOM_match_logo |
|---|---|---|---|---|---|---|
| pos_patterns.pattern_0 | 4606 | | | SP2_HUMAN.H11MO.0.A | 2.553280e-06 | |
| pos_patterns.pattern_1 | 3585 | | | ZNF384_MA1125.1 | 5.533850e-02 | |
| pos_patterns.pattern_2 | 3245 | | | ZNF384_MA1125.1 | 1.415450e-04 | |
| pos_patterns.pattern_3 | 3192 | | | ZNF384_MA1125.1 | 6.577530e-02 | |
| pos_patterns.pattern_4 | 2881 | | | ZNF384_MA1125.1 | 1.269090e-02 | |
| pos_patterns.pattern_5 | 2495 | | | DNASE_2 | 8.046790e-01 | |
| pos_patterns.pattern_6 | 1992 | | | HXB7_HUMAN.H11MO.0.C | 1.993840e-02 | |
| pos_patterns.pattern_7 | 1726 | | | ZNF384_MA1125.1 | 3.657430e-02 | |
| pos_patterns.pattern_8 | 1225 | | | PBX1_MA0070.1 | 1.000000e+00 | |
| pos_patterns.pattern_9 | 824 | | | ONECUT3_CUT_1 | 3.017140e-01 | |
| pos_patterns.pattern_10 | 520 | | | FLI1_ETS_4 | 8.544260e-01 | |
| pos_patterns.pattern_11 | 430 | | | ZBT7A_HUMAN.H11MO.0.A | 4.922340e-02 | |
| pos_patterns.pattern_12 | 427 | | | FOXO3_forkhead_3 | 1.000000e+00 | |
| pos_patterns.pattern_13 | 129 | | | ZBT7A_HUMAN.H11MO.0.A | 4.892730e-01 | |
| pos_patterns.pattern_14 | 128 | | | DNASE_3 | 5.311230e-01 | |
| pos_patterns.pattern_15 | 81 | | | ZSCAN4_C2H2_1 | 1.000000e+00 | |
| pos_patterns.pattern_16 | 112 | | | NaN | NaN | |
| pos_patterns.pattern_17 | 73 | | | ZBT7A_HUMAN.H11MO.0.A | 3.800490e-02 | |
| pos_patterns.pattern_18 | 45 | | | SP1_HUMAN.H11MO.0.A | 5.224730e-01 | |
| pos_patterns.pattern_19 | 25 | | | ZN274_HUMAN.H11MO.0.A | 1.000000e+00 | |
| neg_patterns.pattern_0 | 6245 | | | TN5_7 | 2.294010e-04 | |
| neg_patterns.pattern_1 | 4664 | | | TN5_2 | 3.166030e-06 | |
| neg_patterns.pattern_2 | 2904 | | | TN5_6 | 0.000000e+00 | |
| neg_patterns.pattern_3 | 2847 | | | ZN467_HUMAN.H11MO.0.C | 5.218860e-08 | |
| neg_patterns.pattern_4 | 1883 | | | P53_MOUSE.H11MO.0.A | 4.170980e-02 | |
| neg_patterns.pattern_5 | 1666 | | | DNASE_5 | 2.866080e-04 | |
| neg_patterns.pattern_6 | 1525 | | | TBX1_TBX_1 | 9.801400e-01 | |
| neg_patterns.pattern_7 | 1469 | | | ZN770_HUMAN.H11MO.0.C | 7.061130e-08 | |
| neg_patterns.pattern_8 | 1259 | | | BARX1_MOUSE.H11MO.0.C | 4.074490e-01 | |
| neg_patterns.pattern_9 | 1027 | | | TN5_6 | 1.549250e-01 | |

| pattern | num_seqlets | cwm_fwd | cwm_rev | TOMTOM_match | TOMTOM_qval | TOMTOM_match_logo |
|---|---|---|---|---|---|---|
| neg_patterns.pattern_10 | 840 | 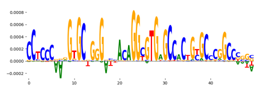 | 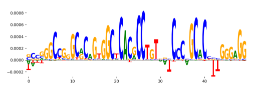 | IKZF1_HUMAN.H11MO.0.C | 6.237710e-01 | 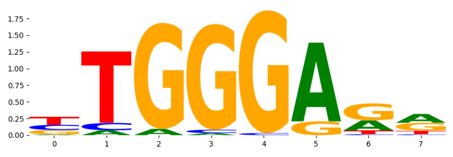 |
| neg_patterns.pattern_11 | 694 | 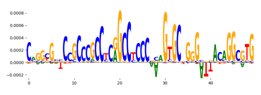 | 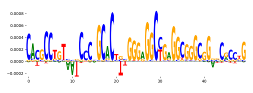 | ZN770_HUMAN.H11MO.0.C | 1.379390e-06 | 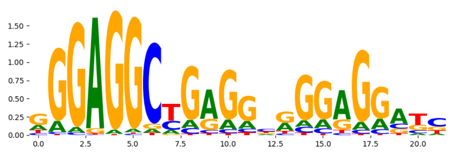 |
| neg_patterns.pattern_12 | 669 | 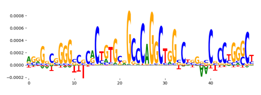 | 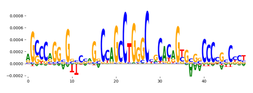 | DNASE_5 | 3.223360e-13 | 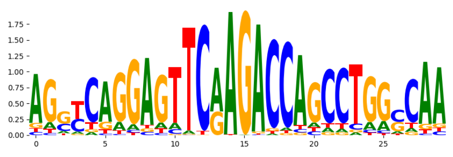 |
| neg_patterns.pattern_13 | 400 | 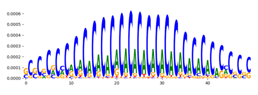 | 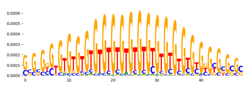 | NaN | NaN | |
| neg_patterns.pattern_14 | 398 | 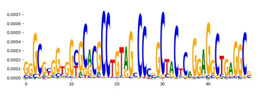 | 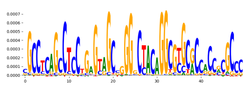 | ITF2_HUMAN.H11MO.0.C | 1.000000e+00 | 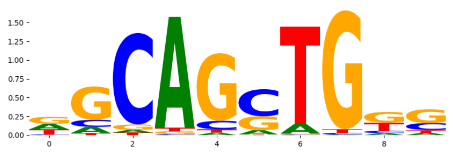 |
| neg_patterns.pattern_15 | 349 | 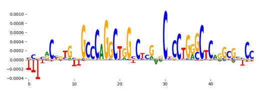 | 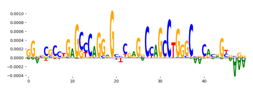 | DNASE_5 | 9.603220e-17 | 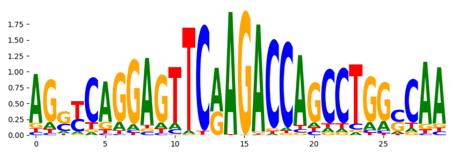 |
| neg_patterns.pattern_16 | 270 | 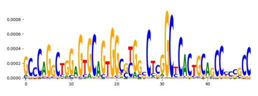 | 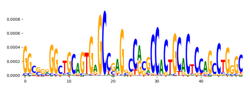 | TN5_6 | 5.215680e-10 | 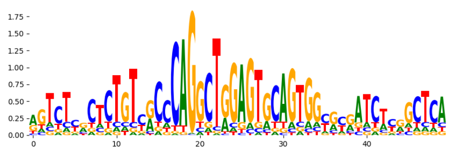 |
| neg_patterns.pattern_17 | 171 | 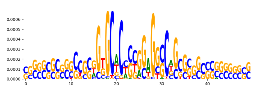 | 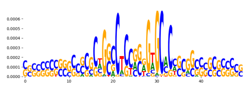 | CTCF_MOUSE.H11MO.0.A | 1.014050e-01 | 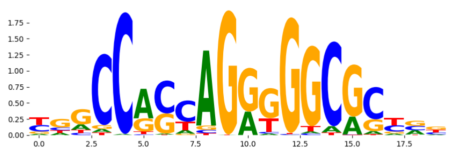 |
| neg_patterns.pattern_18 | 155 | 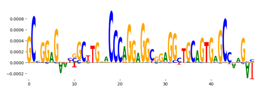 | 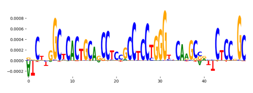 | VDR_MA0693.2 | 1.000000e+00 | 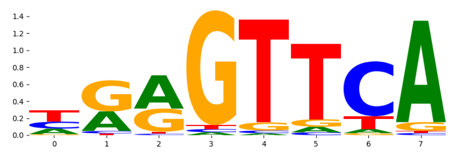 |
| neg_patterns.pattern_19 | 60 | 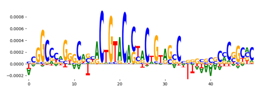 | 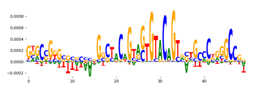 | TN5_4 | 1.000000e+00 | 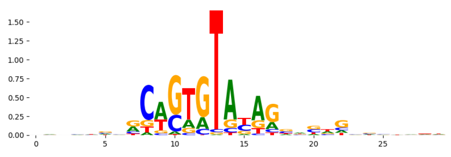 |
| neg_patterns.pattern_20 | 42 | 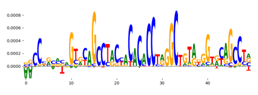 | 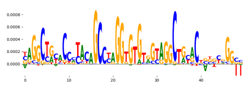 | TBX20_MA0689.1 | 4.554620e-01 | 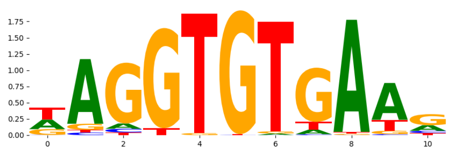 |
| neg_patterns.pattern_21 | 36 | 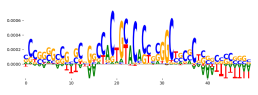 | 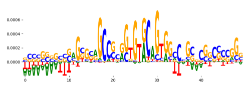 | ZNF232_C2H2_1 | 1.000000e+00 | 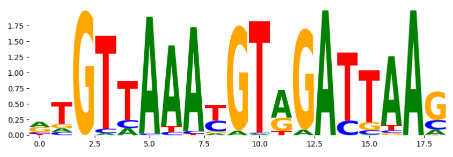 |
| neg_patterns.pattern_22 | 24 | 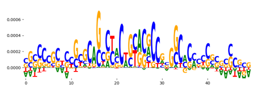 | 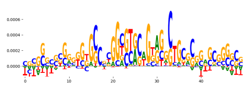 | Ar.mouse_nuclearreceptor_1 | 1.000000e+00 | 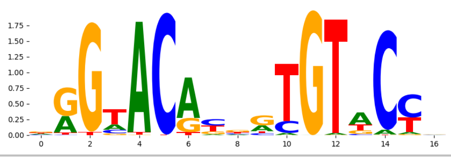 |
