## Supplementary Files 3 for "ChromBPNet: bias factorized, base-resolution deep learning models of chromatin accessibility reveal cis-regulatory sequence syntax, transcription factor footprints and regulatory variants": imr90_ATAC_raw_bpnet_bias_fold0_profile_modisco.pdf

| pattern | num_seqlets | cwm_fwd | cwm_rev | TOMTOM_match | TOMTOM_qval | TOMTOM_match_logo |
|---|---|---|---|---|---|---|
| pos_patterns.pattern_0 | 12385 | 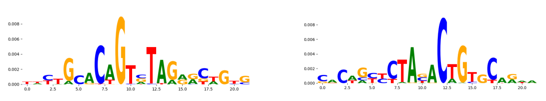 | | TN5_8 | 2.297410e-02 | 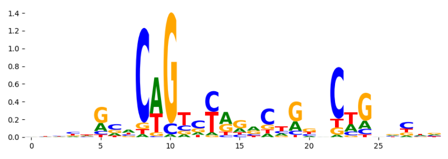 |
| pos_patterns.pattern_1 | 9636 | 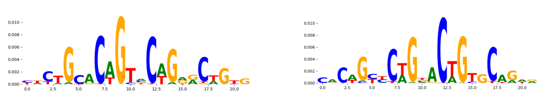 | | TN5_1 | 4.496600e-10 | 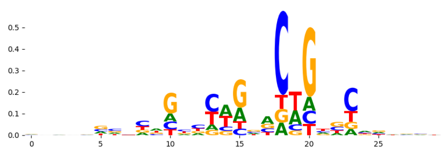 |
| pos_patterns.pattern_2 | 4321 | 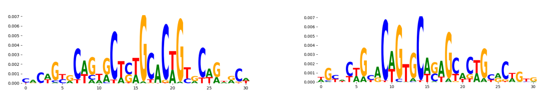 | | TN5_2 | 1.962610e-20 | 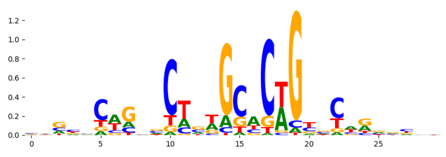 |
| pos_patterns.pattern_3 | 3255 | 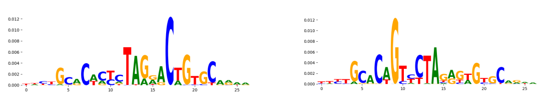 | | TN5_3 | 3.241600e-07 | 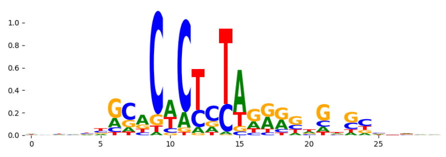 |
| pos_patterns.pattern_4 | 2952 | 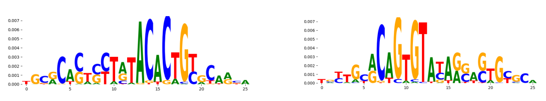 | | TN5_4 | 1.535820e-09 | 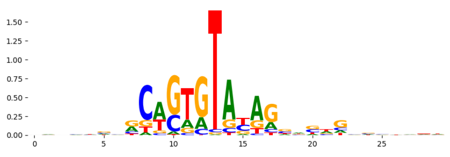 |
| pos_patterns.pattern_5 | 2722 | 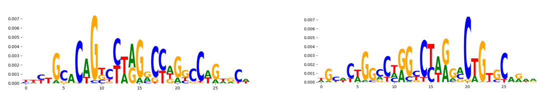 | | TN5_1 | 4.538340e-04 | 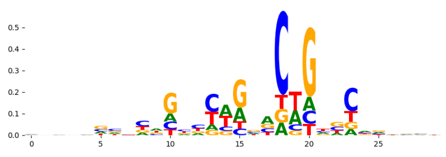 |
| pos_patterns.pattern_6 | 1671 | 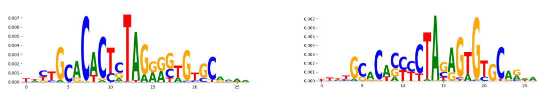 | | TN5_3 | 1.974290e-17 | 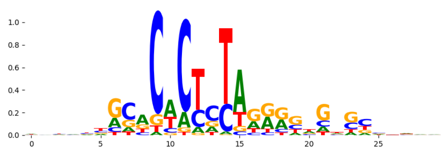 |
| pos_patterns.pattern_7 | 1647 | 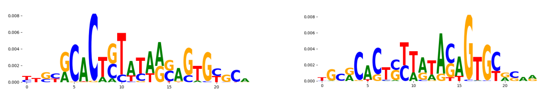 | | TN5_4 | 1.316630e-03 | 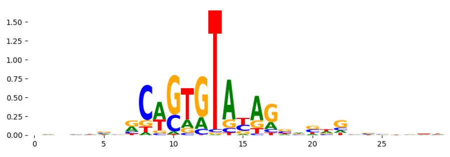 |
| pos_patterns.pattern_8 | 1581 | 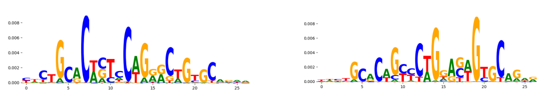 | | TN5_3 | 1.152750e-07 | 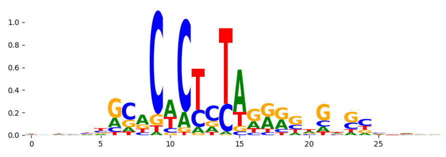 |
| pos_patterns.pattern_9 | 1140 | 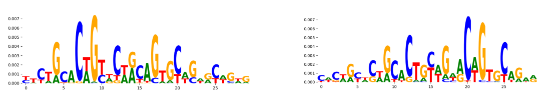 | | TN5_7 | 1.126780e-12 | 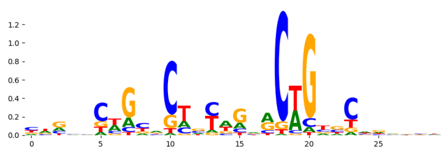 |
| pos_patterns.pattern_10 | 884 | 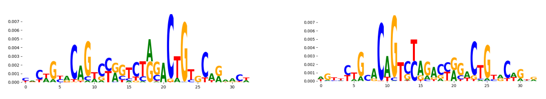 | | TN5_8 | 3.596620e-13 | 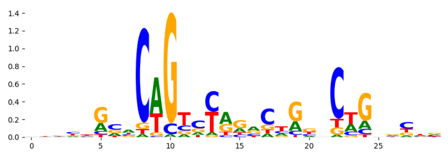 |
| pos_patterns.pattern_11 | 645 | 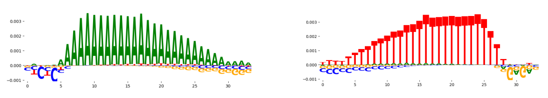 | | ZNF384_MA1125.1 | 7.645990e-02 | 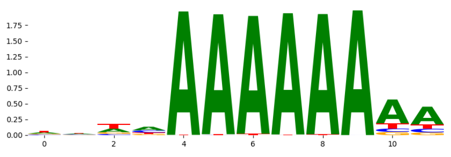 |
| pos_patterns.pattern_12 | 640 | 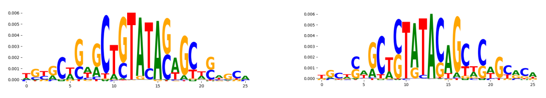 | | TN5_4 | 8.121710e-03 | 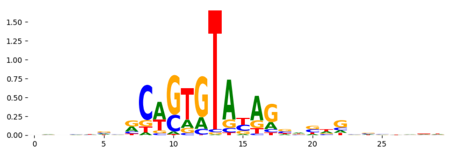 |
| pos_patterns.pattern_13 | 275 | 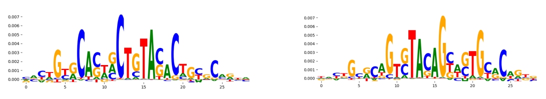 | | TN5_4 | 4.985440e-04 | 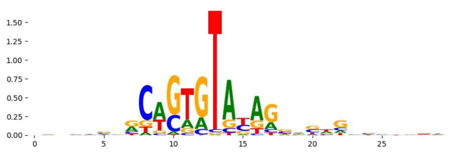 |
| pos_patterns.pattern_14 | 158 | 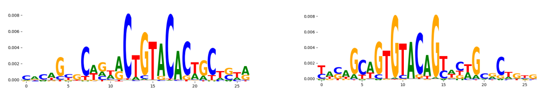 | | TN5_4 | 7.326300e-04 | 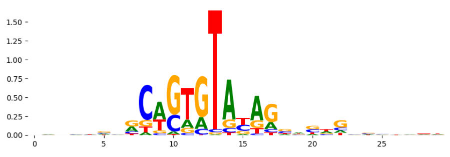 |
| pos_patterns.pattern_15 | 58 | 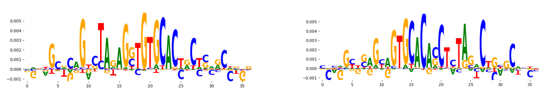 | | TCF4_bHLH_2 | 6.139190e-01 | 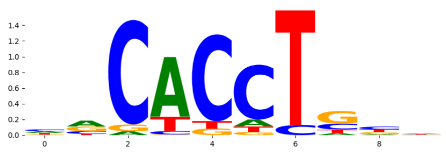 |
| pos_patterns.pattern_16 | 47 | 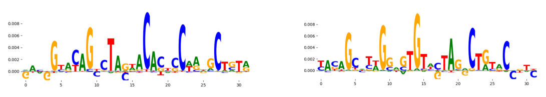 | | TBX20_MA0689.1 | 2.600740e-01 | 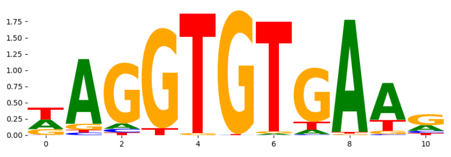 |
| pos_patterns.pattern_17 | 49 | 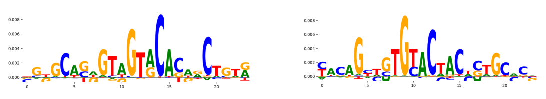 | | TP63_MA0525.2 | 6.005200e-01 | 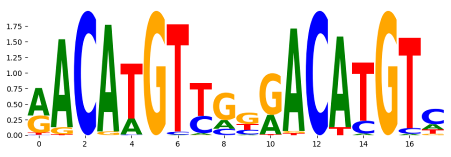 |
| pos_patterns.pattern_18 | 31 | 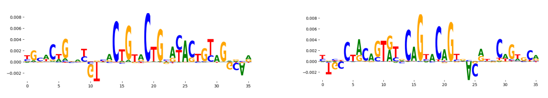 | | OVOL1_HUMAN.H11MO.0.C | 1.000000e+00 | 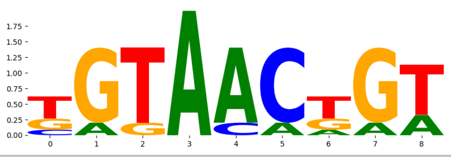 |
