## Supplementary Files 3 for "ChromBPNet: bias factorized, base-resolution deep learning models of chromatin accessibility reveal cis-regulatory sequence syntax, transcription factor footprints and regulatory variants": imr90_ATAC_raw_bpnet_bias_fold1_counts_modisco.pdf

| pattern | num_seqlets | cwm_fwd | cwm_rev | TOMTOM_match | TOMTOM_qval | TOMTOM_match_logo |
|---|---|---|---|---|---|---|
| pos_patterns.pattern_0 | 3832 | | | ZNF384_MA1125.1 | 1.000000e+00 | |
| pos_patterns.pattern_1 | 2085 | | | ZNF384_MA1125.1 | 1.686180e-02 | |
| pos_patterns.pattern_2 | 1909 | | | ZNF384_MA1125.1 | 1.237820e-02 | |
| pos_patterns.pattern_3 | 1604 | | | ZNF384_MA1125.1 | 8.381310e-03 | |
| pos_patterns.pattern_4 | 1484 | | | ZNF384_MA1125.1 | 5.512490e-02 | |
| pos_patterns.pattern_5 | 1294 | | | SP3_HUMAN.H11MO.0.B | 1.095760e-05 | |
| pos_patterns.pattern_6 | 1165 | | | ZNF384_MA1125.1 | 2.662260e-02 | |
| pos_patterns.pattern_7 | 1119 | | | SP1_HUMAN.H11MO.0.A | 5.476220e-02 | |
| pos_patterns.pattern_8 | 1116 | | | ZNF384_MA1125.1 | 6.452350e-02 | |
| pos_patterns.pattern_9 | 988 | | | SP1_HUMAN.H11MO.0.A | 2.510720e-02 | |
| pos_patterns.pattern_10 | 886 | | | SP1_HUMAN.H11MO.0.A | 1.312890e-02 | |
| pos_patterns.pattern_11 | 854 | | | RUNX3_RUNX_2 | 5.595720e-01 | |
| pos_patterns.pattern_12 | 637 | | | ZBTB7A_C2H2_1 | 6.435600e-01 | |
| pos_patterns.pattern_13 | 522 | | | RELA_MA0107.1 | 6.012060e-01 | |
| pos_patterns.pattern_14 | 397 | | | ZFX_MOUSE.H11MO.0.B | 7.219730e-05 | |
| pos_patterns.pattern_15 | 327 | | | SP1_HUMAN.H11MO.0.A | 5.048380e-03 | |
| pos_patterns.pattern_16 | 289 | | | ZN281_MOUSE.H11MO.0.A | 4.556590e-02 | |
| pos_patterns.pattern_17 | 228 | | | ZBT7A_HUMAN.H11MO.0.A | 8.670540e-01 | |
| pos_patterns.pattern_18 | 219 | | | ZNF384_MA1125.1 | 1.155050e-02 | |
| pos_patterns.pattern_19 | 126 | | | ZIC3_C2H2_1 | 1.000000e+00 | |
| pos_patterns.pattern_20 | 80 | | | MXI1_HUMAN.H11MO.0.A | 5.701390e-02 | |
| pos_patterns.pattern_21 | 68 | | | GLIS2_C2H2_1 | 8.487280e-01 | |
| pos_patterns.pattern_22 | 34 | | | CUX2_MOUSE.H11MO.0.C | 8.024920e-02 | |
| pos_patterns.pattern_23 | 26 | | | THA11_MOUSE.H11MO.0.B | 1.000000e+00 | |
| pos_patterns.pattern_24 | 24 | | | PRDM6_HUMAN.H11MO.0.C | 8.004980e-02 | |
| neg_patterns.pattern_0 | 4412 | | | TN5_1 | 1.486920e-02 | |
| neg_patterns.pattern_1 | 3899 | | | TN5_8 | 3.239990e-01 | |
| neg_patterns.pattern_2 | 3716 | | | TN5_2 | 1.172430e-06 | |
| neg_patterns.pattern_3 | 810 | | | TN5_6 | 1.103230e-24 | |

| pattern | num_seqlets | cwm_fwd | cwm_rev | TOMTOM_match | TOMTOM_qval | TOMTOM_match_logo |
|---|---|---|---|---|---|---|
| neg_patterns.pattern_4 | 433 | | | RREB1_MA0073.1 | 1.000000e+00 | |
| neg_patterns.pattern_5 | 396 | | | ZNF18_HUMAN.H11MO.0.C | 3.567590e-01 | |
| neg_patterns.pattern_6 | 384 | | | TN5_6 | 2.267270e-04 | |
| neg_patterns.pattern_7 | 364 | | | EGR1_MOUSE.H11MO.0.A | 7.295390e-03 | |
| neg_patterns.pattern_8 | 364 | | | IKZF1_HUMAN.H11MO.0.C | 9.132780e-01 | |
| neg_patterns.pattern_9 | 363 | | | TN5_2 | 2.505500e-03 | |
| neg_patterns.pattern_10 | 310 | | | TN5_4 | 2.262460e-02 | |
| neg_patterns.pattern_11 | 272 | | | MGA_MA0801.1 | 2.546510e-01 | |
| neg_patterns.pattern_12 | 248 | | | ZN121_HUMAN.H11MO.0.C | 3.498450e-03 | |
| neg_patterns.pattern_13 | 242 | | | Hes1_MA1099.1 | 1.857480e-01 | |
| neg_patterns.pattern_14 | 233 | | | SMAD3_HUMAN.H11MO.0.B | 4.282920e-01 | |
| neg_patterns.pattern_15 | 208 | | | TN5_6 | 7.863910e-03 | |
| neg_patterns.pattern_16 | 202 | | | TN5_2 | 4.674830e-02 | |
| neg_patterns.pattern_17 | 194 | | | TN5_6 | 6.511850e-05 | |
| neg_patterns.pattern_18 | 160 | | | ZSCAN4_C2H2_1 | 1.000000e+00 | |
| neg_patterns.pattern_19 | 157 | | | BRAC_MOUSE.H11MO.0.B | 1.513440e-01 | |
| neg_patterns.pattern_20 | 156 | | | THA_HUMAN.H11MO.0.C | 8.024990e-02 | |
| neg_patterns.pattern_21 | 116 | | | TN5_8 | 3.033490e-02 | |
| neg_patterns.pattern_22 | 71 | | | TN5_6 | 4.697640e-06 | |
| neg_patterns.pattern_23 | 50 | | | TN5_6 | 1.000000e+00 | |
| neg_patterns.pattern_24 | 42 | | | DNASE_5 | 1.609920e-04 | |
| neg_patterns.pattern_25 | 24 | | | HIC2_C2H2_1 | 8.095220e-01 | |
| neg_patterns.pattern_26 | 21 | | | KLF4_MA0039.3 | 1.164220e-01 | |
| neg_patterns.pattern_27 | 20 | | | SOX9_HMG_2 | 4.795950e-01 | |
