## Supplementary Files 3 for "ChromBPNet: bias factorized, base-resolution deep learning models of chromatin accessibility reveal cis-regulatory sequence syntax, transcription factor footprints and regulatory variants": imr90_ATAC_raw_bpnet_bias_fold1_profile_modisco.pdf

| pattern | num_seqlets | cwm_fwd | cwm_rev | TOMTOM_match | TOMTOM_qval | TOMTOM_match_logo |
|---|---|---|---|---|---|---|
| pos_patterns.pattern_0 | 9518 | | | TN5_1 | 2.045570e-08 | |
| pos_patterns.pattern_1 | 5938 | | | TN5_4 | 2.607330e-02 | |
| pos_patterns.pattern_2 | 4189 | | | TN5_1 | 3.137040e-06 | |
| pos_patterns.pattern_3 | 3125 | | | TN5_3 | 1.022420e-03 | |
| pos_patterns.pattern_4 | 3122 | | | KLF4_MA0039.3 | 7.096000e-01 | |
| pos_patterns.pattern_5 | 2641 | | | TN5_3 | 1.260580e-12 | |
| pos_patterns.pattern_6 | 718 | | | TN5_3 | 9.027640e-03 | |
| pos_patterns.pattern_7 | 603 | | | TN5_3 | 7.588830e-09 | |
| pos_patterns.pattern_8 | 591 | | | TN5_3 | 2.416740e-13 | |
| pos_patterns.pattern_9 | 400 | | | TN5_3 | 6.377630e-07 | |
| pos_patterns.pattern_10 | 296 | | | PRDM6_HUMAN.H11MO.0.C | 6.468200e-02 | |
| pos_patterns.pattern_11 | 206 | | | RREB1_MA0073.1 | 1.000000e+00 | |
| pos_patterns.pattern_12 | 132 | | | FOXA2_HUMAN.H11MO.0.A | 9.928580e-01 | |
| pos_patterns.pattern_13 | 92 | | | HOXC13_homeodomain_1 | 6.092230e-01 | |
| pos_patterns.pattern_14 | 82 | | | ZN121_HUMAN.H11MO.0.C | 3.194300e-04 | |
| pos_patterns.pattern_15 | 64 | | | ZN121_HUMAN.H11MO.0.C | 2.965610e-05 | |
| pos_patterns.pattern_16 | 46 | | | ZNF384_MA1125.1 | 2.493950e-02 | |
| pos_patterns.pattern_17 | 20 | | | TN5_3 | 3.860090e-02 | |
