## Supplementary Files 3 for "ChromBPNet: bias factorized, base-resolution deep learning models of chromatin accessibility reveal cis-regulatory sequence syntax, transcription factor footprints and regulatory variants": imr90_ATAC_raw_bpnet_bias_fold2_counts_modisco.pdf

| pattern | num_seqlets | cwm_fwd | cwm_rev | TOMTOM_match | TOMTOM_qval | TOMTOM_match_logo |
|---|---|---|---|---|---|---|
| pos_patterns.pattern_0 | 2930 | | | ZNF384_MA1125.1 | 9.681010e-02 | |
| pos_patterns.pattern_1 | 2920 | | | ZNF384_MA1125.1 | 3.919540e-03 | |
| pos_patterns.pattern_2 | 2908 | | | ZNF384_MA1125.1 | 1.574520e-03 | |
| pos_patterns.pattern_3 | 2902 | | | FOXJ3_HUMAN.H11MO.0.A | 3.902680e-02 | |
| pos_patterns.pattern_4 | 2555 | | | ZNF384_MA1125.1 | 7.314620e-02 | |
| pos_patterns.pattern_5 | 2465 | | | DNASE_2 | 2.578700e-01 | |
| pos_patterns.pattern_6 | 1257 | | | MAZ_HUMAN.H11MO.0.A | 2.058930e-04 | |
| pos_patterns.pattern_7 | 806 | | | ZFX_MOUSE.H11MO.0.B | 8.234770e-03 | |
| pos_patterns.pattern_8 | 752 | | | ZBT7A_HUMAN.H11MO.0.A | 1.660590e-02 | |
| pos_patterns.pattern_9 | 629 | | | DNASE_2 | 1.675300e-01 | |
| pos_patterns.pattern_10 | 572 | | | ZNF384_MA1125.1 | 8.056040e-03 | |
| pos_patterns.pattern_11 | 509 | | | NFIX_NFI_4 | 1.000000e+00 | |
| pos_patterns.pattern_12 | 479 | | | SP3_HUMAN.H11MO.0.B | 2.686780e-04 | |
| pos_patterns.pattern_13 | 360 | | | KLF9_HUMAN.H11MO.0.C | 9.399620e-01 | |
| pos_patterns.pattern_14 | 257 | | | SP2_HUMAN.H11MO.0.A | 1.163550e-04 | |
| pos_patterns.pattern_15 | 252 | | | RREB1_MA0073.1 | 3.139460e-05 | |
| pos_patterns.pattern_16 | 194 | | | SP1_HUMAN.H11MO.0.A | 3.069150e-05 | |
| pos_patterns.pattern_17 | 180 | | | CTCFL_MOUSE.H11MO.0.A | 1.797100e-02 | |
| pos_patterns.pattern_18 | 52 | | | POU2F2_MA0507.1 | 3.067460e-01 | |
| pos_patterns.pattern_19 | 43 | | | FOXD2_forkhead_1 | 5.494860e-02 | |
| pos_patterns.pattern_20 | 25 | | | Nfe2l2_MA0150.2 | 1.000000e+00 | |
| neg_patterns.pattern_0 | 3581 | | | TN5_2 | 2.479720e-03 | |
| neg_patterns.pattern_1 | 3308 | | | TN5_1 | 1.219860e-05 | |
| neg_patterns.pattern_2 | 2277 | | | TN5_1 | 6.221840e-03 | |
| neg_patterns.pattern_3 | 1713 | | | TN5_2 | 3.384470e-04 | |
| neg_patterns.pattern_4 | 806 | | | TN5_6 | 3.592370e-22 | |
| neg_patterns.pattern_5 | 579 | | | TN5_8 | 8.905280e-04 | |
| neg_patterns.pattern_6 | 577 | | | FOS_HUMAN.H11MO.0.A | 1.000000e+00 | |
| neg_patterns.pattern_7 | 478 | | | TN5_4 | 1.598690e-02 | |

| pattern | num_seqlets | cwm_fwd | cwm_rev | TOMTOM_match | TOMTOM_qval | TOMTOM_match_logo |
|---|---|---|---|---|---|---|
| neg_patterns.pattern_8 | 393 | | | ZN770_HUMAN.H11MO.0.C | 1.960270e-07 | |
| neg_patterns.pattern_9 | 352 | | | TN5_2 | 5.511530e-02 | |
| neg_patterns.pattern_10 | 293 | | | RARG_HUMAN.H11MO.0.B | 1.000000e+00 | |
| neg_patterns.pattern_11 | 262 | | | Tcfl5_MA0632.1 | 5.869820e-01 | |
| neg_patterns.pattern_12 | 173 | | | ZN770_HUMAN.H11MO.0.C | 1.042920e-07 | |
| neg_patterns.pattern_13 | 110 | | | DNASE_5 | 1.111950e-07 | |
| neg_patterns.pattern_14 | 92 | | | TN5_1 | 3.020640e-02 | |
| neg_patterns.pattern_15 | 74 | | | TFAP2C_TFAP_6 | 3.242090e-01 | |
| neg_patterns.pattern_16 | 50 | | | RUNX3_MOUSE.H11MO.0.A | 9.944240e-01 | |
| neg_patterns.pattern_17 | 47 | | | EGR2_HUMAN.H11MO.0.A | 3.985070e-02 | |
| neg_patterns.pattern_18 | 36 | | | EGR1_HUMAN.H11MO.0.A | 2.161950e-01 | |
| neg_patterns.pattern_19 | 35 | | | DNASE_5 | 5.545120e-01 | |
| neg_patterns.pattern_20 | 34 | | | SMAD3_HUMAN.H11MO.0.B | 2.377830e-01 | |
| neg_patterns.pattern_21 | 34 | | | DNASE_5 | 4.803090e-01 | |
| neg_patterns.pattern_22 | 28 | | | SOX21_HMG_2 | 6.000370e-01 | |
| neg_patterns.pattern_23 | 28 | | | KLF5_MOUSE.H11MO.0.A | 3.100630e-01 | |
