## Supplementary Files 3 for "ChromBPNet: bias factorized, base-resolution deep learning models of chromatin accessibility reveal cis-regulatory sequence syntax, transcription factor footprints and regulatory variants": imr90_ATAC_raw_bpnet_bias_fold2_profile_modisco.pdf

| pattern | num_seqlets | cwm_fwd | cwm_rev | TOMTOM_match | TOMTOM_qval | TOMTOM_match_logo |
|---|---|---|---|---|---|---|
| pos_patterns.pattern_0 | 8154 | | | TN5_1 | 3.495660e-08 | |
| pos_patterns.pattern_1 | 5724 | | | TN5_4 | 2.522990e-02 | |
| pos_patterns.pattern_2 | 4624 | | | TN5_1 | 7.306880e-05 | |
| pos_patterns.pattern_3 | 3856 | | | KLF4_MA0039.3 | 8.149190e-01 | |
| pos_patterns.pattern_4 | 2580 | | | TN5_2 | 1.341290e-07 | |
| pos_patterns.pattern_5 | 1645 | | | TN5_3 | 7.233370e-11 | |
| pos_patterns.pattern_6 | 1619 | | | TN5_1 | 4.388730e-02 | |
| pos_patterns.pattern_7 | 698 | | | TN5_3 | 7.475330e-05 | |
| pos_patterns.pattern_8 | 684 | | | TN5_3 | 1.464640e-05 | |
| pos_patterns.pattern_9 | 579 | | | TN5_3 | 1.796540e-05 | |
| pos_patterns.pattern_10 | 383 | | | PRDM6_HUMAN.H11MO.0.C | 8.213510e-02 | |
| pos_patterns.pattern_11 | 235 | | | TN5_8 | 1.000000e+00 | |
| pos_patterns.pattern_12 | 225 | | | RREB1_MA0073.1 | 1.000000e+00 | |
| pos_patterns.pattern_13 | 170 | | | TN5_6 | 4.246270e-03 | |
| pos_patterns.pattern_14 | 78 | | | RORA_MA0072.1 | 1.000000e+00 | |
| pos_patterns.pattern_15 | 64 | | | TN5_3 | 1.175170e-02 | |
