## Supplementary Files 3 for "ChromBPNet: bias factorized, base-resolution deep learning models of chromatin accessibility reveal cis-regulatory sequence syntax, transcription factor footprints and regulatory variants": imr90_ATAC_raw_bpnet_bias_fold3_counts_modisco.pdf

| pattern | num_seqlets | cwm_fwd | cwm_rev | TOMTOM_match | TOMTOM_qval | TOMTOM_match_logo |
|---|---|---|---|---|---|---|
| pos_patterns.pattern_0 | 5360 | | | ZNF384_MA1125.1 | 2.032130e-02 | |
| pos_patterns.pattern_1 | 3680 | | | NaN | NaN | |
| pos_patterns.pattern_2 | 3304 | | | CUX2_CUT_1 | 2.000880e-01 | |
| pos_patterns.pattern_3 | 2481 | | | ZNF384_MA1125.1 | 2.314530e-02 | |
| pos_patterns.pattern_4 | 1437 | | | HNF6_HUMAN.H11MO.0.B | 1.088070e-01 | |
| pos_patterns.pattern_5 | 1273 | | | DNASE_2 | 1.070430e-02 | |
| pos_patterns.pattern_6 | 731 | | | ANDR_HUMAN.H11MO.0.A | 3.917290e-02 | |
| pos_patterns.pattern_7 | 631 | | | SP2_HUMAN.H11MO.0.A | 6.045480e-03 | |
| pos_patterns.pattern_8 | 495 | | | SP2_HUMAN.H11MO.0.A | 2.477800e-02 | |
| pos_patterns.pattern_9 | 463 | | | ZNF384_MA1125.1 | 1.283060e-03 | |
| pos_patterns.pattern_10 | 277 | | | PBX3_MOUSE.H11MO.0.A | 5.018230e-01 | |
| pos_patterns.pattern_11 | 231 | | | ZNF384_MA1125.1 | 6.055020e-01 | |
| pos_patterns.pattern_12 | 214 | | | SP3_HUMAN.H11MO.0.B | 8.195580e-03 | |
| pos_patterns.pattern_13 | 152 | | | SP2_HUMAN.H11MO.0.A | 1.597500e-06 | |
| pos_patterns.pattern_14 | 150 | | | DNASE_2 | 4.608560e-01 | |
| pos_patterns.pattern_15 | 135 | | | ANDR_HUMAN.H11MO.0.A | 4.902010e-03 | |
| pos_patterns.pattern_16 | 116 | | | AP2B_HUMAN.H11MO.0.B | 1.213230e-01 | |
| pos_patterns.pattern_17 | 107 | | | INSM1_MA0155.1 | 1.104890e-01 | |
| pos_patterns.pattern_18 | 94 | | | PDX1_HUMAN.H11MO.0.A | 2.116730e-01 | |
| pos_patterns.pattern_19 | 71 | | | FOXL1_forkhead_2 | 1.000000e+00 | |
| pos_patterns.pattern_20 | 69 | | | NFAC1_HUMAN.H11MO.0.B | 3.373280e-01 | |
| pos_patterns.pattern_21 | 46 | | | NFAC2_HUMAN.H11MO.0.B | 2.725480e-01 | |
| pos_patterns.pattern_22 | 36 | | | TBX2_TBX_1 | 4.212570e-02 | |
| pos_patterns.pattern_23 | 34 | | | ZN563_HUMAN.H11MO.0.C | 1.000000e+00 | |
| pos_patterns.pattern_24 | 25 | | | MESP1_bHLH_1 | 1.000000e+00 | |
| pos_patterns.pattern_25 | 21 | | | Foxj3.mouse_forkhead_2 | 8.732700e-01 | |
| neg_patterns.pattern_0 | 2840 | | | TN5_4 | 1.948740e-06 | |
| neg_patterns.pattern_1 | 2695 | | | TN5_8 | 9.045920e-01 | |
| neg_patterns.pattern_2 | 1545 | | | SP5_MOUSE.H11MO.0.C | 3.253620e-05 | |
| neg_patterns.pattern_3 | 951 | | | TN5_1 | 3.702540e-03 | |

| pattern | num_seqlets | cwm_fwd | cwm_rev | TOMTOM_match | TOMTOM_qval | TOMTOM_match_logo |
|---|---|---|---|---|---|---|
| neg_patterns.pattern_4 | 769 | 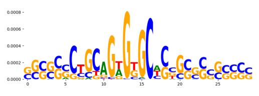 | 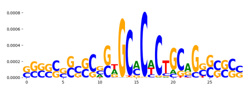 | TN5_2 | 1.276540e-01 | 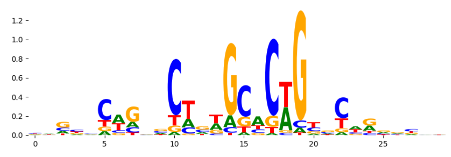 |
| neg_patterns.pattern_5 | 439 | 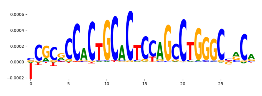 | 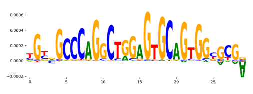 | TN5_6 | 2.175850e-20 | 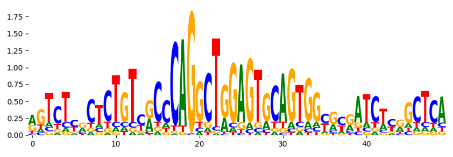 |
| neg_patterns.pattern_6 | 302 | 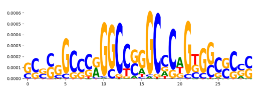 | 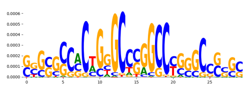 | FOS_HUMAN.H11MO.0.A | 1.000000e+00 | 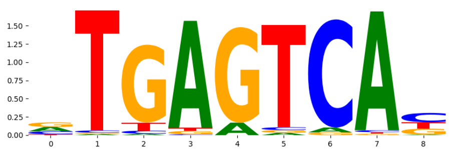 |
| neg_patterns.pattern_7 | 301 | 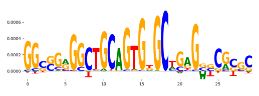 | 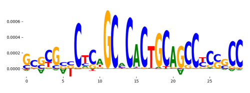 | RARG_HUMAN.H11MO.0.B | 1.000000e+00 | 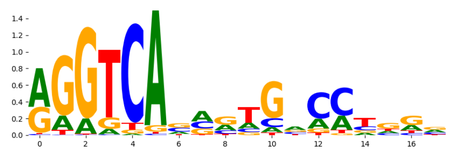 |
| neg_patterns.pattern_8 | 259 | 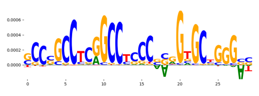 | 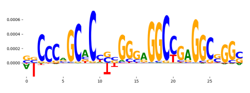 | ZN770_HUMAN.H11MO.0.C | 4.583540e-02 | 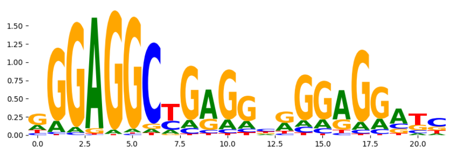 |
| neg_patterns.pattern_9 | 247 | 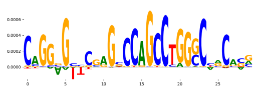 | 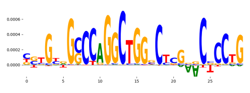 | DNASE_5 | 8.250310e-13 | 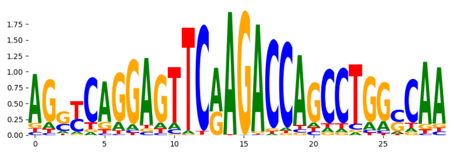 |
| neg_patterns.pattern_10 | 223 | 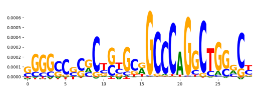 | 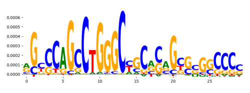 | ZN121_HUMAN.H11MO.0.C | 1.487870e-08 | 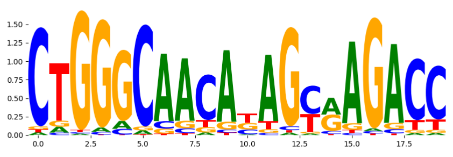 |
| neg_patterns.pattern_11 | 212 | 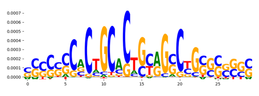 | 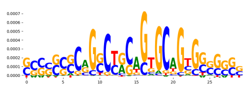 | TN5_6 | 5.425760e-02 | 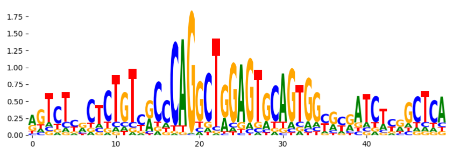 |
| neg_patterns.pattern_12 | 134 | 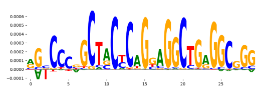 | 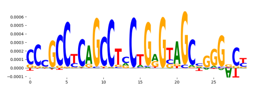 | ZN770_HUMAN.H11MO.0.C | 3.730220e-01 | 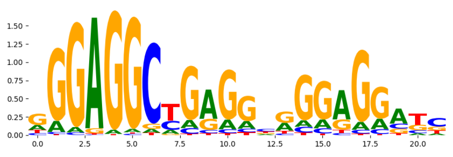 |
| neg_patterns.pattern_13 | 65 | 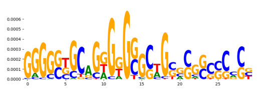 | 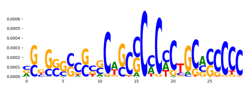 | ZN341_HUMAN.H11MO.0.C | 2.211780e-01 | 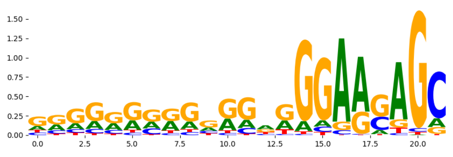 |
| neg_patterns.pattern_14 | 52 | 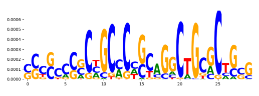 | 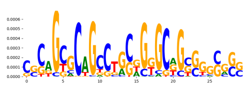 | CTCF_MA0139.1 | 6.181050e-01 | 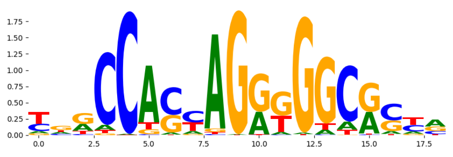 |
| neg_patterns.pattern_15 | 38 | 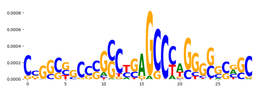 | 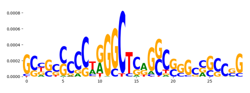 | TN5_1 | 8.383400e-01 | 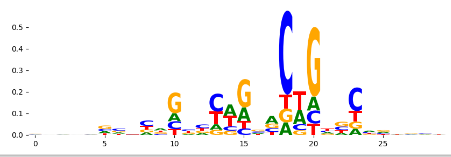 |
