## Supplementary Files 3 for "ChromBPNet: bias factorized, base-resolution deep learning models of chromatin accessibility reveal cis-regulatory sequence syntax, transcription factor footprints and regulatory variants": imr90_ATAC_raw_bpnet_bias_fold3_profile_modisco.pdf

| pattern | num_seqlets | cwm_fwd | cwm_rev | TOMTOM_match | TOMTOM_qval | TOMTOM_match_logo |
|---|---|---|---|---|---|---|
| pos_patterns.pattern_0 | 9972 | 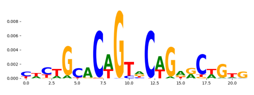 | 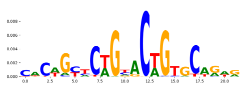 | TN5_2 | 9.959580e-10 | 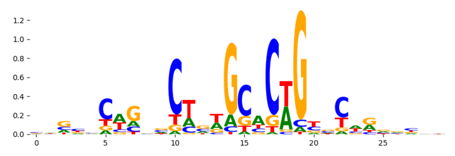 |
| pos_patterns.pattern_1 | 6642 | 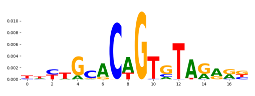 | 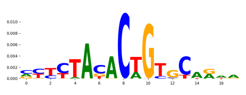 | TN5_4 | 2.181650e-02 | 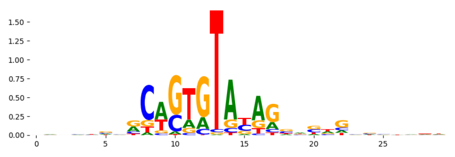 |
| pos_patterns.pattern_2 | 4680 | 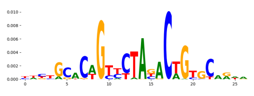 | 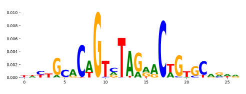 | TN5_1 | 1.621050e-06 | 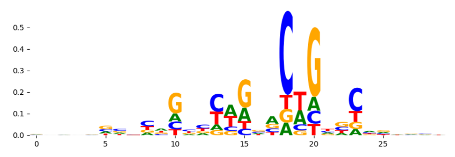 |
| pos_patterns.pattern_3 | 3879 | 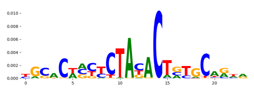 | 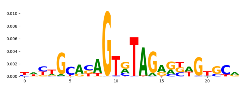 | TN5_3 | 2.762170e-01 | 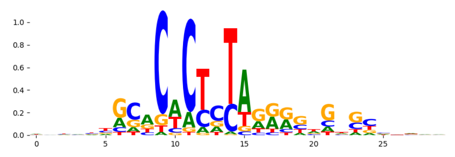 |
| pos_patterns.pattern_4 | 2863 | 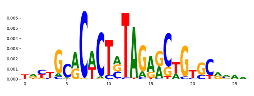 | 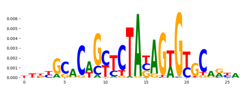 | TN5_3 | 2.195470e-10 | 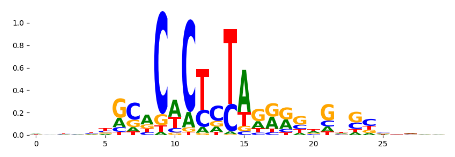 |
| pos_patterns.pattern_5 | 1502 | 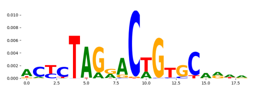 | 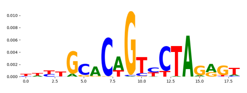 | TN5_7 | 2.439630e-02 | 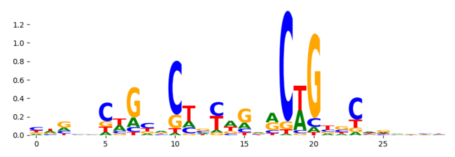 |
| pos_patterns.pattern_6 | 898 | 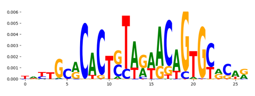 | 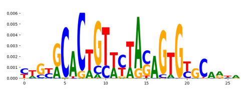 | TN5_3 | 6.512600e-07 | 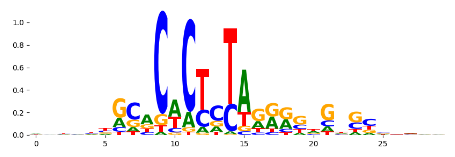 |
| pos_patterns.pattern_7 | 689 | 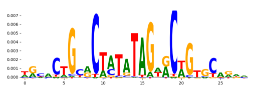 | 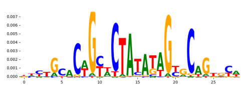 | TN5_3 | 3.450650e-04 | 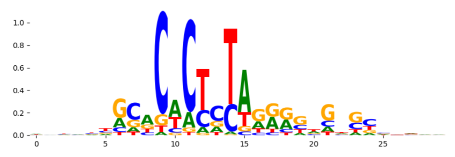 |
| pos_patterns.pattern_8 | 331 | 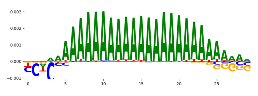 | 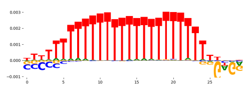 | PRDM6_HUMAN.H11MO.0.C | 7.555640e-02 | 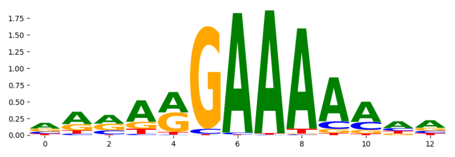 |
| pos_patterns.pattern_9 | 142 | 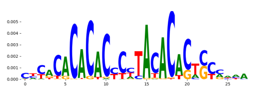 | 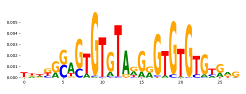 | EGR2_HUMAN.H11MO.0.A | 1.000000e+00 | 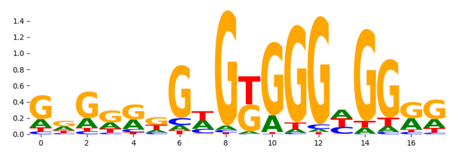 |
| pos_patterns.pattern_10 | 92 | 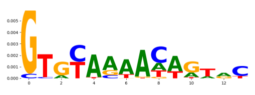 | 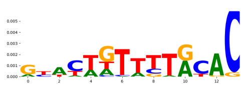 | TBX1_TBX_1 | 5.990980e-01 | 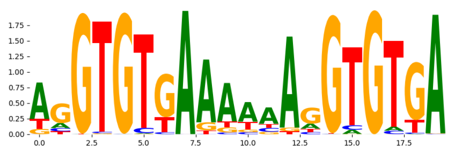 |
| pos_patterns.pattern_11 | 23 | 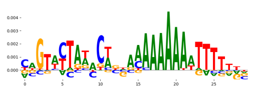 | 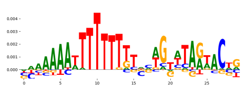 | ZNF384_MA1125.1 | 4.135290e-02 | 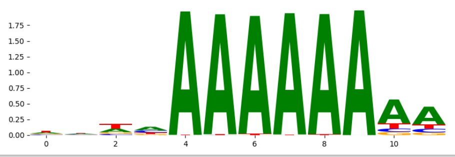 |
