## Supplementary Files 3 for "ChromBPNet: bias factorized, base-resolution deep learning models of chromatin accessibility reveal cis-regulatory sequence syntax, transcription factor footprints and regulatory variants": imr90_ATAC_raw_bpnet_bias_fold4_counts_modisco.pdf

| pattern | num_seqlets | cwm_fwd | cwm_rev | TOMTOM_match | TOMTOM_qval | TOMTOM_match_logo |
|---|---|---|---|---|---|---|
| pos_patterns.pattern_0 | 4588 | | | ONECUT3_CUT_1 | 4.448680e-01 | |
| pos_patterns.pattern_1 | 2414 | | | ZNF384_MA1125.1 | 3.072980e-02 | |
| pos_patterns.pattern_2 | 1550 | | | DNASE_2 | 1.544380e-02 | |
| pos_patterns.pattern_3 | 1404 | | | ZFX_MOUSE.H11MO.0.B | 1.489160e-01 | |
| pos_patterns.pattern_4 | 1279 | | | SP1_HUMAN.H11MO.0.A | 8.089270e-02 | |
| pos_patterns.pattern_5 | 1269 | | | ZFX_MOUSE.H11MO.0.B | 8.266320e-04 | |
| pos_patterns.pattern_6 | 1159 | | | SP2_HUMAN.H11MO.0.A | 2.027550e-03 | |
| pos_patterns.pattern_7 | 992 | | | ZFX_MOUSE.H11MO.0.B | 9.948390e-02 | |
| pos_patterns.pattern_8 | 632 | | | FOXJ3_HUMAN.H11MO.0.A | 1.572650e-01 | |
| pos_patterns.pattern_9 | 573 | | | ZFX_MOUSE.H11MO.0.B | 6.144880e-03 | |
| pos_patterns.pattern_10 | 569 | | | RUNX3_RUNX_3 | 1.301980e-01 | |
| pos_patterns.pattern_11 | 527 | | | TBX1_TBX_4 | 9.865440e-02 | |
| pos_patterns.pattern_12 | 524 | | | SP2_HUMAN.H11MO.0.A | 2.248460e-01 | |
| pos_patterns.pattern_13 | 500 | | | SP2_HUMAN.H11MO.0.A | 6.779970e-04 | |
| pos_patterns.pattern_14 | 467 | | | PRDM6_HUMAN.H11MO.0.C | 4.383070e-01 | |
| pos_patterns.pattern_15 | 431 | | | RUNX2_HUMAN.H11MO.0.A | 4.536790e-01 | |
| pos_patterns.pattern_16 | 424 | | | TBR1_TBX_2 | 3.821530e-01 | |
| pos_patterns.pattern_17 | 364 | | | SIX2_MA1119.1 | 1.000000e+00 | |
| pos_patterns.pattern_18 | 248 | | | TN5_8 | 1.000000e+00 | |
| pos_patterns.pattern_19 | 217 | | | ZSCAN4_C2H2_1 | 5.028780e-01 | |
| pos_patterns.pattern_20 | 184 | | | E2F5_HUMAN.H11MO.0.B | 9.445140e-01 | |
| pos_patterns.pattern_21 | 149 | | | FOXC1_forkhead_1 | 4.307800e-02 | |
| pos_patterns.pattern_22 | 139 | | | TCF7L2_MA0523.1 | 9.547530e-02 | |
| pos_patterns.pattern_23 | 139 | | | LMX1B_MA0703.1 | 2.457080e-02 | |
| pos_patterns.pattern_24 | 136 | | | ZFX_MOUSE.H11MO.0.B | 6.294900e-01 | |
| pos_patterns.pattern_25 | 114 | | | MAFF_MA0495.2 | 3.922880e-01 | |
| pos_patterns.pattern_26 | 74 | | | GLIS2_C2H2_1 | 1.000000e+00 | |
| pos_patterns.pattern_27 | 69 | | | SP2_HUMAN.H11MO.0.A | 2.661840e-04 | |
| pos_patterns.pattern_28 | 50 | | | Nfe2l2_MA0150.2 | 1.000000e+00 | |

| pattern | num_seqlets | cwm_fwd | cwm_rev | TOMTOM_match | TOMTOM_qval | TOMTOM_match_logo |
|---|---|---|---|---|---|---|
| pos_patterns.pattern_29 | 33 | | | TN5_7 | 1.000000e+00 | |
| pos_patterns.pattern_30 | 30 | | | FOXB1_forkhead_2 | 3.978250e-01 | |
| pos_patterns.pattern_31 | 20 | | | SP1_HUMAN.H11MO.0.A | 1.000000e+00 | |
| neg_patterns.pattern_0 | 1216 | | | TN5_8 | 6.350700e-01 | |
| neg_patterns.pattern_1 | 990 | | | TN5_2 | 1.048260e-05 | |
| neg_patterns.pattern_2 | 568 | | | TN5_7 | 7.113750e-04 | |
| neg_patterns.pattern_3 | 410 | | | ZN331_HUMAN.H11MO.0.C | 1.063680e-04 | |
| neg_patterns.pattern_4 | 235 | | | TN5_6 | 1.050940e-23 | |
| neg_patterns.pattern_5 | 196 | | | TAF1_HUMAN.H11MO.0.A | 1.000000e+00 | |
| neg_patterns.pattern_6 | 190 | | | KLF9_HUMAN.H11MO.0.C | 1.618260e-02 | |
| neg_patterns.pattern_7 | 185 | | | ZN770_HUMAN.H11MO.0.C | 5.939930e-13 | |
| neg_patterns.pattern_8 | 183 | | | EGR2_HUMAN.H11MO.0.A | 2.454810e-03 | |
| neg_patterns.pattern_9 | 168 | | | MTF1_C2H2_1 | 3.001770e-01 | |
| neg_patterns.pattern_10 | 106 | | | ZN331_HUMAN.H11MO.0.C | 1.000000e+00 | |
| neg_patterns.pattern_11 | 90 | | | SP2_HUMAN.H11MO.0.A | 1.652660e-04 | |
| neg_patterns.pattern_12 | 82 | | | MAFB_HUMAN.H11MO.0.B | 8.089600e-04 | |
| neg_patterns.pattern_13 | 82 | | | FOSL1_MA0477.1 | 5.134070e-02 | |
| neg_patterns.pattern_14 | 43 | | | SOX9_HMG_2 | 1.000000e+00 | |
| neg_patterns.pattern_15 | 30 | | | THA_HUMAN.H11MO.0.C | 2.503940e-01 | |
