## Supplementary Files 3 for "ChromBPNet: bias factorized, base-resolution deep learning models of chromatin accessibility reveal cis-regulatory sequence syntax, transcription factor footprints and regulatory variants": imr90_ATAC_raw_bpnet_bias_fold4_profile_modisco.pdf

| pattern | num_seqlets | cwm_fwd | cwm_rev | TOMTOM_match | TOMTOM_qval | TOMTOM_match_logo |
|---|---|---|---|---|---|---|
| pos_patterns.pattern_0 | 9822 | 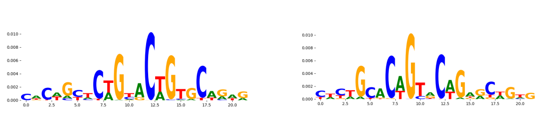 | | TN5_2 | 7.406830e-09 | 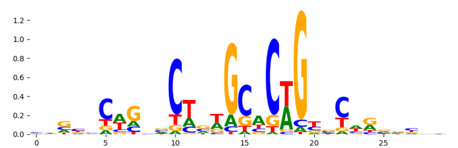 |
| pos_patterns.pattern_1 | 6242 | 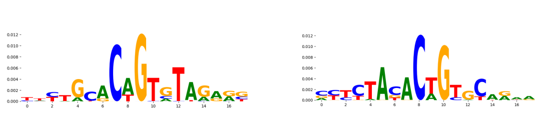 | | TN5_4 | 2.454310e-02 | 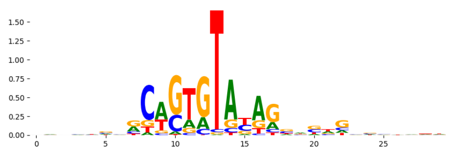 |
| pos_patterns.pattern_2 | 4824 | 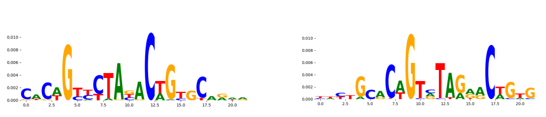 | | TN5_2 | 2.585750e-03 | 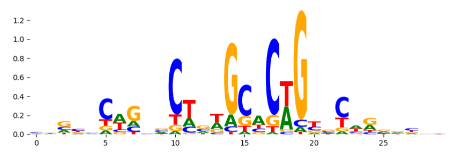 |
| pos_patterns.pattern_3 | 3650 | 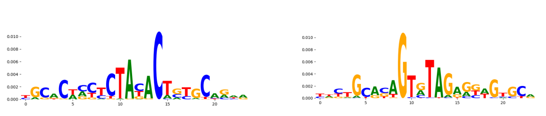 | | TN5_3 | 2.659300e-01 | 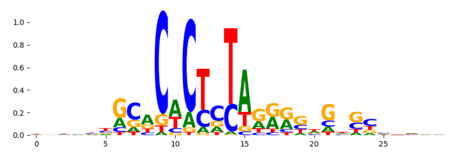 |
| pos_patterns.pattern_4 | 2607 | 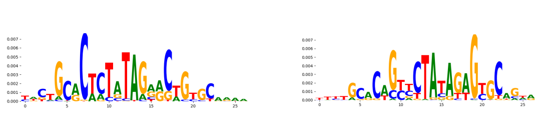 | | TN5_3 | 1.073280e-06 | 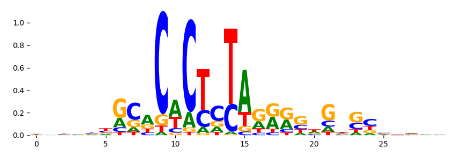 |
| pos_patterns.pattern_5 | 1762 | 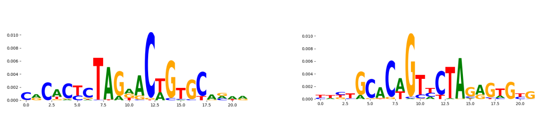 | | TN5_3 | 1.207080e-03 | 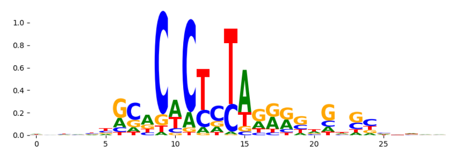 |
| pos_patterns.pattern_6 | 1019 | 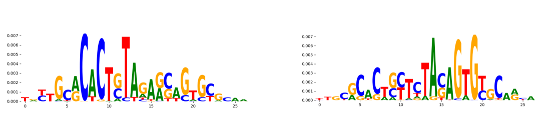 | | TN5_3 | 7.972970e-11 | 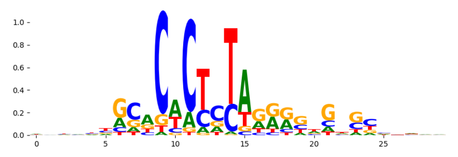 |
| pos_patterns.pattern_7 | 393 | 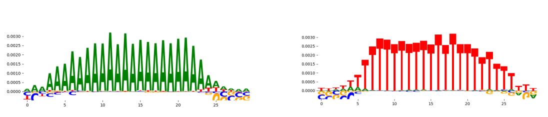 | | PRDM6_HUMAN.H11MO.0.C | 7.185230e-02 | 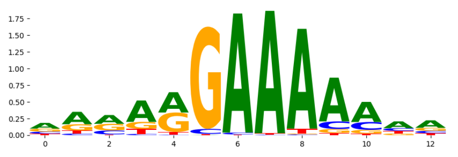 |
| pos_patterns.pattern_8 | 212 | 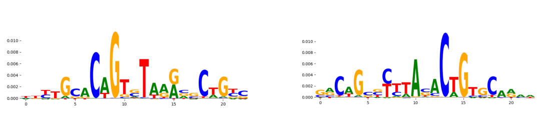 | | FOXA2_HUMAN.H11MO.0.A | 1.000000e+00 | 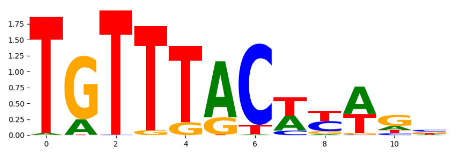 |
| pos_patterns.pattern_9 | 172 | 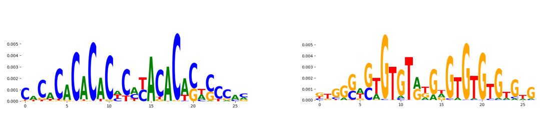 | | RREB1_MA0073.1 | 1.000000e+00 | 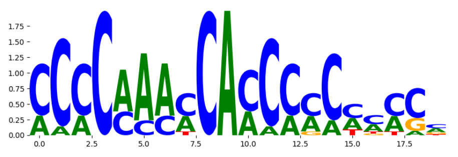 |
| pos_patterns.pattern_10 | 132 | 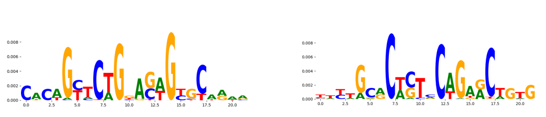 | | TN5_1 | 1.004180e-02 | 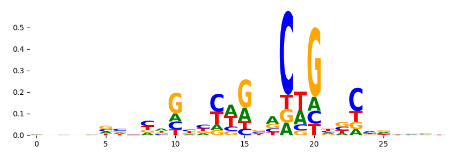 |
| pos_patterns.pattern_11 | 78 | 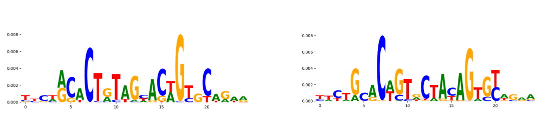 | | TN5_7 | 2.426110e-04 | 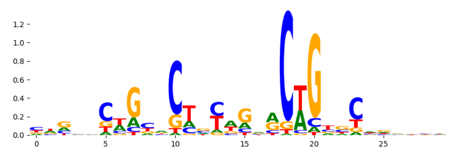 |
| pos_patterns.pattern_12 | 71 | 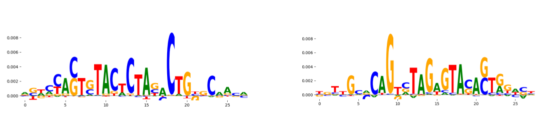 | | TN5_6 | 3.002750e-08 | 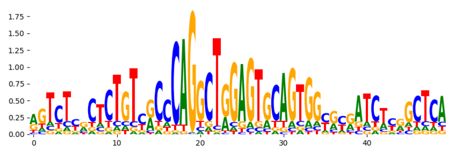 |
| pos_patterns.pattern_13 | 67 | 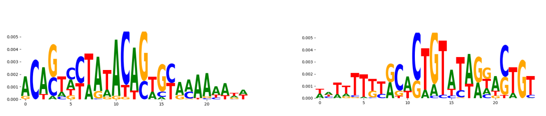 | | TN5_4 | 8.316990e-03 | 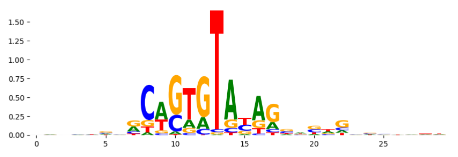 |
| pos_patterns.pattern_14 | 29 | 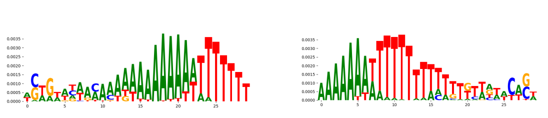 | | ZNF384_MA1125.1 | 2.043820e-02 | 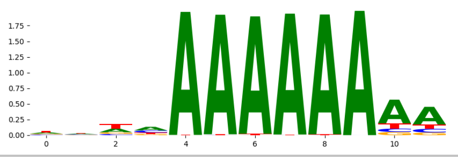 |
