## Supplementary Files 3 for "ChromBPNet: bias factorized, base-resolution deep learning models of chromatin accessibility reveal cis-regulatory sequence syntax, transcription factor footprints and regulatory variants": k562_ATAC_raw_bpnet_bias_fold0_counts_modisco.pdf

| pattern | num_seqlets | cwm_fwd | cwm_rev | TOMTOM_match | TOMTOM_qval | TOMTOM_match_logo |
|---|---|---|---|---|---|---|
| pos_patterns.pattern_0 | 4623 | | | Zfx_MA0146.2 | 7.514190e-02 | |
| pos_patterns.pattern_1 | 4509 | | | SP3_HUMAN.H11MO.0.B | 6.076060e-02 | |
| pos_patterns.pattern_2 | 4452 | | | COE1_HUMAN.H11MO.0.A | 1.612260e-01 | |
| pos_patterns.pattern_3 | 3732 | | | Zfx_MA0146.2 | 7.383070e-02 | |
| pos_patterns.pattern_4 | 1924 | | | PRDM6_HUMAN.H11MO.0.C | 3.509900e-03 | |
| pos_patterns.pattern_5 | 1387 | | | SP1_HUMAN.H11MO.0.A | 6.590080e-03 | |
| pos_patterns.pattern_6 | 1248 | | | NFAC2_HUMAN.H11MO.0.B | 1.463050e-01 | |
| pos_patterns.pattern_7 | 1105 | | | ZFX_MOUSE.H11MO.0.B | 4.500500e-02 | |
| pos_patterns.pattern_8 | 1097 | | | EHF_HUMAN.H11MO.0.B | 5.983360e-01 | |
| pos_patterns.pattern_9 | 981 | | | SP5_MOUSE.H11MO.0.C | 1.013310e-08 | |
| pos_patterns.pattern_10 | 923 | | | HNF6_HUMAN.H11MO.0.B | 1.069850e-01 | |
| pos_patterns.pattern_11 | 895 | | | WT1_HUMAN.H11MO.0.C | 4.402920e-02 | |
| pos_patterns.pattern_12 | 856 | | | PRDM6_HUMAN.H11MO.0.C | 1.328680e-03 | |
| pos_patterns.pattern_13 | 815 | | | PTF1A_HUMAN.H11MO.0.B | 3.291000e-02 | |
| pos_patterns.pattern_14 | 785 | | | SP5_MOUSE.H11MO.0.C | 2.431270e-08 | |
| pos_patterns.pattern_15 | 772 | | | Zfp740.mouse_C2H2_1 | 4.421730e-01 | |
| pos_patterns.pattern_16 | 665 | | | ZNF384_MA1125.1 | 2.182940e-01 | |
| pos_patterns.pattern_17 | 640 | | | NFKB1_HUMAN.H11MO.1.B | 6.221000e-01 | |
| pos_patterns.pattern_18 | 575 | | | COE1_MOUSE.H11MO.0.A | 1.000000e+00 | |
| pos_patterns.pattern_19 | 575 | | | ZN320_HUMAN.H11MO.0.C | 8.931030e-01 | |
| pos_patterns.pattern_20 | 545 | | | Zfx_MA0146.2 | 1.894700e-01 | |
| pos_patterns.pattern_21 | 506 | | | ZN490_HUMAN.H11MO.0.C | 1.000000e+00 | |
| pos_patterns.pattern_22 | 481 | | | GATA1_MOUSE.H11MO.0.A | 8.308370e-01 | |
| pos_patterns.pattern_23 | 466 | | | NaN | NaN | |
| pos_patterns.pattern_24 | 319 | | | NaN | NaN | |
| pos_patterns.pattern_25 | 302 | | | USF2_HUMAN.H11MO.0.A | 3.102670e-01 | |
| pos_patterns.pattern_26 | 256 | | | NR1H3_MOUSE.H11MO.0.A | 9.424250e-01 | |
| pos_patterns.pattern_27 | 201 | | | PAX5_HUMAN.H11MO.0.A | 4.151330e-01 | |
| pos_patterns.pattern_28 | 189 | | | GRHL1_CP2_2 | 1.000000e+00 | |
| pos_patterns.pattern_29 | 185 | | | IRF3_HUMAN.H11MO.0.B | 2.067950e-01 | |

| pattern | num_seqlets | cwm_fwd | cwm_rev | TOMTOM_match | TOMTOM_qval | TOMTOM_match_logo |
|---|---|---|---|---|---|---|
| pos_patterns.pattern_30 | 180 | | | MBD2_HUMAN.H11MO.0.B | 1.000000e+00 | |
| pos_patterns.pattern_31 | 162 | | | MAZ_HUMAN.H11MO.0.A | 1.468100e-01 | |
| pos_patterns.pattern_32 | 121 | | | ZN274_HUMAN.H11MO.0.A | 1.000000e+00 | |
| pos_patterns.pattern_33 | 126 | | | HIC1_HUMAN.H11MO.0.C | 5.295130e-02 | |
| pos_patterns.pattern_34 | 103 | | | TN5_4 | 7.348250e-01 | |
| pos_patterns.pattern_35 | 100 | | | ZFP57_MOUSE.H11MO.0.B | 1.000000e+00 | |
| pos_patterns.pattern_36 | 67 | | | SP1_HUMAN.H11MO.0.A | 1.000000e+00 | |
| pos_patterns.pattern_37 | 60 | | | NR1H4_HUMAN.H11MO.0.B | 4.044930e-02 | |
| pos_patterns.pattern_38 | 61 | | | ZN121_HUMAN.H11MO.0.C | 1.326440e-02 | |
| pos_patterns.pattern_39 | 63 | | | TP53_MA0106.3 | 5.534980e-01 | |
| pos_patterns.pattern_40 | 51 | | | TGIF2_MA0797.1 | 4.873440e-02 | |
| pos_patterns.pattern_41 | 49 | | | SP1_HUMAN.H11MO.0.A | 1.341510e-01 | |
| pos_patterns.pattern_42 | 44 | | | SMAD2_MOUSE.H11MO.0.A | 1.000000e+00 | |
| pos_patterns.pattern_43 | 46 | | | SP5_MOUSE.H11MO.0.C | 8.803520e-03 | |
| pos_patterns.pattern_44 | 46 | | | ZSCAN4_C2H2_1 | 9.999990e-01 | |
| pos_patterns.pattern_45 | 33 | | | VDR_HUMAN.H11MO.0.A | 2.861960e-01 | |
| pos_patterns.pattern_46 | 38 | | | ESR2_HUMAN.H11MO.0.A | 2.129400e-01 | |
| pos_patterns.pattern_47 | 35 | | | ZNF41_HUMAN.H11MO.0.C | 1.000000e+00 | |
| neg_patterns.pattern_0 | 827 | | | SP3_HUMAN.H11MO.0.B | 1.602600e-08 | |
| neg_patterns.pattern_1 | 616 | | | TN5_2 | 6.234200e-04 | |
| neg_patterns.pattern_2 | 368 | | | ZNF384_MA1125.1 | 9.593720e-04 | |
| neg_patterns.pattern_3 | 365 | | | TN5_1 | 1.743260e-03 | |
| neg_patterns.pattern_4 | 312 | | | TN5_7 | 2.519440e-02 | |
| neg_patterns.pattern_5 | 312 | | | SMAD4_HUMAN.H11MO.0.B | 6.954020e-02 | |
| neg_patterns.pattern_6 | 274 | | | NR1D2_MOUSE.H11MO.0.A | 8.128240e-01 | |
| neg_patterns.pattern_7 | 277 | | | Myog_MA0500.1 | 6.288240e-01 | |
| neg_patterns.pattern_8 | 198 | | | SP1_MOUSE.H11MO.0.A | 5.516340e-04 | |
| neg_patterns.pattern_9 | 210 | | | SP3_HUMAN.H11MO.0.B | 2.494550e-03 | |
| neg_patterns.pattern_10 | 211 | | | ELF2_MOUSE.H11MO.0.C | 2.268930e-01 | |
| neg_patterns.pattern_11 | 199 | | | USF2_HUMAN.H11MO.0.A | 1.629330e-01 | |

| pattern | num_seqlets | cwm_fwd | cwm_rev | TOMTOM_match | TOMTOM_qval | TOMTOM_match_logo |
|---|---|---|---|---|---|---|
| neg_patterns.pattern_12 | 138 | 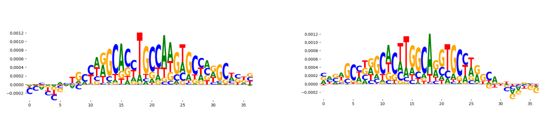 | | HES5_MA0821.1 | 3.995470e-03 | 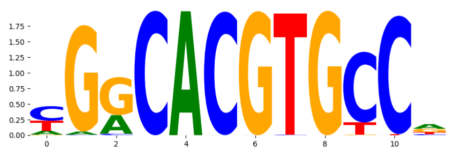 |
| neg_patterns.pattern_13 | 110 | 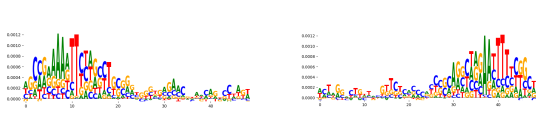 | | SP2_HUMAN.H11MO.0.A | 1.638920e-05 | 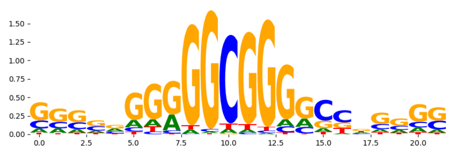 |
| neg_patterns.pattern_14 | 96 | 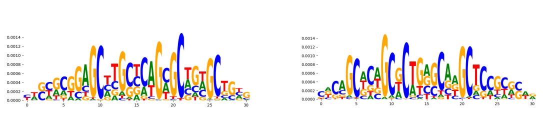 | | THAP1_HUMAN.H11MO.0.C | 1.000000e+00 | 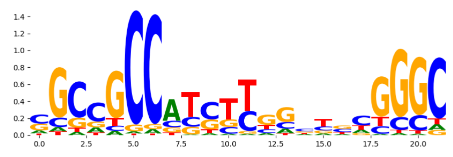 |
| neg_patterns.pattern_15 | 64 | 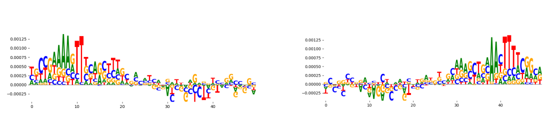 | | SP1_HUMAN.H11MO.0.A | 5.133080e-03 | 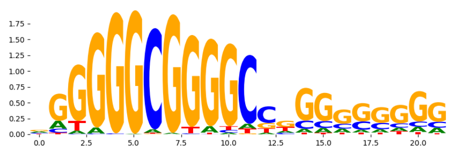 |
| neg_patterns.pattern_16 | 71 | 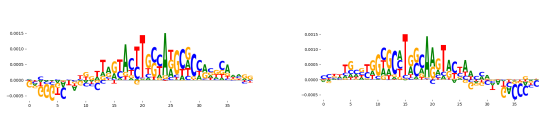 | | NRF1_MOUSE.H11MO.0.A | 4.433670e-02 | 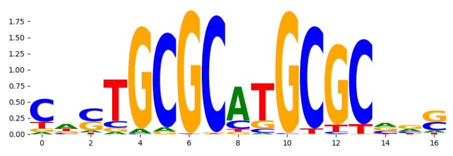 |
| neg_patterns.pattern_17 | 49 | 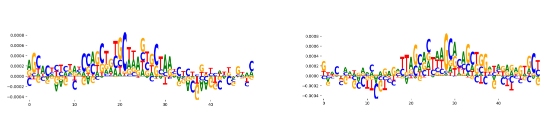 | | SP2_HUMAN.H11MO.0.A | 5.241080e-03 | 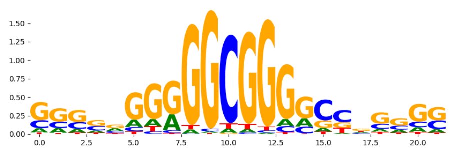 |
| neg_patterns.pattern_18 | 78 | 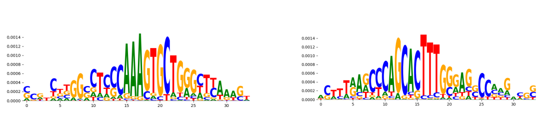 | | IKZF1_HUMAN.H11MO.0.C | 1.000000e+00 | 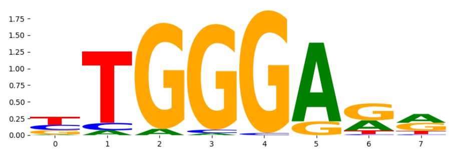 |
| neg_patterns.pattern_19 | 36 | 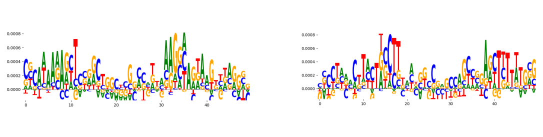 | | ZFX_MOUSE.H11MO.0.B | 4.960440e-03 | 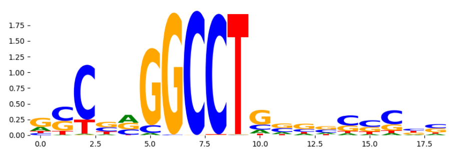 |
| neg_patterns.pattern_20 | 36 | 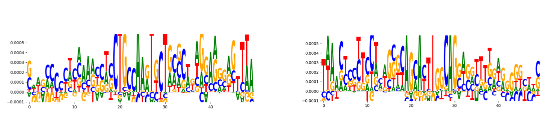 | | NFIB_MOUSE.H11MO.0.C | 8.919150e-03 | 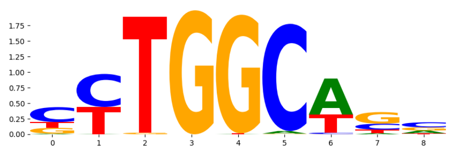 |
| neg_patterns.pattern_21 | 35 | 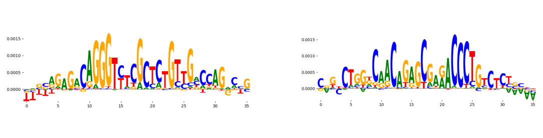 | | ZN121_HUMAN.H11MO.0.C | 1.341930e-08 | 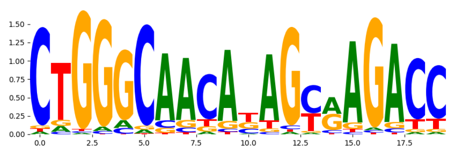 |
| neg_patterns.pattern_22 | 29 | 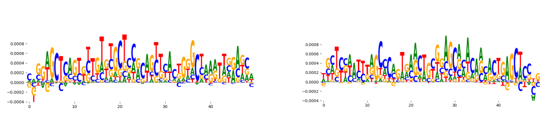 | | TN5_2 | 1.257090e-01 | 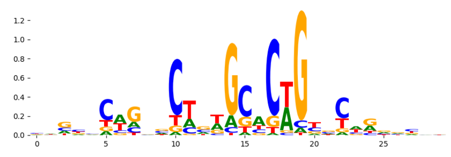 |
| neg_patterns.pattern_23 | 30 | 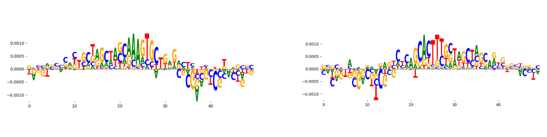 | | TN5_2 | 4.920810e-02 | 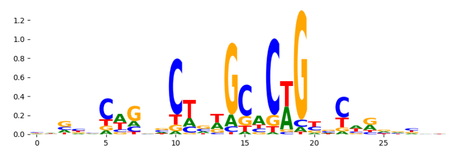 |
| neg_patterns.pattern_24 | 39 | 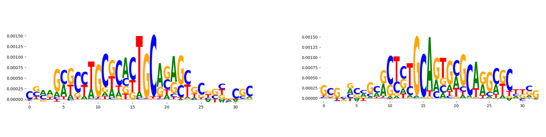 | | NRF1_MA0506.1 | 2.476270e-01 | 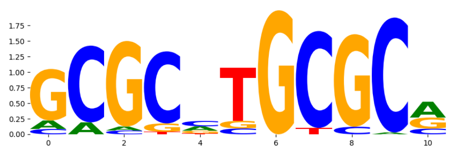 |
| neg_patterns.pattern_25 | 30 | 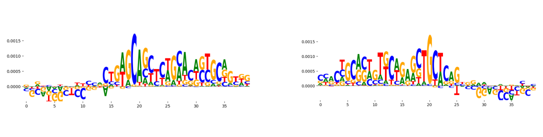 | | TN5_7 | 2.002190e-01 | 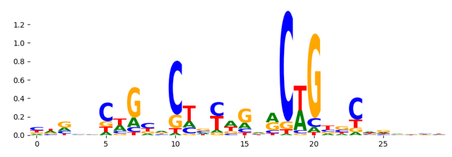 |
| neg_patterns.pattern_26 | 27 | 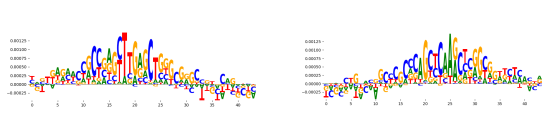 | | KLF3_HUMAN.H11MO.0.B | 2.150620e-01 | 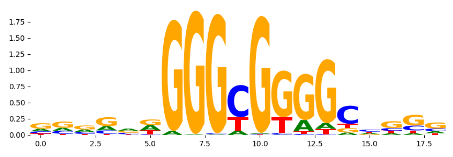 |
| neg_patterns.pattern_27 | 27 | 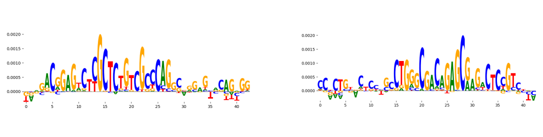 | | TN5_6 | 3.212420e-12 | 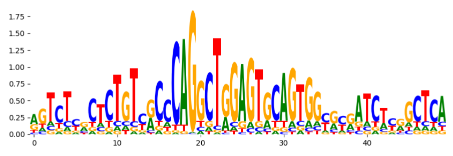 |
| neg_patterns.pattern_28 | 26 | 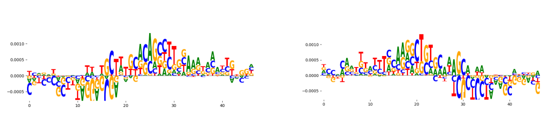 | | ZFX_MOUSE.H11MO.0.B | 3.639610e-02 | 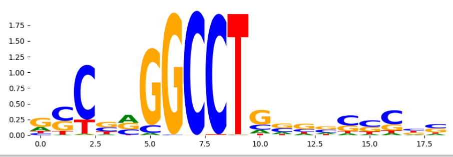 |
