## Supplementary Files 3 for "ChromBPNet: bias factorized, base-resolution deep learning models of chromatin accessibility reveal cis-regulatory sequence syntax, transcription factor footprints and regulatory variants": k562_ATAC_raw_bpnet_bias_fold0_profile_modisco.pdf

| pattern | num_seqlets | cwm_fwd | cwm_rev | TOMTOM_match | TOMTOM_qval | TOMTOM_match_logo |
|---|---|---|---|---|---|---|
| pos_patterns.pattern_0 | 16897 | | | TN5_1 | 3.311610e-26 | |
| pos_patterns.pattern_1 | 4237 | | | TN5_2 | 0.000000e+00 | |
| pos_patterns.pattern_2 | 3666 | | | TN5_1 | 1.592680e-06 | |
| pos_patterns.pattern_3 | 3334 | | | TN5_3 | 1.435220e-29 | |
| pos_patterns.pattern_4 | 3274 | | | TN5_4 | 3.465330e-31 | |
| pos_patterns.pattern_5 | 1997 | | | TN5_6 | 2.943660e-23 | |
| pos_patterns.pattern_6 | 1624 | | | TN5_7 | 3.107550e-35 | |
| pos_patterns.pattern_7 | 1053 | | | TN5_8 | 2.458440e-41 | |
| pos_patterns.pattern_8 | 925 | | | TN5_8 | 2.250910e-06 | |
| pos_patterns.pattern_9 | 551 | | | TN5_4 | 1.000000e+00 | |
| pos_patterns.pattern_10 | 470 | | | PRDM6_HUMAN.H11MO.0.C | 1.122440e-01 | |
| pos_patterns.pattern_11 | 233 | | | TN5_6 | 9.400980e-06 | |
| pos_patterns.pattern_12 | 187 | | | ZNF384_MA1125.1 | 1.718190e-02 | |
| pos_patterns.pattern_13 | 172 | | | ZNF384_MA1125.1 | 3.074120e-02 | |
| pos_patterns.pattern_14 | 89 | | | RREB1_MA0073.1 | 1.000000e+00 | |
| pos_patterns.pattern_15 | 82 | | | EGR2_HUMAN.H11MO.0.A | 1.000000e+00 | |
| pos_patterns.pattern_16 | 34 | | | TEAD2_MOUSE.H11MO.0.C | 9.812700e-01 | |
| pos_patterns.pattern_17 | 32 | | | TN5_7 | 2.705120e-01 | |
| pos_patterns.pattern_18 | 29 | | | NaN | NaN | |
| pos_patterns.pattern_19 | 22 | | | TN5_3 | 1.000000e+00 | |
