## Supplementary Files 3 for "ChromBPNet: bias factorized, base-resolution deep learning models of chromatin accessibility reveal cis-regulatory sequence syntax, transcription factor footprints and regulatory variants": k562_ATAC_raw_bpnet_bias_fold1_counts_modisco.pdf

| pattern | num_seqlets | cwm_fwd | cwm_rev | TOMTOM_match | TOMTOM_qval | TOMTOM_match_logo |
|---|---|---|---|---|---|---|
| pos_patterns.pattern_0 | 4940 | | | KLF12_HUMAN.H11MO.0.C | 1.784760e-01 | |
| pos_patterns.pattern_1 | 3022 | | | SP2_HUMAN.H11MO.0.A | 9.521130e-03 | |
| pos_patterns.pattern_2 | 1786 | | | ZBT7A_HUMAN.H11MO.0.A | 1.350880e-01 | |
| pos_patterns.pattern_3 | 954 | | | RREB1_MA0073.1 | 1.000000e+00 | |
| pos_patterns.pattern_4 | 773 | | | TN5_2 | 1.978000e-01 | |
| pos_patterns.pattern_5 | 764 | | | SOX10_HMG_5 | 1.000000e+00 | |
| pos_patterns.pattern_6 | 539 | | | TN5_1 | 1.000000e+00 | |
| pos_patterns.pattern_7 | 531 | | | ANDR_HUMAN.H11MO.0.A | 2.642490e-01 | |
| pos_patterns.pattern_8 | 531 | | | HIC1_HUMAN.H11MO.0.C | 1.899190e-01 | |
| pos_patterns.pattern_9 | 518 | | | ZFP57_MOUSE.H11MO.0.B | 1.000000e+00 | |
| pos_patterns.pattern_10 | 496 | | | ETS1_HUMAN.H11MO.0.A | 7.790330e-01 | |
| pos_patterns.pattern_11 | 464 | | | SP2_HUMAN.H11MO.0.A | 5.011730e-01 | |
| pos_patterns.pattern_12 | 454 | | | ARI5B_HUMAN.H11MO.0.C | 1.000000e+00 | |
| pos_patterns.pattern_13 | 306 | | | RREB1_MA0073.1 | 2.799840e-05 | |
| pos_patterns.pattern_14 | 271 | | | ESR2_HUMAN.H11MO.0.A | 2.373130e-02 | |
| pos_patterns.pattern_15 | 236 | | | ZNF740_C2H2_2 | 4.588520e-02 | |
| pos_patterns.pattern_16 | 232 | | | OSR2_HUMAN.H11MO.0.C | 3.293840e-03 | |
| pos_patterns.pattern_17 | 214 | | | SP2_HUMAN.H11MO.0.A | 1.313400e-04 | |
| pos_patterns.pattern_18 | 208 | | | ANDR_HUMAN.H11MO.0.A | 8.861060e-01 | |
| pos_patterns.pattern_19 | 165 | | | PATZ1_HUMAN.H11MO.0.C | 1.635340e-03 | |
| pos_patterns.pattern_20 | 160 | | | ZN467_HUMAN.H11MO.0.C | 1.705560e-04 | |
| pos_patterns.pattern_21 | 145 | | | SP1_HUMAN.H11MO.0.A | 3.274330e-07 | |
| pos_patterns.pattern_22 | 117 | | | SP1_MOUSE.H11MO.0.A | 7.407780e-07 | |
| pos_patterns.pattern_23 | 74 | | | P63_HUMAN.H11MO.0.A | 5.037470e-01 | |
| pos_patterns.pattern_24 | 67 | | | SP5_MOUSE.H11MO.0.C | 2.436330e-02 | |
| pos_patterns.pattern_25 | 36 | | | TP63_MA0525.2 | 4.339640e-01 | |
| neg_patterns.pattern_0 | 139 | | | SP2_HUMAN.H11MO.0.A | 8.064840e-07 | |
| neg_patterns.pattern_1 | 135 | | | SP1_HUMAN.H11MO.0.A | 5.436040e-02 | |
| neg_patterns.pattern_2 | 118 | | | ZN331_HUMAN.H11MO.0.C | 1.831570e-03 | |

| pattern | num_seqlets | cwm_fwd | cwm_rev | TOMTOM_match | TOMTOM_qval | TOMTOM_match_logo |
|---|---|---|---|---|---|---|
| neg_patterns.pattern_3 | 115 | 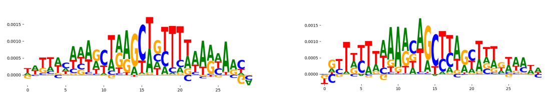 | | ZEB1_MA0103.3 | 1.000000e+00 | 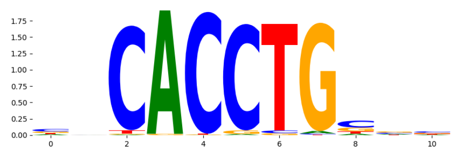 |
| neg_patterns.pattern_4 | 108 | 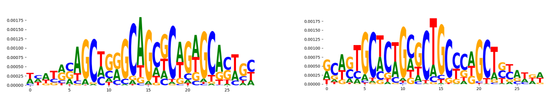 | | SP1_MOUSE.H11MO.0.A | 2.597490e-02 | 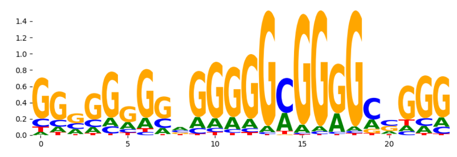 |
| neg_patterns.pattern_5 | 108 | 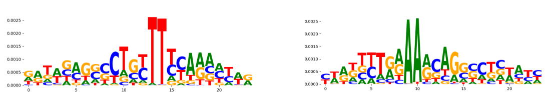 | | RFX6_MOUSE.H11MO.0.C | 1.000000e+00 | 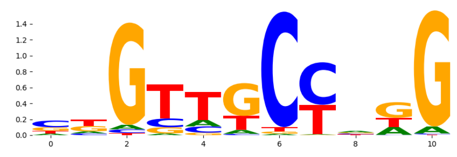 |
| neg_patterns.pattern_6 | 100 | 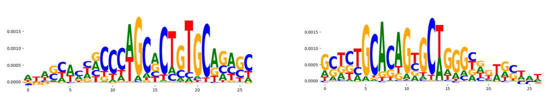 | | ZFX_MOUSE.H11MO.0.B | 1.165950e-02 | 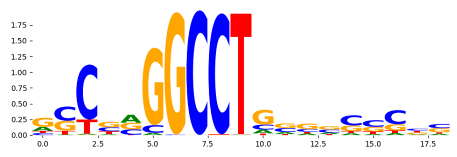 |
| neg_patterns.pattern_7 | 97 | 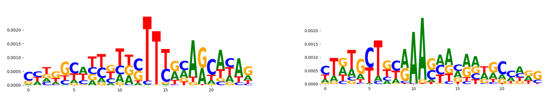 | | RFX1_HUMAN.H11MO.0.B | 8.414950e-02 | 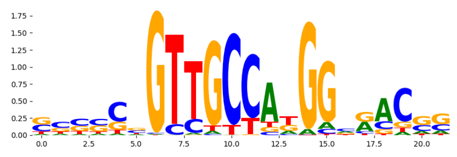 |
| neg_patterns.pattern_8 | 88 | 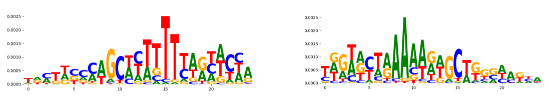 | | SP3_HUMAN.H11MO.0.B | 4.724440e-01 | 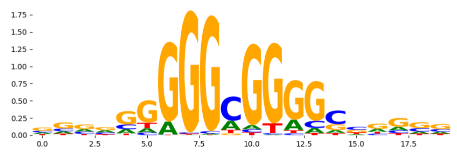 |
| neg_patterns.pattern_9 | 67 | 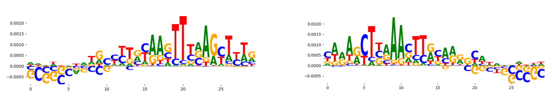 | | Zfx_MA0146.2 | 7.526160e-03 | 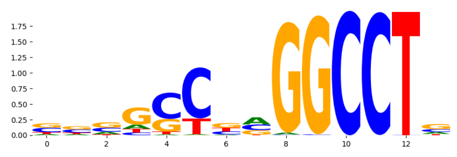 |
| neg_patterns.pattern_10 | 65 | 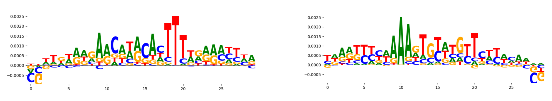 | | SP1_MOUSE.H11MO.0.A | 4.318450e-02 | 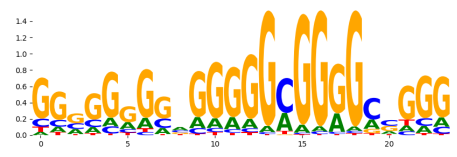 |
| neg_patterns.pattern_11 | 49 | 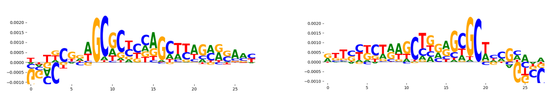 | | SP2_HUMAN.H11MO.0.A | 3.653230e-03 | 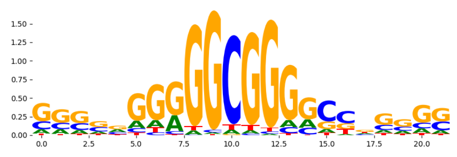 |
| neg_patterns.pattern_12 | 45 | 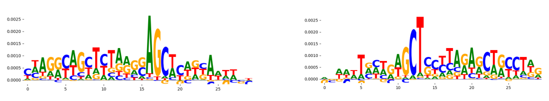 | | TN5_7 | 3.167920e-01 | 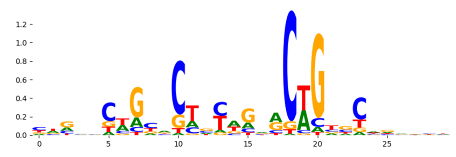 |
| neg_patterns.pattern_13 | 43 | 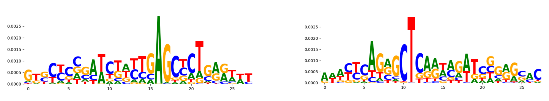 | | SP1_HUMAN.H11MO.0.A | 1.831900e-01 | 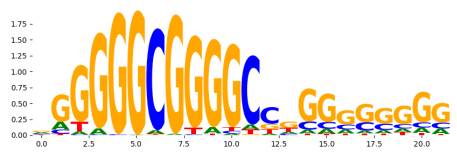 |
| neg_patterns.pattern_14 | 42 | 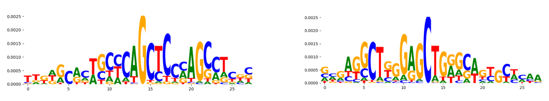 | | SP1_HUMAN.H11MO.0.A | 3.663470e-04 | 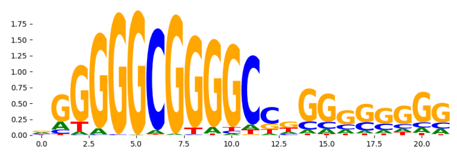 |
| neg_patterns.pattern_15 | 38 | 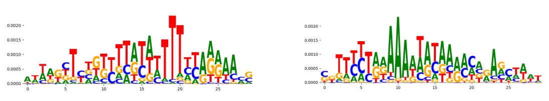 | | BCL6_HUMAN.H11MO.0.A | 1.759410e-01 | 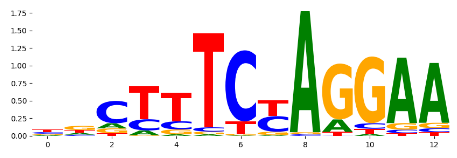 |
| neg_patterns.pattern_16 | 38 | 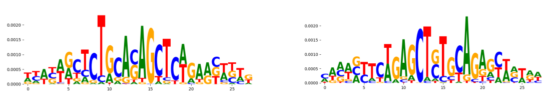 | | TN5_2 | 3.021440e-02 | 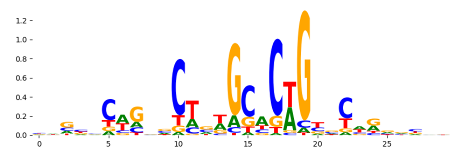 |
| neg_patterns.pattern_17 | 35 | 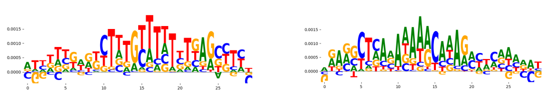 | | Zic3.mouse_C2H2_1 | 5.698560e-01 | 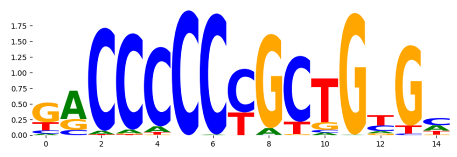 |
| neg_patterns.pattern_18 | 29 | 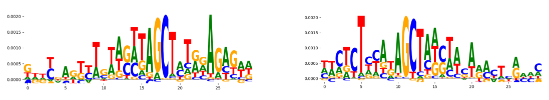 | | ZFX_MOUSE.H11MO.0.B | 2.038440e-01 | 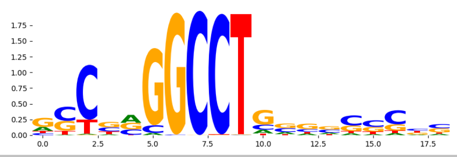 |
