## Supplementary Files 3 for "ChromBPNet: bias factorized, base-resolution deep learning models of chromatin accessibility reveal cis-regulatory sequence syntax, transcription factor footprints and regulatory variants": k562_ATAC_raw_bpnet_bias_fold1_profile_modisco.pdf

| pattern | num_seqlets | cwm_fwd | cwm_rev | TOMTOM_match | TOMTOM_qval | TOMTOM_match_logo |
|---|---|---|---|---|---|---|
| pos_patterns.pattern_0 | 13018 | 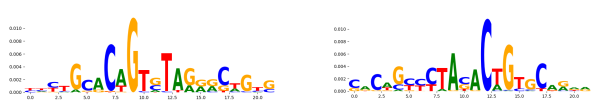 | | TN5_2 | 8.201700e-04 | 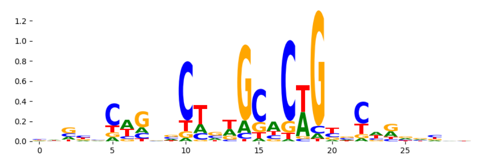 |
| pos_patterns.pattern_1 | 7535 | 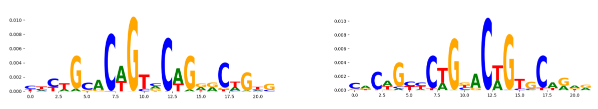 | | TN5_1 | 6.700460e-10 | 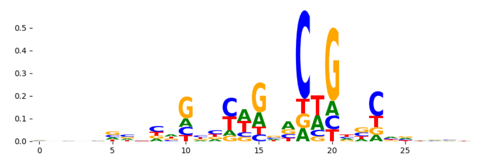 |
| pos_patterns.pattern_2 | 2572 | 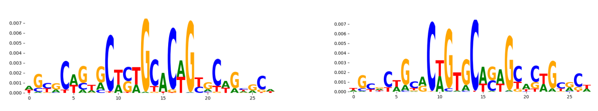 | | TN5_2 | 1.089560e-20 | 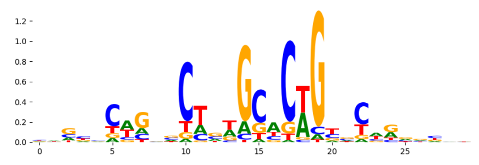 |
| pos_patterns.pattern_3 | 2294 | 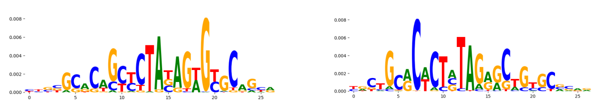 | | TN5_3 | 5.824520e-09 | 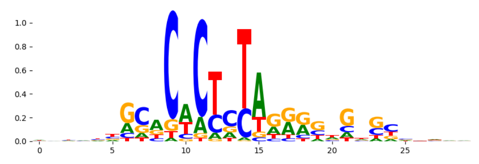 |
| pos_patterns.pattern_4 | 1032 | 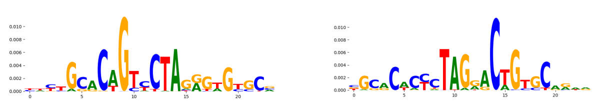 | | TN5_3 | 4.443830e-04 | 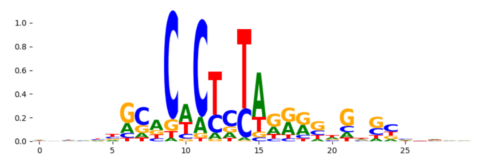 |
| pos_patterns.pattern_5 | 1029 | 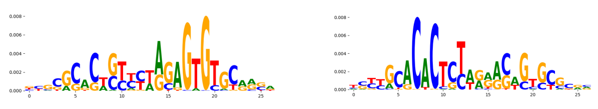 | | TN5_3 | 5.368410e-11 | 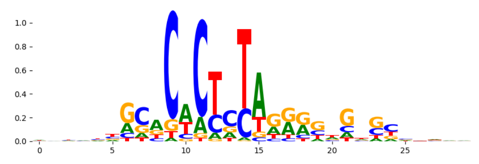 |
| pos_patterns.pattern_6 | 742 | 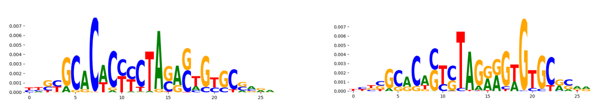 | | TN5_3 | 7.226360e-14 | 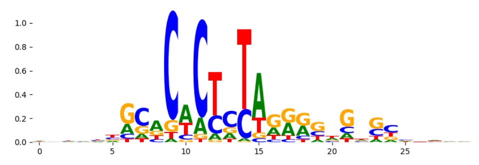 |
| pos_patterns.pattern_7 | 695 | 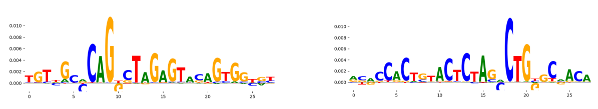 | | TN5_6 | 1.512940e-17 | 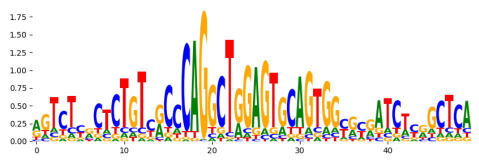 |
| pos_patterns.pattern_8 | 374 | 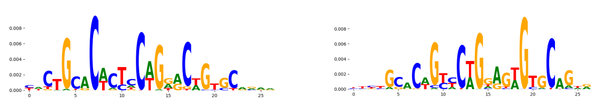 | | TN5_1 | 3.391360e-08 | 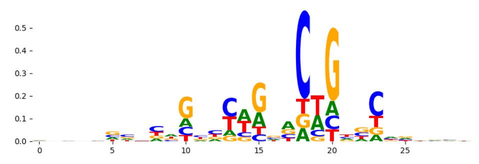 |
| pos_patterns.pattern_9 | 371 | 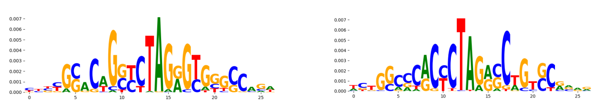 | | TN5_3 | 6.608430e-04 | 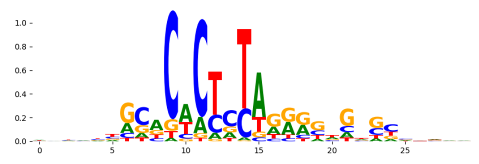 |
| pos_patterns.pattern_10 | 104 | 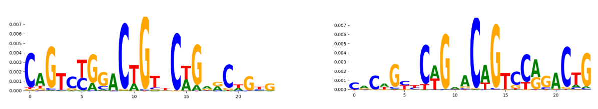 | | TN5_8 | 1.819300e-06 | 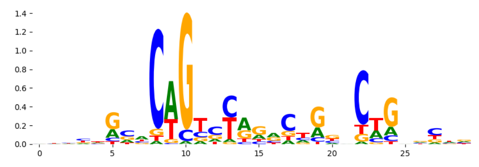 |
| pos_patterns.pattern_11 | 100 | 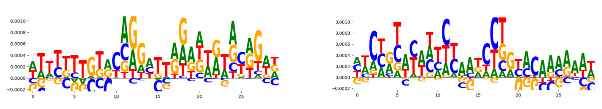 | | TN5_6 | 1.213600e-02 | 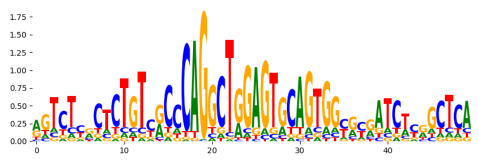 |
| pos_patterns.pattern_12 | 39 | 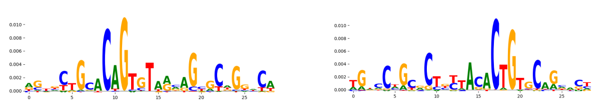 | | TN5_7 | 7.886330e-04 | 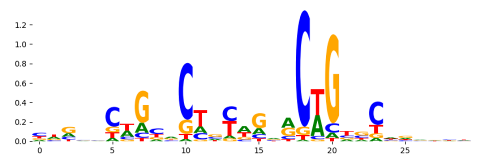 |
| pos_patterns.pattern_13 | 20 | 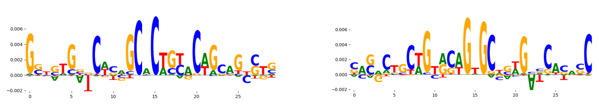 | | TN5_6 | 4.468860e-05 | 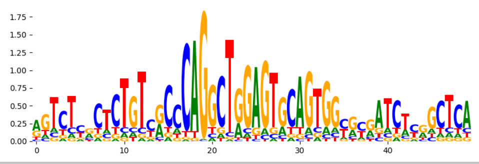 |
