## Supplementary Files 3 for "ChromBPNet: bias factorized, base-resolution deep learning models of chromatin accessibility reveal cis-regulatory sequence syntax, transcription factor footprints and regulatory variants": k562_ATAC_raw_bpnet_bias_fold2_counts_modisco.pdf

| pattern | num_seqlets | cwm_fwd | cwm_rev | TOMTOM_match | TOMTOM_qval | TOMTOM_match_logo |
|---|---|---|---|---|---|---|
| pos_patterns.pattern_0 | 7909 | | | SP2_HUMAN.H11MO.0.A | 1.000000 | |
| pos_patterns.pattern_1 | 4005 | | | ZFX_MOUSE.H11MO.0.B | 0.040608 | |
| pos_patterns.pattern_2 | 827 | | | ZNF384_MA1125.1 | 0.006267 | |
| pos_patterns.pattern_3 | 785 | | | NFATC1_NFAT_1 | 1.000000 | |
| pos_patterns.pattern_4 | 733 | | | ZNF524_C2H2_1 | 1.000000 | |
| pos_patterns.pattern_5 | 681 | | | NFAC3_HUMAN.H11MO.0.B | 0.096131 | |
| pos_patterns.pattern_6 | 550 | | | ZN214_HUMAN.H11MO.0.C | 1.000000 | |
| pos_patterns.pattern_7 | 525 | | | SP3_HUMAN.H11MO.0.B | 0.000057 | |
| pos_patterns.pattern_8 | 502 | | | ZBT7A_HUMAN.H11MO.0.A | 0.439502 | |
| pos_patterns.pattern_9 | 484 | | | PBX1_MA0070.1 | 1.000000 | |
| pos_patterns.pattern_10 | 458 | | | PATZ1_HUMAN.H11MO.0.C | 0.000444 | |
| pos_patterns.pattern_11 | 433 | | | SP1_HUMAN.H11MO.0.A | 0.029421 | |
| pos_patterns.pattern_12 | 283 | | | ESR1_HUMAN.H11MO.0.A | 1.000000 | |
| pos_patterns.pattern_13 | 226 | | | EGR2_HUMAN.H11MO.0.A | 0.001542 | |
| pos_patterns.pattern_14 | 199 | | | SP2_HUMAN.H11MO.0.A | 0.419817 | |
| pos_patterns.pattern_15 | 198 | | | AP2C_MOUSE.H11MO.0.A | 0.190258 | |
| pos_patterns.pattern_16 | 183 | | | ZN281_MOUSE.H11MO.0.A | 0.002343 | |
| pos_patterns.pattern_17 | 174 | | | TCF7L2_MA0523.1 | 0.410400 | |
| pos_patterns.pattern_18 | 124 | | | ARI5B_HUMAN.H11MO.0.C | 1.000000 | |
| pos_patterns.pattern_19 | 22 | | | ZN320_HUMAN.H11MO.0.C | 0.415961 | |
| neg_patterns.pattern_0 | 272 | | | TN5_2 | 0.300609 | |
| neg_patterns.pattern_1 | 160 | | | NHLH1_bHLH_1 | 0.006659 | |
| neg_patterns.pattern_2 | 141 | | | ZFX_MOUSE.H11MO.0.B | 0.039905 | |
| neg_patterns.pattern_3 | 139 | | | THAP1_HUMAN.H11MO.0.C | 0.000705 | |
| neg_patterns.pattern_4 | 124 | | | KLF6_HUMAN.H11MO.0.A | 0.444607 | |
| neg_patterns.pattern_5 | 112 | | | ZN331_HUMAN.H11MO.0.C | 0.100857 | |
| neg_patterns.pattern_6 | 96 | | | THAP1_HUMAN.H11MO.0.C | 0.089762 | |
| neg_patterns.pattern_7 | 94 | | | TN5_2 | 0.215890 | |
| neg_patterns.pattern_8 | 94 | | | SP2_HUMAN.H11MO.0.A | 0.003688 | |

| pattern | num_seqlets | cwm_fwd | cwm_rev | TOMTOM_match | TOMTOM_qval | TOMTOM_match_logo |
|---|---|---|---|---|---|---|
| neg_patterns.pattern_9 | 85 | 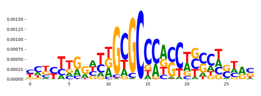 | 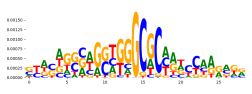 | SP2_HUMAN.H11MO.0.A | 0.028890 | 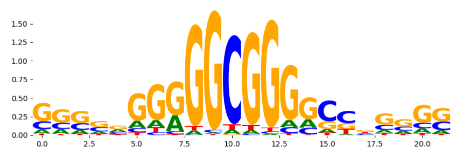 |
| neg_patterns.pattern_10 | 83 | 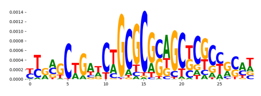 | 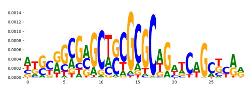 | SP2_HUMAN.H11MO.0.A | 0.001120 | 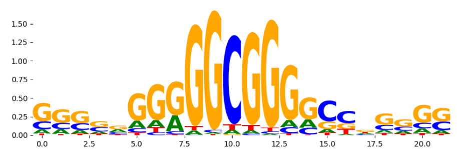 |
| neg_patterns.pattern_11 | 81 | 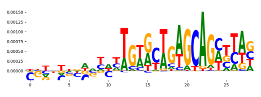 | 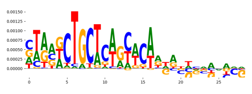 | ZN331_HUMAN.H11MO.0.C | 0.688227 | 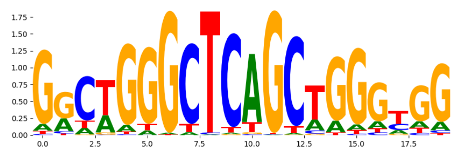 |
| neg_patterns.pattern_12 | 80 | 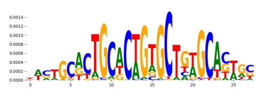 | 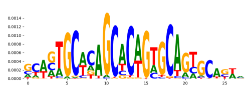 | TN5_2 | 0.004024 | 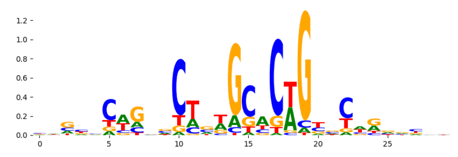 |
| neg_patterns.pattern_13 | 70 | 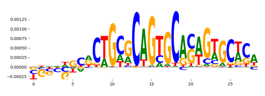 | 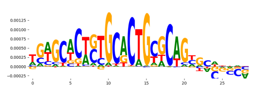 | TN5_2 | 0.001492 | 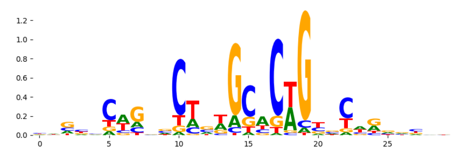 |
| neg_patterns.pattern_14 | 70 | 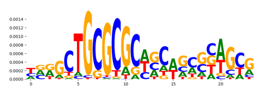 | 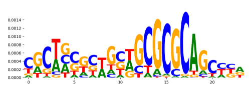 | ZBT14_HUMAN.H11MO.0.C | 0.274724 | 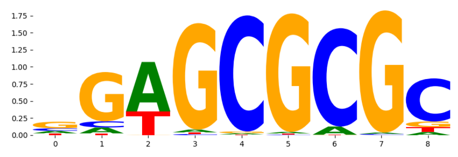 |
| neg_patterns.pattern_15 | 68 | 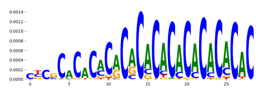 | 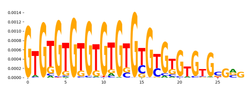 | RREB1_MA0073.1 | 1.000000 | 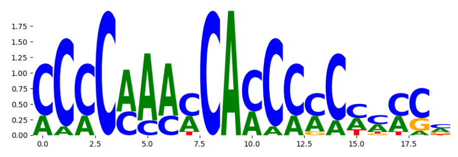 |
| neg_patterns.pattern_16 | 66 | 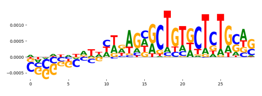 | 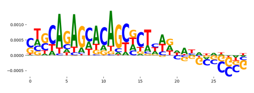 | TN5_2 | 0.007280 | 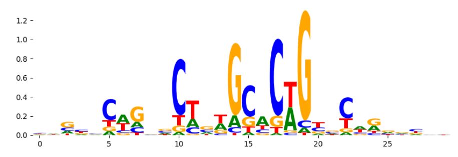 |
| neg_patterns.pattern_17 | 64 | 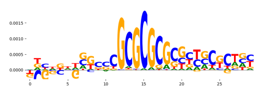 | 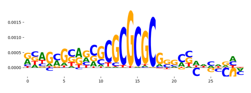 | SP2_HUMAN.H11MO.0.A | 0.000072 | 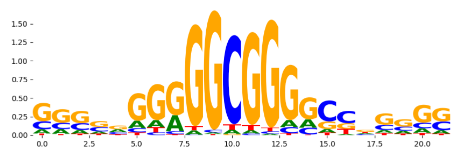 |
| neg_patterns.pattern_18 | 61 | 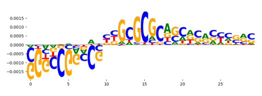 | 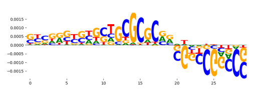 | SP2_HUMAN.H11MO.0.A | 0.000105 | 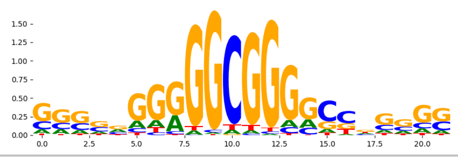 |
