## Supplementary Files 3 for "ChromBPNet: bias factorized, base-resolution deep learning models of chromatin accessibility reveal cis-regulatory sequence syntax, transcription factor footprints and regulatory variants": k562_ATAC_raw_bpnet_bias_fold2_profile_modisco.pdf

| pattern | num_seqlets | cwm_fwd | cwm_rev | TOMTOM_match | TOMTOM_qval | TOMTOM_match_logo |
|---|---|---|---|---|---|---|
| pos_patterns.pattern_0 | 8012 | 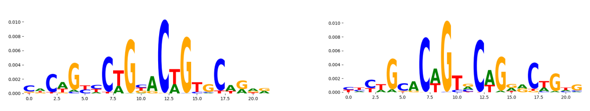 | | TN5_1 | 9.469040e-09 | 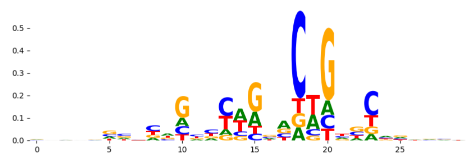 |
| pos_patterns.pattern_1 | 5392 | 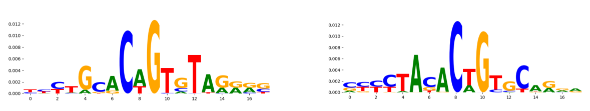 | | TN5_4 | 2.114050e-02 | 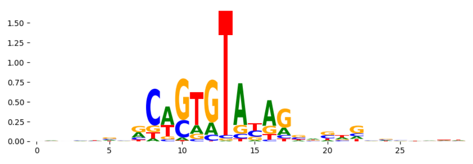 |
| pos_patterns.pattern_2 | 4319 | 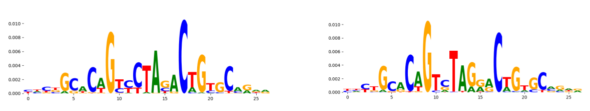 | | TN5_1 | 7.896670e-08 | 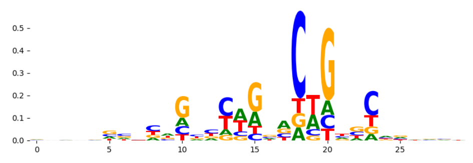 |
| pos_patterns.pattern_3 | 2950 | 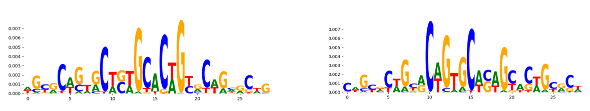 | | TN5_2 | 1.235040e-30 | 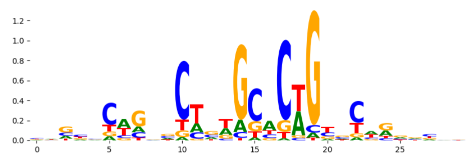 |
| pos_patterns.pattern_4 | 2641 | 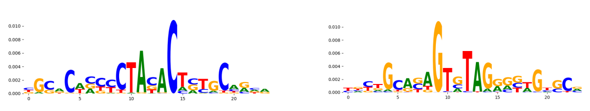 | | TN5_3 | 1.274990e-01 | 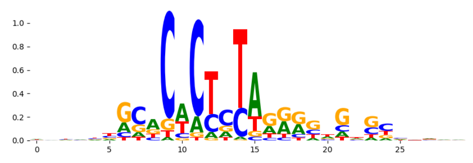 |
| pos_patterns.pattern_5 | 994 | 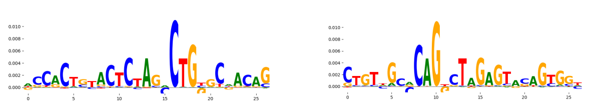 | | TN5_6 | 4.125130e-19 | 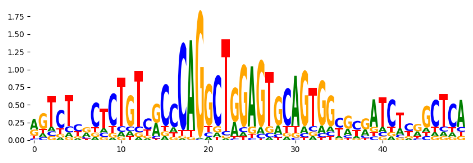 |
| pos_patterns.pattern_6 | 938 | 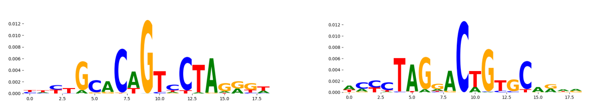 | | TN5_1 | 4.500240e-02 | 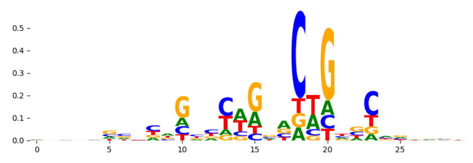 |
| pos_patterns.pattern_7 | 815 | 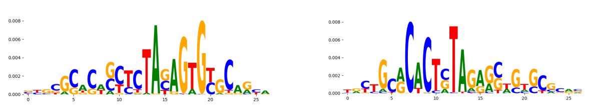 | | TN5_3 | 1.840150e-14 | 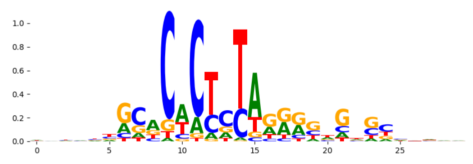 |
| pos_patterns.pattern_8 | 775 | 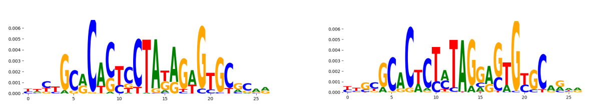 | | TN5_3 | 1.889700e-08 | 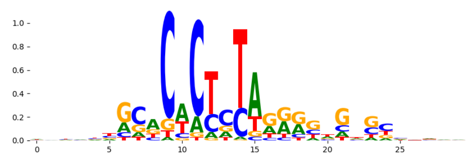 |
| pos_patterns.pattern_9 | 756 | 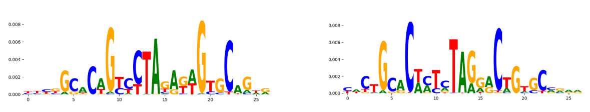 | | TN5_3 | 7.552660e-07 | 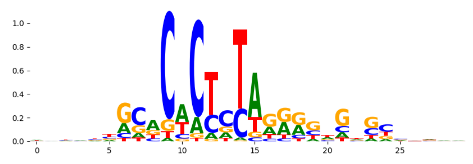 |
| pos_patterns.pattern_10 | 496 | 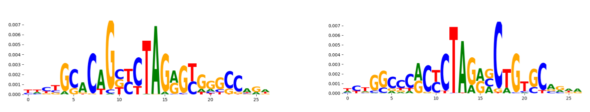 | | TN5_3 | 3.811370e-04 | 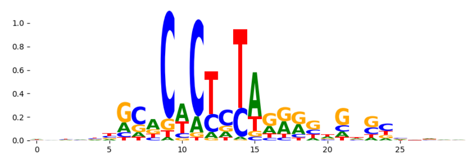 |
| pos_patterns.pattern_11 | 484 | 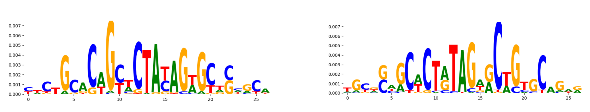 | | TN5_7 | 1.254610e-06 | 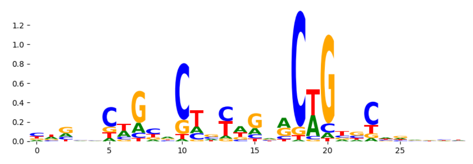 |
| pos_patterns.pattern_12 | 398 | 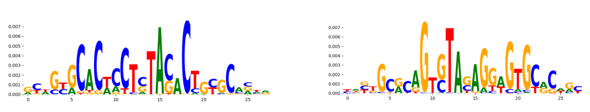 | | TN5_4 | 2.208020e-05 | 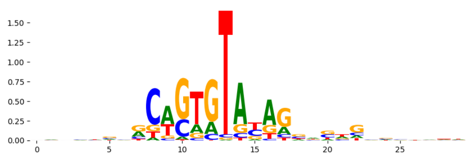 |
| pos_patterns.pattern_13 | 386 | 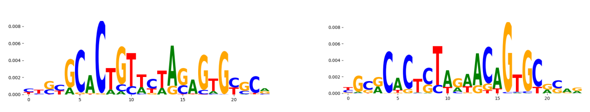 | | TN5_3 | 8.902310e-04 | 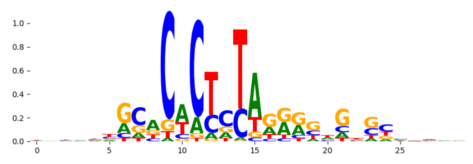 |
| pos_patterns.pattern_14 | 321 | 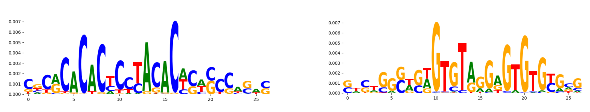 | | TN5_4 | 6.381050e-01 | 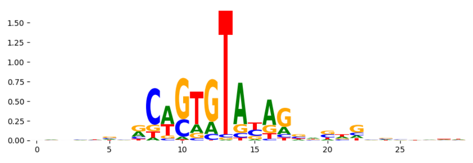 |
| pos_patterns.pattern_15 | 190 | 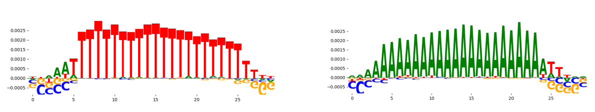 | | ZNF384_MA1125.1 | 6.461390e-02 | 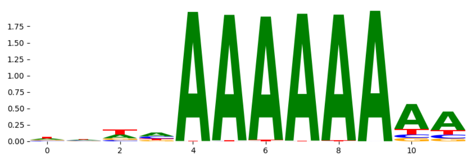 |
| pos_patterns.pattern_16 | 35 | 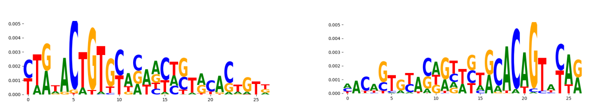 | | FOXB1_forkhead_2 | 8.769330e-01 | 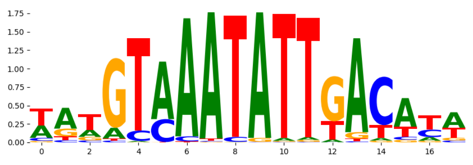 |
| pos_patterns.pattern_17 | 34 | 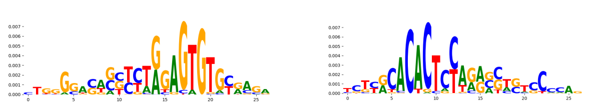 | | TN5_3 | 2.496710e-05 | 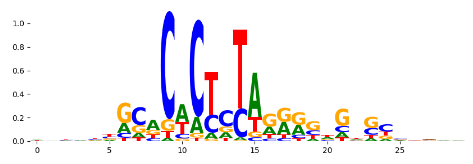 |
| pos_patterns.pattern_18 | 30 | 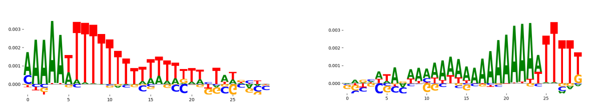 | | ZNF384_MA1125.1 | 9.537570e-02 | 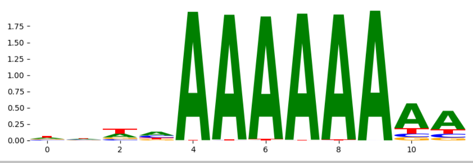 |
