## Supplementary Files 3 for "ChromBPNet: bias factorized, base-resolution deep learning models of chromatin accessibility reveal cis-regulatory sequence syntax, transcription factor footprints and regulatory variants": k562_ATAC_raw_bpnet_bias_fold3_counts_modisco.pdf

| pattern | num_seqlets | cwm_fwd | cwm_rev | TOMTOM_match | TOMTOM_qval | TOMTOM_match_logo |
|---|---|---|---|---|---|---|
| pos_patterns.pattern_0 | 6027 | | | NaN | NaN | |
| pos_patterns.pattern_1 | 4358 | | | ZFX_MOUSE.H11MO.0.B | 1.327300e-01 | |
| pos_patterns.pattern_2 | 2839 | | | ZBT7A_HUMAN.H11MO.0.A | 7.431000e-02 | |
| pos_patterns.pattern_3 | 925 | | | HSF2_HSF_1 | 1.000000e+00 | |
| pos_patterns.pattern_4 | 813 | | | RREB1_MA0073.1 | 4.534670e-02 | |
| pos_patterns.pattern_5 | 808 | | | ZN436_HUMAN.H11MO.0.C | 1.000000e+00 | |
| pos_patterns.pattern_6 | 608 | | | ZNF384_MA1125.1 | 3.313860e-02 | |
| pos_patterns.pattern_7 | 539 | | | DNASE_4 | 1.000000e+00 | |
| pos_patterns.pattern_8 | 418 | | | Zfx_MA0146.2 | 1.000000e+00 | |
| pos_patterns.pattern_9 | 371 | | | EGR2_HUMAN.H11MO.0.A | 1.294610e-03 | |
| pos_patterns.pattern_10 | 355 | | | RUNX2_HUMAN.H11MO.0.A | 6.378170e-01 | |
| pos_patterns.pattern_11 | 199 | | | SOX9_HMG_1 | 1.000000e+00 | |
| pos_patterns.pattern_12 | 188 | | | VEZF1_HUMAN.H11MO.0.C | 1.167430e-04 | |
| pos_patterns.pattern_13 | 181 | | | SP1_HUMAN.H11MO.0.A | 1.860660e-06 | |
| pos_patterns.pattern_14 | 160 | | | SP2_HUMAN.H11MO.0.A | 2.389050e-11 | |
| pos_patterns.pattern_15 | 160 | | | EGR2_HUMAN.H11MO.0.A | 1.797250e-04 | |
| pos_patterns.pattern_16 | 156 | | | NFAC1_HUMAN.H11MO.0.B | 2.777270e-01 | |
| pos_patterns.pattern_17 | 150 | | | SP5_MOUSE.H11MO.0.C | 4.528580e-05 | |
| pos_patterns.pattern_18 | 55 | | | HNF4A_nuclearreceptor_1 | 4.168720e-01 | |
| pos_patterns.pattern_19 | 54 | | | SP3_HUMAN.H11MO.0.B | 5.632300e-04 | |
| pos_patterns.pattern_20 | 53 | | | SP3_HUMAN.H11MO.0.B | 3.547680e-05 | |
| pos_patterns.pattern_21 | 22 | | | ZN263_HUMAN.H11MO.0.A | 1.255920e-01 | |
| neg_patterns.pattern_0 | 110 | | | TN5_2 | 1.074060e-03 | |
| neg_patterns.pattern_1 | 99 | | | SP2_HUMAN.H11MO.0.A | 5.572160e-06 | |
| neg_patterns.pattern_2 | 91 | | | ZFX_MOUSE.H11MO.0.B | 1.269110e-03 | |
| neg_patterns.pattern_3 | 89 | | | TN5_2 | 3.501280e-01 | |
| neg_patterns.pattern_4 | 82 | | | DNASE_3 | 1.000000e+00 | |
| neg_patterns.pattern_5 | 82 | | | ZN335_HUMAN.H11MO.0.A | 1.000000e+00 | |
| neg_patterns.pattern_6 | 81 | | | ZN331_HUMAN.H11MO.0.C | 6.314360e-01 | |
| neg_patterns.pattern_7 | 80 | | | SUH_HUMAN.H11MO.0.A | 9.042350e-01 | |

| pattern | num_seqlets | cwm_fwd | cwm_rev | TOMTOM_match | TOMTOM_qval | TOMTOM_match_logo |
|---|---|---|---|---|---|---|
| neg_patterns.pattern_8 | 75 | 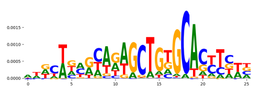 | 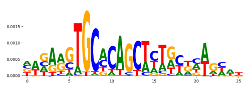 | ZFX_MOUSE.H11MO.0.B | 1.537290e-02 | 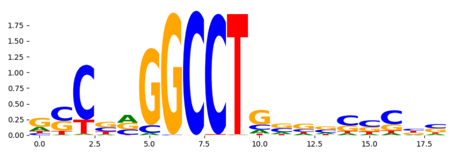 |
| neg_patterns.pattern_9 | 74 | 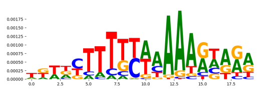 | 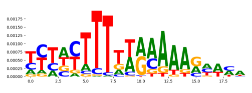 | Stat4_MA0518.1 | 5.268070e-01 | 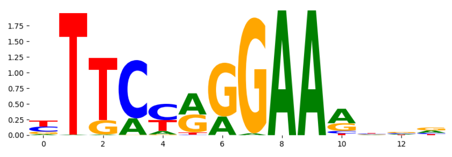 |
| neg_patterns.pattern_10 | 62 | 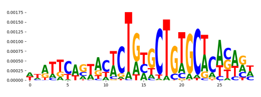 | 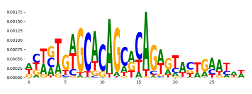 | TN5_2 | 3.466260e-03 | 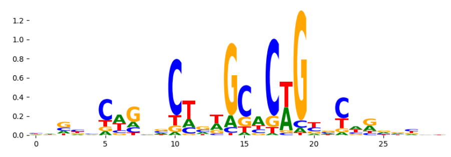 |
| neg_patterns.pattern_11 | 59 | 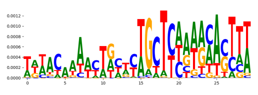 | 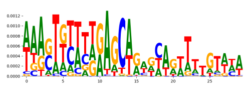 | ZEB1_MA0103.3 | 8.601340e-02 | 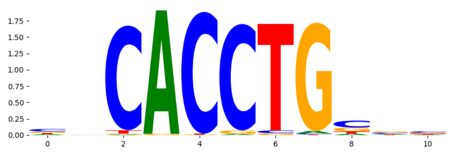 |
| neg_patterns.pattern_12 | 56 | 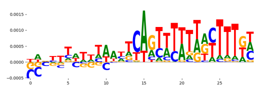 | 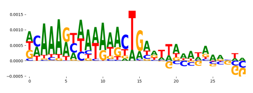 | ZN281_MOUSE.H11MO.0.A | 1.935400e-01 | 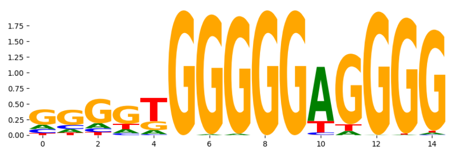 |
| neg_patterns.pattern_13 | 50 | 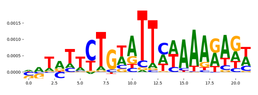 | 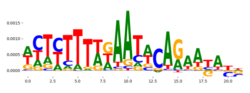 | ZN490_HUMAN.H11MO.0.C | 8.361060e-01 | 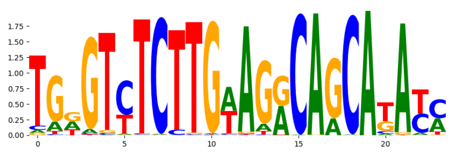 |
| neg_patterns.pattern_14 | 26 | 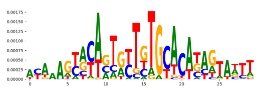 | 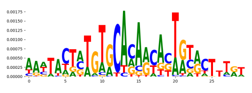 | TN5_2 | 4.132160e-04 | 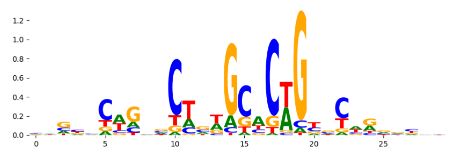 |
| neg_patterns.pattern_15 | 23 | 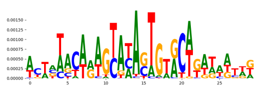 | 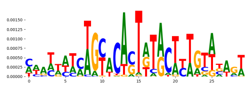 | TN5_2 | 3.918880e-02 | 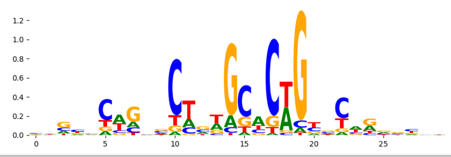 |
