## Supplementary Files 3 for "ChromBPNet: bias factorized, base-resolution deep learning models of chromatin accessibility reveal cis-regulatory sequence syntax, transcription factor footprints and regulatory variants": k562_ATAC_raw_bpnet_bias_fold3_profile_modisco.pdf

| pattern | num_seqlets | cwm_fwd | cwm_rev | TOMTOM_match | TOMTOM_qval | TOMTOM_match_logo |
|---|---|---|---|---|---|---|
| pos_patterns.pattern_0 | 8950 | 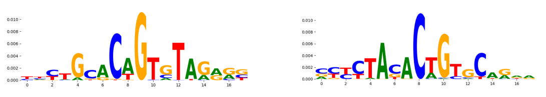 | | TN5_4 | 2.675730e-02 | 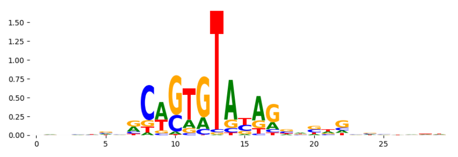 |
| pos_patterns.pattern_1 | 8338 | 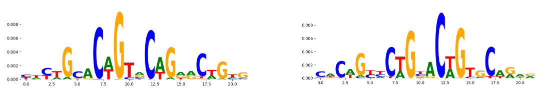 | | TN5_1 | 3.927120e-10 | 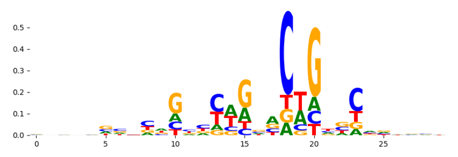 |
| pos_patterns.pattern_2 | 4280 | 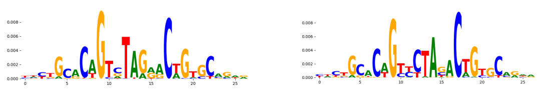 | | TN5_1 | 6.434130e-07 | 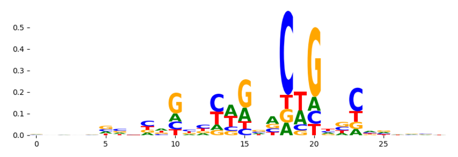 |
| pos_patterns.pattern_3 | 2223 | 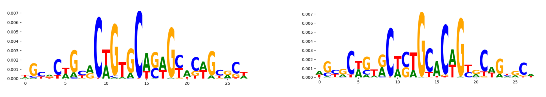 | | TN5_2 | 3.441800e-17 | 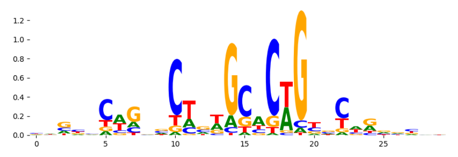 |
| pos_patterns.pattern_4 | 1042 | 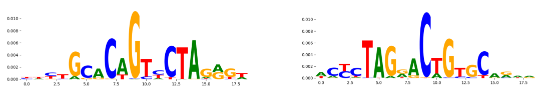 | | TN5_7 | 3.651520e-02 | 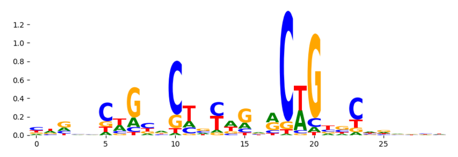 |
| pos_patterns.pattern_5 | 991 | 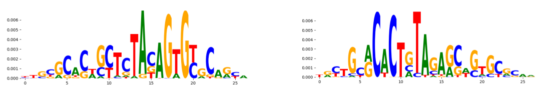 | | TN5_3 | 1.173050e-09 | 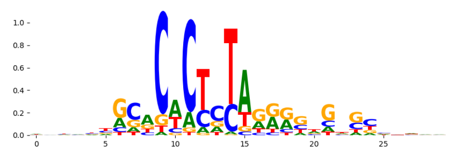 |
| pos_patterns.pattern_6 | 877 | 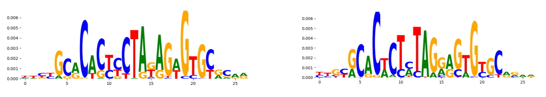 | | TN5_3 | 2.156950e-09 | 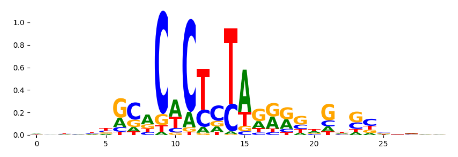 |
| pos_patterns.pattern_7 | 800 | 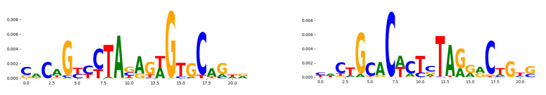 | | TN5_3 | 1.574800e-04 | 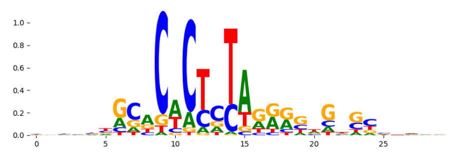 |
| pos_patterns.pattern_8 | 773 | 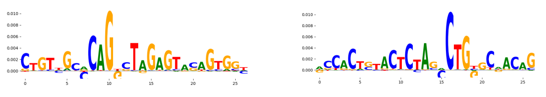 | | TN5_6 | 2.033860e-20 | 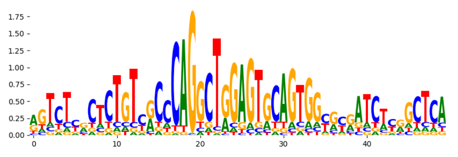 |
| pos_patterns.pattern_9 | 502 | 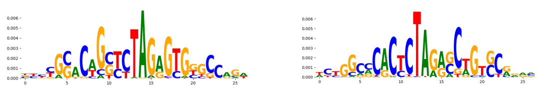 | | TN5_3 | 3.141760e-05 | 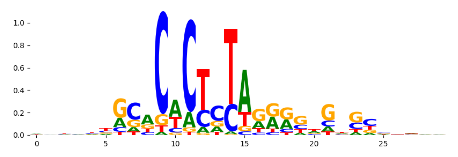 |
| pos_patterns.pattern_10 | 363 | 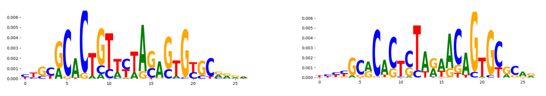 | | TN5_3 | 7.863180e-06 | 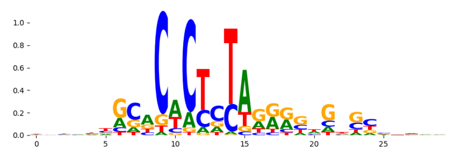 |
| pos_patterns.pattern_11 | 343 | 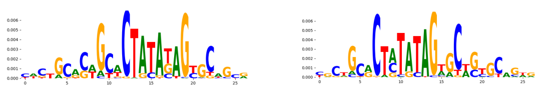 | | TN5_3 | 1.842520e-03 | 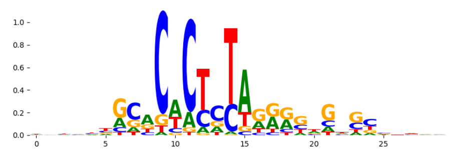 |
| pos_patterns.pattern_12 | 318 | 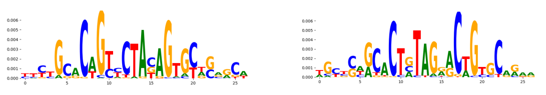 | | TN5_7 | 5.053040e-08 | 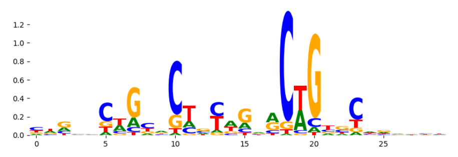 |
| pos_patterns.pattern_13 | 279 | 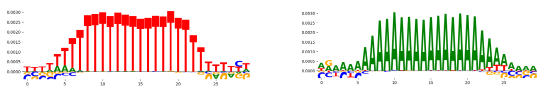 | | PRDM6_HUMAN.H11MO.0.C | 6.748890e-02 | 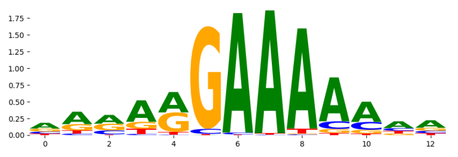 |
| pos_patterns.pattern_14 | 69 | 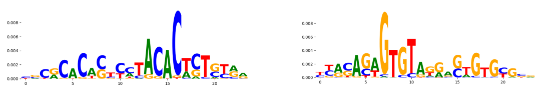 | | TN5_4 | 4.792600e-01 | 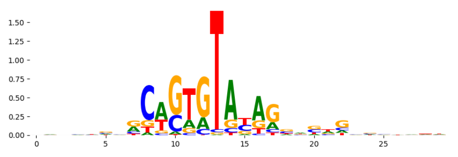 |
| pos_patterns.pattern_15 | 28 | 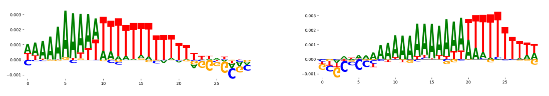 | | ZNF384_MA1125.1 | 2.338480e-02 | 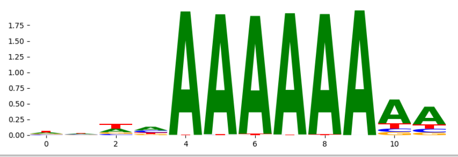 |
