## Supplementary Files 3 for "ChromBPNet: bias factorized, base-resolution deep learning models of chromatin accessibility reveal cis-regulatory sequence syntax, transcription factor footprints and regulatory variants": k562_ATAC_raw_bpnet_bias_fold4_counts_modisco.pdf

| pattern | num_seqlets | cwm_fwd | cwm_rev | TOMTOM_match | TOMTOM_qval | TOMTOM_match_logo |
|---|---|---|---|---|---|---|
| pos_patterns.pattern_0 | 10471 | | | NaN | NaN | |
| pos_patterns.pattern_1 | 2449 | | | ZFX_MOUSE.H11MO.0.B | 1.862750e-02 | |
| pos_patterns.pattern_2 | 1351 | | | NR2F1_nuclearreceptor_3 | 5.930350e-01 | |
| pos_patterns.pattern_3 | 813 | | | NaN | NaN | |
| pos_patterns.pattern_4 | 638 | | | ZN436_HUMAN.H11MO.0.C | 1.000000e+00 | |
| pos_patterns.pattern_5 | 531 | | | DNASE_4 | 1.000000e+00 | |
| pos_patterns.pattern_6 | 417 | | | RUNX2_HUMAN.H11MO.0.A | 2.899030e-01 | |
| pos_patterns.pattern_7 | 353 | | | ZN708_HUMAN.H11MO.0.C | 8.424740e-01 | |
| pos_patterns.pattern_8 | 314 | | | RUNX3_MA0684.1 | 2.569040e-01 | |
| pos_patterns.pattern_9 | 298 | | | LHX6_MOUSE.H11MO.0.C | 9.999880e-01 | |
| pos_patterns.pattern_10 | 288 | | | ZNF384_MA1125.1 | 3.859580e-02 | |
| pos_patterns.pattern_11 | 244 | | | MAZ_HUMAN.H11MO.0.A | 4.578520e-09 | |
| pos_patterns.pattern_12 | 183 | | | RUNX2_HUMAN.H11MO.0.A | 1.896700e-01 | |
| pos_patterns.pattern_13 | 183 | | | ZBTB6_HUMAN.H11MO.0.C | 4.113100e-02 | |
| pos_patterns.pattern_14 | 168 | | | KLF15_HUMAN.H11MO.0.A | 1.795350e-01 | |
| pos_patterns.pattern_15 | 165 | | | RREB1_MA0073.1 | 2.428990e-01 | |
| pos_patterns.pattern_16 | 138 | | | OSR2_HUMAN.H11MO.0.C | 1.003230e-01 | |
| pos_patterns.pattern_17 | 95 | | | ZBT17_MOUSE.H11MO.0.A | 9.470300e-03 | |
| neg_patterns.pattern_0 | 1352 | | | SP3_HUMAN.H11MO.0.B | 3.699310e-01 | |
| neg_patterns.pattern_1 | 56 | | | SP2_HUMAN.H11MO.0.A | 1.945760e-03 | |
