## Supplementary Files 3 for "ChromBPNet: bias factorized, base-resolution deep learning models of chromatin accessibility reveal cis-regulatory sequence syntax, transcription factor footprints and regulatory variants": k562_ATAC_raw_bpnet_bias_fold4_profile_modisco.pdf

| pattern | num_seqlets | cwm_fwd | cwm_rev | TOMTOM_match | TOMTOM_qval | TOMTOM_match_logo |
|---|---|---|---|---|---|---|
| pos_patterns.pattern_0 | 8647 | 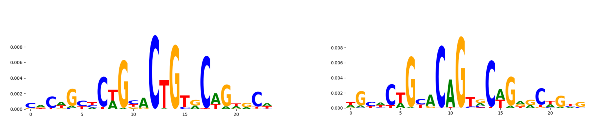 | | TN5_1 | 4.637900e-08 | 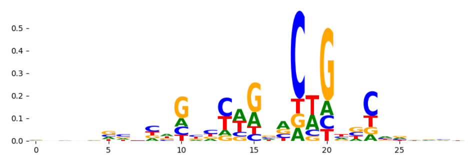 |
| pos_patterns.pattern_1 | 8511 | 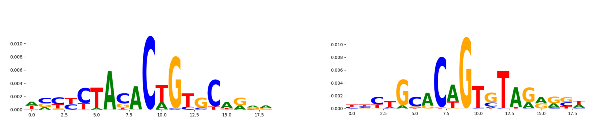 | | TN5_8 | 2.637640e-02 | 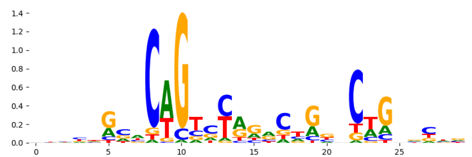 |
| pos_patterns.pattern_2 | 4263 | 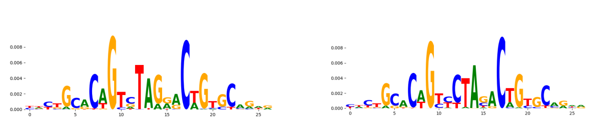 | | TN5_1 | 5.456290e-07 | 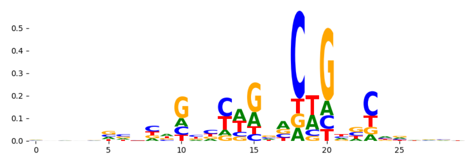 |
| pos_patterns.pattern_3 | 2571 | 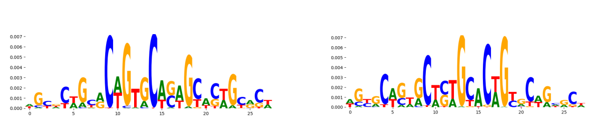 | | TN5_2 | 1.712550e-18 | 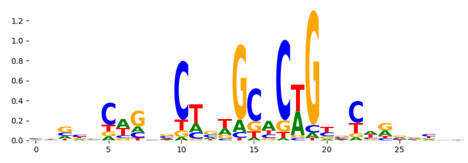 |
| pos_patterns.pattern_4 | 989 | 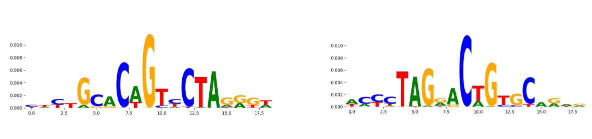 | | TN5_1 | 3.126240e-02 | 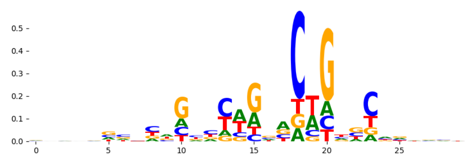 |
| pos_patterns.pattern_5 | 894 | 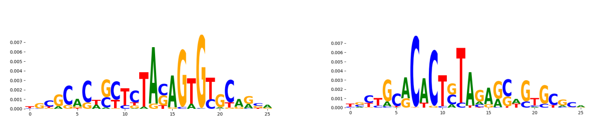 | | TN5_3 | 2.745060e-13 | 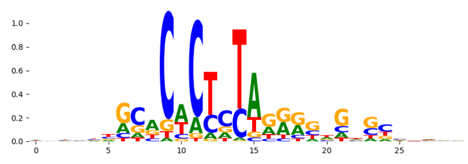 |
| pos_patterns.pattern_6 | 769 | 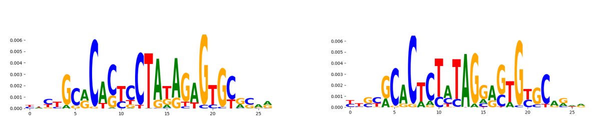 | | TN5_3 | 2.952300e-07 | 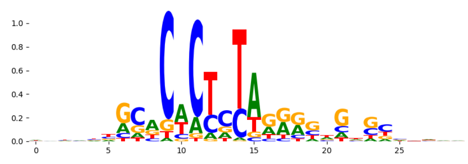 |
| pos_patterns.pattern_7 | 737 | 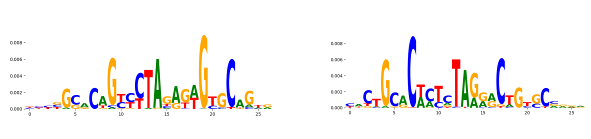 | | TN5_3 | 1.392260e-05 | 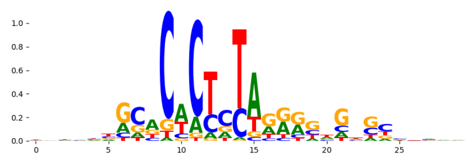 |
| pos_patterns.pattern_8 | 679 | 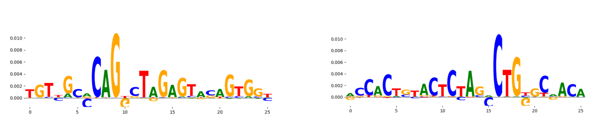 | | TN5_6 | 7.419670e-15 | 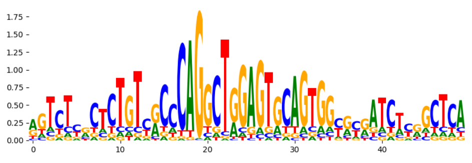 |
| pos_patterns.pattern_9 | 488 | 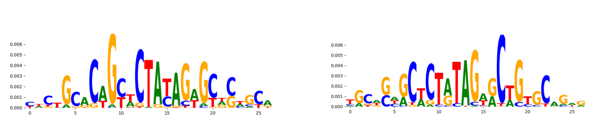 | | TN5_7 | 1.228760e-04 | 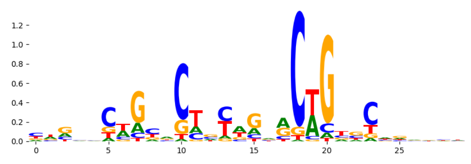 |
| pos_patterns.pattern_10 | 463 | 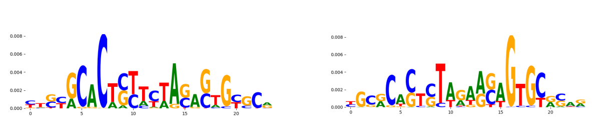 | | TN5_3 | 5.550970e-06 | 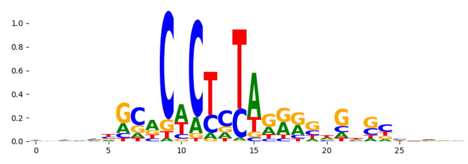 |
| pos_patterns.pattern_11 | 324 | 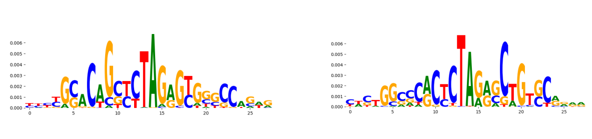 | | TN5_1 | 4.269920e-04 | 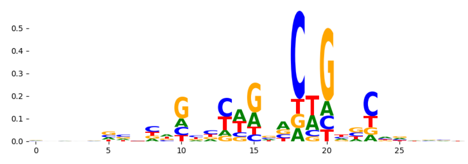 |
| pos_patterns.pattern_12 | 226 | 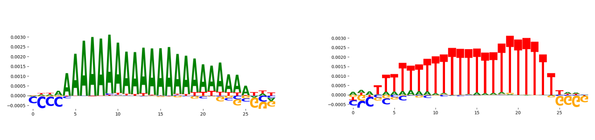 | | ZNF384_MA1125.1 | 4.853100e-02 | 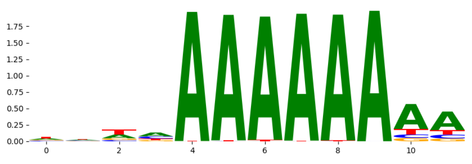 |
| pos_patterns.pattern_13 | 174 | 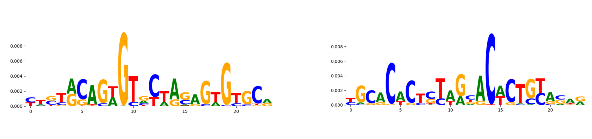 | | TN5_4 | 5.679640e-02 | 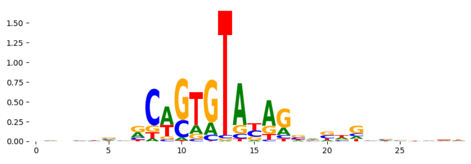 |
| pos_patterns.pattern_14 | 68 | 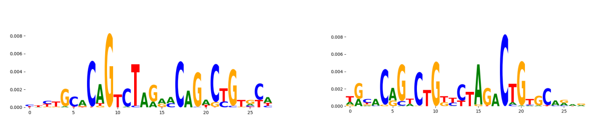 | | TN5_8 | 7.577830e-06 | 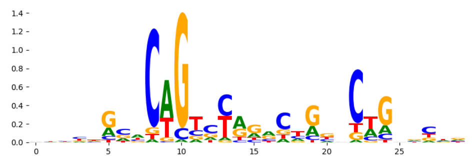 |
| pos_patterns.pattern_15 | 55 | 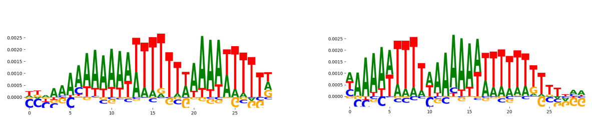 | | ZNF384_MA1125.1 | 1.971230e-01 | 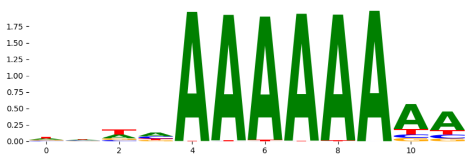 |
| pos_patterns.pattern_16 | 52 | 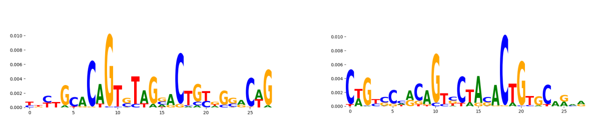 | | TN5_7 | 4.432020e-04 | 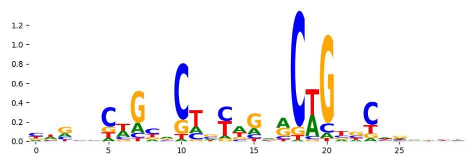 |
| pos_patterns.pattern_17 | 51 | 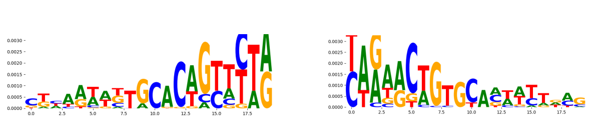 | | TN5_1 | 1.654870e-03 | 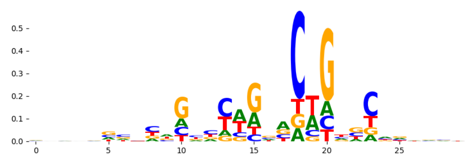 |
| pos_patterns.pattern_18 | 21 | 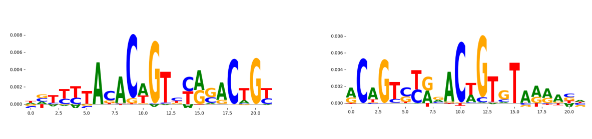 | | TN5_8 | 6.126630e-03 | 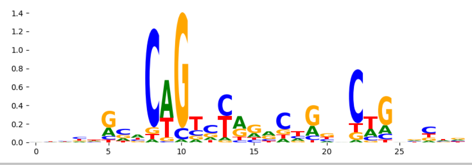 |
