## Supplementary Files 3 for "ChromBPNet: bias factorized, base-resolution deep learning models of chromatin accessibility reveal cis-regulatory sequence syntax, transcription factor footprints and regulatory variants": k562_DNASE_raw_bpnet_bias_fold0_counts_modisco.pdf

| pattern | num_seqlets | cwm_fwd | cwm_rev | TOMTOM_match | TOMTOM_qval | TOMTOM_match_logo |
|---|---|---|---|---|---|---|
| pos_patterns.pattern_0 | 3334 | | | SP2_HUMAN.H11MO.0.A | 3.533940e-09 | |
| pos_patterns.pattern_1 | 2362 | | | ZNF384_MA1125.1 | 4.767260e-02 | |
| pos_patterns.pattern_2 | 1336 | | | DNASE_2 | 4.345240e-01 | |
| pos_patterns.pattern_3 | 999 | | | SP2_HUMAN.H11MO.0.A | 3.643640e-05 | |
| pos_patterns.pattern_4 | 616 | | | SP2_HUMAN.H11MO.0.A | 3.055720e-04 | |
| pos_patterns.pattern_5 | 520 | | | PRDM6_HUMAN.H11MO.0.C | 1.157740e-02 | |
| pos_patterns.pattern_6 | 491 | | | FOXB1_forkhead_2 | 1.576520e-01 | |
| pos_patterns.pattern_7 | 447 | | | FOXG1_forkhead_1 | 5.712610e-01 | |
| pos_patterns.pattern_8 | 238 | | | IRF1_HUMAN.H11MO.0.A | 2.938620e-02 | |
| pos_patterns.pattern_9 | 227 | | | STAT1_MOUSE.H11MO.0.A | 3.865050e-01 | |
| pos_patterns.pattern_10 | 219 | | | NaN | NaN | |
| pos_patterns.pattern_11 | 144 | | | ZIM3_HUMAN.H11MO.0.C | 1.000000e+00 | |
| pos_patterns.pattern_12 | 55 | | | SOX8_HMG_3 | 1.000000e+00 | |
| neg_patterns.pattern_0 | 1544 | | | DNASE_1 | 1.000000e+00 | |
| neg_patterns.pattern_1 | 1309 | | | MYB_HUMAN.H11MO.0.A | 2.222720e-01 | |
| neg_patterns.pattern_2 | 1001 | | | NFIC_MA0161.2 | 8.809010e-01 | |
| neg_patterns.pattern_3 | 634 | | | CTCF_C2H2_1 | 2.439580e-01 | |
| neg_patterns.pattern_4 | 490 | | | DNASE_2 | 8.722250e-02 | |
| neg_patterns.pattern_5 | 457 | | | SALL4_HUMAN.H11MO.0.B | 8.932590e-02 | |
| neg_patterns.pattern_6 | 371 | | | COE1_MOUSE.H11MO.0.A | 6.161280e-04 | |
| neg_patterns.pattern_7 | 291 | | | NHLH1_MA0048.2 | 5.711200e-02 | |
| neg_patterns.pattern_8 | 274 | | | NR2E1_nuclearreceptor_2 | 1.000000e+00 | |
| neg_patterns.pattern_9 | 264 | | | NFIC_HUMAN.H11MO.0.A | 3.037560e-01 | |
| neg_patterns.pattern_10 | 247 | | | P73_HUMAN.H11MO.0.A | 4.394040e-03 | |
| neg_patterns.pattern_11 | 166 | | | ZN257_HUMAN.H11MO.0.C | 6.461030e-01 | |
| neg_patterns.pattern_12 | 103 | | | ZN770_HUMAN.H11MO.0.C | 5.231240e-01 | |
| neg_patterns.pattern_13 | 56 | | | ZFX_MOUSE.H11MO.0.B | 3.834680e-01 | |
| neg_patterns.pattern_14 | 28 | | | ZN547_HUMAN.H11MO.0.C | 7.633560e-02 | |
