## Supplementary Files 3 for "ChromBPNet: bias factorized, base-resolution deep learning models of chromatin accessibility reveal cis-regulatory sequence syntax, transcription factor footprints and regulatory variants": k562_DNASE_raw_bpnet_bias_fold0_profile_modisco.pdf

| pattern | num_seqlets | cwm_fwd | cwm_rev | TOMTOM_match | TOMTOM_qval | TOMTOM_match_logo |
|---|---|---|---|---|---|---|
| pos_patterns.pattern_0 | 5693 | | | DNASE_1 | 2.635070e-01 | |
| pos_patterns.pattern_1 | 3621 | | | CREM_HUMAN.H11MO.0.C | 1.000000e+00 | |
| pos_patterns.pattern_2 | 2711 | | | DNASE_2 | 3.161020e-04 | |
| pos_patterns.pattern_3 | 1749 | | | ZNF384_MA1125.1 | 1.901360e-01 | |
| pos_patterns.pattern_4 | 921 | | | PRDM6_HUMAN.H11MO.0.C | 7.562760e-02 | |
| pos_patterns.pattern_5 | 921 | | | NaN | NaN | |
| pos_patterns.pattern_6 | 859 | | | MEF2B_HUMAN.H11MO.0.A | 2.146410e-01 | |
| pos_patterns.pattern_7 | 630 | | | KLF3_HUMAN.H11MO.0.B | 3.518420e-01 | |
| pos_patterns.pattern_8 | 516 | | | EGR1_MOUSE.H11MO.0.A | 1.014840e-01 | |
| pos_patterns.pattern_9 | 497 | | | ZFX_MOUSE.H11MO.0.B | 2.755700e-01 | |
| pos_patterns.pattern_10 | 455 | | | MXI1_HUMAN.H11MO.0.A | 7.342290e-01 | |
| pos_patterns.pattern_11 | 270 | | | SP1_HUMAN.H11MO.0.A | 6.910940e-01 | |
| pos_patterns.pattern_12 | 268 | | | THAP1_HUMAN.H11MO.0.C | 3.162780e-01 | |
| pos_patterns.pattern_13 | 224 | | | TYY1_HUMAN.H11MO.0.A | 1.322730e-02 | |
| pos_patterns.pattern_14 | 209 | | | TN5_6 | 9.054130e-21 | |
| pos_patterns.pattern_15 | 190 | | | ZN770_HUMAN.H11MO.0.C | 2.604730e-08 | |
| pos_patterns.pattern_16 | 161 | | | FOXB1_forkhead_2 | 1.557050e-01 | |
| pos_patterns.pattern_17 | 139 | | | ZN770_HUMAN.H11MO.0.C | 6.701660e-08 | |
| pos_patterns.pattern_18 | 133 | | | CTCFL_HUMAN.H11MO.0.A | 9.706390e-04 | |
| pos_patterns.pattern_19 | 122 | | | ZNF384_MA1125.1 | 1.861490e-01 | |
| pos_patterns.pattern_20 | 119 | | | TAF1_HUMAN.H11MO.0.A | 4.224020e-01 | |
| pos_patterns.pattern_21 | 90 | | | DNASE_5 | 5.536750e-01 | |
| pos_patterns.pattern_22 | 86 | | | STAT1_MOUSE.H11MO.0.A | 6.668460e-02 | |
| pos_patterns.pattern_23 | 86 | | | SP1_MOUSE.H11MO.0.A | 3.532660e-08 | |
| pos_patterns.pattern_24 | 55 | | | ZN121_HUMAN.H11MO.0.C | 1.017460e-03 | |
| pos_patterns.pattern_25 | 41 | | | ZN770_HUMAN.H11MO.0.C | 6.074630e-02 | |
| pos_patterns.pattern_26 | 33 | | | MAF_MOUSE.H11MO.0.A | 5.617860e-01 | |
| pos_patterns.pattern_27 | 32 | | | RARA_HUMAN.H11MO.0.A | 1.537110e-01 | |
| pos_patterns.pattern_28 | 31 | | | DNASE_6 | 3.557460e-01 | |
| neg_patterns.pattern_0 | 24 | | | PRDM6_HUMAN.H11MO.0.C | 3.875060e-01 | |
