## Supplementary Files 3 for "ChromBPNet: bias factorized, base-resolution deep learning models of chromatin accessibility reveal cis-regulatory sequence syntax, transcription factor footprints and regulatory variants": k562_DNASE_raw_bpnet_bias_fold1_counts_modisco.pdf

| pattern | num_seqlets | cwm_fwd | cwm_rev | TOMTOM_match | TOMTOM_qval | TOMTOM_match_logo |
|---|---|---|---|---|---|---|
| pos_patterns.pattern_0 | 2926 | | | SP1_MOUSE.H11MO.0.A | 4.141100e-10 | |
| pos_patterns.pattern_1 | 2716 | | | SP2_HUMAN.H11MO.0.A | 6.082560e-07 | |
| pos_patterns.pattern_2 | 1847 | | | PRDM6_HUMAN.H11MO.0.C | 8.515960e-03 | |
| pos_patterns.pattern_3 | 1091 | | | ZNF740_C2H2_1 | 1.877430e-02 | |
| pos_patterns.pattern_4 | 933 | | | RREB1_MA0073.1 | 5.720710e-01 | |
| pos_patterns.pattern_5 | 933 | | | DNASE_2 | 1.457250e-01 | |
| pos_patterns.pattern_6 | 740 | | | SOX17_HUMAN.H11MO.0.C | 1.000000e+00 | |
| pos_patterns.pattern_7 | 595 | | | SP2_HUMAN.H11MO.0.A | 1.906920e-05 | |
| pos_patterns.pattern_8 | 384 | | | RUNX3_RUNX_3 | 1.000000e+00 | |
| pos_patterns.pattern_9 | 275 | | | RELB_MA1117.1 | 1.000000e+00 | |
| pos_patterns.pattern_10 | 255 | | | NFAC4_HUMAN.H11MO.0.C | 6.388840e-01 | |
| pos_patterns.pattern_11 | 207 | | | ZN431_MOUSE.H11MO.0.C | 8.170400e-01 | |
| pos_patterns.pattern_12 | 188 | | | LYL1_HUMAN.H11MO.0.A | 5.938620e-01 | |
| pos_patterns.pattern_13 | 180 | | | ZN257_HUMAN.H11MO.0.C | 1.000000e+00 | |
| pos_patterns.pattern_14 | 72 | | | ISX_homeodomain_1 | 1.460030e-01 | |
| pos_patterns.pattern_15 | 49 | | | TN5_6 | 5.986190e-04 | |
| pos_patterns.pattern_16 | 42 | | | EWSR1-FLI1_MA0149.1 | 5.623700e-11 | |
| pos_patterns.pattern_17 | 40 | | | VDR_MA0693.2 | 1.000000e+00 | |
| pos_patterns.pattern_18 | 24 | | | SMAD2+SMAD3+SMAD4_MA0513.1 | 7.998340e-02 | |
| pos_patterns.pattern_19 | 24 | | | ZN770_HUMAN.H11MO.0.C | 7.065100e-07 | |
| pos_patterns.pattern_20 | 21 | | | TN5_6 | 2.675810e-06 | |
| neg_patterns.pattern_0 | 2688 | | | DNASE_2 | 1.934110e-03 | |
| neg_patterns.pattern_1 | 1712 | | | DNASE_2 | 5.153310e-01 | |
| neg_patterns.pattern_2 | 807 | | | EGR1_MOUSE.H11MO.0.A | 5.715480e-03 | |
| neg_patterns.pattern_3 | 463 | | | BARHL2_homeodomain_2 | 1.000000e+00 | |
| neg_patterns.pattern_4 | 410 | | | PRDM6_HUMAN.H11MO.0.C | 2.721510e-02 | |
| neg_patterns.pattern_5 | 373 | | | DNASE_1 | 1.658490e-01 | |
| neg_patterns.pattern_6 | 357 | | | E2F7_HUMAN.H11MO.0.B | 1.000000e+00 | |
| neg_patterns.pattern_7 | 241 | | | PPARG_MA0066.1 | 8.912570e-01 | |
| neg_patterns.pattern_8 | 227 | | | LYL1_HUMAN.H11MO.0.A | 3.529440e-01 | |

| pattern | num_seqlets | cwm_fwd | cwm_rev | TOMTOM_match | TOMTOM_qval | TOMTOM_match_logo |
|---|---|---|---|---|---|---|
| neg_patterns.pattern_9 | 174 | 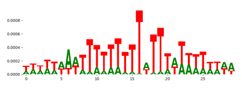 | 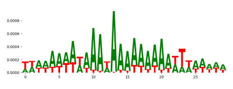 | ELF5_MOUSE.H11MO.0.A | 7.360410e-04 | 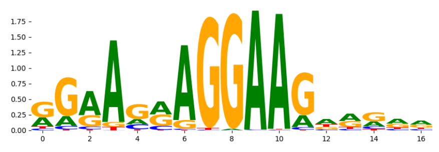 |
| neg_patterns.pattern_10 | 126 | 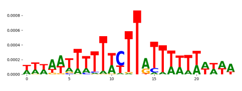 | 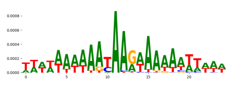 | PRDM6_HUMAN.H11MO.0.C | 9.146250e-02 | 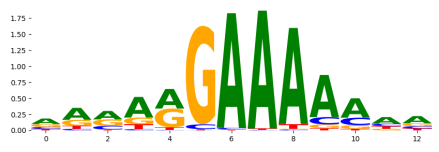 |
| neg_patterns.pattern_11 | 104 | 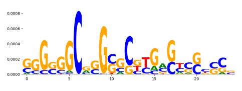 | 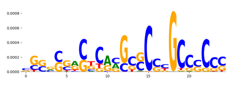 | ZN257_HUMAN.H11MO.0.C | 1.000000e+00 | 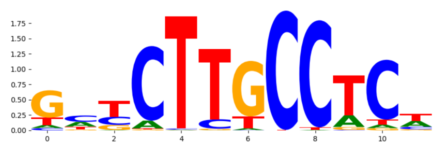 |
| neg_patterns.pattern_12 | 93 | 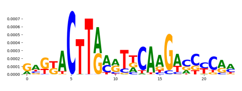 | 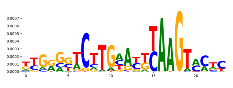 | NKX28_HUMAN.H11MO.0.C | 1.000000e+00 | 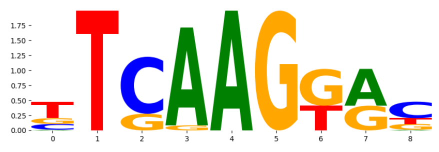 |
| neg_patterns.pattern_13 | 80 | 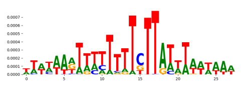 | 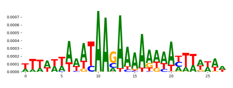 | GATA3_GATA_1 | 5.050270e-02 | 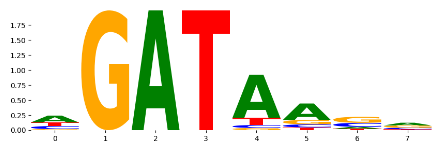 |
| neg_patterns.pattern_14 | 72 | 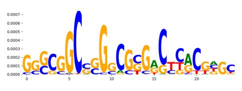 | 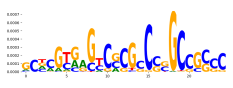 | P73_HUMAN.H11MO.0.A | 2.057810e-01 | 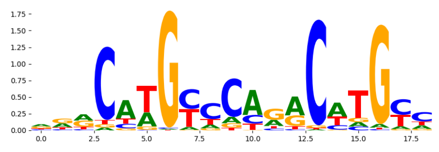 |
| neg_patterns.pattern_15 | 50 | 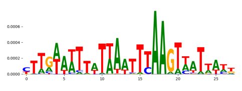 | 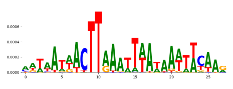 | NKX2-3_MA0672.1 | 1.918850e-01 | 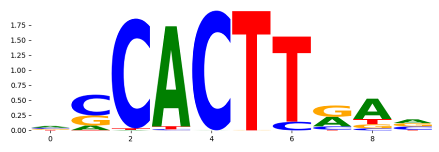 |
| neg_patterns.pattern_16 | 22 | 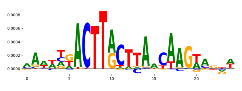 | 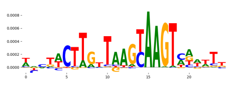 | DNASE_6 | 1.708030e-01 | 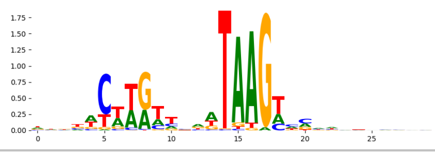 |
