## Supplementary Files 3 for "ChromBPNet: bias factorized, base-resolution deep learning models of chromatin accessibility reveal cis-regulatory sequence syntax, transcription factor footprints and regulatory variants": k562_DNASE_raw_bpnet_bias_fold1_profile_modisco.pdf

| pattern | num_seqlets | cwm_fwd | cwm_rev | TOMTOM_match | TOMTOM_qval | TOMTOM_match_logo |
|---|---|---|---|---|---|---|
| pos_patterns.pattern_0 | 4184 | | | TFAP2A_AP2_4 | 2.532460e-01 | |
| pos_patterns.pattern_1 | 3809 | | | ERR2_HUMAN.H11MO.0.A | 1.000000e+00 | |
| pos_patterns.pattern_2 | 2736 | | | DNASE_2 | 8.434530e-01 | |
| pos_patterns.pattern_3 | 1458 | | | FOXJ3_HUMAN.H11MO.0.A | 1.777370e-01 | |
| pos_patterns.pattern_4 | 1351 | | | CTCFL_HUMAN.H11MO.0.A | 2.710010e-01 | |
| pos_patterns.pattern_5 | 1154 | | | ZFX_MOUSE.H11MO.0.B | 3.428800e-02 | |
| pos_patterns.pattern_6 | 856 | | | DNASE_1 | 1.000000e+00 | |
| pos_patterns.pattern_7 | 850 | | | PRDM6_HUMAN.H11MO.0.C | 8.057040e-02 | |
| pos_patterns.pattern_8 | 807 | | | CTCFL_MOUSE.H11MO.0.A | 7.542130e-04 | |
| pos_patterns.pattern_9 | 610 | | | MEF2B_HUMAN.H11MO.0.A | 2.148970e-01 | |
| pos_patterns.pattern_10 | 331 | | | NFIB_MOUSE.H11MO.0.C | 2.227970e-01 | |
| pos_patterns.pattern_11 | 247 | | | ZN770_HUMAN.H11MO.0.C | 4.280760e-03 | |
| pos_patterns.pattern_12 | 225 | | | SP1_HUMAN.H11MO.0.A | 2.581380e-09 | |
| pos_patterns.pattern_13 | 193 | | | TN5_6 | 2.826920e-16 | |
| pos_patterns.pattern_14 | 171 | | | TN5_6 | 2.260950e-01 | |
| pos_patterns.pattern_15 | 159 | | | MYBA_MOUSE.H11MO.0.C | 1.253610e-01 | |
| pos_patterns.pattern_16 | 152 | | | SMAD3_HUMAN.H11MO.0.B | 6.498270e-01 | |
| pos_patterns.pattern_17 | 148 | | | ZN770_HUMAN.H11MO.0.C | 9.034850e-12 | |
| pos_patterns.pattern_18 | 117 | | | ZN250_HUMAN.H11MO.0.C | 1.000000e+00 | |
| pos_patterns.pattern_19 | 80 | | | FOXB1_forkhead_2 | 2.831350e-01 | |
| pos_patterns.pattern_20 | 62 | | | DNASE_5 | 1.207530e-01 | |
| pos_patterns.pattern_21 | 57 | | | TN5_6 | 3.335100e-09 | |
| pos_patterns.pattern_22 | 49 | | | ZN770_HUMAN.H11MO.0.C | 6.213010e-04 | |
| pos_patterns.pattern_23 | 34 | | | ITF2_HUMAN.H11MO.0.C | 1.000000e+00 | |
| pos_patterns.pattern_24 | 30 | | | YY1_MA0095.2 | 1.565180e-01 | |
| pos_patterns.pattern_25 | 27 | | | DNASE_5 | 1.383860e-10 | |
| pos_patterns.pattern_26 | 25 | | | PO3F1_MOUSE.H11MO.0.C | 1.000000e+00 | |
| pos_patterns.pattern_27 | 25 | | | Arid5a_MA0602.1 | 1.000000e+00 | |
| pos_patterns.pattern_28 | 24 | | | RXRA_MOUSE.H11MO.0.A | 7.124840e-01 | |

| pattern | num_seqlets | cwm_fwd | cwm_rev | TOMTOM_match | TOMTOM_qval | TOMTOM_match_logo |
|---|---|---|---|---|---|---|
| neg_patterns.pattern_0 | 30 | 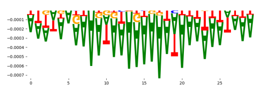 | 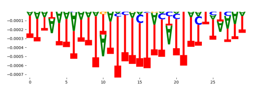 | PRDM6_HUMAN.H11MO.0.C | 5.031330e-03 | 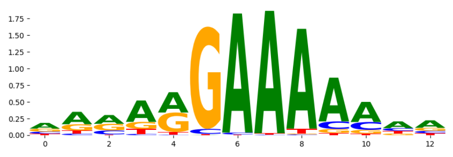 |
| neg_patterns.pattern_1 | 24 | 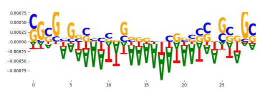 | 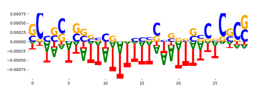 | ZNF384_MA1125.1 | 2.271800e-02 | 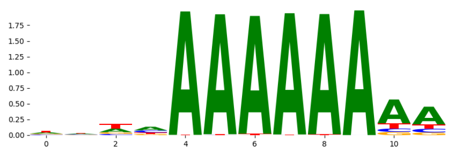 |
| neg_patterns.pattern_2 | 20 | 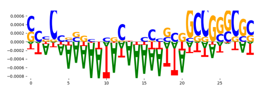 | 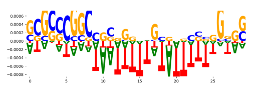 | FOXJ3_HUMAN.H11MO.0.A | 8.812020e-02 | 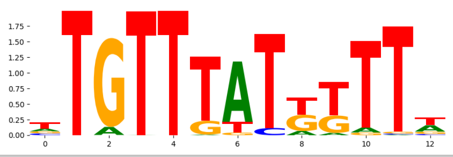 |
