## Supplementary Files 3 for "ChromBPNet: bias factorized, base-resolution deep learning models of chromatin accessibility reveal cis-regulatory sequence syntax, transcription factor footprints and regulatory variants": k562_DNASE_raw_bpnet_bias_fold2_counts_modisco.pdf

| pattern | num_seqlets | cwm_fwd / cwm_rev |
|---|---|---|
| pos_patterns.pattern_0 | 1900 | 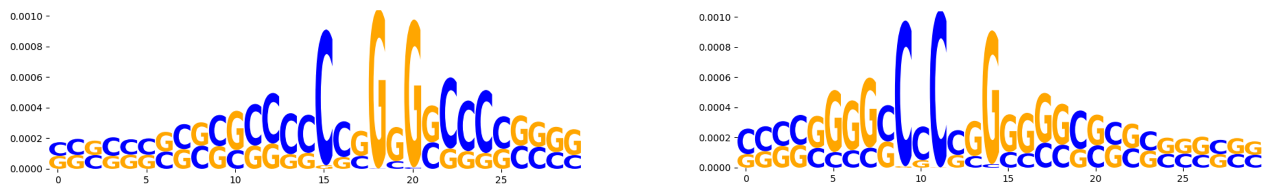 |
| pos_patterns.pattern_1 | 1605 | 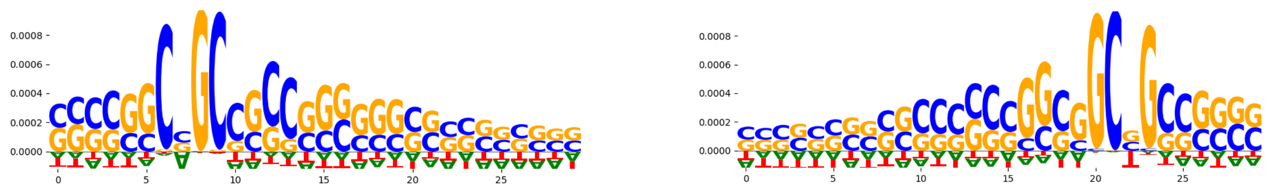 |
| pos_patterns.pattern_2 | 1295 | 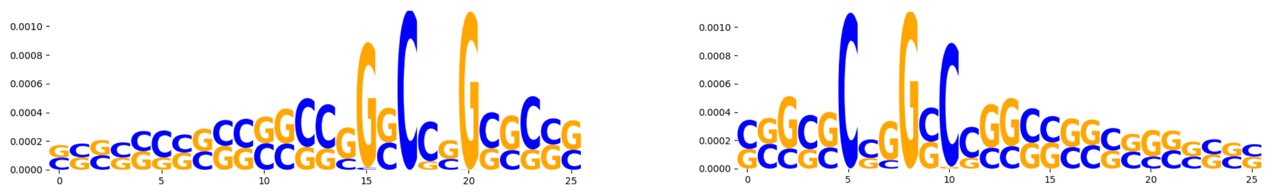 |
| pos_patterns.pattern_3 | 837 | 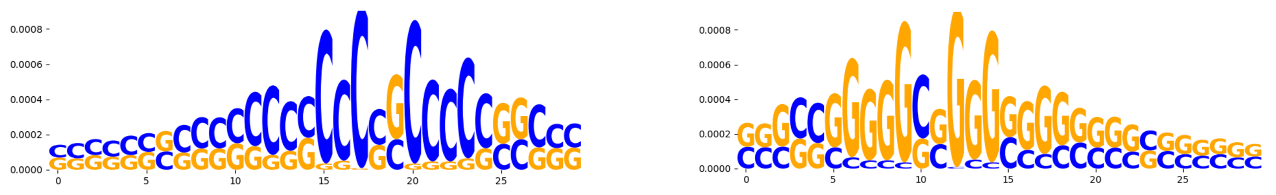 |
| pos_patterns.pattern_4 | 337 | 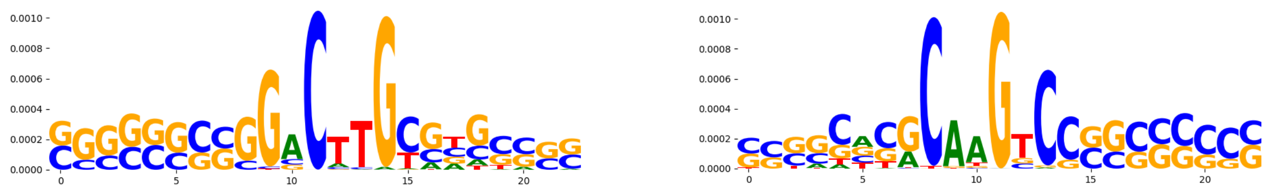 |
| pos_patterns.pattern_5 | 336 | 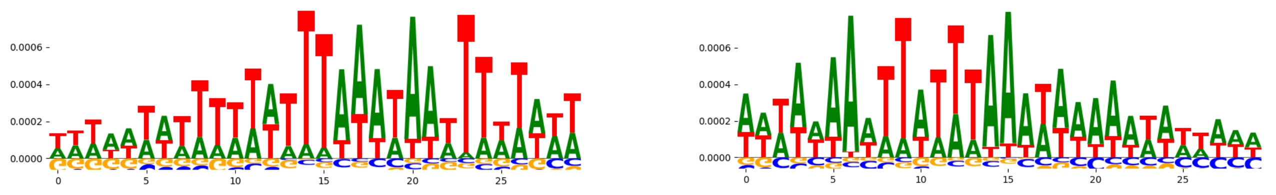 |
| pos_patterns.pattern_6 | 292 | 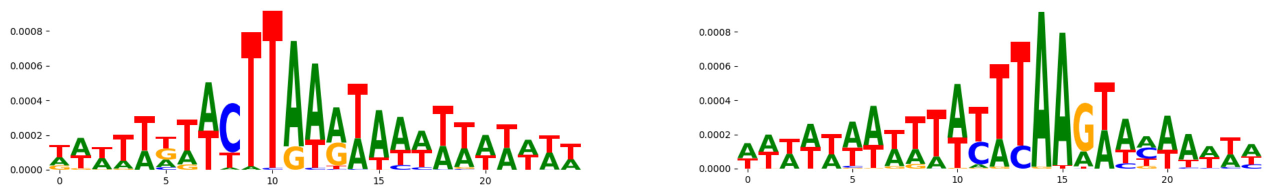 |
| pos_patterns.pattern_7 | 255 | 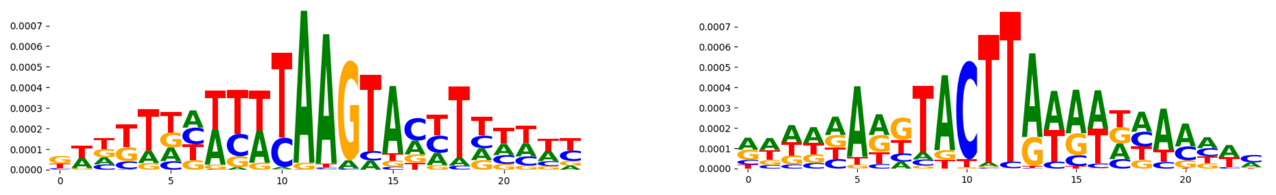 |
| pos_patterns.pattern_8 | 218 | 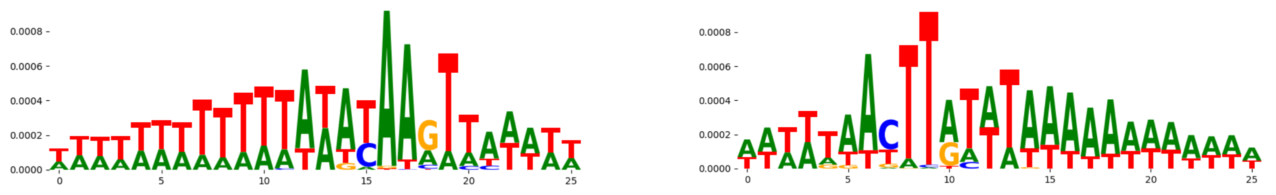 |
| pos_patterns.pattern_9 | 214 | 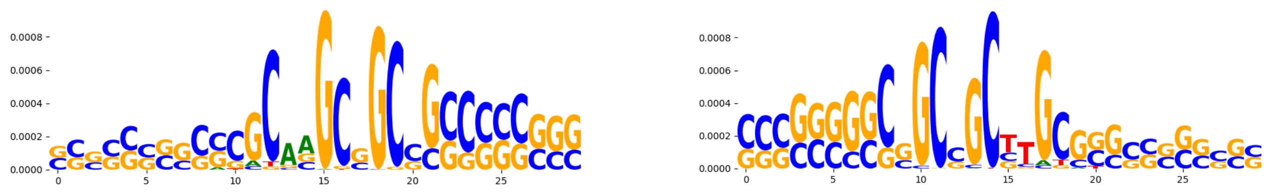 |
| pos_patterns.pattern_10 | 189 | 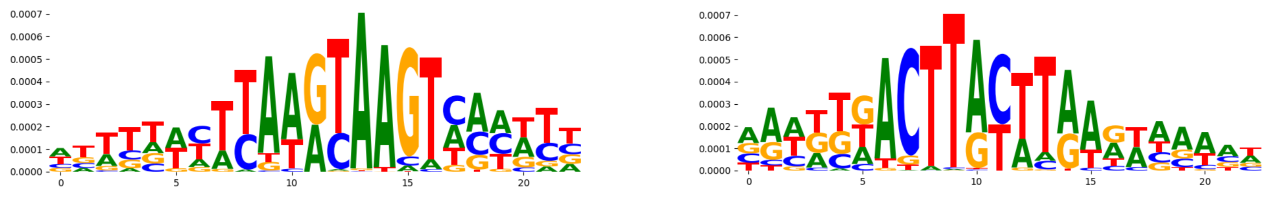 |
| pos_patterns.pattern_11 | 172 | 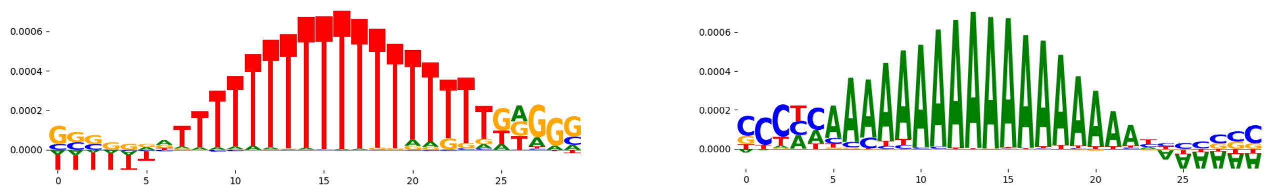 |
| pos_patterns.pattern_12 | 118 | 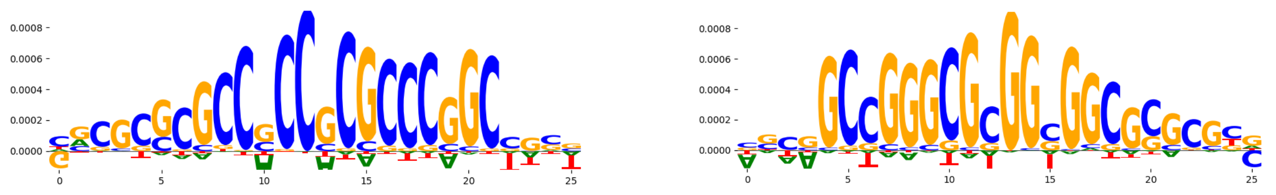 |
| pos_patterns.pattern_13 | 109 | 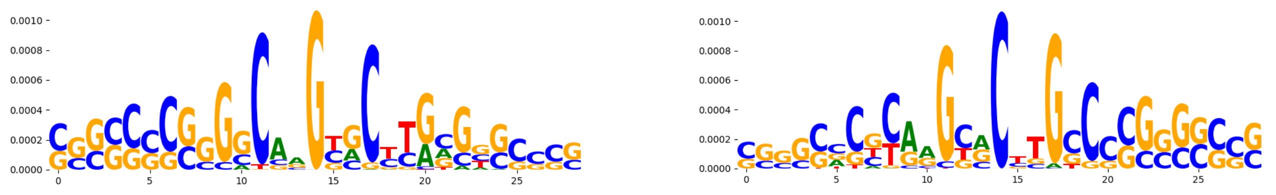 |
| pos_patterns.pattern_14 | 100 | 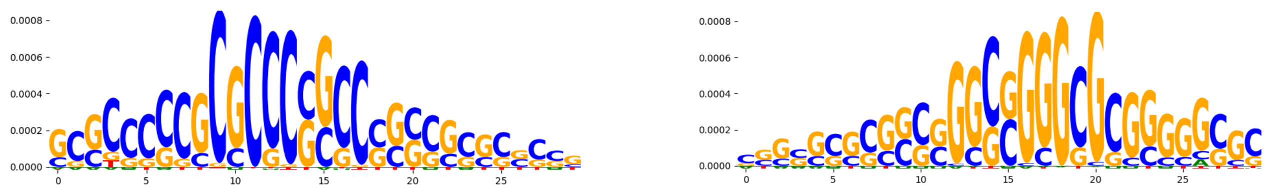 |
| pos_patterns.pattern_15 | 62 | 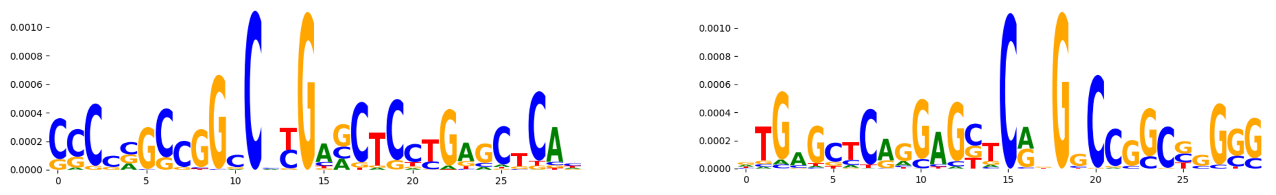 |
| pos_patterns.pattern_16 | 44 | 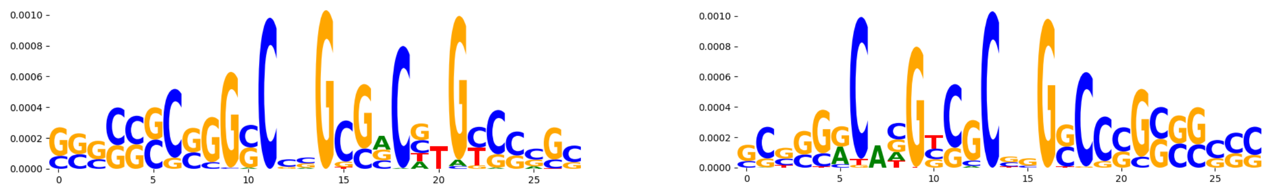 |
| pos_patterns.pattern_17 | 37 | 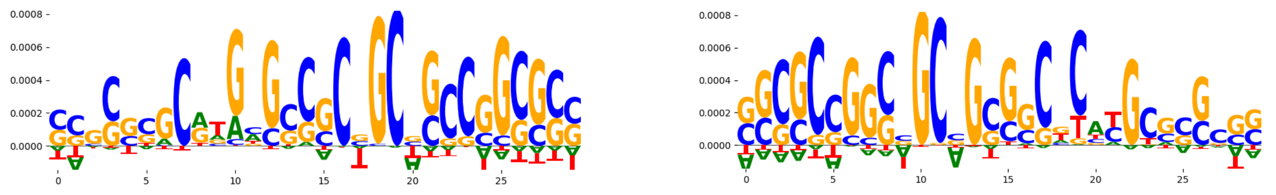 |
| pos_patterns.pattern_18 | 36 | 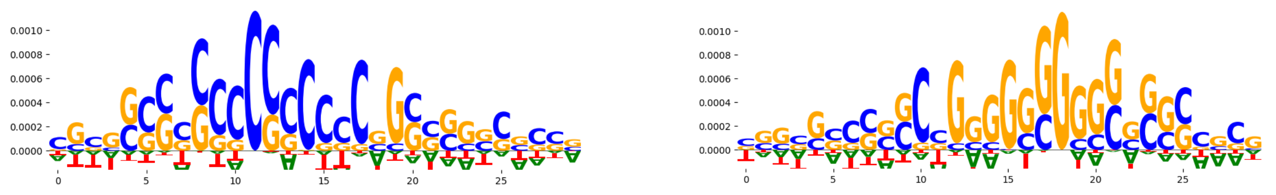 |
| pos_patterns.pattern_19 | 35 | 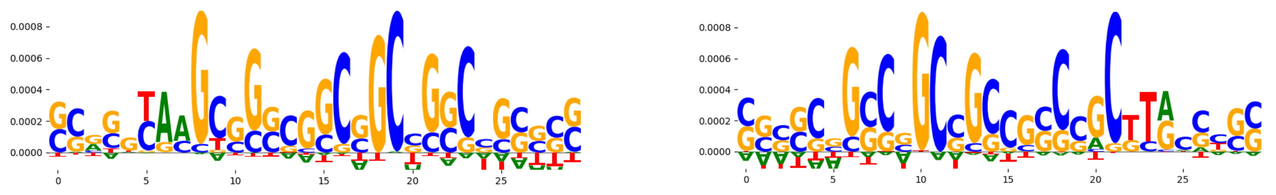 |
| pos_patterns.pattern_20 | 34 | 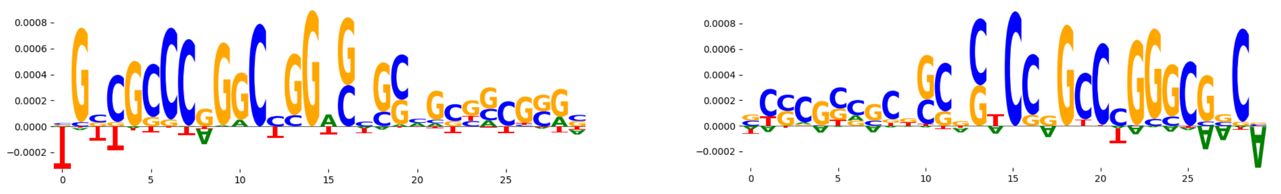 |
| pos_patterns.pattern_21 | 21 | 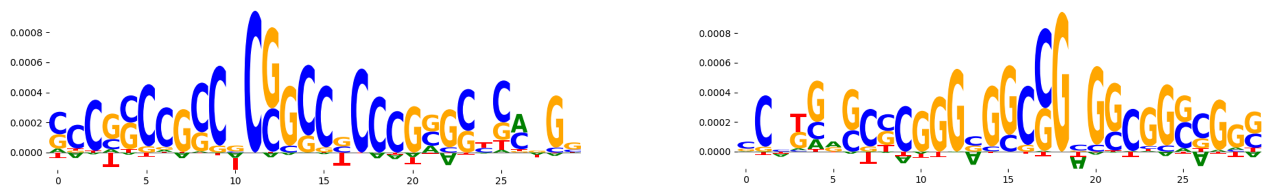 |
| neg_patterns.pattern_0 | 1635 | 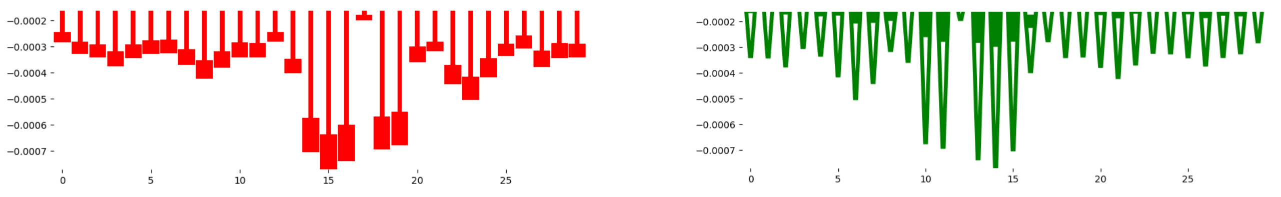 |

| pattern | num_seqlets | cwm_fwd | cwm_rev |
|---|---|---|---|
| neg_patterns.pattern_1 | 1616 | 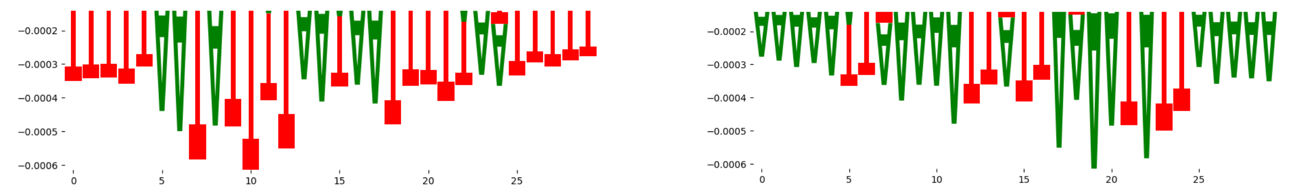 | |
| neg_patterns.pattern_2 | 986 | 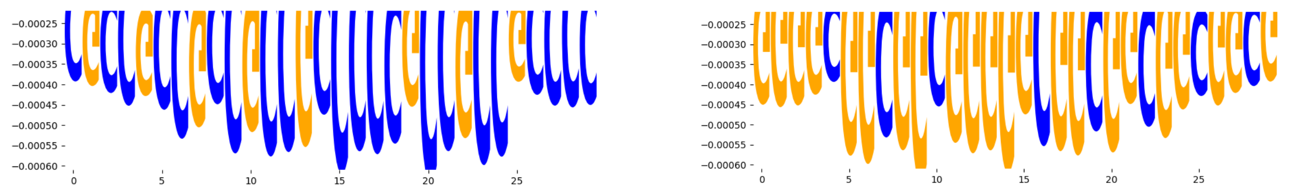 | |
| neg_patterns.pattern_3 | 393 | 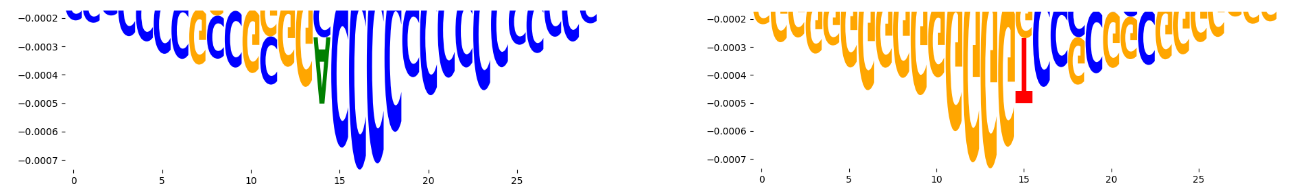 | |
| neg_patterns.pattern_4 | 287 | 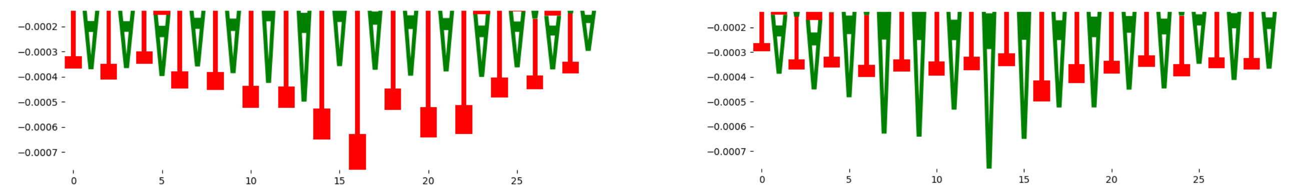 | |
| neg_patterns.pattern_5 | 279 | 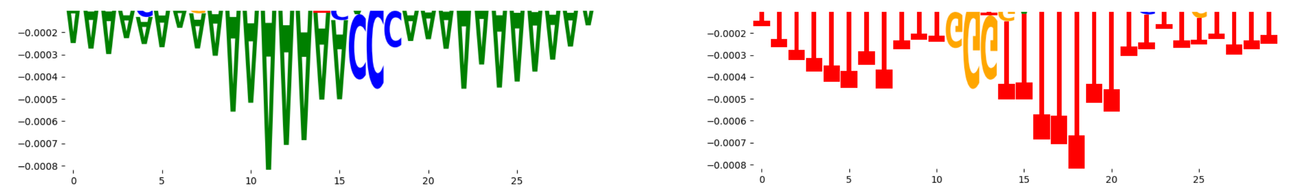 | |
| neg_patterns.pattern_6 | 279 | 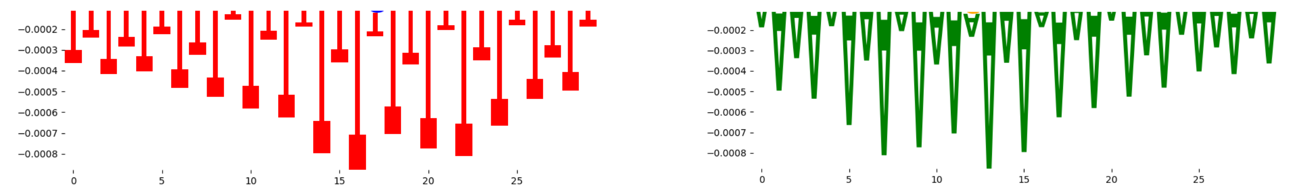 | |
| neg_patterns.pattern_7 | 257 | 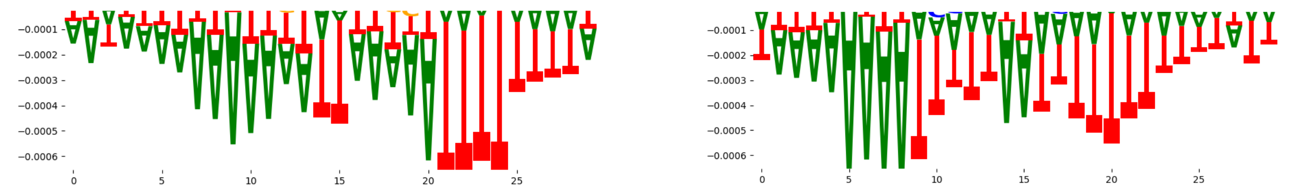 | |
| neg_patterns.pattern_8 | 233 | 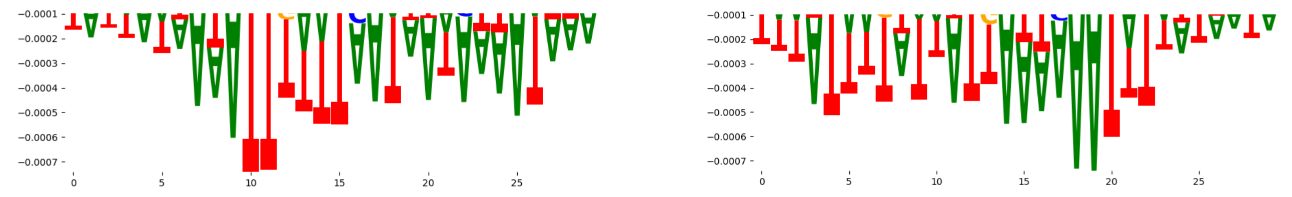 | |
| neg_patterns.pattern_9 | 229 | 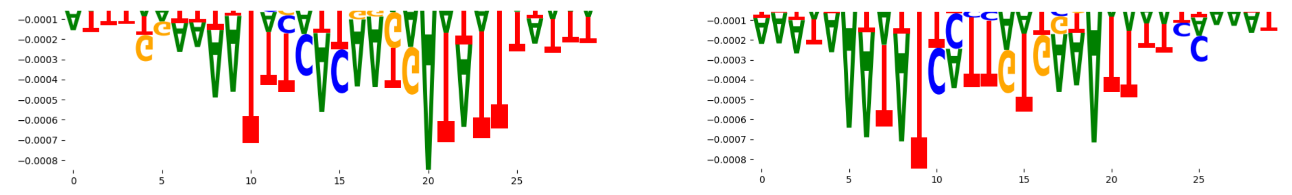 | |
| neg_patterns.pattern_10 | 227 | 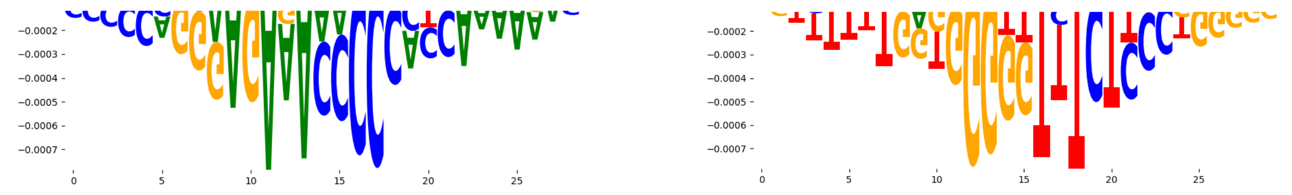 | |
| neg_patterns.pattern_11 | 214 | 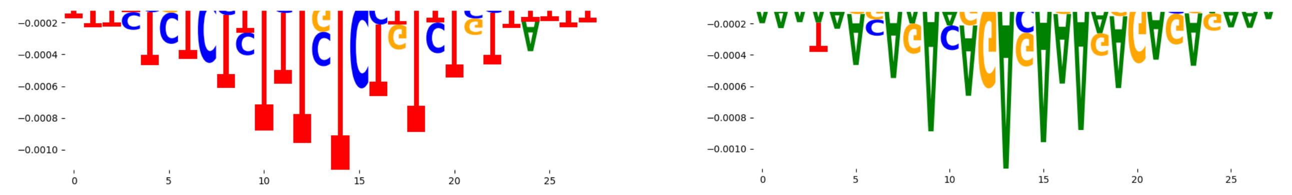 | |
| neg_patterns.pattern_12 | 177 | 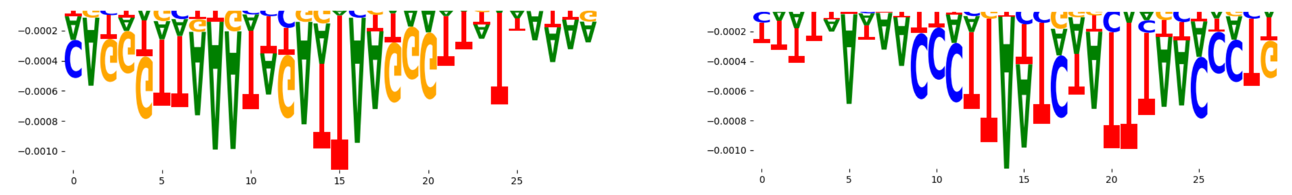 | |
| neg_patterns.pattern_13 | 168 | 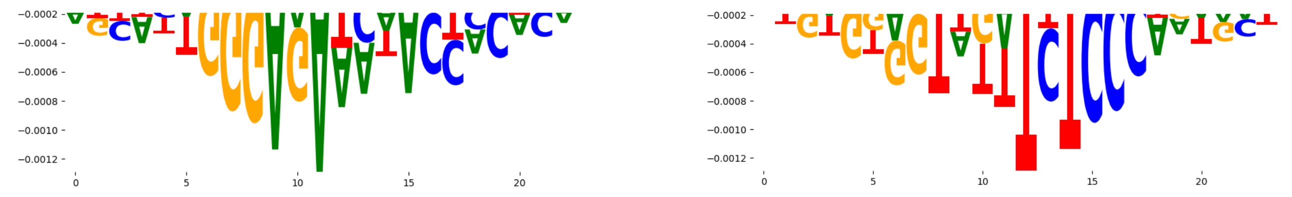 | |
| neg_patterns.pattern_14 | 164 | 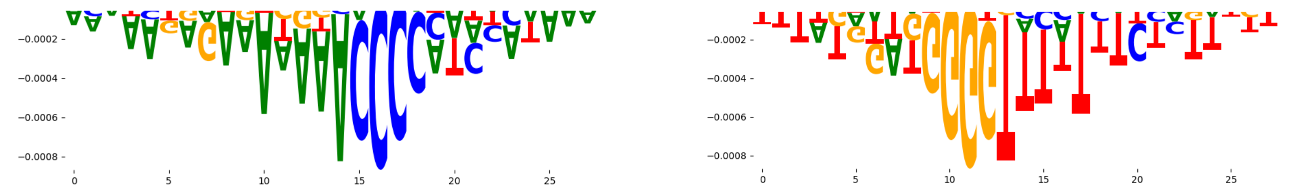 | |
| neg_patterns.pattern_15 | 156 | 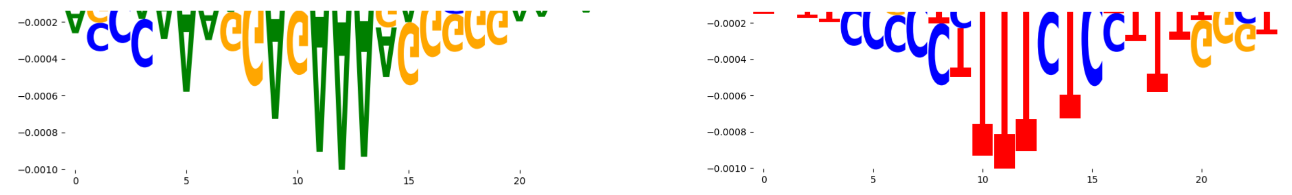 | |
| neg_patterns.pattern_16 | 129 | 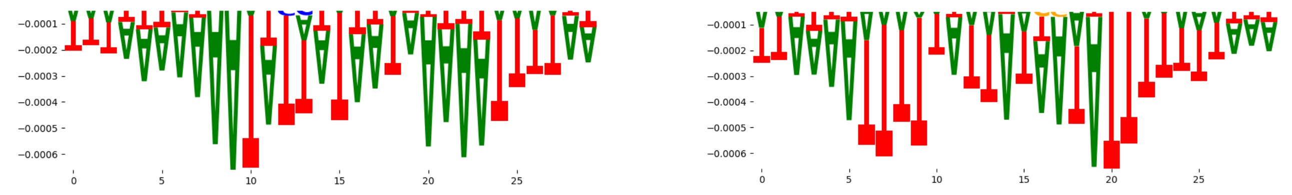 | |
| neg_patterns.pattern_17 | 122 | 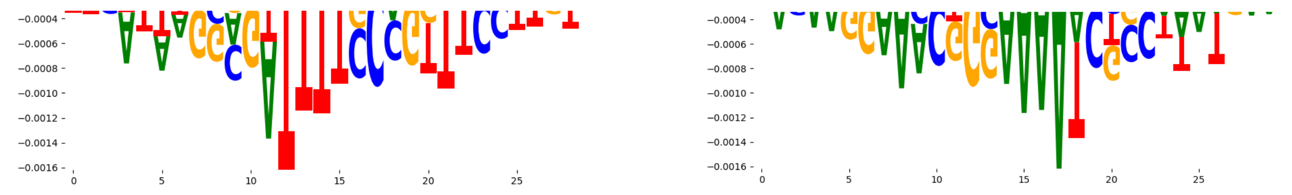 | |
| neg_patterns.pattern_18 | 117 | 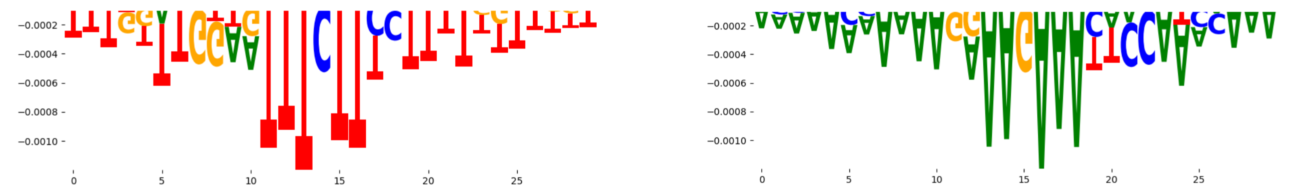 | |
| neg_patterns.pattern_19 | 117 | 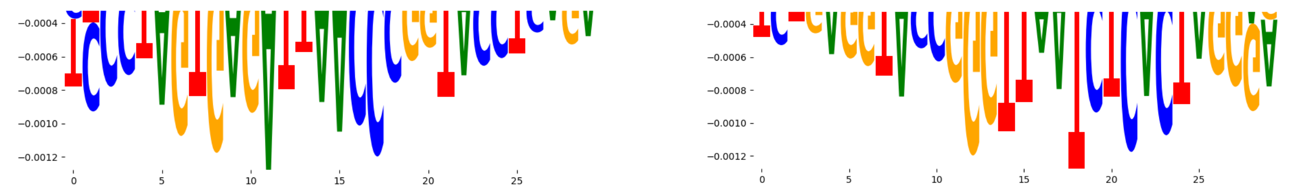 | |
| neg_patterns.pattern_20 | 113 | 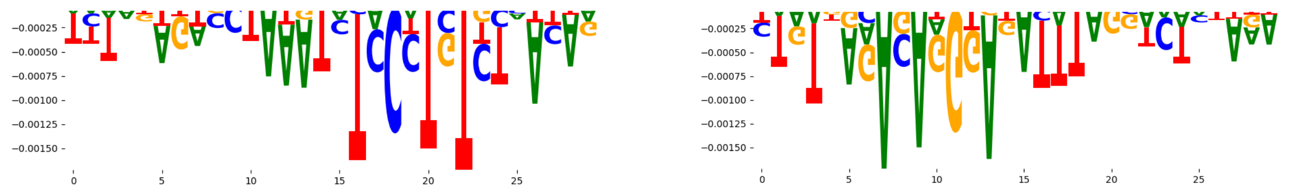 | |
| neg_patterns.pattern_21 | 113 | 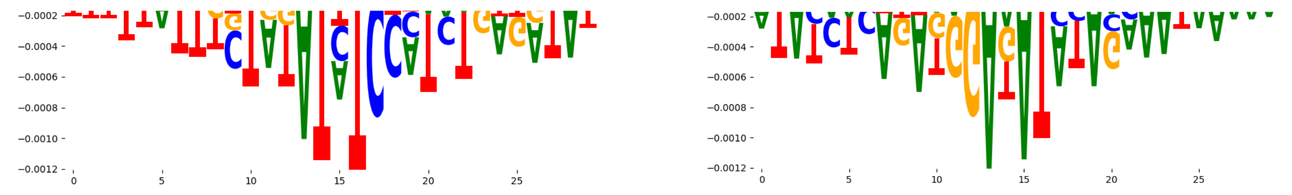 | |
| neg_patterns.pattern_22 | 95 | 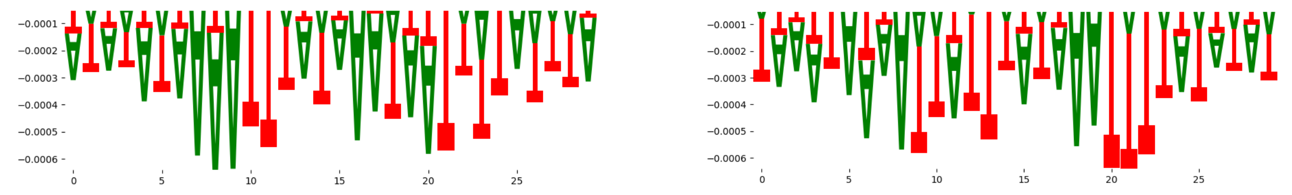 | |
| neg_patterns.pattern_23 | 93 | 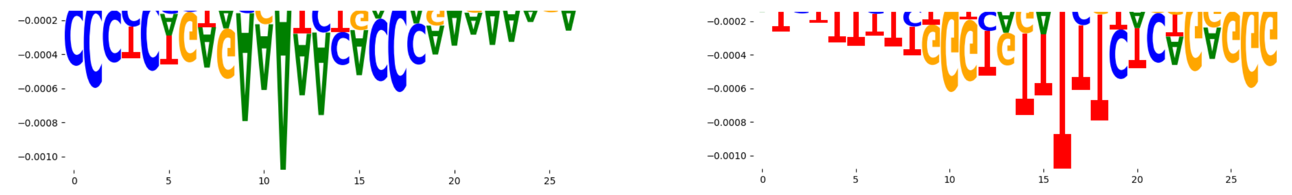 | |

| pattern | num_seqlets | cwm_fwd | cwm_rev |
|---|---|---|---|
| neg_patterns.pattern_24 | 81 | 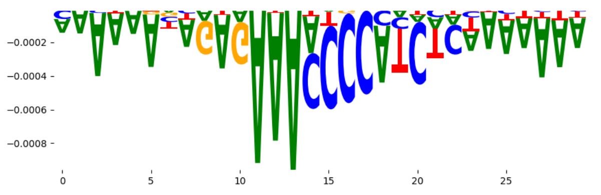 | 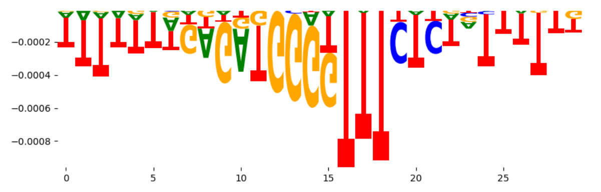 |
| neg_patterns.pattern_25 | 53 | 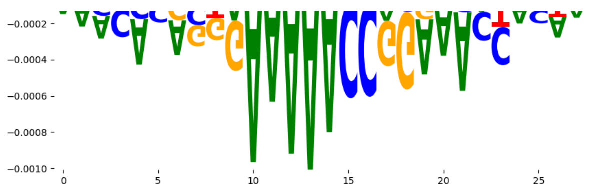 | 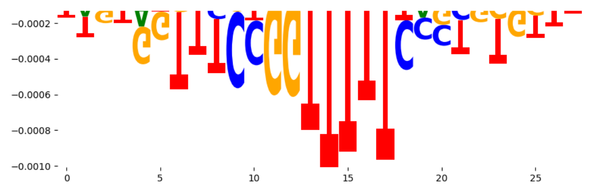 |
| neg_patterns.pattern_26 | 50 | 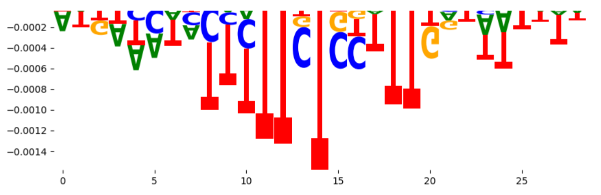 | 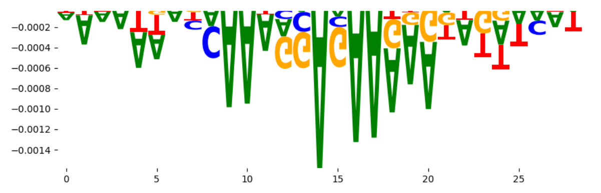 |
| neg_patterns.pattern_27 | 45 | 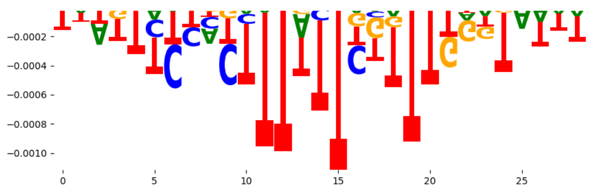 | 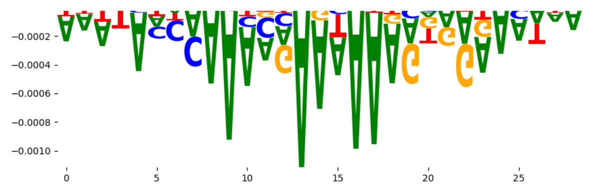 |
| neg_patterns.pattern_28 | 24 | 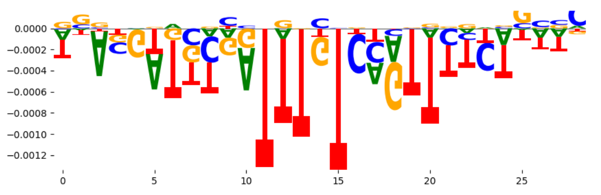 | 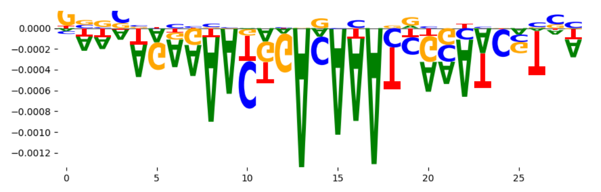 |
| neg_patterns.pattern_29 | 23 | 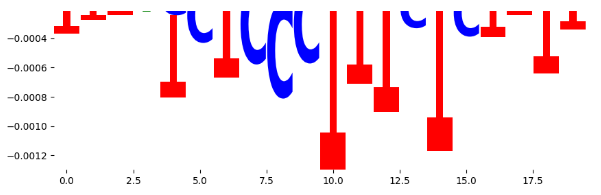 | 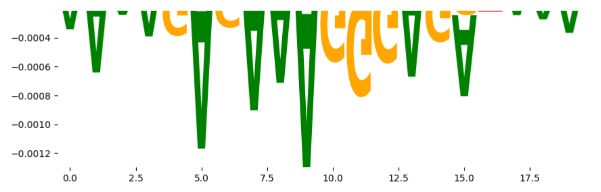 |
| neg_patterns.pattern_30 | 21 | 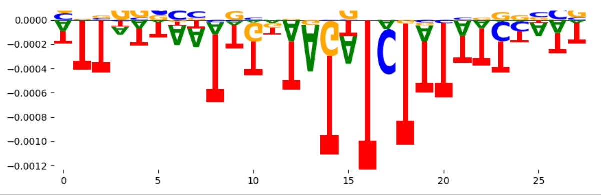 | 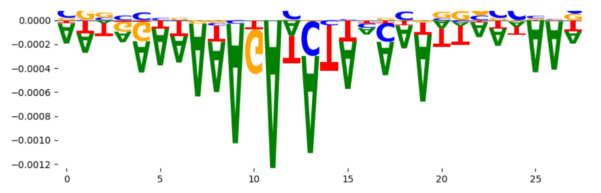 |
