## Supplementary Files 3 for "ChromBPNet: bias factorized, base-resolution deep learning models of chromatin accessibility reveal cis-regulatory sequence syntax, transcription factor footprints and regulatory variants": k562_DNASE_raw_bpnet_bias_fold2_profile_modisco.pdf

| pattern | num_seqlets | cwm_fwd | cwm_rev |
|---|---|---|---|
| pos_patterns.pattern_0 | 5585 | 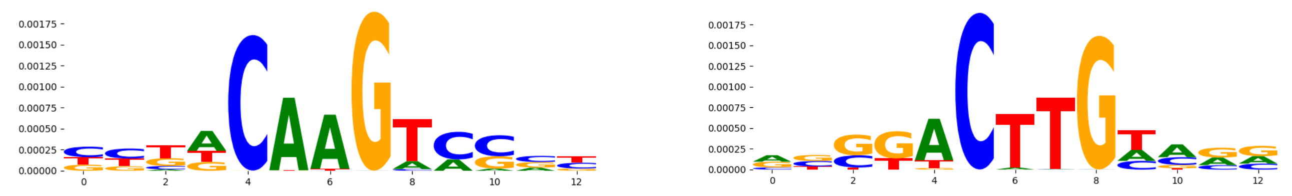 | |
| pos_patterns.pattern_1 | 3642 | 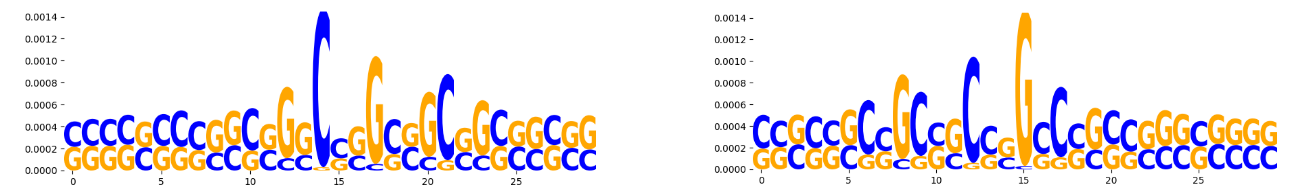 | |
| pos_patterns.pattern_2 | 2531 | 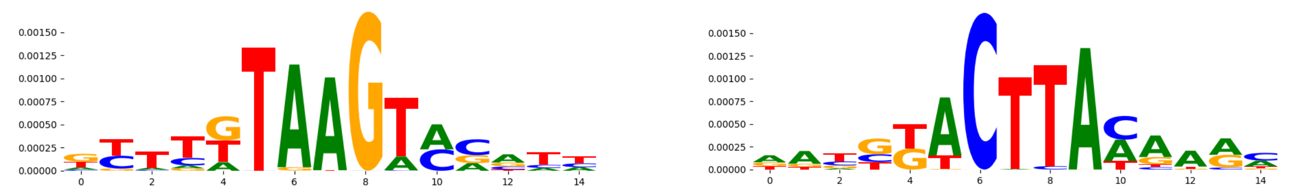 | |
| pos_patterns.pattern_3 | 1830 | 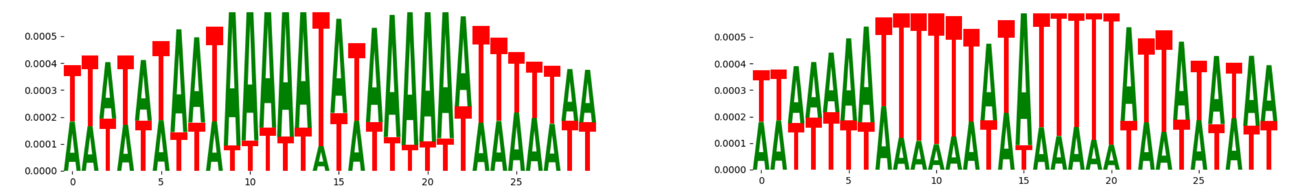 | |
| pos_patterns.pattern_4 | 959 | 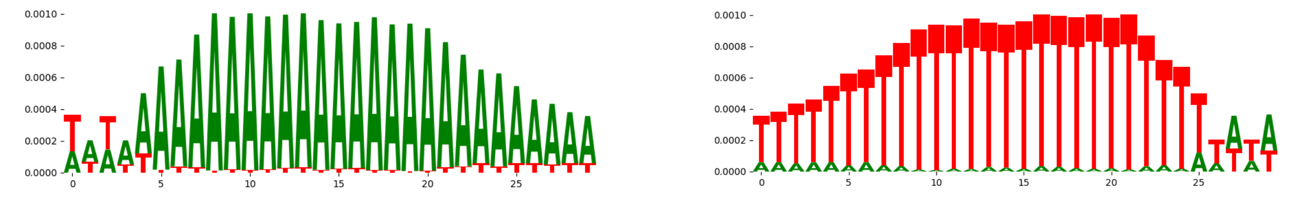 | |
| pos_patterns.pattern_5 | 934 | 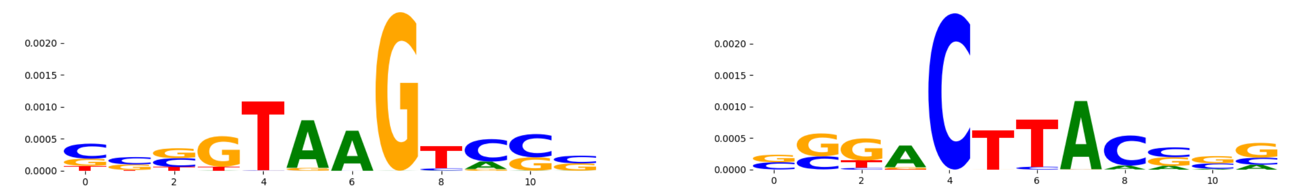 | |
| pos_patterns.pattern_6 | 875 | 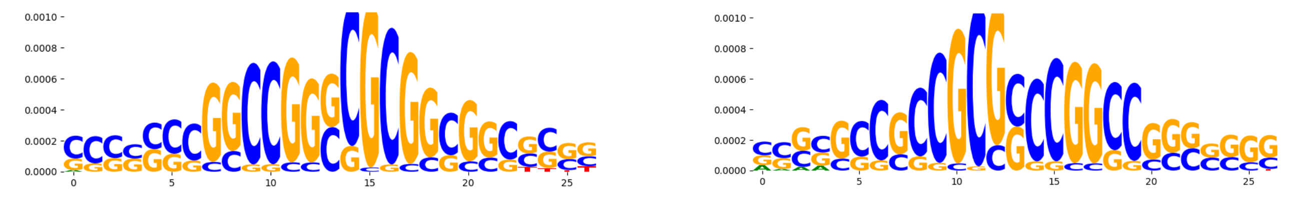 | |
| pos_patterns.pattern_7 | 787 | 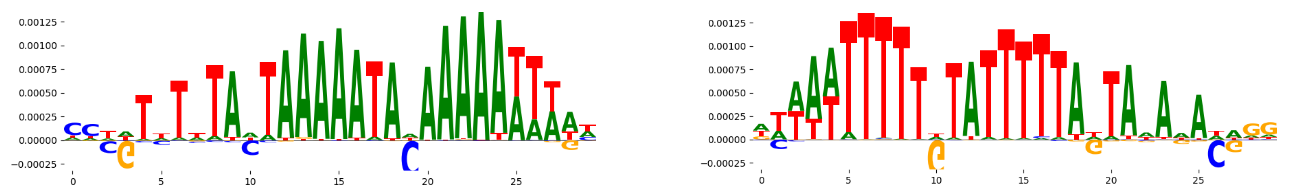 | |
| pos_patterns.pattern_8 | 422 | 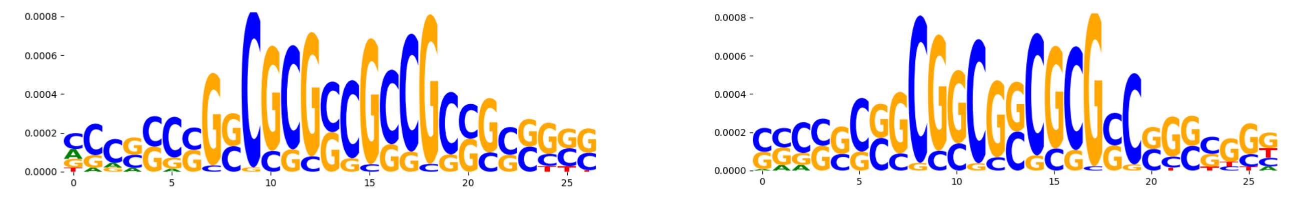 | |
| pos_patterns.pattern_9 | 317 | 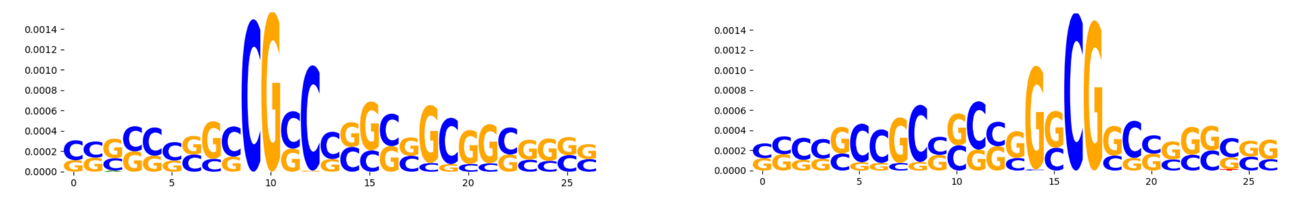 | |
| pos_patterns.pattern_10 | 316 | 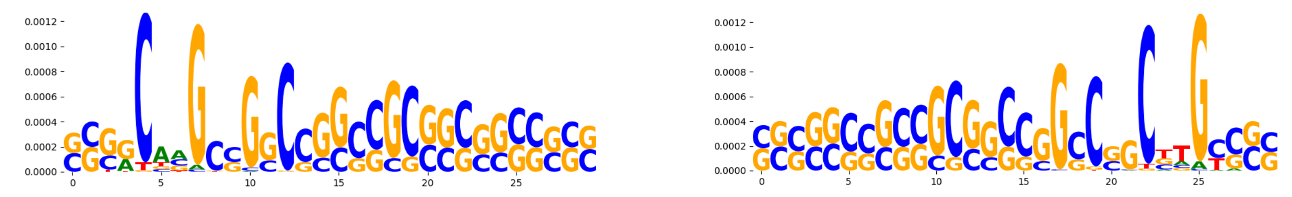 | |
| pos_patterns.pattern_11 | 240 | 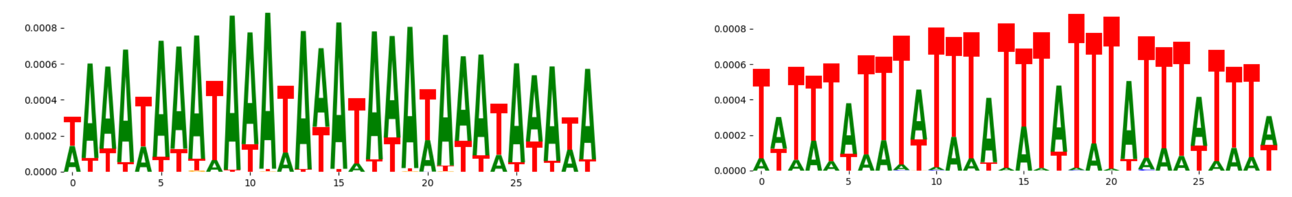 | |
| pos_patterns.pattern_12 | 211 | 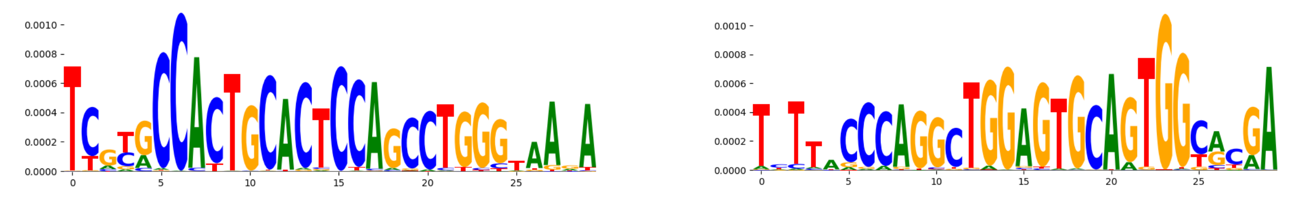 | |
| pos_patterns.pattern_13 | 192 | 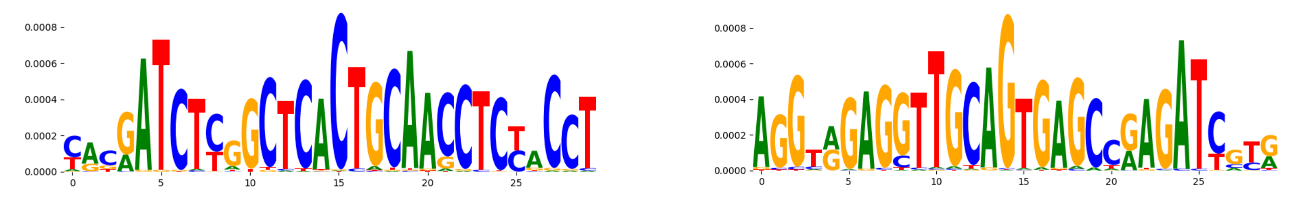 | |
| pos_patterns.pattern_14 | 183 | 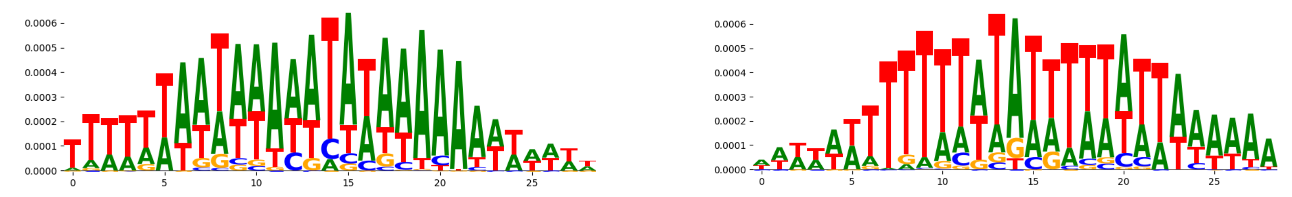 | |
| pos_patterns.pattern_15 | 177 | 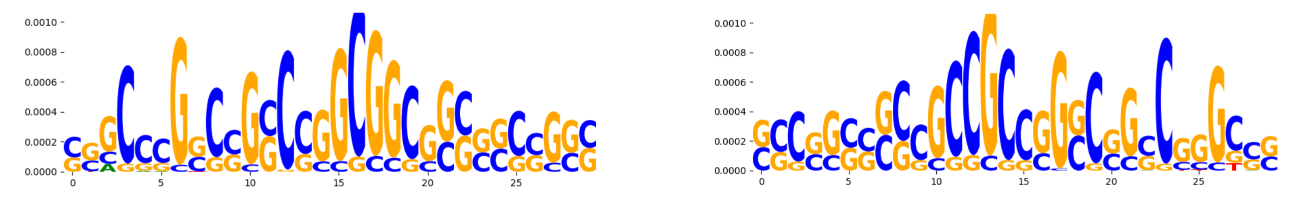 | |
| pos_patterns.pattern_16 | 166 | 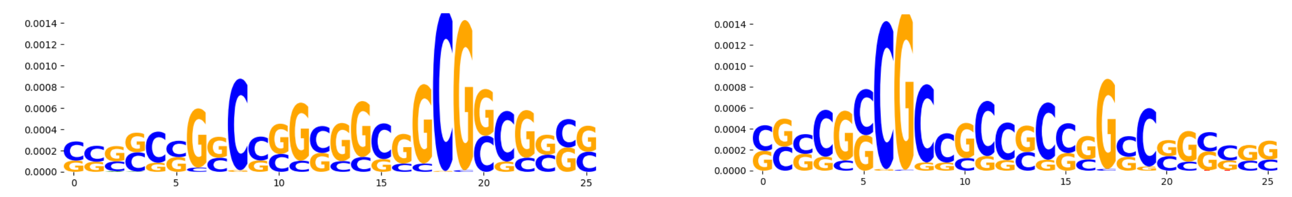 | |
| pos_patterns.pattern_17 | 165 | 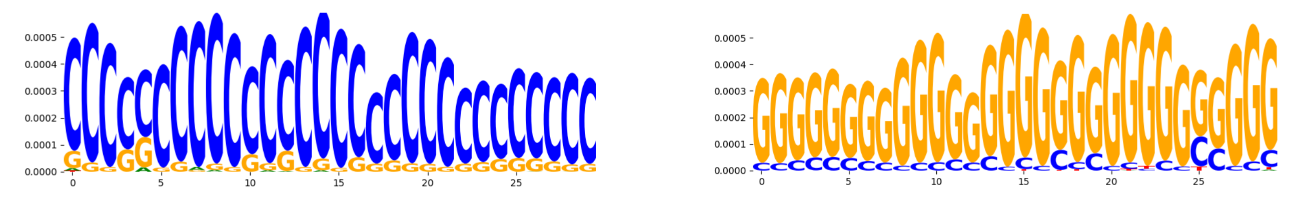 | |
| pos_patterns.pattern_18 | 135 | 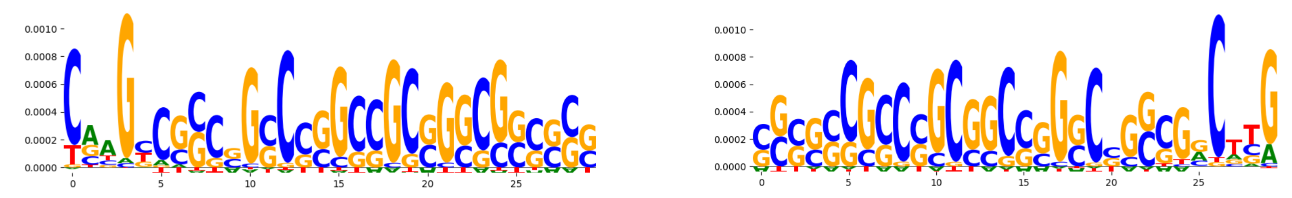 | |
| pos_patterns.pattern_19 | 117 | 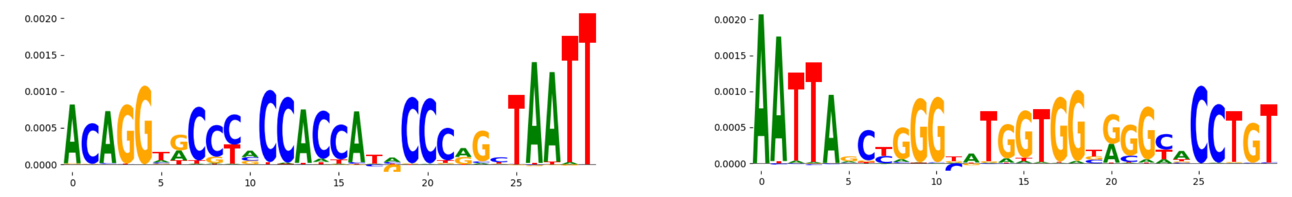 | |
| pos_patterns.pattern_20 | 100 | 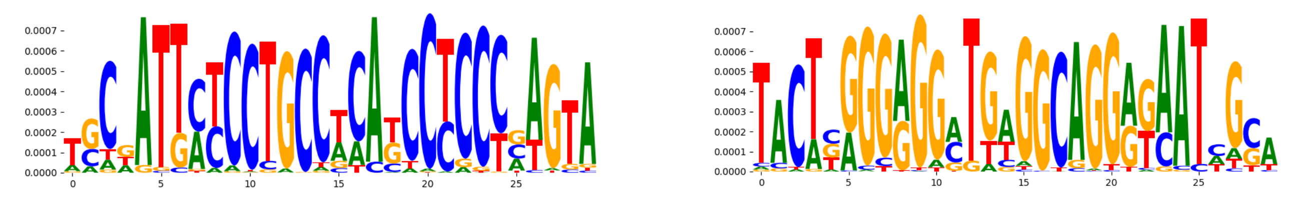 | |
| pos_patterns.pattern_21 | 81 | 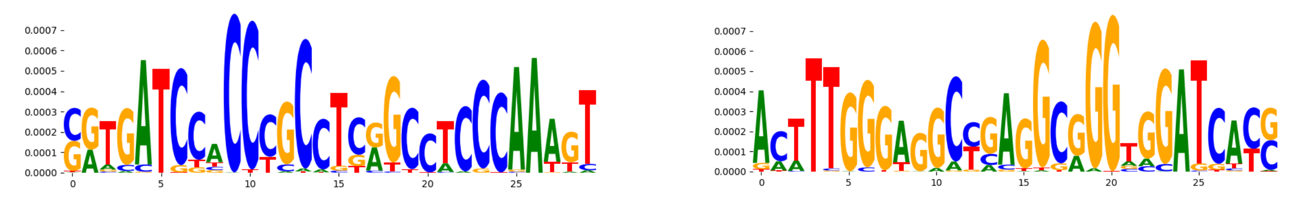 | |
| pos_patterns.pattern_22 | 70 | 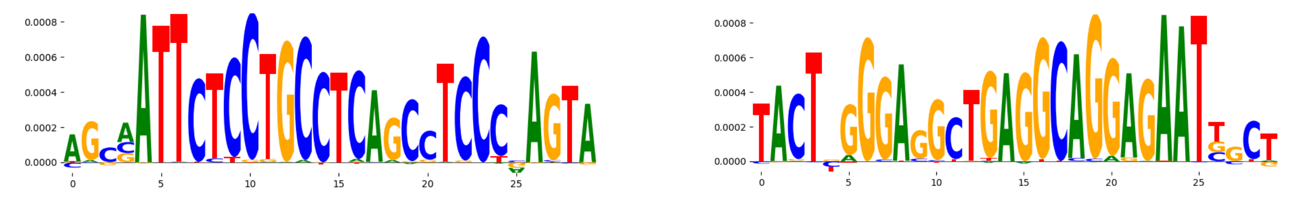 | |

| pattern | num_seqlets | cwm_fwd | cwm_rev |
|---|---|---|---|
| pos_patterns.pattern_23 | 64 | 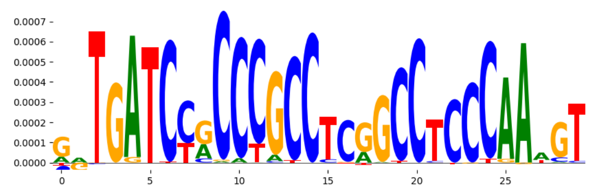 | 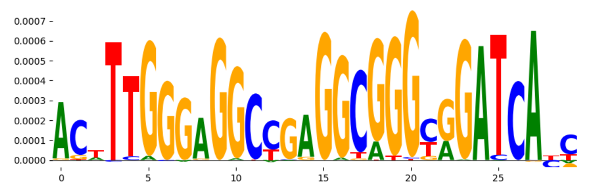 |
| pos_patterns.pattern_24 | 40 | 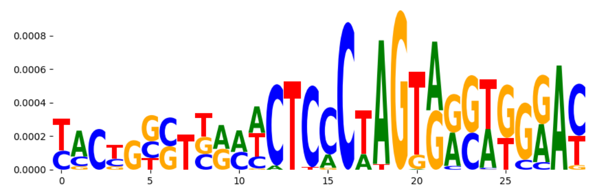 | 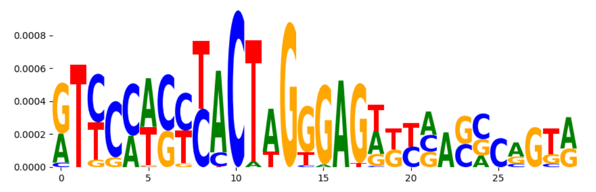 |
| pos_patterns.pattern_25 | 38 | 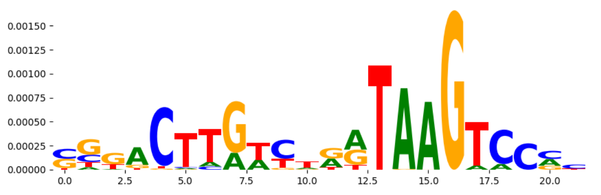 | 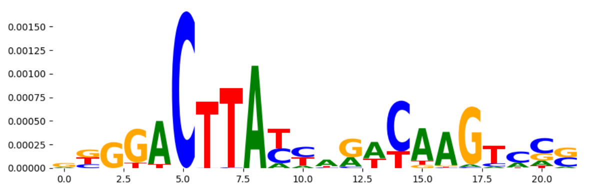 |
| pos_patterns.pattern_26 | 34 | 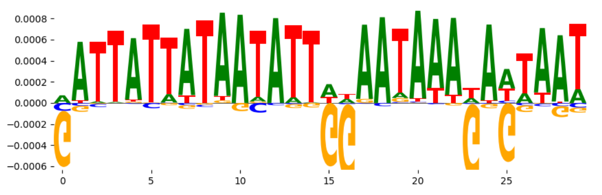 | 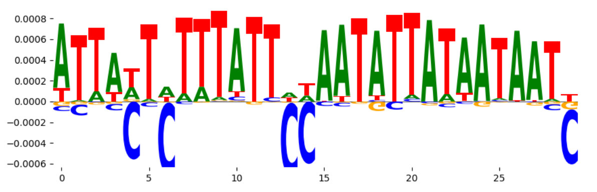 |
| pos_patterns.pattern_27 | 32 | 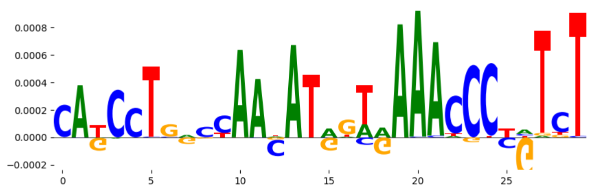 | 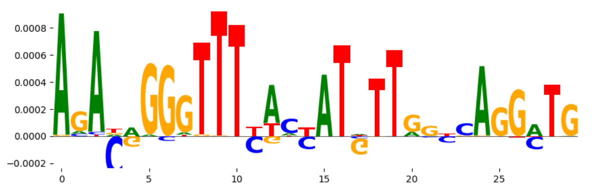 |
| pos_patterns.pattern_28 | 27 | 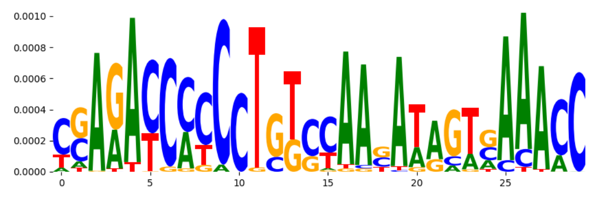 | 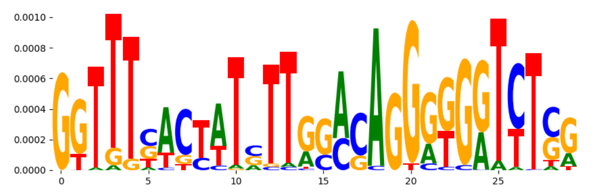 |
| neg_patterns.pattern_0 | 25 | 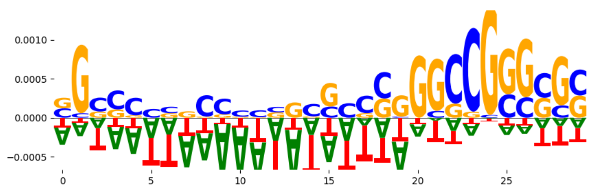 | 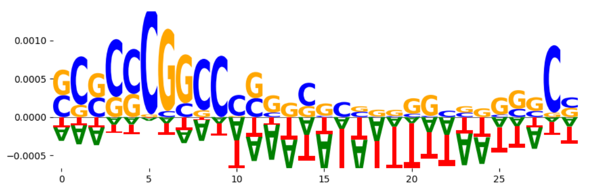 |
| neg_patterns.pattern_1 | 22 | 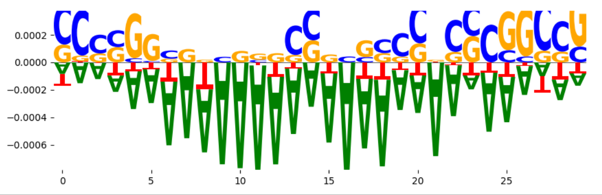 | 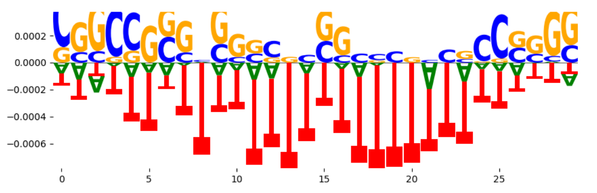 |
