## Supplementary Files 3 for "ChromBPNet: bias factorized, base-resolution deep learning models of chromatin accessibility reveal cis-regulatory sequence syntax, transcription factor footprints and regulatory variants": k562_DNASE_raw_bpnet_bias_fold3_counts_modisco.pdf

| pattern | num_seqlets | cwm_fwd | cwm_rev | TOMTOM_match | TOMTOM_qval | TOMTOM_match_logo |
|---|---|---|---|---|---|---|
| pos_patterns.pattern_0 | 3602 | | | ZNF384_MA1125.1 | 2.739470e-02 | |
| pos_patterns.pattern_1 | 2664 | | | SP1_MOUSE.H11MO.0.A | 1.288410e-10 | |
| pos_patterns.pattern_2 | 1720 | | | SP2_HUMAN.H11MO.0.A | 8.576680e-06 | |
| pos_patterns.pattern_3 | 722 | | | Foxj3_MA0851.1 | 1.000390e-01 | |
| pos_patterns.pattern_4 | 539 | | | SP1_HUMAN.H11MO.0.A | 4.485270e-02 | |
| pos_patterns.pattern_5 | 194 | | | SP2_HUMAN.H11MO.0.A | 1.525860e-02 | |
| neg_patterns.pattern_0 | 1725 | | | DNASE_2 | 6.793600e-03 | |
| neg_patterns.pattern_1 | 1316 | | | INSM1_MA0155.1 | 9.916480e-01 | |
| neg_patterns.pattern_2 | 574 | | | TFE2_HUMAN.H11MO.0.A | 1.000000e+00 | |
| neg_patterns.pattern_3 | 480 | | | DNASE_1 | 2.286400e-01 | |
| neg_patterns.pattern_4 | 350 | | | Hic1.mouse_C2H2_1 | 2.417530e-01 | |
| neg_patterns.pattern_5 | 263 | | | KLF4_MA0039.3 | 1.057400e-01 | |
| neg_patterns.pattern_6 | 241 | | | DNASE_2 | 7.352590e-01 | |
| neg_patterns.pattern_7 | 209 | | | ISL2_MA0914.1 | 9.226180e-01 | |
| neg_patterns.pattern_8 | 201 | | | ZN770_HUMAN.H11MO.0.C | 1.000000e+00 | |
| neg_patterns.pattern_9 | 173 | | | CLOCK_HUMAN.H11MO.0.C | 6.414540e-01 | |
| neg_patterns.pattern_10 | 128 | | | DNASE_5 | 4.153550e-05 | |
| neg_patterns.pattern_11 | 108 | | | DNASE_3 | 7.218670e-01 | |
| neg_patterns.pattern_12 | 97 | | | ZNF384_MA1125.1 | 8.771860e-02 | |
| neg_patterns.pattern_13 | 65 | | | TN5_6 | 4.132790e-15 | |
| neg_patterns.pattern_14 | 45 | | | PTF1A_HUMAN.H11MO.0.B | 2.151590e-01 | |
| neg_patterns.pattern_15 | 44 | | | PATZ1_HUMAN.H11MO.0.C | 2.255090e-02 | |
| neg_patterns.pattern_16 | 42 | | | Hoxd8_MA0910.1 | 1.000000e+00 | |
| neg_patterns.pattern_17 | 34 | | | DNASE_3 | 2.315990e-02 | |
