## Supplementary Files 3 for "ChromBPNet: bias factorized, base-resolution deep learning models of chromatin accessibility reveal cis-regulatory sequence syntax, transcription factor footprints and regulatory variants": k562_DNASE_raw_bpnet_bias_fold3_profile_modisco.pdf

| pattern | num_seqlets | cwm_fwd | cwm_rev | TOMTOM_match | TOMTOM_qval | TOMTOM_match_logo |
|---|---|---|---|---|---|---|
| pos_patterns.pattern_0 | 5468 | | | DNASE_1 | 1.596660e-01 | |
| pos_patterns.pattern_1 | 3011 | | | DNASE_2 | 5.999550e-01 | |
| pos_patterns.pattern_2 | 3000 | | | SMAD3_HUMAN.H11MO.0.B | 3.642250e-01 | |
| pos_patterns.pattern_3 | 1849 | | | ZNF384_MA1125.1 | 1.463050e-01 | |
| pos_patterns.pattern_4 | 1083 | | | CTCFL_HUMAN.H11MO.0.A | 2.250680e-01 | |
| pos_patterns.pattern_5 | 1035 | | | PRDM6_HUMAN.H11MO.0.C | 8.316880e-02 | |
| pos_patterns.pattern_6 | 989 | | | CTCFL_MOUSE.H11MO.0.A | 6.020230e-05 | |
| pos_patterns.pattern_7 | 917 | | | TYY1_HUMAN.H11MO.0.A | 1.500790e-01 | |
| pos_patterns.pattern_8 | 744 | | | MEF2B_HUMAN.H11MO.0.A | 2.134900e-01 | |
| pos_patterns.pattern_9 | 393 | | | CTCFL_HUMAN.H11MO.0.A | 2.425830e-01 | |
| pos_patterns.pattern_10 | 275 | | | SP2_HUMAN.H11MO.0.A | 1.462600e-09 | |
| pos_patterns.pattern_11 | 220 | | | ZN257_HUMAN.H11MO.0.C | 1.000000e+00 | |
| pos_patterns.pattern_12 | 192 | | | RARG_HUMAN.H11MO.0.B | 1.000000e+00 | |
| pos_patterns.pattern_13 | 175 | | | DNASE_5 | 2.742280e-03 | |
| pos_patterns.pattern_14 | 164 | | | AP2B_HUMAN.H11MO.0.B | 1.555480e-01 | |
| pos_patterns.pattern_15 | 138 | | | ITF2_HUMAN.H11MO.0.C | 1.000000e+00 | |
| pos_patterns.pattern_16 | 138 | | | CPEB1_RRM_1 | 3.223090e-01 | |
| pos_patterns.pattern_17 | 134 | | | CTCFL_HUMAN.H11MO.0.A | 6.909920e-01 | |
| pos_patterns.pattern_18 | 70 | | | RARA_HUMAN.H11MO.0.A | 9.558640e-02 | |
| pos_patterns.pattern_19 | 67 | | | ZN250_HUMAN.H11MO.0.C | 1.000000e+00 | |
| pos_patterns.pattern_20 | 62 | | | ZN770_HUMAN.H11MO.0.C | 1.242120e-04 | |
| pos_patterns.pattern_21 | 48 | | | FOXB1_forkhead_2 | 1.000000e+00 | |
| pos_patterns.pattern_22 | 31 | | | DNASE_6 | 7.491160e-02 | |
| neg_patterns.pattern_0 | 32 | | | CPEB1_RRM_1 | 3.004510e-01 | |
