## Supplementary Files 3 for "ChromBPNet: bias factorized, base-resolution deep learning models of chromatin accessibility reveal cis-regulatory sequence syntax, transcription factor footprints and regulatory variants": k562_DNASE_raw_bpnet_bias_fold4_counts_modisco.pdf

| pattern | num_seqlets | cwm_fwd | cwm_rev | TOMTOM_match | TOMTOM_qval | TOMTOM_match_logo |
|---|---|---|---|---|---|---|
| pos_patterns.pattern_0 | 3985 | | | ZNF384_MA1125.1 | 1.917260e-01 | |
| pos_patterns.pattern_1 | 3129 | | | SP2_HUMAN.H11MO.0.A | 4.680670e-11 | |
| pos_patterns.pattern_2 | 1435 | | | SP2_HUMAN.H11MO.0.A | 3.113130e-06 | |
| pos_patterns.pattern_3 | 350 | | | ZNF740_C2H2_2 | 4.076140e-02 | |
| pos_patterns.pattern_4 | 322 | | | PRDM6_HUMAN.H11MO.0.C | 1.000000e+00 | |
| pos_patterns.pattern_5 | 122 | | | ZN770_HUMAN.H11MO.0.C | 4.871910e-06 | |
| pos_patterns.pattern_6 | 67 | | | NFAT5_NFAT_1 | 5.698780e-01 | |
| pos_patterns.pattern_7 | 62 | | | CPEB1_RRM_1 | 1.000000e+00 | |
| neg_patterns.pattern_0 | 1953 | | | MAFK_bZIP_1 | 1.000000e+00 | |
| neg_patterns.pattern_1 | 1683 | | | MYB_HUMAN.H11MO.0.A | 1.000000e+00 | |
| neg_patterns.pattern_2 | 1241 | | | ZFX_MOUSE.H11MO.0.B | 1.111810e-01 | |
| neg_patterns.pattern_3 | 469 | | | CTCF_MA0139.1 | 1.450810e-02 | |
| neg_patterns.pattern_4 | 359 | | | DNASE_2 | 2.738920e-03 | |
| neg_patterns.pattern_5 | 239 | | | NR2C1_HUMAN.H11MO.0.C | 5.474930e-01 | |
| neg_patterns.pattern_6 | 137 | | | KLF5_MOUSE.H11MO.0.A | 3.593870e-03 | |
| neg_patterns.pattern_7 | 109 | | | EGR2_HUMAN.H11MO.0.A | 6.224870e-01 | |
| neg_patterns.pattern_8 | 107 | | | GATA1_HUMAN.H11MO.0.A | 7.771510e-03 | |
| neg_patterns.pattern_9 | 105 | | | EGR2_MOUSE.H11MO.0.A | 6.844440e-03 | |
| neg_patterns.pattern_10 | 102 | | | SALL4_HUMAN.H11MO.0.B | 4.211610e-03 | |
| neg_patterns.pattern_11 | 89 | | | SALL4_HUMAN.H11MO.0.B | 4.561280e-01 | |
| neg_patterns.pattern_12 | 77 | | | ZN281_HUMAN.H11MO.0.A | 2.109040e-02 | |
| neg_patterns.pattern_13 | 69 | | | SP8_C2H2_1 | 6.087330e-01 | |
| neg_patterns.pattern_14 | 31 | | | KLF4_MOUSE.H11MO.0.A | 1.090060e-02 | |
| neg_patterns.pattern_15 | 27 | | | CTCFL_HUMAN.H11MO.0.A | 7.146070e-02 | |
| neg_patterns.pattern_16 | 21 | | | FOS_HUMAN.H11MO.0.A | 2.965190e-01 | |
