## Supplementary Files 3 for "ChromBPNet: bias factorized, base-resolution deep learning models of chromatin accessibility reveal cis-regulatory sequence syntax, transcription factor footprints and regulatory variants": k562_DNASE_raw_bpnet_bias_fold4_profile_modisco.pdf

| pattern | num_seqlets | cwm_fwd | cwm_rev | TOMTOM_match | TOMTOM_qval | TOMTOM_match_logo |
|---|---|---|---|---|---|---|
| pos_patterns.pattern_0 | 6193 | | | PRDM9_MOUSE.H11MO.0.C | 1.716240e-01 | |
| pos_patterns.pattern_1 | 4253 | | | ERR2_HUMAN.H11MO.0.A | 1.000000e+00 | |
| pos_patterns.pattern_2 | 2514 | | | DNASE_2 | 6.439080e-04 | |
| pos_patterns.pattern_3 | 1772 | | | FOXJ3_HUMAN.H11MO.0.A | 2.769370e-01 | |
| pos_patterns.pattern_4 | 905 | | | ZNF384_MA1125.1 | 9.670030e-02 | |
| pos_patterns.pattern_5 | 899 | | | TYY1_HUMAN.H11MO.0.A | 5.954010e-02 | |
| pos_patterns.pattern_6 | 835 | | | Zfx_MA0146.2 | 5.402300e-01 | |
| pos_patterns.pattern_7 | 789 | | | MEF2B_HUMAN.H11MO.0.A | 2.086250e-01 | |
| pos_patterns.pattern_8 | 706 | | | DNASE_1 | 4.744360e-01 | |
| pos_patterns.pattern_9 | 237 | | | TN5_6 | 1.480540e-20 | |
| pos_patterns.pattern_10 | 204 | | | SP1_HUMAN.H11MO.0.A | 9.313590e-08 | |
| pos_patterns.pattern_11 | 198 | | | SP1_MOUSE.H11MO.0.A | 7.499990e-01 | |
| pos_patterns.pattern_12 | 162 | | | ZN770_HUMAN.H11MO.0.C | 4.338470e-09 | |
| pos_patterns.pattern_13 | 133 | | | FOXC1_forkhead_1 | 6.003630e-02 | |
| pos_patterns.pattern_14 | 131 | | | ZN770_HUMAN.H11MO.0.C | 4.245640e-08 | |
| pos_patterns.pattern_15 | 113 | | | KLF4_MA0039.3 | 6.618320e-01 | |
| pos_patterns.pattern_16 | 111 | | | Arid5a_MA0602.1 | 1.000000e+00 | |
| pos_patterns.pattern_17 | 90 | | | IRF3_HUMAN.H11MO.0.B | 5.476980e-02 | |
| pos_patterns.pattern_18 | 81 | | | ZN770_HUMAN.H11MO.0.C | 6.396090e-08 | |
| pos_patterns.pattern_19 | 64 | | | NR4A2_nuclearreceptor_1 | 6.307350e-01 | |
| pos_patterns.pattern_20 | 62 | | | ZN121_HUMAN.H11MO.0.C | 6.004250e-06 | |
| pos_patterns.pattern_21 | 61 | | | NKX31_HUMAN.H11MO.0.C | 9.798240e-01 | |
| pos_patterns.pattern_22 | 36 | | | ZN121_HUMAN.H11MO.0.C | 1.877100e-02 | |
| pos_patterns.pattern_23 | 34 | | | ITF2_HUMAN.H11MO.0.C | 1.000000e+00 | |
| pos_patterns.pattern_24 | 27 | | | MAFK_bZIP_3 | 6.518160e-01 | |
| pos_patterns.pattern_25 | 24 | | | ZN770_HUMAN.H11MO.0.C | 1.622820e-03 | |
| pos_patterns.pattern_26 | 21 | | | RARA_HUMAN.H11MO.0.A | 7.818600e-01 | |
| neg_patterns.pattern_0 | 50 | | | NF2L2_MOUSE.H11MO.0.A | 4.423890e-01 | |
| neg_patterns.pattern_1 | 47 | | | CPEB1_RRM_1 | 7.855680e-01 | |

| pattern | num_seqlets | cwm_fwd | cwm_rev | TOMTOM_match | TOMTOM_qval | TOMTOM_match_logo |
|---|---|---|---|---|---|---|
| neg_patterns.pattern_2 | 45 | 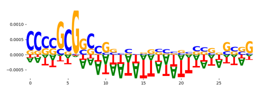 | 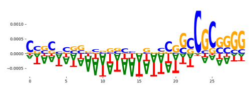 | ZNF384_MA1125.1 | 6.280170e-01 | 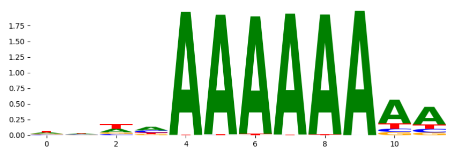 |
| neg_patterns.pattern_3 | 42 | 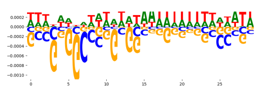 | 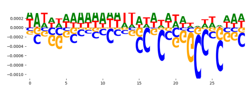 | NFATC1_NFAT_1 | 7.833110e-01 | 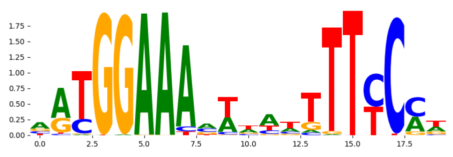 |
| neg_patterns.pattern_4 | 42 | 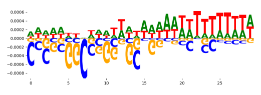 | 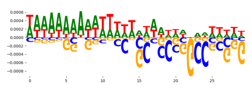 | ZNF384_MA1125.1 | 8.092630e-01 | 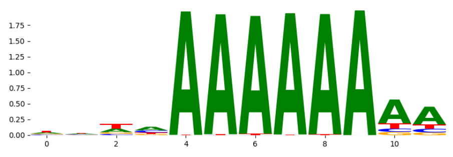 |
| neg_patterns.pattern_5 | 34 | 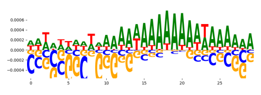 | 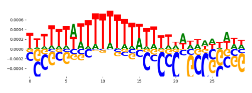 | ZNF384_MA1125.1 | 8.334500e-02 | 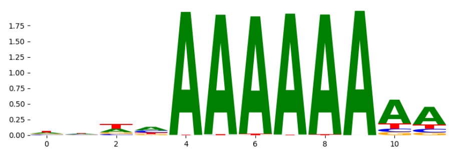 |
| neg_patterns.pattern_6 | 30 | 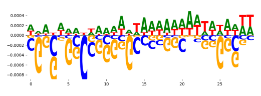 | 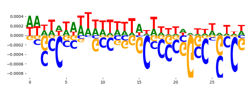 | HOXC11_homeodomain_2 | 2.152600e-01 | 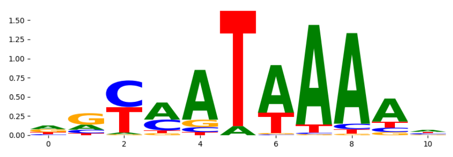 |
| neg_patterns.pattern_7 | 27 | 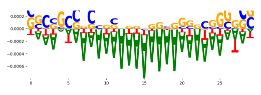 | 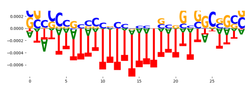 | PRDM6_HUMAN.H11MO.0.C | 3.082330e-02 | 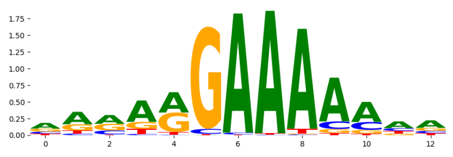 |
| neg_patterns.pattern_8 | 25 | 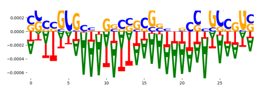 | 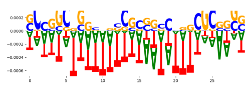 | ZNF384_MA1125.1 | 3.081730e-01 | 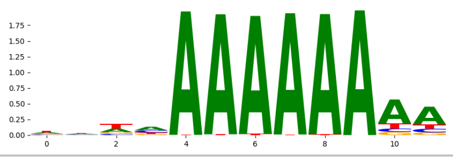 |
