## Supplementary Files 4 for "ChromBPNet: bias factorized, base-resolution deep learning models of chromatin accessibility reveal cis-regulatory sequence syntax, transcription factor footprints and regulatory variants": fig4a_naked_dna_ATAC_raw_bias_model_counts_modisco.pdf

| pattern | num_seqlets | cwm_fwd | cwm_rev | TOMTOM_match | TOMTOM_qval | TOMTOM_match_logo |
|---|---|---|---|---|---|---|
| pos_patterns.pattern_0 | 2702 | 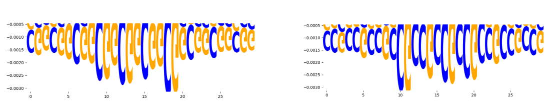 | | SP2_HUMAN.H11MO.0.A | 3.291210e-05 | 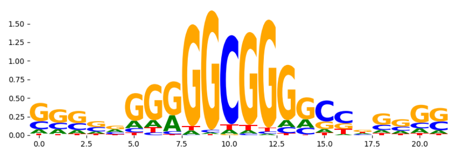 |
| pos_patterns.pattern_1 | 2581 | 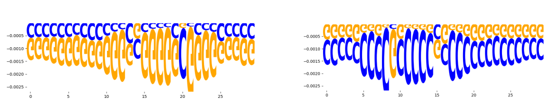 | | SP2_HUMAN.H11MO.0.A | 5.786870e-11 | 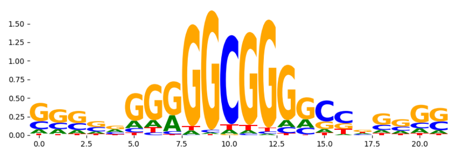 |
| pos_patterns.pattern_2 | 1966 | 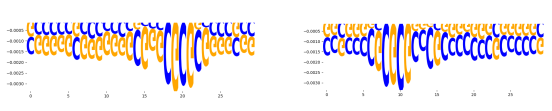 | | SP2_HUMAN.H11MO.0.A | 7.391890e-07 | 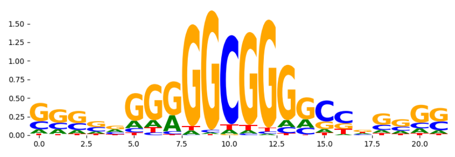 |
| pos_patterns.pattern_3 | 1501 | 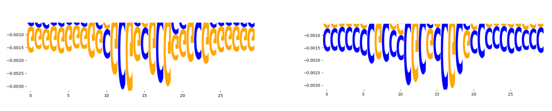 | | SP2_HUMAN.H11MO.0.A | 1.611420e-07 | 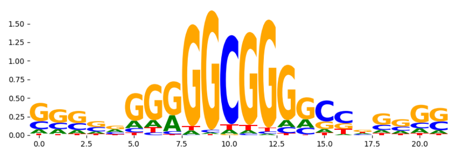 |
| pos_patterns.pattern_4 | 948 | 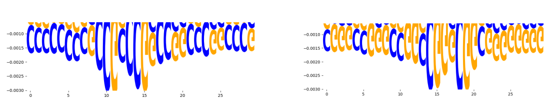 | | SP2_HUMAN.H11MO.0.A | 5.495660e-07 | 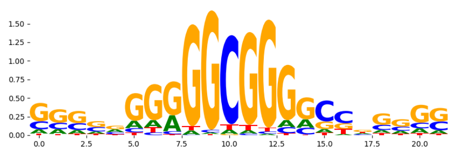 |
| neg_patterns.pattern_0 | 489 | 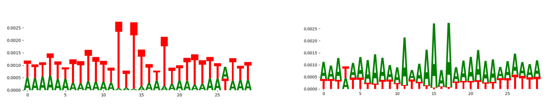 | | ELF5_MOUSE.H11MO.0.A | 7.347790e-04 | 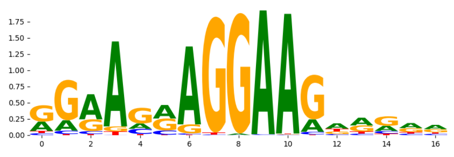 |
| neg_patterns.pattern_1 | 203 | 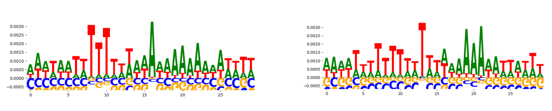 | | ZBT17_MOUSE.H11MO.0.A | 1.915460e-02 | 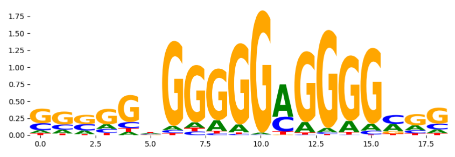 |
| neg_patterns.pattern_2 | 129 | 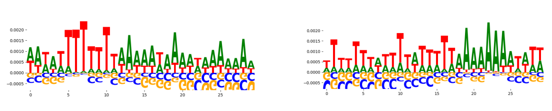 | | TAF1_HUMAN.H11MO.0.A | 3.322220e-01 | 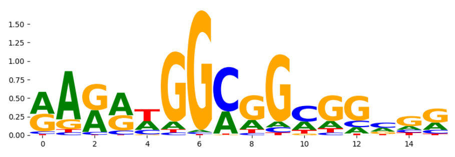 |
| neg_patterns.pattern_3 | 124 | 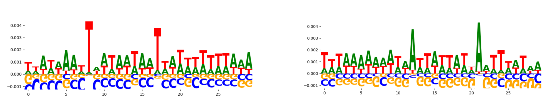 | | SP2_HUMAN.H11MO.0.A | 6.445520e-06 | 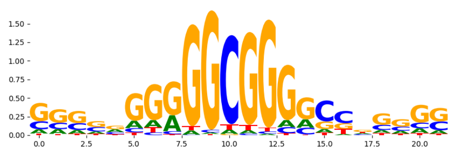 |
| neg_patterns.pattern_4 | 92 | 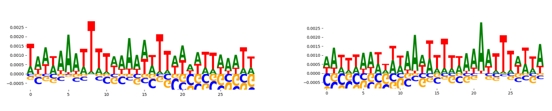 | | Zfx_MA0146.2 | 7.398490e-02 | 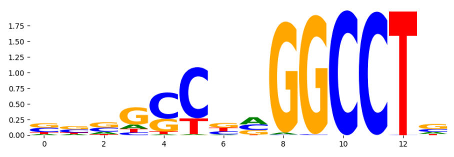 |
| neg_patterns.pattern_5 | 69 | 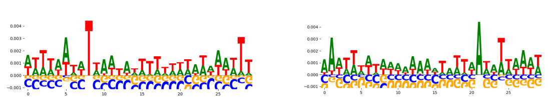 | | SP1_HUMAN.H11MO.0.A | 1.091220e-05 | 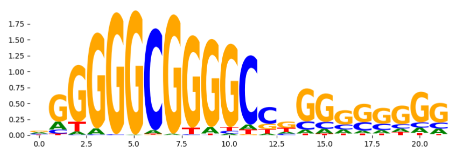 |
| neg_patterns.pattern_6 | 65 | 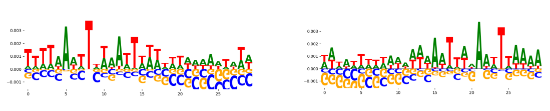 | | SP1_HUMAN.H11MO.0.A | 1.721450e-04 | 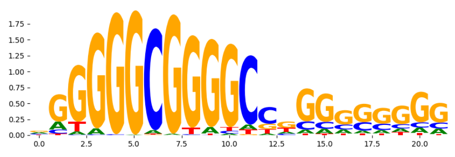 |
| neg_patterns.pattern_7 | 64 | 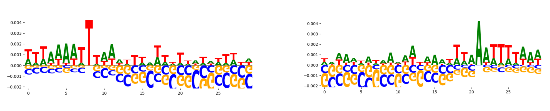 | | SP2_HUMAN.H11MO.0.A | 7.595640e-05 | 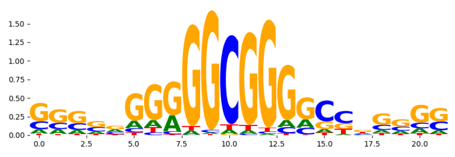 |
| neg_patterns.pattern_8 | 64 | 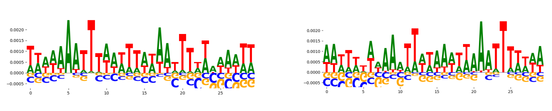 | | BHA15_HUMAN.H11MO.0.B | 3.930840e-01 | 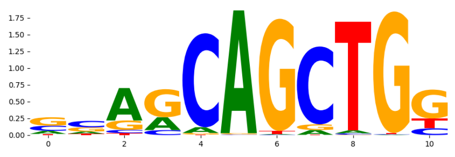 |
| neg_patterns.pattern_9 | 54 | 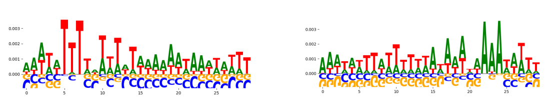 | | SP1_HUMAN.H11MO.0.A | 3.405030e-02 | 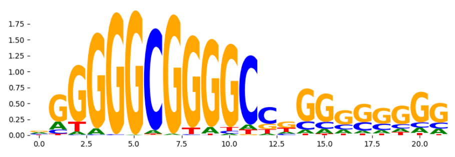 |
| neg_patterns.pattern_10 | 46 | 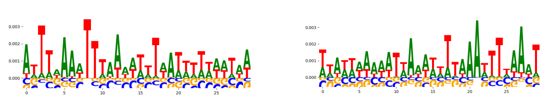 | | RXRG_MOUSE.H11MO.0.B | 1.336440e-01 | 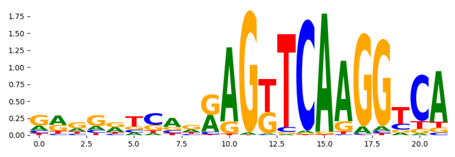 |
| neg_patterns.pattern_11 | 46 | 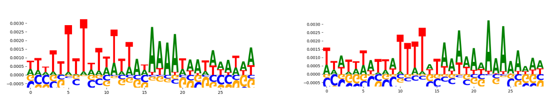 | | TAF1_HUMAN.H11MO.0.A | 1.909640e-01 | 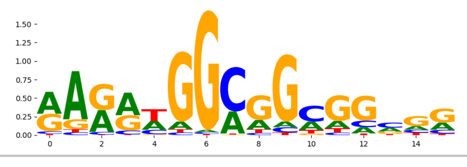 |
