## Supplementary Files 4 for "ChromBPNet: bias factorized, base-resolution deep learning models of chromatin accessibility reveal cis-regulatory sequence syntax, transcription factor footprints and regulatory variants": fig4b_naked_dna_ATAC_raw_bias_model_profile_modisco.pdf

| pattern | num_seqlets | cwm_fwd | cwm_rev | TOMTOM_match | TOMTOM_qval | TOMTOM_match_logo |
|---|---|---|---|---|---|---|
| pos_patterns.pattern_0 | 13284 | 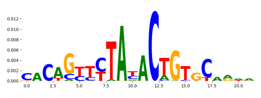 | 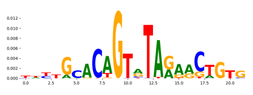 | TN5_2 | 2.585450e-02 | 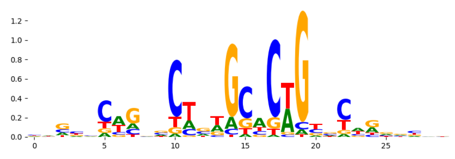 |
| pos_patterns.pattern_1 | 7275 | 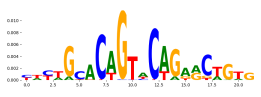 | 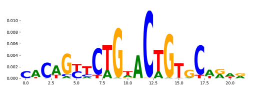 | TN5_2 | 7.139180e-06 | 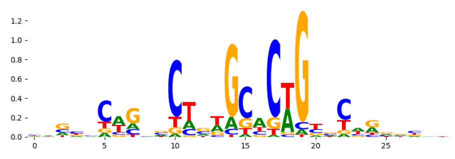 |
| pos_patterns.pattern_2 | 4538 | 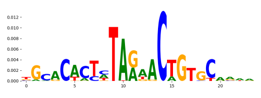 | 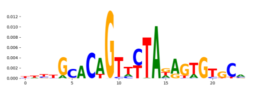 | TN5_3 | 2.477680e-04 | 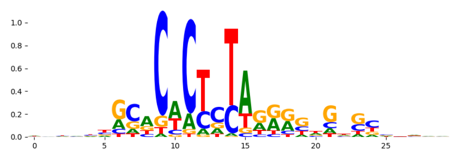 |
| pos_patterns.pattern_3 | 3923 | 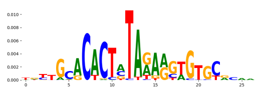 | 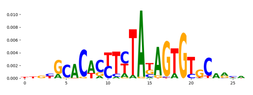 | TN5_3 | 5.278590e-10 | 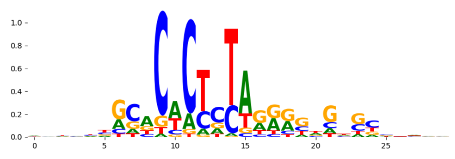 |
| pos_patterns.pattern_4 | 190 | 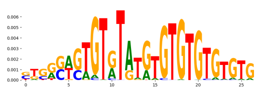 | 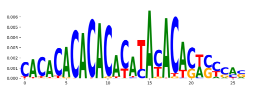 | EGR2_HUMAN.H11MO.0.A | 1.000000e+00 | 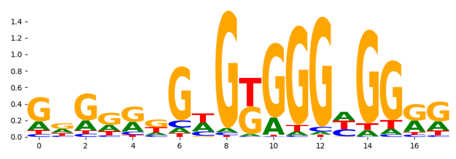 |
| pos_patterns.pattern_5 | 175 | 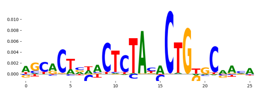 | 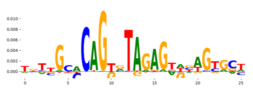 | TN5_6 | 3.491220e-07 | 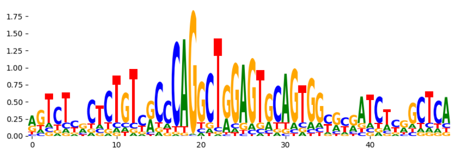 |
| pos_patterns.pattern_6 | 131 | 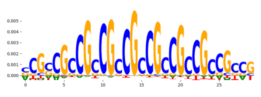 | 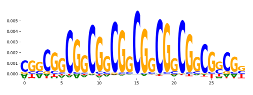 | SP2_HUMAN.H11MO.0.A | 8.263200e-04 | 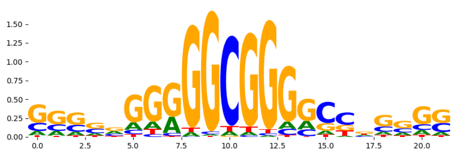 |
| pos_patterns.pattern_7 | 47 | 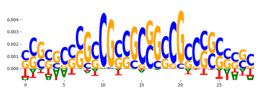 | 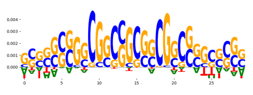 | SP2_HUMAN.H11MO.0.A | 5.127050e-07 | 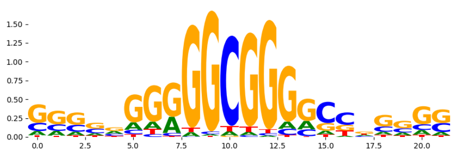 |
| pos_patterns.pattern_8 | 31 | 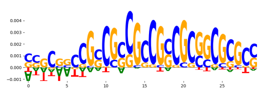 | 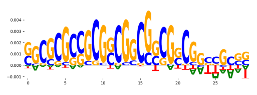 | SP2_HUMAN.H11MO.0.A | 2.250760e-04 | 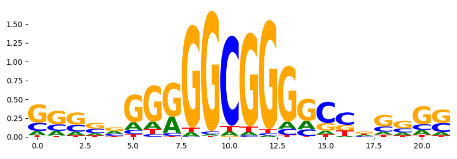 |
| pos_patterns.pattern_9 | 26 | 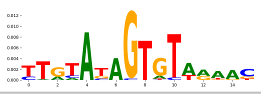 | 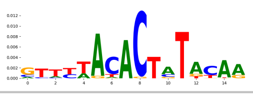 | NaN | NaN | |
