## Supplementary Files 4 for "ChromBPNet: bias factorized, base-resolution deep learning models of chromatin accessibility reveal cis-regulatory sequence syntax, transcription factor footprints and regulatory variants": fig4c_naked_dna_DNASE_raw_bias_model_counts_modisco.pdf

| pattern | num_seqlets | cwm_fwd | cwm_rev | TOMTOM_match | TOMTOM_qval | TOMTOM_match_logo |
|---|---|---|---|---|---|---|
| pos_patterns.pattern_0 | 2213 | | | ZNF384_MA1125.1 | 0.028745 | |
| pos_patterns.pattern_1 | 1516 | | | DNASE_2 | 1.000000 | |
| pos_patterns.pattern_2 | 1056 | | | FOXB1_forkhead_2 | 0.169313 | |
| pos_patterns.pattern_3 | 872 | | | ZNF384_MA1125.1 | 0.026828 | |
| pos_patterns.pattern_4 | 836 | | | DNASE_2 | 0.049511 | |
| pos_patterns.pattern_5 | 547 | | | MEF2B_HUMAN.H11MO.0.A | 0.281251 | |
| pos_patterns.pattern_6 | 324 | | | SP2_HUMAN.H11MO.0.A | 0.010986 | |
| pos_patterns.pattern_7 | 297 | | | PRDM6_HUMAN.H11MO.0.C | 0.038345 | |
| pos_patterns.pattern_8 | 207 | | | BRAC_MOUSE.H11MO.0.B | 1.000000 | |
| pos_patterns.pattern_9 | 94 | | | ZN264_HUMAN.H11MO.0.C | 1.000000 | |
| pos_patterns.pattern_10 | 82 | | | ZN770_HUMAN.H11MO.0.C | 0.512755 | |
| pos_patterns.pattern_11 | 54 | | | ZNF384_MA1125.1 | 1.000000 | |
| pos_patterns.pattern_12 | 22 | | | IRF3_HUMAN.H11MO.0.B | 0.533869 | |
| neg_patterns.pattern_0 | 4087 | | | MYBA_MOUSE.H11MO.0.C | 1.000000 | |
| neg_patterns.pattern_1 | 37 | | | ZNF435_C2H2_1 | 0.354321 | |
| neg_patterns.pattern_2 | 20 | | | FOXH1_MA0479.1 | 1.000000 | |

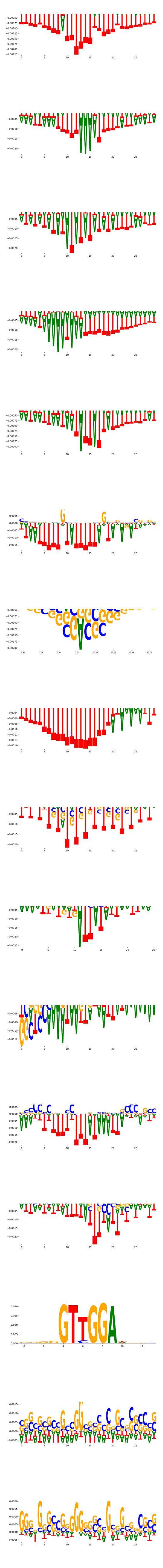

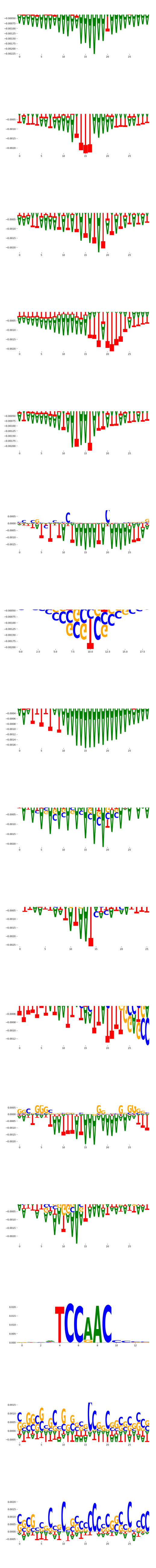

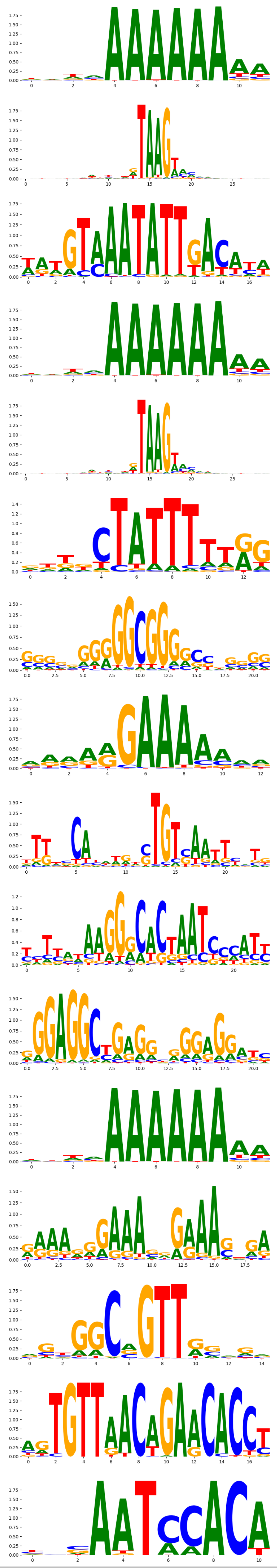
