## Supplementary Files 4 for "ChromBPNet: bias factorized, base-resolution deep learning models of chromatin accessibility reveal cis-regulatory sequence syntax, transcription factor footprints and regulatory variants": fig4d_naked_dna_DNASE_raw_bias_model_profile_modisco.pdf

| pattern | num_seqlets | cwm_fwd | cwm_rev | TOMTOM_match | TOMTOM_qval | TOMTOM_match_logo |
|---|---|---|---|---|---|---|
| pos_patterns.pattern_0 | 2214 | | | ZNF384_MA1125.1 | 7.792200e-02 | |
| pos_patterns.pattern_1 | 1700 | | | SOX8_HMG_8 | 6.104490e-01 | |
| pos_patterns.pattern_2 | 1368 | | | Ar.mouse_nuclearreceptor_1 | 1.000000e+00 | |
| pos_patterns.pattern_3 | 1048 | | | IKZF1_HUMAN.H11MO.0.C | 1.000000e+00 | |
| pos_patterns.pattern_4 | 846 | | | TN5_6 | 4.483670e-23 | |
| pos_patterns.pattern_5 | 793 | | | RARG_HUMAN.H11MO.0.B | 9.358020e-01 | |
| pos_patterns.pattern_6 | 771 | | | ZN770_HUMAN.H11MO.0.C | 8.241370e-09 | |
| pos_patterns.pattern_7 | 584 | | | DNASE_5 | 4.886980e-04 | |
| pos_patterns.pattern_8 | 566 | | | ZN816_HUMAN.H11MO.0.C | 1.000000e+00 | |
| pos_patterns.pattern_9 | 474 | | | MEF2B_HUMAN.H11MO.0.A | 2.864880e-01 | |
| pos_patterns.pattern_10 | 444 | | | DNASE_2 | 3.083670e-01 | |
| pos_patterns.pattern_11 | 404 | | | ONECUT3_CUT_1 | 2.661330e-01 | |
| pos_patterns.pattern_12 | 404 | | | DNASE_2 | 4.653150e-01 | |
| pos_patterns.pattern_13 | 318 | | | RREB1_MA0073.1 | 1.000000e+00 | |
| pos_patterns.pattern_14 | 276 | | | RELB_MA1117.1 | 1.000000e+00 | |
| pos_patterns.pattern_15 | 211 | | | TN5_6 | 2.305050e-04 | |
| pos_patterns.pattern_16 | 207 | | | ZN770_HUMAN.H11MO.0.C | 1.732780e-13 | |
| pos_patterns.pattern_17 | 170 | | | IRF3_HUMAN.H11MO.0.B | 2.484470e-02 | |
| pos_patterns.pattern_18 | 132 | | | MEF2B_HUMAN.H11MO.0.A | 2.757370e-01 | |
| pos_patterns.pattern_19 | 115 | | | E2F7_E2F_1 | 1.462340e-01 | |
| pos_patterns.pattern_20 | 86 | | | SP5_MOUSE.H11MO.0.C | 7.817040e-08 | |
| pos_patterns.pattern_21 | 69 | | | FOXC1_forkhead_1 | 1.533760e-01 | |
| pos_patterns.pattern_22 | 29 | | | ZNF384_MA1125.1 | 1.000000e+00 | |
| pos_patterns.pattern_23 | 20 | | | ZN770_HUMAN.H11MO.0.C | 1.569570e-02 | |

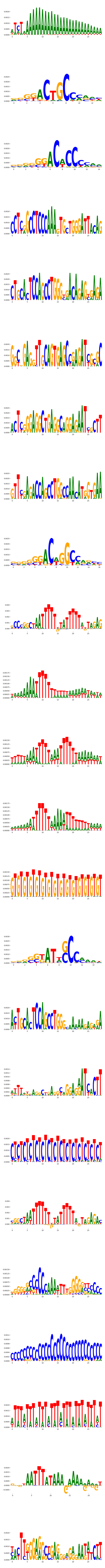
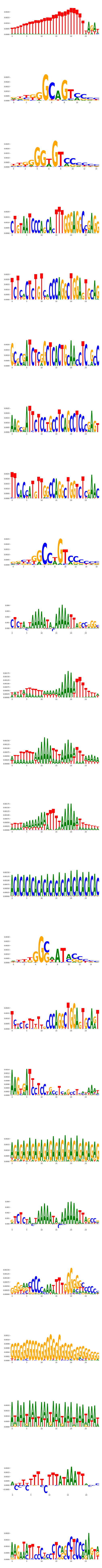
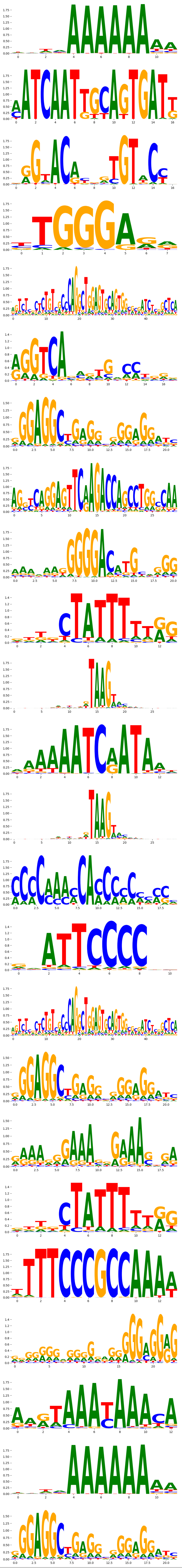
