## Supplementary Files 4 for "ChromBPNet: bias factorized, base-resolution deep learning models of chromatin accessibility reveal cis-regulatory sequence syntax, transcription factor footprints and regulatory variants": fig4e_gm12878_ATAC_raw_bpnet_w_naked_dna_bias_correction_counts_modisco.pdf

| pattern | num_seqlets | cwm_fwd | cwm_rev | TOMTOM_match | TOMTOM_qval | TOMTOM_match_logo |
|---|---|---|---|---|---|---|
| pos_patterns.pattern_0 | 3371 | | | CTCF_MA0139.1 | 3.367020e-15 | |
| pos_patterns.pattern_1 | 3167 | | | ELF5_HUMAN.H11MO.0.A | 1.709640e-05 | |
| pos_patterns.pattern_2 | 2828 | | | IRF4_MOUSE.H11MO.0.A | 4.495630e-02 | |
| pos_patterns.pattern_3 | 2645 | | | FOSL2+JUN_MA1130.1 | 2.656940e-03 | |
| pos_patterns.pattern_4 | 1796 | | | RUNX1_HUMAN.H11MO.0.A | 7.671330e-02 | |
| pos_patterns.pattern_5 | 1713 | | | IRF1_MOUSE.H11MO.0.A | 1.617020e-05 | |
| pos_patterns.pattern_6 | 1520 | | | NFKB2_HUMAN.H11MO.0.B | 5.195800e-06 | |
| pos_patterns.pattern_7 | 734 | | | SP1_HUMAN.H11MO.0.A | 6.772420e-05 | |
| pos_patterns.pattern_8 | 353 | | | NFYB_HUMAN.H11MO.0.A | 1.160880e-02 | |
| pos_patterns.pattern_9 | 353 | | | POU5F1_MA1115.1 | 4.286820e-03 | |
| pos_patterns.pattern_10 | 352 | | | NFKB1_HUMAN.H11MO.1.B | 8.951890e-03 | |
| pos_patterns.pattern_11 | 336 | | | BATF_HUMAN.H11MO.0.A | 1.347630e-07 | |
| pos_patterns.pattern_12 | 300 | | | NRF1_HUMAN.H11MO.0.A | 1.316570e-06 | |
| pos_patterns.pattern_13 | 270 | | | REL_MA0101.1 | 7.345320e-01 | |
| pos_patterns.pattern_14 | 266 | | | IRF4_HUMAN.H11MO.0.A | 1.119760e-03 | |
| pos_patterns.pattern_15 | 251 | | | HNF1A_HUMAN.H11MO.0.C | 1.063610e-07 | |
| pos_patterns.pattern_16 | 241 | | | PAX5_MOUSE.H11MO.0.A | 2.608270e-12 | |
| pos_patterns.pattern_17 | 217 | | | COE1_HUMAN.H11MO.0.A | 2.717330e-07 | |
| pos_patterns.pattern_18 | 156 | | | SPIB_ETS_1 | 6.161940e-03 | |
| pos_patterns.pattern_19 | 143 | | | ATF1_MOUSE.H11MO.0.B | 2.089910e-03 | |
| pos_patterns.pattern_20 | 139 | | | MEF2B_MA0660.1 | 1.317560e-07 | |
| pos_patterns.pattern_21 | 114 | | | ZN143_MOUSE.H11MO.0.A | 4.571410e-17 | |
| pos_patterns.pattern_22 | 97 | | | IRF9_IRF_1 | 2.099090e-03 | |
| pos_patterns.pattern_23 | 78 | | | FOSL2+JUN_MA1131.1 | 7.084870e-05 | |
| pos_patterns.pattern_24 | 74 | | | PAX2_PAX_1 | 8.249670e-09 | |
| pos_patterns.pattern_25 | 53 | | | KAISO_HUMAN.H11MO.0.A | 1.145200e-07 | |
| pos_patterns.pattern_26 | 50 | | | TYY1_HUMAN.H11MO.0.A | 8.027640e-06 | |
| pos_patterns.pattern_27 | 34 | | | THA11_HUMAN.H11MO.0.B | 1.533700e-06 | |
| pos_patterns.pattern_28 | 27 | | | PAX5_MOUSE.H11MO.0.A | 2.136890e-04 | |

| pattern | num_seqlets | cwm_fwd | cwm_rev | TOMTOM_match | TOMTOM_qval | TOMTOM_match_logo |
|---|---|---|---|---|---|---|
| pos_patterns.pattern_29 | 24 | 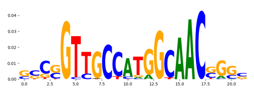 | 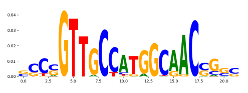 | RFX2_HUMAN.H11MO.0.A | 2.687770e-09 | 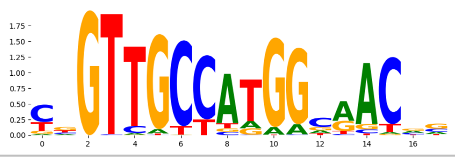 |
