## Supplementary Files 4 for "ChromBPNet: bias factorized, base-resolution deep learning models of chromatin accessibility reveal cis-regulatory sequence syntax, transcription factor footprints and regulatory variants": fig4f_gm12878_ATAC_raw_bpnet_w_naked_dna_bias_correction_profile_modisco.pdf

| pattern | num_seqlets | cwm_fwd | cwm_rev | TOMTOM_match | TOMTOM_qval | TOMTOM_match_logo |
|---|---|---|---|---|---|---|
| pos_patterns.pattern_0 | 4013 | | | CTCF_MA0139.1 | 6.793640e-17 | |
| pos_patterns.pattern_1 | 1975 | | | SP2_HUMAN.H11MO.0.A | 5.910480e-10 | |
| pos_patterns.pattern_2 | 1959 | | | SP2_HUMAN.H11MO.0.A | 7.311640e-05 | |
| pos_patterns.pattern_3 | 1601 | | | ATF3_MOUSE.H11MO.0.A | 1.871830e-03 | |
| pos_patterns.pattern_4 | 1175 | | | ELF5_HUMAN.H11MO.0.A | 7.760780e-05 | |
| pos_patterns.pattern_5 | 1101 | | | IRF1_MOUSE.H11MO.0.A | 2.154140e-03 | |
| pos_patterns.pattern_6 | 863 | | | SP2_HUMAN.H11MO.0.A | 7.617620e-19 | |
| pos_patterns.pattern_7 | 787 | | | RELB_HUMAN.H11MO.0.C | 6.935320e-06 | |
| pos_patterns.pattern_8 | 662 | | | KLF5_MOUSE.H11MO.0.A | 1.060060e-04 | |
| pos_patterns.pattern_9 | 624 | | | IRF4_HUMAN.H11MO.0.A | 1.261150e-10 | |
| pos_patterns.pattern_10 | 622 | | | IRF1_HUMAN.H11MO.0.A | 1.047540e-06 | |
| pos_patterns.pattern_11 | 532 | | | RUNX3_HUMAN.H11MO.0.A | 4.602700e-01 | |
| pos_patterns.pattern_12 | 516 | | | SP2_HUMAN.H11MO.0.A | 3.356900e-06 | |
| pos_patterns.pattern_13 | 377 | | | MYOD1_HUMAN.H11MO.0.A | 2.843030e-03 | |
| pos_patterns.pattern_14 | 343 | | | NFYB_HUMAN.H11MO.0.A | 3.196650e-07 | |
| pos_patterns.pattern_15 | 127 | | | BATF_HUMAN.H11MO.0.A | 1.678740e-05 | |
| pos_patterns.pattern_16 | 92 | | | MSX2_homeodomain_1 | 3.105830e-01 | |
| pos_patterns.pattern_17 | 88 | | | Pou2f2.mouse_POU_2 | 1.832920e-02 | |
| pos_patterns.pattern_18 | 84 | | | NRF1_MA0506.1 | 3.717790e-04 | |
| pos_patterns.pattern_19 | 63 | | | ATF2_HUMAN.H11MO.0.B | 1.401630e-04 | |
| pos_patterns.pattern_20 | 60 | | | CTCF_MA0139.1 | 4.398870e-04 | |
| pos_patterns.pattern_21 | 54 | | | STAT1_MOUSE.H11MO.0.A | 6.753500e-01 | |
| pos_patterns.pattern_22 | 27 | | | JUN_MA0489.1 | 1.196910e-02 | |
| pos_patterns.pattern_23 | 26 | | | TYY1_HUMAN.H11MO.0.A | 2.238340e-05 | |
| pos_patterns.pattern_24 | 25 | | | SPI1_MOUSE.H11MO.0.A | 6.180360e-02 | |
| pos_patterns.pattern_25 | 22 | | | ZNF76_HUMAN.H11MO.0.C | 7.640220e-12 | |
| neg_patterns.pattern_0 | 135 | | | Stat5a+Stat5b_MA0519.1 | 1.000000e+00 | |
| neg_patterns.pattern_1 | 63 | | | ZNF384_MA1125.1 | 5.151300e-01 | |
| neg_patterns.pattern_2 | 48 | | | IRF4_HUMAN.H11MO.0.A | 9.631840e-05 | |

| pattern | num_seqlets | cwm_fwd | cwm_rev | TOMTOM_match | TOMTOM_qval | TOMTOM_match_logo |
|---|---|---|---|---|---|---|
| neg_patterns.pattern_3 | 42 | 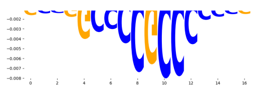 | 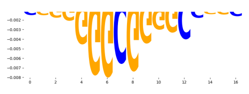 | KLF12_HUMAN.H11MO.0.C | 2.936490e-04 | 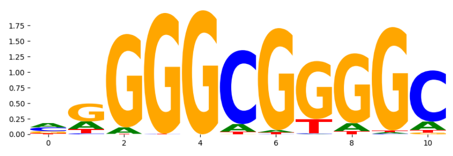 |
| neg_patterns.pattern_4 | 31 | 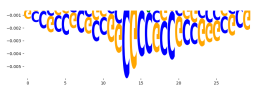 | 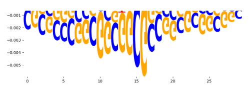 | SP2_HUMAN.H11MO.0.A | 6.515550e-07 | 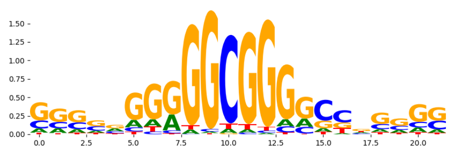 |
| neg_patterns.pattern_5 | 29 | 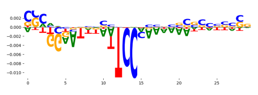 | 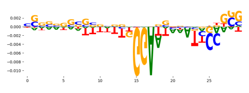 | STAT3_MA0144.2 | 1.178100e-01 | 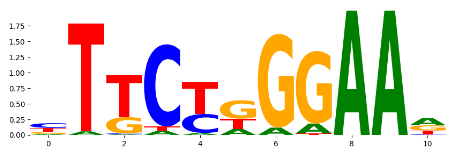 |
| neg_patterns.pattern_6 | 29 | 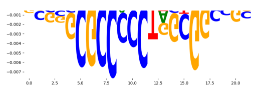 | 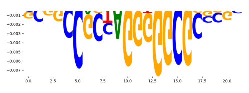 | CTCFL_MOUSE.H11MO.0.A | 5.807460e-07 | 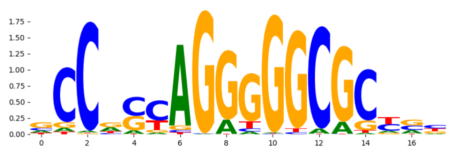 |
| neg_patterns.pattern_7 | 24 | 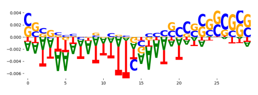 | 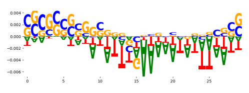 | MXI1_HUMAN.H11MO.0.A | 4.364370e-01 | 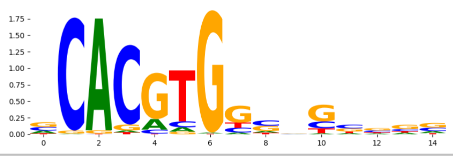 |
