## Supplementary Files 4 for "ChromBPNet: bias factorized, base-resolution deep learning models of chromatin accessibility reveal cis-regulatory sequence syntax, transcription factor footprints and regulatory variants": fig4g_gm12878_ATAC_raw_bpnet_w_heg2_bias_correction_counts_modisco.pdf

| pattern | num_seqlets | cwm_fwd | cwm_rev | TOMTOM_match | TOMTOM_qval | TOMTOM_match_logo |
|---|---|---|---|---|---|---|
| pos_patterns.pattern_0 | 5384 | | | IRF1_MOUSE.H11MO.0.A | 2.976030e-03 | |
| pos_patterns.pattern_1 | 3024 | | | ELF5_HUMAN.H11MO.0.A | 2.347390e-05 | |
| pos_patterns.pattern_2 | 2882 | | | CTCF_MA0139.1 | 1.703230e-15 | |
| pos_patterns.pattern_3 | 2438 | | | FOS+JUN_MA0099.3 | 1.978980e-03 | |
| pos_patterns.pattern_4 | 1920 | | | RUNX3_HUMAN.H11MO.0.A | 7.953870e-02 | |
| pos_patterns.pattern_5 | 1897 | | | NFKB1_HUMAN.H11MO.1.B | 9.715450e-07 | |
| pos_patterns.pattern_6 | 961 | | | IRF4_HUMAN.H11MO.0.A | 2.231930e-05 | |
| pos_patterns.pattern_7 | 713 | | | KLF12_HUMAN.H11MO.0.C | 1.858470e-04 | |
| pos_patterns.pattern_8 | 422 | | | NFYC_HUMAN.H11MO.0.A | 1.439830e-03 | |
| pos_patterns.pattern_9 | 320 | | | NRF1_HUMAN.H11MO.0.A | 2.003370e-07 | |
| pos_patterns.pattern_10 | 255 | | | REL_MA0101.1 | 9.350560e-01 | |
| pos_patterns.pattern_11 | 241 | | | CREB1_HUMAN.H11MO.0.A | 1.915050e-05 | |
| pos_patterns.pattern_12 | 217 | | | POU5F1_MA1115.1 | 3.560680e-03 | |
| pos_patterns.pattern_13 | 208 | | | EBF1_EBF_1 | 4.518600e-06 | |
| pos_patterns.pattern_14 | 194 | | | ETS1_HUMAN.H11MO.0.A | 1.303170e-02 | |
| pos_patterns.pattern_15 | 169 | | | ZN143_MOUSE.H11MO.0.A | 8.036540e-11 | |
| pos_patterns.pattern_16 | 155 | | | SPI1_ETS_1 | 1.712780e-02 | |
| pos_patterns.pattern_17 | 134 | | | PAX5_MOUSE.H11MO.0.A | 4.597220e-14 | |
| pos_patterns.pattern_18 | 94 | | | TYY1_HUMAN.H11MO.0.A | 2.384500e-06 | |
| pos_patterns.pattern_19 | 93 | | | KAISO_HUMAN.H11MO.0.A | 5.783060e-08 | |
| pos_patterns.pattern_20 | 63 | | | PAX5_MOUSE.H11MO.0.A | 9.624550e-06 | |
| pos_patterns.pattern_21 | 42 | | | RFX5_RFX_2 | 5.467310e-06 | |
| pos_patterns.pattern_22 | 36 | | | MEF2B_MA0660.1 | 4.334320e-05 | |
| pos_patterns.pattern_23 | 34 | | | ARNTL_bHLH_1 | 1.825530e-04 | |
