## Supplementary Files 4 for "ChromBPNet: bias factorized, base-resolution deep learning models of chromatin accessibility reveal cis-regulatory sequence syntax, transcription factor footprints and regulatory variants": fig4h_gm12878_ATAC_raw_bpnet_w_hepg2_bias_correction_profile_modisco.pdf

| pattern | num_seqlets | cwm_fwd | cwm_rev | TOMTOM_match | TOMTOM_qval | TOMTOM_match_logo |
|---|---|---|---|---|---|---|
| pos_patterns.pattern_0 | 6200 | | | IRF1_MOUSE.H11MO.0.A | 3.150150e-03 | |
| pos_patterns.pattern_1 | 3999 | | | CTCF_MA0139.1 | 7.824870e-16 | |
| pos_patterns.pattern_2 | 2744 | | | SPIC_ETS_1 | 2.065390e-04 | |
| pos_patterns.pattern_3 | 2579 | | | JDP2_MA0655.1 | 2.878720e-03 | |
| pos_patterns.pattern_4 | 1969 | | | RUNX3_HUMAN.H11MO.0.A | 1.026170e-01 | |
| pos_patterns.pattern_5 | 1527 | | | NFKB1_HUMAN.H11MO.1.B | 4.285830e-06 | |
| pos_patterns.pattern_6 | 1154 | | | KLF12_HUMAN.H11MO.0.C | 1.531090e-04 | |
| pos_patterns.pattern_7 | 990 | | | NFYC_HUMAN.H11MO.0.A | 1.764620e-04 | |
| pos_patterns.pattern_8 | 739 | | | IRF4_HUMAN.H11MO.0.A | 1.152220e-05 | |
| pos_patterns.pattern_9 | 566 | | | SPIB_MA0081.1 | 1.327240e-01 | |
| pos_patterns.pattern_10 | 558 | | | FOSL2+JUN_MA1131.1 | 2.591580e-07 | |
| pos_patterns.pattern_11 | 412 | | | Pou2f2.mouse_POU_2 | 5.607450e-03 | |
| pos_patterns.pattern_12 | 398 | | | BATF_HUMAN.H11MO.0.A | 2.301610e-10 | |
| pos_patterns.pattern_13 | 347 | | | NRF1_MOUSE.H11MO.0.A | 1.509740e-06 | |
| pos_patterns.pattern_14 | 224 | | | TYY1_HUMAN.H11MO.0.A | 2.013350e-04 | |
| pos_patterns.pattern_15 | 210 | | | FOXJ3_HUMAN.H11MO.0.A | 7.092900e-02 | |
| pos_patterns.pattern_16 | 197 | | | ZN143_MOUSE.H11MO.0.A | 3.344770e-17 | |
| pos_patterns.pattern_17 | 119 | | | MITF_HUMAN.H11MO.0.A | 2.876540e-05 | |
| pos_patterns.pattern_18 | 90 | | | PAX2_PAX_1 | 4.858030e-07 | |
| pos_patterns.pattern_19 | 87 | | | PAX5_HUMAN.H11MO.0.A | 1.344530e-11 | |
| pos_patterns.pattern_20 | 86 | | | COE1_HUMAN.H11MO.0.A | 2.232010e-05 | |
| pos_patterns.pattern_21 | 81 | | | RELA_MA0107.1 | 4.932500e-04 | |
| pos_patterns.pattern_22 | 65 | | | Rfx1_MA0509.1 | 5.223400e-08 | |
| pos_patterns.pattern_23 | 61 | | | THAP1_HUMAN.H11MO.0.C | 5.516520e-02 | |
| pos_patterns.pattern_24 | 60 | | | NFKB1_HUMAN.H11MO.1.B | 1.733130e-03 | |
| pos_patterns.pattern_25 | 59 | | | STAT1+STAT2_MA0517.1 | 2.505030e-05 | |
| pos_patterns.pattern_26 | 56 | | | KAISO_HUMAN.H11MO.0.A | 1.214560e-04 | |
| pos_patterns.pattern_27 | 37 | | | PRDM1_C2H2_1 | 3.588410e-01 | |
| pos_patterns.pattern_28 | 29 | | | BATF+JUN_MA0462.1 | 3.354280e-02 | |

| pattern | num_seqlets | cwm_fwd | cwm_rev | TOMTOM_match | TOMTOM_qval | TOMTOM_match_logo |
|---|---|---|---|---|---|---|
| pos_patterns.pattern_29 | 24 | 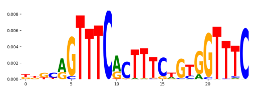 | 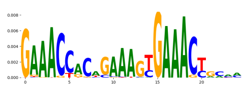 | IRF9_HUMAN.H11MO.0.C | 1.058150e-02 | 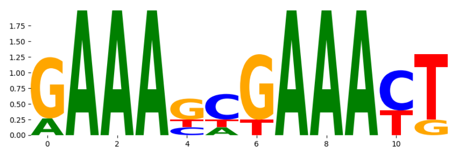 |
| pos_patterns.pattern_30 | 21 | 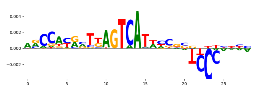 | 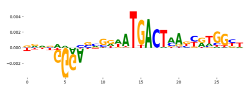 | BATF+JUN_MA0462.1 | 5.976100e-02 | 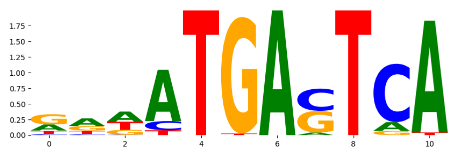 |
| neg_patterns.pattern_0 | 587 | 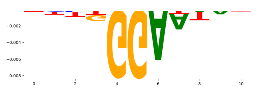 | 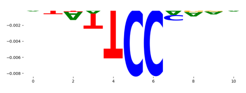 | NaN | NaN | |
| neg_patterns.pattern_1 | 79 | 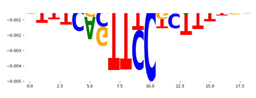 | 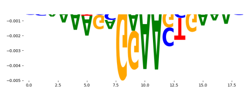 | BC11A_HUMAN.H11MO.0.A | 9.053620e-04 | 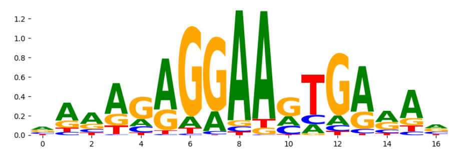 |
| neg_patterns.pattern_2 | 74 | 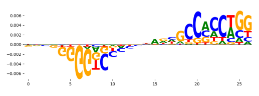 | 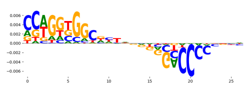 | RARA+RXRG_MA1149.1 | 6.950980e-02 | 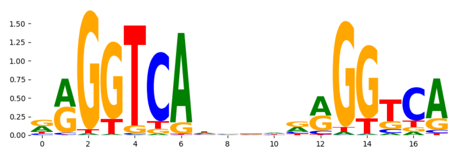 |
| neg_patterns.pattern_3 | 54 | 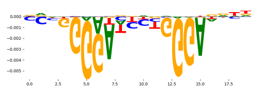 | 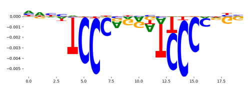 | SIX2_HUMAN.H11MO.0.A | 1.000000e+00 | 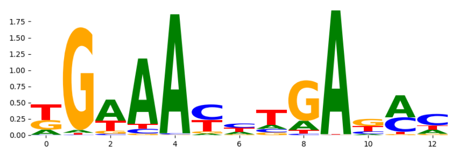 |
| neg_patterns.pattern_4 | 32 | 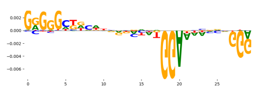 | 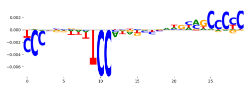 | NFATC2_MA0152.1 | 1.093190e-01 | 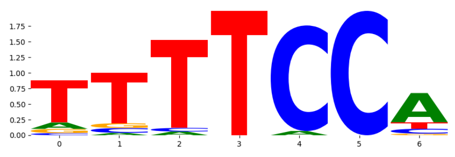 |
| neg_patterns.pattern_5 | 31 | 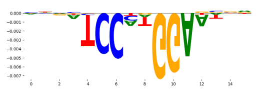 | 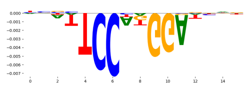 | NFATC1_NFAT_3 | 2.256470e-01 | 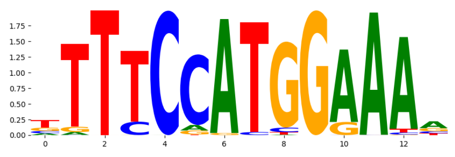 |
| neg_patterns.pattern_6 | 23 | 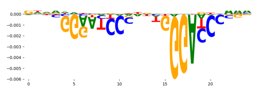 | 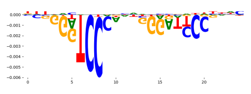 | ZIC3_HUMAN.H11MO.0.B | 4.980120e-01 | 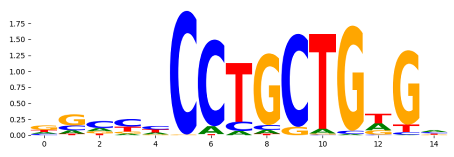 |
| neg_patterns.pattern_7 | 22 | 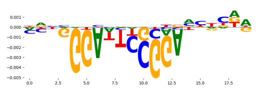 | 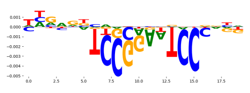 | ZFP82_HUMAN.H11MO.0.C | 1.000000e+00 | 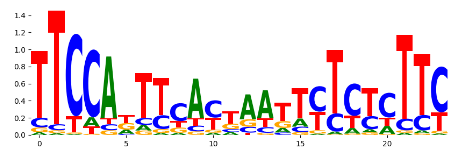 |
| neg_patterns.pattern_8 | 20 | 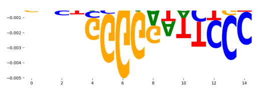 | 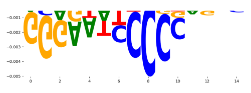 | NFKB2_MA0778.1 | 2.425160e-03 | 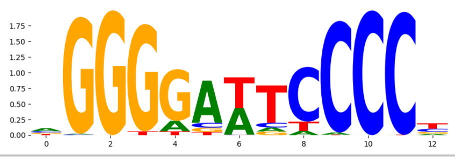 |
