## Supplementary Files 4 for "ChromBPNet: bias factorized, base-resolution deep learning models of chromatin accessibility reveal cis-regulatory sequence syntax, transcription factor footprints and regulatory variants": fig4i_gm12878_DNASE_raw_bpnet_w_naked_dna_bias_correction_counts_modisco.pdf

| pattern | num_seqlets | cwm_fwd | cwm_rev | TOMTOM_match | TOMTOM_qval | TOMTOM_match_logo |
|---|---|---|---|---|---|---|
| pos_patterns.pattern_0 | 5051 | | | IRF1_MOUSE.H11MO.0.A | 3.028240e-04 | |
| pos_patterns.pattern_1 | 4705 | | | ELF5_HUMAN.H11MO.0.A | 2.137510e-05 | |
| pos_patterns.pattern_2 | 3938 | | | CTCF_MA0139.1 | 3.953770e-12 | |
| pos_patterns.pattern_3 | 2445 | | | ATF3_MOUSE.H11MO.0.A | 4.622950e-03 | |
| pos_patterns.pattern_4 | 2227 | | | RUNX3_HUMAN.H11MO.0.A | 1.765460e-02 | |
| pos_patterns.pattern_5 | 1713 | | | NFKB1_HUMAN.H11MO.1.B | 3.333090e-07 | |
| pos_patterns.pattern_6 | 1238 | | | KLF12_HUMAN.H11MO.0.C | 2.043950e-05 | |
| pos_patterns.pattern_7 | 737 | | | NRF1_MA0506.1 | 2.261740e-06 | |
| pos_patterns.pattern_8 | 608 | | | NFYB_HUMAN.H11MO.0.A | 7.535850e-03 | |
| pos_patterns.pattern_9 | 600 | | | ATF1_MOUSE.H11MO.0.B | 1.557860e-03 | |
| pos_patterns.pattern_10 | 475 | | | POU5F1_MA1115.1 | 3.272310e-03 | |
| pos_patterns.pattern_11 | 406 | | | ETS1_HUMAN.H11MO.0.A | 2.050190e-03 | |
| pos_patterns.pattern_12 | 370 | | | COE1_MOUSE.H11MO.0.A | 3.416240e-05 | |
| pos_patterns.pattern_13 | 362 | | | ZN143_MOUSE.H11MO.0.A | 1.135250e-19 | |
| pos_patterns.pattern_14 | 300 | | | NFKB1_HUMAN.H11MO.1.B | 6.442470e-03 | |
| pos_patterns.pattern_15 | 149 | | | PAX1_MA0779.1 | 1.031160e-07 | |
| pos_patterns.pattern_16 | 134 | | | REL_MA0101.1 | 4.615740e-01 | |
| pos_patterns.pattern_17 | 104 | | | SPIB_ETS_1 | 3.048380e-02 | |
| pos_patterns.pattern_18 | 104 | | | ZBTB33_MA0527.1 | 6.383250e-05 | |
| pos_patterns.pattern_19 | 64 | | | MEF2D_MOUSE.H11MO.0.A | 4.916110e-05 | |
| pos_patterns.pattern_20 | 44 | | | IRF4_IRF_1 | 1.493280e-02 | |
| pos_patterns.pattern_21 | 26 | | | RUNX3_RUNX_1 | 4.937990e-02 | |
| pos_patterns.pattern_22 | 23 | | | IRF7_HUMAN.H11MO.0.C | 2.574340e-05 | |
| pos_patterns.pattern_23 | 23 | | | Rfx1_MA0509.1 | 2.090690e-07 | |
| pos_patterns.pattern_24 | 23 | | | TYY1_HUMAN.H11MO.0.A | 9.047930e-05 | |
