## Supplementary Files 4 for "ChromBPNet: bias factorized, base-resolution deep learning models of chromatin accessibility reveal cis-regulatory sequence syntax, transcription factor footprints and regulatory variants": fig4j_gm12878_DNASE_raw_bpnet_w_naked_dna_bias_correction_profile_modisco.pdf

| pattern | num_seqlets | cwm_fwd | cwm_rev | TOMTOM_match | TOMTOM_qval | TOMTOM_match_logo |
|---|---|---|---|---|---|---|
| pos_patterns.pattern_0 | 4270 | | | CTCF_MA0139.1 | 9.019020e-12 | |
| pos_patterns.pattern_1 | 3318 | | | IRF4_MOUSE.H11MO.0.A | 5.473920e-05 | |
| pos_patterns.pattern_2 | 2783 | | | ETS1_HUMAN.H11MO.0.A | 8.936240e-04 | |
| pos_patterns.pattern_3 | 1971 | | | ATF3_MOUSE.H11MO.0.A | 3.759360e-03 | |
| pos_patterns.pattern_4 | 1584 | | | RUNX3_HUMAN.H11MO.0.A | 2.269830e-03 | |
| pos_patterns.pattern_5 | 1058 | | | KLF12_HUMAN.H11MO.0.C | 8.237540e-05 | |
| pos_patterns.pattern_6 | 1025 | | | RELB_HUMAN.H11MO.0.C | 5.611040e-07 | |
| pos_patterns.pattern_7 | 816 | | | ZN667_HUMAN.H11MO.0.C | 1.000000e+00 | |
| pos_patterns.pattern_8 | 770 | | | NFYB_HUMAN.H11MO.0.A | 6.101990e-04 | |
| pos_patterns.pattern_9 | 724 | | | ZNF384_MA1125.1 | 6.522740e-03 | |
| pos_patterns.pattern_10 | 717 | | | ZNF384_MA1125.1 | 8.926680e-02 | |
| pos_patterns.pattern_11 | 590 | | | PRDM6_HUMAN.H11MO.0.C | 9.864940e-02 | |
| pos_patterns.pattern_12 | 579 | | | NRF1_MOUSE.H11MO.0.A | 4.311620e-08 | |
| pos_patterns.pattern_13 | 529 | | | TBX1_TBX_1 | 7.227480e-01 | |
| pos_patterns.pattern_14 | 525 | | | SP5_MOUSE.H11MO.0.C | 1.752400e-07 | |
| pos_patterns.pattern_15 | 429 | | | BATF3_HUMAN.H11MO.0.B | 5.567150e-06 | |
| pos_patterns.pattern_16 | 391 | | | ZNF384_MA1125.1 | 6.292510e-02 | |
| pos_patterns.pattern_17 | 317 | | | TN5_6 | 1.462770e-20 | |
| pos_patterns.pattern_18 | 311 | | | FOSB+JUNB_MA1136.1 | 2.066430e-05 | |
| pos_patterns.pattern_19 | 287 | | | EBF1_EBF_1 | 1.139310e-05 | |
| pos_patterns.pattern_20 | 277 | | | RREB1_MA0073.1 | 1.000000e+00 | |
| pos_patterns.pattern_21 | 244 | | | ZN143_MOUSE.H11MO.0.A | 5.324780e-17 | |
| pos_patterns.pattern_22 | 229 | | | Pou2f2.mouse_POU_2 | 4.285970e-03 | |
| pos_patterns.pattern_23 | 222 | | | SP5_MOUSE.H11MO.0.C | 5.818330e-07 | |
| pos_patterns.pattern_24 | 196 | | | PAX5_MA0014.3 | 1.000000e+00 | |
| pos_patterns.pattern_25 | 178 | | | ARNTL_bHLH_1 | 1.914400e-05 | |
| pos_patterns.pattern_26 | 149 | | | MEF2B_HUMAN.H11MO.0.A | 2.436510e-01 | |
| pos_patterns.pattern_27 | 133 | | | PAX1_MA0779.1 | 5.781960e-05 | |
| pos_patterns.pattern_28 | 108 | | | DNASE_5 | 1.911520e-03 | |

| pattern | num_seqlets | cwm_fwd | cwm_rev | TOMTOM_match | TOMTOM_qval | TOMTOM_match_logo |
|---|---|---|---|---|---|---|
| pos_patterns.pattern_29 | 104 | | | IRF3_HUMAN.H11MO.0.B | 1.353080e-02 | |
| pos_patterns.pattern_30 | 91 | | | CRX_HUMAN.H11MO.0.B | 1.000000e+00 | |
| pos_patterns.pattern_31 | 87 | | | SP1_HUMAN.H11MO.0.A | 7.849410e-10 | |
| pos_patterns.pattern_32 | 72 | | | THA_HUMAN.H11MO.0.C | 1.000000e+00 | |
| pos_patterns.pattern_33 | 71 | | | RUNX1_MOUSE.H11MO.0.A | 1.145040e-02 | |
| pos_patterns.pattern_34 | 68 | | | ZBTB33_MA0527.1 | 7.716610e-05 | |
| pos_patterns.pattern_35 | 52 | | | THA11_HUMAN.H11MO.0.B | 6.607040e-06 | |
| pos_patterns.pattern_36 | 50 | | | Six3_MA0631.1 | 1.000000e+00 | |
| pos_patterns.pattern_37 | 46 | | | RUNX3_MOUSE.H11MO.0.A | 5.178750e-03 | |
| pos_patterns.pattern_38 | 44 | | | ZN121_HUMAN.H11MO.0.C | 3.891050e-03 | |
| pos_patterns.pattern_39 | 39 | | | RFX5_RFX_2 | 1.530100e-08 | |
| pos_patterns.pattern_40 | 26 | | | CTCFL_MA1102.1 | 1.000000e+00 | |
| pos_patterns.pattern_41 | 26 | | | DNASE_2 | 2.725410e-01 | |
| pos_patterns.pattern_42 | 24 | | | ETV6_MOUSE.H11MO.0.C | 1.638190e-01 | |
| pos_patterns.pattern_43 | 21 | | | FOXB1_forkhead_2 | 6.569020e-01 | |
| pos_patterns.pattern_44 | 20 | | | PAX6_PAX_1 | 4.642670e-04 | |
| neg_patterns.pattern_0 | 108 | | | ERG_HUMAN.H11MO.0.A | 5.002480e-06 | |
| neg_patterns.pattern_1 | 83 | | | IRF1_MOUSE.H11MO.0.A | 8.981530e-07 | |
| neg_patterns.pattern_2 | 59 | | | CTCF_MA0139.1 | 5.809740e-10 | |
| neg_patterns.pattern_3 | 59 | | | Egr1.mouse_C2H2_1 | 6.336640e-02 | |
| neg_patterns.pattern_4 | 52 | | | FOSL1_MA0477.1 | 7.326350e-06 | |
| neg_patterns.pattern_5 | 49 | | | ZNF384_MA1125.1 | 5.278780e-01 | |
| neg_patterns.pattern_6 | 49 | | | ZNF384_MA1125.1 | 1.180350e-02 | |
| neg_patterns.pattern_7 | 41 | | | PROP1_homeodomain_2 | 2.710320e-01 | |
| neg_patterns.pattern_8 | 34 | | | NFKB1_HUMAN.H11MO.1.B | 4.702380e-05 | |
| neg_patterns.pattern_9 | 32 | | | TEAD4_MOUSE.H11MO.0.A | 1.000000e+00 | |
| neg_patterns.pattern_10 | 22 | | | MYB_MOUSE.H11MO.0.A | 6.650920e-02 | |
