## Supplementary Files 4 for "ChromBPNet: bias factorized, base-resolution deep learning models of chromatin accessibility reveal cis-regulatory sequence syntax, transcription factor footprints and regulatory variants": fig4k_gm12878_DNASE_raw_bpnet_w_heg2_bias_correction_counts_modisco.pdf

| pattern | num_seqlets | cwm_fwd | cwm_rev | TOMTOM_match | TOMTOM_qval | TOMTOM_match_logo |
|---|---|---|---|---|---|---|
| pos_patterns.pattern_0 | 4730 | | | ELF5_HUMAN.H11MO.0.A | 1.547520e-05 | |
| pos_patterns.pattern_1 | 3426 | | | CTCF_MA0139.1 | 1.524140e-11 | |
| pos_patterns.pattern_2 | 3347 | | | IRF4_HUMAN.H11MO.0.A | 1.119210e-02 | |
| pos_patterns.pattern_3 | 3026 | | | RUNX3_HUMAN.H11MO.0.A | 9.365550e-02 | |
| pos_patterns.pattern_4 | 2229 | | | NFKB1_HUMAN.H11MO.1.B | 7.809360e-07 | |
| pos_patterns.pattern_5 | 2171 | | | JDP2_MA0655.1 | 3.396910e-03 | |
| pos_patterns.pattern_6 | 1777 | | | IRF4_HUMAN.H11MO.0.A | 4.690170e-08 | |
| pos_patterns.pattern_7 | 679 | | | SP1_HUMAN.H11MO.0.A | 3.160590e-05 | |
| pos_patterns.pattern_8 | 614 | | | BATF3_HUMAN.H11MO.0.B | 1.967330e-07 | |
| pos_patterns.pattern_9 | 506 | | | ETS1_HUMAN.H11MO.0.A | 8.071990e-02 | |
| pos_patterns.pattern_10 | 418 | | | NRF1_HUMAN.H11MO.0.A | 3.575780e-08 | |
| pos_patterns.pattern_11 | 324 | | | NFYC_HUMAN.H11MO.0.A | 1.341650e-03 | |
| pos_patterns.pattern_12 | 301 | | | POU5F1_MA1115.1 | 3.380980e-03 | |
| pos_patterns.pattern_13 | 258 | | | ZN143_MOUSE.H11MO.0.A | 3.858330e-15 | |
| pos_patterns.pattern_14 | 238 | | | XBP1_bZIP_1 | 3.835180e-04 | |
| pos_patterns.pattern_15 | 219 | | | EBF1_EBF_1 | 8.724300e-06 | |
| pos_patterns.pattern_16 | 176 | | | SPI1_ETS_1 | 2.157080e-02 | |
| pos_patterns.pattern_17 | 156 | | | PAX2_PAX_1 | 1.332250e-09 | |
| pos_patterns.pattern_18 | 97 | | | HNF1B_HUMAN.H11MO.0.A | 1.083950e-05 | |
| pos_patterns.pattern_19 | 92 | | | ZBTB33_MA0527.1 | 6.556330e-06 | |
| pos_patterns.pattern_20 | 66 | | | TYY1_HUMAN.H11MO.0.A | 1.018530e-04 | |
| pos_patterns.pattern_21 | 55 | | | PAX2_PAX_1 | 2.307710e-05 | |
| pos_patterns.pattern_22 | 54 | | | BATF+JUN_MA0462.1 | 2.162940e-03 | |
| pos_patterns.pattern_23 | 47 | | | Rfx1_MA0509.1 | 1.501770e-08 | |
| pos_patterns.pattern_24 | 40 | | | ZNF524_C2H2_1 | 5.388540e-02 | |
| pos_patterns.pattern_25 | 39 | | | MEF2D_HUMAN.H11MO.0.A | 2.220990e-04 | |
| pos_patterns.pattern_26 | 35 | | | MAZ_HUMAN.H11MO.0.A | 3.010770e-05 | |
| pos_patterns.pattern_27 | 27 | | | RORA_nuclearreceptor_1 | 1.000000e+00 | |
| pos_patterns.pattern_28 | 27 | | | SP2_MA0516.1 | 6.836270e-02 | |

| pattern | num_seqlets | cwm_fwd | cwm_rev | TOMTOM_match | TOMTOM_qval | TOMTOM_match_logo |
|---|---|---|---|---|---|---|
| pos_patterns.pattern_29 | 24 | 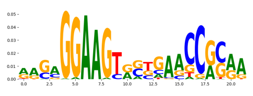 | 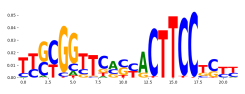 | RUNX3_HUMAN.H11MO.0.A | 1.214550e-01 | 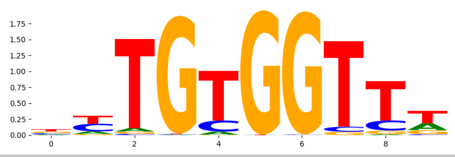 |
