## Supplementary Files 4 for "ChromBPNet: bias factorized, base-resolution deep learning models of chromatin accessibility reveal cis-regulatory sequence syntax, transcription factor footprints and regulatory variants": fig4l_gm12878_DNASE_raw_bpnet_w_hepg2_bias_correction_profile_modisco.pdf

| pattern | num_seqlets | cwm_fwd | cwm_rev | TOMTOM_match | TOMTOM_qval | TOMTOM_match_logo |
|---|---|---|---|---|---|---|
| pos_patterns.pattern_0 | 5249 | | | ELF5_HUMAN.H11MO.0.A | 9.691420e-05 | |
| pos_patterns.pattern_1 | 5033 | | | CTCF_MA0139.1 | 2.194550e-12 | |
| pos_patterns.pattern_2 | 4134 | | | RUNX3_HUMAN.H11MO.0.A | 8.828870e-02 | |
| pos_patterns.pattern_3 | 3013 | | | STAT1+STAT2_MA0517.1 | 1.285860e-01 | |
| pos_patterns.pattern_4 | 2725 | | | JDP2_MA0655.1 | 2.140440e-03 | |
| pos_patterns.pattern_5 | 1650 | | | NFKB1_HUMAN.H11MO.1.B | 2.103800e-05 | |
| pos_patterns.pattern_6 | 1550 | | | KLF12_HUMAN.H11MO.0.C | 3.349660e-06 | |
| pos_patterns.pattern_7 | 1273 | | | IRF4_HUMAN.H11MO.0.A | 2.337670e-07 | |
| pos_patterns.pattern_8 | 1070 | | | STAT2_HUMAN.H11MO.0.A | 4.737680e-06 | |
| pos_patterns.pattern_9 | 947 | | | NFYB_HUMAN.H11MO.0.A | 2.764350e-04 | |
| pos_patterns.pattern_10 | 687 | | | NRF1_HUMAN.H11MO.0.A | 1.436610e-06 | |
| pos_patterns.pattern_11 | 551 | | | SP5_MOUSE.H11MO.0.C | 2.550590e-07 | |
| pos_patterns.pattern_12 | 459 | | | POU5F1_MA1115.1 | 3.140390e-02 | |
| pos_patterns.pattern_13 | 438 | | | FOSB+JUNB_MA1136.1 | 3.041810e-05 | |
| pos_patterns.pattern_14 | 406 | | | ETS1_HUMAN.H11MO.0.A | 3.318090e-04 | |
| pos_patterns.pattern_15 | 316 | | | RFX3_MOUSE.H11MO.0.C | 8.473740e-09 | |
| pos_patterns.pattern_16 | 309 | | | MITF_MA0620.2 | 9.471860e-06 | |
| pos_patterns.pattern_17 | 270 | | | SP5_MOUSE.H11MO.0.C | 4.327930e-08 | |
| pos_patterns.pattern_18 | 263 | | | ZN143_MOUSE.H11MO.0.A | 2.385520e-19 | |
| pos_patterns.pattern_19 | 262 | | | EBF1_EBF_1 | 1.091940e-05 | |
| pos_patterns.pattern_20 | 214 | | | ZN281_MOUSE.H11MO.0.A | 2.328410e-04 | |
| pos_patterns.pattern_21 | 207 | | | PAX2_PAX_1 | 1.733700e-09 | |
| pos_patterns.pattern_22 | 119 | | | ETV6_ETS_1 | 8.093770e-02 | |
| pos_patterns.pattern_23 | 111 | | | ZBTB33_MA0527.1 | 1.473020e-04 | |
| pos_patterns.pattern_24 | 96 | | | HNF1A_HUMAN.H11MO.0.C | 8.929480e-06 | |
| pos_patterns.pattern_25 | 74 | | | ZN143_HUMAN.H11MO.0.A | 1.830400e-07 | |
| pos_patterns.pattern_26 | 70 | | | PAX2_PAX_1 | 4.364550e-04 | |
| pos_patterns.pattern_27 | 68 | | | IKZF1_HUMAN.H11MO.0.C | 8.640450e-01 | |
| pos_patterns.pattern_28 | 68 | | | SP1_MOUSE.H11MO.0.A | 1.626670e-05 | |

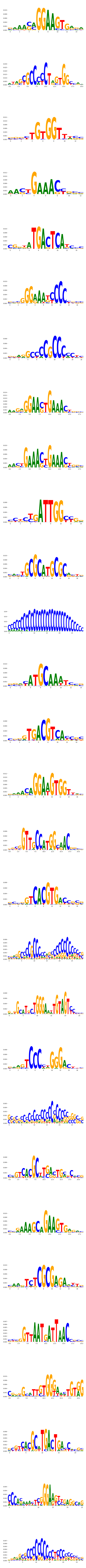

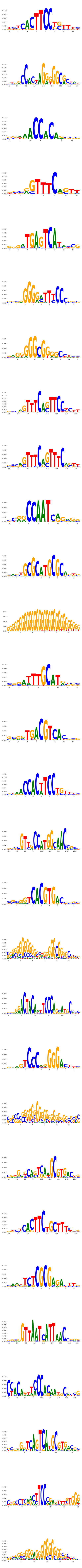

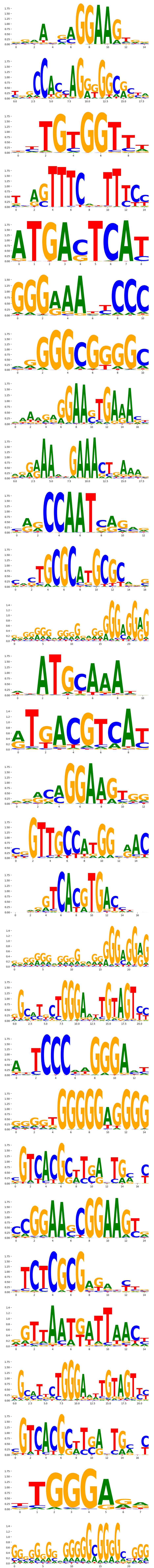

| pattern | num_seqlets | cwm_fwd | cwm_rev | TOMTOM_match | TOMTOM_qval | TOMTOM_match_logo |
|---|---|---|---|---|---|---|
| pos_patterns.pattern_29 | 52 | | | BATF+JUN_MA0462.1 | 1.290240e-03 | |
| pos_patterns.pattern_30 | 42 | | | BATF+JUN_MA0462.1 | 7.203670e-03 | |
| pos_patterns.pattern_31 | 39 | | | ZN549_HUMAN.H11MO.0.C | 1.000000e+00 | |
| pos_patterns.pattern_32 | 38 | | | Gfi1b_MA0483.1 | 1.000000e+00 | |
| pos_patterns.pattern_33 | 34 | | | TFCP2_CP2_2 | 1.000000e+00 | |
| pos_patterns.pattern_34 | 33 | | | MEF2D_MA0773.1 | 4.503870e-02 | |
| pos_patterns.pattern_35 | 26 | | | NFIA_HUMAN.H11MO.0.C | 3.381870e-04 | |
| pos_patterns.pattern_36 | 22 | | | DNASE_5 | 2.572300e-04 | |
| neg_patterns.pattern_0 | 112 | | | TEAD3_MA0808.1 | 8.774540e-01 | |
| neg_patterns.pattern_1 | 90 | | | ETS1_HUMAN.H11MO.0.A | 8.614510e-06 | |
| neg_patterns.pattern_2 | 75 | | | PROP1_MA0715.1 | 1.022970e-03 | |
| neg_patterns.pattern_3 | 69 | | | IRF1_MOUSE.H11MO.0.A | 4.594720e-05 | |
| neg_patterns.pattern_4 | 33 | | | ZFX_MOUSE.H11MO.0.B | 2.394470e-01 | |
| neg_patterns.pattern_5 | 28 | | | NFKB2_HUMAN.H11MO.0.B | 9.446830e-03 | |
| neg_patterns.pattern_6 | 25 | | | SP4_HUMAN.H11MO.0.A | 3.061250e-04 | |
| neg_patterns.pattern_7 | 24 | | | STA5A_HUMAN.H11MO.0.A | 7.568250e-02 | |
| neg_patterns.pattern_8 | 22 | | | JUNB_MOUSE.H11MO.0.A | 8.143200e-03 | |
| neg_patterns.pattern_9 | 20 | | | RUNX3_HUMAN.H11MO.0.A | 9.070560e-02 | |
